# A unique cerebellar pattern of microglia activation in a mouse model of encephalopathy of prematurity

**DOI:** 10.1101/2021.06.26.449853

**Authors:** Luisa Klein, Juliette Van Steenwinckel, Bobbi Fleiss, Till Scheuer, Christoph Bührer, Valerie Faivre, Cindy Bokobza, Sophie Lemoine, Corinne Blugeon, Leslie Schwendimann, Zsolt Csaba, Dulcie A. Vousden, Jason P. Lerch, Anthony C. Vernon, Pierre Gressens, Thomas Schmitz

## Abstract

Preterm infants often show pathologies of the cerebellum, which are associated with impaired motor performance, lower IQ and poor language skills at school ages. Because 1 in 10 babies is born preterm cerebellar injury is a significant clinical problem. The causes of cerebellar damage are yet to be fully explained. Herein, we tested the hypothesis that perinatal inflammatory stimuli may play a key role in cerebellar injury of preterm infants. We undertook our studies in an established mouse model of inflammation-induced encephalopathy of prematurity driven by systemic administration of the prototypic pro-inflammatory cytokine interleukin-1β (IL-1β). Inflammation is induced between postnatal day (P) 1 to day 5, timing equivalent to the last trimester for brain development in humans the period of vulnerability to preterm birth related brain injury. We investigated acute and long-term consequences for the cerebellum on brain volume expansion, oligodendroglial maturation, myelin levels and the microglial transcriptome. Perinatal inflammation induced global mouse brain volume reductions, including specific grey and white matter volume reductions in cerebellar lobules I and II (5% FDR) in IL-1β versus control treated mice from P15 onwards. Oligodendroglia damage preceded the MRI-detectable volume changes, as evidenced by a reduced proliferation of OLIG2+ cells at P10 and reduced levels of the myelin proteins MOG, MBP and MAG at P10 and P15. Increased density of Iba1+ cerebellar microglia was observed at P5 and P45, with evidence for increased microglial proliferation at P5 and P10. Comparison of the transcriptome of microglia isolated from P5 cerebelli and cerebrum revealed significant enrichment of pro-inflammatory markers in microglia from both regions, but in the cerebellum microglia displayed a unique type I interferon signalling dysregulation. Collectively, these data suggest that in our model that systemic inflammation causes chronic activation of microglia and maldevelopment of cerebellum that includes myelin deficits which is driven in the cerebellum by type I interferon signalling. Future protective strategies for preterm infants should consider sustained type I interferon signalling driven cerebellar inflammation as an important target.

## Introduction

Maldevelopment and injury to the developing cerebellum are increasingly linked to cognitive impairment including specific deficiencies in executive function, deficits of attention, and increased risk for psychiatric disorders with a neurodevelopmental origin such as autism spectrum disorder (ASD)^1–7^ and schizophrenia^8^. Babies born preterm (at less than 37 weeks of gestation) have a pronounced vulnerability to cerebellar injury^9, 10^, it is however still under-acknowledged, and cerebellar pathologies are likely to be underdiagnosed ^11^. Because approximately 15 million babies are born premature every year (−10% of all births)^12^ there is a significant need to understand how cerebellar maldevelopment and injury develops in these infants.

Exposure to systemic inflammation in the foetal and neonatal periods has been demonstrated to have a key role in driving brain maldevelopment and in causing injury in clinical studies and using pre-clinical models^13–16^. Brain damage occurring in preterm born infants is termed encephalopathy of prematurity (EoP), and it is characterised by neuroinflammation, oligodendrocyte maturation arrest and hypomyelination, axonopathy, reduced fractional anisotropy and cortical volume determined by magnetic resonance imaging (MRI), and eventually, significant cognitive deficits ^17^. The damaging effects of systemic inflammation are linked to two main processes. Firstly, that it causes preterm birth depriving the infant of trophic support including from the placenta and exposing it to the noxious extra-uterine environment^18^. Secondly, that it prevents glial cells from performing important brain building functions and instead forces them into a damaging inflammatory reactivity^19, 20^. A common source of inflammatory exposure is chorioamnionitis, which has specifically been identified as a risk factor for a diagnosis of cerebral palsy^19, 21^ or ASD^22–24^. Although strong inflammatory insults such as chorioamnionitis or postnatal sepsis are known drivers of brain changes, importantly there is also subtle, low grade perinatal/postnatal inflammation occurring in response to mechanical ventilation^18, 20^, medical interventions and medications^25^, that have also been shown to cause brain injury in experimental models^26^.

The brain’s resident innate immune cells, microglia, are a key mediator of brain injury in models of encephalopathy of prematurity ^27–30^ and in human causes of EoP ^31, 32^. Furthermore, the importance of typically functioning microglia for the development of the brain is underlined by their production of growth factors, such as insulin-like growth factor 1 (IGF1) ^33^, brain-derived neurotrophic factor (BDNF) ^34^and transforming growth factor beta (TGFβ) ^35^, that promote and regulate proliferation and maturation of immature neurons and oligodendrocytes^36, 37^. Microglia are also involved in in regulating neuronal death and shaping connectivity by regulating synaptic pruning during brain development^38^, including in the cerebellum ^39^.

It can therefore be hypothesized that one mechanism by which early exposure to perinatal systemic inflammation could have adverse effects on the developing cerebellum is by effects on microglia. Specifically, by interfering with the physiological roles of microglia and inducing inflammatory activation resulting in perturbed oligodendrocyte development and myelination. To test this hypothesis, we used an established mouse model of encephalopathy of prematurity triggered by systemic exposure to the pro-inflammatory cytokine interleukin-1β (IL-1β) during the developmental stage of the brain (postnatal day 1-5) corresponding to a window of vulnerability to human preterm birth (23-32 weeks’ gestation). We have already established in this model that in the forebrain there is a oligodendroglial dysmaturation ^25, 27, 40^ that is specifically caused by microglial reactivity ^25^ and that there are also deficits in grey matter gene expression profiles ^41^ and interneurons^42^. We have now expanded this work and have investigated the impact of neonatal systemic inflammation using magnetic resonance imaging (MRI), and uncovered a specific effect on cerebellar development, characterised by reduced cerebellar grey and white matter. Guided by these data we carried out focused *in vivo* and *ex vivo* studies on the cerebellar oligodendrocytes and microglia to link these macroscale changes to their cellular and molecular correlates. The results indicated that in addition to oligodendrocyte damage and myelin deficits caused by systemic inflammation, there was a persistent increase in microglial density and microglial activation in the cerebellum accompanied by a cerebella specific pattern of microglial inflammation driven by interferon type 1. The results of this study highlight brain region specific consequences of perinatal systemic inflammation on the developmental program of oligodendrocytes and microglia in the cerebellum and uncover a novel pathway regulating this region-specific outcome.

## Methods

Experimental protocols were approved by the institutional guidelines of the Institut National de la Santé et de la Recherche Medicale (Inserm, France) (Approval 2012-15/676-0079 and 2012-15/676-0083), the Ethics Committee and the services of the French Ministry in charge of Higher Education and Research according to the directive 2010/63/EU of the European Parliament (#9286-2016090617132750).

### Mouse model of Encephalopathy of Prematurity

Experiments were performed on OF1 strain mice (Charles River, France). As previously described^25, 27, 28, 40^, a 5 µl volume of phosphate-buffered PBS (PBS) containing 10 μg/kg/injection of recombinant mouse IL-1β (Miltenyi Biotec, Bergisch Gladbach, Germany) or of PBS alone (control) was injected intraperitoneal (i.p.) twice a day on days P1 to P4 and once a day on day P5. The timing of IL-1β injections (P1–P5) was chosen to mimic a chronic exposure to circulating cytokines at a developmental stage of the brain corresponding to human preterm birth (23 to 32-weeks gestational age). In this mouse model, male pups have more effected white matter compared to the females^27^, mimicking the higher vulnerability of preterm born males to poor neurodevelopmental outcome^43, 44^. Based on that, in the present study, we only used male pups. For proliferation analyses, pups received bromo-deoxyuridine (BrdU; BD Biosciences, San Jose, CA, USA) (10 mg/kg per day, P1 to P4) in conjunction with IL-1β or PBS-injections.

### Magnetic resonance imaging (MRI)

#### Perfusions for magnetic resonance imaging

For *ex vivo* magnetic resonance imaging (MRI), two cohorts of individual animals sampled randomly from 5 independent litters were sacrificed at P15 (PBS, n=9; IL-1β, n=9) and P60 (PBS, n=8; IL-1β, n=11). Littermates from these mice were assessed at P10 with qRT-PCR to ascertain they had the requisite oligodendrocyte maturation delay that is characteristic of the model. An additional cohort of PBS and IL-1β-exposed mice were sacrificed at P15 and P60 for *post-mortem* follow up. Details of the perfusion protocol have been described at length elsewhere^45^. Briefly, mice were anesthetized with pentobarbital and then transcardially perfused with 30 mL of 0.1M PBS containing 10 U/mL heparin and 2 mM Prohance (a gadolinium based contrast agent; Bracco Inc., Bucks, UK), followed by 30 mL of 4% paraformaldehyde (PFA) also containing 2 mM Prohance. Post-perfusion mice were decapitated and the skin, ears and lower jaw removed, but the brain was left inside the skull to minimize distortions or deformations from dissection. Brain tissues were then incubated in 4% PFA+Prohance overnight and prior to scanning, for an additional 7-14 days in 0.1M PBS containing 2 mM Prohance and 0.05% (w/v) sodium azide to equalise the contrast agent.

#### Data acquisition by MRI

A 7 T horizontal small-bore magnet and (Agilent Technologies Inc, Santa Clara, USA) and a quadrature volume radiofrequency coil (39 mm internal diameter, Rapid Biomedical GmbH) were used for all MRI acquisitions. Fixed brain samples were placed securely one a time in an MR-compatible holder immersed in proton-free susceptibility matching fluid (FluorinertTM FC-70; Sigma-Aldrich, St-Louis, MO, USA). In order to assess volume differences throughout the mouse brain, a T2-weighted 3D Fast Spin-Echo (FSE) sequence was used, with the following sequence parameters: TR=1000 ms, echo train length (ETL) =16, echo spacing=5.68 ms, TE_eff_ = 34 ms, field of view (FOV) = 19.2 mm x 19.2 mm x 19.2 mm and a matrix size of 192 x 192 x 192, yielding an isotropic (3D) resolution of 100 µm. Total imaging time for each brain was 38 minutes.

#### Image registration and statistical analysis

Image registration and analysis were carried out at the Mouse Imaging Centre (MICe) at the University of Toronto, ON, Canada. Deformation based morphometry (DBM) was used to analyse volume differences between PBS and IL-1β-exposed mice at P15 and P60. DBM is performed by registering the individual T2-weighted MR images from each mouse brain together via a series of linear (6, then 12-parameter fit) and non-linear registration steps using a fully automated pipeline^46 47^. Post-registration, all scans are resampled with an appropriate transform and averaged to create a population atlas, which represents the average anatomy of the mice in the study. Registration was performed using a combination of mni_autoreg tools^48^ and advanced normalization tools^49, 50^. The registration brings all the scans into alignment via deformation in an unbiased manner, allowing the experiment to then analyze the deformations requited to take each individual mouse’s anatomy into a final atlas space. The jacobian determinants of the deformation fields are utilized as the measures of apparent volume change on a per voxel basis, corrected for inter-individual mouse differences in total brain volume^51^. Regional volume differences are then calculated by warping a pre-existing classified MRI atlas onto the population atlas, permitting the calculation of regional volumes across the brain^52^. This MRI atlas includes 159 different structures incorporating three separate pre-existing atlases: (1) 62 different structures throughout the mouse brain including subcortical white and grey matter structures, corpus callosum, striatum, and thalamus^52^ (2) further divides the cerebellum into its various regions, individual lobules, white and grey matter, and the deep cerebellar nuclei^53^; and (3) divides the cortex into 64 different regions, including areas of the cingulate cortex and primary motor and somatosensory cortices^54^. The total brain volume is also included. Differences in brain volume were analysed using atlas-derived regional brain volumes using mixed linear modelling with age as within group factor (P15, P60), treatment (PBS or IL-1β) as between group factor and age x treatment interactions^46^. Multiple comparisons were controlled for using the false discovery rate (FDR) at 5% (q<0.05)^55^.

### Tissue preparation

Brain tissue samples were collected at P3, P5, P10, P45, P60 and P150 (Supplementary Figure 1) For molecular studies, cerebellums were snap-frozen in liquid nitrogen and stored at −80 °C until further analysis. For immunohistochemical studies, the animals were perfused with 0.1 M PBS followed by 4 % paraformaldehyde. The brains were then postfixed at 4 °C for 3 days, embedded in paraffin, and processed for histological staining. Sagittal paraffin sections of 10 μm were obtained and mounted onto Super Frost plus^TM^-coated slides (R. Langenbrinck, Emmendingen, Germany). Sections were deparaffinised using Roti-Histol Solution (Carl Roth, Karlsruhe, Germany) twice for 10min each and rehydrated in descendant ethanol concentrations (from 100%, 100%, 90%, 80%, 70%; 3min each) and finally distilled water (3min). For immunostainings, sagittal sections were selected according to sagittal positions 186 to 195 of the Allen mouse reference atlas (developing brain) (https://mouse.brain-map.org/static/atlas).

### Protein extraction

Snap-frozen whole cerebella were homogenized in radioimmunoprecipitation assay buffer (RIPA Buffer; Sigma–Aldrich) with complete Mini EDTA-free Protease Inhibitor Cocktail Tablets (Roche Diagnostics, Mannheim, Germany). The homogenate was centrifuged at 13,000 *g* (4 °C) for 20 min before collecting the supernatant. Protein concentrations were determined using the Bicinchoninic Acid Protein Assay kit (Pierce/Thermo Scientific, Rockford, IL, USA).

### Immunoblotting

Immunoblotting was performed as previously described^56^. Briefly, samples were diluted in Laemmli sample loading buffer (Bio-Rad, Munich, Germany) denatured (95 °C for 5 min) and protein extract (20 µg per lane) were cooled on ice, separated electrophoretically on 10-12% Mini-PROTEAN TGX precast gels (Bio-Rad), and transferred onto nitrocellulose membrane (0.2 µm pore; Bio-Rad) using a semidry electrotransfer unit at 15 V for 5 min. By staining the membranes with Ponceau S solution (Fluka, Buchs, Switzerland) equal loading and transfer of proteins was confirmed. After blocking the membranes with 5 % bovine serum albumin (BSA) in Tris-buffered PBS/ 0.1 % Tween 20 for 1 h at room temperature they were incubated overnight at 4 °C with the following antibodies: rabbit polyclonal anti-2′,3′-cyclic nucleotide 3′-phosphodiesterase (CNPase; 49 kDa; 1:1000; Thermo Fisher Scientific, Waltham, MA, USA), mouse monoclonal anti-myelin-associated glycoprotein (MAG, 63 and 69 kDa; 1:500; Abcam, Cambridge, UK), rabbit polyclonal anti-myelin oligodendrocyte protein (MOG, 27 kDa; 1:1000; Abcam), rabbit polyclonal anti-myelin proteolipid protein (PLP; 26 and 30 kDa; 1:1000; Abcam), mouse monoclonal anti-myelin basic protein (MBP, 18 and 24 kDa; 1:1000; Covance, Princeton, NJ, USA) or mouse monoclonal anti-β-actinin (42 kDa; 1:5000; Sigma Aldrich), respectively. Secondary incubations were performed with horseradish peroxidase-linked polyclonal goat anti-rabbit (1:5000; Dako, Glostrup, Denmark) or polyclonal rabbit anti-mouse (1:5000; Dako) antibodies. Positive signals were visualized using enhanced chemiluminescence (Amersham Biosciences, Freiburg, Germany) and quantified using a ChemiDoc XRS+ system and the software Image Lab (Bio-Rad). To ensure the equal loading and accuracy of changes in protein abundance, the protein levels were normalized to β-actin.

### Immunostaining

To increase cell permeability slides were microwaved for 10 min at 600 W in a pH 6.0 citrate buffer and afterwards left for 30 min at room temperature. For staining with BRDU it was necessary to apply hydrochloric acid for 20 min and then neutralize it with a borate buffer. Slides were cooled and washed using PBS solution before being blocked for one hour at room temperature with either blocking solution I (1 % BSA, 10 % normal goat serum (NGS), 0.05 % Tween-20 in PBS) for oligodendrocyte transcription factor/ proliferating cell nuclear antigen (OLIG2/PCNA), OLIG2/cleaved caspase 3 (Casp3), OLIG2/BrdU, ionized calcium binding adaptor molecule 1 (IBA1)/BRDU and OLIG1/ adenomatous polyposis coli (APC) or blocking solution II (3 % BSA, 1 % NGS, 0.05 % TW-20 in PBS) for MAG and MOG. Primary antibodies (Supplementary Table 1A) were diluted in DAKO Antibody diluent (Dako Deutschland, Hamburg, Germany) and subsequently incubated at 4 °C for 24-48 h. Secondary fluorescent antibodies (Supplementary Table 1B) were diluted 1:200 in Dako Antibody diluent and incubated at room temperature for 1h. Sections were counterstained with 4,6-diamidino-2-phenylindole (DAPI, 10 ng/ml, Sigma Aldrich, diluted 1:2000 in PBS, incubation time 10 min). Slides were mounted using mounting medium (Vectashield HardSet Mounting Media, Vector Laboratories, Burlingame, CA, USA).

Sections were viewed with Keyence BZ-900 microscope (BIOREVO) with 20-40x magnification and confocal z-stacks were merged and analyzed with BZII-analyser software (Keyence Deutschland GmbH, Neu-Isenburg, Germany). Region of interest were found in lobules I-VI in the proximal parts within lobules, which in mice start being myelinated at postnatal ages ^57^. Three photos per lobule were taken. Cell counts have been performed with Adobe Photoshop CS6 Extended Version 13.0 x64 (Adobe Systems Incorporated, San Jose, CA, USA). For statistical analyses, the mean value of all sections per animal was used. All images were acquired by an experimenter who was blinded to the treatment group.

### RNA extraction and quantitative real-time PCR

RNA isolation was performed by acidic phenol/chloroform extraction (peqGOLD RNApure, Peqlab Biotechnologie, Erlangen, Germany) from snap frozen tissue and 2 µg of total RNA was reverse transcribed. Real time quantification of PCR products of *Olig2*, *Cnp*, inducible nitric oxide synthetase (*Nos2)*, major histocompatibility complex II (*MhcII)*, superoxide dismutase 2 (*Sod2)*, glutamate cysteine ligase (*Gclc)*, histocompatibility 2, class II A (*H2-Ab),* and Cyclin D2 (*Ccnd2)* was performed using dye-labelled fluorogenic reporter oligonucleotide probes or SYBR^TM^Green. Sequences of primers are given in Supplementary Table 1C. PCR was performed in triplicate. The reaction mixture consisted of 5 µl of qPCRBIO Probe Mix High ROX (Thermo Fisher), 2.5 µl (1.25 µM) oligonucleotide mix, 0.5 µl (0.2 µM) of probe (BioTeZ, Berlin, Germany), and 3 µl of cDNA template with hypoxanthine phosphoribosyltransferase 1 (*Hprt)* used as an internal reference. For SYBR^TM^Green the reaction mixture consisted of 5 µl of qPCRBIO SYGreen Mix High ROX (Thermo Fisher), 2.5 µl (1.25 µM) oligonucleotide mix and 3 µl of cDNA template. 96-well reaction plates were used for PCR amplification for 40 cycles, each cycle at 95 °C for 15 seconds and 60 °C for 1 min. Analysis of expression profiles was done using StepOnePlus^TM^ real-time PCR system (Applied Biosystems, Life Technologies, Carlsbad, CA, USA) using the 2^−ΔΔ*C*T^ method^56^.

### Cell sorting and RNA extraction from isolated cells

Cerebrum and cerebellum at P5 were collected for cell dissociation using Neural tissue Dissociation (Miltenyi Biotec) kit with papain and double cell isolation using magnetic coupled antibodies anti-CD11B (Microglia) according to the manufacturer’s protocol (Miltenyi Biotec) and as previously described^27, 28^. Cells were pelleted and conserved at - 80 °C. RNA from CD11B+ cells from brains of mice exposed to IL-1β or PBS at P5 were extracted (RNA XS plus, Macherey-Nagel, Dueren, Germany).

### RNA sequencing and bioinformatics analysis

Library preparation and Illumina sequencing were performed at the Ecole Normale Supérieure genomic core facility (Paris, France). Messenger (polyA+) RNAs were purified from 100 ng of total RNA using oligo(dT). Libraries were prepared using the strand specific RNA-Seq library preparation TruSeq Stranded mRNA kit (Illumina, Évry-Courcouronnes, France). Libraries were multiplexed by 8 on 4 high-output flow cells. Four 75 bp single read sequencing were performed on a NextSeq 500 (Illumina). The analyses were performed using the Eoulsan pipeline^58^, including read filtering, mapping, alignment filtering, read quantification, normalisation and differential analysis: Before mapping, poly N read tails were trimmed, reads ≤40 bases were removed, and reads with quality mean ≤30 were discarded. Reads were then aligned against the Mus musculus genome from Ensembl version 91 using STAR (version 2.6.1b)^59^. Alignments from reads matching more than once on the reference genome were removed using Java version of samtools^60^. To compute gene expression, Mus musculus GTF genome annotation version 91 from Ensembl database was used. All overlapping regions between alignments and referenced exons were counted using HTSeq-count 0.5.3^61^ and then aggregated to each referenced gene. The sample counts were normalized using DESeq2 1.8.1^62^. Statistical treatments and differential analyses were also performed using DESeq2 1.8.1. Heatmap and hierarchical clustering were realized with one minus Pearson correlation and was generated using Morpheus (https://software.broadinstitute.org/morpheus/). Significant differentially expressed (DE) genes under IL-1β condition in CD11B+ cells were identified with Benjamini and Hochberg adjusted p-values <0.05. Functional analyses were done using DAVID 6.8 (https://david.ncifcrf.gov). Analysis of predicted protein interactions was undertaken using STRING (https://string-db.org) with an interaction stringency of 0.400 (medium confidence) and first and second shell interactions set to none to allow the significance of the interactions to be accurately assessed. To access WikiPathways (https://www.wikipathways.org) we used the Enrichr tool (https://maayanlab.cloud/Enrichr/;Kuleshov). Significant representative GO terms, STRING local network clusters and Wikigene pathways (mouse) were identified with Benjamini and Hochberg adjusted p-values <0.05.

### Fluorescence Activated Cell Sorting (FACS) analysis

Cerebrum and cerebellum at P5 were collected for cell dissociation using Neural Tissue Dissociation (Miltenyi Biotec). The cells were counted (Nucleocounter NC-200, Chemometec, Allerod, Denmark) and re-suspended at 10.10^6^ cells/ml in FACs buffer (Dulbeccos’s Phosphate Buffer Saline (Gibco Life Technologies, Paisly, UK), 2mM EDTA (Sigma Aldrich, Saint Louis, MO, USA), 0,5% Bovine Serum Albumin (Miltenyi Biotec). After Fc blocking (BD Biosciences, Le Pont De Claix, France), cells were incubated with viability probe (FVS780, BD Biosciences) and fluorophore-conjugated antibodies against mouse CD45, CD11B (CD11B BV421, clone MI/70, CD45BV510, clone 30-F11; Sony Biotechnology, San Jose, CA, USA), CD18 (CD18 APC, clone C71/16) BD Biosciences, Le Pont De Claix, France) or their corresponding control isotypes (Sony Biotechnology and BD Biosciences), at concentrations recommended by the manufacturers or calculated after titration. Cells were washed with FACs buffer, resuspended in PBS and FACs analysis was done within 24hrs. After doublets and dead cells exclusion based respectively on morphological parameters and FVS780 staining, gating strategy selected microglia as live/CD11B^+/^CD45^lo^. Expression of CD18 was analysed in microglia. Myeloid cells (including polymorphonuclear neutrophils, monocytes and macrophages) were defined as CD11B^+^/CD45^high^. For oligodendrocyte marker O4 (O4)/platelet derived growth factor receptor alpha (PDGFRa) analysis, only P5 cerebellum cells were analysed. After Fc blocking (BD Biosciences), cells were incubated with viability probe (FVS780, BD Biosciences) and fluorophore-conjugated antibodies against mouse O4 (O4 VioBright 515, Miltenyi Biotech) and PDGFRa (CD140a-PE, Miltenyi Biotech).

### Statistical analysis of neuropathology analyses

All quantitative data are expressed as the mean ± standard error of the mean (SEM) for each treatment group. Groups were compared pairwise by using either t-test or nonparametric Mann-Whitney-U test (GraphPad Prism 5.0; GraphPad Software, San Diego, CA, USA) as appropriate. A two-sided p value <0.05 was considered statistically significant.

## Results

### Systemic inflammation reduced total and regional mouse brain volumes especially in the cerebellum

As expected, total mouse brain volume increased as a function of age between P15 and P60 (F (1,33) = 59.4; p<0.0001) (Supplementary Figure 2). A statistically significant main effect of systemic IL-1β exposure was also found (F (1,33) = 6.7; p<0.01). Although the age x treatment interaction term did not reach statistical significance (F (1,33) = 0.99; p=0.34), *post-hoc* testing on the main effect of treatment revealed a statistically significant reduction in mouse total brain volume at P60 (−5.3%; p=0.02; q=0.02), but not at P15 (−2.7%; p=0.27; q=0.14) in those mice exposed to systemic inflammation relative to controls (Supplementary Figure 2).

We next probed which brain regions contribute to the global effect of systemic inflammation on mouse brain volume, using atlas-based segmentation (ABS) to examine the effects of age, treatment and age x treatment interaction with linear mixed effect models, corrected for multiple comparisons using the false discovery rate (FDR) at 5% (q<0.05) (Figure 1). Using absolute volumes (mm^3^), after correction for multiple comparisons we observed that 64% (102/159) of mouse brain atlas regions of interest (ROI) were statistically significantly affected by increasing postnatal age across both treatment group. Also, it was noted that 27% (43/159) of atlas ROIs were statistically significantly different as a function of IL-1β exposure, relative to PBS-exposed mice (Supplementary Table 2) and each of these 43 brain regions were smaller in IL-1β-exposed mice, relative to PBS-exposed mice at both P15 and P60 (Figure 1A). Whilst these areas covered a range of cortical and sub-cortical regions as well as ventricular structures, the effects of systemic inflammation were particularly concentrated in the cerebellar grey and white matter, which also displayed the largest apparent volume reductions (% change; Supplementary Table 2). Across all atlas ROIs however, there were no statistically significant age x treatment interactions (Supplementary Table 2).

**Figure 1:**
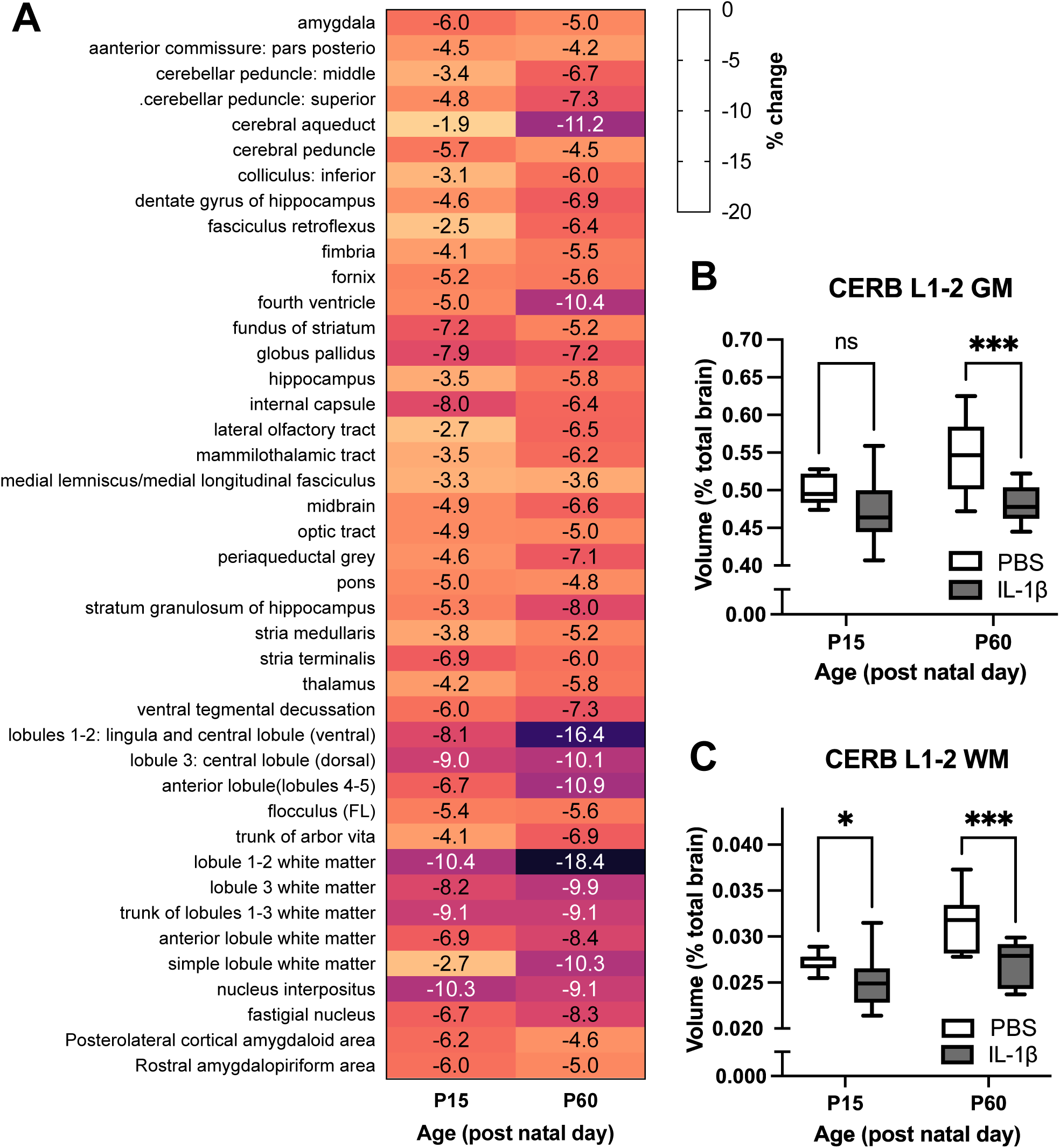
Impact of systemic inflammation on brain volumes. **A**: Heat map to illustrate the % change in absolute volumes (mm^3^) between IL-1β-exposed mice relative to PBS controls at P15 and P60, derived from atlas based segmentation. Only brain regions that were significantly different after multiple comparisons correction (5% FDR) are shown. Relative volumes (expressed as a % of total brain volume) for **B**: cerebella grey matter (GM) volumes and **C**: cerebella white matter (WM) volumes in lobules I-II. Horizontal line indicates group mean. *p<0.05; ***p<0.001 PBS vs. IL-1β-exposed mice, post-hoc analysis corrected for multiple comparisons (2-step FDR method at 5%); ns, not significant.

Absolute volume analysis however, cannot determine whether brain regions were smaller but normally scaled, or if any individual brain regions were abnormally scaled relative to whole brain volume change following systemic inflammation^63^. Put another way; are the brains smaller following systemic inflammation, but normally scaled? This is relevant since abnormal scaling could indicate atypical connectivity, or enervation of other brain areas, which could contribute to behavioural dysfunction^64^. Evidence of abnormal scaling could also indicate developmental timing of the effects of systemic inflammation. To examine the scaling of individual brain regions, we therefore calculated the relative volume of each brain region ([(brain region volume)/(whole brain volume) × 100]^65^. This analysis revealed that 74% (118/159) of mouse atlas brain regions were altered with increasing post-natal age across both treatment groups (Supplementary Table 2). In contrast to the absolute volume data, only 1% (2/159) of atlas ROIs were statistically significantly different in IL-1β-exposed mice relative to PBS-exposed mice (FDR q<0.05) these being the grey and white matter ROIs for cerebellar lobules 1-2 (Figure 1B-C; Supplementary Table 2). These regions also showed the largest % changes in the absolute volume dataset (Supplementary Table 2). *Post-hoc* testing revealed that these changes in the lobule 1-2 grey matter were only statistically significant at P60, but not P15, but at both time-points for the cerebellar white matter, suggestive of differential developmental timing (Figure 1B-C). There were however no statistically significant age x treatment interactions after correction for multiple comparisons (5% FDR). Collectively, these data suggest that systemic inflammation induces apparent global effects on mouse brain volume but controlling for this suggests a more selective alteration of cerebellar white matter growth from at least P15 onwards, which is maintained at least until P60, with grey matter effects becoming apparent from P60 onwards.

### Systemic inflammation altered myelination and oligodendrocyte maturation in the cerebellum

Guided by the aforementioned MRI data we next carried out focused post-mortem investigations in the cerebellum to establish the cellular correlates of the apparent macroscale volume changes in this region following IL-1β exposure focused on oligodendrocytes. Firstly, we examined the levels of the myelin proteins MBP and MAG at ages P10, P15 and P60 via western blotting (Figure 2, Supplementary Figure 3). At P10 and P15, (but not P60) there was a significant reduction in the levels of MAG (Figure 2A-B) in the mice exposed to systemic inflammation. At P15 there was a marked reduction MBP expression, in comparison to control litters (Figure 2C-D) although levels were visibly lower at each age tested. In agreement with this western blotting data, immunohistochemical analyses revealed that the numbers of oligodendrocytes co-labelled with the mature oligodendrocyte marker adenomatous polyposis coli (APC) and the oligodendrocyte transcription factor 1 (OLIG1) was significantly lower at P10 in mice exposed to systemic inflammation as compared to controls (Figure 2E-F). Specifically, in the control group the vast majority of OLIG1+ oligodendroglia co-expressed (mean = 82.46 %, ± 2.14 %) but there was a significant reduction in these + mature oligodendrocyte numbers in IL-1β-exposed mice (mean = 56.67 %, ± 3.86 %) (p<0.01, n=5; Figure 3D). Of important note, there was no significant difference in the density of OLIG2+ cells between the two experimental groups, both at P5 (89.50 ± 13.57 cells/ 0.2 mm^2^ in controls vs 88.25 ± 16.03 in IL-1β-exposed mice; p>0.05, n=5) and P10 (117.20 ± 15.53 cells/ 0.2 mm^2^ in controls vs 115.70 ± 12.19 in IL-1β-exposed mice; p>0.05, n=5).

**Figure 2:**
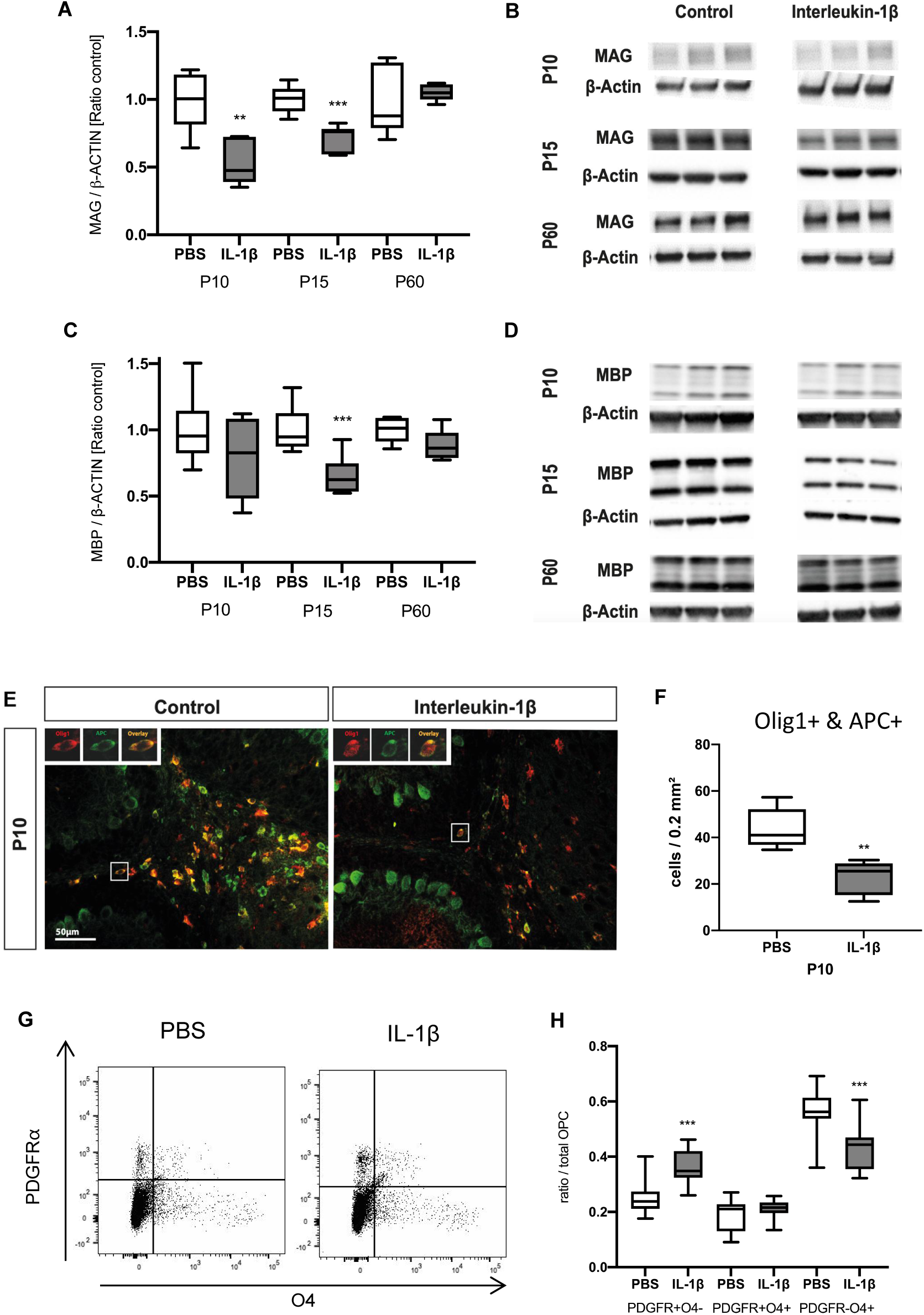
Impact of systemic inflammation on myelin and oligodendroglial maturation. **A-D**: Western blotting of MAG (A-B) and MBP (C-D) of P10, P15, and P60 cerebellar tissues from PBS vs IL-1β-exposed mice (**p<0.01, ***p<0.001; t test). **E-F**: Representative micrographs of APC+ (green) and OLIG1+ (red) cells at P10 in cerebellar slices from PBS vs IL-1β-exposed pups (E; scale bar = 50µm) and corresponding quantification (F; **P<0.01, t test). **G-H**: Representative scatter plots of the FACS analysis of total numbers of PDGFRa+/O4+ oligodendrocytes in P10 cerebellum from PBS or IL-1β-exposed pups and corresponding quantification (F; ***p<0.001, 2-way ANOVA).

**Figure 3:**
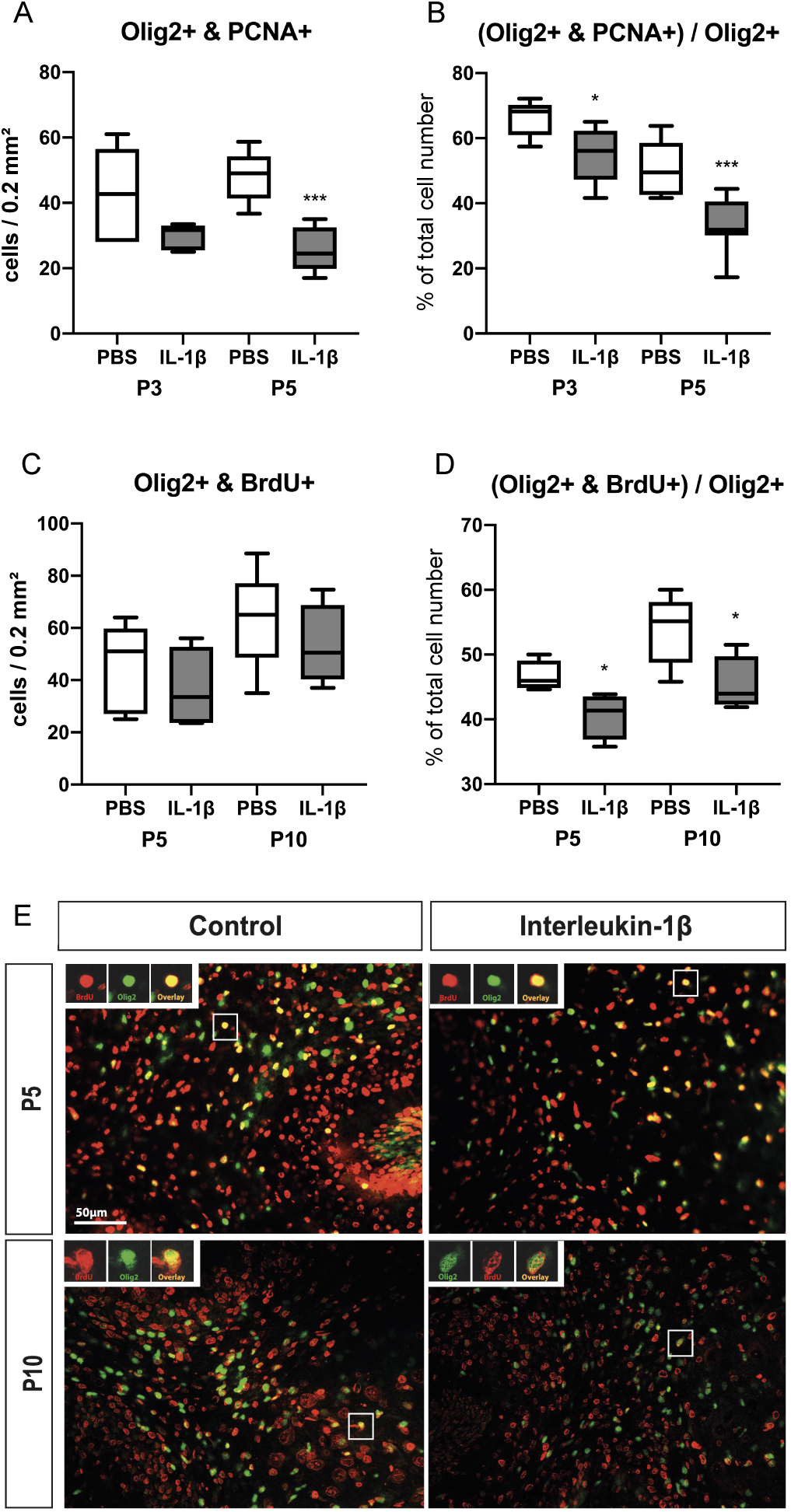
Impact of systemic inflammation on oligodendroglial proliferation. **A-B**: Quantification of OLIG2+/PCNA+ cell numbers and OLIG2+/PCNA+ over OLIG2+ ratios in the cerebellum at P3 and P5 in PBS and IL-1β-exposed P5 pups. **C-D**: Quantification of OLIG2+/BRDU+ cell number and OLIG2/BRDU+ over OLIG2+ ratios in the cerebellum at P5 and P10 in PBS and IL-1β-exposed P5 pups. *p<0.05, ***p<0.001; t-test. **E**: Representative micrographs of proliferating (BRDU+, red) OLIG2+ (green) cells at P5 and P10 in cerebellar slices from IL-1β-exposed pups.

We then sought to determine if the reduction in mature oligodendrocytes could be explained by increased levels of cell death and/or modification of progenitor pools. We assessed cerebellar cell death at P3, P5 and P10 and found only three to six CASP3+ cells (apoptotic cell death marker) per 0.2 mm^2^ at P3 and P5, and three or less at P10, with no detectable impact of IL-1β exposure (Supplementary Figure 4). ‘

We then explored the effect of systemic inflammation on oligodendroglial lineage progression using FACS at P5 to determine the numbers of PDGFR+/O4-OPCs, PDGFR+/O4+ intermediate precursor / immature oligodendrocytes, and PDGFR-/O4+ immature oligodendrocytes after perinatal exposure to IL-1β compared to controls injected with PBS. A clear increase in PDGFR+/O4-OPC cell number was noted in the cerebellum of pups with systemic inflammation in comparison to control pups (Figure 2G-H). At the same time, numbers of immature oligodendrocytes were reduced in pups after inflammatory exposure when compared to controls (Figure 2G-H). The population of PDGFR+O4+ intermediate stage cells was not altered.

### Cerebellar oligodendrocyte proliferation was inhibited by systemic inflammation

Having determined that there was a dysmaturation of oligodendrocytes, we also sought to determine if our macroscale observations might also be related to changes in oligodendrocyte proliferation. The rapid expansion of the cerebellar volume during development is characterized by high proliferation of oligodendrocyte precursor cells (OPCs) and immature oligodendrocytes in the cerebellar white matter, while during further development, maturation is enhanced and numbers of mature, post-mitotic oligodendrocytes increase^66–68^. To determine proliferation of oligodendroglial lineage cells in our study, firstly we performed immunohistochemistry for OLIG2 in combination with proliferation marker PCNA. The absolute numbers of OLIG2+/PCNA+ cells in the cerebellum of IL-1β-exposed mice at postnatal age P5 was lower than in control littermates (Figure 3A). At P3, there was a 23% reduction of OLIG2+/PCNA+ cells but this was not statistically significant. As a ratio of all OLIG2+ oligodendroglial lineage cells in the cerebellar white matter (OLIG2+/PCNA+ over all OLIG2+ cells), the proportion of proliferating oligodendroglia was very high with 68.25 ± 2.49 % in control pups at P3 and 49.50 ± 3.07 % at P5 (Figure 3B). In IL-1β-treated pups, this ratio of OLIG2+/PCNA+ over all OLIG2+ cells was significantly decreased at P3 and P5 in mice exposed to systemic inflammation (Figure 3B). To assess the cumulative proliferation over P1-P5, we delivered daily BrdU injections (P1 – P5) and performed co-labelling of OLIG2 and BrdU at P5 and also after 5-days of recovery at P10 (Figure 3C, E). In contrast to the *snap-shot in time* PCNA analysis (OLIG2+/PCNA+) which showed a reduction caused by systemic inflammation exposure, the absolute counts for OLIG2+/BrdU+ cells showed no effect of systemic inflammation (Figure 3C). However, when expressed as a ratio to all OLIG2+ cells, the OLIG2+/BrdU+ population agreed with the PCNA data and revealed significantly reduced proportions of OLIG2+/PCNA+ cells in animals with systemic inflammation (Figure 3D).

### Microglia were the predominant CD11B cell type in the cerebellum in control mice and in mice exposed to systemic inflammation

To explore what cell types were responding to systemic inflammation we undertook cell type characterization of the CD11B+ expressing cells at P5 by FACS (Figure 4). This analysis demonstrated high enrichment of CD11B+CD45^lo^microglia in the cerebellum (97% and 88% in PBS and IL-1β exposed mice, respectively) and also in the cerebrum (98% and 90% in PBS and IL-1β group, respectively) (Figure 4A-B). Notably, the percentage of CD11B+/CD45^lo^ microglia was 6 times lower in the cerebellum than in the cerebrum (Figure 4B). The numbers of CD11+/CD45^hi^ macrophage^69^ in both brain structures increased slightly with IL-1β injections (Figure 4A-B). This increase reflects modest myeloid cell infiltration including neutrophils, monocytes and macrophages, as previously described in our model^27^. Significant increase of the Mean Fluorescence Intensities (MFIs) of CD11B and CD18, which form the phagocytic receptors CR3 (complement receptor 3) in CD11B+CD45^lo^ microglia in the cerebellum and also in the cerebrum of IL-1β exposed mice clearly indicated microglial activation (Figure 4A, C). Of note, the MFI of CD11B and CD18 in mice exposed to systemic inflammation was significantly higher in the cerebellum than in the cerebrum, suggesting a higher activation of cerebellar microglia compared with those in the cerebrum (Figure 4C). In addition, we undertook immunostaining for CD3 in P10 cerebellar tissue to rule out leucocyte/lymphocyte origin of the cells found after IL-1β injections. Very few CD3+ cells (between 1 and 2 cells per total cerebellum section) were identified in the cerebellum of both experimental groups (Supplementary Figure 5), supporting that there is minimal invasion of blood-derived cells in the cerebellum of the present model. This data indicates that microglia are the predominant phagocytic cells in the cerebellum during the time of insult.

**Figure 4:**
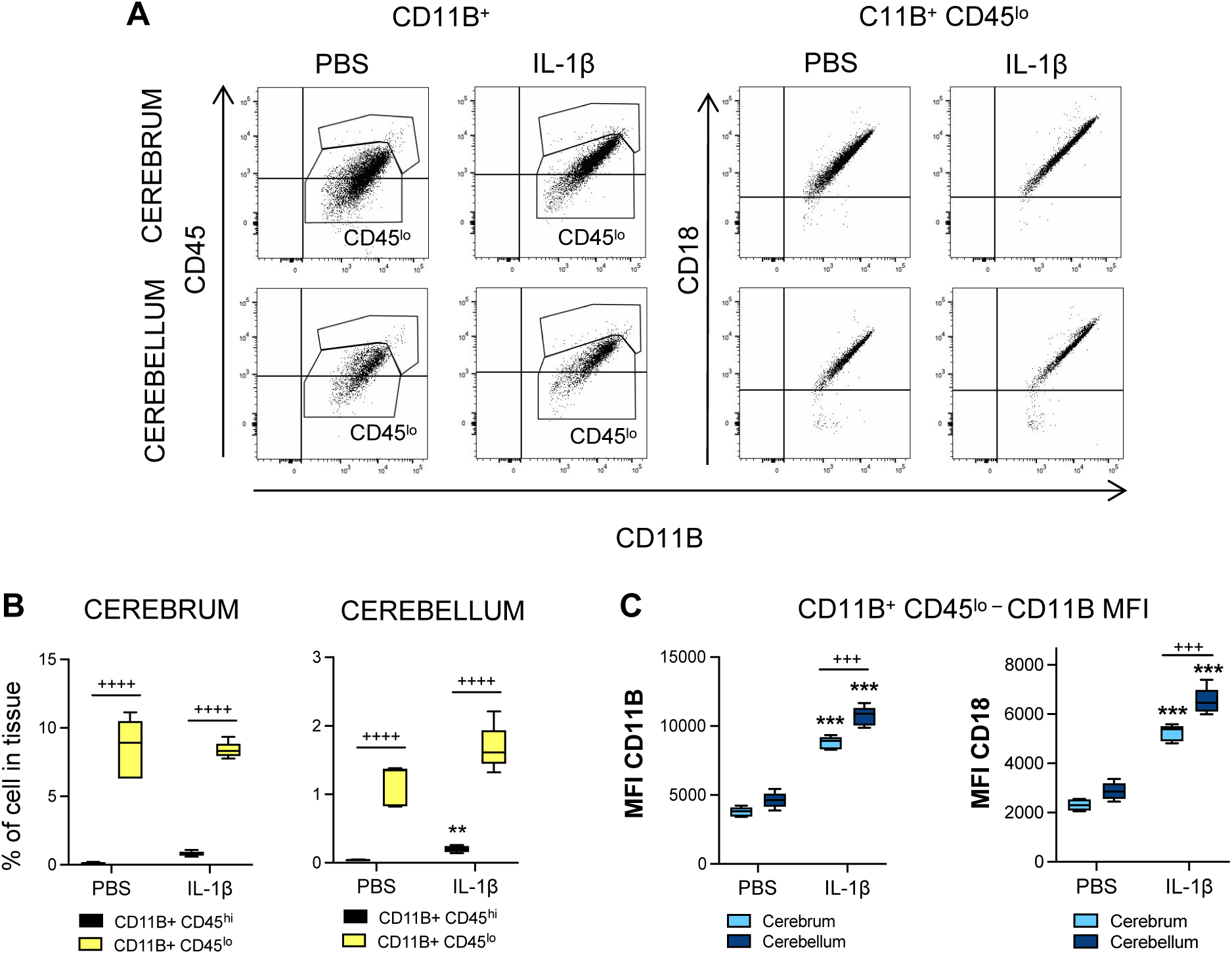
Impact of systemic inflammation on microglia. **A**: Representative picture of flow cytometry gating strategies to analyse brain CD11B+ and CD11B+/CD45^lo^ cells from the cerebrum and cerebellum of P5 IL-1β-treated and PBS treated mice and **B**, the quantification of this data. **C**: Quantification of dot plot analysis by flow cytometry of CD11B/CD18 MFI in CD11B+/CD45^lo^ microglia in cerebellum and cerebrum from P5 IL-1β-treated and control mice. Analysed via 2-Way ANOVA, ++++ = cerebrum vs cerebellum or CD45^hi^ versus CD45^lo^, p<0.001; **** = PBS versus IL-1 β, p<0.0001; *** = PBS versus IL-1 β, p<0.001; ** = PBS versus IL-1β, p<0.01.

### Systemic inflammation induced a sustained microglial activation in the cerebellum

In the present model, we have previously shown that microglia play a key role in the maturation blockade of oligodendrocytes in the cerebrum and that microglial activation displays a characteristic temporal profile in terms of number, morphology and gene expression^25, 27, 28^. We thus explored these microglial characteristics in the cerebellum in order to compare with the cerebrum data.

We analysed IBA1+ microglial cell numbers at P3 and P5 during the exposure to systemic inflammatory challenge, at the juvenile age P45, and at the adult age of P150. In IL-1β-exposed mice, IBA1+ numbers were significantly higher at P3, P5 and even out to P45 in comparison to controls (p<0.01) (Figure 5A-B); there was a non-significant trend at P10 (p=0.10, n=6). At P150, IBA+ numbers were comparable in both groups (Figure 5A-B).

**Figure 5:**
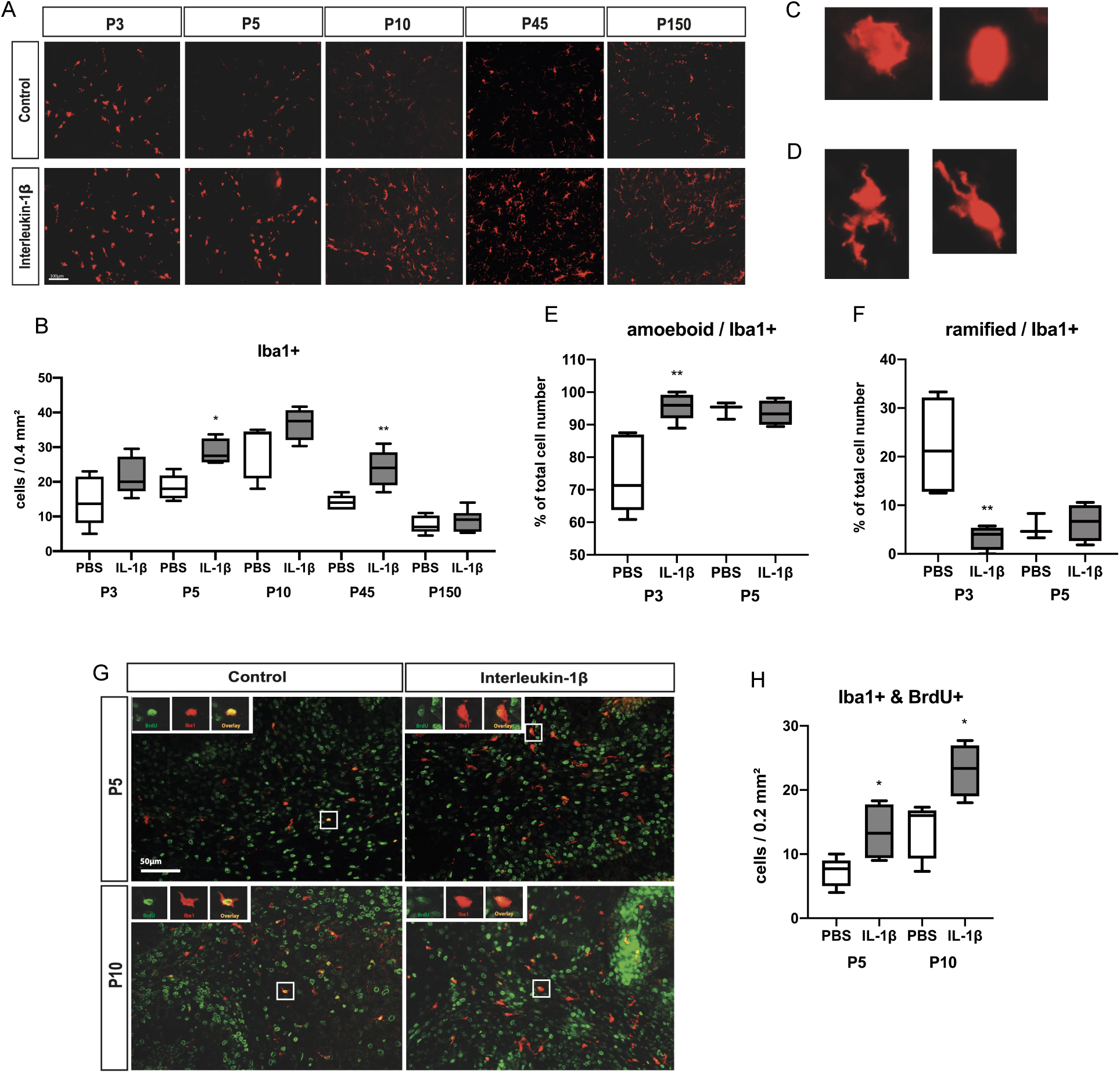
Impact of systemic inflammation on microglia activation and proliferation. **A-B**: Representative micrographs of IBA1 at P3, P5, P10, P45, and P160 in cerebellar sections from PBS vs IL-1β-exposed pups (A; Bar 100 µm) and corresponding quantification (B; *p<0.05, **p<0.01; t-test). **C-D**: Representative micrographs of ameboid (C) and ramified (D) IBA1+ microglia observed at P3 and P5, with corresponding quantification (E and F, **p<0.01, t-test). **G-H**: Representative micrographs of IBA1+ (green) and BRDU+ (red) cells at P5 and P10 in cerebellar sections from PBS vs IL-1β-exposed pups (G; Bar 50 µm) and corresponding quantification (H; *p<0.05; t-test).

Microglia can be schematically described as having an amoeboid, ramified, or intermediate linked to distinct functional activities and although this is a simplification, it captures significant shifts in their functional state^70^. An amoeboid morphology (representative image Figure 5C) is observed during (i) early brain development and it reflects the ongoing proliferation, migration and integration of these cells at that time^71, 72^, or (ii) during a pro-typical pro-inflammatory response wherein they are also often proliferating but performing immune functions. A ramified morphology (a surveying microglia), is observed in the uninjured adult brain and is characterized by small round soma with numerous processes (representative image Figure 5D). When we assessed morphology at P3 and P5 in control (PBS exposed) pups, respectively 71.32 ± 5.33 % and 95.40 ± 1.50 % of IBA1+ cells were of amoeboid morphology (Figure 5E). However, at P3, in pups receiving IL-1β injections the proportion of amoeboid IBA1+ microglia increased more than 20% to 95.97 ± 1.90 % (Figure 5E). Conversely, in P3 PBS exposed mice, 21.14 ± 5.28 % of all IBA1+ cells were ramified, but IL-1β exposure drastically reduced the number to 4.03 ± 1.08 % (Figure 5F). As we had observed increased numbers and more amoeboid morphology of microglia we expected to also find increased proliferation. Indeed, there was a large increase in the number IBA1+/BRDU+ cells in the cerebellum at P5, at the end of IL-1β exposure period, and also at P10, after 5 days of recovery, when compared to control pups (Figure 5G-H). These data altogether show a significant and sustained change in microglia in the cerebellum caused by exposure to systemic inflammation.

### The cerebellar microglial transcriptome in response to systemic inflammation had a specific IFN-pathway signature

Our previous work has shown elaborate transcriptomics changes in microglia linked to oligodendrocyte injury in the cerebrum^25, 28^. To measure this transcriptional profile and determine if the response was the same in the cerebellum and the cerebrum, we analysed the transcriptomic response of both populations of microglia to the inflammatory stimulus. Specifically, we isolated CD11B+ cells (primarily microglia, Figure 4) of the cerebrum and the cerebellum by magnetic cell sorting followed by transcriptome analysis with RNAseq at P5.

Transcriptomic analysis revealed a higher number of differentially expressed (DE) genes induced by IL-1β exposure in the cerebrum than in the cerebellum (3860 and 2523, respectively, with an adjusted p value < 0.05; Figure 6A, Supplementary Table 3). Analysis of microglial phenotype by quantification of validated markers^28, 73^ from normalized read count matrix of transcriptomic analysis (Supplementary Table 4) revealed that in a basal state (PBS only) that microglia from the cerebellum expressed significantly different levels of expression across phenotype associated genes. Of note, the gene for pro-inflammatory associated interleukin 6 (*Il6*) was only 35% lower than that observed in the cerebrum, expression of the gene for anti-inflammatory associated Arginase 1 (*Arg1*) was 90% lower than that observed in the cerebrum, the gene for immunomodulatory interleukin 1 receptor antagonist (*Il1rn*) was 40% higher than the observed in the cerebrum and the gene for immunomodulatory Sphingosine kinase-1 (*Sphk1*) was 70% lower than that observed in the cerebrum. When comparing the response to inflammatory stimuli, this revealed a similar microglial phenotype with significant increases of pro-inflammatory (inducible nitric oxide synthase, *Nos2*, and *Il6* mRNA), immuno-regulator markers (interleukin-4 receptor antagonist, *Il4ra,* and *Il1rn* mRNA), and anti-inflammatory galectin-3 (*Lgals3* mRNA) mRNA marker. In the cerebellum, there was a significant up-regulation of the immuno-regulator marker induced by IL-1β (suppressor of cytokines 3, *Socs3* mRNA) (Figure 6B, Supplementary Table 4).

**Figure 6:**
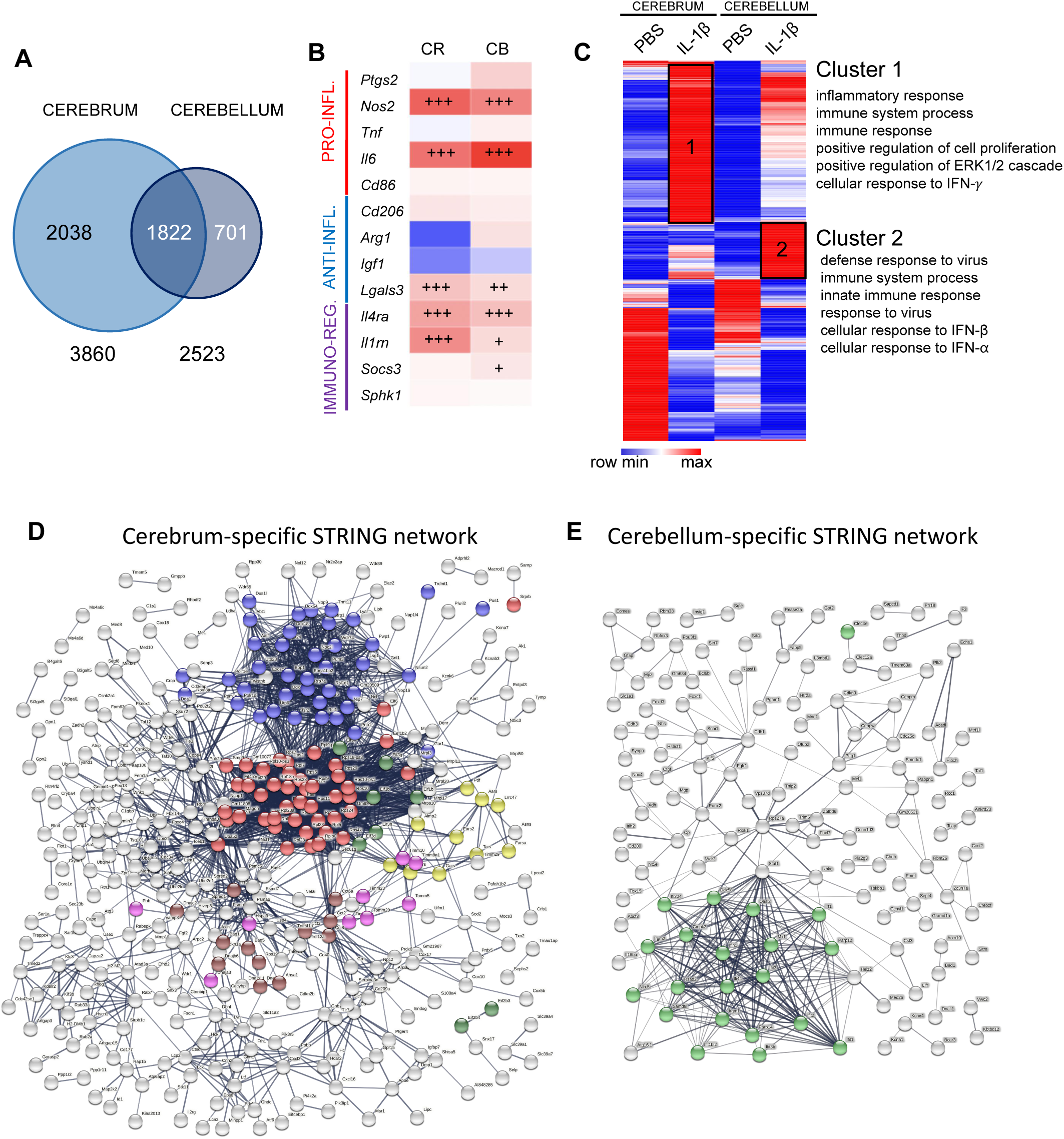
Region specific impact of systemic inflammation on the microglial transcriptome. **A**: Venn diagram of Differentially Expressed (DE) genes in CD11B+ microglia in cerebrum and cerebellum from P5 IL-1 β-treated and control mice. **B**: mRNA expression of gene associated with different phenotypes in the cerebrum and cerebellum: pro-inflammatory (red), anti-inflammatory (blue) and immuno-regulatory (purple) markers (mRNA values in Supplementary Table 4). +p<0.05, ++p<0.01,+++p<0.001; t-test **F**: Heatmap of DE genes in cerebrum and cerebellum, and the Gene Ontology terms significantly associated to cluster 1 and 2. **D-E**. Network representation from STRING showing cerebrum exclusive and cerebellum exclusive networks of predicted protein interactions. Coloured dots indicate protein members of selected significantly (q<0.05) enriched pathways. In (**D**), red, blue, yellow, green for translation processes (107/676 nodes); brown for heat shock responses (12/676); and pink for mitochondrial import processes (10/676). In (**E**), green for viral or interferon signalling (23/218 nodes). Note, viral or interferon signalling was not significantly enriched in (D).

To explore the regional differences and responses to inflammation a heatmap was generated from DE genes induced by IL-1β exposure in the cerebrum and/or in the cerebellum with a minimum of 2-fold increase (LogFC>1) or 2-fold decrease (LogFC<1), representing 929 genes (Figure 6C, Supplementary Table 5). One minus Pearson’s correlation revealed region-specific transcriptional identities of microglia from control (PBS-treated) mice and those subjected to systemic inflammation (IL-1β-treated). IL-1β-treatment revealed two relevant gene clusters with significant enrichment in Gene Ontology (GO) Biological Processes (BP), Cluster 1 was composed of genes with a higher expression level in the cerebrum than in the cerebellum but with a similar fold change between PBS and IL-1β in both structures. Cluster 2 was composed of genes with a strong level of induction under IL-1β in the cerebellum compared to the cerebrum. Functional enrichment analysis using DAVID 6.8^74^ revealed that Cluster 1 was enriched for GO terms related to inflammation, cell proliferation, extracellular signal-regulated protein kinase (ERK) 1/2 and interferon (IFN)-γ while Cluster 2 was enriched for GO terms related to response to virus, IFN-β and IFN-α, suggesting additional IFN-pathway activation specifically in cerebellar microglia (Figure 6C, Supplementary Table 6).

### Cell-location specific analysis confirmed a unique type-II interferon response in the cerebellar microglia

To explore the specific differences between the response of the cerebrum and cerebella microglia we identified from our DEG lists what genes were exclusively altered in each population of microglia and also created a list of commonly dysregulated genes. We explored these cell-location-exclusive and overlapping DEG lists using STRING and Wiki Pathways. STRING converted our gene data into known proteins and queried databases of established protein-protein interactions (PPI) to build networks (Supplementary Table 7).

The PPI network for each of the three gene lists had significant (p<1.0e^-16^) connectivity enrichment based on its size (adjusting for innate shell interactions) shown in Figure 6D-E and Supplementary Figure 6. Connectivity enrichment suggests the proteins have biological interactions compared with data sets generated from randomly selected proteins. For the shared gene list (Supplementary Figure 6, Supplementary Table 7), there was a significant enrichment (q<0.05) in 71 known local network cluster groups, the vast majority (41/71) of which were directly related to ribosomes, proliferation, cytoskeleton, chromosomes, DNA replication or chromatin structure. Of note, 14/71 were related to mixed viral defence or inflammation, and of these 4/71 specifically to IFNB signalling. For the cerebrum, specific network there was a significant enrichment (q<0.05, Figure 6D) in 54 known local network cluster groups, the vast majority (39/54) of which were directly related to protein synthesis, especially ribosome function and none were related to viral responses. However, for the PPI network for cerebellum exclusive genes there was a significant enrichment (q<0.05, Figure 6E) in 12 known local network cluster groups, all of which were related specifically to interferon or vial signalling.

When we sought to determine in more detail the differences in the specific lists by querying the Wiki Pathway database (mouse). For the shared gene list, we identified five pathways significantly enriched (q<0.05) and all of these related to cell cycle, replication or cell survival (Supplementary Table 8) but none to viral responses. Our cell type exclusive lists returned only one pathway for each cell type. For the cerebrum data, in agreement with our observations from our PPI network in STRING, there was enrichment in the Cytoplasmic Ribosomal Proteins (WP163) pathway (Supplementary Table 8). Also supporting our data from STRING for the cerebellum data was that this analysis revealed that there was a significant enrichment in the Type II interferon signalling (IFNG, WP1253) pathway in the cerebellum microglia data (Supplementary Table 8).

## Discussion

Despite the prevalence of cerebellar injury in preterm born infants ^5, 6, 24, 75, 76^ and the links between these insults and poor outcomes in preterm born infants ^77, 78^, there is not many studies focused on the cerebellum in models of preterm brain injury. We have addressed this issue and we have documented by MRI that in our model of inflammation-associated encephalopathy of prematurity that there is a global change in absolute volumes of the mouse brain, with a specific loss of cerebellar volume into adulthood. These volumetric changes were accompanied by deficits in oligodendroglial proliferation and maturation, and myelination. A sustained increase in microglial density and proliferation was also noted so we explored the transcriptome of these cerebellar microglia and compared and contrasted the responses to those of cerebrum microglia. The transcriptome analysis in P5 isolated cerebellar microglia revealed a unique pattern of in the transcriptomic profile towards interferon type II pathway-mediated inflammatory and immunological programs compared with the microglia response in cerebrum.

Systemic inflammation in the fetal and neonatal period plays a key role in disturbance of brain development as shown clinically^79–81^ and supported by pre-clinical models^25, 82, 83^, reviewed in^41, 84^. In clinical studies in preterm infants, where exposure to inflammation is very common, MRI anomalies of the cerebellum are correlated with lower outcome scores in neurodevelopmental assessments^85^. Also, lower volume of the cerebellum at term age equivalent was associated with reduced IQ and with poor language and motor performance at school age^19^. Of note, in a classical eyeblink conditioning test, former preterm infants showed cerebellar functional deficits even when they displayed mild or no cerebellar lesions^77^ highlighting the importance of models with a mild phenotype. In this context, a postmortem study of the cerebellum of preterm infants without overt brain and cerebellar destructive damage revealed a significant cerebellar maldevelopment^86^. Our model also produces no overt destructive lesions, cell death itself is very low and the primary injury is driven by dysmaturation^25, 27, 40^. Although infrequent there are studies on focused on the impact of perinatal systemic inflammation on the development of the cerebellum^87, 88^ or that include an analysis of this region^82^. Dean *at al* exposed fetal sheep at 93-96d GA once to 90 ng/kg of LPS and analysed the brain three days later. Tolcos *et al*., exposed fetal sheep at 102d GA once to the same dose of LPS (90 ng/kg) and analysed the brain nine days later. Interestingly, both groups reported cerebellar white matter injury without cell death, in agreement with our current work. However, in the model from Dean *et al* (93-96d GA, +3-day analysis) there were no cerebellar grey matter changes but in the work from Tolcos *et al* (102d GA sheep, +9-day analysis) there were increased densities of mature neurons and the proliferative area of the external granule cell layer was increased. We have previously shown that the time between inflammatory exposure and analysis significantly impacts forebrain outcomes in a sheep model of inflammatory injury^83^. Overall however, our data of decreased cerebellar grey matter volumes supports the previous findings of a vulnerability of the developing grey matter to inflammatory injury reported by Tolcos *et al*., and our findings of reduced molecular layer area in a mouse model of repeated exposure to a specific toll-like receptor (TLR)-2 agonist (Pam3CysSerLys4, Pam3CSK4)^82^. Volume loss of the cerebellum has also been found in a chronic inflammation model caused by GFAP driven transgenic IL-6 production^89^. In this model, elevated number of activated microglia was associated with lower cerebellar volumes at age 3 months, hence underlining the vulnerability of the developing cerebellum to inflammation.

Available data suggest that the density of microglia is lower in the developing cerebellum when compared to the developing forebrain^90^. Our FACS data in the present study support this previous report. However, transient systemic inflammation caused a pronounced and sustained increase of the number of microglia in the cerebellum, characterized by higher number of amoeboid shaped microglia. These cerebellar data are in contrast to our previous data from the cerebrum in this inflammation model, where we did not observe a sustained change in microglia number (numbers returned to normal by P10) or robust change in morphology at any time point^25, 27, 28^. Morphology is often used as a surrogate indictor of the relative level of microglia ‘activation’. However, we have observed comparable levels of injury to oligodendrocytes in the cerebellum in this study and cerebrum in previous work from this model^25, 40^ but this effect is independent of microglia morphology, so further work is required to interpret this finding. The sustained alterations of microglia in the cerebellum could participate to the so-called tertiary long-lasting phase following an acute brain insult^91^ a persisting set of changes that may sensitise to injury or prevent repair that might also underpin why brain volume changes became greater in magnitude with increasing age in this model. Altogether, these data support the hypothesis that microglia from different brain structures respond differently to inflammatory stimuli, shown previously for the grey and white matter in the cerebrum in adults^92^.

As indicated above, microglia are drivers of injury and we have previously demonstrated a causal relationship between pro-inflammatory microglia activation and hypomyelination in this model^28^. Microglia so also play an important role in the maturation and maintenance of oligodendrocyte cell populations^93, 94^. As such, our increased numbers and altered activation of the cerebellar microglial after systemic inflammation is likely responsible for the oligodendrocyte dysmaturation and myelination deficit we observed via direct effects that injure these cells and possibly via disrupted development. This injury to oligodendrocytes was demonstrable as changes in oligodendrocyte maturation and myelin, and also reduced cerebellar white matter volumes on MRI. We cannot also rule out a contribution from other cell populations (i.e., astrocytes, NG2 glia), cell compartments (i.e., axons) and/or extracellular matrix protein abundance. In regards to the reduction of grey matter volumes on MRI, further studies will be needed to validate our hypotheses of the presence of alterations in: i) microglia mediated changes in synaptic and neurite pruning, as these are key functions of microglia^95–97^, and; ii) changes in the density of some neuronal subpopulations, as previously we have shown in this model changes in subclasses of interneurons^42^ and in neuronal-associated gene expression in the cerebrum^41^ and in another inflammatory model volumetric reductions in the molecular layer^82^.

When we investigated the transcriptomics response of microglia systemic-inflammation the cerebrum and cerebellum have a common “activated” signature with an over-representation of terms related to inflammation and cell proliferation. However, analysis of the data across multiple approaches revealed that IFN signalling and especially and type II IFN (IFN-γ) signalling, is specific to the cerebellum and not significantly observed in the cerebrum. This specific response of cerebellar microglia could potentially contribute to long-term neurodevelopmental sequelae of prematurity, including ASD. Indeed, clinical studies have shown that cerebellar abnormalities on MRI of preterm infants are a strong predictor of ASD in this high-risk population, especially in the context of chorioamnionitis^24, 98, 99^. A high activity of IFN-γ has been found in children with ASD in comparison to healthy controls^100^ or to children with non-ASD developmental delay^101^. Treatment of neural progenitors derived from human induced pluripotent stem cells with IFN-γ increased neurite outgrowth in these cells while inducing the expression of genes that have a role in schizophrenia and ASD, hence suggesting convergence of genetic and environmental factors^102^. The specific overexpression of *Socs3* mRNA found in cerebellar microglia in the present study is in agreement with the fact that IFN-induced signalling is regulated via SOCS proteins^103^. Type I IFN is known to mediate recruitment of inflammatory monocytes^104^ suggesting that it would be relevant to realize in the future an in-depth study of myeloid cells infiltrated into the cerebellum in the present model. Altered type I IFN signalling in cerebellar microglia could also explain modification of microglia proliferation observed in the cerebellum in the present study as type I IFN is an important regulator of microglia proliferation^105^.

The cerebellum is important for motor performance and memory, cognition, and emotion, this region is vulnerable to injury in preterm born infants and these infants have increased rates of neurodevelopmental disorders. As such, this project investigates the characteristics of cerebellar impairment after perinatal systemic inflammation to decipher the underlying cellular and molecular mechanisms for the long-term goal of finding ways to improve outcomes in preterm infants. When compared to cerebrum microglia, our study provides novel and intriguing data showing a strikingly sustained microglial activation (tertiary phase microglial dysfunction) combined with a specific microglia transcriptomic profile including altered IFN type I signalling in the cerebellum following neonatal systemic inflammation. This unique pattern of cerebellar microglia activation might play a key role in the pathophysiology of cognitive and behavioural consequences of prematurity, making these cells a potential target for neuroprotection.

## Declaration of authorship

LK, JVS, BF, ChB, VF, DAV, ACV, PG, and ThS designed the research. LK, JVS, BF, TiS, VF, LS, ZC, and ACV performed the research. SL and CoB performed the RNAseq. LK, JVS, BF, TiS, ChB, VF, SL, CoB, LS, ZC, DAV, ACV, PG, and ThS analyzed the data. LK, JVS, BF, ACV, PG, and ThS wrote the manuscript.

## Data availability

All new data are available from the authors on reasonable request.

## Acknowledgements

ThS, TiS and LK’s research was supported by funding from Deutsche Forschungsgemeinschaft – DFG, SCHM3007/3-2 and SCHE 2078/2-1, and from Förderverein für frühgeborene Kinder an der Charité e.V. ACV acknowledges financial support for this study from the Medical Research Council (New Investigator Research Grant (NIRG), MR/N025377/1). The work (at King’s College, London) was also supported by the Medical Research Council (MRC) Centre grant (MR/N026063/1). PG, JVS, BF, LS, and ZC’s research was supported by funding from Inserm, Université de Paris, Fondation de France, Fondation pour la Recherche sur le Cerveau, Fondation Grace de Monaco, Horizon 2020 Framework Program of the European Union (grant agreement no. 874721/PREMSTEM), and an additional grant from Investissement d’Avenir-ANR-11-INBS-0011 NeurATRIS. BF’s research was also funded by the Cerebral Palsy Alliance, Australia.

## Legends of Supplementary Figures

**Supplementary Figure 1: Timeline for injections and analyses. A**: Mice were administered IL-1β (10µg/kg/injection) or PBS twice a day for 4 days and once on day 5. Analyses were performed at various time points. **B**: Animals were administered IL-1β or PBS for 5 days as shown in A. Additionally, BrdU was injected intraperitoneally. PBS, phosphate buffered saline; IHC, immunohistochemistry; WB, western blot; PCR, polymerase chain reaction.

**Supplementary Figure 2: Impact of systemic inflammation on whole brain volume.** Whole brain volume at P15 and P60 in PBS and IL-1β-exposed mice as measured using MRI. Horizontal line indicates group mean. *p<0.05 post-hoc contrast corrected for multiple comparisons (2-step FDR method at 5%); ns, not significant.

**Supplementary Figure 3: Native Western blots.** Western blotting of MAG and MBP of P10, P15, and P60 cerebellar tissues from PBS and IL-1β-exposed mice.

**Supplementary Figure 4: Impact of systemic inflammation on oligodendrocyte cell death. A**: Representative micrographs of cerebellar slices showing OLIG2+ (green) and CASP3+ (green) cells in PBS and IL-1β-exposed P5 pups (DAPI counterstaining in blue). **B**: Quantification of OLIG2+CASP3+ double-stained cells at P3, P5 and P10 in cerebellum of PBS and IL-1β-exposed P5 pups.

**Supplementary Figure 5: Impact of systemic inflammation on blood-borne CD3+ brain invasion.** Representative micrographs of cerebellar slices (DAPI staining in blue) at P3 and P5 showing anecdotic CD3+ cells (red) in PBS and IL-1β-exposed pups.

**Supplementary Figure 6. Network representation from STRING showing the predicted protein interactions network built with the shared gene list.** Coloured dots indicate protein members of selected significantly (q<0.05) enriched pathways. Red = ribosome biogenesis, Blue = mixed anti-viral defence, Magenta = DNA replication, Green = transmembrane receptor tyrosine kinase, yellow = kinetochores signal amplification (cell division).

## Supplementary Tables

**Supplementary Table 1**: List of antibodies and primers. (**A**) Primary antibodies for immunohistochemistry. (**B**) Secondary antibodies for immunohistochemistry. (**C**) Sequences of nucleotides and gene loci.

**Supplementary Table 2:** Atlas-based segmentation dataset, including absolute and relative volumes and statistical analysis of data from PBS and IL-1β exposed mice.

**Supplementary Table 3:** Differentially expressed gene (DEG) lists from RNAseq analysis of CD11B+ microglia from the cerebrum and cerebellum.

**Supplementary Table 4:** mRNA values of phenotype associated genes used to create Figure 6B.

**Supplementary Table 5:** Gene list used to build the DEG heat map and analysis for clusters in Figure 6C.

**Supplementary Table 6:** Outputs from the cluster-based gene analysis.

**Supplementary Table 7**: Specific gene lists used to build the predicted protein networks in STRING.

**Supplementary Table 8:** Wiki pathways outputs from the cerebellum only, cerebrum only and shared genes lists presenting the significantly enriched pathways.

