## Supplemental Figures for "A unique cerebellar pattern of microglia activation in a mouse model of encephalopathy of prematurity"

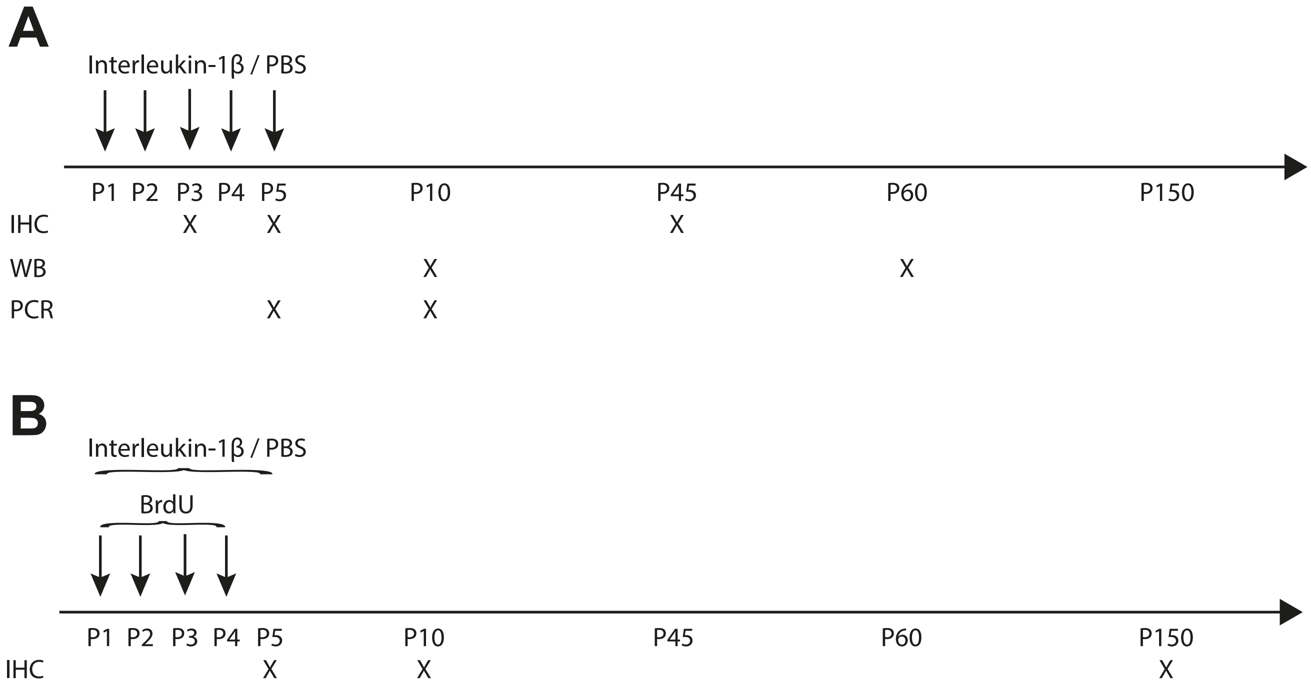


Supplementary Figure 1: Timeline for injections and analyses. A: Mice were administered IL-1β (10μg/kg/injection) or PBS twice a day for 4 days and once on day 5. Analyses were performed at various time points. B: Animals wer e administered IL-1β or PBS for 5 days as shown in A. Additionally, BrdU was injected intraperitoneally.
PBS, phosphate buffered saline; IHC, immunohistochemistry; WB, western blot; PCR, polymerase chain reaction.

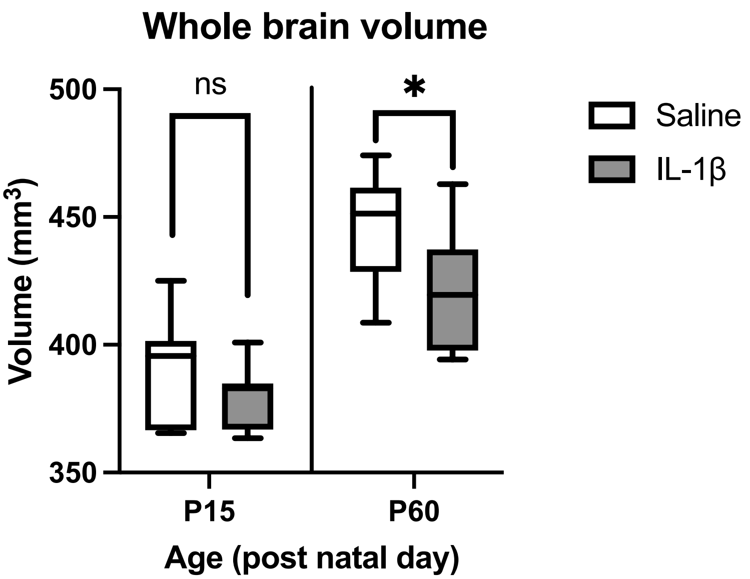


**Supplementary Figure 2: Impact of systemic inflammation on whole brain volume.** Whole brain volume at P15 and P60 in PBS and IL-1β-exposed mice as measured using MRI. Horizontal line indicates group mean. *$p<0.05$ post-hoc contrast corrected for multiple comparisons (2-step FDR method at 5%); ns, not significant.

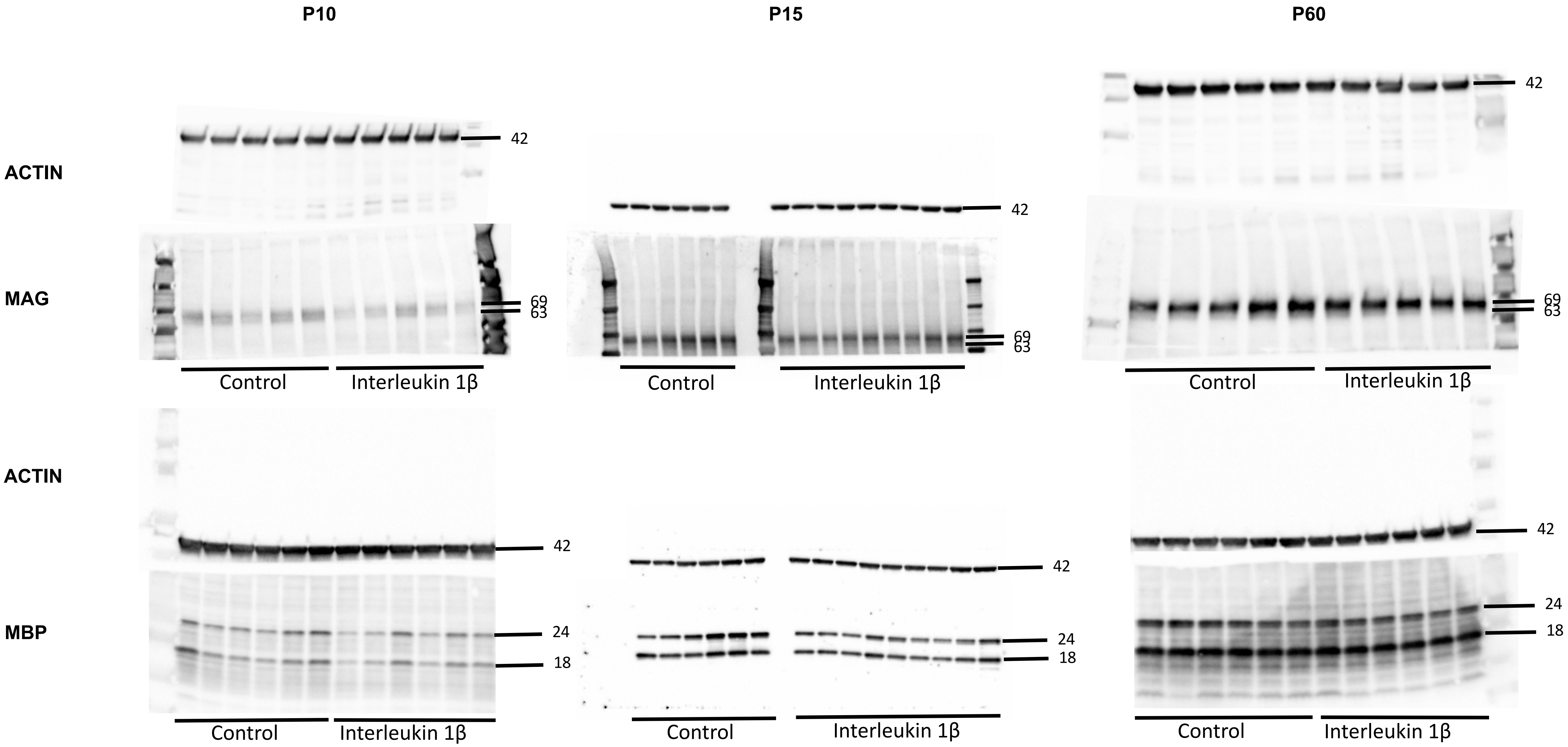


**Supplementary Figure 3: Native Western blots.** Western blotting of MAG and MBP of P10, P15, and P60 cerebellar tissues from PBS and IL-1β-exposed mice.

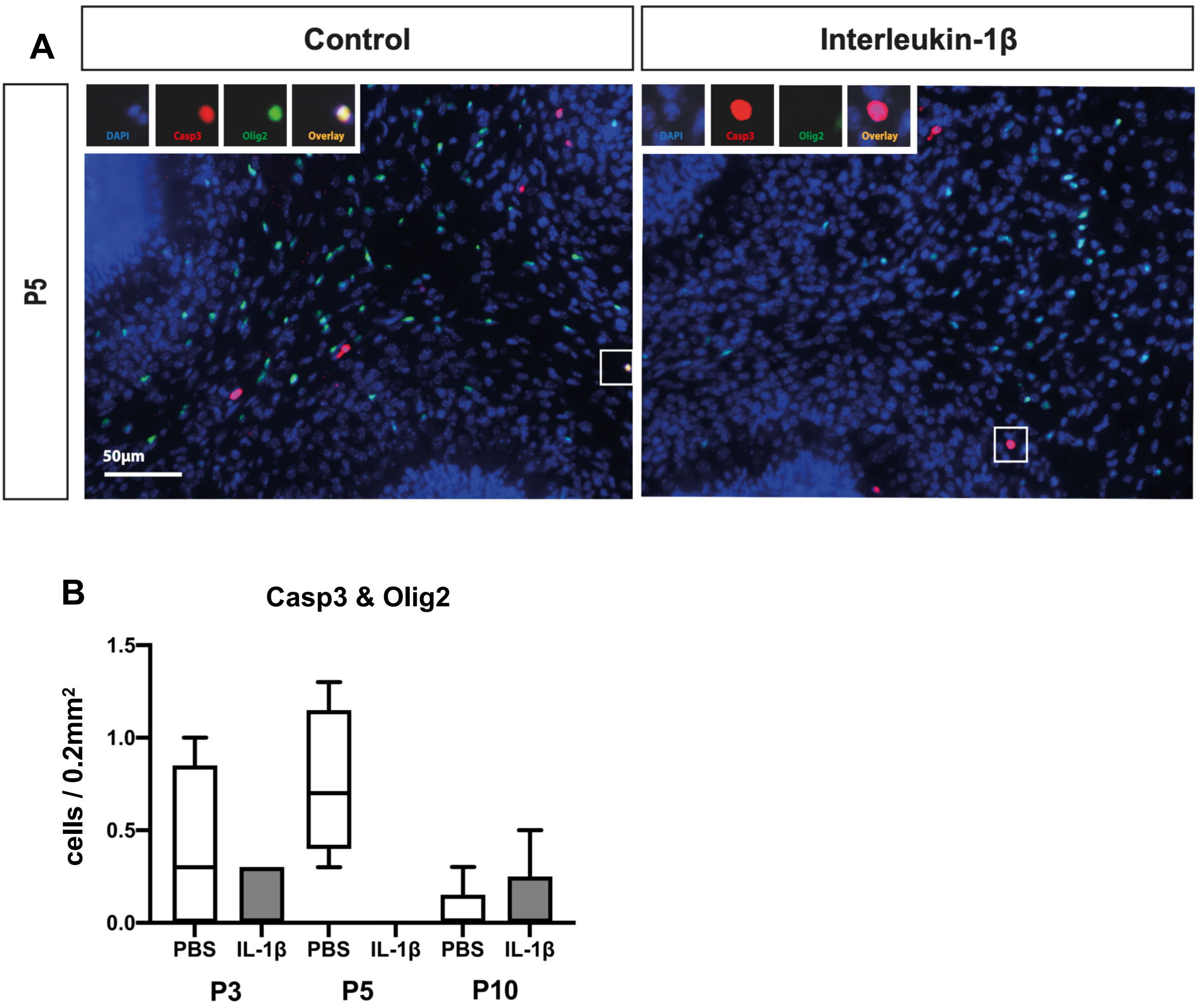


**Supplementary Figure 4: Impact of systemic inflammation on oligodendrocyte cell death. A**: Representative micrographs of cerebellar slices showing OLIG2+ (green) and CASP3+ (green) cells in PBS and IL-1β-exposed P5 pups (DAPI counterstaining in blue). **B**: Quantification of OLIG2+CASP3+ double-stained cells at P3, P5 and P10 in cerebellum of PBS and IL-1β-exposed P5 pups.

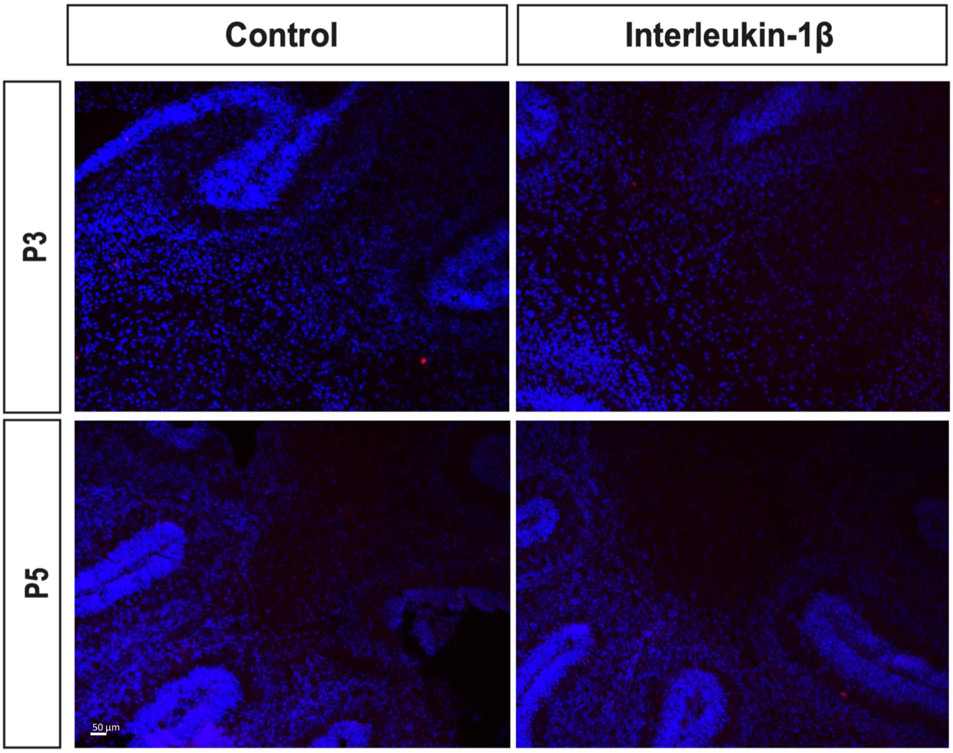


**Supplementary Figure 5: Impact of systemic inflammation on blood-borne CD3+ brain invasion.** Representative micrographs of cerebellar slices (DAPI staining in blue) at P3 and P5 showing anecdotic CD3+ cells (red) in PBS and IL-1β-exposed pups.

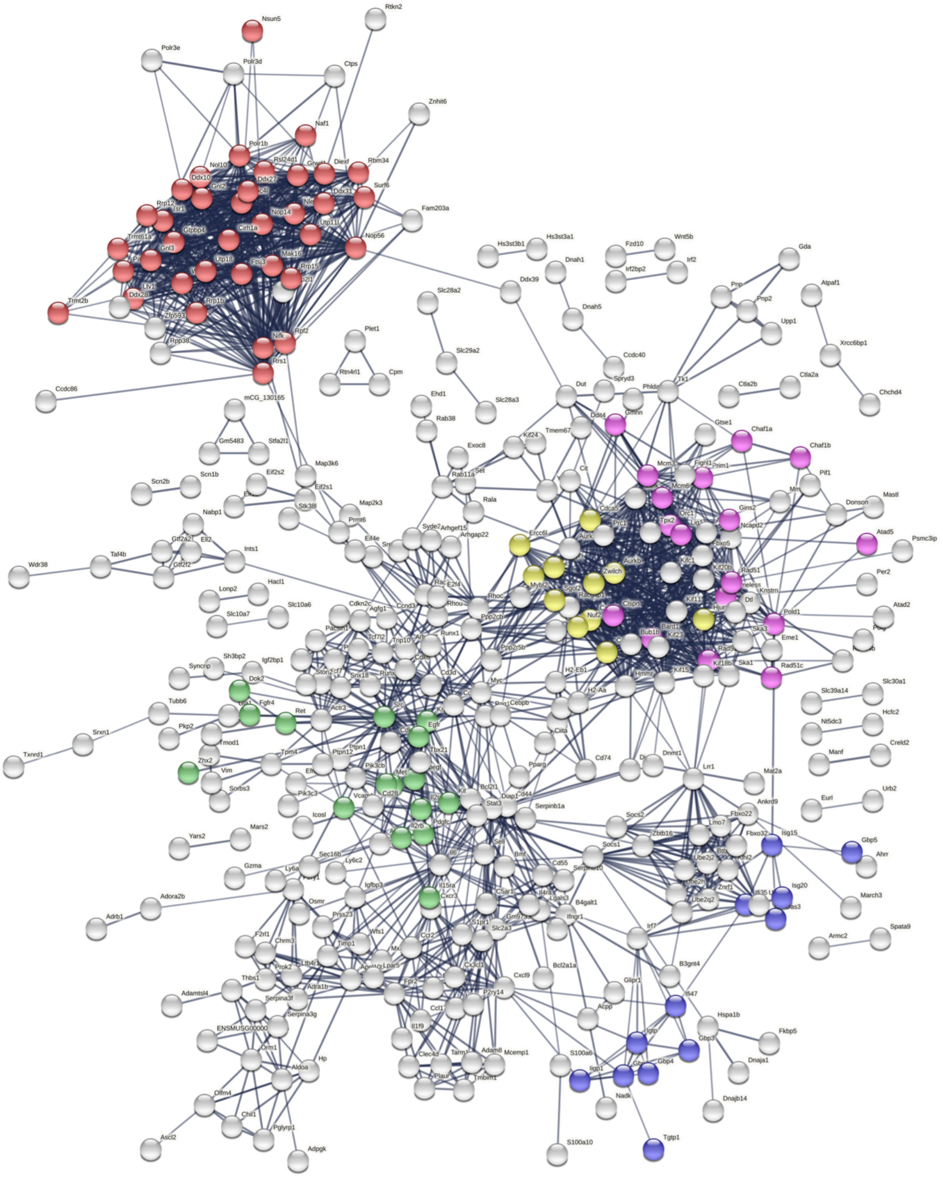

**Supplementary Figure 6. Network representation from STRING showing the predicted protein interactions network built with the shared gene list.** Coloured dots indicate protein members of selected significantly (q<0.05) enriched pathways.  Red = ribosome biogenesis, Blue = mixed anti-viral defence, Magenta = DNA replication, Green = transmembrane receptor tyrosine kinase, yellow = kinetochore signal amplification (cell division).
