## Supplemental Tables for "A unique cerebellar pattern of microglia activation in a mouse model of encephalopathy of prematurity"

**Supplementary table 1: List of antibodies and primers**

**Table 1A: Primary antibodies for immunohistochemistry**

| Antigen | Specification | Company | Number | Dilution | Incubation Time |
|---|---|---|---|---|---|
| PAX6 | Mouse Anti Pax6 | Millipore | MAB5552 | 1:100 | 24h |
| KI67 | Rabbit Anti Ki67 | Abcam | Ab16667 | 1:200 | 24h |
| OLIG2 | Mouse Anti Olig2 | Abcam | Ab56643 | 1:500 | 24h |
| OLIG1 | Rabbit Anti Olig1 | Millipore | AB15620 | 1:750 | 24h |
| PCNA | Mouse Anti PCNA | Abcam | Ab29 | 1:1000 | 24h |
| PCNA | Rabbit Anti PCNA | Abcam | Ab15497 | 1:500 | 24h |
| CASP3 | Rabbit Anti Casp3 | Cell Signaling | 9664 | 1:200 | 24h |
| IBA1 | Rabbit Anti Iba1 | Wako Chemicals | 019-19741 | 1:750 | 48h |
| MAG | Mouse Anti MAG | Abcam | Ab89780 | 1:1000 | 24h |
| MOG | Rabbit Anti MOG | Abcam | Ab32760 | 1:1500 | 24h |
| BRDU | Rabbit Anti Brdu | Bioss | Bs-0489R | 1:50 | 24h |
| BRDU | Chicken Anti Brdu | Abcam | Ab92837 | 1:300 | 48h |
| APC | Mouse Anti CC1 | Calbiochem | OP86 | 1:250 | 24h |

PAX6, Paired Box 6; KI67, Marker of proliferation KI-67; OLIG2, Oligodendrocyte Transcription Factor 2; OLIG1, Oligodendrocyte Transcription Factor 1; PCNA, Proliferating Cell Nuclear Antigen; CASP3, Cleaved Caspase 3; IBA1, Ionized Calcium-Binding Adapter Molecule 1; MAG, Myelin-Associated Glycoprotein; MOG, Myelin Oligodendrocyte Glycoprotein; BRDU, Bromo-Desoxyuridine; APC, Anti-Adenomatous Polyposis Coli Clone 1.

**Table 1B: Secondary antibodies for immunohistochemistry**

| Antigen | Specification | Company | Number | Dilution | IT |
|---|---|---|---|---|---|
| Rabbit | Goat Anti Rabbit, Alexa Fluor, 488nm | Invitrogen | A11034 | 1:200 | 1h |
| Chicken | Goat Anti Chicken, Alexa Fluor, 594nm | Molecular Probes | A11042 | 1:200 | 1h |
| Mouse | Goat Anti Mouse, Alexa Fluor, 594nm | Molecular Probes | A11032 | 1:200 | 1h |
| Mouse | Goat Anti Mouse, Alexa Fluor, 488nm | Molecular Probes | A11029 | 1:200 | 1h |
| Rabbit | Goat Anti Rabbit, Alexa Fluor, 594nm | Molecular Probes | A11037 | 1:200 | 1h |
| Rabbit | Donkey Anti Rabbit, Alexa Fluor, 546nm | Molecular Probes | A10040 | 1:200 | 1h |

IT, incubation time

**Table 1c: Sequences of oligonucleotides and gene loci**

| Gene | Oligonucleotide Sequences 5'-3' | | Gene Bank Sequence |
|---|---|---|---|
| *Hprt* | Forward | TGCTGACCTGCTGGATTACATT | NM_013556.2 |
| | Reverse | TCCCCCGTTGACTGATCATT | |
| | Probe | TCAGACTGAAGAGCTACTG | |
| *Olig2* | Forward | ATCTTCCTCCAGCACCTCCT | NM_016967.2 |
| | Reverse | GGGCTCAGTCATCTGCTTCT | |
| | Probe | CCACGTCTTCCACCAAGAAA | |
| *Cnp* | Forward | AGCAGGAGGTGGTGAAGAGA | NM_001146318.1 |
| | Reverse | GATGGCTTGTCCAGATCA | |
| | Probe | TTTGTGACACCCAAGACAGC | |
| *Gclc* | Forward | GTGGAGGCCAATATGAGGAA | NM_010295.2 |
| | Reverse | CTGGGTTGGGTCTGTGTTCT | |
| | Probe | AGGCTCTCTGCACCATCACT | |
| *Sod2* | Forward | CCGAGGAGAAGTACCACGAG | NM_013671.3 |
| | Reverse | ATATGTCCCCCACCATTGAA | |
| | Probe | ACAACTCAGGTCGCTCTTCAG | |
| *H2-Ab (MchII)* | Forward | CATCAAGGCCGACCACATG | NM_008206.2 |
| | Reverse | TCCCCGTCAAATTCGTGTGT | |
| | Probe | CTTCTACCAATCTTACGACGCT | |
| *Nos2* | Forward | CATCGACCCGTCCACAGTATG | NM_001313922.1 |
| | Reverse | GAAATCCGATGTGGCCTTGT | |
| | Probe | CAGTGGAGAGATTTTGCATG | |
| *Igf1* | Forward | GCTGGTGGATGCTCTTCAGTT | NM_010512.5 |
| | Reverse | AGGTGCCCTCCGAATGCT | |
| | Probe | Detected With SYBR Green | |
| *Ccnd2* | Forward | TCGATGATTGCAACTGGAAG | NM_009829.3 |
| | Reverse | TCCACATCAGTGTGGGTGAT | |
| | Probe | ACACACTCACGTGTGATGCC | |

*Hprt,* hypoxanthine-guanine phosphoribosyl-transferase; *Olig2*, oligodendrocyte transcription factor 2; *Cnp*, 2',3'-cyclic nucleotide 3'-phosphodiesterase; *Gclc*, glutamate-cysteine ligase catalytic subunit; *Sod2*, superoxide dismutase 2; *H2-Ab,* MHC class II antigen; Nos2 = nitric oxide synthase 2; *Igf1*, insulin-like growth factor 1; *Ccnd2*, cyclin D2

**Supplementary table 2:**
**Atlas-based segmentation dataset, including absolute and relative volumes and statistical analysis of data from PBS and IL-1β exposed mice.**

| | Absolute volumes (mm3) | | | | | | Absolute volumes (mm3) | | | | | |
|---|---|---|---|---|---|---|---|---|---|---|---|---|
| | P15 | | | | | | P60 | | | | | |
| | Saline | | IL-1β | | | | Saline | | IL-1β | | | |
| Atlas ROI | Mean | SD | Mean | SD | % change | Effect (SD) | Mean | SD | Mean | SD | % change | Effect (SD) |
| amygdala | 8,797 | 0,667 | 8,265 | 0,833 | -6,0 | -0,80 | 10,107 | 0,673 | 9,602 | 0,614 | -5,0 | -0,75 |
| anterior commissure: pars anterior | 1,004 | 0,053 | 0,990 | 0,070 | -1,5 | -0,28 | 1,201 | 0,090 | 1,138 | 0,079 | -5,2 | -0,70 |
| anterior commissure: pars posterior | 0,342 | 0,020 | 0,326 | 0,027 | -4,5 | -0,78 | 0,417 | 0,035 | 0,399 | 0,020 | -4,2 | -0,50 |
| basal forebrain | 4,873 | 0,375 | 4,623 | 0,443 | -5,1 | -0,67 | 5,545 | 0,425 | 5,228 | 0,324 | -5,7 | -0,75 |
| bed nucleus of stria terminalis | 1,161 | 0,099 | 1,122 | 0,106 | -3,4 | -0,40 | 1,352 | 0,141 | 1,283 | 0,087 | -5,1 | -0,49 |
| cerebellar peduncle: inferior | 0,690 | 0,053 | 0,663 | 0,029 | -4,0 | -0,51 | 0,818 | 0,024 | 0,802 | 0,040 | -2,0 | -0,66 |
| cerebellar peduncle: middle | 0,977 | 0,060 | 0,944 | 0,045 | -3,4 | -0,56 | 1,249 | 0,082 | 1,166 | 0,098 | -6,7 | -1,02 |
| .cerebellar peduncle: superior | 0,781 | 0,070 | 0,744 | 0,043 | -4,8 | -0,53 | 0,993 | 0,094 | 0,921 | 0,061 | -7,3 | -0,77 |
| cerebral aqueduct | 0,477 | 0,023 | 0,468 | 0,044 | -1,9 | -0,40 | 0,529 | 0,053 | 0,470 | 0,041 | -11,2 | -1,12 |
| cerebral peduncle | 1,693 | 0,128 | 1,596 | 0,065 | -5,7 | -0,75 | 2,216 | 0,178 | 2,116 | 0,150 | -4,5 | -0,56 |
| colliculus: inferior | 4,798 | 0,366 | 4,648 | 0,170 | -3,1 | -0,41 | 5,755 | 0,392 | 5,412 | 0,306 | -6,0 | -0,87 |
| colliculus: superior | 7,749 | 0,615 | 7,472 | 0,331 | -3,6 | -0,45 | 8,933 | 0,528 | 8,515 | 0,579 | -4,7 | -0,79 |
| corpus callosum | 11,430 | 0,611 | 11,130 | 0,473 | -2,6 | -0,49 | 13,556 | 1,190 | 12,733 | 1,074 | -6,1 | -0,69 |
| corticospinal tract/pyramids | 1,309 | 0,142 | 1,262 | 0,080 | -3,6 | -0,33 | 1,562 | 0,077 | 1,539 | 0,090 | -1,5 | -0,30 |
| cuneate nucleus | 0,266 | 0,024 | 0,265 | 0,020 | -0,4 | -0,05 | 0,266 | 0,026 | 0,269 | 0,027 | 0,9 | 0,10 |
| dentate gyrus of hippocampus | 2,743 | 0,131 | 2,617 | 0,103 | -4,6 | -0,96 | 3,353 | 0,225 | 3,120 | 0,292 | -6,9 | -1,03 |
| facial nerve (cranial nerve 7) | 0,188 | 0,014 | 0,181 | 0,009 | -3,7 | -0,51 | 0,235 | 0,016 | 0,226 | 0,019 | -3,6 | -0,53 |
| fasciculus retroflexus | 0,232 | 0,013 | 0,227 | 0,010 | -2,5 | -0,44 | 0,280 | 0,025 | 0,262 | 0,018 | -6,4 | -0,72 |
| fimbria | 2,706 | 0,191 | 2,595 | 0,172 | -4,1 | -0,58 | 3,243 | 0,300 | 3,066 | 0,249 | -5,5 | -0,59 |
| fornix | 0,569 | 0,041 | 0,540 | 0,040 | -5,2 | -0,72 | 0,671 | 0,066 | 0,633 | 0,035 | -5,6 | -0,57 |
| fourth ventricle | 0,841 | 0,094 | 0,799 | 0,092 | -5,0 | -0,45 | 0,936 | 0,056 | 0,839 | 0,059 | -10,4 | -1,73 |

| | | | | | | | | | | | | |
|---|---|---|---|---|---|---|---|---|---|---|---|---|
| fundus of striatum | 0,163 | 0,011 | 0,152 | 0,013 | -7,2 | -1,03 | 0,186 | 0,016 | 0,176 | 0,013 | -5,2 | -0,59 |
| globus pallidus | 2,265 | 0,187 | 2,087 | 0,069 | -7,9 | -0,95 | 3,048 | 0,340 | 2,827 | 0,168 | -7,2 | -0,65 |
| habenular commissure | 0,030 | 0,001 | 0,030 | 0,002 | -0,5 | -0,10 | 0,034 | 0,004 | 0,034 | 0,004 | 1,6 | 0,12 |
| hippocampus | 17,503 | 0,974 | 16,890 | 0,583 | -3,5 | -0,63 | 21,314 | 1,119 | 20,072 | 1,606 | -5,8 | -1,11 |
| hypothalamus | 8,958 | 0,615 | 8,588 | 0,753 | -4,1 | -0,60 | 10,245 | 0,884 | 9,783 | 0,515 | -4,5 | -0,52 |
| inferior olivary complex | 0,222 | 0,030 | 0,217 | 0,017 | -2,0 | -0,15 | 0,266 | 0,019 | 0,263 | 0,017 | -1,1 | -0,15 |
| internal capsule | 2,158 | 0,188 | 1,985 | 0,086 | -8,0 | -0,92 | 2,934 | 0,325 | 2,748 | 0,229 | -6,4 | -0,57 |
| interpeduncular nucleus | 0,215 | 0,025 | 0,202 | 0,024 | -6,0 | -0,52 | 0,258 | 0,022 | 0,244 | 0,030 | -5,2 | -0,61 |
| lateral olfactory tract | 0,775 | 0,034 | 0,754 | 0,043 | -2,7 | -0,62 | 0,852 | 0,043 | 0,797 | 0,061 | -6,5 | -1,30 |
| lateral septum | 2,509 | 0,226 | 2,419 | 0,180 | -3,6 | -0,40 | 2,920 | 0,246 | 2,770 | 0,184 | -5,2 | -0,61 |
| lateral ventricle | 2,537 | 0,202 | 2,479 | 0,219 | -2,3 | -0,29 | 2,939 | 0,253 | 2,760 | 0,231 | -6,1 | -0,71 |
| mammillary bodies | 0,390 | 0,029 | 0,377 | 0,036 | -3,5 | -0,47 | 0,473 | 0,043 | 0,477 | 0,036 | 1,0 | 0,11 |
| mammilothalamic tract | 0,240 | 0,013 | 0,231 | 0,011 | -3,5 | -0,63 | 0,268 | 0,022 | 0,251 | 0,014 | -6,2 | -0,77 |
| medial lemniscus/medial longitudinal fasciculus | 2,145 | 0,165 | 2,074 | 0,060 | -3,3 | -0,43 | 2,626 | 0,143 | 2,532 | 0,124 | -3,6 | -0,66 |
| medial septum | 1,155 | 0,090 | 1,094 | 0,111 | -5,3 | -0,68 | 1,335 | 0,106 | 1,276 | 0,065 | -4,4 | -0,55 |
| medulla | 22,692 | 1,765 | 22,037 | 0,973 | -2,9 | -0,37 | 25,365 | 1,352 | 24,728 | 1,671 | -2,5 | -0,47 |
| midbrain | 11,177 | 0,823 | 10,628 | 0,525 | -4,9 | -0,67 | 14,179 | 1,241 | 13,245 | 0,975 | -6,6 | -0,75 |
| nucleus accumbens | 3,652 | 0,274 | 3,537 | 0,280 | -3,2 | -0,42 | 4,274 | 0,314 | 4,120 | 0,262 | -3,6 | -0,49 |
| olfactory bulbs | 19,548 | 0,898 | 18,551 | 0,919 | -5,1 | -1,11 | 20,108 | 1,394 | 18,559 | 1,938 | -7,7 | -1,11 |
| olfactory tubercle | 2,739 | 0,153 | 2,678 | 0,186 | -2,2 | -0,40 | 3,340 | 0,259 | 3,233 | 0,246 | -3,2 | -0,41 |
| optic tract | 1,444 | 0,083 | 1,373 | 0,076 | -4,9 | -0,85 | 1,794 | 0,137 | 1,704 | 0,096 | -5,0 | -0,66 |
| periaqueductal grey | 3,751 | 0,249 | 3,578 | 0,201 | -4,6 | -0,69 | 4,311 | 0,301 | 4,006 | 0,271 | -7,1 | -1,01 |
| pons | 13,094 | 0,938 | 12,437 | 0,570 | -5,0 | -0,70 | 16,533 | 0,986 | 15,735 | 0,920 | -4,8 | -0,81 |
| pontine nucleus | 0,626 | 0,057 | 0,609 | 0,045 | -2,6 | -0,29 | 0,805 | 0,050 | 0,773 | 0,067 | -3,9 | -0,63 |
| posterior commissure | 0,138 | 0,009 | 0,136 | 0,009 | -1,3 | -0,19 | 0,156 | 0,011 | 0,145 | 0,011 | -7,4 | -1,05 |
| pre-para subiculum | 1,985 | 0,217 | 1,929 | 0,101 | -2,8 | -0,25 | 2,266 | 0,111 | 2,119 | 0,149 | -6,5 | -1,33 |
| stratum granulosum of hippocampus | 0,697 | 0,032 | 0,660 | 0,033 | -5,3 | -1,15 | 0,843 | 0,053 | 0,775 | 0,070 | -8,0 | -1,28 |
| stria medullaris | 0,653 | 0,035 | 0,628 | 0,028 | -3,8 | -0,72 | 0,734 | 0,046 | 0,696 | 0,047 | -5,2 | -0,83 |

| | | | | | | | | | | | | |
|---|---|---|---|---|---|---|---|---|---|---|---|---|
| stria terminalis | 0,705 | 0,072 | 0,657 | 0,046 | -6,9 | -0,68 | 0,915 | 0,094 | 0,860 | 0,081 | -6,0 | -0,59 |
| striatum | 16,749 | 1,580 | 16,285 | 1,023 | -2,8 | -0,29 | 21,450 | 1,796 | 20,677 | 1,019 | -3,6 | -0,43 |
| subependymale zone / rhinocele | 0,070 | 0,004 | 0,068 | 0,005 | -2,5 | -0,39 | 0,071 | 0,003 | 0,066 | 0,005 | -6,8 | -1,42 |
| superior olivary complex | 0,490 | 0,053 | 0,476 | 0,035 | -2,8 | -0,26 | 0,610 | 0,041 | 0,585 | 0,051 | -4,1 | -0,62 |
| thalamus | 14,682 | 1,052 | 14,059 | 0,642 | -4,2 | -0,59 | 17,157 | 1,255 | 16,157 | 0,953 | -5,8 | -0,80 |
| third ventricle | 0,910 | 0,054 | 0,878 | 0,055 | -3,4 | -0,58 | 1,000 | 0,073 | 0,952 | 0,061 | -4,9 | -0,67 |
| ventral tegmental decussation | 0,122 | 0,009 | 0,115 | 0,012 | -6,0 | -0,84 | 0,149 | 0,014 | 0,138 | 0,015 | -7,3 | -0,80 |
| lobules 1-2: lingula and central lobule (ventral) | 1,950 | 0,123 | 1,792 | 0,153 | -8,1 | -1,29 | 2,430 | 0,297 | 2,031 | 0,143 | -16,4 | -1,34 |
| lobule 3: central lobule (dorsal) | 1,979 | 0,230 | 1,801 | 0,161 | -9,0 | -0,77 | 2,584 | 0,273 | 2,324 | 0,165 | -10,1 | -0,95 |
| lobules 4-5: culmen (ventral and dorsal) | 3,631 | 0,441 | 3,570 | 0,286 | -1,7 | -0,14 | 4,552 | 0,370 | 4,322 | 0,286 | -5,1 | -0,62 |
| lobule 6: declive | 2,041 | 0,214 | 2,114 | 0,212 | 3,6 | 0,34 | 2,382 | 0,158 | 2,253 | 0,194 | -5,4 | -0,82 |
| lobule 7: tuber (or folium) | 0,825 | 0,097 | 0,858 | 0,095 | 3,9 | 0,33 | 0,880 | 0,108 | 0,814 | 0,094 | -7,5 | -0,61 |
| lobule 8: pyramis | 1,253 | 0,120 | 1,288 | 0,142 | 2,8 | 0,29 | 1,301 | 0,184 | 1,213 | 0,172 | -6,7 | -0,48 |
| lobule 9: uvula | 2,819 | 0,202 | 2,835 | 0,276 | 0,5 | 0,08 | 2,810 | 0,282 | 2,654 | 0,200 | -5,5 | -0,55 |
| lobule 10: nodulus | 1,651 | 0,199 | 1,651 | 0,281 | 0,0 | 0,00 | 1,711 | 0,083 | 1,637 | 0,093 | -4,3 | -0,88 |
| anterior lobule (lobules 4-5) | 1,675 | 0,148 | 1,563 | 0,125 | -6,7 | -0,76 | 2,137 | 0,205 | 1,904 | 0,200 | -10,9 | -1,14 |
| simple lobule (lobule 6) | 5,119 | 0,560 | 5,015 | 0,457 | -2,0 | -0,19 | 6,286 | 0,381 | 5,723 | 0,477 | -9,0 | -1,48 |
| crus 1: ansiform lobule (lobule 6) | 4,703 | 0,493 | 4,650 | 0,564 | -1,1 | -0,11 | 5,309 | 0,462 | 4,863 | 0,373 | -8,4 | -0,96 |
| crus 2: ansiform lobule (lobule 7) | 3,269 | 0,302 | 3,326 | 0,387 | 1,7 | 0,19 | 3,491 | 0,385 | 3,287 | 0,302 | -5,8 | -0,53 |
| paramedian lobule (lobule 7) | 3,368 | 0,281 | 3,489 | 0,363 | 3,6 | 0,43 | 3,461 | 0,312 | 3,479 | 0,364 | 0,5 | 0,06 |
| copula: pyramis (lobule 8) | 2,063 | 0,175 | 2,127 | 0,219 | 3,1 | 0,37 | 2,200 | 0,162 | 2,136 | 0,205 | -2,9 | -0,39 |
| flocculus (FL) | 0,994 | 0,053 | 0,941 | 0,043 | -5,4 | -1,01 | 1,098 | 0,106 | 1,036 | 0,083 | -5,6 | -0,58 |
| paraflocculus (PFL) | 4,192 | 0,212 | 4,093 | 0,104 | -2,4 | -0,47 | 4,253 | 0,272 | 4,113 | 0,254 | -3,3 | -0,51 |
| trunk of arbor vita | 3,202 | 0,330 | 3,072 | 0,332 | -4,1 | -0,39 | 4,413 | 0,477 | 4,111 | 0,279 | -6,9 | -0,63 |
| lobule 1-2 white matter | 0,106 | 0,008 | 0,095 | 0,010 | -10,4 | -1,41 | 0,141 | 0,019 | 0,115 | 0,010 | -18,4 | -1,38 |
| lobule 3 white matter | 0,265 | 0,029 | 0,243 | 0,023 | -8,2 | -0,74 | 0,348 | 0,044 | 0,314 | 0,027 | -9,9 | -0,79 |
| trunk of lobules 1-3 white matter | 0,138 | 0,014 | 0,125 | 0,011 | -9,1 | -0,89 | 0,205 | 0,031 | 0,186 | 0,016 | -9,1 | -0,59 |
| lobules 4-5 white matter | 0,701 | 0,080 | 0,708 | 0,061 | 0,9 | 0,08 | 0,880 | 0,090 | 0,841 | 0,063 | -4,5 | -0,44 |

| | | | | | | | | | | | | |
|---|---|---|---|---|---|---|---|---|---|---|---|---|
| lobules 6-7 white matter | 0,702 | 0,092 | 0,759 | 0,103 | 8,1 | 0,62 | 0,862 | 0,094 | 0,797 | 0,071 | -7,6 | -0,70 |
| lobule 8 white matter | 0,165 | 0,018 | 0,171 | 0,021 | 3,3 | 0,31 | 0,177 | 0,027 | 0,161 | 0,024 | -8,7 | -0,57 |
| trunk of lobules 6-8 white matter | 0,118 | 0,019 | 0,127 | 0,016 | 6,9 | 0,43 | 0,139 | 0,014 | 0,128 | 0,015 | -7,5 | -0,74 |
| lobule 9 white matter | 0,386 | 0,038 | 0,387 | 0,037 | 0,2 | 0,02 | 0,393 | 0,051 | 0,365 | 0,032 | -7,0 | -0,54 |
| lobule 10 white matter | 0,124 | 0,016 | 0,124 | 0,023 | -0,1 | -0,01 | 0,129 | 0,008 | 0,124 | 0,008 | -4,1 | -0,65 |
| anterior lobule white matter | 0,100 | 0,008 | 0,093 | 0,009 | -6,9 | -0,83 | 0,132 | 0,015 | 0,121 | 0,015 | -8,4 | -0,73 |
| simple lobule white matter | 0,571 | 0,064 | 0,555 | 0,053 | -2,7 | -0,24 | 0,724 | 0,052 | 0,650 | 0,066 | -10,3 | -1,43 |
| crus 1 white matter | 0,499 | 0,064 | 0,491 | 0,072 | -1,7 | -0,13 | 0,614 | 0,066 | 0,548 | 0,054 | -10,8 | -1,01 |
| trunk of simple and crus 1 white matter | 0,145 | 0,021 | 0,147 | 0,024 | 0,9 | 0,06 | 0,206 | 0,027 | 0,183 | 0,020 | -11,4 | -0,87 |
| crus 2 white matter | 0,316 | 0,036 | 0,328 | 0,048 | 3,9 | 0,34 | 0,347 | 0,047 | 0,322 | 0,033 | -7,3 | -0,54 |
| paramedian lobule | 0,231 | 0,027 | 0,246 | 0,031 | 6,1 | 0,53 | 0,241 | 0,027 | 0,242 | 0,027 | 0,3 | 0,03 |
| trunk of crus 2 and paramedian white matter | 0,338 | 0,041 | 0,359 | 0,053 | 6,3 | 0,51 | 0,381 | 0,042 | 0,370 | 0,028 | -3,0 | -0,27 |
| copula white matter | 0,111 | 0,010 | 0,113 | 0,013 | 2,1 | 0,24 | 0,118 | 0,011 | 0,109 | 0,010 | -8,3 | -0,88 |
| paraflocculus white matter | 0,360 | 0,021 | 0,353 | 0,010 | -2,0 | -0,34 | 0,369 | 0,032 | 0,357 | 0,025 | -3,4 | -0,39 |
| flocculus white matter | 0,065 | 0,006 | 0,060 | 0,004 | -7,0 | -0,81 | 0,074 | 0,008 | 0,070 | 0,006 | -5,7 | -0,52 |
| dentate nucleus | 0,246 | 0,032 | 0,233 | 0,024 | -5,1 | -0,39 | 0,347 | 0,033 | 0,328 | 0,026 | -5,4 | -0,58 |
| nucleus interpositus | 0,270 | 0,034 | 0,242 | 0,023 | -10,3 | -0,83 | 0,429 | 0,063 | 0,390 | 0,034 | -9,1 | -0,62 |
| fastigial nucleus | 0,246 | 0,032 | 0,229 | 0,029 | -6,7 | -0,51 | 0,369 | 0,054 | 0,338 | 0,027 | -8,3 | -0,57 |
| Cingulate cortex: area 24a | 1,472 | 0,070 | 1,451 | 0,117 | -1,4 | -0,30 | 1,538 | 0,199 | 1,491 | 0,141 | -3,1 | -0,24 |
| Cingulate cortex: area 24a' | 0,648 | 0,031 | 0,644 | 0,076 | -0,6 | -0,12 | 0,681 | 0,063 | 0,669 | 0,060 | -1,9 | -0,20 |
| Cingulate cortex: area 24b | 1,259 | 0,055 | 1,234 | 0,090 | -2,0 | -0,46 | 1,247 | 0,131 | 1,203 | 0,088 | -3,5 | -0,34 |
| Cingulate cortex: area 24b' | 0,456 | 0,021 | 0,453 | 0,054 | -0,7 | -0,15 | 0,475 | 0,040 | 0,465 | 0,035 | -2,0 | -0,23 |
| Cingulate cortex: area 25 | 0,489 | 0,037 | 0,497 | 0,039 | 1,6 | 0,20 | 0,514 | 0,032 | 0,488 | 0,029 | -5,1 | -0,82 |
| Cingulate cortex: area 29a | 0,696 | 0,052 | 0,690 | 0,058 | -0,8 | -0,10 | 0,730 | 0,050 | 0,724 | 0,060 | -0,8 | -0,12 |
| Cingulate cortex: area 29b | 0,393 | 0,031 | 0,383 | 0,032 | -2,5 | -0,32 | 0,414 | 0,030 | 0,402 | 0,034 | -3,1 | -0,43 |
| Cingulate cortex: area 29c | 1,503 | 0,080 | 1,498 | 0,161 | -0,4 | -0,07 | 1,549 | 0,100 | 1,581 | 0,116 | 2,1 | 0,32 |
| Cingulate cortex: area 30 | 2,615 | 0,163 | 2,640 | 0,245 | 1,0 | 0,15 | 2,822 | 0,211 | 2,709 | 0,254 | -4,0 | -0,53 |
| Cingulate cortex: area 32 | 2,000 | 0,064 | 1,994 | 0,173 | -0,3 | -0,09 | 1,969 | 0,168 | 1,909 | 0,130 | -3,0 | -0,35 |

| | | | | | | | | | | | | |
|---|---|---|---|---|---|---|---|---|---|---|---|---|
| Amygdalopiriform transition area | 0,698 | 0,054 | 0,670 | 0,042 | -4,1 | -0,53 | 0,830 | 0,057 | 0,795 | 0,049 | -4,2 | -0,62 |
| Primary auditory cortex | 1,366 | 0,105 | 1,367 | 0,082 | 0,1 | 0,02 | 1,471 | 0,145 | 1,387 | 0,143 | -5,7 | -0,58 |
| Secondary auditory cortex: dorsal area | 1,535 | 0,112 | 1,520 | 0,110 | -1,0 | -0,13 | 1,644 | 0,141 | 1,554 | 0,145 | -5,5 | -0,63 |
| Secondary auditory cortex: ventral area | 1,543 | 0,106 | 1,541 | 0,097 | -0,1 | -0,02 | 1,653 | 0,176 | 1,540 | 0,162 | -6,8 | -0,64 |
| Caudomedial entorhinal cortex | 3,888 | 0,362 | 3,856 | 0,159 | -0,8 | -0,09 | 4,832 | 0,291 | 4,438 | 0,337 | -8,2 | -1,35 |
| Cingulum | 0,624 | 0,023 | 0,615 | 0,045 | -1,5 | -0,40 | 0,714 | 0,060 | 0,682 | 0,057 | -4,5 | -0,53 |
| Claustrum | 0,232 | 0,021 | 0,232 | 0,017 | -0,1 | -0,01 | 0,268 | 0,021 | 0,261 | 0,021 | -2,8 | -0,36 |
| Cortex-amygdala transition zones | 0,893 | 0,049 | 0,876 | 0,099 | -2,0 | -0,36 | 1,053 | 0,096 | 1,017 | 0,106 | -3,4 | -0,37 |
| Claustrum: dorsal part | 0,309 | 0,028 | 0,302 | 0,024 | -2,3 | -0,26 | 0,328 | 0,028 | 0,317 | 0,021 | -3,2 | -0,38 |
| Dorsal nucleus of the endopiriform | 1,266 | 0,103 | 1,231 | 0,129 | -2,8 | -0,34 | 1,396 | 0,083 | 1,346 | 0,098 | -3,6 | -0,60 |
| Dorsal intermediate entorhinal cortex | 1,496 | 0,119 | 1,462 | 0,093 | -2,3 | -0,29 | 1,740 | 0,107 | 1,659 | 0,088 | -4,7 | -0,76 |
| Dorsolateral entorhinal cortex | 2,036 | 0,168 | 2,018 | 0,121 | -0,9 | -0,11 | 2,340 | 0,153 | 2,224 | 0,129 | -5,0 | -0,76 |
| Dorsolateral orbital cortex | 0,760 | 0,042 | 0,765 | 0,066 | 0,6 | 0,12 | 0,852 | 0,063 | 0,824 | 0,088 | -3,3 | -0,45 |
| Dorsal tenia tecta | 0,714 | 0,053 | 0,710 | 0,049 | -0,6 | -0,08 | 0,781 | 0,049 | 0,741 | 0,050 | -5,1 | -0,81 |
| Ectorhinal cortex | 2,590 | 0,195 | 2,575 | 0,134 | -0,6 | -0,08 | 2,818 | 0,186 | 2,617 | 0,210 | -7,1 | -1,08 |
| Frontal cortex: area 3 | 0,677 | 0,038 | 0,682 | 0,034 | 0,7 | 0,12 | 0,722 | 0,051 | 0,702 | 0,048 | -2,8 | -0,40 |
| Frontal association cortex | 6,432 | 0,259 | 6,439 | 0,374 | 0,1 | 0,03 | 6,630 | 0,370 | 6,383 | 0,523 | -3,7 | -0,67 |
| Intermediate nucleus of the endopiriform claustrum | 0,539 | 0,043 | 0,530 | 0,053 | -1,7 | -0,21 | 0,635 | 0,037 | 0,605 | 0,048 | -4,7 | -0,82 |
| Insular region: not subdivided | 7,212 | 0,586 | 7,160 | 0,639 | -0,7 | -0,09 | 7,625 | 0,459 | 7,323 | 0,501 | -4,0 | -0,66 |
| Lateral orbital cortex | 2,724 | 0,211 | 2,748 | 0,233 | 0,9 | 0,11 | 2,955 | 0,143 | 2,878 | 0,268 | -2,6 | -0,54 |
| Lateral parietal association cortex | 0,183 | 0,010 | 0,182 | 0,018 | -0,9 | -0,17 | 0,191 | 0,017 | 0,178 | 0,020 | -6,6 | -0,73 |
| Primary motor cortex | 6,111 | 0,222 | 6,100 | 0,353 | -0,2 | -0,05 | 6,495 | 0,524 | 6,251 | 0,491 | -3,8 | -0,47 |
| Secondary motor cortex | 6,056 | 0,200 | 6,085 | 0,418 | 0,5 | 0,14 | 6,226 | 0,511 | 5,937 | 0,455 | -4,6 | -0,57 |
| Medial entorhinal cortex | 0,528 | 0,049 | 0,518 | 0,018 | -2,0 | -0,21 | 0,667 | 0,050 | 0,627 | 0,045 | -6,1 | -0,82 |
| Medial orbital cortex | 1,411 | 0,083 | 1,419 | 0,120 | 0,6 | 0,10 | 1,394 | 0,090 | 1,346 | 0,106 | -3,4 | -0,53 |
| Medial parietal association cortex | 0,329 | 0,017 | 0,331 | 0,042 | 0,8 | 0,15 | 0,357 | 0,031 | 0,333 | 0,039 | -6,8 | -0,78 |
| Piriform cortex | 9,134 | 0,674 | 8,885 | 0,812 | -2,7 | -0,37 | 10,430 | 0,494 | 9,947 | 0,722 | -4,6 | -0,98 |
| Posterolateral cortical amygdaloid area | 1,002 | 0,074 | 0,940 | 0,080 | -6,2 | -0,83 | 1,184 | 0,059 | 1,130 | 0,050 | -4,6 | -0,91 |

| | | | | | | | | | | | | |
|---|---|---|---|---|---|---|---|---|---|---|---|---|
| Posteromedial cortical amygdaloid area | 0,967 | 0,075 | 0,929 | 0,074 | -4,0 | -0,51 | 1,147 | 0,060 | 1,120 | 0,053 | -2,3 | -0,44 |
| Perirhinal cortex | 2,380 | 0,185 | 2,352 | 0,128 | -1,2 | -0,15 | 2,647 | 0,162 | 2,487 | 0,183 | -6,0 | -0,98 |
| Parietal cortex: posterior area: rostral part | 0,116 | 0,007 | 0,113 | 0,006 | -3,0 | -0,48 | 0,122 | 0,010 | 0,113 | 0,012 | -7,5 | -0,89 |
| Rostral amygdalopiriform area | 0,526 | 0,042 | 0,494 | 0,032 | -6,0 | -0,75 | 0,618 | 0,032 | 0,588 | 0,031 | -5,0 | -0,95 |
| Primary somatosensory cortex | 4,109 | 0,253 | 4,099 | 0,203 | -0,2 | -0,04 | 4,405 | 0,317 | 4,229 | 0,315 | -4,0 | -0,56 |
| Primary somatosensory cortex: barrel field | 9,881 | 0,728 | 9,697 | 0,524 | -1,9 | -0,25 | 10,869 | 0,980 | 10,253 | 0,928 | -5,7 | -0,63 |
| Primary somatosensory cortex: dysgranular zone | 0,302 | 0,020 | 0,296 | 0,012 | -1,9 | -0,28 | 0,324 | 0,026 | 0,307 | 0,026 | -5,2 | -0,67 |
| Primary somatosensory cortex: forelimb region | 3,558 | 0,208 | 3,504 | 0,154 | -1,5 | -0,26 | 3,906 | 0,345 | 3,702 | 0,335 | -5,2 | -0,59 |
| Primary somatosensory cortex: hindlimb region | 2,418 | 0,113 | 2,388 | 0,208 | -1,3 | -0,27 | 2,615 | 0,234 | 2,466 | 0,224 | -5,7 | -0,63 |
| Primary somatosensory cortex: jaw region | 0,706 | 0,044 | 0,709 | 0,030 | 0,4 | 0,06 | 0,771 | 0,062 | 0,744 | 0,061 | -3,4 | -0,42 |
| Primary somatosensory cortex: shoulder region | 0,206 | 0,014 | 0,199 | 0,012 | -3,2 | -0,49 | 0,218 | 0,019 | 0,202 | 0,019 | -7,2 | -0,83 |
| Primary somatosensory cortex: trunk region | 0,461 | 0,025 | 0,454 | 0,040 | -1,6 | -0,29 | 0,483 | 0,042 | 0,451 | 0,049 | -6,7 | -0,78 |
| Primary somatosensory cortex: upper lip region | 5,418 | 0,343 | 5,385 | 0,218 | -0,6 | -0,10 | 5,824 | 0,459 | 5,567 | 0,422 | -4,4 | -0,56 |
| Secondary somatosensory cortex | 6,276 | 0,474 | 6,192 | 0,420 | -1,3 | -0,18 | 6,659 | 0,601 | 6,289 | 0,485 | -5,6 | -0,61 |
| Temporal association area | 2,499 | 0,183 | 2,499 | 0,118 | 0,0 | 0,00 | 2,692 | 0,198 | 2,498 | 0,209 | -7,2 | -0,98 |
| Primary visual cortex | 2,129 | 0,179 | 2,075 | 0,112 | -2,5 | -0,30 | 2,303 | 0,160 | 2,138 | 0,196 | -7,2 | -1,03 |
| Primary visual cortex: binocular area | 1,891 | 0,138 | 1,835 | 0,118 | -3,0 | -0,41 | 1,984 | 0,129 | 1,875 | 0,202 | -5,5 | -0,85 |
| Primary visual cortex: monocular area | 1,965 | 0,128 | 1,890 | 0,067 | -3,8 | -0,59 | 2,010 | 0,161 | 1,907 | 0,185 | -5,1 | -0,64 |
| Secondary visual cortex: lateral area | 2,430 | 0,195 | 2,393 | 0,142 | -1,5 | -0,19 | 2,617 | 0,170 | 2,461 | 0,249 | -6,0 | -0,92 |
| Secondary visual cortex: mediolateral area | 0,944 | 0,061 | 0,922 | 0,054 | -2,3 | -0,36 | 0,959 | 0,092 | 0,912 | 0,093 | -5,0 | -0,52 |
| Secondary visual cortex: mediomedial area | 1,335 | 0,081 | 1,317 | 0,086 | -1,3 | -0,22 | 1,392 | 0,121 | 1,332 | 0,131 | -4,3 | -0,50 |
| Claustrum: ventral part | 0,556 | 0,044 | 0,538 | 0,051 | -3,3 | -0,41 | 0,584 | 0,042 | 0,571 | 0,042 | -2,3 | -0,32 |
| Ventral nucleus of the endopiriform claustrum | 0,388 | 0,032 | 0,368 | 0,042 | -5,1 | -0,61 | 0,440 | 0,031 | 0,420 | 0,031 | -4,4 | -0,62 |
| Ventral intermediate entorhinal cortex | 0,787 | 0,063 | 0,767 | 0,040 | -2,5 | -0,32 | 0,951 | 0,067 | 0,905 | 0,058 | -4,9 | -0,70 |
| Ventral orbital cortex | 1,254 | 0,065 | 1,261 | 0,090 | 0,6 | 0,12 | 1,311 | 0,075 | 1,279 | 0,117 | -2,4 | -0,42 |
| Ventral tenia tecta | 0,120 | 0,010 | 0,121 | 0,009 | 0,4 | 0,06 | 0,140 | 0,011 | 0,132 | 0,012 | -5,7 | -0,73 |

| Atlas ROI | Absolute volumes (mm3) Mixed-model ANOVA F-statistic Age | Tx | Age:Tx | FDR-q value q-Age | q-Tx | q-Age:Tx | Relative volumes (% total brain volume) Mixed-model ANOVA F-statistic Age | Tx | Age:Tx | FDR-q value q-Age | q-Tx | q-Age:Tx |
|---|---|---|---|---|---|---|---|---|---|---|---|---|
| amygdala | 35,386 | 7,594 | 0,239 | 0,000 | 0,041 | 0,719 | 2,141 | 1,306 | 0,992 | 0,188 | 0,805 | 0,988 |
| anterior commissure: pars anterior | 57,793 | 5,809 | 3,175 | 0,000 | 0,064 | 0,523 | 8,814 | 0,001 | 0,609 | 0,009 | 0,992 | 0,988 |
| anterior commissure: pars posterior | 109,944 | 9,452 | 1,022 | 0,000 | 0,029 | 0,535 | 40,929 | 0,578 | 0,557 | 0,000 | 0,805 | 0,988 |
| basal forebrain | 26,268 | 6,407 | 0,292 | 0,000 | 0,058 | 0,714 | 0,002 | 0,388 | 0,478 | 0,979 | 0,820 | 0,988 |
| bed nucleus of stria terminalis | 30,635 | 4,472 | 0,840 | 0,000 | 0,094 | 0,576 | 1,089 | 0,067 | 0,016 | 0,356 | 0,949 | 0,988 |
| cerebellar peduncle: inferior | 111,017 | 4,710 | 0,010 | 0,000 | 0,090 | 0,939 | 10,533 | 0,423 | 1,792 | 0,005 | 0,805 | 0,988 |
| cerebellar peduncle: middle | 119,818 | 11,450 | 3,802 | 0,000 | 0,022 | 0,523 | 56,930 | 1,728 | 0,669 | 0,000 | 0,805 | 0,988 |
| .cerebellar peduncle: superior | 98,921 | 11,529 | 2,314 | 0,000 | 0,022 | 0,523 | 31,577 | 1,819 | 0,098 | 0,000 | 0,805 | 0,988 |
| cerebral aqueduct | 3,437 | 7,629 | 4,191 | 0,101 | 0,041 | 0,523 | 10,948 | 0,916 | 1,508 | 0,004 | 0,805 | 0,988 |
| cerebral peduncle | 177,074 | 9,342 | 0,434 | 0,000 | 0,029 | 0,646 | 114,035 | 0,622 | 1,176 | 0,000 | 0,805 | 0,988 |
| colliculus: inferior | 72,194 | 8,207 | 1,957 | 0,000 | 0,035 | 0,523 | 8,479 | 0,058 | 0,001 | 0,010 | 0,949 | 0,988 |
| colliculus: superior | 43,544 | 5,764 | 0,624 | 0,000 | 0,064 | 0,592 | 0,807 | 0,007 | 0,377 | 0,417 | 0,991 | 0,988 |
| corpus callosum | 41,058 | 6,272 | 2,303 | 0,000 | 0,060 | 0,523 | 7,150 | 0,599 | 0,655 | 0,016 | 0,805 | 0,988 |
| corticospinal tract/pyramids | 64,439 | 1,844 | 0,000 | 0,000 | 0,258 | 0,983 | 8,667 | 0,716 | 1,297 | 0,009 | 0,805 | 0,988 |
| cuneate nucleus | 0,022 | 0,063 | 0,148 | 0,899 | 0,814 | 0,782 | 14,301 | 2,962 | 0,963 | 0,001 | 0,793 | 0,988 |
| dentate gyrus of hippocampus | 76,885 | 10,771 | 1,753 | 0,000 | 0,022 | 0,523 | 17,298 | 1,695 | 0,003 | 0,001 | 0,805 | 0,988 |
| facial nerve (cranial nerve 7) | 97,738 | 4,956 | 0,628 | 0,000 | 0,086 | 0,592 | 23,375 | 0,001 | 0,240 | 0,000 | 0,992 | 0,988 |
| fasciculus retroflexus | 67,379 | 8,861 | 3,517 | 0,000 | 0,032 | 0,523 | 11,107 | 0,483 | 0,695 | 0,004 | 0,805 | 0,988 |
| fimbria | 56,652 | 7,924 | 1,567 | 0,000 | 0,037 | 0,523 | 12,786 | 1,084 | 0,015 | 0,002 | 0,805 | 0,988 |
| fornix | 52,937 | 8,752 | 0,676 | 0,000 | 0,032 | 0,586 | 5,903 | 0,961 | 0,315 | 0,029 | 0,805 | 0,988 |
| fourth ventricle | 6,909 | 9,567 | 2,094 | 0,021 | 0,029 | 0,523 | 2,427 | 1,801 | 0,177 | 0,163 | 0,805 | 0,988 |
| fundus of striatum | 30,168 | 8,489 | 0,072 | 0,000 | 0,033 | 0,824 | 0,899 | 2,259 | 1,733 | 0,398 | 0,805 | 0,988 |

| | | | | | | | | | | | | |
|---|---|---|---|---|---|---|---|---|---|---|---|---|
| globus pallidus | 177,803 | 18,208 | 1,560 | 0,000 | 0,008 | 0,523 | 212,493 | 12,178 | 0,160 | 0,000 | 0,074 | 0,988 |
| habenular commissure | 21,359 | 0,009 | 0,004 | 0,000 | 0,933 | 0,963 | 0,084 | 2,946 | 0,835 | 0,804 | 0,793 | 0,988 |
| hippocampus | 93,494 | 9,325 | 1,912 | 0,000 | 0,029 | 0,523 | 26,530 | 0,437 | 0,001 | 0,000 | 0,805 | 0,988 |
| hypothalamus | 34,219 | 4,909 | 0,243 | 0,000 | 0,086 | 0,719 | 0,168 | 0,000 | 0,524 | 0,721 | 0,995 | 0,988 |
| inferior olivary complex | 43,669 | 0,639 | 0,033 | 0,000 | 0,495 | 0,878 | 5,328 | 0,973 | 0,445 | 0,037 | 0,805 | 0,988 |
| internal capsule | 135,420 | 11,324 | 0,735 | 0,000 | 0,022 | 0,585 | 146,064 | 5,561 | 0,352 | 0,000 | 0,648 | 0,988 |
| interpedunclar nucleus | 26,136 | 3,314 | 0,083 | 0,000 | 0,147 | 0,823 | 3,613 | 0,212 | 0,272 | 0,086 | 0,896 | 0,988 |
| lateral olfactory tract | 13,859 | 7,973 | 2,466 | 0,001 | 0,037 | 0,523 | 20,307 | 0,741 | 0,466 | 0,000 | 0,805 | 0,988 |
| lateral septum | 39,107 | 5,968 | 1,048 | 0,000 | 0,064 | 0,530 | 1,718 | 0,141 | 0,018 | 0,240 | 0,918 | 0,988 |
| lateral ventricle | 24,221 | 4,768 | 1,899 | 0,000 | 0,090 | 0,523 | 0,125 | 0,142 | 0,330 | 0,759 | 0,918 | 0,988 |
| mammillary bodies | 69,565 | 0,492 | 0,227 | 0,000 | 0,535 | 0,724 | 13,037 | 1,459 | 2,123 | 0,002 | 0,805 | 0,988 |
| mammilothalamic tract | 24,866 | 8,802 | 1,473 | 0,000 | 0,032 | 0,523 | 4,571 | 0,176 | 0,039 | 0,053 | 0,908 | 0,988 |
| medial lemniscus/medial longitudinal fasciculus | 135,201 | 6,969 | 0,870 | 0,000 | 0,049 | 0,569 | 37,272 | 0,187 | 0,661 | 0,000 | 0,908 | 0,988 |
| medial septum | 42,156 | 5,896 | 0,086 | 0,000 | 0,064 | 0,823 | 1,459 | 0,094 | 0,816 | 0,280 | 0,938 | 0,988 |
| medulla | 29,122 | 2,307 | 0,055 | 0,000 | 0,209 | 0,843 | 0,251 | 0,782 | 0,948 | 0,665 | 0,805 | 0,988 |
| midbrain | 108,476 | 11,023 | 1,766 | 0,000 | 0,022 | 0,523 | 43,658 | 1,688 | 0,000 | 0,000 | 0,805 | 0,988 |
| nucleus accumbens | 53,218 | 4,819 | 0,760 | 0,000 | 0,089 | 0,585 | 5,491 | 0,017 | 0,231 | 0,035 | 0,981 | 0,988 |
| olfactory bulbs | 0,389 | 8,473 | 0,818 | 0,593 | 0,033 | 0,577 | 69,413 | 3,424 | 0,024 | 0,000 | 0,728 | 0,988 |
| olfactory tubercle | 79,156 | 3,175 | 0,739 | 0,000 | 0,147 | 0,585 | 14,420 | 0,339 | 0,154 | 0,001 | 0,836 | 0,988 |
| optic tract | 140,678 | 11,016 | 0,806 | 0,000 | 0,022 | 0,577 | 49,918 | 0,516 | 0,823 | 0,000 | 0,805 | 0,988 |
| periaqueductal grey | 37,033 | 11,059 | 1,481 | 0,000 | 0,022 | 0,523 | 0,000 | 1,124 | 0,011 | 0,993 | 0,805 | 0,988 |
| pons | 180,490 | 13,293 | 1,253 | 0,000 | 0,018 | 0,524 | 75,371 | 0,897 | 0,494 | 0,000 | 0,805 | 0,988 |
| pontine nucleus | 99,663 | 3,204 | 0,708 | 0,000 | 0,147 | 0,586 | 27,831 | 0,118 | 0,064 | 0,000 | 0,925 | 0,988 |
| posterior commissure | 16,618 | 5,335 | 3,054 | 0,001 | 0,075 | 0,523 | 3,175 | 0,001 | 0,640 | 0,108 | 0,992 | 0,988 |
| pre-para subiculum | 20,776 | 4,632 | 1,101 | 0,000 | 0,090 | 0,530 | 0,974 | 0,000 | 0,005 | 0,384 | 0,995 | 0,988 |
| stratum granulosum of hippocampus | 68,893 | 14,105 | 2,015 | 0,000 | 0,015 | 0,523 | 8,385 | 2,880 | 0,013 | 0,010 | 0,793 | 0,988 |
| stria medullaris | 35,737 | 8,648 | 0,968 | 0,000 | 0,033 | 0,541 | 1,579 | 0,118 | 0,350 | 0,260 | 0,925 | 0,988 |
| stria terminalis | 71,188 | 6,988 | 0,529 | 0,000 | 0,049 | 0,615 | 46,153 | 1,980 | 0,139 | 0,000 | 0,805 | 0,988 |

| | | | | | | | | | | | | |
|---|---|---|---|---|---|---|---|---|---|---|---|---|
| striatum | 133,162 | 5,145 | 1,276 | 0,000 | 0,079 | 0,524 | 85,640 | 0,104 | 0,107 | 0,000 | 0,931 | 0,988 |
| subependymale zone / rhinocele | 0,010 | 6,009 | 1,805 | 0,927 | 0,064 | 0,523 | 58,970 | 0,174 | 0,175 | 0,000 | 0,908 | 0,988 |
| superior olivary complex | 59,215 | 2,739 | 0,564 | 0,000 | 0,176 | 0,612 | 13,335 | 0,085 | 0,057 | 0,002 | 0,938 | 0,988 |
| thalamus | 51,586 | 8,992 | 1,186 | 0,000 | 0,032 | 0,527 | 4,101 | 0,734 | 0,156 | 0,067 | 0,805 | 0,988 |
| third ventricle | 18,623 | 5,609 | 0,534 | 0,000 | 0,068 | 0,615 | 4,476 | 0,002 | 0,295 | 0,056 | 0,992 | 0,988 |
| ventral tegmental decussation | 41,500 | 6,989 | 0,689 | 0,000 | 0,049 | 0,586 | 7,963 | 1,417 | 0,030 | 0,012 | 0,805 | 0,988 |
| lobules 1-2: lingula and central lobule (ventral) | 35,030 | 26,846 | 6,312 | 0,000 | 0,001 | 0,523 | 3,214 | 15,725 | 2,627 | 0,106 | 0,029 | 0,988 |
| lobule 3: central lobule (dorsal) | 73,122 | 14,055 | 1,160 | 0,000 | 0,015 | 0,529 | 27,219 | 5,242 | 0,002 | 0,000 | 0,649 | 0,988 |
| lobules 4-5: culmen (ventral and dorsal) | 54,517 | 3,017 | 1,442 | 0,000 | 0,159 | 0,523 | 12,015 | 0,031 | 0,056 | 0,003 | 0,972 | 0,988 |
| lobule 6: declive | 14,183 | 0,778 | 3,682 | 0,001 | 0,463 | 0,523 | 0,209 | 0,854 | 1,254 | 0,690 | 0,805 | 0,988 |
| lobule 7: tuber (or folium) | 0,039 | 0,566 | 2,807 | 0,866 | 0,516 | 0,523 | 10,488 | 0,420 | 1,269 | 0,005 | 0,805 | 0,988 |
| lobule 8: pyramis | 0,129 | 0,306 | 1,457 | 0,759 | 0,619 | 0,523 | 13,026 | 0,605 | 0,456 | 0,002 | 0,805 | 0,988 |
| lobule 9: uvula | 2,276 | 0,603 | 0,922 | 0,185 | 0,503 | 0,552 | 33,517 | 0,988 | 0,028 | 0,000 | 0,805 | 0,988 |
| lobule 10: nodulus | 0,087 | 0,404 | 0,395 | 0,800 | 0,573 | 0,663 | 8,222 | 0,429 | 0,001 | 0,011 | 0,805 | 0,988 |
| anterior lobule (lobules 4-5) | 49,888 | 11,140 | 1,742 | 0,000 | 0,022 | 0,523 | 12,158 | 3,539 | 0,143 | 0,003 | 0,728 | 0,988 |
| simple lobule (lobule 6) | 33,671 | 5,821 | 3,062 | 0,000 | 0,064 | 0,523 | 2,972 | 0,440 | 0,875 | 0,120 | 0,805 | 0,988 |
| crus 1: ansiform lobule (lobule 6) | 6,369 | 3,179 | 2,060 | 0,027 | 0,147 | 0,523 | 2,297 | 0,037 | 0,446 | 0,174 | 0,964 | 0,988 |
| crus 2: ansiform lobule (lobule 7) | 0,612 | 0,632 | 1,607 | 0,503 | 0,495 | 0,523 | 9,402 | 0,580 | 0,371 | 0,007 | 0,805 | 0,988 |
| paramedian lobule (lobule 7) | 0,251 | 0,208 | 0,365 | 0,679 | 0,682 | 0,667 | 10,850 | 3,997 | 0,003 | 0,004 | 0,714 | 0,988 |
| copula: pyramis (lobule 8) | 1,516 | 0,064 | 1,446 | 0,278 | 0,814 | 0,523 | 7,850 | 2,003 | 0,182 | 0,012 | 0,805 | 0,988 |
| flocculus (FL) | 15,441 | 6,870 | 0,431 | 0,001 | 0,049 | 0,646 | 0,856 | 0,479 | 0,226 | 0,408 | 0,805 | 0,988 |
| paraflocculus (PFL) | 0,168 | 2,692 | 0,078 | 0,725 | 0,177 | 0,823 | 39,957 | 1,264 | 0,884 | 0,000 | 0,805 | 0,988 |
| trunk of arbor vita | 129,942 | 8,424 | 2,615 | 0,000 | 0,033 | 0,523 | 49,581 | 0,926 | 0,164 | 0,000 | 0,805 | 0,988 |
| lobule 1-2 white matter | 48,075 | 28,398 | 6,052 | 0,000 | 0,001 | 0,523 | 13,287 | 17,488 | 2,417 | 0,002 | 0,029 | 0,988 |
| lobule 3 white matter | 67,199 | 12,269 | 1,558 | 0,000 | 0,022 | 0,523 | 23,538 | 4,164 | 0,050 | 0,000 | 0,714 | 0,988 |
| trunk of lobules 1-3 white matter | 178,403 | 15,607 | 2,044 | 0,000 | 0,013 | 0,523 | 91,584 | 4,849 | 0,032 | 0,000 | 0,667 | 0,988 |
| lobules 4-5 white matter | 45,391 | 1,414 | 2,072 | 0,000 | 0,310 | 0,523 | 8,583 | 0,495 | 0,369 | 0,010 | 0,805 | 0,988 |
| lobules 6-7 white matter | 12,022 | 0,328 | 5,874 | 0,003 | 0,609 | 0,523 | 0,004 | 0,869 | 3,143 | 0,968 | 0,805 | 0,988 |

| | | | | | | | | | | | | |
|---|---|---|---|---|---|---|---|---|---|---|---|---|
| lobule 8 white matter | 0,002 | 0,499 | 1,999 | 0,968 | 0,535 | 0,523 | 8,576 | 0,297 | 0,828 | 0,010 | 0,859 | 0,988 |
| trunk of lobules 6-8 white matter | 4,343 | 0,199 | 3,787 | 0,067 | 0,684 | 0,523 | 0,961 | 0,689 | 1,537 | 0,385 | 0,805 | 0,988 |
| lobule 9 white matter | 0,890 | 0,672 | 0,759 | 0,412 | 0,495 | 0,585 | 21,360 | 0,501 | 0,041 | 0,000 | 0,805 | 0,988 |
| lobule 10 white matter | 0,194 | 0,236 | 0,200 | 0,712 | 0,664 | 0,736 | 6,097 | 0,469 | 0,017 | 0,026 | 0,805 | 0,988 |
| anterior lobule white matter | 64,164 | 7,006 | 0,696 | 0,000 | 0,049 | 0,586 | 23,878 | 1,519 | 0,009 | 0,000 | 0,805 | 0,988 |
| simple lobule white matter | 38,779 | 6,914 | 3,356 | 0,000 | 0,049 | 0,523 | 6,910 | 1,153 | 1,161 | 0,018 | 0,805 | 0,988 |
| crus 1 white matter | 16,466 | 4,354 | 2,792 | 0,001 | 0,097 | 0,523 | 0,765 | 0,614 | 1,003 | 0,428 | 0,805 | 0,988 |
| trunk of simple and crus 1 white matter | 48,088 | 4,603 | 5,257 | 0,000 | 0,090 | 0,523 | 16,991 | 0,664 | 2,562 | 0,001 | 0,805 | 0,988 |
| crus 2 white matter | 0,733 | 0,382 | 2,212 | 0,459 | 0,581 | 0,523 | 5,193 | 0,496 | 0,899 | 0,039 | 0,805 | 0,988 |
| paramedian lobule | 0,172 | 0,423 | 0,683 | 0,725 | 0,566 | 0,586 | 7,428 | 3,845 | 0,078 | 0,015 | 0,714 | 0,988 |
| trunk of crus 2 and paramedian white matter | 4,957 | 0,007 | 2,411 | 0,050 | 0,936 | 0,523 | 1,148 | 1,648 | 0,780 | 0,344 | 0,805 | 0,988 |
| copula white matter | 0,123 | 1,325 | 3,077 | 0,762 | 0,325 | 0,523 | 12,354 | 0,239 | 1,021 | 0,003 | 0,884 | 0,988 |
| paraflocculus white matter | 0,583 | 1,749 | 0,137 | 0,508 | 0,263 | 0,788 | 26,322 | 1,166 | 0,512 | 0,000 | 0,805 | 0,988 |
| flocculus white matter | 19,886 | 5,977 | 0,250 | 0,000 | 0,064 | 0,719 | 0,731 | 0,990 | 0,202 | 0,437 | 0,805 | 0,988 |
| dentate nucleus | 135,214 | 6,440 | 1,240 | 0,000 | 0,058 | 0,524 | 67,314 | 0,672 | 0,000 | 0,000 | 0,805 | 0,988 |
| nucleus interpositus | 214,477 | 15,550 | 1,884 | 0,000 | 0,013 | 0,523 | 151,789 | 6,439 | 0,022 | 0,000 | 0,511 | 0,988 |
| fastigial nucleus | 145,929 | 9,938 | 2,224 | 0,000 | 0,028 | 0,523 | 71,392 | 2,372 | 0,135 | 0,000 | 0,805 | 0,988 |
| Cingulate cortex: area 24a | 1,731 | 1,183 | 0,439 | 0,243 | 0,354 | 0,646 | 11,056 | 0,372 | 0,017 | 0,004 | 0,820 | 0,988 |
| Cingulate cortex: area 24a' | 2,766 | 0,657 | 0,387 | 0,141 | 0,495 | 0,663 | 8,464 | 0,962 | 0,030 | 0,010 | 0,805 | 0,988 |
| Cingulate cortex: area 24b | 0,214 | 1,865 | 0,365 | 0,703 | 0,257 | 0,667 | 47,946 | 0,446 | 0,141 | 0,000 | 0,805 | 0,988 |
| Cingulate cortex: area 24b' | 1,746 | 0,551 | 0,251 | 0,243 | 0,519 | 0,719 | 12,996 | 1,314 | 0,117 | 0,002 | 0,805 | 0,988 |
| Cingulate cortex: area 25 | 0,869 | 1,606 | 3,479 | 0,415 | 0,281 | 0,523 | 30,473 | 1,239 | 1,284 | 0,000 | 0,805 | 0,988 |
| Cingulate cortex: area 29a | 3,519 | 0,114 | 0,002 | 0,098 | 0,757 | 0,972 | 14,215 | 4,020 | 0,919 | 0,001 | 0,714 | 0,988 |
| Cingulate cortex: area 29b | 3,549 | 1,458 | 0,082 | 0,097 | 0,307 | 0,823 | 18,775 | 1,037 | 0,700 | 0,000 | 0,805 | 0,988 |
| Cingulate cortex: area 29c | 2,957 | 0,125 | 0,238 | 0,129 | 0,750 | 0,719 | 13,753 | 6,528 | 2,081 | 0,002 | 0,511 | 0,988 |
| Cingulate cortex: area 30 | 3,405 | 0,540 | 1,115 | 0,102 | 0,520 | 0,530 | 11,084 | 1,796 | 0,125 | 0,004 | 0,805 | 0,988 |
| Cingulate cortex: area 32 | 0,902 | 1,053 | 0,799 | 0,411 | 0,379 | 0,577 | 58,449 | 1,471 | 0,000 | 0,000 | 0,805 | 0,988 |
| Amygdalopiriform transition area | 62,197 | 5,781 | 0,519 | 0,000 | 0,064 | 0,615 | 11,873 | 0,009 | 0,575 | 0,003 | 0,987 | 0,988 |

| | | | | | | | | | | | | |
|---|---|---|---|---|---|---|---|---|---|---|---|---|
| Primary auditory cortex | 2,537 | 1,725 | 1,786 | 0,160 | 0,265 | 0,523 | 21,793 | 0,547 | 0,914 | 0,000 | 0,805 | 0,988 |
| Secondary auditory cortex: dorsal area | 3,028 | 2,349 | 1,428 | 0,125 | 0,208 | 0,523 | 22,915 | 0,218 | 0,413 | 0,000 | 0,896 | 0,988 |
| Secondary auditory cortex: ventral area | 1,354 | 2,188 | 1,997 | 0,305 | 0,223 | 0,523 | 25,745 | 0,174 | 1,019 | 0,000 | 0,908 | 0,988 |
| Caudomedial entorhinal cortex | 55,742 | 6,040 | 4,472 | 0,000 | 0,064 | 0,523 | 12,943 | 0,090 | 2,607 | 0,002 | 0,938 | 0,988 |
| Cingulum | 24,736 | 3,267 | 1,539 | 0,000 | 0,147 | 0,523 | 0,020 | 0,150 | 0,095 | 0,911 | 0,918 | 0,988 |
| Claustrum | 25,661 | 1,075 | 0,971 | 0,000 | 0,377 | 0,541 | 0,383 | 1,216 | 0,010 | 0,584 | 0,805 | 0,988 |
| Cortex-amygdala transition zones | 29,751 | 1,959 | 0,636 | 0,000 | 0,247 | 0,592 | 1,852 | 0,062 | 0,010 | 0,224 | 0,949 | 0,988 |
| Claustrum: dorsal part | 4,818 | 1,822 | 0,271 | 0,053 | 0,260 | 0,719 | 13,289 | 0,629 | 0,314 | 0,002 | 0,805 | 0,988 |
| Dorsal nucleus of the endopiriform | 13,760 | 2,906 | 0,544 | 0,001 | 0,163 | 0,615 | 1,833 | 0,041 | 0,105 | 0,224 | 0,962 | 0,988 |
| Dorsal intermediate entorhinal cortex | 42,869 | 4,017 | 0,963 | 0,000 | 0,110 | 0,541 | 0,830 | 0,371 | 0,100 | 0,413 | 0,820 | 0,988 |
| Dorsolateral entorhinal cortex | 28,085 | 2,388 | 1,304 | 0,000 | 0,207 | 0,524 | 0,383 | 1,475 | 0,001 | 0,584 | 0,805 | 0,988 |
| Dorsolateral orbital cortex | 12,938 | 1,071 | 1,505 | 0,002 | 0,377 | 0,523 | 0,910 | 0,786 | 0,211 | 0,397 | 0,805 | 0,988 |
| Dorsal tenia tecta | 10,380 | 3,546 | 2,594 | 0,005 | 0,138 | 0,523 | 11,167 | 0,275 | 0,502 | 0,004 | 0,863 | 0,988 |
| Ectorhinal cortex | 4,043 | 3,473 | 2,556 | 0,077 | 0,140 | 0,523 | 46,912 | 0,509 | 1,470 | 0,000 | 0,805 | 0,988 |
| Frontal cortex: area 3 | 5,747 | 1,190 | 1,817 | 0,035 | 0,354 | 0,523 | 34,806 | 4,684 | 0,347 | 0,000 | 0,667 | 0,988 |
| Frontal association cortex | 0,588 | 1,758 | 1,831 | 0,508 | 0,263 | 0,523 | 46,920 | 1,573 | 0,249 | 0,000 | 0,805 | 0,988 |
| Intermediate nucleus of the endopiriform claustrum | 35,029 | 3,628 | 1,650 | 0,000 | 0,134 | 0,523 | 2,142 | 0,005 | 0,115 | 0,188 | 0,992 | 0,988 |
| Insular region: not subdivided | 2,925 | 1,796 | 1,125 | 0,130 | 0,262 | 0,530 | 24,807 | 0,924 | 0,078 | 0,000 | 0,805 | 0,988 |
| Lateral orbital cortex | 6,868 | 0,632 | 1,182 | 0,022 | 0,495 | 0,527 | 8,385 | 2,328 | 0,142 | 0,010 | 0,805 | 0,988 |
| Lateral parietal association cortex | 0,210 | 2,333 | 1,487 | 0,703 | 0,208 | 0,523 | 19,304 | 0,015 | 0,323 | 0,000 | 0,981 | 0,988 |
| Primary motor cortex | 4,267 | 1,868 | 1,626 | 0,069 | 0,257 | 0,523 | 24,624 | 1,275 | 0,181 | 0,000 | 0,805 | 0,988 |
| Secondary motor cortex | 0,076 | 1,765 | 2,260 | 0,810 | 0,263 | 0,523 | 43,066 | 1,028 | 0,481 | 0,000 | 0,805 | 0,988 |
| Medial entorhinal cortex | 78,834 | 4,701 | 2,019 | 0,000 | 0,090 | 0,523 | 26,923 | 0,018 | 0,134 | 0,000 | 0,981 | 0,988 |
| Medial orbital cortex | 1,441 | 0,681 | 1,085 | 0,290 | 0,495 | 0,530 | 58,176 | 1,939 | 0,042 | 0,000 | 0,805 | 0,988 |
| Medial parietal association cortex | 1,838 | 1,415 | 1,959 | 0,233 | 0,310 | 0,523 | 8,268 | 0,165 | 0,723 | 0,010 | 0,911 | 0,988 |
| Piriform cortex | 27,896 | 4,362 | 1,048 | 0,000 | 0,097 | 0,530 | 0,035 | 0,000 | 0,007 | 0,880 | 0,995 | 0,988 |
| Posterolateral cortical amygdaloid area | 82,658 | 10,832 | 0,131 | 0,000 | 0,022 | 0,790 | 12,501 | 0,833 | 1,892 | 0,002 | 0,805 | 0,988 |
| Posteromedial cortical amygdaloid area | 78,750 | 4,077 | 0,070 | 0,000 | 0,108 | 0,824 | 16,244 | 0,427 | 2,128 | 0,001 | 0,805 | 0,988 |

| | | | | | | | | | | | | |
|---|---|---|---|---|---|---|---|---|---|---|---|---|
| Perirhinal cortex | 11,946 | 3,204 | 1,627 | 0,003 | 0,147 | 0,523 | 17,812 | 0,928 | 0,200 | 0,000 | 0,805 | 0,988 |
| Parietal cortex: posterior area: rostral part | 1,006 | 5,263 | 1,475 | 0,384 | 0,076 | 0,523 | 25,269 | 0,441 | 0,191 | 0,000 | 0,805 | 0,988 |
| Rostral amygdalopiriform area | 69,771 | 9,885 | 0,123 | 0,000 | 0,028 | 0,793 | 9,032 | 0,707 | 1,979 | 0,008 | 0,805 | 0,988 |
| Primary somatosensory cortex | 5,770 | 2,054 | 1,732 | 0,035 | 0,238 | 0,523 | 37,759 | 2,371 | 0,404 | 0,000 | 0,805 | 0,988 |
| Primary somatosensory cortex: barrel field | 8,110 | 3,235 | 1,325 | 0,013 | 0,147 | 0,524 | 9,473 | 0,012 | 0,203 | 0,007 | 0,981 | 0,988 |
| Primary somatosensory cortex: dysgranular zone | 5,629 | 3,524 | 1,311 | 0,037 | 0,138 | 0,524 | 25,059 | 0,080 | 0,106 | 0,000 | 0,939 | 0,988 |
| Primary somatosensory cortex: forelimb region | 9,008 | 3,219 | 1,540 | 0,009 | 0,147 | 0,523 | 9,826 | 0,061 | 0,261 | 0,006 | 0,949 | 0,988 |
| Primary somatosensory cortex: hindlimb region | 4,496 | 2,822 | 1,534 | 0,062 | 0,170 | 0,523 | 10,705 | 0,027 | 0,254 | 0,004 | 0,974 | 0,988 |
| Primary somatosensory cortex: jaw region | 9,093 | 1,411 | 1,736 | 0,008 | 0,310 | 0,523 | 15,740 | 2,869 | 0,519 | 0,001 | 0,793 | 0,988 |
| Primary somatosensory cortex: shoulder region | 2,174 | 5,442 | 1,399 | 0,192 | 0,072 | 0,523 | 19,542 | 0,517 | 0,136 | 0,000 | 0,805 | 0,988 |
| Primary somatosensory cortex: trunk region | 0,528 | 2,989 | 1,395 | 0,529 | 0,159 | 0,523 | 20,004 | 0,002 | 0,236 | 0,000 | 0,992 | 0,988 |
| Primary somatosensory cortex: upper lip region | 5,878 | 2,405 | 1,669 | 0,034 | 0,207 | 0,523 | 34,897 | 1,561 | 0,346 | 0,000 | 0,805 | 0,988 |
| Secondary somatosensory cortex | 2,225 | 2,699 | 1,297 | 0,188 | 0,177 | 0,524 | 32,679 | 0,268 | 0,185 | 0,000 | 0,863 | 0,988 |
| Temporal association area | 2,233 | 3,273 | 3,175 | 0,188 | 0,147 | 0,523 | 57,279 | 0,444 | 2,855 | 0,000 | 0,805 | 0,988 |
| Primary visual cortex | 3,932 | 4,325 | 1,283 | 0,081 | 0,098 | 0,524 | 23,260 | 0,046 | 0,155 | 0,000 | 0,962 | 0,988 |
| Primary visual cortex: binocular area | 1,796 | 3,402 | 0,653 | 0,237 | 0,144 | 0,592 | 25,128 | 0,012 | 0,001 | 0,000 | 0,981 | 0,988 |
| Primary visual cortex: monocular area | 0,441 | 4,083 | 0,261 | 0,568 | 0,108 | 0,719 | 44,209 | 0,013 | 0,348 | 0,000 | 0,981 | 0,988 |
| Secondary visual cortex: lateral area | 3,731 | 2,971 | 1,391 | 0,089 | 0,159 | 0,523 | 20,702 | 0,042 | 0,344 | 0,000 | 0,962 | 0,988 |
| Secondary visual cortex: mediolateral area | 0,015 | 2,372 | 0,497 | 0,914 | 0,207 | 0,623 | 37,121 | 0,116 | 0,014 | 0,000 | 0,925 | 0,988 |
| Secondary visual cortex: mediomedial area | 1,093 | 1,627 | 0,613 | 0,363 | 0,280 | 0,592 | 23,912 | 0,597 | 0,001 | 0,000 | 0,805 | 0,988 |
| Claustrum: ventral part | 4,984 | 2,048 | 0,091 | 0,050 | 0,238 | 0,823 | 12,289 | 0,283 | 0,811 | 0,003 | 0,863 | 0,988 |
| Ventral nucleus of the endopiriform claustrum | 22,152 | 4,592 | 0,218 | 0,000 | 0,090 | 0,726 | 0,218 | 0,333 | 0,553 | 0,687 | 0,836 | 0,988 |
| Ventral intermediate entorhinal cortex | 64,518 | 4,582 | 1,198 | 0,000 | 0,090 | 0,527 | 13,484 | 0,113 | 0,023 | 0,002 | 0,925 | 0,988 |
| Ventral orbital cortex | 2,107 | 0,634 | 1,063 | 0,198 | 0,495 | 0,530 | 33,950 | 3,600 | 0,038 | 0,000 | 0,728 | 0,988 |
| Ventral tenia tecta | 23,189 | 1,984 | 2,387 | 0,000 | 0,246 | 0,523 | 0,001 | 0,502 | 0,605 | 0,984 | 0,805 | 0,988 |

**Supplementary table 3:**
**Differentially expressed gene (DEG) lists from RNAseq analysis of CD11B+ microglia from the cerebrum and cerebellum.**

| *CEREBRUM IL1 vs PBS* | *log2FoldC* | *padj* | *Gene name* | *CEREBELLUM IL1 vs PBS* | *log2FoldCh* | *padj* | *Gene name* |
|---|---|---|---|---|---|---|---|
| ENSMUSG00000067219 | 5,362 | 0,011 | Nipal1 | ENSMUSG00000067219 | 6,013 | 0,005 | Nipal1 |
| ENSMUSG00000041481 | 4,892 | 0,000 | Serpina3g | ENSMUSG00000041481 | 5,087 | 0,000 | Serpina3g |
| ENSMUSG00000048534 | 4,745 | 0,000 | Jaml | ENSMUSG00000032726 | 4,850 | 0,016 | Bmp8a |
| ENSMUSG00000032068 | 4,679 | 0,000 | Plet1 | ENSMUSG00000054072 | 4,840 | 0,000 | Iigp1 |
| ENSMUSG00000006179 | 4,622 | 0,000 | Prss16 | ENSMUSG00000078921 | 4,732 | 0,000 | Tgtp2 |
| ENSMUSG00000041449 | 4,498 | 0,000 | Serpina3h | ENSMUSG00000006179 | 4,704 | 0,000 | Prss16 |
| ENSMUSG00000031780 | 4,469 | 0,000 | Ccl17 | ENSMUSG00000031780 | 4,537 | 0,000 | Ccl17 |
| ENSMUSG00000079014 | 4,340 | 0,000 | Serpina3i | ENSMUSG00000067261 | 4,397 | 0,040 | Foxd3 |
| ENSMUSG00000066363 | 4,275 | 0,000 | Serpina3f | ENSMUSG00000032068 | 4,364 | 0,000 | Plet1 |
| ENSMUSG00000085913 | 4,234 | 0,026 | Gm15601 | ENSMUSG00000068303 | 4,180 | 0,017 | Spr-ps1 |
| ENSMUSG00000091553 | 4,202 | 0,000 | Serpina3e-ps | ENSMUSG00000078922 | 4,180 | 0,000 | Tgtp1 |
| ENSMUSG00000078922 | 4,110 | 0,000 | Tgtp1 | ENSMUSG00000092920 | 3,831 | 0,001 | Mirt2 |
| ENSMUSG00000021553 | 4,088 | 0,000 | Slc28a3 | ENSMUSG00000056569 | 3,728 | 0,000 | Mpz |
| ENSMUSG00000054072 | 3,905 | 0,000 | Iigp1 | ENSMUSG00000021553 | 3,695 | 0,001 | Slc28a3 |
| ENSMUSG00000032726 | 3,862 | 0,038 | Bmp8a | ENSMUSG00000041449 | 3,652 | 0,000 | Serpina3h |
| ENSMUSG00000012428 | 3,677 | 0,000 | Steap4 | ENSMUSG00000048534 | 3,632 | 0,000 | Jaml |
| ENSMUSG00000101304 | 3,568 | 0,000 | Plet1os | ENSMUSG00000023132 | 3,599 | 0,000 | Gzma |
| ENSMUSG00000026073 | 3,433 | 0,000 | Il1r2 | ENSMUSG00000091553 | 3,551 | 0,000 | Serpina3e-ps |
| ENSMUSG00000097848 | 3,357 | 0,000 | Gm807 | ENSMUSG00000012428 | 3,445 | 0,000 | Steap4 |
| ENSMUSG00000039196 | 3,205 | 0,000 | Orm1 | ENSMUSG00000056290 | 3,425 | 0,000 | Ms4a4b |
| ENSMUSG00000085810 | 3,189 | 0,024 | Gm16325 | ENSMUSG00000094733 | 3,387 | 0,000 | Gm5416 |
| ENSMUSG00000048752 | 3,132 | 0,023 | Prss50 | ENSMUSG00000079014 | 3,350 | 0,000 | Serpina3i |
| ENSMUSG00000045534 | 3,055 | 0,000 | Kcna5 | ENSMUSG00000050014 | 3,319 | 0,000 | Apol10b |
| ENSMUSG00000078921 | 3,025 | 0,000 | Tgtp2 | ENSMUSG00000066363 | 3,293 | 0,000 | Serpina3f |
| ENSMUSG00000035896 | 2,987 | 0,003 | Rnase1 | ENSMUSG00000079363 | 3,291 | 0,000 | Gbp4 |
| ENSMUSG00000020407 | 2,895 | 0,000 | Upp1 | ENSMUSG00000060183 | 3,139 | 0,000 | Cxcl11 |
| ENSMUSG00000029379 | 2,893 | 0,002 | Cxcl3 | ENSMUSG00000105096 | 3,114 | 0,000 | Gbp10 |
| ENSMUSG00000060183 | 2,849 | 0,000 | Cxcl11 | ENSMUSG00000047222 | 3,114 | 0,020 | Rnase2a |
| ENSMUSG00000094733 | 2,821 | 0,000 | Gm5416 | ENSMUSG00000026073 | 3,075 | 0,000 | Il1r2 |
| ENSMUSG00000066687 | 2,814 | 0,000 | Zbtb16 | ENSMUSG00000066687 | 3,069 | 0,000 | Zbtb16 |
| ENSMUSG00000020183 | 2,795 | 0,000 | Cpm | ENSMUSG00000068227 | 3,060 | 0,000 | Il2rb |
| ENSMUSG00000032311 | 2,729 | 0,000 | Nrg4 | ENSMUSG00000029417 | 3,023 | 0,000 | Cxcl9 |
| ENSMUSG00000070407 | 2,713 | 0,000 | Hs3st3b1 | ENSMUSG00000101304 | 2,965 | 0,000 | Plet1os |
| ENSMUSG00000066170 | 2,660 | 0,000 | E230001N04Rik | ENSMUSG00000032311 | 2,957 | 0,000 | Nrg4 |
| ENSMUSG00000047798 | 2,651 | 0,000 | Cd300lf | ENSMUSG00000097848 | 2,929 | 0,000 | Gm807 |
| ENSMUSG00000033863 | 2,629 | 0,000 | Klf9 | ENSMUSG00000095620 | 2,912 | 0,002 | 2010005H15Rik |
| ENSMUSG00000027962 | 2,579 | 0,000 | Vcam1 | ENSMUSG00000064147 | 2,893 | 0,000 | Rab44 |
| ENSMUSG00000020027 | 2,538 | 0,000 | Socs2 | ENSMUSG00000020183 | 2,787 | 0,000 | Cpm |
| ENSMUSG00000027670 | 2,520 | 0,000 | Ocstamp | ENSMUSG00000078783 | 2,750 | 0,000 | Gm9733 |
| ENSMUSG00000024222 | 2,500 | 0,000 | Fkbp5 | ENSMUSG00000066170 | 2,729 | 0,000 | E230001N04Rik |

| | | | | | | | |
|---|---|---|---|---|---|---|---|
| ENSMUSG00000084956 | 2,478 | 0,000 | Gm16194 | ENSMUSG00000020407 | 2,726 | 0,000 | Upp1 |
| ENSMUSG00000022479 | 2,466 | 0,000 | Vdr | ENSMUSG00000033863 | 2,707 | 0,000 | Klf9 |
| ENSMUSG00000031779 | 2,462 | 0,001 | Ccl22 | ENSMUSG00000025739 | 2,622 | 0,027 | Gng13 |
| ENSMUSG00000022651 | 2,451 | 0,000 | Retnlg | ENSMUSG00000027962 | 2,621 | 0,000 | Vcam1 |
| ENSMUSG00000064147 | 2,441 | 0,000 | Rab44 | ENSMUSG00000039196 | 2,613 | 0,001 | Orm1 |
| ENSMUSG00000002847 | 2,439 | 0,000 | Pla1a | ENSMUSG00000062488 | 2,598 | 0,000 | Ifit3b |
| ENSMUSG00000027360 | 2,424 | 0,000 | Hdc | ENSMUSG00000035896 | 2,597 | 0,017 | Rnase1 |
| ENSMUSG00000031762 | 2,410 | 0,000 | Mt2 | ENSMUSG00000029378 | 2,564 | 0,001 | Areg |
| ENSMUSG00000026770 | 2,389 | 0,003 | Il2ra | ENSMUSG00000064032 | 2,532 | 0,015 | Gm10143 |
| ENSMUSG00000030144 | 2,365 | 0,000 | Clec4d | ENSMUSG00000026581 | 2,527 | 0,000 | Sell |
| ENSMUSG00000078763 | 2,363 | 0,000 | Slfn1 | ENSMUSG00000045534 | 2,513 | 0,000 | Kcna5 |
| ENSMUSG00000102189 | 2,362 | 0,006 | Gm37194 | ENSMUSG00000034271 | 2,476 | 0,000 | Jdp2 |
| ENSMUSG00000053007 | 2,360 | 0,000 | Creb5 | ENSMUSG00000078817 | 2,471 | 0,000 | Nlrp12 |
| ENSMUSG00000034271 | 2,359 | 0,000 | Jdp2 | ENSMUSG00000047976 | 2,429 | 0,048 | Kcna1 |
| ENSMUSG00000031778 | 2,331 | 0,000 | Cx3cl1 | ENSMUSG00000099137 | 2,415 | 0,000 | Gm10603 |
| ENSMUSG00000009248 | 2,302 | 0,000 | Ascl2 | ENSMUSG00000038067 | 2,409 | 0,041 | Csf3 |
| ENSMUSG00000105096 | 2,295 | 0,000 | Gbp10 | ENSMUSG00000002847 | 2,409 | 0,000 | Pla1a |
| ENSMUSG00000095620 | 2,261 | 0,017 | 2010005H15Rik | ENSMUSG00000026770 | 2,406 | 0,016 | Il2ra |
| ENSMUSG00000096278 | 2,234 | 0,010 | Dcpp2 | ENSMUSG00000022651 | 2,385 | 0,000 | Retnlg |
| ENSMUSG00000039956 | 2,221 | 0,000 | Mrap | ENSMUSG00000007480 | 2,384 | 0,018 | Mc5r |
| ENSMUSG00000005824 | 2,209 | 0,000 | Tnfsf14 | ENSMUSG00000022602 | 2,367 | 0,000 | Arc |
| ENSMUSG00000050578 | 2,208 | 0,002 | Mmp13 | ENSMUSG00000052403 | 2,353 | 0,016 | Fcnaos |
| ENSMUSG00000035352 | 2,208 | 0,000 | Ccl12 | ENSMUSG00000034459 | 2,352 | 0,000 | Ifit1 |
| ENSMUSG00000097472 | 2,197 | 0,010 | Gm26586 | ENSMUSG00000078630 | 2,338 | 0,016 | Tomt |
| ENSMUSG00000020940 | 2,188 | 0,013 | 1700023F06Rik | ENSMUSG00000032327 | 2,310 | 0,000 | Stra6 |
| ENSMUSG00000058626 | 2,180 | 0,021 | Capn11 | ENSMUSG00000033368 | 2,304 | 0,000 | Trim69 |
| ENSMUSG00000029417 | 2,180 | 0,001 | Cxcl9 | ENSMUSG00000097141 | 2,302 | 0,036 | Gm10524 |
| ENSMUSG00000029378 | 2,172 | 0,001 | Areg | ENSMUSG00000082292 | 2,299 | 0,000 | Gm12250 |
| ENSMUSG00000079363 | 2,165 | 0,000 | Gbp4 | ENSMUSG00000004328 | 2,294 | 0,000 | Hif3a |
| ENSMUSG00000032094 | 2,157 | 0,000 | Cd3d | ENSMUSG00000096719 | 2,264 | 0,031 | Mrgpra2b |
| ENSMUSG00000004328 | 2,156 | 0,000 | Hif3a | ENSMUSG00000052270 | 2,247 | 0,000 | Fpr2 |
| ENSMUSG00000028270 | 2,124 | 0,000 | Gbp2 | ENSMUSG00000084956 | 2,234 | 0,002 | Gm16194 |
| ENSMUSG00000022584 | 2,122 | 0,000 | Ly6c2 | ENSMUSG00000032094 | 2,224 | 0,000 | Cd3d |
| ENSMUSG00000086742 | 2,119 | 0,000 | Gm16201 | ENSMUSG00000024222 | 2,203 | 0,000 | Fkbp5 |
| ENSMUSG00000022102 | 2,117 | 0,000 | Dok2 | ENSMUSG00000028270 | 2,198 | 0,000 | Gbp2 |
| ENSMUSG00000097141 | 2,111 | 0,022 | Gm10524 | ENSMUSG00000039956 | 2,197 | 0,000 | Mrap |
| ENSMUSG00000092680 | 2,106 | 0,000 | Snord55 | ENSMUSG00000076498 | 2,179 | 0,005 | Trbc2 |
| ENSMUSG00000028862 | 2,095 | 0,000 | Map3k6 | ENSMUSG00000028862 | 2,175 | 0,000 | Map3k6 |
| ENSMUSG00000022602 | 2,095 | 0,000 | Arc | ENSMUSG00000025216 | 2,157 | 0,000 | Lbx1 |
| ENSMUSG00000020108 | 2,089 | 0,000 | Ddit4 | ENSMUSG00000080413 | 2,157 | 0,000 | Gm26473 |
| ENSMUSG00000076258 | 2,076 | 0,003 | Gm23935 | ENSMUSG00000025746 | 2,153 | 0,000 | Il6 |
| ENSMUSG00000026581 | 2,052 | 0,000 | Sell | ENSMUSG00000086825 | 2,139 | 0,005 | Gm15675 |
| ENSMUSG00000041046 | 2,048 | 0,000 | Ramp3 | ENSMUSG00000032661 | 2,133 | 0,000 | Oas3 |
| ENSMUSG00000026580 | 2,039 | 0,017 | Selp | ENSMUSG00000035042 | 2,126 | 0,000 | Ccl5 |
| ENSMUSG00000022126 | 2,031 | 0,001 | Acod1 | ENSMUSG00000058626 | 2,117 | 0,041 | Capn11 |
| ENSMUSG00000032724 | 2,022 | 0,000 | Abtb2 | ENSMUSG00000050830 | 2,116 | 0,007 | Vwc2 |
| ENSMUSG00000025216 | 2,017 | 0,000 | Lbx1 | ENSMUSG00000027360 | 2,103 | 0,000 | Hdc |

| | | | | | |
|---|---|---|---|---|---|
| ENSMUSG00000078817 | 2,015 | 0,000 Nlrp12 | ENSMUSG00000021123 | 2,097 | 0,000 Rdh12 |
| ENSMUSG00000038357 | 2,014 | 0,004 Camp | ENSMUSG00000070407 | 2,096 | 0,000 Hs3st3b1 |
| ENSMUSG00000110588 | 2,012 | 0,003 Gm45774 | ENSMUSG00000092572 | 2,082 | 0,000 Serpinb10 |
| ENSMUSG00000039405 | 1,994 | 0,000 Prss23 | ENSMUSG00000106609 | 2,079 | 0,001 Gm43181 |
| ENSMUSG00000070691 | 1,992 | 0,000 Runx3 | ENSMUSG00000078763 | 2,079 | 0,000 Slfn1 |
| ENSMUSG00000092920 | 1,990 | 0,002 Mirt2 | ENSMUSG00000061048 | 2,078 | 0,031 Cdh3 |
| ENSMUSG00000024424 | 1,989 | 0,000 Ttc39c | ENSMUSG00000040264 | 2,069 | 0,000 Gbp2b |
| ENSMUSG00000038793 | 1,970 | 0,002 Lefty1 | ENSMUSG00000044103 | 2,067 | 0,001 Il1f9 |
| ENSMUSG00000022582 | 1,958 | 0,000 Ly6g | ENSMUSG00000022534 | 2,065 | 0,001 Mefv |
| ENSMUSG00000031444 | 1,946 | 0,000 F10 | ENSMUSG00000053007 | 2,055 | 0,000 Creb5 |
| ENSMUSG00000047293 | 1,940 | 0,005 Gpr15 | ENSMUSG00000100625 | 2,051 | 0,004 1700016G22Rik |
| ENSMUSG00000023132 | 1,922 | 0,008 Gzma | ENSMUSG00000022586 | 2,049 | 0,007 Ly6i |
| ENSMUSG00000097165 | 1,917 | 0,003 2210008F06Rik | ENSMUSG00000022479 | 2,044 | 0,000 Vdr |
| ENSMUSG00000100113 | 1,917 | 0,018 Gm28592 | ENSMUSG00000007107 | 2,043 | 0,022 Atp1a4 |
| ENSMUSG00000025473 | 1,909 | 0,000 Adam8 | ENSMUSG00000079362 | 2,041 | 0,000 Gm43302 |
| ENSMUSG00000027978 | 1,907 | 0,000 Prss12 | ENSMUSG00000020108 | 2,036 | 0,000 Ddit4 |
| ENSMUSG00000032578 | 1,902 | 0,000 Cish | ENSMUSG00000039405 | 2,015 | 0,000 Prss23 |
| ENSMUSG00000021123 | 1,901 | 0,000 Rdh12 | ENSMUSG00000097714 | 2,011 | 0,001 Gm20109 |
| ENSMUSG00000086815 | 1,901 | 0,019 3110082J24Rik | ENSMUSG00000097472 | 2,007 | 0,038 Gm26586 |
| ENSMUSG00000022902 | 1,895 | 0,023 Stfa2 | ENSMUSG00000054203 | 2,006 | 0,001 Ifi205 |
| ENSMUSG00000040121 | 1,893 | 0,035 Rep15 | ENSMUSG00000110588 | 2,004 | 0,010 Gm45774 |
| ENSMUSG00000082292 | 1,884 | 0,000 Gm12250 | ENSMUSG00000021750 | 2,002 | 0,000 Fam107a |
| ENSMUSG00000079597 | 1,876 | 0,010 Gm5483 | ENSMUSG00000109560 | 1,997 | 0,010 Gm8463 |
| ENSMUSG00000020826 | 1,867 | 0,000 Nos2 | ENSMUSG00000024424 | 1,987 | 0,000 Ttc39c |
| ENSMUSG00000113701 | 1,867 | 0,015 B230303A05Rik | ENSMUSG00000102051 | 1,983 | 0,000 I830127L07Rik |
| ENSMUSG00000058624 | 1,866 | 0,000 Gda | ENSMUSG00000063767 | 1,968 | 0,046 S100a7a |
| ENSMUSG00000052188 | 1,864 | 0,001 Gm14964 | ENSMUSG00000027978 | 1,964 | 0,000 Prss12 |
| ENSMUSG00000074052 | 1,861 | 0,049 BC048644 | ENSMUSG00000047798 | 1,963 | 0,000 Cd300lf |
| ENSMUSG00000038201 | 1,856 | 0,000 Kcna7 | ENSMUSG00000105504 | 1,960 | 0,000 Gbp5 |
| ENSMUSG00000080413 | 1,856 | 0,001 Gm26473 | ENSMUSG00000060808 | 1,960 | 0,038 B9d1os |
| ENSMUSG00000049130 | 1,848 | 0,000 C5ar1 | ENSMUSG00000052749 | 1,959 | 0,000 Trim30b |
| ENSMUSG00000097714 | 1,842 | 0,002 Gm20109 | ENSMUSG00000022582 | 1,956 | 0,000 Ly6g |
| ENSMUSG00000001227 | 1,838 | 0,000 Sema6b | ENSMUSG00000025473 | 1,952 | 0,000 Adam8 |
| ENSMUSG00000050370 | 1,837 | 0,024 Ch25h | ENSMUSG00000062157 | 1,940 | 0,001 Ifnlr1 |
| ENSMUSG00000096719 | 1,835 | 0,032 Mrgpra2b | ENSMUSG00000058624 | 1,936 | 0,000 Gda |
| ENSMUSG00000068227 | 1,832 | 0,001 Il2rb | ENSMUSG00000021903 | 1,935 | 0,000 Galnt15 |
| ENSMUSG00000103164 | 1,829 | 0,001 Gm38150 | ENSMUSG00000022102 | 1,922 | 0,000 Dok2 |
| ENSMUSG00000095562 | 1,822 | 0,026 Gm21887 | ENSMUSG00000035352 | 1,920 | 0,000 Ccl12 |
| ENSMUSG00000020310 | 1,810 | 0,000 Madcam1 | ENSMUSG00000031778 | 1,914 | 0,000 Cx3cl1 |
| ENSMUSG00000032484 | 1,796 | 0,008 Ngp | ENSMUSG00000021125 | 1,914 | 0,000 Arg2 |
| ENSMUSG00000051397 | 1,795 | 0,005 Tacstd2 | ENSMUSG00000100113 | 1,904 | 0,038 Gm28592 |
| ENSMUSG00000026822 | 1,793 | 0,023 Lcn2 | ENSMUSG00000005800 | 1,898 | 0,000 Mmp8 |
| ENSMUSG00000053214 | 1,773 | 0,008 Gm9899 | ENSMUSG00000026399 | 1,884 | 0,000 Cd55 |
| ENSMUSG00000086481 | 1,773 | 0,044 Gm11707 | ENSMUSG00000104713 | 1,884 | 0,000 Gbp6 |
| ENSMUSG00000040711 | 1,768 | 0,000 Sh3pxd2b | ENSMUSG00000074896 | 1,883 | 0,000 Ifit3 |
| ENSMUSG00000054342 | 1,768 | 0,000 Kcnn4 | ENSMUSG00000032446 | 1,860 | 0,047 Eomes |
| ENSMUSG00000065145 | 1,758 | 0,028 Vaultrc5 | ENSMUSG00000041046 | 1,852 | 0,000 Ramp3 |

| | | | | | |
|---|---|---|---|---|---|
| ENSMUSG00000032204 | 1,753 | 0,000 Aqp9 | ENSMUSG00000034997 | 1,850 | 0,043 Htr2a |
| ENSMUSG00000048572 | 1,751 | 0,000 Tmem252 | ENSMUSG00000023206 | 1,847 | 0,000 Il15ra |
| ENSMUSG00000106609 | 1,751 | 0,000 Gm43181 | ENSMUSG00000030144 | 1,843 | 0,002 Clec4d |
| ENSMUSG00000025746 | 1,741 | 0,000 Il6 | ENSMUSG00000045094 | 1,841 | 0,004 Arhgef37 |
| ENSMUSG00000023206 | 1,734 | 0,000 Il15ra | ENSMUSG00000022126 | 1,831 | 0,008 Acod1 |
| ENSMUSG00000032860 | 1,722 | 0,000 P2ry2 | ENSMUSG00000053101 | 1,824 | 0,000 Gpr141 |
| ENSMUSG00000038917 | 1,721 | 0,000 3930402G23Rik | ENSMUSG00000046159 | 1,821 | 0,005 Chrm3 |
| ENSMUSG00000102051 | 1,721 | 0,001 I830127L07Rik | ENSMUSG00000051748 | 1,821 | 0,004 Wfdc21 |
| ENSMUSG00000001131 | 1,717 | 0,000 Timp1 | ENSMUSG00000030142 | 1,813 | 0,001 Clec4e |
| ENSMUSG00000099137 | 1,716 | 0,012 Gm10603 | ENSMUSG00000032420 | 1,808 | 0,000 Nt5e |
| ENSMUSG00000021125 | 1,713 | 0,000 Arg2 | ENSMUSG00000043873 | 1,806 | 0,003 Chil5 |
| ENSMUSG00000053338 | 1,710 | 0,006 Tarm1 | ENSMUSG00000032204 | 1,787 | 0,000 Aqp9 |
| ENSMUSG00000030069 | 1,705 | 0,002 Prok2 | ENSMUSG00000076309 | 1,786 | 0,032 Mir147 |
| ENSMUSG00000029372 | 1,702 | 0,003 Ppbp | ENSMUSG00000079339 | 1,780 | 0,006 Ifit1bl1 |
| ENSMUSG00000007107 | 1,696 | 0,033 Atp1a4 | ENSMUSG00000037681 | 1,778 | 0,045 Esyt3 |
| ENSMUSG00000025573 | 1,690 | 0,000 6030468B19Rik | ENSMUSG00000108985 | 1,773 | 0,005 A930030B08Rik |
| ENSMUSG00000044103 | 1,690 | 0,004 Il1f9 | ENSMUSG00000064380 | 1,768 | 0,022 Gm26448 |
| ENSMUSG00000076498 | 1,689 | 0,012 Trbc2 | ENSMUSG00000020027 | 1,768 | 0,000 Socs2 |
| ENSMUSG00000062515 | 1,680 | 0,000 Fabp4 | ENSMUSG00000059657 | 1,767 | 0,001 Stfa2l1 |
| ENSMUSG00000102037 | 1,678 | 0,000 Bcl2a1a | ENSMUSG00000036863 | 1,762 | 0,003 Syde2 |
| ENSMUSG00000063767 | 1,676 | 0,037 S100a7a | ENSMUSG00000027670 | 1,760 | 0,000 Ocstamp |
| ENSMUSG00000022094 | 1,674 | 0,000 Slc39a14 | ENSMUSG00000053338 | 1,758 | 0,011 Tarm1 |
| ENSMUSG00000090877 | 1,671 | 0,039 Hspa1b | ENSMUSG00000096929 | 1,743 | 0,036 A330023F24Rik |
| ENSMUSG00000030878 | 1,669 | 0,000 Cdr2 | ENSMUSG00000024730 | 1,742 | 0,048 Ms4a8a |
| ENSMUSG00000018500 | 1,663 | 0,000 Adora2b | ENSMUSG00000032724 | 1,741 | 0,000 Abtb2 |
| ENSMUSG00000054203 | 1,658 | 0,006 Ifi205 | ENSMUSG00000026220 | 1,735 | 0,005 Slc16a14 |
| ENSMUSG00000031104 | 1,649 | 0,000 Rab33a | ENSMUSG00000032105 | 1,729 | 0,001 Pdzd3 |
| ENSMUSG00000028019 | 1,643 | 0,000 Pdgfc | ENSMUSG00000048572 | 1,722 | 0,001 Tmem252 |
| ENSMUSG00000052270 | 1,641 | 0,000 Fpr2 | ENSMUSG00000028037 | 1,720 | 0,000 Ifi44 |
| ENSMUSG00000030325 | 1,640 | 0,007 Klrb1c | ENSMUSG00000031444 | 1,719 | 0,000 F10 |
| ENSMUSG00000042265 | 1,633 | 0,000 Trem1 | ENSMUSG00000070691 | 1,713 | 0,007 Runx3 |
| ENSMUSG00000076309 | 1,625 | 0,021 Mir147 | ENSMUSG00000030069 | 1,713 | 0,004 Prok2 |
| ENSMUSG00000027068 | 1,620 | 0,005 Dhrs9 | ENSMUSG00000090877 | 1,700 | 0,049 Hspa1b |
| ENSMUSG00000024772 | 1,609 | 0,000 Ehd1 | ENSMUSG00000103164 | 1,699 | 0,007 Gm38150 |
| ENSMUSG00000009633 | 1,605 | 0,000 G0s2 | ENSMUSG00000105813 | 1,698 | 0,043 Gm42928 |
| ENSMUSG00000092572 | 1,598 | 0,000 Serpinb10 | ENSMUSG00000079597 | 1,696 | 0,030 Gm5483 |
| ENSMUSG00000025044 | 1,595 | 0,000 Msr1 | ENSMUSG00000040711 | 1,687 | 0,000 Sh3pxd2b |
| ENSMUSG00000078161 | 1,594 | 0,000 Erich3 | ENSMUSG00000000386 | 1,676 | 0,043 Mx1 |
| ENSMUSG00000058603 | 1,593 | 0,000 Rpl28-ps1 | ENSMUSG00000052188 | 1,674 | 0,006 Gm14964 |
| ENSMUSG00000001493 | 1,589 | 0,001 Meox1 | ENSMUSG00000081965 | 1,673 | 0,021 Gm11620 |
| ENSMUSG00000001025 | 1,585 | 0,000 S100a6 | ENSMUSG00000070304 | 1,659 | 0,000 Scn2b |
| ENSMUSG00000046159 | 1,581 | 0,006 Chrm3 | ENSMUSG00000022094 | 1,653 | 0,000 Slc39a14 |
| ENSMUSG00000068262 | 1,577 | 0,032 Gm5879 | ENSMUSG00000009185 | 1,653 | 0,021 Ccl8 |
| ENSMUSG00000074361 | 1,573 | 0,000 C5ar2 | ENSMUSG00000015970 | 1,651 | 0,050 Chdh |
| ENSMUSG00000079362 | 1,573 | 0,000 Gm43302 | ENSMUSG00000100147 | 1,647 | 0,035 1700047M11Rik |
| ENSMUSG00000005864 | 1,569 | 0,020 Cnga2 | ENSMUSG00000021614 | 1,638 | 0,000 Vcan |
| ENSMUSG00000021903 | 1,566 | 0,002 Galnt15 | ENSMUSG00000020638 | 1,634 | 0,000 Cmpk2 |

| | | | | | | | |
|---|---|---|---|---|---|---|---|
| ENSMUSG00000032496 | 1,566 | 0,036 | Ltf | ENSMUSG00000076258 | 1,632 | 0,037 | Gm23935 |
| ENSMUSG00000078783 | 1,565 | 0,001 | Gm9733 | ENSMUSG00000027338 | 1,631 | 0,000 | Prnd |
| ENSMUSG00000051314 | 1,558 | 0,016 | Ffar2 | ENSMUSG00000000791 | 1,628 | 0,000 | Il12rb1 |
| ENSMUSG00000050896 | 1,555 | 0,041 | Rtn4rl2 | ENSMUSG00000041827 | 1,626 | 0,000 | Oasl1 |
| ENSMUSG00000031722 | 1,551 | 0,000 | Hp | ENSMUSG00000038357 | 1,626 | 0,036 | Camp |
| ENSMUSG00000026874 | 1,550 | 0,019 | Hc | ENSMUSG00000022512 | 1,624 | 0,011 | Cldn1 |
| ENSMUSG00000100147 | 1,549 | 0,019 | 1700047M11Rik | ENSMUSG00000030218 | 1,620 | 0,000 | Mgp |
| ENSMUSG00000114608 | 1,547 | 0,003 | AC238811.2 | ENSMUSG00000063820 | 1,617 | 0,004 | Arl9 |
| ENSMUSG00000060962 | 1,544 | 0,001 | Dmkn | ENSMUSG00000056966 | 1,614 | 0,007 | Gjc3 |
| ENSMUSG00000043165 | 1,535 | 0,005 | Lor | ENSMUSG00000020826 | 1,603 | 0,000 | Nos2 |
| ENSMUSG00000034226 | 1,532 | 0,000 | Rhov | ENSMUSG00000046908 | 1,597 | 0,000 | Ltb4r1 |
| ENSMUSG00000060019 | 1,531 | 0,000 | Gm10073 | ENSMUSG00000038793 | 1,594 | 0,022 | Lefty1 |
| ENSMUSG00000026981 | 1,530 | 0,000 | Il1rn | ENSMUSG00000055866 | 1,593 | 0,000 | Per2 |
| ENSMUSG00000032327 | 1,517 | 0,000 | Stra6 | ENSMUSG00000018500 | 1,591 | 0,000 | Adora2b |
| ENSMUSG00000029321 | 1,505 | 0,002 | Slc10a6 | ENSMUSG00000051054 | 1,582 | 0,022 | 1700014D04Rik |
| ENSMUSG00000021750 | 1,501 | 0,000 | Fam107a | ENSMUSG00000028019 | 1,582 | 0,000 | Pdgfc |
| ENSMUSG00000060131 | 1,496 | 0,000 | Atp8b4 | ENSMUSG00000022584 | 1,580 | 0,011 | Ly6c2 |
| ENSMUSG00000029314 | 1,494 | 0,000 | Gpat3 | ENSMUSG00000027219 | 1,577 | 0,007 | Slc28a2 |
| ENSMUSG00000047986 | 1,492 | 0,021 | Palm3 | ENSMUSG00000043448 | 1,571 | 0,000 | Gjc2 |
| ENSMUSG00000059657 | 1,489 | 0,007 | Stfa2l1 | ENSMUSG00000053461 | 1,571 | 0,004 | Hhipl2 |
| ENSMUSG00000050014 | 1,488 | 0,022 | Apol10b | ENSMUSG00000053541 | 1,569 | 0,013 | Gm4759 |
| ENSMUSG00000021091 | 1,483 | 0,002 | Serpina3n | ENSMUSG00000027068 | 1,566 | 0,012 | Dhrs9 |
| ENSMUSG00000017144 | 1,480 | 0,000 | Rnd3 | ENSMUSG00000029321 | 1,563 | 0,003 | Slc10a6 |
| ENSMUSG00000025355 | 1,477 | 0,000 | Mmp19 | ENSMUSG00000053214 | 1,553 | 0,033 | Gm9899 |
| ENSMUSG00000037872 | 1,476 | 0,044 | Ackr1 | ENSMUSG00000003484 | 1,550 | 0,003 | Cyp4f18 |
| ENSMUSG00000053101 | 1,476 | 0,000 | Gpr141 | ENSMUSG00000013974 | 1,547 | 0,000 | Mcemp1 |
| ENSMUSG00000040473 | 1,466 | 0,000 | Cfap69 | ENSMUSG00000051397 | 1,546 | 0,037 | Tacstd2 |
| ENSMUSG00000045664 | 1,466 | 0,000 | Cdc42ep2 | ENSMUSG00000020641 | 1,545 | 0,001 | Rsad2 |
| ENSMUSG00000022952 | 1,465 | 0,000 | Runx1 | ENSMUSG00000001227 | 1,544 | 0,000 | Sema6b |
| ENSMUSG00000027219 | 1,456 | 0,008 | Slc28a2 | ENSMUSG00000045664 | 1,540 | 0,000 | Cdc42ep2 |
| ENSMUSG00000021614 | 1,452 | 0,000 | Vcan | ENSMUSG00000030878 | 1,533 | 0,000 | Cdr2 |
| ENSMUSG00000022534 | 1,450 | 0,009 | Mefv | ENSMUSG00000002831 | 1,532 | 0,048 | Plin4 |
| ENSMUSG00000026068 | 1,440 | 0,000 | Il18rap | ENSMUSG00000042265 | 1,527 | 0,000 | Trem1 |
| ENSMUSG00000071001 | 1,437 | 0,001 | Hrct1 | ENSMUSG00000035873 | 1,526 | 0,000 | Pawr |
| ENSMUSG00000105504 | 1,436 | 0,000 | Gbp5 | ENSMUSG00000033508 | 1,517 | 0,000 | Asprv1 |
| ENSMUSG00000022408 | 1,433 | 0,012 | Fam83f | ENSMUSG00000038917 | 1,512 | 0,016 | 3930402G23Rik |
| ENSMUSG00000026220 | 1,430 | 0,017 | Slc16a14 | ENSMUSG00000031722 | 1,510 | 0,000 | Hp |
| ENSMUSG00000032661 | 1,419 | 0,001 | Oas3 | ENSMUSG00000050953 | 1,504 | 0,000 | Gja1 |
| ENSMUSG00000037446 | 1,418 | 0,016 | Tulp1 | ENSMUSG00000030107 | 1,499 | 0,000 | Usp18 |
| ENSMUSG00000029307 | 1,416 | 0,007 | Dmp1 | ENSMUSG00000028841 | 1,494 | 0,036 | Cnksr1 |
| ENSMUSG00000047759 | 1,413 | 0,000 | Hs3st3a1 | ENSMUSG00000042549 | 1,483 | 0,000 | Map2k3os |
| ENSMUSG00000030208 | 1,411 | 0,000 | Emp1 | ENSMUSG00000030187 | 1,478 | 0,000 | Klra2 |
| ENSMUSG00000001473 | 1,408 | 0,000 | Tubb6 | ENSMUSG00000033213 | 1,475 | 0,005 | AA467197 |
| ENSMUSG00000032105 | 1,407 | 0,010 | Pdzd3 | ENSMUSG00000020122 | 1,470 | 0,000 | Egfr |
| ENSMUSG00000058443 | 1,405 | 0,001 | Rpl10-ps3 | ENSMUSG00000032578 | 1,470 | 0,000 | Cish |
| ENSMUSG00000020077 | 1,400 | 0,000 | Srgn | ENSMUSG00000009633 | 1,469 | 0,000 | G0s2 |
| ENSMUSG00000026303 | 1,397 | 0,006 | Mlph | ENSMUSG00000001131 | 1,469 | 0,002 | Timp1 |

| | | | | | |
|---|---|---|---|---|---|
| ENSMUSG00000026399 | 1,396 | 0,000 Cd55 | ENSMUSG00000024772 | 1,461 | 0,000 Ehd1 |
| ENSMUSG00000104713 | 1,394 | 0,000 Gbp6 | ENSMUSG00000007877 | 1,461 | 0,016 Tcap |
| ENSMUSG00000064246 | 1,388 | 0,000 Chil1 | ENSMUSG00000055093 | 1,460 | 0,032 Gm8430 |
| ENSMUSG00000020122 | 1,387 | 0,000 Egfr | ENSMUSG00000020427 | 1,459 | 0,000 Igfbp3 |
| ENSMUSG00000085196 | 1,383 | 0,049 Gm14963 | ENSMUSG00000007033 | 1,458 | 0,001 Hspa1l |
| ENSMUSG00000042549 | 1,382 | 0,000 Map2k3os | ENSMUSG00000009376 | 1,453 | 0,000 Met |
| ENSMUSG00000051748 | 1,382 | 0,028 Wfdc21 | ENSMUSG00000108738 | 1,453 | 0,017 Gm44777 |
| ENSMUSG00000070304 | 1,376 | 0,000 Scn2b | ENSMUSG00000067297 | 1,452 | 0,049 Ifit1bl2 |
| ENSMUSG00000025856 | 1,368 | 0,000 Pdgfa | ENSMUSG00000001444 | 1,450 | 0,040 Tbx21 |
| ENSMUSG00000112567 | 1,365 | 0,015 Gm32899 | ENSMUSG00000030208 | 1,450 | 0,000 Emp1 |
| ENSMUSG00000068240 | 1,363 | 0,003 Gm11808 | ENSMUSG00000060131 | 1,446 | 0,001 Atp8b4 |
| ENSMUSG00000007659 | 1,358 | 0,000 Bcl2l1 | ENSMUSG00000031762 | 1,445 | 0,001 Mt2 |
| ENSMUSG00000033213 | 1,357 | 0,004 AA467197 | ENSMUSG00000092680 | 1,436 | 0,020 Snord55 |
| ENSMUSG00000005800 | 1,356 | 0,005 Mmp8 | ENSMUSG00000035283 | 1,432 | 0,000 Adrb1 |
| ENSMUSG00000020427 | 1,355 | 0,000 Igfbp3 | ENSMUSG00000089726 | 1,426 | 0,000 Mir17hg |
| ENSMUSG00000001444 | 1,351 | 0,036 Tbx21 | ENSMUSG00000035692 | 1,425 | 0,000 Isg15 |
| ENSMUSG00000084159 | 1,349 | 0,010 Gm12696 | ENSMUSG00000030303 | 1,424 | 0,000 Far2 |
| ENSMUSG00000056290 | 1,348 | 0,013 Ms4a4b | ENSMUSG00000105338 | 1,415 | 0,000 Gm43802 |
| ENSMUSG00000097558 | 1,346 | 0,014 Gm26902 | ENSMUSG00000071347 | 1,407 | 0,003 C1qtnf9 |
| ENSMUSG00000111966 | 1,345 | 0,000 Gm5781 | ENSMUSG00000044461 | 1,406 | 0,022 Shisa2 |
| ENSMUSG00000022512 | 1,343 | 0,013 Cldn1 | ENSMUSG00000088634 | 1,403 | 0,041 Gm24958 |
| ENSMUSG00000023087 | 1,330 | 0,000 Noct | ENSMUSG00000104880 | 1,400 | 0,042 AL645522.1 |
| ENSMUSG00000030748 | 1,328 | 0,000 Il4ra | ENSMUSG00000078853 | 1,394 | 0,000 Igtp |
| ENSMUSG00000079419 | 1,327 | 0,009 Ms4a6c | ENSMUSG00000064441 | 1,388 | 0,003 Snord37 |
| ENSMUSG00000110686 | 1,312 | 0,006 Gm45855 | ENSMUSG00000027514 | 1,384 | 0,000 Zbp1 |
| ENSMUSG00000027338 | 1,311 | 0,001 Prnd | ENSMUSG00000025573 | 1,381 | 0,006 6030468B19Rik |
| ENSMUSG00000024164 | 1,306 | 0,004 C3 | ENSMUSG00000008307 | 1,370 | 0,010 1700109H08Rik |
| ENSMUSG00000030303 | 1,298 | 0,000 Far2 | ENSMUSG00000045932 | 1,370 | 0,000 Ifit2 |
| ENSMUSG00000027849 | 1,292 | 0,039 Syt6 | ENSMUSG00000027849 | 1,366 | 0,040 Syt6 |
| ENSMUSG00000073293 | 1,284 | 0,004 Nudt10 | ENSMUSG00000036185 | 1,361 | 0,029 Sapcd1 |
| ENSMUSG00000053111 | 1,283 | 0,025 Fank1 | ENSMUSG00000034438 | 1,360 | 0,001 Gbp8 |
| ENSMUSG00000094497 | 1,283 | 0,010 Gm8210 | ENSMUSG00000031710 | 1,350 | 0,016 Ucp1 |
| ENSMUSG00000105699 | 1,281 | 0,000 Gm43703 | ENSMUSG00000037820 | 1,346 | 0,000 Tgm2 |
| ENSMUSG00000030187 | 1,275 | 0,000 Klra2 | ENSMUSG00000001493 | 1,345 | 0,016 Meox1 |
| ENSMUSG00000050621 | 1,274 | 0,003 Rps27rt | ENSMUSG00000009248 | 1,345 | 0,001 Ascl2 |
| ENSMUSG00000046223 | 1,273 | 0,000 Plaur | ENSMUSG00000029314 | 1,340 | 0,000 Gpat3 |
| ENSMUSG00000074874 | 1,267 | 0,000 Ctla2b | ENSMUSG00000038037 | 1,339 | 0,000 Socs1 |
| ENSMUSG00000036863 | 1,266 | 0,015 Syde2 | ENSMUSG00000064246 | 1,334 | 0,000 Chil1 |
| ENSMUSG00000055866 | 1,266 | 0,000 Per2 | ENSMUSG00000030214 | 1,334 | 0,000 Plbd1 |
| ENSMUSG00000079445 | 1,254 | 0,000 B3gnt7 | ENSMUSG00000049130 | 1,330 | 0,009 C5ar1 |
| ENSMUSG00000040264 | 1,254 | 0,001 Gbp2b | ENSMUSG00000106258 | 1,329 | 0,031 Rpl7a-ps7 |
| ENSMUSG00000026833 | 1,253 | 0,000 Olfm1 | ENSMUSG00000001025 | 1,315 | 0,000 S100a6 |
| ENSMUSG00000115358 | 1,250 | 0,002 AC127578.1 | ENSMUSG00000031958 | 1,313 | 0,000 Ldhd |
| ENSMUSG00000045094 | 1,245 | 0,048 Arhgef37 | ENSMUSG00000039699 | 1,309 | 0,000 Batf2 |
| ENSMUSG00000000791 | 1,236 | 0,003 Il12rb1 | ENSMUSG00000073739 | 1,309 | 0,023 Gm16287 |
| ENSMUSG00000070348 | 1,233 | 0,000 Ccnd1 | ENSMUSG00000024066 | 1,308 | 0,005 Xdh |
| ENSMUSG00000041959 | 1,228 | 0,000 S100a10 | ENSMUSG00000097572 | 1,304 | 0,005 Gm26797 |

| | | | | | |
|---|---|---|---|---|---|
| ENSMUSG00000013974 | 1,227 | 0,002 Mcemp1 | ENSMUSG00000073489 | 1,303 | 0,000 Ifi204 |
| ENSMUSG00000035283 | 1,227 | 0,000 Adrb1 | ENSMUSG00000002204 | 1,302 | 0,000 Napsa |
| ENSMUSG00000114432 | 1,226 | 0,001 AC160962.1 | ENSMUSG00000100465 | 1,300 | 0,002 Gm29264 |
| ENSMUSG00000000440 | 1,225 | 0,000 Pparg | ENSMUSG00000003070 | 1,292 | 0,000 Efna2 |
| ENSMUSG00000086825 | 1,224 | 0,043 Gm15675 | ENSMUSG00000078920 | 1,291 | 0,000 Ifi47 |
| ENSMUSG00000104603 | 1,223 | 0,011 AL845457.1 | ENSMUSG00000098754 | 1,284 | 0,000 Prn |
| ENSMUSG00000073421 | 1,223 | 0,000 H2-Ab1 | ENSMUSG00000051444 | 1,281 | 0,000 Bbs12 |
| ENSMUSG00000049892 | 1,218 | 0,002 Rasd1 | ENSMUSG00000064791 | 1,276 | 0,006 Snord14e |
| ENSMUSG00000038037 | 1,217 | 0,000 Socs1 | ENSMUSG00000029410 | 1,271 | 0,000 Ppef2 |
| ENSMUSG00000050549 | 1,213 | 0,000 Fam241a | ENSMUSG00000078161 | 1,269 | 0,009 Erich3 |
| ENSMUSG00000015850 | 1,209 | 0,001 Adamtsl4 | ENSMUSG00000031997 | 1,268 | 0,032 Trpc6 |
| ENSMUSG00000068105 | 1,207 | 0,001 Tnfrsf13c | ENSMUSG00000065987 | 1,254 | 0,013 Cd209b |
| ENSMUSG00000041608 | 1,202 | 0,032 Entpd3 | ENSMUSG00000114724 | 1,253 | 0,000 CT009713.7 |
| ENSMUSG00000081984 | 1,201 | 0,001 Dnajb3 | ENSMUSG00000040473 | 1,252 | 0,001 Cfap69 |
| ENSMUSG00000052749 | 1,200 | 0,001 Trim30b | ENSMUSG00000030562 | 1,248 | 0,013 Nox4 |
| ENSMUSG00000031997 | 1,198 | 0,021 Trpc6 | ENSMUSG00000025498 | 1,248 | 0,000 Irf7 |
| ENSMUSG00000026180 | 1,198 | 0,000 Cxcr2 | ENSMUSG00000066058 | 1,248 | 0,026 Cldn19 |
| ENSMUSG00000044734 | 1,194 | 0,004 Serpinb1a | ENSMUSG00000054342 | 1,245 | 0,000 Kcnn4 |
| ENSMUSG00000026558 | 1,194 | 0,000 Uck2 | ENSMUSG00000086124 | 1,242 | 0,016 A530076I17Rik |
| ENSMUSG00000040714 | 1,192 | 0,007 Klc3 | ENSMUSG00000110697 | 1,240 | 0,000 Gm31718 |
| ENSMUSG00000038059 | 1,189 | 0,000 Smim3 | ENSMUSG00000049892 | 1,238 | 0,004 Rasd1 |
| ENSMUSG00000045106 | 1,188 | 0,036 Ccdc73 | ENSMUSG00000034226 | 1,237 | 0,005 Rhov |
| ENSMUSG00000052212 | 1,185 | 0,046 Cd177 | ENSMUSG00000017144 | 1,234 | 0,000 Rnd3 |
| ENSMUSG00000033644 | 1,181 | 0,001 Piwil2 | ENSMUSG00000033032 | 1,233 | 0,000 Afap1l1 |
| ENSMUSG00000016283 | 1,170 | 0,020 H2-M2 | ENSMUSG00000037428 | 1,231 | 0,004 Vgf |
| ENSMUSG00000002204 | 1,170 | 0,000 Napsa | ENSMUSG00000022952 | 1,224 | 0,000 Runx1 |
| ENSMUSG00000028480 | 1,164 | 0,000 Glipr2 | ENSMUSG00000040152 | 1,220 | 0,005 Thbs1 |
| ENSMUSG00000045502 | 1,164 | 0,001 Hcar2 | ENSMUSG00000062515 | 1,220 | 0,034 Fabp4 |
| ENSMUSG00000040274 | 1,159 | 0,000 Cdk6 | ENSMUSG00000097494 | 1,219 | 0,045 4933406C10Rik |
| ENSMUSG00000028874 | 1,155 | 0,000 Fgr | ENSMUSG00000041959 | 1,213 | 0,001 S100a10 |
| ENSMUSG00000058905 | 1,155 | 0,027 Gm10051 | ENSMUSG00000020077 | 1,205 | 0,000 Srgn |
| ENSMUSG00000087412 | 1,154 | 0,000 Gm15501 | ENSMUSG00000025355 | 1,205 | 0,000 Mmp19 |
| ENSMUSG00000015501 | 1,152 | 0,000 Hivep2 | ENSMUSG00000005672 | 1,202 | 0,000 Kit |
| ENSMUSG00000050953 | 1,151 | 0,000 Gja1 | ENSMUSG00000026068 | 1,201 | 0,004 Il18rap |
| ENSMUSG00000005087 | 1,150 | 0,000 Cd44 | ENSMUSG00000030711 | 1,196 | 0,001 Sult1a1 |
| ENSMUSG00000106895 | 1,135 | 0,025 Gm4754 | ENSMUSG00000003032 | 1,195 | 0,000 Klf4 |
| ENSMUSG00000024910 | 1,133 | 0,013 Ctsw | ENSMUSG00000026303 | 1,194 | 0,029 Mlph |
| ENSMUSG00000048807 | 1,126 | 0,000 Slc35e4 | ENSMUSG00000047759 | 1,191 | 0,002 Hs3st3a1 |
| ENSMUSG00000056749 | 1,121 | 0,000 Nfil3 | ENSMUSG00000074874 | 1,190 | 0,002 Ctla2b |
| ENSMUSG00000055632 | 1,120 | 0,011 Hmcn2 | ENSMUSG00000038354 | 1,189 | 0,036 Ankrd35 |
| ENSMUSG00000076441 | 1,119 | 0,000 Ass1 | ENSMUSG00000015850 | 1,188 | 0,001 Adamtsl4 |
| ENSMUSG00000033508 | 1,119 | 0,009 Asprv1 | ENSMUSG00000109032 | 1,187 | 0,004 Gm7972 |
| ENSMUSG00000071347 | 1,116 | 0,020 C1qtnf9 | ENSMUSG00000044734 | 1,186 | 0,007 Serpinb1a |
| ENSMUSG00000036594 | 1,110 | 0,000 H2-Aa | ENSMUSG00000104517 | 1,186 | 0,047 Gm18407 |
| ENSMUSG00000031700 | 1,104 | 0,000 Gpt2 | ENSMUSG00000079681 | 1,185 | 0,011 Zglp1 |
| ENSMUSG00000032643 | 1,101 | 0,000 Fhl3 | ENSMUSG00000050612 | 1,185 | 0,028 Txndc2 |
| ENSMUSG00000031958 | 1,101 | 0,002 Ldhd | ENSMUSG00000114432 | 1,183 | 0,002 AC160962.1 |

| | | | | | |
|---|---|---|---|---|---|
| ENSMUSG00000035873 | 1,100 | 0,003 Pawr | ENSMUSG00000024087 | 1,168 | 0,000 Cyp1b1 |
| ENSMUSG00000098754 | 1,095 | 0,001 Prn | ENSMUSG00000069874 | 1,165 | 0,000 Irgm2 |
| ENSMUSG00000024526 | 1,095 | 0,038 Cidea | ENSMUSG00000020836 | 1,164 | 0,000 Coro6 |
| ENSMUSG00000032470 | 1,095 | 0,000 Mras | ENSMUSG00000000303 | 1,161 | 0,008 Cdh1 |
| ENSMUSG00000092837 | 1,095 | 0,002 Rpph1 | ENSMUSG00000022528 | 1,156 | 0,000 Hes1 |
| ENSMUSG00000037820 | 1,095 | 0,000 Tgm2 | ENSMUSG00000028268 | 1,153 | 0,000 Gbp3 |
| ENSMUSG00000062456 | 1,090 | 0,001 Rpl9-ps6 | ENSMUSG00000020310 | 1,147 | 0,040 Madcam1 |
| ENSMUSG00000039960 | 1,090 | 0,000 Rhou | ENSMUSG00000061531 | 1,147 | 0,004 Tmem236 |
| ENSMUSG00000070780 | 1,086 | 0,000 Rbm47 | ENSMUSG00000041754 | 1,147 | 0,014 Trem3 |
| ENSMUSG00000086124 | 1,083 | 0,023 A530076I17Rik | ENSMUSG00000002108 | 1,144 | 0,000 Nr1h3 |
| ENSMUSG00000030214 | 1,082 | 0,000 Plbd1 | ENSMUSG00000024430 | 1,143 | 0,010 Cabyr |
| ENSMUSG00000070305 | 1,080 | 0,005 Mpzl3 | ENSMUSG00000027456 | 1,135 | 0,012 Sdcbp2 |
| ENSMUSG00000051444 | 1,077 | 0,000 Bbs12 | ENSMUSG00000030170 | 1,127 | 0,002 Wnt5b |
| ENSMUSG00000009376 | 1,075 | 0,003 Met | ENSMUSG00000032193 | 1,127 | 0,000 Ldlr |
| ENSMUSG00000060586 | 1,074 | 0,000 H2-Eb1 | ENSMUSG00000028480 | 1,127 | 0,000 Glipr2 |
| ENSMUSG00000031765 | 1,071 | 0,001 Mt1 | ENSMUSG00000024526 | 1,126 | 0,042 Cidea |
| ENSMUSG00000034591 | 1,066 | 0,013 Slc41a2 | ENSMUSG00000003271 | 1,122 | 0,035 Sult2b1 |
| ENSMUSG00000011305 | 1,064 | 0,028 Plin5 | ENSMUSG00000024659 | 1,121 | 0,007 Anxa1 |
| ENSMUSG00000042608 | 1,064 | 0,000 Stk40 | ENSMUSG00000040274 | 1,121 | 0,000 Cdk6 |
| ENSMUSG00000114635 | 1,061 | 0,000 CT010498.2 | ENSMUSG00000074361 | 1,118 | 0,003 C5ar2 |
| ENSMUSG00000033032 | 1,060 | 0,000 Afap1l1 | ENSMUSG00000047330 | 1,116 | 0,024 Kcne4 |
| ENSMUSG00000078193 | 1,060 | 0,028 Gm2000 | ENSMUSG00000086742 | 1,115 | 0,019 Gm16201 |
| ENSMUSG00000037225 | 1,059 | 0,004 Fgf2 | ENSMUSG00000097365 | 1,112 | 0,037 C030034L19Rik |
| ENSMUSG00000070343 | 1,055 | 0,011 Gm10288 | ENSMUSG00000020419 | 1,110 | 0,047 Hormad2 |
| ENSMUSG00000073802 | 1,054 | 0,000 Cdkn2b | ENSMUSG00000031896 | 1,110 | 0,005 Ctrl |
| ENSMUSG00000074227 | 1,054 | 0,043 Spint2 | ENSMUSG00000027377 | 1,107 | 0,050 Mall |
| ENSMUSG00000110697 | 1,052 | 0,001 Gm31718 | ENSMUSG00000034156 | 1,106 | 0,000 Tspoap1 |
| ENSMUSG00000061531 | 1,050 | 0,002 Tmem236 | ENSMUSG00000073293 | 1,105 | 0,018 Nudt10 |
| ENSMUSG00000060950 | 1,046 | 0,000 Trmt61a | ENSMUSG00000092626 | 1,101 | 0,034 9130230N09Rik |
| ENSMUSG00000083337 | 1,045 | 0,010 Gm11539 | ENSMUSG00000040253 | 1,101 | 0,001 Gbp7 |
| ENSMUSG00000030413 | 1,044 | 0,011 Pglyrp1 | ENSMUSG00000000792 | 1,101 | 0,000 Slc5a5 |
| ENSMUSG00000029322 | 1,044 | 0,000 Plac8 | ENSMUSG00000009378 | 1,099 | 0,001 Slc16a12 |
| ENSMUSG00000028328 | 1,040 | 0,000 Tmod1 | ENSMUSG00000074151 | 1,099 | 0,002 Nlrc5 |
| ENSMUSG00000056888 | 1,038 | 0,000 Glipr1 | ENSMUSG00000001473 | 1,096 | 0,000 Tubb6 |
| ENSMUSG00000025591 | 1,036 | 0,000 Tma16 | ENSMUSG00000023087 | 1,095 | 0,000 Noct |
| ENSMUSG00000078920 | 1,036 | 0,000 Ifi47 | ENSMUSG00000035295 | 1,094 | 0,002 Wdr38 |
| ENSMUSG00000069805 | 1,034 | 0,007 Fbp1 | ENSMUSG00000101462 | 1,091 | 0,026 Gm3052 |
| ENSMUSG00000032207 | 1,034 | 0,004 Lipc | ENSMUSG00000050621 | 1,090 | 0,017 Rps27rt |
| ENSMUSG00000046908 | 1,033 | 0,008 Ltb4r1 | ENSMUSG00000104671 | 1,090 | 0,042 Gm43062 |
| ENSMUSG00000031506 | 1,031 | 0,001 Ptpn7 | ENSMUSG00000046733 | 1,086 | 0,012 Gprc5a |
| ENSMUSG00000081926 | 1,028 | 0,034 Gm15536 | ENSMUSG00000033967 | 1,085 | 0,000 Rnf225 |
| ENSMUSG00000027514 | 1,028 | 0,001 Zbp1 | ENSMUSG00000099876 | 1,082 | 0,046 Gm29650 |
| ENSMUSG00000057657 | 1,028 | 0,022 Rps18-ps3 | ENSMUSG00000022237 | 1,078 | 0,002 Ankrd33b |
| ENSMUSG00000097365 | 1,027 | 0,040 C030034L19Rik | ENSMUSG00000039960 | 1,075 | 0,001 Rhou |
| ENSMUSG00000041754 | 1,016 | 0,018 Trem3 | ENSMUSG00000029561 | 1,072 | 0,000 Oasl2 |
| ENSMUSG00000022218 | 1,013 | 0,000 Tgm1 | ENSMUSG00000022026 | 1,070 | 0,003 Olfm4 |
| ENSMUSG00000105388 | 1,009 | 0,026 Rpl36a-ps2 | ENSMUSG00000029322 | 1,063 | 0,000 Plac8 |

| | | | | | |
|---|---|---|---|---|---|
| ENSMUSG00000036305 | 1,009 | 0,031 Rpl39-ps | ENSMUSG00000023341 | 1,062 | 0,000 Mx2 |
| ENSMUSG00000061024 | 1,007 | 0,000 Rrs1 | ENSMUSG00000019997 | 1,061 | 0,033 Ctgf |
| ENSMUSG00000024053 | 1,003 | 0,001 Emilin2 | ENSMUSG00000034591 | 1,059 | 0,020 Slc41a2 |
| ENSMUSG00000050335 | 1,001 | 0,000 Lgals3 | ENSMUSG00000001281 | 1,056 | 0,038 Itgb7 |
| ENSMUSG00000047844 | 1,000 | 0,007 Bex4 | ENSMUSG00000061815 | 1,055 | 0,006 Rufy4 |
| ENSMUSG00000039232 | 0,999 | 0,004 Stx11 | ENSMUSG00000005087 | 1,052 | 0,001 Cd44 |
| ENSMUSG00000037434 | 0,995 | 0,000 Slc30a1 | ENSMUSG00000110419 | 1,049 | 0,037 AW046200 |
| ENSMUSG00000024732 | 0,992 | 0,000 Ccdc86 | ENSMUSG00000070305 | 1,047 | 0,010 Mpzl3 |
| ENSMUSG00000071470 | 0,991 | 0,006 Ccnb1ip1 | ENSMUSG00000107331 | 1,046 | 0,001 Gm42732 |
| ENSMUSG00000060419 | 0,989 | 0,003 Rps16-ps2 | ENSMUSG00000074796 | 1,044 | 0,015 Slc4a11 |
| ENSMUSG00000044258 | 0,987 | 0,028 Ctla2a | ENSMUSG00000044258 | 1,044 | 0,028 Ctla2a |
| ENSMUSG00000028069 | 0,981 | 0,000 Gpatch4 | ENSMUSG00000090125 | 1,043 | 0,036 Pou3f1 |
| ENSMUSG00000021149 | 0,981 | 0,000 Gtpbp4 | ENSMUSG00000022346 | 1,042 | 0,000 Myc |
| ENSMUSG00000029343 | 0,979 | 0,001 Crybb1 | ENSMUSG00000022218 | 1,042 | 0,000 Tgm1 |
| ENSMUSG00000001542 | 0,978 | 0,000 Ell2 | ENSMUSG00000024910 | 1,041 | 0,040 Ctsw |
| ENSMUSG00000013415 | 0,978 | 0,011 Igf2bp1 | ENSMUSG00000035576 | 1,041 | 0,046 L3mbtl1 |
| ENSMUSG00000039942 | 0,976 | 0,044 Ptger4 | ENSMUSG00000040605 | 1,039 | 0,001 Bace2 |
| ENSMUSG00000078853 | 0,976 | 0,000 Igtp | ENSMUSG00000030748 | 1,037 | 0,000 Il4ra |
| ENSMUSG00000049999 | 0,976 | 0,000 Ppp1r3d | ENSMUSG00000097386 | 1,032 | 0,046 9130213A22Rik |
| ENSMUSG00000038335 | 0,975 | 0,000 Tsr1 | ENSMUSG00000024935 | 1,031 | 0,037 Slc1a1 |
| ENSMUSG00000002108 | 0,974 | 0,000 Nr1h3 | ENSMUSG00000027318 | 1,030 | 0,024 Adam33 |
| ENSMUSG00000109941 | 0,971 | 0,000 Exosc6 | ENSMUSG00000013415 | 1,026 | 0,010 Igf2bp1 |
| ENSMUSG00000031494 | 0,970 | 0,015 Cd209a | ENSMUSG00000026558 | 1,024 | 0,000 Uck2 |
| ENSMUSG00000025492 | 0,968 | 0,000 Ifitm3 | ENSMUSG00000049103 | 1,024 | 0,002 Ccr2 |
| ENSMUSG00000052040 | 0,967 | 0,000 Klf13 | ENSMUSG00000003469 | 1,021 | 0,021 Phyhip |
| ENSMUSG00000024610 | 0,959 | 0,000 Cd74 | ENSMUSG00000074743 | 1,020 | 0,024 Thbd |
| ENSMUSG00000024679 | 0,957 | 0,026 Ms4a6d | ENSMUSG00000109602 | 1,018 | 0,004 Gm31597 |
| ENSMUSG00000035042 | 0,954 | 0,049 Ccl5 | ENSMUSG00000002897 | 1,018 | 0,000 Il17ra |
| ENSMUSG00000070960 | 0,949 | 0,038 Gm19680 | ENSMUSG00000051065 | 1,015 | 0,004 Mb21d2 |
| ENSMUSG00000045725 | 0,948 | 0,011 Prr15 | ENSMUSG00000103839 | 1,015 | 0,034 Gm37607 |
| ENSMUSG00000057135 | 0,945 | 0,024 Scimp | ENSMUSG00000024843 | 1,014 | 0,000 Chka |
| ENSMUSG00000097346 | 0,945 | 0,049 Gm26619 | ENSMUSG00000068105 | 1,014 | 0,010 Tnfrsf13c |
| ENSMUSG00000018543 | 0,944 | 0,028 1700001P01Rik | ENSMUSG00000027580 | 1,014 | 0,000 Helz2 |
| ENSMUSG00000114884 | 0,943 | 0,023 AC168059.2 | ENSMUSG00000024899 | 1,014 | 0,003 Papss2 |
| ENSMUSG00000022951 | 0,939 | 0,000 Rcan1 | ENSMUSG00000020121 | 1,011 | 0,000 Srgap1 |
| ENSMUSG00000093677 | 0,937 | 0,001 Gm20712 | ENSMUSG00000028328 | 1,009 | 0,000 Tmod1 |
| ENSMUSG00000060438 | 0,936 | 0,003 Rps10-ps1 | ENSMUSG00000039232 | 1,009 | 0,006 Stx11 |
| ENSMUSG00000106847 | 0,934 | 0,000 Peg13 | ENSMUSG00000032561 | 1,003 | 0,006 Acpp |
| ENSMUSG00000020961 | 0,931 | 0,001 Ston2 | ENSMUSG00000055833 | 1,000 | 0,007 1700034H15Rik |
| ENSMUSG00000024899 | 0,929 | 0,004 Papss2 | ENSMUSG00000114884 | 0,999 | 0,036 AC168059.2 |
| ENSMUSG00000068762 | 0,926 | 0,016 Gstm6 | ENSMUSG00000030413 | 0,998 | 0,022 Pglyrp1 |
| ENSMUSG00000038521 | 0,926 | 0,050 C1s1 | ENSMUSG00000054435 | 0,994 | 0,022 Gimap4 |
| ENSMUSG00000053801 | 0,926 | 0,000 Grwd1 | ENSMUSG00000070427 | 0,989 | 0,000 Il18bp |
| ENSMUSG00000067288 | 0,925 | 0,005 Rps28 | ENSMUSG00000068762 | 0,987 | 0,030 Gstm6 |
| ENSMUSG00000029082 | 0,923 | 0,002 Bst1 | ENSMUSG00000024610 | 0,987 | 0,000 Cd74 |
| ENSMUSG00000096883 | 0,922 | 0,028 Shisa8 | ENSMUSG00000026180 | 0,986 | 0,000 Cxcr2 |
| ENSMUSG00000058761 | 0,920 | 0,000 Rnf169 | ENSMUSG00000045319 | 0,985 | 0,007 Proser2 |

| | | | | | | | |
|---|---|---|---|---|---|---|---|
| ENSMUSG00000046791 | 0,919 | 0,000 | Riox1 | ENSMUSG00000026107 | 0,983 | 0,002 | Nabp1 |
| ENSMUSG00000033255 | 0,919 | 0,030 | Gm5134 | ENSMUSG00000114487 | 0,982 | 0,024 | CT010498.1 |
| ENSMUSG00000027395 | 0,918 | 0,000 | Polr1b | ENSMUSG00000007659 | 0,982 | 0,000 | Bcl2l1 |
| ENSMUSG00000043421 | 0,916 | 0,000 | Hilpda | ENSMUSG00000015501 | 0,976 | 0,001 | Hivep2 |
| ENSMUSG00000022305 | 0,916 | 0,000 | Lrp12 | ENSMUSG00000075602 | 0,975 | 0,013 | Ly6a |
| ENSMUSG00000074796 | 0,914 | 0,016 | Slc4a11 | ENSMUSG00000050549 | 0,970 | 0,000 | Fam241a |
| ENSMUSG00000022237 | 0,911 | 0,005 | Ankrd33b | ENSMUSG00000060568 | 0,969 | 0,014 | Fam78b |
| ENSMUSG00000053175 | 0,910 | 0,027 | Bcl3 | ENSMUSG00000029082 | 0,968 | 0,002 | Bst1 |
| ENSMUSG00000040152 | 0,905 | 0,040 | Thbs1 | ENSMUSG00000020205 | 0,967 | 0,000 | Phlda1 |
| ENSMUSG00000079547 | 0,905 | 0,042 | H2-DMb1 | ENSMUSG00000054404 | 0,967 | 0,005 | Slfn5 |
| ENSMUSG00000093327 | 0,903 | 0,000 | Mir5107 | ENSMUSG00000056501 | 0,964 | 0,004 | Cebpb |
| ENSMUSG00000036381 | 0,900 | 0,002 | P2ry14 | ENSMUSG00000039236 | 0,961 | 0,003 | Isg20 |
| ENSMUSG00000073775 | 0,900 | 0,000 | Kti12 | ENSMUSG00000070348 | 0,957 | 0,000 | Ccnd1 |
| ENSMUSG00000030711 | 0,899 | 0,016 | Sult1a1 | ENSMUSG00000029605 | 0,955 | 0,005 | Oas1b |
| ENSMUSG00000020798 | 0,897 | 0,000 | Spns3 | ENSMUSG00000022504 | 0,955 | 0,026 | Ciita |
| ENSMUSG00000102737 | 0,897 | 0,009 | 1700016A09Rik | ENSMUSG00000098850 | 0,954 | 0,003 | Gm27390 |
| ENSMUSG00000044576 | 0,895 | 0,000 | Garem2 | ENSMUSG00000022951 | 0,954 | 0,000 | Rcan1 |
| ENSMUSG00000051065 | 0,895 | 0,009 | Mb21d2 | ENSMUSG00000032860 | 0,950 | 0,008 | P2ry2 |
| ENSMUSG00000105338 | 0,893 | 0,021 | Gm43802 | ENSMUSG00000000440 | 0,949 | 0,000 | Pparg |
| ENSMUSG00000040605 | 0,893 | 0,002 | Bace2 | ENSMUSG00000001588 | 0,943 | 0,015 | Acap1 |
| ENSMUSG00000022018 | 0,892 | 0,007 | Rgcc | ENSMUSG00000032470 | 0,939 | 0,000 | Mras |
| ENSMUSG00000027907 | 0,891 | 0,003 | S100a11 | ENSMUSG00000018543 | 0,936 | 0,021 | 1700001P01Rik |
| ENSMUSG00000022346 | 0,891 | 0,000 | Myc | ENSMUSG00000061024 | 0,936 | 0,000 | Rrs1 |
| ENSMUSG00000001707 | 0,890 | 0,000 | Eef1e1 | ENSMUSG00000025492 | 0,935 | 0,000 | Ifitm3 |
| ENSMUSG00000071637 | 0,890 | 0,001 | Cebpd | ENSMUSG00000036594 | 0,933 | 0,001 | H2-Aa |
| ENSMUSG00000020224 | 0,889 | 0,002 | Llph | ENSMUSG00000036390 | 0,933 | 0,000 | Gadd45a |
| ENSMUSG00000002897 | 0,888 | 0,000 | Il17ra | ENSMUSG00000115426 | 0,930 | 0,034 | AC129931.1 |
| ENSMUSG00000095687 | 0,886 | 0,030 | Rnaset2a | ENSMUSG00000039629 | 0,929 | 0,014 | Strip2 |
| ENSMUSG00000032656 | 0,886 | 0,016 | March3 | ENSMUSG00000022146 | 0,924 | 0,011 | Osmr |
| ENSMUSG00000110010 | 0,884 | 0,006 | Gm45629 | ENSMUSG00000020044 | 0,924 | 0,002 | Timp3 |
| ENSMUSG00000037904 | 0,883 | 0,000 | Ankrd9 | ENSMUSG00000005148 | 0,924 | 0,005 | Klf5 |
| ENSMUSG00000098430 | 0,882 | 0,023 | Gm27494 | ENSMUSG00000032643 | 0,923 | 0,000 | Fhl3 |
| ENSMUSG00000008734 | 0,882 | 0,000 | Gprc5b | ENSMUSG00000102037 | 0,922 | 0,036 | Bcl2a1a |
| ENSMUSG00000022026 | 0,882 | 0,014 | Olfm4 | ENSMUSG00000058761 | 0,922 | 0,000 | Rnf169 |
| ENSMUSG00000047657 | 0,879 | 0,047 | Crry-ps | ENSMUSG00000060470 | 0,920 | 0,015 | Adgrg3 |
| ENSMUSG00000032014 | 0,877 | 0,000 | Oaf | ENSMUSG00000026020 | 0,920 | 0,000 | Nop58 |
| ENSMUSG00000045179 | 0,875 | 0,025 | Sox3 | ENSMUSG00000089774 | 0,917 | 0,000 | Slc5a3 |
| ENSMUSG00000022504 | 0,875 | 0,019 | Ciita | ENSMUSG00000097467 | 0,914 | 0,017 | Gm26737 |
| ENSMUSG00000034203 | 0,874 | 0,000 | Chchd4 | ENSMUSG00000001542 | 0,909 | 0,002 | Ell2 |
| ENSMUSG00000057561 | 0,874 | 0,000 | Eif1a | ENSMUSG00000114025 | 0,909 | 0,032 | AC154636.3 |
| ENSMUSG00000022528 | 0,873 | 0,000 | Hes1 | ENSMUSG00000059493 | 0,907 | 0,048 | Nhs |
| ENSMUSG00000045709 | 0,872 | 0,007 | Smkr-ps | ENSMUSG00000053414 | 0,906 | 0,007 | Hunk |
| ENSMUSG00000056737 | 0,871 | 0,002 | Capg | ENSMUSG00000039116 | 0,904 | 0,019 | Adgrg6 |
| ENSMUSG00000060143 | 0,870 | 0,008 | Gm10076 | ENSMUSG00000020363 | 0,901 | 0,035 | Gfpt2 |
| ENSMUSG00000038888 | 0,870 | 0,000 | Ctu1 | ENSMUSG00000024486 | 0,901 | 0,000 | Hbegf |
| ENSMUSG00000028771 | 0,869 | 0,000 | Ptpn12 | ENSMUSG00000021203 | 0,900 | 0,000 | Otub2 |
| ENSMUSG00000026020 | 0,863 | 0,000 | Nop58 | ENSMUSG00000027868 | 0,899 | 0,021 | Tbx15 |

| | | | | | |
|---|---|---|---|---|---|
| ENSMUSG00000032369 | 0,863 | 0,001 Plscr1 | ENSMUSG00000030790 | 0,897 | 0,048 Adm |
| ENSMUSG00000060568 | 0,863 | 0,009 Fam78b | ENSMUSG00000105161 | 0,895 | 0,007 Gm42595 |
| ENSMUSG00000028268 | 0,861 | 0,000 Gbp3 | ENSMUSG00000110404 | 0,894 | 0,041 Rnf223 |
| ENSMUSG00000056501 | 0,858 | 0,009 Cebpb | ENSMUSG00000020656 | 0,894 | 0,000 Grhl1 |
| ENSMUSG00000017929 | 0,854 | 0,000 B4galt5 | ENSMUSG00000024871 | 0,892 | 0,022 Doc2g |
| ENSMUSG00000040253 | 0,852 | 0,014 Gbp7 | ENSMUSG00000020812 | 0,891 | 0,000 1810032O08Rik |
| ENSMUSG00000026377 | 0,850 | 0,000 Nifk | ENSMUSG00000028063 | 0,891 | 0,000 Lmna |
| ENSMUSG00000023153 | 0,849 | 0,021 Tmem52 | ENSMUSG00000026435 | 0,886 | 0,013 Slc45a3 |
| ENSMUSG00000088088 | 0,848 | 0,008 Rmrp | ENSMUSG00000025591 | 0,882 | 0,000 Tma16 |
| ENSMUSG00000115420 | 0,848 | 0,008 AL732506.1 | ENSMUSG00000058794 | 0,880 | 0,004 Nfe2 |
| ENSMUSG00000035692 | 0,845 | 0,002 Isg15 | ENSMUSG00000027395 | 0,880 | 0,000 Polr1b |
| ENSMUSG00000046229 | 0,844 | 0,017 Scand1 | ENSMUSG00000049241 | 0,878 | 0,002 Hcar1 |
| ENSMUSG00000074800 | 0,843 | 0,028 Gm4149 | ENSMUSG00000040957 | 0,878 | 0,000 Cables1 |
| ENSMUSG00000057605 | 0,842 | 0,023 Gm6807 | ENSMUSG00000069805 | 0,877 | 0,041 Fbp1 |
| ENSMUSG00000097327 | 0,841 | 0,001 E030030I06Rik | ENSMUSG00000023153 | 0,877 | 0,023 Tmem52 |
| ENSMUSG00000022900 | 0,840 | 0,003 Ildr1 | ENSMUSG00000036452 | 0,876 | 0,000 Arhgap26 |
| ENSMUSG00000039236 | 0,839 | 0,008 Isg20 | ENSMUSG00000055865 | 0,875 | 0,036 Fam19a3 |
| ENSMUSG00000087590 | 0,838 | 0,013 Epb41l4aos | ENSMUSG00000024053 | 0,873 | 0,010 Emilin2 |
| ENSMUSG00000062353 | 0,834 | 0,020 Gm15772 | ENSMUSG00000044033 | 0,873 | 0,020 Ccdc141 |
| ENSMUSG00000038046 | 0,833 | 0,000 Mrm3 | ENSMUSG00000038335 | 0,871 | 0,000 Tsr1 |
| ENSMUSG00000063354 | 0,833 | 0,002 Slc39a4 | ENSMUSG00000025856 | 0,869 | 0,000 Pdgfa |
| ENSMUSG00000049103 | 0,830 | 0,010 Ccr2 | ENSMUSG00000045180 | 0,867 | 0,006 Shroom2 |
| ENSMUSG00000032322 | 0,830 | 0,045 Pstpip1 | ENSMUSG00000024085 | 0,864 | 0,001 Man2a1 |
| ENSMUSG00000067870 | 0,830 | 0,019 Rpl31-ps8 | ENSMUSG00000024769 | 0,862 | 0,000 Cdc42bpg |
| ENSMUSG00000021958 | 0,829 | 0,000 Pinx1 | ENSMUSG00000021149 | 0,861 | 0,000 Gtpbp4 |
| ENSMUSG00000028381 | 0,828 | 0,000 Ugcg | ENSMUSG00000027907 | 0,859 | 0,007 S100a11 |
| ENSMUSG00000030170 | 0,826 | 0,024 Wnt5b | ENSMUSG00000026875 | 0,857 | 0,006 Traf1 |
| ENSMUSG00000024843 | 0,826 | 0,000 Chka | ENSMUSG00000065124 | 0,853 | 0,001 Snora65 |
| ENSMUSG00000028413 | 0,825 | 0,000 B4galt1 | ENSMUSG00000040328 | 0,848 | 0,026 Olfr56 |
| ENSMUSG00000031605 | 0,825 | 0,000 Klhl2 | ENSMUSG00000031700 | 0,845 | 0,000 Gpt2 |
| ENSMUSG00000009378 | 0,824 | 0,010 Slc16a12 | ENSMUSG00000020961 | 0,843 | 0,004 Ston2 |
| ENSMUSG00000044583 | 0,824 | 0,038 Tlr7 | ENSMUSG00000042608 | 0,841 | 0,000 Stk40 |
| ENSMUSG00000020599 | 0,824 | 0,006 Rgs9 | ENSMUSG00000060586 | 0,841 | 0,004 H2-Eb1 |
| ENSMUSG00000065922 | 0,823 | 0,008 n-R5-8s1 | ENSMUSG00000026833 | 0,841 | 0,006 Olfm1 |
| ENSMUSG00000004100 | 0,823 | 0,000 Ppan | ENSMUSG00000114635 | 0,838 | 0,012 CT010498.2 |
| ENSMUSG00000066975 | 0,817 | 0,002 Cryba4 | ENSMUSG00000056749 | 0,837 | 0,000 Nfil3 |
| ENSMUSG00000114487 | 0,812 | 0,049 CT010498.1 | ENSMUSG00000106847 | 0,834 | 0,000 Peg13 |
| ENSMUSG00000023988 | 0,812 | 0,000 Bysl | ENSMUSG00000080058 | 0,825 | 0,012 Gm11175 |
| ENSMUSG00000071757 | 0,812 | 0,000 Zhx2 | ENSMUSG00000037434 | 0,825 | 0,000 Slc30a1 |
| ENSMUSG00000015176 | 0,809 | 0,000 Nolc1 | ENSMUSG00000111291 | 0,824 | 0,005 AC124577.2 |
| ENSMUSG00000113165 | 0,809 | 0,001 AC115037.1 | ENSMUSG00000042349 | 0,824 | 0,016 Ikbke |
| ENSMUSG00000048216 | 0,809 | 0,014 Gpr85 | ENSMUSG00000086429 | 0,822 | 0,001 Gt(ROSA)26Sor |
| ENSMUSG00000030609 | 0,808 | 0,000 Aen | ENSMUSG00000032656 | 0,817 | 0,041 March3 |
| ENSMUSG00000033538 | 0,807 | 0,015 Casp4 | ENSMUSG00000091996 | 0,817 | 0,008 BC049352 |
| ENSMUSG00000000184 | 0,805 | 0,001 Ccnd2 | ENSMUSG00000046204 | 0,813 | 0,006 Pnma2 |
| ENSMUSG00000061815 | 0,803 | 0,031 Rufy4 | ENSMUSG00000109015 | 0,811 | 0,007 Gm31024 |
| ENSMUSG00000032193 | 0,801 | 0,003 Ldlr | ENSMUSG00000070780 | 0,808 | 0,000 Rbm47 |

| | | | | | |
|---|---|---|---|---|---|
| ENSMUSG00000109536 | 0,801 | 0,043 9330162G02Rik | ENSMUSG00000094626 | 0,808 | 0,032 Tmem121b |
| ENSMUSG00000030521 | 0,801 | 0,000 Mphosph10 | ENSMUSG00000028771 | 0,806 | 0,000 Ptpn12 |
| ENSMUSG00000001588 | 0,799 | 0,030 Acap1 | ENSMUSG00000037254 | 0,803 | 0,045 Itih2 |
| ENSMUSG00000017652 | 0,799 | 0,012 Cd40 | ENSMUSG00000093327 | 0,802 | 0,003 Mir5107 |
| ENSMUSG00000008540 | 0,797 | 0,000 Mgst1 | ENSMUSG00000073421 | 0,801 | 0,008 H2-Ab1 |
| ENSMUSG00000021265 | 0,796 | 0,001 Slc25a29 | ENSMUSG00000058883 | 0,801 | 0,043 Zfp708 |
| ENSMUSG00000073599 | 0,796 | 0,005 Ecscr | ENSMUSG00000028874 | 0,800 | 0,015 Fgr |
| ENSMUSG00000050299 | 0,796 | 0,011 Gm9843 | ENSMUSG00000026576 | 0,799 | 0,009 Atp1b1 |
| ENSMUSG00000020205 | 0,795 | 0,000 Phlda1 | ENSMUSG00000052040 | 0,797 | 0,000 Klf13 |
| ENSMUSG00000019194 | 0,795 | 0,000 Scn1b | ENSMUSG00000110945 | 0,796 | 0,022 Gm9856 |
| ENSMUSG00000038279 | 0,795 | 0,000 Nop2 | ENSMUSG00000068417 | 0,794 | 0,000 Pnp2 |
| ENSMUSG00000075602 | 0,794 | 0,041 Ly6a | ENSMUSG00000020599 | 0,792 | 0,014 Rgs9 |
| ENSMUSG00000002983 | 0,794 | 0,000 Relb | ENSMUSG00000022371 | 0,792 | 0,005 Col14a1 |
| ENSMUSG00000000776 | 0,792 | 0,000 Polr3d | ENSMUSG00000074876 | 0,787 | 0,000 Spata5l1 |
| ENSMUSG00000021285 | 0,791 | 0,000 Ppp1r13b | ENSMUSG00000097327 | 0,781 | 0,002 E030030I06Rik |
| ENSMUSG00000039699 | 0,791 | 0,010 Batf2 | ENSMUSG00000076441 | 0,779 | 0,020 Ass1 |
| ENSMUSG00000032561 | 0,790 | 0,018 Acpp | ENSMUSG00000022661 | 0,776 | 0,016 Cd200 |
| ENSMUSG00000063193 | 0,786 | 0,011 Cd300lb | ENSMUSG00000113165 | 0,775 | 0,004 AC115037.1 |
| ENSMUSG00000024486 | 0,784 | 0,001 Hbegf | ENSMUSG00000110275 | 0,772 | 0,003 Gm5905 |
| ENSMUSG00000084416 | 0,782 | 0,019 Rpl10a-ps1 | ENSMUSG00000078490 | 0,770 | 0,001 Cfap74 |
| ENSMUSG00000090439 | 0,780 | 0,040 Gm17455 | ENSMUSG00000030203 | 0,768 | 0,000 Dusp16 |
| ENSMUSG00000041057 | 0,780 | 0,000 Wdr43 | ENSMUSG00000036381 | 0,767 | 0,036 P2ry14 |
| ENSMUSG00000038296 | 0,778 | 0,000 Galnt18 | ENSMUSG00000039782 | 0,766 | 0,023 Cpeb2 |
| ENSMUSG00000028063 | 0,775 | 0,001 Lmna | ENSMUSG00000097743 | 0,765 | 0,001 Gm16973 |
| ENSMUSG00000031578 | 0,773 | 0,000 Mak16 | ENSMUSG00000048807 | 0,765 | 0,000 Slc35e4 |
| ENSMUSG00000024087 | 0,772 | 0,018 Cyp1b1 | ENSMUSG00000050860 | 0,758 | 0,001 Phospho1 |
| ENSMUSG00000090069 | 0,772 | 0,038 E430024P14Rik | ENSMUSG00000015843 | 0,755 | 0,020 Rxrg |
| ENSMUSG00000002233 | 0,770 | 0,000 Rhoc | ENSMUSG00000027508 | 0,754 | 0,018 Pag1 |
| ENSMUSG00000026728 | 0,769 | 0,000 Vim | ENSMUSG00000039153 | 0,752 | 0,046 Runx2 |
| ENSMUSG00000024754 | 0,769 | 0,008 Tmem2 | ENSMUSG00000085156 | 0,750 | 0,000 Snhg15 |
| ENSMUSG00000018012 | 0,768 | 0,000 Rac3 | ENSMUSG00000008734 | 0,749 | 0,000 Gprc5b |
| ENSMUSG00000003153 | 0,768 | 0,000 Slc2a3 | ENSMUSG00000046791 | 0,748 | 0,000 Riox1 |
| ENSMUSG00000078974 | 0,767 | 0,008 Sec61g | ENSMUSG00000015176 | 0,746 | 0,000 Nolc1 |
| ENSMUSG00000021453 | 0,767 | 0,003 Gadd45g | ENSMUSG00000054263 | 0,746 | 0,018 Lifr |
| ENSMUSG00000028840 | 0,766 | 0,000 Zfp593 | ENSMUSG00000035673 | 0,745 | 0,001 Sbno2 |
| ENSMUSG00000035673 | 0,766 | 0,000 Sbno2 | ENSMUSG00000063193 | 0,744 | 0,028 Cd300lb |
| ENSMUSG00000049225 | 0,764 | 0,000 Pdp1 | ENSMUSG00000042821 | 0,741 | 0,003 Snai1 |
| ENSMUSG00000040093 | 0,764 | 0,003 Bmf | ENSMUSG00000027765 | 0,739 | 0,031 P2ry1 |
| ENSMUSG00000042417 | 0,763 | 0,047 Ccno | ENSMUSG00000049999 | 0,738 | 0,000 Ppp1r3d |
| ENSMUSG00000068099 | 0,763 | 0,002 1500009C09Rik | ENSMUSG00000004100 | 0,737 | 0,000 Ppan |
| ENSMUSG00000006360 | 0,763 | 0,042 Crip1 | ENSMUSG00000040451 | 0,736 | 0,003 Sgms1 |
| ENSMUSG00000026796 | 0,763 | 0,000 Fam129b | ENSMUSG00000099109 | 0,733 | 0,046 Mir7115 |
| ENSMUSG00000040447 | 0,763 | 0,003 Spns2 | ENSMUSG00000044576 | 0,733 | 0,003 Garem2 |
| ENSMUSG00000074466 | 0,763 | 0,015 Gm15417 | ENSMUSG00000052776 | 0,731 | 0,005 Oas1a |
| ENSMUSG00000104367 | 0,761 | 0,018 Gm38079 | ENSMUSG00000092035 | 0,730 | 0,007 Peg10 |
| ENSMUSG00000031837 | 0,761 | 0,019 Necab2 | ENSMUSG00000087365 | 0,730 | 0,022 C430049B03Rik |
| ENSMUSG00000000957 | 0,760 | 0,000 Mmp14 | ENSMUSG00000071637 | 0,728 | 0,013 Cebpd |

| | | | | | | | |
|---|---|---|---|---|---|---|---|
| ENSMUSG00000044197 | 0,758 | 0,000 | Gpr146 | ENSMUSG00000057561 | 0,728 | 0,000 | Eif1a |
| ENSMUSG00000069899 | 0,757 | 0,018 | Gm12166 | ENSMUSG00000028967 | 0,727 | 0,000 | Errfi1 |
| ENSMUSG00000015476 | 0,755 | 0,039 | Prrt1 | ENSMUSG00000002881 | 0,726 | 0,000 | Nab1 |
| ENSMUSG00000022146 | 0,752 | 0,034 | Osmr | ENSMUSG00000044345 | 0,725 | 0,001 | Marveld1 |
| ENSMUSG00000079508 | 0,752 | 0,017 | Apoo | ENSMUSG00000079508 | 0,722 | 0,039 | Apoo |
| ENSMUSG00000054934 | 0,750 | 0,045 | Kcnmb4 | ENSMUSG00000005237 | 0,722 | 0,014 | Dnah2 |
| ENSMUSG00000027236 | 0,749 | 0,000 | Eif3j1 | ENSMUSG00000085385 | 0,720 | 0,000 | Snhg17 |
| ENSMUSG00000002881 | 0,748 | 0,000 | Nab1 | ENSMUSG00000028381 | 0,719 | 0,000 | Ugcg |
| ENSMUSG00000031490 | 0,748 | 0,005 | Eif4ebp1 | ENSMUSG00000049225 | 0,718 | 0,000 | Pdp1 |
| ENSMUSG00000096974 | 0,748 | 0,028 | Gm26881 | ENSMUSG00000021453 | 0,718 | 0,009 | Gadd45g |
| ENSMUSG00000039450 | 0,747 | 0,000 | Dcxr | ENSMUSG00000033721 | 0,718 | 0,002 | Vav3 |
| ENSMUSG00000030203 | 0,747 | 0,000 | Dusp16 | ENSMUSG00000038264 | 0,717 | 0,035 | Sema7a |
| ENSMUSG00000011008 | 0,746 | 0,047 | Mcoln2 | ENSMUSG00000079445 | 0,716 | 0,004 | B3gnt7 |
| ENSMUSG00000035295 | 0,746 | 0,037 | Wdr38 | ENSMUSG00000046794 | 0,713 | 0,004 | Ppp1r3b |
| ENSMUSG00000097906 | 0,745 | 0,000 | Gm9625 | ENSMUSG00000024754 | 0,711 | 0,023 | Tmem2 |
| ENSMUSG00000085355 | 0,745 | 0,029 | 3010003L21Rik | ENSMUSG00000038279 | 0,710 | 0,000 | Nop2 |
| ENSMUSG00000017715 | 0,743 | 0,000 | Pgs1 | ENSMUSG00000046879 | 0,708 | 0,000 | Irgm1 |
| ENSMUSG00000027076 | 0,743 | 0,000 | Timm10 | ENSMUSG00000067931 | 0,707 | 0,012 | Zfp948 |
| ENSMUSG00000033777 | 0,743 | 0,001 | Tlr13 | ENSMUSG00000033355 | 0,707 | 0,000 | Rtp4 |
| ENSMUSG00000045538 | 0,742 | 0,000 | Ddx28 | ENSMUSG00000087412 | 0,706 | 0,043 | Gm15501 |
| ENSMUSG00000050860 | 0,741 | 0,001 | Phospho1 | ENSMUSG00000024732 | 0,703 | 0,000 | Ccdc86 |
| ENSMUSG00000001630 | 0,740 | 0,000 | Stk38l | ENSMUSG00000038046 | 0,703 | 0,001 | Mrm3 |
| ENSMUSG00000051435 | 0,738 | 0,000 | Fhad1 | ENSMUSG00000000776 | 0,703 | 0,000 | Polr3d |
| ENSMUSG00000036390 | 0,736 | 0,000 | Gadd45a | ENSMUSG00000040093 | 0,701 | 0,011 | Bmf |
| ENSMUSG00000037966 | 0,734 | 0,000 | Ninj1 | ENSMUSG00000030521 | 0,699 | 0,000 | Mphosph10 |
| ENSMUSG00000027508 | 0,733 | 0,016 | Pag1 | ENSMUSG00000022305 | 0,697 | 0,007 | Lrp12 |
| ENSMUSG00000028010 | 0,733 | 0,000 | Gar1 | ENSMUSG00000025357 | 0,697 | 0,000 | Dgka |
| ENSMUSG00000047090 | 0,732 | 0,000 | Tmem198b | ENSMUSG00000002617 | 0,695 | 0,003 | Zfp40 |
| ENSMUSG00000038489 | 0,730 | 0,002 | Polr2l | ENSMUSG00000008540 | 0,695 | 0,001 | Mgst1 |
| ENSMUSG00000029570 | 0,730 | 0,000 | Lfng | ENSMUSG00000038342 | 0,693 | 0,002 | Mlxip |
| ENSMUSG00000040957 | 0,729 | 0,000 | Cables1 | ENSMUSG00000024042 | 0,692 | 0,014 | Sik1 |
| ENSMUSG00000091509 | 0,728 | 0,002 | Gm17066 | ENSMUSG00000034570 | 0,691 | 0,018 | Inpp5j |
| ENSMUSG00000085436 | 0,728 | 0,010 | Zfp335os | ENSMUSG00000085492 | 0,687 | 0,000 | Trmt61b |
| ENSMUSG00000031304 | 0,727 | 0,019 | Il2rg | ENSMUSG00000022957 | 0,687 | 0,000 | Itsn1 |
| ENSMUSG00000060032 | 0,727 | 0,016 | H2afj | ENSMUSG00000085316 | 0,686 | 0,039 | D330050G23Rik |
| ENSMUSG00000040451 | 0,727 | 0,002 | Sgms1 | ENSMUSG00000024985 | 0,686 | 0,000 | Tcf7l2 |
| ENSMUSG00000041506 | 0,727 | 0,000 | Rrp9 | ENSMUSG00000027375 | 0,685 | 0,027 | Mal |
| ENSMUSG00000053414 | 0,724 | 0,029 | Hunk | ENSMUSG00000003153 | 0,682 | 0,001 | Slc2a3 |
| ENSMUSG00000020656 | 0,724 | 0,000 | Grhl1 | ENSMUSG00000040829 | 0,681 | 0,038 | Zmynd15 |
| ENSMUSG00000028108 | 0,724 | 0,001 | Ecm1 | ENSMUSG00000071757 | 0,680 | 0,000 | Zhx2 |
| ENSMUSG00000016664 | 0,723 | 0,000 | Pacsin2 | ENSMUSG00000047735 | 0,679 | 0,017 | Samd9l |
| ENSMUSG00000018920 | 0,720 | 0,013 | Cxcl16 | ENSMUSG00000030609 | 0,679 | 0,000 | Aen |
| ENSMUSG00000052253 | 0,718 | 0,000 | Zfp622 | ENSMUSG00000046223 | 0,678 | 0,005 | Plaur |
| ENSMUSG00000027596 | 0,716 | 0,010 | a | ENSMUSG00000024778 | 0,678 | 0,020 | Fas |
| ENSMUSG00000026875 | 0,716 | 0,016 | Traf1 | ENSMUSG00000031578 | 0,678 | 0,000 | Mak16 |
| ENSMUSG00000050212 | 0,716 | 0,002 | Eva1b | ENSMUSG00000045672 | 0,675 | 0,000 | Col27a1 |
| ENSMUSG00000027221 | 0,715 | 0,000 | Chst1 | ENSMUSG00000093677 | 0,675 | 0,033 | Gm20712 |

| | | | | | | | |
|---|---|---|---|---|---|---|---|
| ENSMUSG00000064307 | 0,715 | 0,045 | Lrrc51 | ENSMUSG00000032119 | 0,674 | 0,000 | Hinfp |
| ENSMUSG00000034614 | 0,714 | 0,006 | Pik3ip1 | ENSMUSG00000026981 | 0,673 | 0,044 | Il1rn |
| ENSMUSG00000022372 | 0,713 | 0,012 | Sla | ENSMUSG00000028069 | 0,673 | 0,001 | Gpatch4 |
| ENSMUSG00000068417 | 0,713 | 0,000 | Pnp2 | ENSMUSG00000010358 | 0,672 | 0,004 | Ifi35 |
| ENSMUSG00000048007 | 0,710 | 0,007 | Timm8a1 | ENSMUSG00000031825 | 0,672 | 0,011 | Crispld2 |
| ENSMUSG00000031839 | 0,710 | 0,000 | Hsbp1 | ENSMUSG00000030409 | 0,671 | 0,038 | Dmpk |
| ENSMUSG00000042854 | 0,709 | 0,000 | Trp53rkb | ENSMUSG00000031880 | 0,670 | 0,025 | Rrad |
| ENSMUSG00000025290 | 0,709 | 0,012 | Rps24 | ENSMUSG00000025887 | 0,670 | 0,025 | Casp12 |
| ENSMUSG00000025402 | 0,709 | 0,000 | Nab2 | ENSMUSG00000031605 | 0,669 | 0,000 | Klhl2 |
| ENSMUSG00000025171 | 0,708 | 0,000 | Ubtd1 | ENSMUSG00000056888 | 0,668 | 0,000 | Glipr1 |
| ENSMUSG00000020121 | 0,704 | 0,011 | Srgap1 | ENSMUSG00000037904 | 0,667 | 0,027 | Ankrd9 |
| ENSMUSG00000039084 | 0,703 | 0,031 | Chad | ENSMUSG00000002058 | 0,667 | 0,006 | Unc119 |
| ENSMUSG00000022371 | 0,703 | 0,010 | Col14a1 | ENSMUSG00000050335 | 0,666 | 0,003 | Lgals3 |
| ENSMUSG00000000732 | 0,702 | 0,000 | Icosl | ENSMUSG00000049950 | 0,663 | 0,004 | Rpp38 |
| ENSMUSG00000021687 | 0,700 | 0,000 | Scamp1 | ENSMUSG00000097772 | 0,662 | 0,001 | 5430416N02Rik |
| ENSMUSG00000028948 | 0,700 | 0,000 | Nol9 | ENSMUSG00000053063 | 0,661 | 0,035 | Clec12a |
| ENSMUSG00000030403 | 0,699 | 0,000 | Vasp | ENSMUSG00000028128 | 0,657 | 0,046 | F3 |
| ENSMUSG00000015337 | 0,699 | 0,006 | Endog | ENSMUSG00000037692 | 0,657 | 0,000 | Ahdc1 |
| ENSMUSG00000042745 | 0,698 | 0,011 | Id1 | ENSMUSG00000054855 | 0,656 | 0,005 | Rnd1 |
| ENSMUSG00000001020 | 0,697 | 0,032 | S100a4 | ENSMUSG00000041057 | 0,652 | 0,000 | Wdr43 |
| ENSMUSG00000036278 | 0,696 | 0,003 | Macrod1 | ENSMUSG00000038888 | 0,652 | 0,000 | Ctu1 |
| ENSMUSG00000083621 | 0,694 | 0,045 | Gm14586 | ENSMUSG00000021958 | 0,651 | 0,000 | Pinx1 |
| ENSMUSG00000042729 | 0,694 | 0,000 | Wdr74 | ENSMUSG00000020798 | 0,651 | 0,022 | Spns3 |
| ENSMUSG00000035397 | 0,694 | 0,000 | Klf16 | ENSMUSG00000101628 | 0,650 | 0,040 | Gm28177 |
| ENSMUSG00000025869 | 0,694 | 0,000 | Nop16 | ENSMUSG00000016552 | 0,650 | 0,045 | Foxred2 |
| ENSMUSG00000000409 | 0,692 | 0,027 | Lck | ENSMUSG00000036948 | 0,649 | 0,000 | BC037034 |
| ENSMUSG00000097772 | 0,692 | 0,000 | 5430416N02Rik | ENSMUSG00000022351 | 0,648 | 0,007 | Sqle |
| ENSMUSG00000020484 | 0,692 | 0,000 | Xbp1 | ENSMUSG00000003617 | 0,648 | 0,027 | Cp |
| ENSMUSG00000002058 | 0,692 | 0,002 | Unc119 | ENSMUSG00000043079 | 0,648 | 0,010 | Synpo |
| ENSMUSG00000023272 | 0,690 | 0,000 | Creld2 | ENSMUSG00000031146 | 0,645 | 0,014 | Plp2 |
| ENSMUSG00000021493 | 0,690 | 0,012 | Pdlim7 | ENSMUSG00000021493 | 0,644 | 0,033 | Pdlim7 |
| ENSMUSG00000110423 | 0,689 | 0,000 | Gm45736 | ENSMUSG00000089762 | 0,641 | 0,022 | Ier5l |
| ENSMUSG00000039725 | 0,688 | 0,000 | Trp53rka | ENSMUSG00000036533 | 0,641 | 0,012 | Cdc42ep3 |
| ENSMUSG00000022864 | 0,688 | 0,003 | D16Ertd472e | ENSMUSG00000026377 | 0,639 | 0,000 | Nifk |
| ENSMUSG00000021508 | 0,688 | 0,008 | Cxcl14 | ENSMUSG00000037260 | 0,639 | 0,000 | Hgsnat |
| ENSMUSG00000063694 | 0,687 | 0,000 | Cycs | ENSMUSG00000000732 | 0,637 | 0,001 | Icosl |
| ENSMUSG00000045690 | 0,686 | 0,003 | Wdr89 | ENSMUSG00000043421 | 0,637 | 0,000 | Hilpda |
| ENSMUSG00000033721 | 0,685 | 0,003 | Vav3 | ENSMUSG00000001707 | 0,636 | 0,000 | Eef1e1 |
| ENSMUSG00000024608 | 0,684 | 0,007 | Rps14 | ENSMUSG00000019846 | 0,635 | 0,003 | Lama4 |
| ENSMUSG00000039985 | 0,684 | 0,009 | Fam60a | ENSMUSG00000021285 | 0,633 | 0,000 | Ppp1r13b |
| ENSMUSG00000085492 | 0,683 | 0,000 | Trmt61b | ENSMUSG00000028413 | 0,633 | 0,000 | B4galt1 |
| ENSMUSG00000017765 | 0,683 | 0,000 | Slc12a4 | ENSMUSG00000067653 | 0,631 | 0,001 | Ankrd23 |
| ENSMUSG00000027215 | 0,683 | 0,000 | Cd82 | ENSMUSG00000014778 | 0,629 | 0,000 | Fhod1 |
| ENSMUSG00000022247 | 0,682 | 0,000 | Brix1 | ENSMUSG00000053801 | 0,629 | 0,000 | Grwd1 |
| ENSMUSG00000020812 | 0,681 | 0,002 | 1810032O08Rik | ENSMUSG00000051435 | 0,628 | 0,001 | Fhad1 |
| ENSMUSG00000046410 | 0,681 | 0,017 | Kcnk6 | ENSMUSG00000034818 | 0,627 | 0,045 | Celf5 |
| ENSMUSG00000026107 | 0,678 | 0,042 | Nabp1 | ENSMUSG00000097392 | 0,627 | 0,000 | D930016D06Rik |

| | | | | | |
|---|---|---|---|---|---|
| ENSMUSG00000013846 | 0,677 | 0,020 St3gal1 | ENSMUSG00000002983 | 0,626 | 0,001 Relb |
| ENSMUSG00000032841 | 0,676 | 0,004 Prr5l | ENSMUSG00000025650 | 0,625 | 0,010 Col7a1 |
| ENSMUSG00000003070 | 0,676 | 0,036 Efna2 | ENSMUSG00000041438 | 0,624 | 0,000 Utp4 |
| ENSMUSG00000027333 | 0,676 | 0,000 Smox | ENSMUSG00000001630 | 0,623 | 0,002 Stk38l |
| ENSMUSG00000020399 | 0,675 | 0,037 Havcr2 | ENSMUSG00000007617 | 0,623 | 0,004 Homer1 |
| ENSMUSG00000115338 | 0,675 | 0,000 Pnp | ENSMUSG00000026104 | 0,622 | 0,000 Stat1 |
| ENSMUSG00000023068 | 0,674 | 0,000 Nus1 | ENSMUSG00000091509 | 0,622 | 0,015 Gm17066 |
| ENSMUSG00000049744 | 0,673 | 0,034 Arhgap15 | ENSMUSG00000026970 | 0,620 | 0,000 Rbms1 |
| ENSMUSG00000018339 | 0,673 | 0,000 Gpx3 | ENSMUSG00000028717 | 0,619 | 0,008 Tal1 |
| ENSMUSG00000003283 | 0,672 | 0,004 Hck | ENSMUSG00000069049 | 0,617 | 0,005 Eif2s3y |
| ENSMUSG00000021871 | 0,671 | 0,000 Pnp | ENSMUSG00000032802 | 0,617 | 0,000 Srxn1 |
| ENSMUSG00000038510 | 0,670 | 0,000 Rpf2 | ENSMUSG00000051495 | 0,617 | 0,000 Irf2bp2 |
| ENSMUSG00000025436 | 0,670 | 0,000 Atp23 | ENSMUSG00000114771 | 0,616 | 0,043 AC163667.1 |
| ENSMUSG00000026069 | 0,667 | 0,002 Il1rl1 | ENSMUSG00000038296 | 0,616 | 0,006 Galnt18 |
| ENSMUSG00000026576 | 0,667 | 0,023 Atp1b1 | ENSMUSG00000074622 | 0,616 | 0,000 Mafb |
| ENSMUSG00000074876 | 0,666 | 0,000 Spata5l1 | ENSMUSG00000017715 | 0,614 | 0,000 Pgs1 |
| ENSMUSG00000031431 | 0,666 | 0,001 Tsc22d3 | ENSMUSG00000073775 | 0,613 | 0,000 Kti12 |
| ENSMUSG00000041390 | 0,664 | 0,003 Mdfic | ENSMUSG00000000184 | 0,613 | 0,028 Ccnd2 |
| ENSMUSG00000022554 | 0,663 | 0,000 Hgh1 | ENSMUSG00000022960 | 0,611 | 0,000 Donson |
| ENSMUSG00000098178 | 0,662 | 0,042 Gm42418 | ENSMUSG00000027583 | 0,610 | 0,000 Zbtb46 |
| ENSMUSG00000038274 | 0,662 | 0,015 Fau | ENSMUSG00000021365 | 0,608 | 0,003 Nedd9 |
| ENSMUSG00000064109 | 0,660 | 0,035 Hcst | ENSMUSG00000040447 | 0,607 | 0,036 Spns2 |
| ENSMUSG00000033294 | 0,659 | 0,000 Noc4l | ENSMUSG00000028840 | 0,606 | 0,005 Zfp593 |
| ENSMUSG00000023905 | 0,659 | 0,011 Tnfrsf12a | ENSMUSG00000097906 | 0,606 | 0,000 Gm9625 |
| ENSMUSG00000026305 | 0,659 | 0,000 Lrrfip1 | ENSMUSG00000046667 | 0,606 | 0,001 Rbm12b1 |
| ENSMUSG00000043439 | 0,659 | 0,002 Epop | ENSMUSG00000040033 | 0,606 | 0,003 Stat2 |
| ENSMUSG00000070284 | 0,659 | 0,000 Gmppb | ENSMUSG00000028948 | 0,604 | 0,000 Nol9 |
| ENSMUSG00000044345 | 0,658 | 0,001 Marveld1 | ENSMUSG00000034579 | 0,601 | 0,038 Pla2g3 |
| ENSMUSG00000032109 | 0,657 | 0,000 Nlrx1 | ENSMUSG00000104950 | 0,600 | 0,004 4833413G10Rik |
| ENSMUSG00000058794 | 0,657 | 0,031 Nfe2 | ENSMUSG00000063972 | 0,599 | 0,010 Nr6a1 |
| ENSMUSG00000038845 | 0,657 | 0,004 Phb | ENSMUSG00000075585 | 0,599 | 0,001 6330403L08Rik |
| ENSMUSG00000055137 | 0,656 | 0,001 Sugct | ENSMUSG00000067212 | 0,598 | 0,000 H2-T23 |
| ENSMUSG00000024085 | 0,654 | 0,016 Man2a1 | ENSMUSG00000022864 | 0,597 | 0,017 D16Ertd472e |
| ENSMUSG00000054720 | 0,653 | 0,001 Lrrc8c | ENSMUSG00000026305 | 0,594 | 0,000 Lrrfip1 |
| ENSMUSG00000038264 | 0,653 | 0,044 Sema7a | ENSMUSG00000098465 | 0,594 | 0,033 Mir1668 |
| ENSMUSG00000020806 | 0,652 | 0,019 Rhbdf2 | ENSMUSG00000025813 | 0,593 | 0,036 Homer2 |
| ENSMUSG00000049482 | 0,651 | 0,000 Ctu2 | ENSMUSG00000115338 | 0,593 | 0,000 Pnp |
| ENSMUSG00000060216 | 0,651 | 0,000 Arrb2 | ENSMUSG00000009741 | 0,593 | 0,000 Ubp1 |
| ENSMUSG00000050244 | 0,651 | 0,000 Heatr1 | ENSMUSG00000017929 | 0,593 | 0,000 B4galt5 |
| ENSMUSG00000070462 | 0,650 | 0,000 Tlnrd1 | ENSMUSG00000019487 | 0,593 | 0,000 Trip10 |
| ENSMUSG00000027765 | 0,649 | 0,039 P2ry1 | ENSMUSG00000097000 | 0,592 | 0,016 Gm17435 |
| ENSMUSG00000089762 | 0,648 | 0,014 Ier5l | ENSMUSG00000098794 | 0,591 | 0,001 Gm28038 |
| ENSMUSG00000034686 | 0,648 | 0,034 Prr7 | ENSMUSG00000021871 | 0,590 | 0,000 Pnp |
| ENSMUSG00000032802 | 0,648 | 0,000 Srxn1 | ENSMUSG00000032841 | 0,589 | 0,024 Prr5l |
| ENSMUSG00000017679 | 0,648 | 0,000 Ttpal | ENSMUSG00000060950 | 0,588 | 0,000 Trmt61a |
| ENSMUSG00000015843 | 0,647 | 0,040 Rxrg | ENSMUSG00000039470 | 0,588 | 0,000 Zdhhc2 |
| ENSMUSG00000006442 | 0,646 | 0,000 Srm | ENSMUSG00000043183 | 0,587 | 0,001 Simc1 |

| | | | | | | | |
|---|---|---|---|---|---|---|---|
| ENSMUSG00000054871 | 0,645 | 0,022 | Tmem158 | ENSMUSG00000049939 | 0,587 | 0,003 | Lrrc4 |
| ENSMUSG00000029642 | 0,645 | 0,002 | Polr1d | ENSMUSG00000027236 | 0,585 | 0,000 | Eif3j1 |
| ENSMUSG00000087651 | 0,644 | 0,024 | 1500009L16Rik | ENSMUSG00000002885 | 0,585 | 0,007 | Adgre5 |
| ENSMUSG00000003206 | 0,644 | 0,006 | Ebi3 | ENSMUSG00000031024 | 0,585 | 0,000 | St5 |
| ENSMUSG00000036256 | 0,644 | 0,047 | Igfbp7 | ENSMUSG00000086150 | 0,585 | 0,037 | Bach2os |
| ENSMUSG00000041203 | 0,644 | 0,003 | Trir | ENSMUSG00000017679 | 0,583 | 0,000 | Ttpal |
| ENSMUSG00000037260 | 0,642 | 0,000 | Hgsnat | ENSMUSG00000026728 | 0,583 | 0,002 | Vim |
| ENSMUSG00000020075 | 0,641 | 0,000 | Ddx21 | ENSMUSG00000033909 | 0,582 | 0,000 | Usp36 |
| ENSMUSG00000112429 | 0,640 | 0,001 | AC131081.1 | ENSMUSG00000042364 | 0,581 | 0,000 | Snx18 |
| ENSMUSG00000015950 | 0,640 | 0,020 | Ncf1 | ENSMUSG00000044149 | 0,580 | 0,000 | Nkrf |
| ENSMUSG00000070476 | 0,638 | 0,000 | Fam217b | ENSMUSG00000033545 | 0,579 | 0,000 | Znrf1 |
| ENSMUSG00000050796 | 0,637 | 0,000 | B3galt6 | ENSMUSG00000043243 | 0,578 | 0,011 | Fam129c |
| ENSMUSG00000035049 | 0,637 | 0,000 | Rrp12 | ENSMUSG00000031939 | 0,578 | 0,002 | Taf1d |
| ENSMUSG00000022876 | 0,636 | 0,002 | Samsn1 | ENSMUSG00000039633 | 0,577 | 0,009 | Lonrf1 |
| ENSMUSG00000087305 | 0,636 | 0,007 | A430035B10Rik | ENSMUSG00000078651 | 0,576 | 0,005 | Aoc2 |
| ENSMUSG00000020044 | 0,634 | 0,038 | Timp3 | ENSMUSG00000044197 | 0,576 | 0,001 | Gpr146 |
| ENSMUSG00000031511 | 0,633 | 0,000 | Arhgef7 | ENSMUSG00000028261 | 0,575 | 0,000 | Ndufaf4 |
| ENSMUSG00000105161 | 0,633 | 0,049 | Gm42595 | ENSMUSG00000042354 | 0,574 | 0,000 | Gnl3 |
| ENSMUSG00000063480 | 0,632 | 0,000 | Snu13 | ENSMUSG00000109941 | 0,571 | 0,032 | Exosc6 |
| ENSMUSG00000033467 | 0,631 | 0,006 | Crlf2 | ENSMUSG00000046768 | 0,570 | 0,019 | Rhoj |
| ENSMUSG00000007617 | 0,631 | 0,002 | Homer1 | ENSMUSG00000026158 | 0,569 | 0,000 | Ogfrl1 |
| ENSMUSG00000033147 | 0,630 | 0,037 | Slc22a15 | ENSMUSG00000000957 | 0,567 | 0,001 | Mmp14 |
| ENSMUSG00000043388 | 0,629 | 0,043 | Tmem130 | ENSMUSG00000020075 | 0,566 | 0,000 | Ddx21 |
| ENSMUSG00000023017 | 0,629 | 0,013 | Asic1 | ENSMUSG00000000317 | 0,566 | 0,049 | Bcl6b |
| ENSMUSG00000083992 | 0,625 | 0,027 | Gm11478 | ENSMUSG00000034842 | 0,562 | 0,026 | Art3 |
| ENSMUSG00000027540 | 0,623 | 0,000 | Ptpn1 | ENSMUSG00000032103 | 0,561 | 0,000 | Pus3 |
| ENSMUSG00000029552 | 0,622 | 0,017 | Tes | ENSMUSG00000034709 | 0,560 | 0,000 | Ppp1r21 |
| ENSMUSG00000003848 | 0,622 | 0,000 | Nob1 | ENSMUSG00000025359 | 0,559 | 0,005 | Pmel |
| ENSMUSG00000025508 | 0,621 | 0,046 | Rplp2 | ENSMUSG00000030701 | 0,559 | 0,035 | Plekhb1 |
| ENSMUSG00000002699 | 0,620 | 0,003 | Lcp2 | ENSMUSG00000028248 | 0,559 | 0,046 | Pnisr |
| ENSMUSG00000051652 | 0,618 | 0,014 | Lrrc3 | ENSMUSG00000023988 | 0,556 | 0,000 | Bysl |
| ENSMUSG00000042364 | 0,617 | 0,000 | Snx18 | ENSMUSG00000018899 | 0,554 | 0,028 | Irf1 |
| ENSMUSG00000051518 | 0,617 | 0,005 | Rps19bp1 | ENSMUSG00000099041 | 0,554 | 0,024 | Gm28035 |
| ENSMUSG00000082433 | 0,616 | 0,037 | Gm9025 | ENSMUSG00000101166 | 0,553 | 0,004 | Gm28496 |
| ENSMUSG00000049950 | 0,616 | 0,006 | Rpp38 | ENSMUSG00000054321 | 0,552 | 0,006 | Taf4b |
| ENSMUSG00000031799 | 0,615 | 0,000 | Tpm4 | ENSMUSG00000039361 | 0,550 | 0,001 | Picalm |
| ENSMUSG00000005374 | 0,615 | 0,000 | Tbl2 | ENSMUSG00000017765 | 0,550 | 0,000 | Slc12a4 |
| ENSMUSG00000034842 | 0,612 | 0,007 | Art3 | ENSMUSG00000000916 | 0,549 | 0,000 | Nsun5 |
| ENSMUSG00000114429 | 0,612 | 0,037 | AC113060.1 | ENSMUSG00000020484 | 0,548 | 0,000 | Xbp1 |
| ENSMUSG00000029752 | 0,611 | 0,010 | Asns | ENSMUSG00000026069 | 0,547 | 0,023 | Il1rl1 |
| ENSMUSG00000028633 | 0,611 | 0,000 | Ctps | ENSMUSG00000046994 | 0,547 | 0,016 | Mars2 |
| ENSMUSG00000001305 | 0,610 | 0,000 | Rrp15 | ENSMUSG00000036560 | 0,546 | 0,010 | Lgi4 |
| ENSMUSG00000036948 | 0,609 | 0,000 | BC037034 | ENSMUSG00000055945 | 0,546 | 0,027 | Prr18 |
| ENSMUSG00000071568 | 0,609 | 0,029 | Gm5874 | ENSMUSG00000016664 | 0,545 | 0,000 | Pacsin2 |
| ENSMUSG00000005672 | 0,609 | 0,042 | Kit | ENSMUSG00000047090 | 0,545 | 0,000 | Tmem198b |
| ENSMUSG00000062093 | 0,608 | 0,031 | Gm10110 | ENSMUSG00000025762 | 0,543 | 0,000 | Larp1b |
| ENSMUSG00000099764 | 0,608 | 0,043 | Rps10-ps2 | ENSMUSG00000029657 | 0,542 | 0,003 | Hsph1 |

| | | | | | |
|---|---|---|---|---|---|
| ENSMUSG00000026158 | 0,608 | 0,000 Ogfrl1 | ENSMUSG00000053907 | 0,541 | 0,000 Mat2a |
| ENSMUSG00000074825 | 0,607 | 0,000 Itpripl1 | ENSMUSG00000028633 | 0,540 | 0,000 Ctps |
| ENSMUSG00000044968 | 0,606 | 0,007 Napepld | ENSMUSG00000034203 | 0,537 | 0,000 Chchd4 |
| ENSMUSG00000037960 | 0,605 | 0,000 Card19 | ENSMUSG00000042766 | 0,537 | 0,041 Trim46 |
| ENSMUSG00000022255 | 0,605 | 0,001 Mtdh | ENSMUSG00000038510 | 0,536 | 0,002 Rpf2 |
| ENSMUSG00000031604 | 0,605 | 0,000 Msmo1 | ENSMUSG00000027368 | 0,535 | 0,026 Dusp2 |
| ENSMUSG00000000916 | 0,605 | 0,000 Nsun5 | ENSMUSG00000027405 | 0,535 | 0,000 Nop56 |
| ENSMUSG00000049232 | 0,604 | 0,000 Tigd2 | ENSMUSG00000037447 | 0,535 | 0,018 Arid5a |
| ENSMUSG00000031608 | 0,603 | 0,015 Galnt7 | ENSMUSG00000028645 | 0,534 | 0,001 Slc2a1 |
| ENSMUSG00000062373 | 0,603 | 0,000 Tmem65 | ENSMUSG00000012429 | 0,534 | 0,017 Mplkip |
| ENSMUSG00000028261 | 0,601 | 0,000 Ndufaf4 | ENSMUSG00000025236 | 0,534 | 0,000 Adpgk |
| ENSMUSG00000028741 | 0,600 | 0,002 Mrto4 | ENSMUSG00000030403 | 0,533 | 0,000 Vasp |
| ENSMUSG00000020307 | 0,598 | 0,001 Cdc34 | ENSMUSG00000028108 | 0,533 | 0,024 Ecm1 |
| ENSMUSG00000043998 | 0,598 | 0,000 Mgat2 | ENSMUSG00000040723 | 0,533 | 0,000 Rcsd1 |
| ENSMUSG00000024985 | 0,595 | 0,001 Tcf7l2 | ENSMUSG00000032109 | 0,532 | 0,000 Nlrx1 |
| ENSMUSG00000085894 | 0,593 | 0,003 Gm15832 | ENSMUSG00000031431 | 0,532 | 0,013 Tsc22d3 |
| ENSMUSG00000041438 | 0,593 | 0,000 Utp4 | ENSMUSG00000034165 | 0,532 | 0,001 Ccnd3 |
| ENSMUSG00000063506 | 0,592 | 0,003 Arhgap22 | ENSMUSG00000034422 | 0,531 | 0,023 Parp14 |
| ENSMUSG00000044768 | 0,592 | 0,000 D1Ertd622e | ENSMUSG00000044968 | 0,531 | 0,031 Napepld |
| ENSMUSG00000033099 | 0,591 | 0,006 Nol12 | ENSMUSG00000112429 | 0,531 | 0,015 AC131081.1 |
| ENSMUSG00000026159 | 0,590 | 0,000 Agfg1 | ENSMUSG00000040511 | 0,531 | 0,002 Pvr |
| ENSMUSG00000031841 | 0,589 | 0,028 Cdh13 | ENSMUSG00000079036 | 0,530 | 0,000 Alkbh1 |
| ENSMUSG00000034321 | 0,589 | 0,000 Exosc1 | ENSMUSG00000022255 | 0,529 | 0,005 Mtdh |
| ENSMUSG00000026974 | 0,589 | 0,000 Zmynd19 | ENSMUSG00000026896 | 0,529 | 0,027 Ifih1 |
| ENSMUSG00000048027 | 0,588 | 0,004 Rgmb | ENSMUSG00000018012 | 0,528 | 0,033 Rac3 |
| ENSMUSG00000115219 | 0,587 | 0,004 Eef1akmt4 | ENSMUSG00000035640 | 0,528 | 0,003 Cbarp |
| ENSMUSG00000054855 | 0,587 | 0,009 Rnd1 | ENSMUSG00000023017 | 0,528 | 0,047 Asic1 |
| ENSMUSG00000044231 | 0,587 | 0,000 Nhlrc1 | ENSMUSG00000008318 | 0,527 | 0,008 Relt |
| ENSMUSG00000022792 | 0,585 | 0,000 Yars2 | ENSMUSG00000020692 | 0,526 | 0,001 Nle1 |
| ENSMUSG00000010358 | 0,585 | 0,012 Ifi35 | ENSMUSG00000050295 | 0,524 | 0,035 Foxc1 |
| ENSMUSG00000047260 | 0,585 | 0,000 Emc6 | ENSMUSG00000020648 | 0,523 | 0,004 Dus4l |
| ENSMUSG00000044763 | 0,583 | 0,000 Trmt10c | ENSMUSG00000005374 | 0,523 | 0,000 Tbl2 |
| ENSMUSG00000107176 | 0,582 | 0,009 Gm9794 | ENSMUSG00000055065 | 0,522 | 0,000 Ddx17 |
| ENSMUSG00000032575 | 0,582 | 0,000 Manf | ENSMUSG00000063480 | 0,522 | 0,000 Snu13 |
| ENSMUSG00000032103 | 0,582 | 0,000 Pus3 | ENSMUSG00000074873 | 0,522 | 0,021 AI606181 |
| ENSMUSG00000074576 | 0,582 | 0,001 Mocs3 | ENSMUSG00000039191 | 0,521 | 0,004 Rbpj |
| ENSMUSG00000091996 | 0,582 | 0,047 BC049352 | ENSMUSG00000085894 | 0,521 | 0,017 Gm15832 |
| ENSMUSG00000027722 | 0,582 | 0,000 Spata5 | ENSMUSG00000031976 | 0,518 | 0,041 Urb2 |
| ENSMUSG00000049241 | 0,581 | 0,033 Hcar1 | ENSMUSG00000019194 | 0,518 | 0,039 Scn1b |
| ENSMUSG00000056121 | 0,580 | 0,000 Fez2 | ENSMUSG00000024891 | 0,517 | 0,001 Slc29a2 |
| ENSMUSG00000006941 | 0,580 | 0,000 Eif1b | ENSMUSG00000048027 | 0,516 | 0,021 Rgmb |
| ENSMUSG00000095464 | 0,579 | 0,041 Gm21987 | ENSMUSG00000031604 | 0,513 | 0,000 Msmo1 |
| ENSMUSG00000035828 | 0,577 | 0,001 Pim3 | ENSMUSG00000003746 | 0,512 | 0,047 Man1a |
| ENSMUSG00000026970 | 0,576 | 0,001 Rbms1 | ENSMUSG00000028060 | 0,512 | 0,005 2810403A07Rik |
| ENSMUSG00000028634 | 0,576 | 0,003 Hivep3 | ENSMUSG00000044763 | 0,511 | 0,000 Trmt10c |
| ENSMUSG00000015766 | 0,576 | 0,026 Eps8 | ENSMUSG00000005373 | 0,511 | 0,005 Mlxipl |
| ENSMUSG00000031024 | 0,576 | 0,000 St5 | ENSMUSG00000035828 | 0,510 | 0,006 Pim3 |

| | | | | | | | |
|---|---|---|---|---|---|---|---|
| ENSMUSG00000108732 | 0,575 | 0,028 | 2310043P16Rik | ENSMUSG00000021036 | 0,509 | 0,000 | Sptlc2 |
| ENSMUSG00000021036 | 0,575 | 0,000 | Sptlc2 | ENSMUSG00000022194 | 0,507 | 0,003 | Pabpn1 |
| ENSMUSG00000037692 | 0,574 | 0,001 | Ahdc1 | ENSMUSG00000035397 | 0,506 | 0,001 | Klf16 |
| ENSMUSG00000021326 | 0,574 | 0,000 | Trim27 | ENSMUSG00000063506 | 0,504 | 0,023 | Arhgap22 |
| ENSMUSG00000025790 | 0,574 | 0,049 | Slco3a1 | ENSMUSG00000059897 | 0,504 | 0,012 | Zfp930 |
| ENSMUSG00000089989 | 0,574 | 0,031 | Gm45713 | ENSMUSG00000022340 | 0,504 | 0,050 | Sybu |
| ENSMUSG00000045962 | 0,573 | 0,040 | Wnk1 | ENSMUSG00000028339 | 0,504 | 0,043 | Col15a1 |
| ENSMUSG00000024777 | 0,573 | 0,000 | Ppp2r5b | ENSMUSG00000002233 | 0,501 | 0,001 | Rhoc |
| ENSMUSG00000045777 | 0,572 | 0,041 | Ifitm10 | ENSMUSG00000031153 | 0,500 | 0,000 | Gripap1 |
| ENSMUSG00000030888 | 0,571 | 0,000 | Rrp8 | ENSMUSG00000045538 | 0,499 | 0,012 | Ddx28 |
| ENSMUSG00000029344 | 0,570 | 0,000 | Tpst2 | ENSMUSG00000026159 | 0,499 | 0,002 | Agfg1 |
| ENSMUSG00000002885 | 0,569 | 0,006 | Adgre5 | ENSMUSG00000043998 | 0,497 | 0,000 | Mgat2 |
| ENSMUSG00000058126 | 0,569 | 0,031 | Tpm3-rs7 | ENSMUSG00000060216 | 0,497 | 0,001 | Arrb2 |
| ENSMUSG00000051495 | 0,569 | 0,000 | Irf2bp2 | ENSMUSG00000026289 | 0,497 | 0,000 | Atg16l1 |
| ENSMUSG00000110275 | 0,569 | 0,037 | Gm5905 | ENSMUSG00000030545 | 0,495 | 0,000 | Pex11a |
| ENSMUSG00000023106 | 0,568 | 0,002 | Denr | ENSMUSG00000037242 | 0,495 | 0,022 | Clic4 |
| ENSMUSG00000075585 | 0,568 | 0,001 | 6330403L08Rik | ENSMUSG00000090015 | 0,495 | 0,043 | Gm15446 |
| ENSMUSG00000031146 | 0,567 | 0,019 | Plp2 | ENSMUSG00000023272 | 0,493 | 0,002 | Creld2 |
| ENSMUSG00000028540 | 0,567 | 0,000 | Dph2 | ENSMUSG00000001467 | 0,493 | 0,004 | Cyp51 |
| ENSMUSG00000074677 | 0,567 | 0,038 | Sirpb1c | ENSMUSG00000037573 | 0,492 | 0,042 | Tob1 |
| ENSMUSG00000023067 | 0,566 | 0,013 | Cdkn1a | ENSMUSG00000053113 | 0,491 | 0,020 | Socs3 |
| ENSMUSG00000074129 | 0,563 | 0,034 | Rpl13a | ENSMUSG00000029570 | 0,491 | 0,000 | Lfng |
| ENSMUSG00000019487 | 0,563 | 0,000 | Trip10 | ENSMUSG00000055491 | 0,490 | 0,000 | Pprc1 |
| ENSMUSG00000039886 | 0,563 | 0,000 | Tmem120a | ENSMUSG00000063694 | 0,489 | 0,009 | Cycs |
| ENSMUSG00000026520 | 0,563 | 0,002 | Pycr2 | ENSMUSG00000020248 | 0,487 | 0,000 | Nfyb |
| ENSMUSG00000058355 | 0,562 | 0,000 | Abce1 | ENSMUSG00000021701 | 0,487 | 0,012 | Plk2 |
| ENSMUSG00000036966 | 0,561 | 0,000 | Spryd3 | ENSMUSG00000042719 | 0,485 | 0,030 | Naa25 |
| ENSMUSG00000034587 | 0,561 | 0,001 | 8430429K09Rik | ENSMUSG00000049482 | 0,484 | 0,008 | Ctu2 |
| ENSMUSG00000031976 | 0,561 | 0,017 | Urb2 | ENSMUSG00000039176 | 0,482 | 0,000 | Polg |
| ENSMUSG00000026596 | 0,561 | 0,046 | Abl2 | ENSMUSG00000029708 | 0,481 | 0,042 | Gcc1 |
| ENSMUSG00000022557 | 0,561 | 0,000 | Bop1 | ENSMUSG00000003134 | 0,480 | 0,000 | Tbc1d8 |
| ENSMUSG00000033545 | 0,560 | 0,000 | Znrf1 | ENSMUSG00000066152 | 0,480 | 0,000 | Slc31a2 |
| ENSMUSG00000032112 | 0,558 | 0,000 | Trappc4 | ENSMUSG00000020527 | 0,479 | 0,000 | Myo19 |
| ENSMUSG00000009293 | 0,556 | 0,000 | Ube2g2 | ENSMUSG00000023068 | 0,478 | 0,000 | Nus1 |
| ENSMUSG00000044533 | 0,556 | 0,025 | Rps2 | ENSMUSG00000052253 | 0,477 | 0,029 | Zfp622 |
| ENSMUSG00000002393 | 0,556 | 0,000 | Nr2f6 | ENSMUSG00000025616 | 0,477 | 0,000 | Usp16 |
| ENSMUSG00000092035 | 0,554 | 0,045 | Peg10 | ENSMUSG00000025602 | 0,476 | 0,021 | Zfp202 |
| ENSMUSG00000031917 | 0,553 | 0,000 | Nip7 | ENSMUSG00000024999 | 0,475 | 0,012 | Noc3l |
| ENSMUSG00000045128 | 0,553 | 0,033 | Rpl18a | ENSMUSG00000026121 | 0,475 | 0,002 | Sema4c |
| ENSMUSG00000036452 | 0,553 | 0,028 | Arhgap26 | ENSMUSG00000028559 | 0,475 | 0,000 | Osbpl9 |
| ENSMUSG00000048277 | 0,552 | 0,001 | Syngr2 | ENSMUSG00000054027 | 0,475 | 0,032 | Nt5dc3 |
| ENSMUSG00000021930 | 0,552 | 0,000 | Spryd7 | ENSMUSG00000026974 | 0,474 | 0,000 | Zmynd19 |
| ENSMUSG00000046516 | 0,552 | 0,045 | Cox17 | ENSMUSG00000043702 | 0,474 | 0,020 | Pde12 |
| ENSMUSG00000034709 | 0,552 | 0,000 | Ppp1r21 | ENSMUSG00000029484 | 0,474 | 0,007 | Anxa3 |
| ENSMUSG00000039191 | 0,551 | 0,001 | Rbpj | ENSMUSG00000020706 | 0,474 | 0,000 | Ftsj3 |
| ENSMUSG00000058546 | 0,551 | 0,039 | Rpl23a | ENSMUSG00000022974 | 0,473 | 0,005 | Paxbp1 |
| ENSMUSG00000078490 | 0,551 | 0,026 | Cfap74 | ENSMUSG00000051451 | 0,473 | 0,004 | Crebzf |

| | | | | | | | |
|---|---|---|---|---|---|---|---|
| ENSMUSG00000040659 | 0,549 | 0,000 | Efhd2 | ENSMUSG00000003848 | 0,472 | 0,000 | Nob1 |
| ENSMUSG00000020553 | 0,548 | 0,013 | Pctp | ENSMUSG00000018339 | 0,472 | 0,003 | Gpx3 |
| ENSMUSG00000085385 | 0,548 | 0,006 | Snhg17 | ENSMUSG00000056260 | 0,472 | 0,041 | Lrif1 |
| ENSMUSG00000074892 | 0,547 | 0,006 | B3galt5 | ENSMUSG00000022247 | 0,472 | 0,008 | Brix1 |
| ENSMUSG00000084319 | 0,546 | 0,000 | Tpt1-ps3 | ENSMUSG00000035047 | 0,471 | 0,000 | Kri1 |
| ENSMUSG00000028645 | 0,545 | 0,001 | Slc2a1 | ENSMUSG00000022792 | 0,471 | 0,001 | Yars2 |
| ENSMUSG00000054079 | 0,545 | 0,000 | Utp18 | ENSMUSG00000045294 | 0,470 | 0,047 | Insig1 |
| ENSMUSG00000039640 | 0,544 | 0,024 | Mrpl12 | ENSMUSG00000057329 | 0,469 | 0,033 | Bcl2 |
| ENSMUSG00000024953 | 0,543 | 0,015 | Prdx5 | ENSMUSG00000041025 | 0,469 | 0,000 | Iffo2 |
| ENSMUSG00000018470 | 0,543 | 0,019 | Kcnab3 | ENSMUSG00000023009 | 0,469 | 0,001 | Nckap5l |
| ENSMUSG00000027613 | 0,542 | 0,001 | Eif6 | ENSMUSG00000048174 | 0,468 | 0,005 | Tmem81 |
| ENSMUSG00000004771 | 0,541 | 0,000 | Rab11a | ENSMUSG00000024777 | 0,468 | 0,000 | Ppp2r5b |
| ENSMUSG00000105913 | 0,540 | 0,017 | Mir5121 | ENSMUSG00000035649 | 0,467 | 0,001 | Zcchc7 |
| ENSMUSG00000054000 | 0,539 | 0,030 | Tusc1 | ENSMUSG00000016477 | 0,466 | 0,043 | E2f3 |
| ENSMUSG00000031825 | 0,539 | 0,040 | Crispld2 | ENSMUSG00000028952 | 0,466 | 0,005 | Zbtb48 |
| ENSMUSG00000027646 | 0,539 | 0,000 | Src | ENSMUSG00000043556 | 0,465 | 0,040 | Fbxl7 |
| ENSMUSG00000098465 | 0,538 | 0,031 | Mir1668 | ENSMUSG00000092232 | 0,464 | 0,001 | Gm20521 |
| ENSMUSG00000038855 | 0,538 | 0,005 | Itpkb | ENSMUSG00000000555 | 0,464 | 0,013 | Itga5 |
| ENSMUSG00000033931 | 0,536 | 0,000 | Rbm34 | ENSMUSG00000025995 | 0,463 | 0,000 | Wdr75 |
| ENSMUSG00000022674 | 0,536 | 0,000 | Ube2v2 | ENSMUSG00000021687 | 0,462 | 0,001 | Scamp1 |
| ENSMUSG00000035640 | 0,536 | 0,002 | Cbarp | ENSMUSG00000027221 | 0,462 | 0,036 | Chst1 |
| ENSMUSG00000029657 | 0,536 | 0,002 | Hsph1 | ENSMUSG00000040296 | 0,462 | 0,032 | Ddx58 |
| ENSMUSG00000059277 | 0,536 | 0,013 | R74862 | ENSMUSG00000025402 | 0,462 | 0,007 | Nab2 |
| ENSMUSG00000047649 | 0,535 | 0,018 | Cd3eap | ENSMUSG00000064289 | 0,461 | 0,001 | Tank |
| ENSMUSG00000030291 | 0,535 | 0,036 | Med21 | ENSMUSG00000034484 | 0,461 | 0,000 | Snx2 |
| ENSMUSG00000062169 | 0,535 | 0,000 | Cnih4 | ENSMUSG00000044224 | 0,461 | 0,000 | Dnajc21 |
| ENSMUSG00000032215 | 0,535 | 0,000 | Rsl24d1 | ENSMUSG00000025436 | 0,461 | 0,008 | Atp23 |
| ENSMUSG00000074622 | 0,535 | 0,000 | Mafb | ENSMUSG00000031799 | 0,460 | 0,000 | Tpm4 |
| ENSMUSG00000042766 | 0,534 | 0,022 | Trim46 | ENSMUSG00000029344 | 0,459 | 0,000 | Tpst2 |
| ENSMUSG00000034892 | 0,534 | 0,047 | Rps29 | ENSMUSG00000097440 | 0,457 | 0,041 | Gm6277 |
| ENSMUSG00000049939 | 0,533 | 0,007 | Lrrc4 | ENSMUSG00000031565 | 0,456 | 0,048 | Fgfr1 |
| ENSMUSG00000047945 | 0,533 | 0,005 | Marcksl1 | ENSMUSG00000041506 | 0,455 | 0,007 | Rrp9 |
| ENSMUSG00000052146 | 0,532 | 0,038 | Rps10 | ENSMUSG00000058355 | 0,455 | 0,001 | Abce1 |
| ENSMUSG00000025762 | 0,530 | 0,000 | Larp1b | ENSMUSG00000047446 | 0,455 | 0,012 | Arl4a |
| ENSMUSG00000020692 | 0,529 | 0,001 | Nle1 | ENSMUSG00000028540 | 0,451 | 0,000 | Dph2 |
| ENSMUSG00000031897 | 0,529 | 0,031 | Psmb10 | ENSMUSG00000032115 | 0,450 | 0,001 | Hyou1 |
| ENSMUSG00000020107 | 0,528 | 0,000 | Anapc16 | ENSMUSG00000045140 | 0,450 | 0,035 | Pigw |
| ENSMUSG00000079037 | 0,528 | 0,006 | Prnp | ENSMUSG00000024037 | 0,450 | 0,000 | Wdr4 |
| ENSMUSG00000018565 | 0,528 | 0,000 | Elp5 | ENSMUSG00000021692 | 0,450 | 0,001 | Dimt1 |
| ENSMUSG00000043445 | 0,527 | 0,001 | Pgp | ENSMUSG00000036634 | 0,449 | 0,046 | Mag |
| ENSMUSG00000024661 | 0,527 | 0,014 | Fth1 | ENSMUSG00000031839 | 0,448 | 0,014 | Hsbp1 |
| ENSMUSG00000006301 | 0,527 | 0,000 | Tmbim1 | ENSMUSG00000034587 | 0,448 | 0,016 | 8430429K09Rik |
| ENSMUSG00000025498 | 0,526 | 0,026 | Irf7 | ENSMUSG00000014905 | 0,447 | 0,043 | Dnajb9 |
| ENSMUSG00000046768 | 0,525 | 0,024 | Rhoj | ENSMUSG00000026796 | 0,447 | 0,003 | Fam129b |
| ENSMUSG00000020397 | 0,524 | 0,000 | Med7 | ENSMUSG00000020057 | 0,446 | 0,047 | Dram1 |
| ENSMUSG00000020901 | 0,523 | 0,003 | Pik3r5 | ENSMUSG00000083012 | 0,446 | 0,000 | Fam220a |
| ENSMUSG00000027787 | 0,522 | 0,000 | Nmd3 | ENSMUSG00000039725 | 0,446 | 0,005 | Trp53rka |

| | | | | | | | |
|---|---|---|---|---|---|---|---|
| ENSMUSG00000027583 | 0,522 | 0,001 | Zbtb46 | ENSMUSG00000033543 | 0,446 | 0,048 | Gtf2a2 |
| ENSMUSG00000027351 | 0,521 | 0,014 | Spred1 | ENSMUSG00000029504 | 0,445 | 0,000 | Ddx51 |
| ENSMUSG00000067274 | 0,521 | 0,001 | Rplp0 | ENSMUSG00000022489 | 0,445 | 0,004 | Pde1b |
| ENSMUSG00000029711 | 0,520 | 0,003 | Epo | ENSMUSG00000002325 | 0,445 | 0,000 | Irf9 |
| ENSMUSG00000026959 | 0,519 | 0,050 | Grin1 | ENSMUSG00000010048 | 0,443 | 0,005 | Ifrd2 |
| ENSMUSG00000057497 | 0,518 | 0,000 | Fam136a | ENSMUSG00000070643 | 0,443 | 0,000 | Sox13 |
| ENSMUSG00000097290 | 0,517 | 0,004 | 1300002E11Rik | ENSMUSG00000029415 | 0,441 | 0,041 | Sdad1 |
| ENSMUSG00000047368 | 0,517 | 0,000 | Abhd17b | ENSMUSG00000028064 | 0,440 | 0,015 | Sema4a |
| ENSMUSG00000083064 | 0,517 | 0,029 | Gm7785 | ENSMUSG00000004127 | 0,440 | 0,000 | Trmt10a |
| ENSMUSG00000008393 | 0,516 | 0,000 | Carhsp1 | ENSMUSG00000052783 | 0,440 | 0,015 | Grk4 |
| ENSMUSG00000039361 | 0,516 | 0,001 | Picalm | ENSMUSG00000027333 | 0,439 | 0,000 | Smox |
| ENSMUSG00000020706 | 0,516 | 0,000 | Ftsj3 | ENSMUSG00000028907 | 0,439 | 0,002 | Utp11 |
| ENSMUSG00000023791 | 0,515 | 0,000 | Pigx | ENSMUSG00000070462 | 0,439 | 0,000 | Tlnrd1 |
| ENSMUSG00000065124 | 0,515 | 0,042 | Snora65 | ENSMUSG00000071645 | 0,439 | 0,000 | Tut1 |
| ENSMUSG00000034165 | 0,513 | 0,001 | Ccnd3 | ENSMUSG00000027540 | 0,438 | 0,002 | Ptpn1 |
| ENSMUSG00000012429 | 0,513 | 0,017 | Mplkip | ENSMUSG00000074480 | 0,438 | 0,017 | Mex3a |
| ENSMUSG00000044477 | 0,512 | 0,000 | Zfand3 | ENSMUSG00000070972 | 0,436 | 0,001 | Dnajc25 |
| ENSMUSG00000046711 | 0,512 | 0,009 | Hmga1 | ENSMUSG00000050244 | 0,435 | 0,006 | Heatr1 |
| ENSMUSG00000039208 | 0,511 | 0,006 | Metrnl | ENSMUSG00000074825 | 0,435 | 0,001 | Itpripl1 |
| ENSMUSG00000062619 | 0,511 | 0,043 | 2310039H08Rik | ENSMUSG00000026819 | 0,435 | 0,029 | Slc25a25 |
| ENSMUSG00000024790 | 0,510 | 0,016 | Sac3d1 | ENSMUSG00000034321 | 0,434 | 0,001 | Exosc1 |
| ENSMUSG00000024429 | 0,510 | 0,001 | Gnl1 | ENSMUSG00000050796 | 0,434 | 0,003 | B3galt6 |
| ENSMUSG00000031029 | 0,510 | 0,010 | Eif3f | ENSMUSG00000056917 | 0,432 | 0,000 | Sipa1 |
| ENSMUSG00000024091 | 0,510 | 0,000 | Vapa | ENSMUSG00000049957 | 0,431 | 0,000 | Ccdc137 |
| ENSMUSG00000032231 | 0,510 | 0,005 | Anxa2 | ENSMUSG00000017057 | 0,430 | 0,038 | Il13ra1 |
| ENSMUSG00000001467 | 0,509 | 0,002 | Cyp51 | ENSMUSG00000025316 | 0,430 | 0,042 | Banp |
| ENSMUSG00000039470 | 0,508 | 0,001 | Zdhhc2 | ENSMUSG00000035049 | 0,430 | 0,001 | Rrp12 |
| ENSMUSG00000033909 | 0,508 | 0,000 | Usp36 | ENSMUSG00000092541 | 0,429 | 0,002 | Gm20537 |
| ENSMUSG00000104950 | 0,508 | 0,017 | 4833413G10Rik | ENSMUSG00000020009 | 0,428 | 0,000 | Ifngr1 |
| ENSMUSG00000049307 | 0,507 | 0,006 | Fut4 | ENSMUSG00000030681 | 0,427 | 0,005 | Mvp |
| ENSMUSG00000042719 | 0,506 | 0,015 | Naa25 | ENSMUSG00000093485 | 0,426 | 0,036 | Gm20708 |
| ENSMUSG00000022442 | 0,505 | 0,000 | Ttll1 | ENSMUSG00000048100 | 0,425 | 0,023 | Taf13 |
| ENSMUSG00000054676 | 0,505 | 0,000 | 1600014C10Rik | ENSMUSG00000037960 | 0,425 | 0,011 | Card19 |
| ENSMUSG00000060579 | 0,505 | 0,027 | Fhit | ENSMUSG00000024330 | 0,425 | 0,045 | Col11a2 |
| ENSMUSG00000061518 | 0,504 | 0,032 | Cox5b | ENSMUSG00000027787 | 0,422 | 0,000 | Nmd3 |
| ENSMUSG00000029860 | 0,504 | 0,000 | Zyx | ENSMUSG00000092203 | 0,421 | 0,042 | 1110038B12Rik |
| ENSMUSG00000049300 | 0,504 | 0,000 | Prmt6 | ENSMUSG00000058392 | 0,421 | 0,000 | Rrp1b |
| ENSMUSG00000003746 | 0,502 | 0,038 | Man1a | ENSMUSG00000018932 | 0,421 | 0,018 | Map2k3 |
| ENSMUSG00000060126 | 0,502 | 0,001 | Tpt1 | ENSMUSG00000005483 | 0,420 | 0,000 | Dnajb1 |
| ENSMUSG00000079677 | 0,502 | 0,040 | Fdx1l | ENSMUSG00000027804 | 0,419 | 0,000 | Ppid |
| ENSMUSG00000018932 | 0,502 | 0,002 | Map2k3 | ENSMUSG00000054079 | 0,419 | 0,000 | Utp18 |
| ENSMUSG00000025995 | 0,500 | 0,000 | Wdr75 | ENSMUSG00000022119 | 0,419 | 0,008 | Rbm26 |
| ENSMUSG00000026470 | 0,499 | 0,000 | Stx6 | ENSMUSG00000030880 | 0,418 | 0,000 | Polr3e |
| ENSMUSG00000090946 | 0,498 | 0,000 | Ccdc71l | ENSMUSG00000033931 | 0,418 | 0,000 | Rbm34 |
| ENSMUSG00000045671 | 0,498 | 0,030 | Spred2 | ENSMUSG00000024081 | 0,415 | 0,000 | Cebpz |
| ENSMUSG00000023286 | 0,497 | 0,001 | Ube2j2 | ENSMUSG00000020932 | 0,415 | 0,023 | Gfap |
| ENSMUSG00000043284 | 0,497 | 0,005 | Tmem11 | ENSMUSG00000109510 | 0,415 | 0,003 | Gm42417 |

| | | | | | |
|---|---|---|---|---|---|
| ENSMUSG00000038214 | 0,497 | 0,016 Bend3 | ENSMUSG00000044231 | 0,414 | 0,021 Nhlrc1 |
| ENSMUSG00000021024 | 0,497 | 0,012 Psma6 | ENSMUSG00000028034 | 0,414 | 0,035 Fubp1 |
| ENSMUSG00000002395 | 0,497 | 0,040 Use1 | ENSMUSG00000054676 | 0,412 | 0,007 1600014C10Rik |
| ENSMUSG00000046994 | 0,497 | 0,020 Mars2 | ENSMUSG00000019774 | 0,412 | 0,012 Mtrf1l |
| ENSMUSG00000028559 | 0,496 | 0,000 Osbpl9 | ENSMUSG00000044768 | 0,412 | 0,001 D1Ertd622e |
| ENSMUSG00000040723 | 0,495 | 0,000 Rcsd1 | ENSMUSG00000027722 | 0,411 | 0,000 Spata5 |
| ENSMUSG00000056124 | 0,495 | 0,000 B4galt6 | ENSMUSG00000032010 | 0,411 | 0,005 Usp2 |
| ENSMUSG00000025477 | 0,494 | 0,000 Inpp5a | ENSMUSG00000006301 | 0,410 | 0,001 Tmbim1 |
| ENSMUSG00000030880 | 0,493 | 0,000 Polr3e | ENSMUSG00000022554 | 0,410 | 0,037 Hgh1 |
| ENSMUSG00000027746 | 0,492 | 0,000 Ufm1 | ENSMUSG00000110423 | 0,410 | 0,048 Gm45736 |
| ENSMUSG00000024436 | 0,491 | 0,001 Mrps18b | ENSMUSG00000038930 | 0,410 | 0,007 Rccd1 |
| ENSMUSG00000012848 | 0,490 | 0,035 Rps5 | ENSMUSG00000045411 | 0,408 | 0,000 2410002F23Rik |
| ENSMUSG00000024456 | 0,490 | 0,000 Diaph1 | ENSMUSG00000028869 | 0,407 | 0,000 Gnl2 |
| ENSMUSG00000031652 | 0,490 | 0,000 N4bp1 | ENSMUSG00000068114 | 0,407 | 0,016 Ccdc134 |
| ENSMUSG00000033543 | 0,489 | 0,018 Gtf2a2 | ENSMUSG00000048787 | 0,407 | 0,030 Dcun1d3 |
| ENSMUSG00000022960 | 0,489 | 0,000 Donson | ENSMUSG00000020424 | 0,407 | 0,049 Gatsl3 |
| ENSMUSG00000025967 | 0,489 | 0,023 Eef1b2 | ENSMUSG00000030888 | 0,406 | 0,001 Rrp8 |
| ENSMUSG00000039633 | 0,488 | 0,025 Lonrf1 | ENSMUSG00000001065 | 0,405 | 0,000 Zfp276 |
| ENSMUSG00000067367 | 0,488 | 0,019 Lyar | ENSMUSG00000027865 | 0,404 | 0,002 Gdap2 |
| ENSMUSG00000056917 | 0,487 | 0,000 Sipa1 | ENSMUSG00000010067 | 0,404 | 0,027 Rassf1 |
| ENSMUSG00000032803 | 0,487 | 0,000 Cdv3 | ENSMUSG00000002733 | 0,404 | 0,010 Plekha3 |
| ENSMUSG00000074182 | 0,487 | 0,003 Znhit6 | ENSMUSG00000018143 | 0,403 | 0,010 Mafk |
| ENSMUSG00000006589 | 0,487 | 0,045 Aprt | ENSMUSG00000025422 | 0,402 | 0,017 Agap2 |
| ENSMUSG00000028898 | 0,485 | 0,025 Trnau1ap | ENSMUSG00000021326 | 0,402 | 0,001 Trim27 |
| ENSMUSG00000109556 | 0,485 | 0,038 Gm38843 | ENSMUSG00000090946 | 0,402 | 0,008 Ccdc71l |
| ENSMUSG00000006333 | 0,484 | 0,038 Rps9 | ENSMUSG00000070476 | 0,401 | 0,034 Fam217b |
| ENSMUSG00000045140 | 0,484 | 0,014 Pigw | ENSMUSG00000048277 | 0,399 | 0,027 Syngr2 |
| ENSMUSG00000067995 | 0,483 | 0,000 Gtf2f2 | ENSMUSG00000002625 | 0,399 | 0,007 Akap8l |
| ENSMUSG00000062742 | 0,482 | 0,029 Gm5239 | ENSMUSG00000022463 | 0,398 | 0,041 Srebf2 |
| ENSMUSG00000054027 | 0,481 | 0,021 Nt5dc3 | ENSMUSG00000039474 | 0,397 | 0,004 Wfs1 |
| ENSMUSG00000026663 | 0,480 | 0,023 Atf6 | ENSMUSG00000031701 | 0,397 | 0,000 Dnaja2 |
| ENSMUSG00000085156 | 0,480 | 0,031 Snhg15 | ENSMUSG00000069844 | 0,396 | 0,003 Sco1 |
| ENSMUSG00000034484 | 0,480 | 0,000 Snx2 | ENSMUSG00000031511 | 0,396 | 0,003 Arhgef7 |
| ENSMUSG00000092203 | 0,480 | 0,012 1110038B12Rik | ENSMUSG00000004040 | 0,396 | 0,041 Stat3 |
| ENSMUSG00000097039 | 0,478 | 0,042 Pvt1 | ENSMUSG00000037752 | 0,395 | 0,002 Xkr8 |
| ENSMUSG00000020116 | 0,478 | 0,000 Pno1 | ENSMUSG00000022434 | 0,394 | 0,002 Fam118a |
| ENSMUSG00000030744 | 0,477 | 0,024 Rps3 | ENSMUSG00000007476 | 0,394 | 0,000 Lrrc8a |
| ENSMUSG00000002985 | 0,477 | 0,005 Apoe | ENSMUSG00000044847 | 0,393 | 0,002 Lsm11 |
| ENSMUSG00000024248 | 0,477 | 0,036 Cox7a2l | ENSMUSG00000028793 | 0,393 | 0,001 Rnf19b |
| ENSMUSG00000031701 | 0,476 | 0,000 Dnaja2 | ENSMUSG00000024055 | 0,393 | 0,018 Cyp4f13 |
| ENSMUSG00000020009 | 0,475 | 0,000 Ifngr1 | ENSMUSG00000039242 | 0,392 | 0,016 B3galnt2 |
| ENSMUSG00000071415 | 0,475 | 0,047 Rpl23 | ENSMUSG00000020107 | 0,391 | 0,007 Anapc16 |
| ENSMUSG00000090137 | 0,474 | 0,036 Uba52 | ENSMUSG00000040010 | 0,391 | 0,032 Slc7a5 |
| ENSMUSG00000020255 | 0,474 | 0,000 D10Wsu102e | ENSMUSG00000109243 | 0,391 | 0,000 Gm45867 |
| ENSMUSG00000014778 | 0,474 | 0,000 Fhod1 | ENSMUSG00000000881 | 0,390 | 0,000 Dlg3 |
| ENSMUSG00000028907 | 0,473 | 0,000 Utp11 | ENSMUSG00000036966 | 0,390 | 0,000 Spryd3 |
| ENSMUSG00000029415 | 0,473 | 0,018 Sdad1 | ENSMUSG00000020397 | 0,388 | 0,008 Med7 |

| | | | | | |
|---|---|---|---|---|---|
| ENSMUSG00000044715 | 0,471 | 0,000 Gskip | ENSMUSG00000023249 | 0,387 | 0,049 Parp3 |
| ENSMUSG00000005481 | 0,470 | 0,000 Ddx39 | ENSMUSG00000028410 | 0,387 | 0,002 Dnaja1 |
| ENSMUSG00000036430 | 0,470 | 0,022 Tbcc | ENSMUSG00000001774 | 0,387 | 0,016 Chordc1 |
| ENSMUSG00000031781 | 0,470 | 0,000 Ciapin1 | ENSMUSG00000029860 | 0,387 | 0,005 Zyx |
| ENSMUSG00000041629 | 0,470 | 0,000 Fam104a | ENSMUSG00000023286 | 0,387 | 0,014 Ube2j2 |
| ENSMUSG00000018293 | 0,469 | 0,010 Pfn1 | ENSMUSG00000021930 | 0,386 | 0,000 Spryd7 |
| ENSMUSG00000020057 | 0,469 | 0,023 Dram1 | ENSMUSG00000055334 | 0,385 | 0,020 Snupn |
| ENSMUSG00000020424 | 0,469 | 0,013 Gatsl3 | ENSMUSG00000004771 | 0,385 | 0,000 Rab11a |
| ENSMUSG00000037348 | 0,469 | 0,000 Paqr7 | ENSMUSG00000001305 | 0,385 | 0,033 Rrp15 |
| ENSMUSG00000024769 | 0,468 | 0,038 Cdc42bpg | ENSMUSG00000025931 | 0,384 | 0,016 Paqr8 |
| ENSMUSG00000026857 | 0,468 | 0,014 Ntmt1 | ENSMUSG00000033106 | 0,384 | 0,001 Slc7a6os |
| ENSMUSG00000051346 | 0,468 | 0,000 Spryd4 | ENSMUSG00000038213 | 0,383 | 0,000 Tapbpl |
| ENSMUSG00000036299 | 0,467 | 0,010 BC031181 | ENSMUSG00000056121 | 0,381 | 0,000 Fez2 |
| ENSMUSG00000027368 | 0,467 | 0,046 Dusp2 | ENSMUSG00000027646 | 0,380 | 0,007 Src |
| ENSMUSG00000097440 | 0,467 | 0,026 Gm6277 | ENSMUSG00000032309 | 0,380 | 0,000 Fbxo22 |
| ENSMUSG00000091462 | 0,467 | 0,009 Gm17084 | ENSMUSG00000103906 | 0,380 | 0,037 Tigd5 |
| ENSMUSG00000039648 | 0,466 | 0,036 Kyat1 | ENSMUSG00000063268 | 0,377 | 0,003 Parp10 |
| ENSMUSG00000020250 | 0,466 | 0,000 Txnrd1 | ENSMUSG00000036160 | 0,377 | 0,000 Surf6 |
| ENSMUSG00000004530 | 0,466 | 0,000 Coro1c | ENSMUSG00000115293 | 0,375 | 0,004 Ece2 |
| ENSMUSG00000022957 | 0,465 | 0,018 Itsn1 | ENSMUSG00000074182 | 0,375 | 0,046 Znhit6 |
| ENSMUSG00000030057 | 0,464 | 0,000 Cnbp | ENSMUSG00000074030 | 0,375 | 0,013 Exoc8 |
| ENSMUSG00000071078 | 0,463 | 0,017 Nr2c2ap | ENSMUSG00000039804 | 0,375 | 0,000 Ncoa5 |
| ENSMUSG00000024818 | 0,463 | 0,001 Slc25a45 | ENSMUSG00000039156 | 0,374 | 0,011 Stim2 |
| ENSMUSG00000029610 | 0,463 | 0,016 Aimp2 | ENSMUSG00000028653 | 0,374 | 0,000 Trit1 |
| ENSMUSG00000043866 | 0,463 | 0,025 Taf10 | ENSMUSG00000041264 | 0,374 | 0,000 Uspl1 |
| ENSMUSG00000071180 | 0,463 | 0,000 Smim15 | ENSMUSG00000054520 | 0,372 | 0,001 Sh3bp2 |
| ENSMUSG00000071072 | 0,463 | 0,001 Ptges3 | ENSMUSG00000030265 | 0,372 | 0,020 Kras |
| ENSMUSG00000053907 | 0,462 | 0,001 Mat2a | ENSMUSG00000033294 | 0,372 | 0,001 Noc4l |
| ENSMUSG00000029507 | 0,462 | 0,007 Pus1 | ENSMUSG00000030232 | 0,371 | 0,000 Aebp2 |
| ENSMUSG00000027405 | 0,461 | 0,000 Nop56 | ENSMUSG00000031627 | 0,370 | 0,013 Irf2 |
| ENSMUSG00000061477 | 0,461 | 0,028 Rps7 | ENSMUSG00000114763 | 0,368 | 0,000 AC134869.3 |
| ENSMUSG00000024037 | 0,460 | 0,000 Wdr4 | ENSMUSG00000062169 | 0,367 | 0,000 Cnih4 |
| ENSMUSG00000005514 | 0,460 | 0,000 Por | ENSMUSG00000031711 | 0,367 | 0,000 Zfp330 |
| ENSMUSG00000042228 | 0,459 | 0,001 Lyn | ENSMUSG00000024188 | 0,366 | 0,000 Luc7l |
| ENSMUSG00000020248 | 0,457 | 0,000 Nfyb | ENSMUSG00000067995 | 0,366 | 0,002 Gtf2f2 |
| ENSMUSG00000003429 | 0,457 | 0,041 Rps11 | ENSMUSG00000000628 | 0,365 | 0,038 Hk2 |
| ENSMUSG00000054520 | 0,457 | 0,000 Sh3bp2 | ENSMUSG00000014907 | 0,364 | 0,012 Naf1 |
| ENSMUSG00000030980 | 0,457 | 0,000 Knop1 | ENSMUSG00000070871 | 0,364 | 0,003 Ccnyl1 |
| ENSMUSG00000034265 | 0,457 | 0,013 Zdhhc14 | ENSMUSG00000038975 | 0,362 | 0,026 Rabggtb |
| ENSMUSG00000017756 | 0,457 | 0,045 Slc12a7 | ENSMUSG00000025939 | 0,361 | 0,018 Ube2w |
| ENSMUSG00000070972 | 0,457 | 0,000 Dnajc25 | ENSMUSG00000032497 | 0,361 | 0,000 Lrrfip2 |
| ENSMUSG00000038034 | 0,456 | 0,040 Igsf8 | ENSMUSG00000041354 | 0,361 | 0,003 Rgl2 |
| ENSMUSG00000028710 | 0,456 | 0,002 Atpaf1 | ENSMUSG00000032462 | 0,361 | 0,033 Pik3cb |
| ENSMUSG00000036992 | 0,456 | 0,014 Nxt1 | ENSMUSG00000027215 | 0,360 | 0,005 Cd82 |
| ENSMUSG00000022009 | 0,455 | 0,000 Nufip1 | ENSMUSG00000002957 | 0,360 | 0,003 Ap2a2 |
| ENSMUSG00000032834 | 0,455 | 0,000 Pwp2 | ENSMUSG00000038578 | 0,359 | 0,020 Susd1 |
| ENSMUSG00000028410 | 0,454 | 0,000 Dnaja1 | ENSMUSG00000045216 | 0,358 | 0,044 Hs6st1 |

| | | | | | | | |
|---|---|---|---|---|---|---|---|
| ENSMUSG00000024081 | 0,454 | 0,000 | Cebpz | ENSMUSG00000044792 | 0,357 | 0,035 | Isca1 |
| ENSMUSG00000000555 | 0,453 | 0,012 | Itga5 | ENSMUSG00000044636 | 0,356 | 0,037 | Csrnp2 |
| ENSMUSG00000046058 | 0,453 | 0,023 | Eid2 | ENSMUSG00000027091 | 0,356 | 0,000 | Zc3h15 |
| ENSMUSG00000048174 | 0,452 | 0,003 | Tmem81 | ENSMUSG00000026927 | 0,355 | 0,000 | Sdccag3 |
| ENSMUSG00000054091 | 0,452 | 0,032 | 1810037I17Rik | ENSMUSG00000037321 | 0,355 | 0,012 | Tap1 |
| ENSMUSG00000021591 | 0,452 | 0,000 | Glrx | ENSMUSG00000029068 | 0,354 | 0,034 | Ccnl2 |
| ENSMUSG00000025103 | 0,451 | 0,000 | Btbd1 | ENSMUSG00000020250 | 0,354 | 0,014 | Txnrd1 |
| ENSMUSG00000030681 | 0,451 | 0,002 | Mvp | ENSMUSG00000016181 | 0,353 | 0,002 | Diexf |
| ENSMUSG00000097059 | 0,450 | 0,008 | Fam120aos | ENSMUSG00000032803 | 0,353 | 0,000 | Cdv3 |
| ENSMUSG00000083012 | 0,450 | 0,000 | Fam220a | ENSMUSG00000045980 | 0,352 | 0,005 | Tmem104 |
| ENSMUSG00000026749 | 0,449 | 0,000 | Nek6 | ENSMUSG00000030057 | 0,352 | 0,004 | Cnbp |
| ENSMUSG00000050777 | 0,449 | 0,019 | Tmem37 | ENSMUSG00000042133 | 0,351 | 0,003 | Ppig |
| ENSMUSG00000042190 | 0,448 | 0,036 | Cmklr1 | ENSMUSG00000074102 | 0,351 | 0,004 | Rbm15b |
| ENSMUSG00000032220 | 0,448 | 0,030 | Myo1e | ENSMUSG00000002147 | 0,351 | 0,002 | Stat6 |
| ENSMUSG00000028416 | 0,448 | 0,032 | Bag1 | ENSMUSG00000029924 | 0,350 | 0,010 | Slc37a3 |
| ENSMUSG00000044709 | 0,447 | 0,029 | Gemin7 | ENSMUSG00000031781 | 0,349 | 0,001 | Ciapin1 |
| ENSMUSG00000029390 | 0,447 | 0,032 | Tmed2 | ENSMUSG00000022575 | 0,349 | 0,011 | Gsdmd |
| ENSMUSG00000049686 | 0,445 | 0,000 | Orai1 | ENSMUSG00000036873 | 0,349 | 0,013 | 2410004B18Rik |
| ENSMUSG00000019868 | 0,445 | 0,000 | Vta1 | ENSMUSG00000047098 | 0,348 | 0,000 | Rnf31 |
| ENSMUSG00000029581 | 0,445 | 0,000 | Fscn1 | ENSMUSG00000024592 | 0,348 | 0,019 | C330018D20Rik |
| ENSMUSG00000038683 | 0,443 | 0,000 | Pak1ip1 | ENSMUSG00000032575 | 0,346 | 0,036 | Manf |
| ENSMUSG00000028683 | 0,443 | 0,002 | Eif2b3 | ENSMUSG00000028889 | 0,346 | 0,001 | Yrdc |
| ENSMUSG00000044149 | 0,443 | 0,002 | Nkrf | ENSMUSG00000036693 | 0,345 | 0,000 | Nop14 |
| ENSMUSG00000015656 | 0,443 | 0,004 | Hspa8 | ENSMUSG00000024436 | 0,344 | 0,035 | Mrps18b |
| ENSMUSG00000016554 | 0,442 | 0,000 | Eif3d | ENSMUSG00000068742 | 0,344 | 0,025 | Cry2 |
| ENSMUSG00000037242 | 0,442 | 0,035 | Clic4 | ENSMUSG00000024456 | 0,343 | 0,009 | Diaph1 |
| ENSMUSG00000008859 | 0,442 | 0,000 | Rala | ENSMUSG00000033285 | 0,342 | 0,001 | Wdr3 |
| ENSMUSG00000037563 | 0,442 | 0,049 | Rps16 | ENSMUSG00000059970 | 0,341 | 0,042 | Hspa2 |
| ENSMUSG00000020087 | 0,441 | 0,000 | Tysnd1 | ENSMUSG00000032285 | 0,341 | 0,001 | Dnaja4 |
| ENSMUSG00000022639 | 0,440 | 0,002 | Dubr | ENSMUSG00000025171 | 0,341 | 0,034 | Ubtd1 |
| ENSMUSG00000021018 | 0,440 | 0,043 | Polr2h | ENSMUSG00000024818 | 0,341 | 0,033 | Slc25a45 |
| ENSMUSG00000027091 | 0,440 | 0,000 | Zc3h15 | ENSMUSG00000028896 | 0,341 | 0,044 | Rcc1 |
| ENSMUSG00000041025 | 0,440 | 0,000 | Iffo2 | ENSMUSG00000001248 | 0,341 | 0,004 | Gramd1a |
| ENSMUSG00000038467 | 0,439 | 0,000 | Chmp4b | ENSMUSG00000071180 | 0,341 | 0,002 | Smim15 |
| ENSMUSG00000038412 | 0,439 | 0,006 | Higd1a | ENSMUSG00000023791 | 0,340 | 0,022 | Pigx |
| ENSMUSG00000078713 | 0,438 | 0,033 | Tomm5 | ENSMUSG00000021501 | 0,339 | 0,000 | Caml |
| ENSMUSG00000042684 | 0,438 | 0,033 | Npl | ENSMUSG00000041707 | 0,339 | 0,010 | 1810011H11Rik |
| ENSMUSG00000069844 | 0,438 | 0,000 | Sco1 | ENSMUSG00000045576 | 0,338 | 0,002 | St7l |
| ENSMUSG00000054766 | 0,438 | 0,000 | Set | ENSMUSG00000030980 | 0,337 | 0,002 | Knop1 |
| ENSMUSG00000018217 | 0,438 | 0,002 | Pmp22 | ENSMUSG00000031878 | 0,337 | 0,002 | Nae1 |
| ENSMUSG00000039680 | 0,438 | 0,021 | Mrps6 | ENSMUSG00000071072 | 0,337 | 0,034 | Ptges3 |
| ENSMUSG00000075701 | 0,437 | 0,000 | Selenos | ENSMUSG00000031652 | 0,337 | 0,035 | N4bp1 |
| ENSMUSG00000049957 | 0,435 | 0,000 | Ccdc137 | ENSMUSG00000061458 | 0,337 | 0,000 | Nol10 |
| ENSMUSG00000017404 | 0,435 | 0,047 | Rpl19 | ENSMUSG00000021591 | 0,336 | 0,018 | Glrx |
| ENSMUSG00000033434 | 0,435 | 0,041 | Gtpbp6 | ENSMUSG00000024091 | 0,336 | 0,006 | Vapa |
| ENSMUSG00000049396 | 0,435 | 0,001 | Gemin4 | ENSMUSG00000027422 | 0,336 | 0,000 | Rrbp1 |
| ENSMUSG00000036398 | 0,434 | 0,031 | Ppp1r11 | ENSMUSG00000024613 | 0,335 | 0,001 | Tcof1 |

| | | | | | |
|---|---|---|---|---|---|
| ENSMUSG00000030265 | 0,434 | 0,003 Kras | ENSMUSG00000025103 | 0,335 | 0,000 Btbd1 |
| ENSMUSG00000010047 | 0,433 | 0,000 Hyal2 | ENSMUSG00000074656 | 0,334 | 0,001 Eif2s2 |
| ENSMUSG00000013701 | 0,433 | 0,000 Timm23 | ENSMUSG00000066735 | 0,334 | 0,021 Vkorc1l1 |
| ENSMUSG00000029484 | 0,432 | 0,012 Anxa3 | ENSMUSG00000028710 | 0,333 | 0,049 Atpaf1 |
| ENSMUSG00000055553 | 0,432 | 0,046 Kxd1 | ENSMUSG00000072258 | 0,333 | 0,037 Taf1a |
| ENSMUSG00000028988 | 0,430 | 0,002 Ctnnbip1 | ENSMUSG00000030619 | 0,332 | 0,005 Eed |
| ENSMUSG00000066152 | 0,430 | 0,001 Slc31a2 | ENSMUSG00000027510 | 0,331 | 0,029 Rbm38 |
| ENSMUSG00000036160 | 0,429 | 0,000 Surf6 | ENSMUSG00000059288 | 0,331 | 0,036 Cdyl |
| ENSMUSG00000019188 | 0,429 | 0,000 H13 | ENSMUSG00000037251 | 0,331 | 0,000 Pomk |
| ENSMUSG00000032010 | 0,429 | 0,002 Usp2 | ENSMUSG00000005481 | 0,331 | 0,030 Ddx39 |
| ENSMUSG00000051256 | 0,427 | 0,001 Jagn1 | ENSMUSG00000061689 | 0,329 | 0,000 Dlgap4 |
| ENSMUSG00000039804 | 0,427 | 0,000 Ncoa5 | ENSMUSG00000050821 | 0,326 | 0,029 Fam131a |
| ENSMUSG00000032463 | 0,427 | 0,001 Faim | ENSMUSG00000020282 | 0,326 | 0,004 Rhbdf1 |
| ENSMUSG00000057134 | 0,426 | 0,000 Ado | ENSMUSG00000008393 | 0,325 | 0,014 Carhsp1 |
| ENSMUSG00000095597 | 0,425 | 0,020 Gm6472 | ENSMUSG00000020361 | 0,324 | 0,000 Hspa4 |
| ENSMUSG00000028670 | 0,425 | 0,004 Lypla2 | ENSMUSG00000020076 | 0,324 | 0,000 Ddx50 |
| ENSMUSG00000059714 | 0,424 | 0,011 Flot1 | ENSMUSG00000024513 | 0,324 | 0,013 Mbd2 |
| ENSMUSG00000048234 | 0,423 | 0,001 Rnf149 | ENSMUSG00000048234 | 0,322 | 0,029 Rnf149 |
| ENSMUSG00000018559 | 0,423 | 0,001 Ctdnep1 | ENSMUSG00000020680 | 0,322 | 0,000 Taf15 |
| ENSMUSG00000024539 | 0,423 | 0,000 Ptpn2 | ENSMUSG00000040025 | 0,322 | 0,001 Ythdf2 |
| ENSMUSG00000033192 | 0,422 | 0,004 Lpcat2 | ENSMUSG00000073771 | 0,321 | 0,049 Btbd19 |
| ENSMUSG00000036560 | 0,422 | 0,047 Lgi4 | ENSMUSG00000054766 | 0,321 | 0,002 Set |
| ENSMUSG00000070319 | 0,422 | 0,018 Eif3g | ENSMUSG00000032834 | 0,321 | 0,001 Pwp2 |
| ENSMUSG00000064289 | 0,421 | 0,003 Tank | ENSMUSG00000028194 | 0,320 | 0,019 Ddah1 |
| ENSMUSG00000020708 | 0,421 | 0,028 Psmc5 | ENSMUSG00000028042 | 0,320 | 0,016 Zbtb7b |
| ENSMUSG00000007836 | 0,419 | 0,000 Hnrnpa0 | ENSMUSG00000034575 | 0,320 | 0,013 Papd7 |
| ENSMUSG00000037251 | 0,419 | 0,000 Pomk | ENSMUSG00000017999 | 0,319 | 0,000 Ddx27 |
| ENSMUSG00000033499 | 0,419 | 0,000 Larp4b | ENSMUSG00000026519 | 0,319 | 0,031 Tmem63a |
| ENSMUSG00000021458 | 0,418 | 0,006 2010111I01Rik | ENSMUSG00000025880 | 0,318 | 0,046 Smad7 |
| ENSMUSG00000029014 | 0,418 | 0,001 Dnajc2 | ENSMUSG00000018583 | 0,318 | 0,000 G3bp1 |
| ENSMUSG00000079036 | 0,417 | 0,000 Alkbh1 | ENSMUSG00000019814 | 0,318 | 0,002 Ltv1 |
| ENSMUSG00000091811 | 0,417 | 0,049 Inafm1 | ENSMUSG00000023044 | 0,318 | 0,000 Csad |
| ENSMUSG00000060860 | 0,417 | 0,042 Ube2s | ENSMUSG00000034252 | 0,318 | 0,000 Senp6 |
| ENSMUSG00000058558 | 0,417 | 0,041 Rpl5 | ENSMUSG00000041130 | 0,317 | 0,000 Zfp598 |
| ENSMUSG00000054321 | 0,416 | 0,032 Taf4b | ENSMUSG00000059866 | 0,317 | 0,022 Tnip2 |
| ENSMUSG00000038539 | 0,416 | 0,012 Atf5 | ENSMUSG00000001833 | 0,317 | 0,043 Sept7 |
| ENSMUSG00000029504 | 0,416 | 0,000 Ddx51 | ENSMUSG00000029063 | 0,317 | 0,002 Nadk |
| ENSMUSG00000034974 | 0,415 | 0,017 Dapk3 | ENSMUSG00000050846 | 0,316 | 0,035 Zfp623 |
| ENSMUSG00000048486 | 0,415 | 0,000 Fitm2 | ENSMUSG00000024287 | 0,316 | 0,041 Thoc1 |
| ENSMUSG00000032115 | 0,414 | 0,003 Hyou1 | ENSMUSG00000039159 | 0,316 | 0,000 Ube2h |
| ENSMUSG00000014859 | 0,414 | 0,000 E2f4 | ENSMUSG00000042208 | 0,315 | 0,041 0610010F05Rik |
| ENSMUSG00000090996 | 0,412 | 0,011 Gm20458 | ENSMUSG00000036934 | 0,315 | 0,015 4921524J17Rik |
| ENSMUSG00000020133 | 0,412 | 0,008 2310011J03Rik | ENSMUSG00000058503 | 0,314 | 0,000 Fam133b |
| ENSMUSG00000044792 | 0,412 | 0,008 Isca1 | ENSMUSG00000022257 | 0,314 | 0,005 Laptm4b |
| ENSMUSG00000024725 | 0,411 | 0,010 Ostf1 | ENSMUSG00000024045 | 0,313 | 0,002 Akap8 |
| ENSMUSG00000087589 | 0,411 | 0,042 D430040D24Rik | ENSMUSG00000061950 | 0,313 | 0,013 Ppp4r1 |
| ENSMUSG00000004040 | 0,410 | 0,024 Stat3 | ENSMUSG00000026341 | 0,313 | 0,017 Actr3 |

| | | | | | | | |
|---|---|---|---|---|---|---|---|
| ENSMUSG00000030019 | 0,410 | 0,001 | Fbxl14 | ENSMUSG00000005514 | 0,312 | 0,001 | Por |
| ENSMUSG00000022752 | 0,410 | 0,000 | Tomm70a | ENSMUSG00000032215 | 0,311 | 0,031 | Rsl24d1 |
| ENSMUSG00000046731 | 0,409 | 0,008 | Kctd11 | ENSMUSG00000025477 | 0,310 | 0,001 | Inpp5a |
| ENSMUSG00000004665 | 0,409 | 0,017 | Cnn2 | ENSMUSG00000009535 | 0,310 | 0,000 | Rnmt |
| ENSMUSG00000010048 | 0,408 | 0,007 | Ifrd2 | ENSMUSG00000026806 | 0,310 | 0,004 | Ddx31 |
| ENSMUSG00000020935 | 0,408 | 0,000 | Dcakd | ENSMUSG00000036054 | 0,310 | 0,001 | Sugp2 |
| ENSMUSG00000031320 | 0,408 | 0,033 | Rps4x | ENSMUSG00000002845 | 0,309 | 0,010 | Tmem39a |
| ENSMUSG00000044496 | 0,408 | 0,000 | 2510039O18Rik | ENSMUSG00000009293 | 0,309 | 0,001 | Ube2g2 |
| ENSMUSG00000056076 | 0,408 | 0,000 | Eif3b | ENSMUSG00000049232 | 0,308 | 0,034 | Tigd2 |
| ENSMUSG00000074247 | 0,407 | 0,001 | Dda1 | ENSMUSG00000024966 | 0,308 | 0,003 | Stip1 |
| ENSMUSG00000024513 | 0,407 | 0,001 | Mbd2 | ENSMUSG00000038517 | 0,307 | 0,001 | Tbkbp1 |
| ENSMUSG00000002957 | 0,407 | 0,000 | Ap2a2 | ENSMUSG00000020246 | 0,307 | 0,041 | Hcfc2 |
| ENSMUSG00000031513 | 0,407 | 0,000 | Leprotl1 | ENSMUSG00000019082 | 0,307 | 0,012 | Slc25a22 |
| ENSMUSG00000022257 | 0,406 | 0,000 | Laptm4b | ENSMUSG00000107881 | 0,305 | 0,045 | Gm44250 |
| ENSMUSG00000002733 | 0,406 | 0,007 | Plekha3 | ENSMUSG00000055239 | 0,305 | 0,000 | Kcmf1 |
| ENSMUSG00000074102 | 0,406 | 0,000 | Rbm15b | ENSMUSG00000032423 | 0,304 | 0,018 | Syncrip |
| ENSMUSG00000030879 | 0,405 | 0,011 | Mrpl17 | ENSMUSG00000041459 | 0,304 | 0,006 | Tardbp |
| ENSMUSG00000055334 | 0,405 | 0,009 | Snupn | ENSMUSG00000028156 | 0,303 | 0,002 | Eif4e |
| ENSMUSG00000033918 | 0,403 | 0,005 | Parl | ENSMUSG00000022498 | 0,303 | 0,000 | Txndc11 |
| ENSMUSG00000024613 | 0,403 | 0,000 | Tcof1 | ENSMUSG00000038507 | 0,303 | 0,033 | Parp12 |
| ENSMUSG00000035047 | 0,403 | 0,001 | Kri1 | ENSMUSG00000030942 | 0,302 | 0,000 | Thumpd1 |
| ENSMUSG00000020358 | 0,403 | 0,002 | Hnrnpab | ENSMUSG00000026864 | 0,302 | 0,006 | Hspa5 |
| ENSMUSG00000028194 | 0,403 | 0,001 | Ddah1 | ENSMUSG00000025138 | 0,301 | 0,034 | Sirt7 |
| ENSMUSG00000026203 | 0,402 | 0,001 | Dnajb2 | ENSMUSG00000000532 | 0,300 | 0,049 | Acvr1b |
| ENSMUSG00000067212 | 0,402 | 0,023 | H2-T23 | ENSMUSG00000045248 | 0,300 | 0,009 | Med26 |
| ENSMUSG00000004460 | 0,402 | 0,000 | Dnajb11 | ENSMUSG00000039630 | 0,300 | 0,000 | Hnrnpu |
| ENSMUSG00000046722 | 0,402 | 0,000 | Cdc42se1 | ENSMUSG00000045624 | 0,299 | 0,018 | Esf1 |
| ENSMUSG00000025792 | 0,401 | 0,001 | Slc25a10 | ENSMUSG00000021392 | 0,299 | 0,026 | Nol8 |
| ENSMUSG00000055491 | 0,401 | 0,001 | Pprc1 | ENSMUSG00000008859 | 0,299 | 0,001 | Rala |
| ENSMUSG00000024966 | 0,401 | 0,000 | Stip1 | ENSMUSG00000051391 | 0,298 | 0,035 | Ywhag |
| ENSMUSG00000039067 | 0,400 | 0,002 | Psmd7 | ENSMUSG00000010047 | 0,298 | 0,004 | Hyal2 |
| ENSMUSG00000032309 | 0,400 | 0,000 | Fbxo22 | ENSMUSG00000032307 | 0,297 | 0,012 | Ube2q2 |
| ENSMUSG00000045980 | 0,399 | 0,001 | Tmem104 | ENSMUSG00000021938 | 0,297 | 0,040 | Pspc1 |
| ENSMUSG00000024785 | 0,399 | 0,002 | Rcl1 | ENSMUSG00000037348 | 0,296 | 0,007 | Paqr7 |
| ENSMUSG00000016181 | 0,399 | 0,000 | Diexf | ENSMUSG00000022442 | 0,294 | 0,016 | Ttll1 |
| ENSMUSG00000037805 | 0,398 | 0,047 | Rpl10a | ENSMUSG00000024219 | 0,293 | 0,017 | Anks1 |
| ENSMUSG00000022463 | 0,398 | 0,029 | Srebf2 | ENSMUSG00000014859 | 0,293 | 0,000 | E2f4 |
| ENSMUSG00000020955 | 0,398 | 0,049 | Ap4s1 | ENSMUSG00000023007 | 0,293 | 0,034 | Prpf40b |
| ENSMUSG00000041012 | 0,398 | 0,015 | Cmtm8 | ENSMUSG00000019302 | 0,292 | 0,001 | Atp6v0a1 |
| ENSMUSG00000074030 | 0,398 | 0,004 | Exoc8 | ENSMUSG00000014177 | 0,292 | 0,006 | Tvp23b |
| ENSMUSG00000038387 | 0,397 | 0,040 | Rras | ENSMUSG00000074781 | 0,291 | 0,010 | Ube2n |
| ENSMUSG00000020648 | 0,397 | 0,022 | Dus4l | ENSMUSG00000044477 | 0,291 | 0,000 | Zfand3 |
| ENSMUSG00000030224 | 0,397 | 0,000 | Strap | ENSMUSG00000030533 | 0,290 | 0,039 | Unc45a |
| ENSMUSG00000001416 | 0,396 | 0,005 | Cct3 | ENSMUSG00000039294 | 0,287 | 0,004 | BC017643 |
| ENSMUSG00000039159 | 0,396 | 0,000 | Ube2h | ENSMUSG00000050088 | 0,287 | 0,001 | 1600012H06Rik |
| ENSMUSG00000025316 | 0,396 | 0,049 | Banp | ENSMUSG00000011958 | 0,287 | 0,000 | Bnip2 |
| ENSMUSG00000074656 | 0,396 | 0,000 | Eif2s2 | ENSMUSG00000010453 | 0,287 | 0,013 | Kansl3 |

| | | | | | | | |
|---|---|---|---|---|---|---|---|
| ENSMUSG00000039244 | 0,396 | 0,000 | E130309D02Rik | ENSMUSG00000019927 | 0,286 | 0,001 | Ube2d1 |
| ENSMUSG00000078427 | 0,396 | 0,011 | Sarnp | ENSMUSG00000002017 | 0,286 | 0,016 | Fam98a |
| ENSMUSG00000020644 | 0,395 | 0,026 | Id2 | ENSMUSG00000022752 | 0,286 | 0,018 | Tomm70a |
| ENSMUSG00000039156 | 0,395 | 0,004 | Stim2 | ENSMUSG00000049300 | 0,286 | 0,011 | Prmt6 |
| ENSMUSG00000095567 | 0,395 | 0,000 | Noc2l | ENSMUSG00000026470 | 0,286 | 0,000 | Stx6 |
| ENSMUSG00000036873 | 0,394 | 0,003 | 2410004B18Rik | ENSMUSG00000038683 | 0,285 | 0,031 | Pak1ip1 |
| ENSMUSG00000018040 | 0,393 | 0,000 | Rrp7a | ENSMUSG00000029071 | 0,285 | 0,003 | Dvl1 |
| ENSMUSG00000066735 | 0,392 | 0,003 | Vkorc1l1 | ENSMUSG00000041426 | 0,285 | 0,015 | Hibch |
| ENSMUSG00000034201 | 0,391 | 0,000 | Gas2l1 | ENSMUSG00000078786 | 0,284 | 0,047 | BC024978 |
| ENSMUSG00000092310 | 0,391 | 0,049 | Gm20509 | ENSMUSG00000036707 | 0,284 | 0,009 | Cab39 |
| ENSMUSG00000092541 | 0,390 | 0,005 | Gm20537 | ENSMUSG00000025337 | 0,283 | 0,017 | Sbds |
| ENSMUSG00000026121 | 0,390 | 0,010 | Sema4c | ENSMUSG00000038612 | 0,282 | 0,012 | Mcl1 |
| ENSMUSG00000030199 | 0,390 | 0,014 | Etv6 | ENSMUSG00000026283 | 0,281 | 0,043 | Ing5 |
| ENSMUSG00000018433 | 0,388 | 0,000 | Nol11 | ENSMUSG00000021428 | 0,280 | 0,000 | Riok1 |
| ENSMUSG00000022517 | 0,388 | 0,000 | Mgrn1 | ENSMUSG00000041629 | 0,280 | 0,037 | Fam104a |
| ENSMUSG00000028609 | 0,388 | 0,036 | Magoh | ENSMUSG00000091896 | 0,279 | 0,002 | Ube2d2a |
| ENSMUSG00000029924 | 0,387 | 0,002 | Slc37a3 | ENSMUSG00000053253 | 0,279 | 0,026 | Ndfip2 |
| ENSMUSG00000027475 | 0,387 | 0,010 | Kif3b | ENSMUSG00000024150 | 0,278 | 0,001 | Mcfd2 |
| ENSMUSG00000032078 | 0,387 | 0,018 | Zpr1 | ENSMUSG00000068798 | 0,278 | 0,004 | Rap1a |
| ENSMUSG00000040482 | 0,386 | 0,018 | Dxo | ENSMUSG00000024539 | 0,278 | 0,043 | Ptpn2 |
| ENSMUSG00000059897 | 0,386 | 0,046 | Zfp930 | ENSMUSG00000009630 | 0,278 | 0,000 | Ppp2cb |
| ENSMUSG00000029066 | 0,385 | 0,032 | Mrpl20 | ENSMUSG00000022855 | 0,277 | 0,001 | Senp2 |
| ENSMUSG00000103906 | 0,385 | 0,018 | Tigd5 | ENSMUSG00000032557 | 0,277 | 0,000 | Uba5 |
| ENSMUSG00000064267 | 0,385 | 0,042 | Hvcn1 | ENSMUSG00000033499 | 0,276 | 0,001 | Larp4b |
| ENSMUSG00000067722 | 0,385 | 0,015 | BC003965 | ENSMUSG00000036737 | 0,276 | 0,005 | Oxsr1 |
| ENSMUSG00000078652 | 0,385 | 0,000 | Psme3 | ENSMUSG00000021116 | 0,275 | 0,002 | Eif2s1 |
| ENSMUSG00000029063 | 0,385 | 0,000 | Nadk | ENSMUSG00000047866 | 0,274 | 0,000 | Lonp2 |
| ENSMUSG00000071379 | 0,385 | 0,007 | Hpcal1 | ENSMUSG00000030400 | 0,274 | 0,000 | Ercc2 |
| ENSMUSG00000006818 | 0,385 | 0,003 | Sod2 | ENSMUSG00000075701 | 0,274 | 0,047 | Selenos |
| ENSMUSG00000017781 | 0,384 | 0,000 | Pitpna | ENSMUSG00000033953 | 0,273 | 0,007 | Ppp3r1 |
| ENSMUSG00000029246 | 0,383 | 0,004 | Ppat | ENSMUSG00000063785 | 0,272 | 0,002 | Utp14a |
| ENSMUSG00000025236 | 0,383 | 0,001 | Adpgk | ENSMUSG00000025024 | 0,271 | 0,002 | Smndc1 |
| ENSMUSG00000049091 | 0,383 | 0,002 | Sephs2 | ENSMUSG00000018433 | 0,271 | 0,001 | Nol11 |
| ENSMUSG00000017734 | 0,382 | 0,016 | Dbndd2 | ENSMUSG00000030224 | 0,270 | 0,000 | Strap |
| ENSMUSG00000030545 | 0,382 | 0,006 | Pex11a | ENSMUSG00000009566 | 0,270 | 0,004 | Fpgs |
| ENSMUSG00000017747 | 0,382 | 0,000 | Ghdc | ENSMUSG00000073639 | 0,270 | 0,045 | Rab18 |
| ENSMUSG00000014402 | 0,382 | 0,018 | Tsg101 | ENSMUSG00000032745 | 0,269 | 0,002 | Gpbp1 |
| ENSMUSG00000060708 | 0,381 | 0,039 | Bloc1s4 | ENSMUSG00000045598 | 0,269 | 0,050 | Zfp553 |
| ENSMUSG00000029145 | 0,381 | 0,019 | Eif2b4 | ENSMUSG00000034595 | 0,266 | 0,024 | Ppp1r18 |
| ENSMUSG00000032280 | 0,381 | 0,011 | Tle3 | ENSMUSG00000078652 | 0,266 | 0,000 | Psme3 |
| ENSMUSG00000032119 | 0,380 | 0,035 | Hinfp | ENSMUSG00000053289 | 0,265 | 0,007 | Ddx10 |
| ENSMUSG00000042354 | 0,380 | 0,006 | Gnl3 | ENSMUSG00000030036 | 0,265 | 0,029 | Mogs |
| ENSMUSG00000033953 | 0,379 | 0,000 | Ppp3r1 | ENSMUSG00000037965 | 0,264 | 0,024 | Zc3h7a |
| ENSMUSG00000097729 | 0,379 | 0,034 | 2310015A10Rik | ENSMUSG00000059939 | 0,264 | 0,003 | 9430015G10Rik |
| ENSMUSG00000010054 | 0,378 | 0,001 | Tusc2 | ENSMUSG00000089917 | 0,264 | 0,012 | Uckl1 |
| ENSMUSG00000020561 | 0,378 | 0,014 | Twistnb | ENSMUSG00000030847 | 0,262 | 0,041 | Bag3 |
| ENSMUSG00000041645 | 0,377 | 0,000 | Ddx24 | ENSMUSG00000020530 | 0,261 | 0,005 | Ggnbp2 |

| | | | | | |
|---|---|---|---|---|---|
| ENSMUSG00000021116 | 0,377 | 0,000 Eif2s1 | ENSMUSG00000032212 | 0,261 | 0,002 Sltm |
| ENSMUSG00000045211 | 0,377 | 0,003 Nudt18 | ENSMUSG00000019254 | 0,260 | 0,031 Ppp1r12c |
| ENSMUSG00000026341 | 0,377 | 0,002 Actr3 | ENSMUSG00000024862 | 0,260 | 0,007 Klc2 |
| ENSMUSG00000033685 | 0,377 | 0,005 Ucp2 | ENSMUSG00000020935 | 0,259 | 0,030 Dcakd |
| ENSMUSG00000038374 | 0,377 | 0,036 Rbm8a | ENSMUSG00000022300 | 0,259 | 0,035 Dcaf13 |
| ENSMUSG00000109243 | 0,377 | 0,000 Gm45867 | ENSMUSG00000036002 | 0,259 | 0,009 Fam214b |
| ENSMUSG00000028729 | 0,376 | 0,007 Ebna1bp2 | ENSMUSG00000006373 | 0,258 | 0,012 Pgrmc1 |
| ENSMUSG00000045503 | 0,376 | 0,026 Sys1 | ENSMUSG00000000384 | 0,258 | 0,011 Tbrg4 |
| ENSMUSG00000027865 | 0,376 | 0,003 Gdap2 | ENSMUSG00000024404 | 0,257 | 0,029 Riok3 |
| ENSMUSG00000051391 | 0,376 | 0,003 Ywhag | ENSMUSG00000057497 | 0,257 | 0,020 Fam136a |
| ENSMUSG00000054793 | 0,376 | 0,010 Cadm4 | ENSMUSG00000057554 | 0,256 | 0,040 Lgals8 |
| ENSMUSG00000030942 | 0,375 | 0,000 Thumpd1 | ENSMUSG00000026353 | 0,255 | 0,003 Ubxn4 |
| ENSMUSG00000006304 | 0,375 | 0,000 Arpc2 | ENSMUSG00000025609 | 0,255 | 0,017 Mkln1 |
| ENSMUSG00000061458 | 0,375 | 0,000 Nol10 | ENSMUSG00000027176 | 0,254 | 0,004 Cstf3 |
| ENSMUSG00000014907 | 0,375 | 0,006 Naf1 | ENSMUSG00000040250 | 0,253 | 0,034 Ints13 |
| ENSMUSG00000025264 | 0,375 | 0,000 Tsr2 | ENSMUSG00000024528 | 0,253 | 0,017 Srfbp1 |
| ENSMUSG00000054237 | 0,374 | 0,015 Fra10ac1 | ENSMUSG00000022634 | 0,251 | 0,006 Yaf2 |
| ENSMUSG00000057788 | 0,374 | 0,018 Ddx49 | ENSMUSG00000027746 | 0,251 | 0,015 Ufm1 |
| ENSMUSG00000035505 | 0,374 | 0,011 Cox18 | ENSMUSG00000038524 | 0,251 | 0,004 Fchsd1 |
| ENSMUSG00000030232 | 0,374 | 0,000 Aebp2 | ENSMUSG00000095567 | 0,250 | 0,028 Noc2l |
| ENSMUSG00000074781 | 0,374 | 0,000 Ube2n | ENSMUSG00000022052 | 0,250 | 0,025 Ppp2r2a |
| ENSMUSG00000004451 | 0,373 | 0,000 Ralb | ENSMUSG00000022241 | 0,249 | 0,000 Tars |
| ENSMUSG00000066553 | 0,372 | 0,006 Gm6969 | ENSMUSG00000019868 | 0,248 | 0,048 Vta1 |
| ENSMUSG00000092595 | 0,372 | 0,016 Gm20427 | ENSMUSG00000044496 | 0,248 | 0,000 2510039O18Rik |
| ENSMUSG00000024999 | 0,372 | 0,050 Noc3l | ENSMUSG00000020283 | 0,248 | 0,008 Pex13 |
| ENSMUSG00000070003 | 0,372 | 0,046 Ssbp4 | ENSMUSG00000020432 | 0,248 | 0,025 Tcn2 |
| ENSMUSG00000006392 | 0,371 | 0,013 Med8 | ENSMUSG00000041645 | 0,247 | 0,008 Ddx24 |
| ENSMUSG00000028563 | 0,370 | 0,013 Tm2d1 | ENSMUSG00000028149 | 0,247 | 0,003 Rap1gds1 |
| ENSMUSG00000040010 | 0,370 | 0,033 Slc7a5 | ENSMUSG00000052144 | 0,246 | 0,047 Ppp4r2 |
| ENSMUSG00000078931 | 0,369 | 0,001 Pdf | ENSMUSG00000041623 | 0,246 | 0,037 D11Wsu47e |
| ENSMUSG00000031586 | 0,369 | 0,019 Rbpms | ENSMUSG00000003808 | 0,245 | 0,012 Farsa |
| ENSMUSG00000079658 | 0,369 | 0,022 Eloc | ENSMUSG00000024350 | 0,244 | 0,002 Dnajc18 |
| ENSMUSG00000024327 | 0,368 | 0,002 Slc39a7 | ENSMUSG00000008604 | 0,244 | 0,001 Ubqln4 |
| ENSMUSG00000028318 | 0,368 | 0,033 Polr1e | ENSMUSG00000032422 | 0,244 | 0,005 Snx14 |
| ENSMUSG00000020549 | 0,368 | 0,000 Elac2 | ENSMUSG00000003813 | 0,244 | 0,048 Rad23a |
| ENSMUSG00000036693 | 0,367 | 0,000 Nop14 | ENSMUSG00000029528 | 0,243 | 0,011 Pxn |
| ENSMUSG00000000532 | 0,366 | 0,008 Acvr1b | ENSMUSG00000017747 | 0,243 | 0,042 Ghdc |
| ENSMUSG00000024891 | 0,366 | 0,014 Slc29a2 | ENSMUSG00000018965 | 0,243 | 0,005 Ywhah |
| ENSMUSG00000039474 | 0,366 | 0,005 Wfs1 | ENSMUSG00000047714 | 0,243 | 0,003 Ppp1r2 |
| ENSMUSG00000021905 | 0,365 | 0,005 Dph3 | ENSMUSG00000020630 | 0,242 | 0,046 Rnaseh1 |
| ENSMUSG00000020088 | 0,365 | 0,000 Sar1a | ENSMUSG00000066150 | 0,241 | 0,019 Slc31a1 |
| ENSMUSG00000013150 | 0,365 | 0,013 Gfod2 | ENSMUSG00000004069 | 0,241 | 0,000 Dnaja3 |
| ENSMUSG00000022313 | 0,365 | 0,008 Utp23 | ENSMUSG00000104953 | 0,241 | 0,009 AC147560.1 |
| ENSMUSG00000001674 | 0,364 | 0,000 Ddx18 | ENSMUSG00000017781 | 0,240 | 0,004 Pitpna |
| ENSMUSG00000044018 | 0,364 | 0,002 Mrpl50 | ENSMUSG00000070730 | 0,240 | 0,001 Rmdn3 |
| ENSMUSG00000022241 | 0,364 | 0,000 Tars | ENSMUSG00000001674 | 0,240 | 0,037 Ddx18 |
| ENSMUSG00000022300 | 0,364 | 0,001 Dcaf13 | ENSMUSG00000031422 | 0,239 | 0,011 Morf4l2 |

| | | | | | |
|---|---|---|---|---|---|
| ENSMUSG00000055239 | 0,363 | 0,000 Kcmf1 | ENSMUSG00000045795 | 0,239 | 0,013 Whamm |
| ENSMUSG00000031711 | 0,363 | 0,000 Zfp330 | ENSMUSG00000067787 | 0,239 | 0,001 Blcap |
| ENSMUSG00000035027 | 0,363 | 0,041 Map2k2 | ENSMUSG00000004460 | 0,239 | 0,018 Dnajb11 |
| ENSMUSG00000053094 | 0,363 | 0,000 Tmem248 | ENSMUSG00000030138 | 0,238 | 0,005 Bms1 |
| ENSMUSG00000026806 | 0,363 | 0,000 Ddx31 | ENSMUSG00000049327 | 0,237 | 0,007 Kmt5a |
| ENSMUSG00000071645 | 0,363 | 0,001 Tut1 | ENSMUSG00000032320 | 0,235 | 0,040 Rcn2 |
| ENSMUSG00000074480 | 0,363 | 0,046 Mex3a | ENSMUSG00000023883 | 0,235 | 0,002 Phf10 |
| ENSMUSG00000024800 | 0,362 | 0,015 Rpp30 | ENSMUSG00000029474 | 0,233 | 0,028 Rnf34 |
| ENSMUSG00000040025 | 0,362 | 0,000 Ythdf2 | ENSMUSG00000026930 | 0,232 | 0,035 Gpsm1 |
| ENSMUSG00000061979 | 0,362 | 0,038 Rcc1l | ENSMUSG00000052681 | 0,232 | 0,045 Rap1b |
| ENSMUSG00000042558 | 0,362 | 0,015 Adprhl2 | ENSMUSG00000040659 | 0,231 | 0,002 Efhd2 |
| ENSMUSG00000024387 | 0,361 | 0,018 Csnk2b | ENSMUSG00000031931 | 0,231 | 0,028 Ankrd49 |
| ENSMUSG00000064037 | 0,361 | 0,010 Gpn1 | ENSMUSG00000030846 | 0,231 | 0,030 Tial1 |
| ENSMUSG00000029599 | 0,360 | 0,000 Ddx54 | ENSMUSG00000074405 | 0,231 | 0,021 Zfp865 |
| ENSMUSG00000044224 | 0,360 | 0,000 Dnajc21 | ENSMUSG00000020255 | 0,231 | 0,018 D10Wsu102e |
| ENSMUSG00000030036 | 0,360 | 0,001 Mogs | ENSMUSG00000022538 | 0,230 | 0,013 Lsg1 |
| ENSMUSG00000020458 | 0,358 | 0,024 Rtn4 | ENSMUSG00000028849 | 0,230 | 0,018 Map7d1 |
| ENSMUSG00000037347 | 0,358 | 0,001 Chst7 | ENSMUSG00000040028 | 0,230 | 0,012 Elavl1 |
| ENSMUSG00000014177 | 0,357 | 0,000 Tvp23b | ENSMUSG00000025287 | 0,230 | 0,039 Acot9 |
| ENSMUSG00000031904 | 0,357 | 0,024 Slc7a6 | ENSMUSG00000053094 | 0,230 | 0,001 Tmem248 |
| ENSMUSG00000028608 | 0,355 | 0,012 0610037L13Rik | ENSMUSG00000044715 | 0,230 | 0,010 Gskip |
| ENSMUSG00000049401 | 0,354 | 0,016 Ogfr | ENSMUSG00000024896 | 0,228 | 0,010 Minpp1 |
| ENSMUSG00000019302 | 0,353 | 0,000 Atp6v0a1 | ENSMUSG00000020288 | 0,228 | 0,050 Ahsa2 |
| ENSMUSG00000067787 | 0,353 | 0,000 Blcap | ENSMUSG00000011179 | 0,227 | 0,038 Odc1 |
| ENSMUSG00000035278 | 0,353 | 0,038 Plekhj1 | ENSMUSG00000058440 | 0,227 | 0,000 Nrf1 |
| ENSMUSG00000028042 | 0,352 | 0,004 Zbtb7b | ENSMUSG00000029291 | 0,227 | 0,000 Rufy3 |
| ENSMUSG00000000384 | 0,352 | 0,000 Tbrg4 | ENSMUSG00000038848 | 0,226 | 0,015 Ythdf1 |
| ENSMUSG00000047866 | 0,352 | 0,000 Lonp2 | ENSMUSG00000020132 | 0,226 | 0,033 Rab21 |
| ENSMUSG00000032977 | 0,352 | 0,003 Fam207a | ENSMUSG00000026917 | 0,225 | 0,005 Wdr5 |
| ENSMUSG00000032462 | 0,352 | 0,028 Pik3cb | ENSMUSG00000064145 | 0,225 | 0,003 Arih2 |
| ENSMUSG00000093904 | 0,352 | 0,001 Tomm20 | ENSMUSG00000034120 | 0,223 | 0,002 Srsf2 |
| ENSMUSG00000028149 | 0,352 | 0,000 Rap1gds1 | ENSMUSG00000029036 | 0,223 | 0,045 Atad3a |
| ENSMUSG00000028899 | 0,351 | 0,035 Taf12 | ENSMUSG00000035764 | 0,223 | 0,035 Fbxo45 |
| ENSMUSG00000068114 | 0,351 | 0,032 Ccdc134 | ENSMUSG00000015733 | 0,223 | 0,023 Capza2 |
| ENSMUSG00000019804 | 0,351 | 0,002 Snx3 | ENSMUSG00000026889 | 0,222 | 0,007 Rbm18 |
| ENSMUSG00000022538 | 0,350 | 0,000 Lsg1 | ENSMUSG00000022556 | 0,222 | 0,047 Hsf1 |
| ENSMUSG00000056692 | 0,350 | 0,000 D17Wsu92e | ENSMUSG00000015714 | 0,222 | 0,001 Cers2 |
| ENSMUSG00000091896 | 0,350 | 0,000 Ube2d2a | ENSMUSG00000018442 | 0,222 | 0,003 Derl2 |
| ENSMUSG00000028889 | 0,350 | 0,001 Yrdc | ENSMUSG00000028041 | 0,221 | 0,022 Adam15 |
| ENSMUSG00000029038 | 0,350 | 0,049 Ssu72 | ENSMUSG00000025264 | 0,220 | 0,031 Tsr2 |
| ENSMUSG00000037788 | 0,349 | 0,006 Vopp1 | ENSMUSG00000052915 | 0,220 | 0,006 Msl1 |
| ENSMUSG00000028869 | 0,349 | 0,001 Gnl2 | ENSMUSG00000022517 | 0,220 | 0,015 Mgrn1 |
| ENSMUSG00000037752 | 0,349 | 0,005 Xkr8 | ENSMUSG00000005813 | 0,219 | 0,001 Metap1 |
| ENSMUSG00000025871 | 0,348 | 0,000 4833439L19Rik | ENSMUSG00000029629 | 0,219 | 0,002 Phf14 |
| ENSMUSG00000061032 | 0,348 | 0,031 Rrp1 | ENSMUSG00000024097 | 0,218 | 0,029 Srsf7 |
| ENSMUSG00000058392 | 0,347 | 0,001 Rrp1b | ENSMUSG00000041360 | 0,218 | 0,004 Pum3 |
| ENSMUSG00000050821 | 0,347 | 0,012 Fam131a | ENSMUSG00000029439 | 0,217 | 0,044 Sfswap |

| | | | | | | | |
|---|---|---|---|---|---|---|---|
| ENSMUSG00000021692 | 0,347 | 0,008 | Dimt1 | ENSMUSG00000024359 | 0,216 | 0,001 | Hspa9 |
| ENSMUSG00000032418 | 0,347 | 0,044 | Me1 | ENSMUSG00000020549 | 0,216 | 0,020 | Elac2 |
| ENSMUSG00000029447 | 0,347 | 0,000 | Cct6a | ENSMUSG00000030374 | 0,216 | 0,001 | Strn4 |
| ENSMUSG00000026820 | 0,347 | 0,032 | Ptges2 | ENSMUSG00000020116 | 0,215 | 0,010 | Pno1 |
| ENSMUSG00000017999 | 0,347 | 0,000 | Ddx27 | ENSMUSG00000031568 | 0,214 | 0,050 | Rwdd4a |
| ENSMUSG00000033106 | 0,346 | 0,002 | Slc7a6os | ENSMUSG00000021621 | 0,214 | 0,030 | Zcchc9 |
| ENSMUSG00000022634 | 0,346 | 0,000 | Yaf2 | ENSMUSG00000031984 | 0,214 | 0,001 | 2810004N23Rik |
| ENSMUSG00000036733 | 0,344 | 0,039 | Rbm42 | ENSMUSG00000004530 | 0,213 | 0,019 | Coro1c |
| ENSMUSG00000045411 | 0,344 | 0,001 | 2410002F23Rik | ENSMUSG00000005204 | 0,213 | 0,000 | Senp3 |
| ENSMUSG00000024150 | 0,344 | 0,000 | Mcfd2 | ENSMUSG00000068882 | 0,211 | 0,024 | Ssb |
| ENSMUSG00000070953 | 0,344 | 0,004 | Rabepk | ENSMUSG00000026234 | 0,209 | 0,011 | Ncl |
| ENSMUSG00000031885 | 0,344 | 0,041 | Cbfb | ENSMUSG00000038299 | 0,208 | 0,020 | Wdr36 |
| ENSMUSG00000026917 | 0,343 | 0,000 | Wdr5 | ENSMUSG00000030304 | 0,208 | 0,029 | Ergic2 |
| ENSMUSG00000041360 | 0,343 | 0,000 | Pum3 | ENSMUSG00000063334 | 0,208 | 0,001 | Krr1 |
| ENSMUSG00000032788 | 0,343 | 0,010 | Pdxk | ENSMUSG00000025646 | 0,207 | 0,026 | Atrip |
| ENSMUSG00000033809 | 0,342 | 0,023 | Alg3 | ENSMUSG00000006019 | 0,206 | 0,041 | Dhx34 |
| ENSMUSG00000027488 | 0,342 | 0,018 | Snta1 | ENSMUSG00000032612 | 0,206 | 0,000 | Usp4 |
| ENSMUSG00000099681 | 0,342 | 0,044 | 1700052K11Rik | ENSMUSG00000011257 | 0,205 | 0,003 | Pabpc4 |
| ENSMUSG00000003808 | 0,342 | 0,000 | Farsa | ENSMUSG00000023307 | 0,203 | 0,003 | March5 |
| ENSMUSG00000028936 | 0,342 | 0,049 | Rpl22 | ENSMUSG00000020115 | 0,203 | 0,014 | Tbk1 |
| ENSMUSG00000021969 | 0,341 | 0,009 | Zdhhc20 | ENSMUSG00000041837 | 0,203 | 0,036 | Pdcd7 |
| ENSMUSG00000021598 | 0,341 | 0,039 | Med10 | ENSMUSG00000027374 | 0,203 | 0,024 | Mrps5 |
| ENSMUSG00000043716 | 0,341 | 0,045 | Rpl7 | ENSMUSG00000024182 | 0,201 | 0,004 | Axin1 |
| ENSMUSG00000034259 | 0,340 | 0,011 | Exosc4 | ENSMUSG00000037553 | 0,200 | 0,018 | Zdhhc18 |
| ENSMUSG00000019297 | 0,340 | 0,001 | Nop9 | ENSMUSG00000038563 | 0,197 | 0,021 | Efl1 |
| ENSMUSG00000058440 | 0,339 | 0,000 | Nrf1 | ENSMUSG00000026571 | 0,196 | 0,024 | Dcaf6 |
| ENSMUSG00000066150 | 0,338 | 0,000 | Slc31a1 | ENSMUSG00000028086 | 0,196 | 0,040 | Fbxw7 |
| ENSMUSG00000024048 | 0,338 | 0,014 | Myl12a | ENSMUSG00000032042 | 0,195 | 0,039 | Srpr |
| ENSMUSG00000018583 | 0,338 | 0,000 | G3bp1 | ENSMUSG00000005312 | 0,193 | 0,044 | Ubqln1 |
| ENSMUSG00000059248 | 0,337 | 0,000 | Sept9 | ENSMUSG00000029064 | 0,193 | 0,009 | Gnb1 |
| ENSMUSG00000032497 | 0,336 | 0,001 | Lrrfip2 | ENSMUSG00000029028 | 0,189 | 0,004 | Lrrc47 |
| ENSMUSG00000045934 | 0,335 | 0,043 | Mtmr11 | ENSMUSG00000029447 | 0,189 | 0,040 | Cct6a |
| ENSMUSG00000022035 | 0,335 | 0,009 | Ccdc25 | ENSMUSG00000056050 | 0,189 | 0,048 | Mia3 |
| ENSMUSG00000028156 | 0,334 | 0,000 | Eif4e | ENSMUSG00000025178 | 0,188 | 0,007 | Pi4k2a |
| ENSMUSG00000026817 | 0,334 | 0,014 | Ak1 | ENSMUSG00000056820 | 0,188 | 0,007 | Tsnax |
| ENSMUSG00000029780 | 0,334 | 0,007 | Nt5c3 | ENSMUSG00000029500 | 0,187 | 0,030 | Pgam5 |
| ENSMUSG00000020733 | 0,334 | 0,005 | Slc9a3r1 | ENSMUSG00000042901 | 0,187 | 0,043 | Aida |
| ENSMUSG00000032557 | 0,333 | 0,000 | Uba5 | ENSMUSG00000007411 | 0,186 | 0,001 | Mark3 |
| ENSMUSG00000003813 | 0,333 | 0,002 | Rad23a | ENSMUSG00000020088 | 0,184 | 0,016 | Sar1a |
| ENSMUSG00000032563 | 0,333 | 0,000 | Mrpl3 | ENSMUSG00000023075 | 0,184 | 0,050 | Akirin1 |
| ENSMUSG00000005846 | 0,333 | 0,005 | Rsl1d1 | ENSMUSG00000079562 | 0,184 | 0,001 | Maea |
| ENSMUSG00000071256 | 0,333 | 0,030 | Zfp213 | ENSMUSG00000021096 | 0,183 | 0,001 | Ppm1a |
| ENSMUSG00000026049 | 0,332 | 0,027 | Tex30 | ENSMUSG00000056692 | 0,183 | 0,001 | D17Wsu92e |
| ENSMUSG00000040699 | 0,332 | 0,041 | Limd2 | ENSMUSG00000071646 | 0,182 | 0,044 | Mta2 |
| ENSMUSG00000034024 | 0,332 | 0,001 | Cct2 | ENSMUSG00000032870 | 0,181 | 0,025 | Smap2 |
| ENSMUSG00000048707 | 0,331 | 0,002 | Tprn | ENSMUSG00000014601 | 0,180 | 0,001 | Strip1 |
| ENSMUSG00000050017 | 0,330 | 0,000 | Pitpnb | ENSMUSG00000001785 | 0,177 | 0,012 | Pwp1 |

| | | | | | | | |
|---|---|---|---|---|---|---|---|
| ENSMUSG00000073771 | 0,330 | 0,027 | Btbd19 | ENSMUSG00000038990 | 0,177 | 0,045 | Cables2 |
| ENSMUSG00000022184 | 0,330 | 0,016 | Fbxo4 | ENSMUSG00000028161 | 0,177 | 0,025 | Ppp3ca |
| ENSMUSG00000027968 | 0,330 | 0,014 | Larp7 | ENSMUSG00000047731 | 0,170 | 0,018 | Wbp1l |
| ENSMUSG00000037296 | 0,330 | 0,015 | Lsm1 | ENSMUSG00000027706 | 0,170 | 0,048 | Sec62 |
| ENSMUSG00000025155 | 0,329 | 0,007 | Dus1l | ENSMUSG00000026926 | 0,169 | 0,000 | Pmpca |
| ENSMUSG00000022434 | 0,328 | 0,008 | Fam118a | ENSMUSG00000029227 | 0,169 | 0,045 | Fip1l1 |
| ENSMUSG00000025580 | 0,328 | 0,015 | Eif4a3 | ENSMUSG00000033335 | 0,165 | 0,043 | Dnm2 |
| ENSMUSG00000045598 | 0,328 | 0,007 | Zfp553 | ENSMUSG00000032582 | 0,163 | 0,005 | Rbm6 |
| ENSMUSG00000052151 | 0,328 | 0,049 | Plpp2 | ENSMUSG00000024527 | 0,162 | 0,015 | Afg3l2 |
| ENSMUSG00000115293 | 0,327 | 0,009 | Ece2 | ENSMUSG00000059119 | 0,159 | 0,027 | Nap1l4 |
| ENSMUSG00000104953 | 0,327 | 0,000 | AC147560.1 | ENSMUSG00000031575 | 0,158 | 0,004 | Ash2l |
| ENSMUSG00000020092 | 0,327 | 0,012 | Pald1 | ENSMUSG00000046722 | 0,157 | 0,010 | Cdc42se1 |
| ENSMUSG00000021242 | 0,327 | 0,019 | Npc2 | ENSMUSG00000014959 | 0,153 | 0,018 | Gorasp2 |
| ENSMUSG00000029050 | 0,327 | 0,048 | Ski | ENSMUSG00000051934 | 0,152 | 0,048 | Spats2 |
| ENSMUSG00000021645 | 0,326 | 0,011 | Smn1 | ENSMUSG00000037669 | 0,151 | 0,043 | Ldah |
| ENSMUSG00000045624 | 0,326 | 0,006 | Esf1 | ENSMUSG00000062270 | 0,141 | 0,019 | Morf4l1 |
| ENSMUSG00000042660 | 0,326 | 0,025 | Wdr55 | ENSMUSG00000024587 | 0,139 | 0,035 | Nars |
| ENSMUSG00000078768 | 0,325 | 0,047 | Zfp566 | ENSMUSG00000029518 | 0,129 | 0,019 | Rab35 |
| ENSMUSG00000028848 | 0,325 | 0,044 | Gpn2 | ENSMUSG00000026848 | 0,126 | 0,019 | Tor1b |
| ENSMUSG00000002147 | 0,325 | 0,003 | Stat6 | ENSMUSG00000079477 | 0,121 | 0,046 | Rab7 |
| ENSMUSG00000056091 | 0,325 | 0,008 | St3gal5 | ENSMUSG00000024853 | -0,117 | 0,046 | Sf3b2 |
| ENSMUSG00000003068 | 0,324 | 0,021 | Stk11 | ENSMUSG00000017721 | -0,126 | 0,048 | Pigt |
| ENSMUSG00000018848 | 0,324 | 0,001 | Rars | ENSMUSG00000111497 | -0,136 | 0,013 | Raver1 |
| ENSMUSG00000031627 | 0,324 | 0,027 | Irf2 | ENSMUSG00000013787 | -0,140 | 0,016 | Ehmt2 |
| ENSMUSG00000029064 | 0,324 | 0,000 | Gnb1 | ENSMUSG00000026269 | -0,144 | 0,037 | Rnpepl1 |
| ENSMUSG00000028152 | 0,323 | 0,000 | Tspan5 | ENSMUSG00000029103 | -0,146 | 0,049 | Lrpap1 |
| ENSMUSG00000027339 | 0,323 | 0,045 | Rassf2 | ENSMUSG00000034951 | -0,148 | 0,045 | Cog7 |
| ENSMUSG00000061689 | 0,322 | 0,000 | Dlgap4 | ENSMUSG00000020265 | -0,161 | 0,037 | Sumo3 |
| ENSMUSG00000028796 | 0,322 | 0,001 | Phc2 | ENSMUSG00000026554 | -0,164 | 0,041 | Dcaf8 |
| ENSMUSG00000002257 | 0,322 | 0,024 | Def6 | ENSMUSG00000010936 | -0,164 | 0,007 | Vac14 |
| ENSMUSG00000054302 | 0,322 | 0,004 | Eapp | ENSMUSG00000024096 | -0,164 | 0,027 | Ralbp1 |
| ENSMUSG00000014959 | 0,322 | 0,000 | Gorasp2 | ENSMUSG00000001424 | -0,164 | 0,011 | Snd1 |
| ENSMUSG00000020283 | 0,322 | 0,000 | Pex13 | ENSMUSG00000030884 | -0,165 | 0,001 | Uqcrc2 |
| ENSMUSG00000024213 | 0,321 | 0,014 | Nudt3 | ENSMUSG00000028656 | -0,167 | 0,047 | Cap1 |
| ENSMUSG00000018697 | 0,321 | 0,004 | Aatf | ENSMUSG00000026853 | -0,168 | 0,021 | Crat |
| ENSMUSG00000003134 | 0,321 | 0,012 | Tbc1d8 | ENSMUSG00000025534 | -0,169 | 0,009 | Gusb |
| ENSMUSG00000039176 | 0,321 | 0,001 | Polg | ENSMUSG00000038861 | -0,169 | 0,019 | Pi4kb |
| ENSMUSG00000039183 | 0,319 | 0,015 | Nubp2 | ENSMUSG00000020792 | -0,173 | 0,024 | Exoc7 |
| ENSMUSG00000021501 | 0,319 | 0,001 | Caml | ENSMUSG00000107283 | -0,177 | 0,017 | Mpv17 |
| ENSMUSG00000042111 | 0,319 | 0,006 | Ccdc115 | ENSMUSG00000053333 | -0,177 | 0,043 | Dis3l2 |
| ENSMUSG00000025647 | 0,318 | 0,010 | Shisa5 | ENSMUSG00000049339 | -0,178 | 0,019 | Retreg2 |
| ENSMUSG00000032171 | 0,318 | 0,048 | Pin1 | ENSMUSG00000034371 | -0,179 | 0,023 | Tkfc |
| ENSMUSG00000028041 | 0,317 | 0,000 | Adam15 | ENSMUSG00000000552 | -0,180 | 0,044 | Zfp385a |
| ENSMUSG00000091780 | 0,317 | 0,041 | Sco2 | ENSMUSG00000032187 | -0,181 | 0,032 | Smarca4 |
| ENSMUSG00000020246 | 0,317 | 0,022 | Hcfc2 | ENSMUSG00000024740 | -0,182 | 0,036 | Ddb1 |
| ENSMUSG00000037197 | 0,317 | 0,008 | Rbm17 | ENSMUSG00000032388 | -0,183 | 0,021 | Spg21 |
| ENSMUSG00000047714 | 0,317 | 0,000 | Ppp1r2 | ENSMUSG00000015889 | -0,184 | 0,018 | Lta4h |

| | | | | | | | |
|---|---|---|---|---|---|---|---|
| ENSMUSG00000035890 | 0,316 | 0,029 | Rnf126 | ENSMUSG00000025041 | -0,185 | 0,026 | Nt5c2 |
| ENSMUSG00000059820 | 0,316 | 0,006 | AU019823 | ENSMUSG00000025736 | -0,188 | 0,013 | Jmjd8 |
| ENSMUSG00000028955 | 0,316 | 0,000 | Vamp3 | ENSMUSG00000020375 | -0,189 | 0,033 | Rufy1 |
| ENSMUSG00000024583 | 0,316 | 0,004 | Txnl1 | ENSMUSG00000028757 | -0,190 | 0,048 | Ddost |
| ENSMUSG00000089832 | 0,316 | 0,002 | Shkbp1 | ENSMUSG00000029345 | -0,190 | 0,025 | Tfip11 |
| ENSMUSG00000024875 | 0,316 | 0,019 | Yif1a | ENSMUSG00000022051 | -0,190 | 0,011 | Bnip3l |
| ENSMUSG00000018442 | 0,316 | 0,000 | Derl2 | ENSMUSG00000022678 | -0,191 | 0,036 | Nde1 |
| ENSMUSG00000053253 | 0,316 | 0,006 | Ndfip2 | ENSMUSG00000005469 | -0,191 | 0,000 | Prkaca |
| ENSMUSG00000020432 | 0,316 | 0,002 | Tcn2 | ENSMUSG00000034189 | -0,191 | 0,026 | Hsdl1 |
| ENSMUSG00000006705 | 0,315 | 0,005 | Pknox1 | ENSMUSG00000021218 | -0,191 | 0,003 | Gdi2 |
| ENSMUSG00000025940 | 0,315 | 0,008 | Tmem70 | ENSMUSG00000031864 | -0,196 | 0,017 | Ints10 |
| ENSMUSG00000029028 | 0,314 | 0,000 | Lrrc47 | ENSMUSG00000028954 | -0,196 | 0,039 | Nub1 |
| ENSMUSG00000036376 | 0,314 | 0,050 | Abt1 | ENSMUSG00000052395 | -0,197 | 0,007 | Rft1 |
| ENSMUSG00000030189 | 0,314 | 0,018 | Ybx3 | ENSMUSG00000036980 | -0,197 | 0,026 | Taf6 |
| ENSMUSG00000032567 | 0,314 | 0,043 | Aste1 | ENSMUSG00000027810 | -0,198 | 0,019 | Eif2a |
| ENSMUSG00000021774 | 0,314 | 0,001 | Ube2e1 | ENSMUSG00000068335 | -0,202 | 0,038 | Dok1 |
| ENSMUSG00000029036 | 0,314 | 0,001 | Atad3a | ENSMUSG00000027809 | -0,202 | 0,004 | Etfdh |
| ENSMUSG00000020476 | 0,313 | 0,000 | Dbnl | ENSMUSG00000038417 | -0,202 | 0,022 | Fig4 |
| ENSMUSG00000025337 | 0,312 | 0,005 | Sbds | ENSMUSG00000032527 | -0,202 | 0,017 | Pccb |
| ENSMUSG00000027357 | 0,312 | 0,047 | Crls1 | ENSMUSG00000029536 | -0,203 | 0,030 | Gatc |
| ENSMUSG00000038213 | 0,312 | 0,001 | Tapbpl | ENSMUSG00000032513 | -0,204 | 0,028 | Gorasp1 |
| ENSMUSG00000015363 | 0,310 | 0,005 | Trabd | ENSMUSG00000024507 | -0,206 | 0,004 | Hsd17b4 |
| ENSMUSG00000009566 | 0,310 | 0,000 | Fpgs | ENSMUSG00000063801 | -0,207 | 0,003 | Ap3s2 |
| ENSMUSG00000041354 | 0,310 | 0,010 | Rgl2 | ENSMUSG00000000171 | -0,207 | 0,047 | Sdhd |
| ENSMUSG00000037331 | 0,310 | 0,028 | Larp1 | ENSMUSG00000014748 | -0,207 | 0,003 | Tex261 |
| ENSMUSG00000008496 | 0,310 | 0,022 | Pou2f2 | ENSMUSG00000022965 | -0,208 | 0,007 | Ifngr2 |
| ENSMUSG00000020922 | 0,310 | 0,050 | Lsm12 | ENSMUSG00000001082 | -0,209 | 0,048 | Mfsd10 |
| ENSMUSG00000079111 | 0,310 | 0,001 | Kdelr2 | ENSMUSG00000022620 | -0,210 | 0,029 | Arsa |
| ENSMUSG00000025227 | 0,309 | 0,016 | Mfsd13a | ENSMUSG00000042590 | -0,212 | 0,007 | Ipo11 |
| ENSMUSG00000035764 | 0,309 | 0,001 | Fbxo45 | ENSMUSG00000073096 | -0,215 | 0,040 | Lrrc61 |
| ENSMUSG00000020361 | 0,308 | 0,000 | Hspa4 | ENSMUSG00000002871 | -0,215 | 0,047 | Tpra1 |
| ENSMUSG00000063229 | 0,308 | 0,045 | Ldha | ENSMUSG00000032264 | -0,217 | 0,044 | Zw10 |
| ENSMUSG00000025384 | 0,308 | 0,001 | Faap100 | ENSMUSG00000042032 | -0,217 | 0,003 | Mat2b |
| ENSMUSG00000055065 | 0,308 | 0,047 | Ddx17 | ENSMUSG00000022889 | -0,217 | 0,035 | Mrpl39 |
| ENSMUSG00000083282 | 0,307 | 0,007 | Ctsf | ENSMUSG00000050043 | -0,218 | 0,011 | Tmx2 |
| ENSMUSG00000038712 | 0,307 | 0,013 | Mindy1 | ENSMUSG00000028291 | -0,218 | 0,018 | Akirin2 |
| ENSMUSG00000013698 | 0,306 | 0,001 | Pea15a | ENSMUSG00000032547 | -0,218 | 0,009 | Ryk |
| ENSMUSG00000026723 | 0,306 | 0,045 | Trdmt1 | ENSMUSG00000029007 | -0,219 | 0,021 | Agtrap |
| ENSMUSG00000032479 | 0,306 | 0,011 | Map4 | ENSMUSG00000021646 | -0,219 | 0,016 | Mccc2 |
| ENSMUSG00000057130 | 0,305 | 0,018 | Txnl4a | ENSMUSG00000000594 | -0,219 | 0,026 | Gm2a |
| ENSMUSG00000032290 | 0,305 | 0,049 | Ptpn9 | ENSMUSG00000029550 | -0,220 | 0,000 | Sppl3 |
| ENSMUSG00000027422 | 0,305 | 0,000 | Rrbp1 | ENSMUSG00000049287 | -0,221 | 0,014 | Iba57 |
| ENSMUSG00000041264 | 0,304 | 0,001 | Uspl1 | ENSMUSG00000025722 | -0,222 | 0,027 | Wdr73 |
| ENSMUSG00000025178 | 0,304 | 0,000 | Pi4k2a | ENSMUSG00000020818 | -0,222 | 0,011 | Mfsd11 |
| ENSMUSG00000036707 | 0,304 | 0,003 | Cab39 | ENSMUSG00000033933 | -0,224 | 0,008 | Vhl |
| ENSMUSG00000009630 | 0,304 | 0,000 | Ppp2cb | ENSMUSG00000005981 | -0,225 | 0,020 | Trap1 |
| ENSMUSG00000070705 | 0,303 | 0,037 | Eid2b | ENSMUSG00000020538 | -0,227 | 0,049 | Srebf1 |

| | | | | | |
|---|---|---|---|---|---|
| ENSMUSG00000032285 | 0,303 | 0,004 Dnaja4 | ENSMUSG00000029592 | -0,228 | 0,032 Usp30 |
| ENSMUSG00000062580 | 0,303 | 0,010 Timm17a | ENSMUSG00000029073 | -0,228 | 0,028 Cptp |
| ENSMUSG00000029500 | 0,302 | 0,000 Pgam5 | ENSMUSG00000009079 | -0,228 | 0,005 Ewsr1 |
| ENSMUSG00000020869 | 0,302 | 0,000 Lrrc59 | ENSMUSG00000039637 | -0,229 | 0,044 Coro7 |
| ENSMUSG00000043683 | 0,302 | 0,002 Fem1a | ENSMUSG00000031161 | -0,229 | 0,003 Hdac6 |
| ENSMUSG00000005312 | 0,302 | 0,000 Ubqln1 | ENSMUSG00000024197 | -0,229 | 0,025 Plin3 |
| ENSMUSG00000020392 | 0,301 | 0,030 Cdkn2aipnl | ENSMUSG00000029578 | -0,230 | 0,003 Wipi2 |
| ENSMUSG00000024712 | 0,301 | 0,001 Rfk | ENSMUSG00000021048 | -0,233 | 0,031 Mthfd1 |
| ENSMUSG00000030871 | 0,301 | 0,004 Ears2 | ENSMUSG00000001034 | -0,234 | 0,013 Mapk7 |
| ENSMUSG00000048429 | 0,301 | 0,000 Timm29 | ENSMUSG00000022750 | -0,234 | 0,006 Klhl22 |
| ENSMUSG00000019927 | 0,301 | 0,000 Ube2d1 | ENSMUSG00000035206 | -0,235 | 0,005 Sppl2b |
| ENSMUSG00000033728 | 0,301 | 0,001 Lrrc14 | ENSMUSG00000029705 | -0,236 | 0,016 Cux1 |
| ENSMUSG00000029147 | 0,301 | 0,001 Ppm1g | ENSMUSG00000027523 | -0,236 | 0,042 Gnas |
| ENSMUSG00000015733 | 0,301 | 0,001 Capza2 | ENSMUSG00000037344 | -0,237 | 0,019 Slc12a9 |
| ENSMUSG00000002017 | 0,300 | 0,007 Fam98a | ENSMUSG00000005505 | -0,239 | 0,007 Kbtbd4 |
| ENSMUSG00000049090 | 0,300 | 0,024 Zadh2 | ENSMUSG00000006288 | -0,239 | 0,011 Ttc5 |
| ENSMUSG00000025287 | 0,299 | 0,002 Acot9 | ENSMUSG00000026792 | -0,239 | 0,017 Lrsam1 |
| ENSMUSG00000046792 | 0,299 | 0,010 Zfp787 | ENSMUSG00000031077 | -0,239 | 0,048 Fadd |
| ENSMUSG00000053289 | 0,298 | 0,001 Ddx10 | ENSMUSG00000029175 | -0,241 | 0,040 Slc35f6 |
| ENSMUSG00000032553 | 0,298 | 0,000 Srprb | ENSMUSG00000013646 | -0,241 | 0,006 Sh3bp5l |
| ENSMUSG00000037553 | 0,298 | 0,000 Zdhhc18 | ENSMUSG00000038545 | -0,242 | 0,006 Cul7 |
| ENSMUSG00000025939 | 0,298 | 0,048 Ube2w | ENSMUSG00000079334 | -0,242 | 0,029 Nat6 |
| ENSMUSG00000034729 | 0,297 | 0,036 Mrps10 | ENSMUSG00000029247 | -0,243 | 0,000 Paics |
| ENSMUSG00000024359 | 0,297 | 0,000 Hspa9 | ENSMUSG00000003810 | -0,243 | 0,017 Mast2 |
| ENSMUSG00000040028 | 0,296 | 0,000 Elavl1 | ENSMUSG00000016619 | -0,243 | 0,021 Nup50 |
| ENSMUSG00000038268 | 0,296 | 0,020 Ovca2 | ENSMUSG00000079316 | -0,243 | 0,032 Rab9 |
| ENSMUSG00000029048 | 0,296 | 0,018 Rer1 | ENSMUSG00000013833 | -0,244 | 0,002 Med16 |
| ENSMUSG00000021938 | 0,295 | 0,030 Pspc1 | ENSMUSG00000026404 | -0,244 | 0,046 Ddx59 |
| ENSMUSG00000018446 | 0,294 | 0,028 C1qbp | ENSMUSG00000025950 | -0,244 | 0,040 Idh1 |
| ENSMUSG00000001785 | 0,294 | 0,000 Pwp1 | ENSMUSG00000037278 | -0,244 | 0,045 Tmem97 |
| ENSMUSG00000021595 | 0,294 | 0,001 Nsun2 | ENSMUSG00000029781 | -0,245 | 0,008 Fkbp9 |
| ENSMUSG00000026701 | 0,294 | 0,016 Prdx6 | ENSMUSG00000039231 | -0,247 | 0,004 Suv39h1 |
| ENSMUSG00000031984 | 0,293 | 0,000 2810004N23Rik | ENSMUSG00000026816 | -0,249 | 0,015 Gtf3c5 |
| ENSMUSG00000005354 | 0,292 | 0,046 Txn2 | ENSMUSG00000049643 | -0,249 | 0,004 2310022A10Rik |
| ENSMUSG00000022615 | 0,292 | 0,045 Tymp | ENSMUSG00000037012 | -0,249 | 0,010 Hk1 |
| ENSMUSG00000059119 | 0,292 | 0,000 Nap1l4 | ENSMUSG00000036120 | -0,251 | 0,010 Rfxank |
| ENSMUSG00000029672 | 0,291 | 0,000 Fam3c | ENSMUSG00000079442 | -0,252 | 0,041 St6galnac4 |
| ENSMUSG00000000881 | 0,291 | 0,008 Dlg3 | ENSMUSG00000027010 | -0,253 | 0,015 Slc25a12 |
| ENSMUSG00000025016 | 0,291 | 0,048 Tm9sf3 | ENSMUSG00000022898 | -0,253 | 0,000 Dscr3 |
| ENSMUSG00000007891 | 0,290 | 0,048 Ctsd | ENSMUSG00000008140 | -0,253 | 0,036 Emc10 |
| ENSMUSG00000053091 | 0,289 | 0,003 Lins1 | ENSMUSG00000062376 | -0,256 | 0,001 Borcs7 |
| ENSMUSG00000022498 | 0,289 | 0,000 Txndc11 | ENSMUSG00000026174 | -0,258 | 0,002 Cnot9 |
| ENSMUSG00000002210 | 0,289 | 0,038 Smg9 | ENSMUSG00000028439 | -0,259 | 0,023 Fam219a |
| ENSMUSG00000027509 | 0,289 | 0,003 Rae1 | ENSMUSG00000018651 | -0,259 | 0,001 Tada2a |
| ENSMUSG00000018965 | 0,288 | 0,000 Ywhah | ENSMUSG00000022414 | -0,260 | 0,022 Tab1 |
| ENSMUSG00000020386 | 0,288 | 0,010 Sar1b | ENSMUSG00000020472 | -0,260 | 0,034 Zkscan17 |
| ENSMUSG00000052310 | 0,288 | 0,000 Slc39a1 | ENSMUSG00000018841 | -0,261 | 0,022 Rad51d |

| | | | | | |
|---|---|---|---|---|---|
| ENSMUSG00000009535 | 0,287 | 0,000 Rnmt | ENSMUSG00000003038 | -0,261 | 0,014 Hmgn2 |
| ENSMUSG00000068798 | 0,287 | 0,002 Rap1a | ENSMUSG00000031672 | -0,263 | 0,007 Got2 |
| ENSMUSG00000038299 | 0,287 | 0,000 Wdr36 | ENSMUSG00000041491 | -0,263 | 0,037 Cep78 |
| ENSMUSG00000029131 | 0,287 | 0,002 Dnajb6 | ENSMUSG00000028799 | -0,264 | 0,002 Zfp362 |
| ENSMUSG00000021615 | 0,286 | 0,046 Xrcc4 | ENSMUSG00000068551 | -0,266 | 0,028 Zfp467 |
| ENSMUSG00000031153 | 0,286 | 0,011 Gripap1 | ENSMUSG00000033628 | -0,268 | 0,002 Pik3c3 |
| ENSMUSG00000044847 | 0,286 | 0,027 Lsm11 | ENSMUSG00000016541 | -0,268 | 0,004 Atxn10 |
| ENSMUSG00000028932 | 0,286 | 0,047 Psmc2 | ENSMUSG00000026709 | -0,269 | 0,028 Dars2 |
| ENSMUSG00000047187 | 0,286 | 0,004 Rab2a | ENSMUSG00000058388 | -0,273 | 0,015 Phtf1 |
| ENSMUSG00000044030 | 0,286 | 0,013 Irf2bp1 | ENSMUSG00000021900 | -0,274 | 0,000 Btd |
| ENSMUSG00000011958 | 0,286 | 0,000 Bnip2 | ENSMUSG00000025465 | -0,274 | 0,006 Echs1 |
| ENSMUSG00000021037 | 0,286 | 0,027 Ahsa1 | ENSMUSG00000030614 | -0,275 | 0,033 Tmem126b |
| ENSMUSG00000014226 | 0,286 | 0,025 Cacybp | ENSMUSG00000074811 | -0,275 | 0,010 Hps6 |
| ENSMUSG00000019814 | 0,286 | 0,004 Ltv1 | ENSMUSG00000011752 | -0,275 | 0,010 Pgam1 |
| ENSMUSG00000030120 | 0,285 | 0,013 Mlf2 | ENSMUSG00000085793 | -0,276 | 0,015 Lin52 |
| ENSMUSG00000049327 | 0,285 | 0,000 Kmt5a | ENSMUSG00000001751 | -0,277 | 0,013 Naglu |
| ENSMUSG00000034620 | 0,284 | 0,003 Tmem5 | ENSMUSG00000016757 | -0,278 | 0,018 Ttll12 |
| ENSMUSG00000005103 | 0,284 | 0,000 Wdr1 | ENSMUSG00000008855 | -0,280 | 0,005 Hdac5 |
| ENSMUSG00000027502 | 0,284 | 0,019 Rtfdc1 | ENSMUSG00000001445 | -0,280 | 0,046 Mrpl10 |
| ENSMUSG00000031657 | 0,284 | 0,014 Heatr3 | ENSMUSG00000030872 | -0,281 | 0,021 Gga2 |
| ENSMUSG00000056820 | 0,283 | 0,000 Tsnax | ENSMUSG00000002297 | -0,282 | 0,029 Dbf4 |
| ENSMUSG00000026927 | 0,282 | 0,000 Sdccag3 | ENSMUSG00000000149 | -0,283 | 0,024 Gna12 |
| ENSMUSG00000029203 | 0,282 | 0,011 Ube2k | ENSMUSG00000031539 | -0,283 | 0,042 Ap3m2 |
| ENSMUSG00000031982 | 0,281 | 0,041 Arv1 | ENSMUSG00000040997 | -0,284 | 0,012 Abhd4 |
| ENSMUSG00000005204 | 0,281 | 0,000 Senp3 | ENSMUSG00000028973 | -0,284 | 0,000 Abcb8 |
| ENSMUSG00000023030 | 0,281 | 0,010 Slc11a2 | ENSMUSG00000039328 | -0,284 | 0,041 Rnf122 |
| ENSMUSG00000052419 | 0,281 | 0,007 2610001J05Rik | ENSMUSG00000036561 | -0,285 | 0,026 Ppp6r2 |
| ENSMUSG00000032307 | 0,280 | 0,014 Ube2q2 | ENSMUSG00000026003 | -0,285 | 0,003 Acadl |
| ENSMUSG00000032612 | 0,280 | 0,000 Usp4 | ENSMUSG00000021477 | -0,285 | 0,012 Ctsl |
| ENSMUSG00000031960 | 0,280 | 0,001 Aars | ENSMUSG00000033065 | -0,288 | 0,006 Pfkm |
| ENSMUSG00000036432 | 0,280 | 0,047 Siah2 | ENSMUSG00000028646 | -0,288 | 0,002 Rragc |
| ENSMUSG00000079477 | 0,279 | 0,000 Rab7 | ENSMUSG00000051007 | -0,288 | 0,029 Gatd1 |
| ENSMUSG00000114763 | 0,279 | 0,004 AC134869.3 | ENSMUSG00000057156 | -0,288 | 0,028 Homez |
| ENSMUSG00000032870 | 0,279 | 0,000 Smap2 | ENSMUSG00000032245 | -0,289 | 0,007 Cln6 |
| ENSMUSG00000034708 | 0,278 | 0,006 Grn | ENSMUSG00000028089 | -0,290 | 0,011 Chd1l |
| ENSMUSG00000036934 | 0,278 | 0,022 4921524J17Rik | ENSMUSG00000031684 | -0,291 | 0,019 Slc10a7 |
| ENSMUSG00000021392 | 0,278 | 0,031 Nol8 | ENSMUSG00000029547 | -0,291 | 0,024 Ints1 |
| ENSMUSG00000035297 | 0,278 | 0,002 Cops4 | ENSMUSG00000007950 | -0,291 | 0,047 Abhd8 |
| ENSMUSG00000022108 | 0,278 | 0,000 Itm2b | ENSMUSG00000034714 | -0,292 | 0,035 Ttyh2 |
| ENSMUSG00000031007 | 0,278 | 0,024 Atp6ap2 | ENSMUSG00000044573 | -0,292 | 0,004 Acp1 |
| ENSMUSG00000021876 | 0,278 | 0,034 Rnase4 | ENSMUSG00000038206 | -0,292 | 0,002 Fbxo8 |
| ENSMUSG00000074698 | 0,278 | 0,010 Csnk2a1 | ENSMUSG00000028484 | -0,293 | 0,001 Psip1 |
| ENSMUSG00000042148 | 0,277 | 0,007 Cox10 | ENSMUSG00000027227 | -0,293 | 0,021 Sord |
| ENSMUSG00000021087 | 0,277 | 0,031 Rtn1 | ENSMUSG00000002803 | -0,293 | 0,009 Btbd6 |
| ENSMUSG00000022663 | 0,277 | 0,001 Atg3 | ENSMUSG00000099519 | -0,293 | 0,000 Gm29253 |
| ENSMUSG00000106561 | 0,276 | 0,016 AC151967.1 | ENSMUSG00000036622 | -0,295 | 0,001 Atp13a2 |
| ENSMUSG00000034343 | 0,276 | 0,005 Ube2f | ENSMUSG00000019715 | -0,296 | 0,000 Gle1 |

| | | | | | |
|---|---|---|---|---|---|
| ENSMUSG00000025532 | 0,276 | 0,009 Crcp | ENSMUSG00000028164 | -0,296 | 0,011 Manba |
| ENSMUSG00000028849 | 0,275 | 0,002 Map7d1 | ENSMUSG00000074093 | -0,296 | 0,043 Svip |
| ENSMUSG00000041623 | 0,275 | 0,010 D11Wsu47e | ENSMUSG00000037622 | -0,297 | 0,015 Wdtc1 |
| ENSMUSG00000026234 | 0,274 | 0,000 Ncl | ENSMUSG00000028779 | -0,297 | 0,018 Pef1 |
| ENSMUSG00000025646 | 0,274 | 0,001 Atrip | ENSMUSG00000034487 | -0,298 | 0,003 Kdelc2 |
| ENSMUSG00000016257 | 0,274 | 0,004 Prelid3b | ENSMUSG00000024165 | -0,298 | 0,046 Jpt2 |
| ENSMUSG00000070730 | 0,274 | 0,000 Rmdn3 | ENSMUSG00000070425 | -0,298 | 0,047 Xntrpc |
| ENSMUSG00000008604 | 0,274 | 0,000 Ubqln4 | ENSMUSG00000024975 | -0,299 | 0,016 Pdcd4 |
| ENSMUSG00000052681 | 0,273 | 0,010 Rap1b | ENSMUSG00000057778 | -0,299 | 0,035 Cyb5d2 |
| ENSMUSG00000030890 | 0,273 | 0,007 Ilk | ENSMUSG00000060703 | -0,299 | 0,049 Cd302 |
| ENSMUSG00000022560 | 0,272 | 0,001 Slc52a2 | ENSMUSG00000020605 | -0,300 | 0,004 Hs1bp3 |
| ENSMUSG00000019792 | 0,272 | 0,032 Trmt11 | ENSMUSG00000037049 | -0,300 | 0,000 Smpd1 |
| ENSMUSG00000007415 | 0,272 | 0,002 Gatad1 | ENSMUSG00000049670 | -0,301 | 0,038 Morn4 |
| ENSMUSG00000031568 | 0,271 | 0,005 Rwdd4a | ENSMUSG00000024556 | -0,302 | 0,041 Me2 |
| ENSMUSG00000050088 | 0,270 | 0,001 1600012H06Rik | ENSMUSG00000029270 | -0,303 | 0,028 Fam69a |
| ENSMUSG00000022312 | 0,270 | 0,050 Eif3h | ENSMUSG00000000959 | -0,303 | 0,004 Oxa1l |
| ENSMUSG00000025616 | 0,269 | 0,020 Usp16 | ENSMUSG00000067369 | -0,304 | 0,008 Trmt2b |
| ENSMUSG00000078317 | 0,269 | 0,042 F8a | ENSMUSG00000041199 | -0,304 | 0,037 Rpusd1 |
| ENSMUSG00000025613 | 0,268 | 0,001 Cct8 | ENSMUSG00000021023 | -0,304 | 0,034 1110008L16Rik |
| ENSMUSG00000024896 | 0,268 | 0,001 Minpp1 | ENSMUSG00000043964 | -0,305 | 0,001 Orai3 |
| ENSMUSG00000001518 | 0,267 | 0,021 Itfg2 | ENSMUSG00000026198 | -0,306 | 0,013 Abcb6 |
| ENSMUSG00000024182 | 0,267 | 0,000 Axin1 | ENSMUSG00000069255 | -0,306 | 0,001 Dusp22 |
| ENSMUSG00000073982 | 0,266 | 0,049 Rhog | ENSMUSG00000030766 | -0,307 | 0,013 Arhgap17 |
| ENSMUSG00000029146 | 0,266 | 0,013 Snx17 | ENSMUSG00000034006 | -0,308 | 0,013 Pqlc1 |
| ENSMUSG00000003131 | 0,266 | 0,001 Pafah1b2 | ENSMUSG00000062590 | -0,309 | 0,030 Armc9 |
| ENSMUSG00000020225 | 0,265 | 0,040 Tmbim4 | ENSMUSG00000024588 | -0,310 | 0,000 Fech |
| ENSMUSG00000004069 | 0,264 | 0,000 Dnaja3 | ENSMUSG00000078517 | -0,311 | 0,028 Emc1 |
| ENSMUSG00000032423 | 0,264 | 0,036 Syncrip | ENSMUSG00000035790 | -0,311 | 0,048 Cep19 |
| ENSMUSG00000013089 | 0,264 | 0,041 Etv5 | ENSMUSG00000060376 | -0,311 | 0,029 Bckdha |
| ENSMUSG00000029777 | 0,264 | 0,033 Gars | ENSMUSG00000006920 | -0,312 | 0,020 Ezh1 |
| ENSMUSG00000026353 | 0,264 | 0,001 Ubxn4 | ENSMUSG00000030802 | -0,312 | 0,007 Bckdk |
| ENSMUSG00000027429 | 0,264 | 0,019 Sec23b | ENSMUSG00000027198 | -0,313 | 0,000 Ext2 |
| ENSMUSG00000030341 | 0,263 | 0,003 Tnfrsf1a | ENSMUSG00000002031 | -0,314 | 0,031 Ift46 |
| ENSMUSG00000070643 | 0,263 | 0,048 Sox13 | ENSMUSG00000022106 | -0,314 | 0,021 Rcbtb2 |
| ENSMUSG00000070934 | 0,263 | 0,018 Rraga | ENSMUSG00000041654 | -0,314 | 0,013 Slc39a11 |
| ENSMUSG00000028330 | 0,262 | 0,002 Ncbp1 | ENSMUSG00000029047 | -0,315 | 0,034 Pex10 |
| ENSMUSG00000049792 | 0,262 | 0,026 Bag5 | ENSMUSG00000053128 | -0,315 | 0,008 Rnf26 |
| ENSMUSG00000022500 | 0,262 | 0,018 Litaf | ENSMUSG00000022369 | -0,316 | 0,003 Mtbp |
| ENSMUSG00000035575 | 0,261 | 0,024 Utp6 | ENSMUSG00000003031 | -0,316 | 0,022 Cdkn1b |
| ENSMUSG00000010755 | 0,261 | 0,044 Cars | ENSMUSG00000071454 | -0,316 | 0,034 Dtnb |
| ENSMUSG00000022682 | 0,261 | 0,005 Rrn3 | ENSMUSG00000020973 | -0,316 | 0,009 Dnaaf2 |
| ENSMUSG00000054277 | 0,260 | 0,035 Arfgap3 | ENSMUSG00000063052 | -0,317 | 0,005 Lrrc40 |
| ENSMUSG00000003234 | 0,260 | 0,005 Abcf3 | ENSMUSG00000028057 | -0,317 | 0,013 Rit1 |
| ENSMUSG00000061130 | 0,260 | 0,037 Ppm1b | ENSMUSG00000034875 | -0,318 | 0,019 Nudt19 |
| ENSMUSG00000020923 | 0,259 | 0,000 Ubtf | ENSMUSG00000034341 | -0,319 | 0,000 Wbp2 |
| ENSMUSG00000034252 | 0,259 | 0,001 Senp6 | ENSMUSG00000041625 | -0,320 | 0,048 Ggact |
| ENSMUSG00000031950 | 0,259 | 0,048 Gabarapl2 | ENSMUSG00000050989 | -0,320 | 0,003 Selenon |

| | | | | | |
|---|---|---|---|---|---|
| ENSMUSG00000033161 | 0,259 | 0,002 Atp1a1 | ENSMUSG00000029428 | -0,321 | 0,000 Stx2 |
| ENSMUSG00000038976 | 0,259 | 0,007 Ppp1r9b | ENSMUSG00000001942 | -0,322 | 0,046 Siae |
| ENSMUSG00000036002 | 0,259 | 0,006 Fam214b | ENSMUSG00000046949 | -0,323 | 0,002 Nqo2 |
| ENSMUSG00000001569 | 0,258 | 0,003 Nom1 | ENSMUSG00000029676 | -0,324 | 0,002 Pot1a |
| ENSMUSG00000034211 | 0,258 | 0,012 Mrps17 | ENSMUSG00000021179 | -0,324 | 0,033 Nrde2 |
| ENSMUSG00000068039 | 0,258 | 0,009 Tcp1 | ENSMUSG00000032959 | -0,324 | 0,046 Pebp1 |
| ENSMUSG00000014601 | 0,257 | 0,000 Strip1 | ENSMUSG00000020652 | -0,325 | 0,008 Cenpo |
| ENSMUSG00000025373 | 0,257 | 0,028 Rnf41 | ENSMUSG00000056204 | -0,325 | 0,001 Pgpep1 |
| ENSMUSG00000061950 | 0,257 | 0,039 Ppp4r1 | ENSMUSG00000037461 | -0,325 | 0,017 Ints7 |
| ENSMUSG00000032577 | 0,257 | 0,039 Mapkapk3 | ENSMUSG00000040913 | -0,325 | 0,034 Fbxw4 |
| ENSMUSG00000023110 | 0,256 | 0,002 Prmt5 | ENSMUSG00000028096 | -0,327 | 0,007 Gpr89 |
| ENSMUSG00000032020 | 0,256 | 0,041 Ubash3b | ENSMUSG00000031776 | -0,328 | 0,004 Arl2bp |
| ENSMUSG00000074088 | 0,256 | 0,023 Snrnp40 | ENSMUSG00000028621 | -0,328 | 0,004 Cyb5rl |
| ENSMUSG00000024404 | 0,256 | 0,021 Riok3 | ENSMUSG00000029434 | -0,329 | 0,039 Vps33a |
| ENSMUSG00000026889 | 0,256 | 0,001 Rbm18 | ENSMUSG00000037257 | -0,330 | 0,005 Aagab |
| ENSMUSG00000067713 | 0,256 | 0,040 Prkag1 | ENSMUSG00000047766 | -0,330 | 0,003 Lrrc49 |
| ENSMUSG00000023075 | 0,256 | 0,002 Akirin1 | ENSMUSG00000022553 | -0,330 | 0,004 Maf1 |
| ENSMUSG00000026289 | 0,254 | 0,008 Atg16l1 | ENSMUSG00000021481 | -0,331 | 0,014 Zfp346 |
| ENSMUSG00000024683 | 0,254 | 0,033 Mrpl16 | ENSMUSG00000053436 | -0,332 | 0,000 Mapk14 |
| ENSMUSG00000045795 | 0,254 | 0,004 Whamm | ENSMUSG00000009828 | -0,333 | 0,001 Ick |
| ENSMUSG00000021621 | 0,253 | 0,005 Zcchc9 | ENSMUSG00000035722 | -0,333 | 0,014 Abca7 |
| ENSMUSG00000020720 | 0,252 | 0,029 Psmd12 | ENSMUSG00000032594 | -0,333 | 0,006 Ip6k1 |
| ENSMUSG00000033285 | 0,252 | 0,020 Wdr3 | ENSMUSG00000031198 | -0,334 | 0,000 Fundc2 |
| ENSMUSG00000044600 | 0,251 | 0,022 Smim7 | ENSMUSG00000079109 | -0,335 | 0,000 Pms2 |
| ENSMUSG00000020132 | 0,251 | 0,011 Rab21 | ENSMUSG00000031835 | -0,336 | 0,001 Mbtps1 |
| ENSMUSG00000014349 | 0,250 | 0,000 Ube2z | ENSMUSG00000027180 | -0,336 | 0,026 Fbxo3 |
| ENSMUSG00000044857 | 0,250 | 0,020 Lemd2 | ENSMUSG00000022125 | -0,336 | 0,016 Cln5 |
| ENSMUSG00000038982 | 0,250 | 0,019 Bloc1s5 | ENSMUSG00000040272 | -0,338 | 0,021 Accs |
| ENSMUSG00000021814 | 0,250 | 0,033 Anxa7 | ENSMUSG00000020823 | -0,339 | 0,000 Sec14l1 |
| ENSMUSG00000023044 | 0,249 | 0,002 Csad | ENSMUSG00000027963 | -0,340 | 0,006 Extl2 |
| ENSMUSG00000030619 | 0,248 | 0,038 Eed | ENSMUSG00000037656 | -0,340 | 0,018 Slc20a2 |
| ENSMUSG00000015776 | 0,248 | 0,005 Med22 | ENSMUSG00000039958 | -0,341 | 0,018 Etfbkmt |
| ENSMUSG00000027852 | 0,248 | 0,001 Nras | ENSMUSG00000099481 | -0,342 | 0,003 Xndc1 |
| ENSMUSG00000047731 | 0,247 | 0,000 Wbp1l | ENSMUSG00000037415 | -0,343 | 0,002 Ranbp10 |
| ENSMUSG00000026095 | 0,246 | 0,005 Asnsd1 | ENSMUSG00000006850 | -0,343 | 0,014 Tmco6 |
| ENSMUSG00000028961 | 0,246 | 0,005 Pgd | ENSMUSG00000026179 | -0,344 | 0,001 Pnkd |
| ENSMUSG00000041426 | 0,246 | 0,029 Hibch | ENSMUSG00000048232 | -0,346 | 0,021 Fbxo10 |
| ENSMUSG00000025872 | 0,245 | 0,048 Thoc3 | ENSMUSG00000000751 | -0,347 | 0,000 Rpa1 |
| ENSMUSG00000026279 | 0,245 | 0,039 Thap4 | ENSMUSG00000006728 | -0,347 | 0,008 Cdk4 |
| ENSMUSG00000040463 | 0,245 | 0,013 Mybbp1a | ENSMUSG00000031546 | -0,347 | 0,011 Gins4 |
| ENSMUSG00000045576 | 0,245 | 0,029 St7l | ENSMUSG00000081534 | -0,348 | 0,000 Slc48a1 |
| ENSMUSG00000028161 | 0,245 | 0,001 Ppp3ca | ENSMUSG00000026134 | -0,349 | 0,002 Prim2 |
| ENSMUSG00000047832 | 0,244 | 0,004 Cdca4 | ENSMUSG00000029186 | -0,349 | 0,007 Pi4k2b |
| ENSMUSG00000063785 | 0,244 | 0,005 Utp14a | ENSMUSG00000049657 | -0,353 | 0,022 Zbtb5 |
| ENSMUSG00000003348 | 0,244 | 0,013 Mob3a | ENSMUSG00000053134 | -0,353 | 0,004 Supt7l |
| ENSMUSG00000056941 | 0,243 | 0,039 Commd7 | ENSMUSG00000033983 | -0,355 | 0,025 Coil |
| ENSMUSG00000030168 | 0,243 | 0,011 Adipor2 | ENSMUSG00000029191 | -0,356 | 0,001 Rfc1 |

| | | | | | | | |
|---|---|---|---|---|---|---|---|
| ENSMUSG00000015714 | 0,243 | 0,000 | Cers2 | ENSMUSG00000020534 | -0,356 | 0,046 | Shmt1 |
| ENSMUSG00000006390 | 0,242 | 0,012 | Elovl1 | ENSMUSG00000025486 | -0,356 | 0,037 | Sirt3 |
| ENSMUSG00000038697 | 0,242 | 0,017 | Taf5l | ENSMUSG00000021975 | -0,357 | 0,000 | Ints9 |
| ENSMUSG00000044147 | 0,242 | 0,003 | Arf6 | ENSMUSG00000024158 | -0,357 | 0,029 | Hagh |
| ENSMUSG00000024219 | 0,241 | 0,047 | Anks1 | ENSMUSG00000043067 | -0,358 | 0,043 | Dpy19l1 |
| ENSMUSG00000025024 | 0,241 | 0,005 | Smndc1 | ENSMUSG00000043065 | -0,358 | 0,014 | Spice1 |
| ENSMUSG00000008475 | 0,241 | 0,025 | Arpc5 | ENSMUSG00000068115 | -0,359 | 0,007 | Ninl |
| ENSMUSG00000026864 | 0,241 | 0,029 | Hspa5 | ENSMUSG00000041000 | -0,362 | 0,006 | Trim62 |
| ENSMUSG00000028420 | 0,240 | 0,031 | Tmem38b | ENSMUSG00000009905 | -0,362 | 0,000 | Kdsr |
| ENSMUSG00000022378 | 0,240 | 0,015 | Fam49b | ENSMUSG00000022978 | -0,363 | 0,009 | Mis18a |
| ENSMUSG00000048439 | 0,239 | 0,049 | Nupl2 | ENSMUSG00000038860 | -0,363 | 0,009 | Garnl3 |
| ENSMUSG00000019777 | 0,239 | 0,050 | Hdac2 | ENSMUSG00000022420 | -0,364 | 0,002 | Dnal4 |
| ENSMUSG00000020612 | 0,238 | 0,002 | Prkar1a | ENSMUSG00000025648 | -0,365 | 0,011 | Pfkfb4 |
| ENSMUSG00000025980 | 0,238 | 0,014 | Hspd1 | ENSMUSG00000037419 | -0,365 | 0,041 | Endod1 |
| ENSMUSG00000004897 | 0,238 | 0,005 | Hdgf | ENSMUSG00000035266 | -0,365 | 0,002 | Helq |
| ENSMUSG00000028719 | 0,238 | 0,045 | Cmpk1 | ENSMUSG00000058997 | -0,365 | 0,027 | Vwa8 |
| ENSMUSG00000031467 | 0,238 | 0,010 | Agpat5 | ENSMUSG00000002814 | -0,365 | 0,025 | Top3a |
| ENSMUSG00000021385 | 0,237 | 0,049 | Ippk | ENSMUSG00000021483 | -0,366 | 0,003 | Cdk20 |
| ENSMUSG00000042426 | 0,237 | 0,038 | Dhx29 | ENSMUSG00000041346 | -0,366 | 0,012 | Wrap53 |
| ENSMUSG00000028165 | 0,237 | 0,004 | Cisd2 | ENSMUSG00000050530 | -0,367 | 0,040 | Fam171a1 |
| ENSMUSG00000012422 | 0,236 | 0,044 | Tmem167 | ENSMUSG00000019813 | -0,367 | 0,010 | Cep57l1 |
| ENSMUSG00000035152 | 0,236 | 0,011 | Ap2b1 | ENSMUSG00000040618 | -0,367 | 0,043 | Pck2 |
| ENSMUSG00000047123 | 0,235 | 0,047 | Ticam1 | ENSMUSG00000025151 | -0,367 | 0,000 | Maged1 |
| ENSMUSG00000021610 | 0,234 | 0,000 | Clptm1l | ENSMUSG00000035877 | -0,368 | 0,010 | Zhx3 |
| ENSMUSG00000024668 | 0,234 | 0,045 | Sdhaf2 | ENSMUSG00000036882 | -0,369 | 0,035 | Arhgap33 |
| ENSMUSG00000029534 | 0,234 | 0,011 | St7 | ENSMUSG00000033970 | -0,369 | 0,013 | Rfc3 |
| ENSMUSG00000021266 | 0,234 | 0,001 | Wars | ENSMUSG00000019338 | -0,369 | 0,047 | Zfp687 |
| ENSMUSG00000000561 | 0,233 | 0,012 | Wdr77 | ENSMUSG00000055200 | -0,370 | 0,007 | Sertad3 |
| ENSMUSG00000031715 | 0,233 | 0,045 | Smarca5 | ENSMUSG00000022244 | -0,370 | 0,007 | Amacr |
| ENSMUSG00000030682 | 0,232 | 0,011 | Cdipt | ENSMUSG00000062661 | -0,370 | 0,022 | Ncs1 |
| ENSMUSG00000051934 | 0,232 | 0,000 | Spats2 | ENSMUSG00000038637 | -0,372 | 0,018 | Lrrc56 |
| ENSMUSG00000025156 | 0,232 | 0,001 | Gps1 | ENSMUSG00000021120 | -0,372 | 0,007 | Pigh |
| ENSMUSG00000057522 | 0,231 | 0,030 | Spop | ENSMUSG00000056698 | -0,372 | 0,002 | Elmod3 |
| ENSMUSG00000022556 | 0,231 | 0,025 | Hsf1 | ENSMUSG00000033059 | -0,373 | 0,001 | Pygb |
| ENSMUSG00000028833 | 0,231 | 0,007 | Ncdn | ENSMUSG00000038368 | -0,373 | 0,013 | Focad |
| ENSMUSG00000022052 | 0,231 | 0,032 | Ppp2r2a | ENSMUSG00000007338 | -0,373 | 0,014 | Mrpl49 |
| ENSMUSG00000024535 | 0,230 | 0,018 | Snx24 | ENSMUSG00000052949 | -0,373 | 0,012 | Rnf157 |
| ENSMUSG00000022817 | 0,230 | 0,032 | Itgb5 | ENSMUSG00000059040 | -0,374 | 0,041 | Eno1b |
| ENSMUSG00000031532 | 0,230 | 0,038 | Saraf | ENSMUSG00000068264 | -0,374 | 0,007 | Ap5s1 |
| ENSMUSG00000031913 | 0,229 | 0,001 | Vps4a | ENSMUSG00000029687 | -0,375 | 0,000 | Ezh2 |
| ENSMUSG00000017760 | 0,229 | 0,031 | Ctsa | ENSMUSG00000035401 | -0,376 | 0,001 | Emsy |
| ENSMUSG00000038459 | 0,229 | 0,005 | Abhd17c | ENSMUSG00000018548 | -0,376 | 0,029 | Trim37 |
| ENSMUSG00000098923 | 0,229 | 0,027 | Tmem185b | ENSMUSG00000040312 | -0,376 | 0,041 | Cchcr1 |
| ENSMUSG00000037935 | 0,229 | 0,012 | Smarce1 | ENSMUSG00000026421 | -0,377 | 0,035 | Csrp1 |
| ENSMUSG00000071652 | 0,229 | 0,007 | Ints5 | ENSMUSG00000025235 | -0,378 | 0,001 | Bbs4 |
| ENSMUSG00000010554 | 0,228 | 0,017 | Mettl16 | ENSMUSG00000021252 | -0,378 | 0,000 | Erg28 |
| ENSMUSG00000063888 | 0,228 | 0,011 | Rpl7l1 | ENSMUSG00000087174 | -0,378 | 0,009 | 5530601H04Rik |

| | | | | | | | |
|---|---|---|---|---|---|---|---|
| ENSMUSG00000040782 | 0,228 | 0,024 | Rfwd2 | ENSMUSG00000036957 | -0,379 | 0,004 | Lrfn3 |
| ENSMUSG00000028879 | 0,228 | 0,013 | Stx12 | ENSMUSG00000039914 | -0,380 | 0,034 | Coq10a |
| ENSMUSG00000019173 | 0,228 | 0,000 | Rab5c | ENSMUSG00000039936 | -0,380 | 0,027 | Pik3cd |
| ENSMUSG00000021096 | 0,227 | 0,000 | Ppm1a | ENSMUSG00000031787 | -0,380 | 0,000 | Katnb1 |
| ENSMUSG00000025364 | 0,226 | 0,035 | Pa2g4 | ENSMUSG00000034522 | -0,381 | 0,038 | Zfp395 |
| ENSMUSG00000035517 | 0,226 | 0,046 | Tdrd7 | ENSMUSG00000033124 | -0,381 | 0,000 | Atg9a |
| ENSMUSG00000022855 | 0,226 | 0,009 | Senp2 | ENSMUSG00000039747 | -0,382 | 0,046 | Orai2 |
| ENSMUSG00000029364 | 0,225 | 0,008 | Wsb2 | ENSMUSG00000000693 | -0,382 | 0,004 | Loxl3 |
| ENSMUSG00000056305 | 0,224 | 0,018 | Usp39 | ENSMUSG00000039382 | -0,382 | 0,007 | Wdr45 |
| ENSMUSG00000062397 | 0,224 | 0,023 | Zfp706 | ENSMUSG00000022221 | -0,383 | 0,004 | Ripk3 |
| ENSMUSG00000026926 | 0,224 | 0,000 | Pmpca | ENSMUSG00000032192 | -0,384 | 0,006 | Gnb5 |
| ENSMUSG00000021694 | 0,223 | 0,035 | Ercc8 | ENSMUSG00000000378 | -0,384 | 0,005 | Ccm2 |
| ENSMUSG00000016528 | 0,223 | 0,029 | Mapkapk2 | ENSMUSG00000035891 | -0,386 | 0,004 | Cerk |
| ENSMUSG00000046707 | 0,223 | 0,000 | Csnk2a2 | ENSMUSG00000000876 | -0,388 | 0,000 | Pxmp4 |
| ENSMUSG00000011257 | 0,223 | 0,001 | Pabpc4 | ENSMUSG00000024773 | -0,388 | 0,014 | Atg2a |
| ENSMUSG00000029518 | 0,223 | 0,000 | Rab35 | ENSMUSG00000039050 | -0,388 | 0,000 | Osbpl2 |
| ENSMUSG00000024527 | 0,223 | 0,000 | Afg3l2 | ENSMUSG00000037526 | -0,389 | 0,019 | Atg14 |
| ENSMUSG00000042079 | 0,222 | 0,001 | Hnrnpf | ENSMUSG00000024590 | -0,389 | 0,001 | Lmnb1 |
| ENSMUSG00000079562 | 0,222 | 0,000 | Maea | ENSMUSG00000044629 | -0,389 | 0,000 | Cnrip1 |
| ENSMUSG00000039682 | 0,222 | 0,001 | Lap3 | ENSMUSG00000033126 | -0,390 | 0,045 | Ybey |
| ENSMUSG00000011179 | 0,222 | 0,031 | Odc1 | ENSMUSG00000025190 | -0,391 | 0,000 | Got1 |
| ENSMUSG00000003062 | 0,221 | 0,045 | Stard3nl | ENSMUSG00000027244 | -0,391 | 0,000 | Atg13 |
| ENSMUSG00000039756 | 0,221 | 0,004 | Dnttip2 | ENSMUSG00000027342 | -0,391 | 0,000 | Pcna |
| ENSMUSG00000023353 | 0,221 | 0,025 | Agap3 | ENSMUSG00000021286 | -0,392 | 0,002 | Zfyve21 |
| ENSMUSG00000014867 | 0,221 | 0,000 | Surf4 | ENSMUSG00000035776 | -0,392 | 0,000 | Cd99l2 |
| ENSMUSG00000023010 | 0,220 | 0,000 | Tmbim6 | ENSMUSG00000050144 | -0,393 | 0,003 | Slc25a44 |
| ENSMUSG00000027170 | 0,220 | 0,041 | Eif3m | ENSMUSG00000039693 | -0,394 | 0,027 | Msantd3 |
| ENSMUSG00000028409 | 0,220 | 0,033 | Smu1 | ENSMUSG00000058486 | -0,396 | 0,000 | Wdr91 |
| ENSMUSG00000058704 | 0,219 | 0,014 | Memo1 | ENSMUSG00000025791 | -0,398 | 0,000 | Pgm2 |
| ENSMUSG00000021428 | 0,218 | 0,002 | Riok1 | ENSMUSG00000001270 | -0,398 | 0,027 | Ckb |
| ENSMUSG00000028343 | 0,218 | 0,011 | Erp44 | ENSMUSG00000021594 | -0,400 | 0,007 | Srd5a1 |
| ENSMUSG00000038416 | 0,218 | 0,021 | Cdc16 | ENSMUSG00000013629 | -0,401 | 0,020 | Cad |
| ENSMUSG00000022472 | 0,218 | 0,002 | Desi1 | ENSMUSG00000029998 | -0,401 | 0,015 | Pcyox1 |
| ENSMUSG00000059439 | 0,218 | 0,010 | Bcas3 | ENSMUSG00000020766 | -0,402 | 0,040 | Galk1 |
| ENSMUSG00000036371 | 0,217 | 0,003 | Serbp1 | ENSMUSG00000022885 | -0,402 | 0,003 | St6gal1 |
| ENSMUSG00000029528 | 0,217 | 0,020 | Pxn | ENSMUSG00000028675 | -0,403 | 0,000 | Pnrc2 |
| ENSMUSG00000066148 | 0,217 | 0,037 | Prpf4 | ENSMUSG00000004085 | -0,403 | 0,046 | Map3k20 |
| ENSMUSG00000034889 | 0,216 | 0,025 | Cactin | ENSMUSG00000042476 | -0,404 | 0,008 | Abcb4 |
| ENSMUSG00000062937 | 0,216 | 0,050 | Mtap | ENSMUSG00000030530 | -0,404 | 0,031 | Furin |
| ENSMUSG00000033706 | 0,216 | 0,025 | Smyd5 | ENSMUSG00000021185 | -0,404 | 0,011 | Dglucy |
| ENSMUSG00000001687 | 0,216 | 0,027 | Ubl3 | ENSMUSG00000028199 | -0,406 | 0,004 | Cryz |
| ENSMUSG00000027804 | 0,216 | 0,034 | Ppid | ENSMUSG00000029521 | -0,407 | 0,005 | Chek2 |
| ENSMUSG00000031916 | 0,215 | 0,006 | Cog8 | ENSMUSG00000032652 | -0,408 | 0,015 | Crebl2 |
| ENSMUSG00000033793 | 0,215 | 0,012 | Atp6v1h | ENSMUSG00000032086 | -0,408 | 0,047 | Bace1 |
| ENSMUSG00000069520 | 0,215 | 0,000 | Tmem19 | ENSMUSG00000010307 | -0,409 | 0,006 | Tmem86a |
| ENSMUSG00000009291 | 0,215 | 0,041 | Pttg1ip | ENSMUSG00000026360 | -0,409 | 0,044 | Rgs2 |
| ENSMUSG00000027361 | 0,215 | 0,005 | Gabpb1 | ENSMUSG00000022773 | -0,411 | 0,046 | Ypel1 |

| | | | | | |
|---|---|---|---|---|---|
| ENSMUSG00000022175 | 0,215 | 0,001 Lrp10 | ENSMUSG00000071547 | -0,412 | 0,009 Nt5dc2 |
| ENSMUSG00000026254 | 0,215 | 0,034 Eif4e2 | ENSMUSG00000066357 | -0,413 | 0,000 Wdr6 |
| ENSMUSG00000003518 | 0,214 | 0,028 Dusp3 | ENSMUSG00000041161 | -0,413 | 0,011 Otud3 |
| ENSMUSG00000028745 | 0,214 | 0,036 Capzb | ENSMUSG00000023345 | -0,413 | 0,006 Poc1a |
| ENSMUSG00000000827 | 0,214 | 0,006 Tpd52l2 | ENSMUSG00000000278 | -0,415 | 0,000 Scpep1 |
| ENSMUSG00000031931 | 0,213 | 0,031 Ankrd49 | ENSMUSG00000038520 | -0,415 | 0,017 Tbc1d17 |
| ENSMUSG00000037013 | 0,213 | 0,048 Ss18 | ENSMUSG00000024833 | -0,415 | 0,000 Pola2 |
| ENSMUSG00000028538 | 0,213 | 0,049 St3gal3 | ENSMUSG00000022428 | -0,415 | 0,016 Cby1 |
| ENSMUSG00000038848 | 0,212 | 0,018 Ythdf1 | ENSMUSG00000038776 | -0,415 | 0,015 Ephx1 |
| ENSMUSG00000038503 | 0,212 | 0,001 Mesd | ENSMUSG00000109493 | -0,416 | 0,000 Gm45208 |
| ENSMUSG00000029291 | 0,212 | 0,000 Rufy3 | ENSMUSG00000003546 | -0,417 | 0,000 Klc4 |
| ENSMUSG00000024927 | 0,211 | 0,033 Rela | ENSMUSG00000048200 | -0,418 | 0,043 Cracr2b |
| ENSMUSG00000024816 | 0,211 | 0,023 Frmd8 | ENSMUSG00000025289 | -0,422 | 0,027 Prdx4 |
| ENSMUSG00000031575 | 0,211 | 0,000 Ash2l | ENSMUSG00000050394 | -0,422 | 0,020 Armcx6 |
| ENSMUSG00000026878 | 0,211 | 0,023 Rab14 | ENSMUSG00000063439 | -0,422 | 0,029 B9d2 |
| ENSMUSG00000068267 | 0,211 | 0,034 Cenpb | ENSMUSG00000029463 | -0,422 | 0,015 Fam216a |
| ENSMUSG00000057858 | 0,210 | 0,029 Fam204a | ENSMUSG00000031134 | -0,423 | 0,000 Rbmx |
| ENSMUSG00000000346 | 0,210 | 0,001 Dazap2 | ENSMUSG00000022679 | -0,424 | 0,039 Mpv17l |
| ENSMUSG00000019795 | 0,209 | 0,003 Pcmt1 | ENSMUSG00000027163 | -0,424 | 0,015 Commd9 |
| ENSMUSG00000025609 | 0,209 | 0,049 Mkln1 | ENSMUSG00000074738 | -0,425 | 0,002 Fndc10 |
| ENSMUSG00000003037 | 0,209 | 0,048 Rab8a | ENSMUSG00000027984 | -0,425 | 0,000 Hadh |
| ENSMUSG00000005803 | 0,208 | 0,017 Sqor | ENSMUSG00000020105 | -0,426 | 0,030 Lrig3 |
| ENSMUSG00000040462 | 0,207 | 0,013 Os9 | ENSMUSG00000021811 | -0,428 | 0,002 Dnajc9 |
| ENSMUSG00000002741 | 0,207 | 0,000 Ykt6 | ENSMUSG00000039787 | -0,429 | 0,003 Cercam |
| ENSMUSG00000061559 | 0,207 | 0,049 Wdr61 | ENSMUSG00000040675 | -0,430 | 0,024 Mthfd1l |
| ENSMUSG00000020780 | 0,207 | 0,007 Srp68 | ENSMUSG00000026556 | -0,430 | 0,047 Vangl2 |
| ENSMUSG00000033335 | 0,207 | 0,005 Dnm2 | ENSMUSG00000035232 | -0,430 | 0,000 Pdk3 |
| ENSMUSG00000022285 | 0,206 | 0,048 Ywhaz | ENSMUSG00000042606 | -0,431 | 0,005 Hirip3 |
| ENSMUSG00000025510 | 0,206 | 0,018 Cd151 | ENSMUSG00000049086 | -0,431 | 0,017 Bmyc |
| ENSMUSG00000020680 | 0,206 | 0,014 Taf15 | ENSMUSG00000039686 | -0,432 | 0,000 Zer1 |
| ENSMUSG00000020076 | 0,206 | 0,002 Ddx50 | ENSMUSG00000041840 | -0,432 | 0,049 Haus1 |
| ENSMUSG00000046691 | 0,205 | 0,025 Chtf8 | ENSMUSG00000043419 | -0,432 | 0,048 Chd3os |
| ENSMUSG00000020149 | 0,204 | 0,048 Rab1a | ENSMUSG00000047067 | -0,433 | 0,046 Dusp28 |
| ENSMUSG00000015341 | 0,204 | 0,017 Golga7 | ENSMUSG00000002908 | -0,434 | 0,037 Kcnn1 |
| ENSMUSG00000027374 | 0,204 | 0,015 Mrps5 | ENSMUSG00000023079 | -0,435 | 0,004 Gtf2ird1 |
| ENSMUSG00000063334 | 0,204 | 0,001 Krr1 | ENSMUSG00000035885 | -0,436 | 0,041 Cox8a |
| ENSMUSG00000030876 | 0,203 | 0,023 Mettl9 | ENSMUSG00000034317 | -0,436 | 0,026 Trim59 |
| ENSMUSG00000031987 | 0,202 | 0,000 Egln1 | ENSMUSG00000036880 | -0,436 | 0,001 Acaa2 |
| ENSMUSG00000029110 | 0,201 | 0,019 Rnf4 | ENSMUSG00000039197 | -0,436 | 0,000 Adk |
| ENSMUSG00000064145 | 0,201 | 0,008 Arih2 | ENSMUSG00000024018 | -0,437 | 0,004 Ccdc167 |
| ENSMUSG00000031660 | 0,201 | 0,001 Brd7 | ENSMUSG00000040557 | -0,437 | 0,004 Mettl27 |
| ENSMUSG00000090213 | 0,201 | 0,001 Tmem189 | ENSMUSG00000048924 | -0,437 | 0,032 Ccdc125 |
| ENSMUSG00000001847 | 0,200 | 0,001 Rac1 | ENSMUSG00000029163 | -0,438 | 0,018 Emilin1 |
| ENSMUSG00000025793 | 0,200 | 0,003 Hgs | ENSMUSG00000035172 | -0,439 | 0,000 Plekhh3 |
| ENSMUSG00000034254 | 0,200 | 0,016 Agpat1 | ENSMUSG00000030161 | -0,440 | 0,000 Gabarapl1 |
| ENSMUSG00000077450 | 0,199 | 0,013 Rab11b | ENSMUSG00000057193 | -0,441 | 0,005 Slc44a2 |
| ENSMUSG00000005299 | 0,198 | 0,001 Letm1 | ENSMUSG00000052911 | -0,441 | 0,000 Lamb2 |

| | | | | | |
|---|---|---|---|---|---|
| ENSMUSG00000001175 | 0,197 | 0,020 Calm1 | ENSMUSG00000028394 | -0,441 | 0,000 Pole3 |
| ENSMUSG00000026356 | 0,197 | 0,016 Dars | ENSMUSG00000024151 | -0,442 | 0,000 Msh2 |
| ENSMUSG00000035086 | 0,197 | 0,006 Becn1 | ENSMUSG00000047409 | -0,443 | 0,000 Ctdspl |
| ENSMUSG00000068856 | 0,196 | 0,018 Sf3b4 | ENSMUSG00000036854 | -0,444 | 0,004 Hspb6 |
| ENSMUSG00000027634 | 0,196 | 0,050 Ndrg3 | ENSMUSG00000024118 | -0,446 | 0,000 Tedc2 |
| ENSMUSG00000015290 | 0,196 | 0,002 Ubl4a | ENSMUSG00000090115 | -0,446 | 0,031 Usp49 |
| ENSMUSG00000038563 | 0,196 | 0,015 Efl1 | ENSMUSG00000057229 | -0,447 | 0,002 Dmac2 |
| ENSMUSG00000034850 | 0,196 | 0,030 Tmem127 | ENSMUSG00000040746 | -0,447 | 0,003 Rnf167 |
| ENSMUSG00000020349 | 0,195 | 0,007 Ppp2ca | ENSMUSG00000032411 | -0,447 | 0,007 Tfdp2 |
| ENSMUSG00000030082 | 0,195 | 0,001 Sec61a1 | ENSMUSG00000047205 | -0,449 | 0,043 Dusp18 |
| ENSMUSG00000006699 | 0,194 | 0,004 Cdc42 | ENSMUSG00000028933 | -0,449 | 0,001 Xrcc2 |
| ENSMUSG00000034274 | 0,194 | 0,018 Thoc5 | ENSMUSG00000023947 | -0,450 | 0,031 Nfkbie |
| ENSMUSG00000032745 | 0,194 | 0,028 Gpbp1 | ENSMUSG00000037894 | -0,453 | 0,001 H2afz |
| ENSMUSG00000035596 | 0,193 | 0,024 Mboat7 | ENSMUSG00000041132 | -0,454 | 0,000 N4bp2l1 |
| ENSMUSG00000023944 | 0,193 | 0,006 Hsp90ab1 | ENSMUSG00000028044 | -0,454 | 0,028 Cks1b |
| ENSMUSG00000006418 | 0,193 | 0,012 Rnf114 | ENSMUSG00000029863 | -0,455 | 0,020 Casp2 |
| ENSMUSG00000035772 | 0,193 | 0,022 Mrps2 | ENSMUSG00000041930 | -0,455 | 0,036 Fam222a |
| ENSMUSG00000032123 | 0,193 | 0,018 Dpagt1 | ENSMUSG00000076437 | -0,456 | 0,045 Selenoh |
| ENSMUSG00000047098 | 0,192 | 0,041 Rnf31 | ENSMUSG00000021306 | -0,456 | 0,006 Gpr137b |
| ENSMUSG00000037376 | 0,190 | 0,044 Trmt6 | ENSMUSG00000001138 | -0,456 | 0,000 Cnnm3 |
| ENSMUSG00000023883 | 0,190 | 0,011 Phf10 | ENSMUSG00000037455 | -0,457 | 0,005 Slc18b1 |
| ENSMUSG00000004846 | 0,190 | 0,040 Plod3 | ENSMUSG00000021572 | -0,458 | 0,004 Cep72 |
| ENSMUSG00000025162 | 0,190 | 0,011 Csnk1d | ENSMUSG00000022844 | -0,458 | 0,009 Pdia5 |
| ENSMUSG00000028191 | 0,189 | 0,049 Bcl10 | ENSMUSG00000057137 | -0,458 | 0,033 Tmem140 |
| ENSMUSG00000030397 | 0,189 | 0,020 Mark4 | ENSMUSG00000030522 | -0,459 | 0,003 Mtmr10 |
| ENSMUSG00000036275 | 0,188 | 0,001 9530068E07Rik | ENSMUSG00000021226 | -0,459 | 0,001 Acot2 |
| ENSMUSG00000021504 | 0,188 | 0,033 B4galt7 | ENSMUSG00000037313 | -0,459 | 0,021 Tacc3 |
| ENSMUSG00000042901 | 0,187 | 0,028 Aida | ENSMUSG00000021009 | -0,459 | 0,045 Ptpn21 |
| ENSMUSG00000031545 | 0,187 | 0,007 Gpat4 | ENSMUSG00000029455 | -0,459 | 0,001 Aldh2 |
| ENSMUSG00000025027 | 0,185 | 0,017 Xpnpep1 | ENSMUSG00000034349 | -0,459 | 0,008 Smc4 |
| ENSMUSG00000099913 | 0,185 | 0,003 Gm28551 | ENSMUSG00000019124 | -0,460 | 0,005 Scrn1 |
| ENSMUSG00000027879 | 0,185 | 0,000 Sec22b | ENSMUSG00000060548 | -0,461 | 0,044 Tnfrsf19 |
| ENSMUSG00000039630 | 0,185 | 0,016 Hnrnpu | ENSMUSG00000026873 | -0,461 | 0,017 Phf19 |
| ENSMUSG00000028086 | 0,184 | 0,038 Fbxw7 | ENSMUSG00000064105 | -0,461 | 0,002 Cnnm2 |
| ENSMUSG00000032458 | 0,183 | 0,005 Copb2 | ENSMUSG00000029650 | -0,461 | 0,049 Slc46a3 |
| ENSMUSG00000035227 | 0,183 | 0,002 Spcs2 | ENSMUSG00000020888 | -0,464 | 0,000 Dvl2 |
| ENSMUSG00000041130 | 0,182 | 0,006 Zfp598 | ENSMUSG00000032898 | -0,464 | 0,006 Fbxo21 |
| ENSMUSG00000036955 | 0,182 | 0,008 Kif1bp | ENSMUSG00000028884 | -0,464 | 0,017 Rpa2 |
| ENSMUSG00000047719 | 0,182 | 0,018 Ubiad1 | ENSMUSG00000063605 | -0,464 | 0,020 Ccdc102a |
| ENSMUSG00000026127 | 0,182 | 0,000 Imp4 | ENSMUSG00000027669 | -0,465 | 0,004 Gnb4 |
| ENSMUSG00000002844 | 0,181 | 0,019 Adprh | ENSMUSG00000034285 | -0,466 | 0,005 Nipsnap1 |
| ENSMUSG00000037373 | 0,181 | 0,010 Ctbp1 | ENSMUSG00000016637 | -0,466 | 0,039 Ift27 |
| ENSMUSG00000005813 | 0,180 | 0,004 Metap1 | ENSMUSG00000043487 | -0,467 | 0,027 Acot6 |
| ENSMUSG00000069237 | 0,180 | 0,024 Fam8a1 | ENSMUSG00000020883 | -0,468 | 0,004 Fbxl20 |
| ENSMUSG00000037706 | 0,179 | 0,005 Cd81 | ENSMUSG00000049904 | -0,468 | 0,022 Tmem17 |
| ENSMUSG00000024758 | 0,177 | 0,011 Rtn3 | ENSMUSG00000060288 | -0,469 | 0,023 Ppih |
| ENSMUSG00000024604 | 0,177 | 0,001 Rbm22 | ENSMUSG00000022070 | -0,471 | 0,009 Bora |

| | | | | | |
|---|---|---|---|---|---|
| ENSMUSG00000030374 | 0,177 | 0,007 Strn4 | ENSMUSG00000051950 | -0,472 | 0,005 B3glct |
| ENSMUSG00000027465 | 0,175 | 0,016 Tbc1d20 | ENSMUSG00000020868 | -0,473 | 0,000 Xylt2 |
| ENSMUSG00000027706 | 0,175 | 0,029 Sec62 | ENSMUSG00000034247 | -0,473 | 0,000 Plekhm1 |
| ENSMUSG00000051695 | 0,174 | 0,006 Pcbp1 | ENSMUSG00000024696 | -0,474 | 0,002 Lpxn |
| ENSMUSG00000025858 | 0,174 | 0,043 Get4 | ENSMUSG00000005949 | -0,474 | 0,000 Ctns |
| ENSMUSG00000026869 | 0,173 | 0,036 Psmd5 | ENSMUSG00000033596 | -0,474 | 0,001 Rfwd3 |
| ENSMUSG00000060904 | 0,173 | 0,008 Arl1 | ENSMUSG00000046152 | -0,475 | 0,002 Fut10 |
| ENSMUSG00000037742 | 0,172 | 0,024 Eef1a1 | ENSMUSG00000025747 | -0,475 | 0,016 Tyms |
| ENSMUSG00000027012 | 0,172 | 0,003 Dync1i2 | ENSMUSG00000034216 | -0,475 | 0,000 Vps18 |
| ENSMUSG00000048076 | 0,172 | 0,001 Arf1 | ENSMUSG00000056019 | -0,475 | 0,049 Zfp709 |
| ENSMUSG00000036112 | 0,172 | 0,026 Metap2 | ENSMUSG00000038967 | -0,475 | 0,000 Pdk2 |
| ENSMUSG00000023307 | 0,171 | 0,012 March5 | ENSMUSG00000035547 | -0,476 | 0,017 Capn5 |
| ENSMUSG00000054455 | 0,171 | 0,001 Vapb | ENSMUSG00000051343 | -0,476 | 0,006 Rab11fip5 |
| ENSMUSG00000022426 | 0,170 | 0,036 Josd1 | ENSMUSG00000031974 | -0,476 | 0,003 Abcb10 |
| ENSMUSG00000020925 | 0,166 | 0,047 Ccdc43 | ENSMUSG00000040860 | -0,476 | 0,010 Crocc |
| ENSMUSG00000020111 | 0,164 | 0,018 Micu1 | ENSMUSG00000025758 | -0,476 | 0,038 Plk4 |
| ENSMUSG00000030400 | 0,163 | 0,032 Ercc2 | ENSMUSG00000032397 | -0,476 | 0,001 Tipin |
| ENSMUSG00000027425 | 0,163 | 0,030 Kat14 | ENSMUSG00000068299 | -0,478 | 0,007 Nat8f4 |
| ENSMUSG00000026223 | 0,161 | 0,036 Itm2c | ENSMUSG00000005951 | -0,479 | 0,000 Shpk |
| ENSMUSG00000039887 | 0,161 | 0,046 Alg14 | ENSMUSG00000071604 | -0,479 | 0,008 Fam189a2 |
| ENSMUSG00000029227 | 0,160 | 0,045 Fip1l1 | ENSMUSG00000031666 | -0,479 | 0,034 Rbl2 |
| ENSMUSG00000026426 | 0,160 | 0,010 Arl8a | ENSMUSG00000000901 | -0,480 | 0,019 Mmp11 |
| ENSMUSG00000007815 | 0,158 | 0,003 Rhoa | ENSMUSG00000028527 | -0,481 | 0,008 Ak4 |
| ENSMUSG00000042369 | 0,158 | 0,044 Rbm45 | ENSMUSG00000001151 | -0,482 | 0,032 Pcnt |
| ENSMUSG00000029629 | 0,158 | 0,031 Phf14 | ENSMUSG00000023919 | -0,482 | 0,000 Cenpq |
| ENSMUSG00000058407 | 0,155 | 0,027 Txndc9 | ENSMUSG00000035351 | -0,483 | 0,013 Nup37 |
| ENSMUSG00000019087 | 0,155 | 0,007 Atp6ap1 | ENSMUSG00000027353 | -0,483 | 0,031 Mcm8 |
| ENSMUSG00000037669 | 0,145 | 0,039 Ldah | ENSMUSG00000039316 | -0,483 | 0,000 Rftn1 |
| ENSMUSG00000024587 | 0,142 | 0,022 Nars | ENSMUSG00000000253 | -0,484 | 0,011 Gmpr |
| ENSMUSG00000027349 | 0,142 | 0,025 Fam98b | ENSMUSG00000045268 | -0,484 | 0,031 Zfp691 |
| ENSMUSG00000032199 | 0,138 | 0,046 Polr2m | ENSMUSG00000046027 | -0,485 | 0,000 Stard5 |
| ENSMUSG00000032435 | 0,136 | 0,048 Dync1li1 | ENSMUSG00000000056 | -0,487 | 0,000 Narf |
| ENSMUSG00000025228 | 0,136 | 0,006 Actr1a | ENSMUSG00000030086 | -0,487 | 0,048 Chchd6 |
| ENSMUSG00000007458 | 0,135 | 0,005 M6pr | ENSMUSG00000042616 | -0,489 | 0,029 Oscp1 |
| ENSMUSG00000025130 | 0,134 | 0,047 P4hb | ENSMUSG00000076435 | -0,489 | 0,012 Acsf2 |
| ENSMUSG00000062270 | 0,133 | 0,020 Morf4l1 | ENSMUSG00000037295 | -0,492 | 0,000 Ldlrap1 |
| ENSMUSG00000021484 | 0,133 | 0,031 Lman2 | ENSMUSG00000027864 | -0,492 | 0,016 Ptgfrn |
| ENSMUSG00000007411 | 0,132 | 0,021 Mark3 | ENSMUSG00000078794 | -0,493 | 0,001 Dact3 |
| ENSMUSG00000031447 | 0,132 | 0,012 Lamp1 | ENSMUSG00000065979 | -0,494 | 0,000 Cpped1 |
| ENSMUSG00000020936 | 0,130 | 0,025 Nmt1 | ENSMUSG00000014303 | -0,494 | 0,010 Glis2 |
| ENSMUSG00000001017 | 0,128 | 0,022 Chtop | ENSMUSG00000024866 | -0,495 | 0,041 Acy3 |
| ENSMUSG00000017264 | 0,122 | 0,015 Exosc10 | ENSMUSG00000025277 | -0,495 | 0,000 Abhd6 |
| ENSMUSG00000059923 | 0,119 | 0,008 Grb2 | ENSMUSG00000028088 | -0,496 | 0,004 Fmo5 |
| ENSMUSG00000002981 | 0,096 | 0,049 Clptm1 | ENSMUSG00000107879 | -0,497 | 0,024 9330102E08Rik |
| ENSMUSG00000015165 | -0,138 | 0,043 Hnrnpl | ENSMUSG00000022881 | -0,497 | 0,013 Rfc4 |
| ENSMUSG00000029247 | -0,144 | 0,007 Paics | ENSMUSG00000018809 | -0,498 | 0,002 Smyd4 |
| ENSMUSG00000015889 | -0,160 | 0,034 Lta4h | ENSMUSG00000048142 | -0,498 | 0,000 Nat8l |

| | | | | | | | |
|---|---|---|---|---|---|---|---|
| ENSMUSG00000041133 | -0,161 | 0,040 | Smc1a | ENSMUSG00000030729 | -0,499 | 0,036 | Pgm2l1 |
| ENSMUSG00000021193 | -0,162 | 0,045 | Pitrm1 | ENSMUSG00000037235 | -0,499 | 0,000 | Mxd4 |
| ENSMUSG00000040667 | -0,165 | 0,002 | Nup88 | ENSMUSG00000032741 | -0,501 | 0,001 | Tpcn1 |
| ENSMUSG00000062908 | -0,167 | 0,019 | Acadm | ENSMUSG00000089764 | -0,502 | 0,048 | Gm16580 |
| ENSMUSG00000013833 | -0,167 | 0,036 | Med16 | ENSMUSG00000020034 | -0,503 | 0,001 | Tcp11l2 |
| ENSMUSG00000030726 | -0,169 | 0,031 | Pold3 | ENSMUSG00000028980 | -0,503 | 0,002 | H6pd |
| ENSMUSG00000078676 | -0,173 | 0,018 | Casc3 | ENSMUSG00000019942 | -0,503 | 0,003 | Cdk1 |
| ENSMUSG00000036980 | -0,174 | 0,040 | Taf6 | ENSMUSG00000024168 | -0,504 | 0,015 | Tmem204 |
| ENSMUSG00000039254 | -0,174 | 0,012 | Pomt1 | ENSMUSG00000050107 | -0,505 | 0,000 | Haspin |
| ENSMUSG00000029012 | -0,177 | 0,049 | Orc5 | ENSMUSG00000066894 | -0,507 | 0,005 | Vsig10 |
| ENSMUSG00000000441 | -0,177 | 0,016 | Raf1 | ENSMUSG00000029386 | -0,508 | 0,030 | Tctn2 |
| ENSMUSG00000013646 | -0,184 | 0,037 | Sh3bp5l | ENSMUSG00000067199 | -0,508 | 0,028 | Frat1 |
| ENSMUSG00000013787 | -0,184 | 0,000 | Ehmt2 | ENSMUSG00000013495 | -0,508 | 0,005 | Tmem175 |
| ENSMUSG00000024797 | -0,186 | 0,032 | Vps51 | ENSMUSG00000101225 | -0,508 | 0,041 | 1700008J07Rik |
| ENSMUSG00000038225 | -0,187 | 0,049 | Primpol | ENSMUSG00000028073 | -0,509 | 0,003 | Pear1 |
| ENSMUSG00000038838 | -0,191 | 0,028 | Vars2 | ENSMUSG00000005615 | -0,510 | 0,002 | Pcyt1a |
| ENSMUSG00000020776 | -0,191 | 0,018 | Fbf1 | ENSMUSG00000041777 | -0,510 | 0,005 | Cir1 |
| ENSMUSG00000021115 | -0,192 | 0,026 | Vrk1 | ENSMUSG00000022833 | -0,512 | 0,005 | Ccdc14 |
| ENSMUSG00000017210 | -0,193 | 0,045 | Med24 | ENSMUSG00000027330 | -0,512 | 0,006 | Cdc25b |
| ENSMUSG00000006526 | -0,193 | 0,048 | Tmem110 | ENSMUSG00000028696 | -0,512 | 0,027 | Ipp |
| ENSMUSG00000024096 | -0,195 | 0,004 | Ralbp1 | ENSMUSG00000019845 | -0,512 | 0,013 | Tube1 |
| ENSMUSG00000057367 | -0,195 | 0,006 | Birc2 | ENSMUSG00000095930 | -0,513 | 0,016 | Nim1k |
| ENSMUSG00000030822 | -0,195 | 0,044 | Prr14 | ENSMUSG00000020541 | -0,513 | 0,022 | Tom1l1 |
| ENSMUSG00000043964 | -0,196 | 0,036 | Orai3 | ENSMUSG00000027635 | -0,514 | 0,014 | Dsn1 |
| ENSMUSG00000071647 | -0,196 | 0,021 | Eml3 | ENSMUSG00000020532 | -0,515 | 0,001 | Acaca |
| ENSMUSG00000042032 | -0,196 | 0,006 | Mat2b | ENSMUSG00000043157 | -0,516 | 0,000 | Arl11 |
| ENSMUSG00000005899 | -0,197 | 0,017 | Smpd4 | ENSMUSG00000026944 | -0,516 | 0,017 | Abca2 |
| ENSMUSG00000027709 | -0,198 | 0,007 | Mccc1 | ENSMUSG00000022475 | -0,516 | 0,001 | Hdac7 |
| ENSMUSG00000021240 | -0,198 | 0,038 | Abcd4 | ENSMUSG00000022799 | -0,517 | 0,036 | Arhgap31 |
| ENSMUSG00000028291 | -0,198 | 0,026 | Akirin2 | ENSMUSG00000038070 | -0,517 | 0,026 | Cntln |
| ENSMUSG00000039599 | -0,199 | 0,007 | Fam149b | ENSMUSG00000035944 | -0,517 | 0,000 | Ttc38 |
| ENSMUSG00000020375 | -0,199 | 0,015 | Rufy1 | ENSMUSG00000056458 | -0,517 | 0,014 | Mok |
| ENSMUSG00000022678 | -0,199 | 0,018 | Nde1 | ENSMUSG00000026317 | -0,519 | 0,001 | Cln8 |
| ENSMUSG00000021900 | -0,199 | 0,005 | Btd | ENSMUSG00000030598 | -0,519 | 0,010 | Fbxo17 |
| ENSMUSG00000027809 | -0,200 | 0,004 | Etfdh | ENSMUSG00000028066 | -0,519 | 0,015 | Pmf1 |
| ENSMUSG00000087177 | -0,200 | 0,029 | E130307A14Rik | ENSMUSG00000026489 | -0,520 | 0,001 | Coq8a |
| ENSMUSG00000035632 | -0,201 | 0,013 | Cnot3 | ENSMUSG00000022673 | -0,520 | 0,000 | Mcm4 |
| ENSMUSG00000028954 | -0,201 | 0,024 | Nub1 | ENSMUSG00000025083 | -0,523 | 0,037 | Afap1l2 |
| ENSMUSG00000022750 | -0,201 | 0,016 | Klhl22 | ENSMUSG00000029086 | -0,524 | 0,012 | Prom1 |
| ENSMUSG00000028646 | -0,203 | 0,034 | Rragc | ENSMUSG00000037020 | -0,524 | 0,000 | Wdr62 |
| ENSMUSG00000029422 | -0,204 | 0,040 | Rsrc2 | ENSMUSG00000021697 | -0,525 | 0,039 | Depdc1b |
| ENSMUSG00000026003 | -0,204 | 0,037 | Acadl | ENSMUSG00000027784 | -0,526 | 0,035 | Ppm1l |
| ENSMUSG00000013419 | -0,205 | 0,012 | Zfp651 | ENSMUSG00000071064 | -0,526 | 0,007 | Zfp827 |
| ENSMUSG00000046679 | -0,209 | 0,013 | C87436 | ENSMUSG00000032586 | -0,526 | 0,001 | Traip |
| ENSMUSG00000036120 | -0,211 | 0,029 | Rfxank | ENSMUSG00000031478 | -0,527 | 0,004 | Nek3 |
| ENSMUSG00000025188 | -0,211 | 0,015 | Hps1 | ENSMUSG00000039347 | -0,527 | 0,000 | Atp6v0e2 |
| ENSMUSG00000029033 | -0,212 | 0,036 | Acap3 | ENSMUSG00000015944 | -0,527 | 0,000 | Gatsl2 |

| | | | | | | | |
|---|---|---|---|---|---|---|---|
| ENSMUSG00000021646 | -0,215 | 0,013 | Mccc2 | ENSMUSG00000046157 | -0,528 | 0,002 | Tmem229b |
| ENSMUSG00000029781 | -0,216 | 0,018 | Fkbp9 | ENSMUSG00000041809 | -0,529 | 0,031 | Efhc1 |
| ENSMUSG00000026790 | -0,216 | 0,002 | Odf2 | ENSMUSG00000002996 | -0,529 | 0,000 | Hbp1 |
| ENSMUSG00000010110 | -0,216 | 0,040 | Stx5a | ENSMUSG00000027883 | -0,530 | 0,001 | Gpsm2 |
| ENSMUSG00000031378 | -0,217 | 0,031 | Abcd1 | ENSMUSG00000026456 | -0,530 | 0,008 | Cyb5r1 |
| ENSMUSG00000032392 | -0,218 | 0,028 | Parp16 | ENSMUSG00000023908 | -0,531 | 0,005 | Pkmyt1 |
| ENSMUSG00000042709 | -0,218 | 0,018 | Atpaf2 | ENSMUSG00000034685 | -0,531 | 0,002 | Fam171a2 |
| ENSMUSG00000036622 | -0,218 | 0,014 | Atp13a2 | ENSMUSG00000054717 | -0,531 | 0,002 | Hmgb2 |
| ENSMUSG00000036188 | -0,219 | 0,006 | Ankmy2 | ENSMUSG00000021719 | -0,532 | 0,002 | Rgs7bp |
| ENSMUSG00000002803 | -0,220 | 0,049 | Btbd6 | ENSMUSG00000043262 | -0,534 | 0,027 | Uevld |
| ENSMUSG00000027198 | -0,221 | 0,000 | Ext2 | ENSMUSG00000058290 | -0,534 | 0,003 | Espl1 |
| ENSMUSG00000105053 | -0,221 | 0,016 | Gm43064 | ENSMUSG00000041949 | -0,535 | 0,000 | Tango6 |
| ENSMUSG00000021252 | -0,221 | 0,005 | Erg28 | ENSMUSG00000022453 | -0,535 | 0,012 | Naga |
| ENSMUSG00000003038 | -0,222 | 0,033 | Hmgn2 | ENSMUSG00000037890 | -0,537 | 0,020 | Wdr19 |
| ENSMUSG00000042292 | -0,222 | 0,028 | Mkl1 | ENSMUSG00000004891 | -0,537 | 0,001 | Nes |
| ENSMUSG00000028973 | -0,222 | 0,005 | Abcb8 | ENSMUSG00000044317 | -0,538 | 0,039 | Gpr4 |
| ENSMUSG00000028759 | -0,222 | 0,014 | Hp1bp3 | ENSMUSG00000051098 | -0,538 | 0,042 | Mblac2 |
| ENSMUSG00000063052 | -0,226 | 0,048 | Lrrc40 | ENSMUSG00000033031 | -0,538 | 0,048 | C330027C09Rik |
| ENSMUSG00000039536 | -0,227 | 0,013 | Stau1 | ENSMUSG00000063804 | -0,539 | 0,008 | Lin28b |
| ENSMUSG00000030347 | -0,228 | 0,049 | D6Wsu163e | ENSMUSG00000035863 | -0,540 | 0,000 | Palm |
| ENSMUSG00000091512 | -0,229 | 0,038 | Lamtor3 | ENSMUSG00000031767 | -0,540 | 0,013 | Nudt7 |
| ENSMUSG00000115018 | -0,230 | 0,037 | AL732309.1 | ENSMUSG00000041406 | -0,541 | 0,001 | BC055324 |
| ENSMUSG00000019961 | -0,230 | 0,024 | Tmpo | ENSMUSG00000006800 | -0,541 | 0,004 | Sulf2 |
| ENSMUSG00000093445 | -0,231 | 0,047 | Lrch4 | ENSMUSG00000034586 | -0,541 | 0,031 | Hid1 |
| ENSMUSG00000032952 | -0,231 | 0,005 | Ap4b1 | ENSMUSG00000006678 | -0,541 | 0,026 | Pola1 |
| ENSMUSG00000021000 | -0,231 | 0,048 | Ctage5 | ENSMUSG00000055240 | -0,544 | 0,008 | Zfp101 |
| ENSMUSG00000026554 | -0,232 | 0,001 | Dcaf8 | ENSMUSG00000038332 | -0,544 | 0,000 | Sesn1 |
| ENSMUSG00000041890 | -0,232 | 0,019 | Git2 | ENSMUSG00000047003 | -0,545 | 0,020 | Zfp41 |
| ENSMUSG00000022185 | -0,232 | 0,010 | Acin1 | ENSMUSG00000027875 | -0,546 | 0,020 | Hmgcs2 |
| ENSMUSG00000027890 | -0,232 | 0,047 | Gstm4 | ENSMUSG00000014164 | -0,546 | 0,021 | Klhl3 |
| ENSMUSG00000012296 | -0,232 | 0,013 | Tjap1 | ENSMUSG00000002068 | -0,546 | 0,021 | Ccne1 |
| ENSMUSG00000059981 | -0,233 | 0,045 | Taok2 | ENSMUSG00000073434 | -0,546 | 0,001 | Wdr90 |
| ENSMUSG00000031864 | -0,234 | 0,002 | Ints10 | ENSMUSG00000019823 | -0,548 | 0,000 | Mical1 |
| ENSMUSG00000031508 | -0,234 | 0,015 | Ankrd10 | ENSMUSG00000021186 | -0,548 | 0,007 | Fbln5 |
| ENSMUSG00000020451 | -0,235 | 0,010 | Limk2 | ENSMUSG00000029401 | -0,551 | 0,032 | Rilpl2 |
| ENSMUSG00000011832 | -0,236 | 0,008 | Evi5l | ENSMUSG00000001864 | -0,553 | 0,006 | Aif1l |
| ENSMUSG00000037965 | -0,236 | 0,037 | Zc3h7a | ENSMUSG00000036249 | -0,554 | 0,000 | Rbm43 |
| ENSMUSG00000020605 | -0,241 | 0,021 | Hs1bp3 | ENSMUSG00000038618 | -0,554 | 0,010 | Rassf7 |
| ENSMUSG00000058388 | -0,242 | 0,026 | Phtf1 | ENSMUSG00000020492 | -0,555 | 0,000 | Ska2 |
| ENSMUSG00000019295 | -0,243 | 0,012 | Tmem129 | ENSMUSG00000032633 | -0,555 | 0,003 | Flcn |
| ENSMUSG00000029536 | -0,243 | 0,004 | Gatc | ENSMUSG00000030930 | -0,556 | 0,000 | Chst15 |
| ENSMUSG00000029201 | -0,244 | 0,038 | Ugdh | ENSMUSG00000023972 | -0,556 | 0,002 | Ptk7 |
| ENSMUSG00000028737 | -0,245 | 0,024 | Aldh4a1 | ENSMUSG00000009575 | -0,558 | 0,001 | Cbx5 |
| ENSMUSG00000032513 | -0,246 | 0,004 | Gorasp1 | ENSMUSG00000039853 | -0,558 | 0,031 | Trim14 |
| ENSMUSG00000004567 | -0,248 | 0,025 | Mcoln1 | ENSMUSG00000073791 | -0,559 | 0,003 | Efcab7 |
| ENSMUSG00000036057 | -0,248 | 0,036 | Ptpn23 | ENSMUSG00000074793 | -0,559 | 0,013 | Hspa12b |
| ENSMUSG00000020385 | -0,249 | 0,024 | Clk4 | ENSMUSG00000073705 | -0,563 | 0,033 | Cenps |

| | | | | | | | |
|---|---|---|---|---|---|---|---|
| ENSMUSG00000026174 | -0,250 | 0,002 | Cnot9 | ENSMUSG00000040147 | -0,564 | 0,046 | Maob |
| ENSMUSG00000035206 | -0,250 | 0,001 | Sppl2b | ENSMUSG00000006585 | -0,565 | 0,000 | Cdt1 |
| ENSMUSG00000016619 | -0,251 | 0,012 | Nup50 | ENSMUSG00000006235 | -0,565 | 0,000 | Epor |
| ENSMUSG00000019715 | -0,251 | 0,000 | Gle1 | ENSMUSG00000089715 | -0,566 | 0,000 | Cbx6 |
| ENSMUSG00000047412 | -0,251 | 0,031 | Zbtb44 | ENSMUSG00000052302 | -0,569 | 0,043 | Tbc1d30 |
| ENSMUSG00000040365 | -0,252 | 0,002 | Trim41 | ENSMUSG00000062683 | -0,569 | 0,000 | Atp5g2 |
| ENSMUSG00000022898 | -0,252 | 0,000 | Dscr3 | ENSMUSG00000027242 | -0,570 | 0,008 | Wdr76 |
| ENSMUSG00000020941 | -0,253 | 0,038 | Map3k14 | ENSMUSG00000028943 | -0,570 | 0,001 | Espn |
| ENSMUSG00000028096 | -0,253 | 0,039 | Gpr89 | ENSMUSG00000059810 | -0,570 | 0,000 | Rgs3 |
| ENSMUSG00000037012 | -0,253 | 0,006 | Hk1 | ENSMUSG00000031150 | -0,570 | 0,042 | Ccdc120 |
| ENSMUSG00000047766 | -0,254 | 0,019 | Lrrc49 | ENSMUSG00000042628 | -0,572 | 0,000 | Zfyve1 |
| ENSMUSG00000002871 | -0,254 | 0,009 | Tpra1 | ENSMUSG00000024666 | -0,573 | 0,000 | Tmem138 |
| ENSMUSG00000037344 | -0,254 | 0,008 | Slc12a9 | ENSMUSG00000026688 | -0,573 | 0,032 | Mgst3 |
| ENSMUSG00000021483 | -0,254 | 0,042 | Cdk20 | ENSMUSG00000051517 | -0,574 | 0,022 | Arhgef39 |
| ENSMUSG00000036435 | -0,254 | 0,001 | Exoc1 | ENSMUSG00000031669 | -0,574 | 0,000 | Gins3 |
| ENSMUSG00000022793 | -0,254 | 0,018 | B4galt4 | ENSMUSG00000086712 | -0,575 | 0,016 | AI427809 |
| ENSMUSG00000089872 | -0,255 | 0,020 | Rps6kc1 | ENSMUSG00000024742 | -0,575 | 0,000 | Fen1 |
| ENSMUSG00000020818 | -0,255 | 0,002 | Mfsd11 | ENSMUSG00000079304 | -0,575 | 0,040 | Tex52 |
| ENSMUSG00000042726 | -0,256 | 0,006 | Trafd1 | ENSMUSG00000040621 | -0,575 | 0,000 | Gemin8 |
| ENSMUSG00000003810 | -0,256 | 0,008 | Mast2 | ENSMUSG00000006378 | -0,576 | 0,005 | Gcat |
| ENSMUSG00000019143 | -0,257 | 0,000 | Hars2 | ENSMUSG00000072837 | -0,577 | 0,001 | Mir546 |
| ENSMUSG00000000876 | -0,257 | 0,001 | Pxmp4 | ENSMUSG00000005470 | -0,578 | 0,007 | Asf1b |
| ENSMUSG00000027342 | -0,258 | 0,017 | Pcna | ENSMUSG00000078942 | -0,578 | 0,013 | Naip6 |
| ENSMUSG00000032333 | -0,258 | 0,043 | Stoml1 | ENSMUSG00000047786 | -0,578 | 0,008 | Lix1 |
| ENSMUSG00000008226 | -0,258 | 0,041 | Scrn3 | ENSMUSG00000038215 | -0,581 | 0,000 | Cep44 |
| ENSMUSG00000042055 | -0,260 | 0,021 | Wdr11 | ENSMUSG00000078429 | -0,581 | 0,000 | Ctdsp2 |
| ENSMUSG00000022718 | -0,260 | 0,037 | Dgcr8 | ENSMUSG00000021180 | -0,584 | 0,017 | Rps6ka5 |
| ENSMUSG00000027620 | -0,261 | 0,004 | Rbm39 | ENSMUSG00000029406 | -0,585 | 0,005 | Pitpnm2 |
| ENSMUSG00000062110 | -0,261 | 0,026 | Scfd2 | ENSMUSG00000036526 | -0,585 | 0,038 | Card11 |
| ENSMUSG00000021982 | -0,262 | 0,011 | Cdadc1 | ENSMUSG00000025555 | -0,585 | 0,001 | Farp1 |
| ENSMUSG00000024764 | -0,262 | 0,037 | Naa40 | ENSMUSG00000015217 | -0,586 | 0,000 | Hmgb3 |
| ENSMUSG00000021065 | -0,262 | 0,034 | Fut8 | ENSMUSG00000018983 | -0,586 | 0,003 | E2f2 |
| ENSMUSG00000018841 | -0,262 | 0,014 | Rad51d | ENSMUSG00000035683 | -0,586 | 0,016 | Melk |
| ENSMUSG00000040701 | -0,263 | 0,045 | Ap1g2 | ENSMUSG00000076617 | -0,587 | 0,038 | Ighm |
| ENSMUSG00000022414 | -0,263 | 0,014 | Tab1 | ENSMUSG00000029438 | -0,588 | 0,000 | Bcl7a |
| ENSMUSG00000024833 | -0,264 | 0,006 | Pola2 | ENSMUSG00000044783 | -0,588 | 0,001 | Hjurp |
| ENSMUSG00000028799 | -0,265 | 0,001 | Zfp362 | ENSMUSG00000091498 | -0,589 | 0,001 | Mpc1-ps |
| ENSMUSG00000025077 | -0,265 | 0,013 | Dclre1a | ENSMUSG00000062081 | -0,589 | 0,008 | Gm6055 |
| ENSMUSG00000028394 | -0,266 | 0,013 | Pole3 | ENSMUSG00000020415 | -0,592 | 0,040 | Pttg1 |
| ENSMUSG00000024588 | -0,266 | 0,001 | Fech | ENSMUSG00000047881 | -0,593 | 0,000 | Rell1 |
| ENSMUSG00000041528 | -0,268 | 0,014 | Rnf123 | ENSMUSG00000027454 | -0,594 | 0,000 | Gins1 |
| ENSMUSG00000002781 | -0,269 | 0,007 | Tmem143 | ENSMUSG00000107476 | -0,594 | 0,009 | Zfp862-ps |
| ENSMUSG00000027227 | -0,271 | 0,027 | Sord | ENSMUSG00000079559 | -0,596 | 0,034 | Colca2 |
| ENSMUSG00000034042 | -0,272 | 0,000 | Gpbp1l1 | ENSMUSG00000034906 | -0,596 | 0,000 | Ncaph |
| ENSMUSG00000034863 | -0,273 | 0,000 | Ano8 | ENSMUSG00000045625 | -0,596 | 0,018 | Pigz |
| ENSMUSG00000020752 | -0,273 | 0,000 | Recql5 | ENSMUSG00000097415 | -0,597 | 0,003 | AU020206 |
| ENSMUSG00000038506 | -0,273 | 0,049 | Dcun1d2 | ENSMUSG00000021573 | -0,600 | 0,017 | Tppp |

| | | | | | | | |
|---|---|---|---|---|---|---|---|
| ENSMUSG00000028917 | -0,273 | 0,034 | Plekhm2 | ENSMUSG00000054115 | -0,602 | 0,000 | Skp2 |
| ENSMUSG00000028221 | -0,274 | 0,019 | Tmem55a | ENSMUSG00000045930 | -0,602 | 0,013 | Clec14a |
| ENSMUSG00000050989 | -0,275 | 0,011 | Selenon | ENSMUSG00000056394 | -0,603 | 0,000 | Lig1 |
| ENSMUSG00000062906 | -0,275 | 0,010 | Hdac10 | ENSMUSG00000038390 | -0,603 | 0,007 | Gpr162 |
| ENSMUSG00000062040 | -0,276 | 0,047 | Zfp27 | ENSMUSG00000044037 | -0,604 | 0,002 | Als2cl |
| ENSMUSG00000038425 | -0,276 | 0,003 | Poli | ENSMUSG00000034601 | -0,606 | 0,000 | 2700049A03Rik |
| ENSMUSG00000009079 | -0,276 | 0,000 | Ewsr1 | ENSMUSG00000050379 | -0,607 | 0,000 | Sept6 |
| ENSMUSG00000020770 | -0,276 | 0,003 | Unk | ENSMUSG00000026080 | -0,607 | 0,002 | Chst10 |
| ENSMUSG00000027010 | -0,276 | 0,004 | Slc25a12 | ENSMUSG00000032121 | -0,607 | 0,000 | Tmem218 |
| ENSMUSG00000068335 | -0,276 | 0,001 | Dok1 | ENSMUSG00000109523 | -0,607 | 0,031 | Gdf1 |
| ENSMUSG00000029056 | -0,277 | 0,032 | Pank4 | ENSMUSG00000045165 | -0,610 | 0,002 | AI467606 |
| ENSMUSG00000029098 | -0,277 | 0,020 | Acox3 | ENSMUSG00000040936 | -0,610 | 0,027 | Ulk4 |
| ENSMUSG00000067369 | -0,277 | 0,013 | Trmt2b | ENSMUSG00000052688 | -0,611 | 0,007 | Rab7b |
| ENSMUSG00000034485 | -0,277 | 0,018 | Uaca | ENSMUSG00000059713 | -0,611 | 0,000 | Rcan3 |
| ENSMUSG00000037523 | -0,277 | 0,025 | Mavs | ENSMUSG00000066191 | -0,612 | 0,000 | Anks6 |
| ENSMUSG00000031358 | -0,278 | 0,029 | Msl3 | ENSMUSG00000028789 | -0,612 | 0,011 | Azin2 |
| ENSMUSG00000024590 | -0,281 | 0,023 | Lmnb1 | ENSMUSG00000022021 | -0,612 | 0,002 | Diaph3 |
| ENSMUSG00000033628 | -0,281 | 0,001 | Pik3c3 | ENSMUSG00000018931 | -0,613 | 0,000 | Natd1 |
| ENSMUSG00000031161 | -0,281 | 0,000 | Hdac6 | ENSMUSG00000039908 | -0,613 | 0,032 | Slc26a11 |
| ENSMUSG00000037257 | -0,281 | 0,016 | Aagab | ENSMUSG00000020744 | -0,614 | 0,001 | Slc25a19 |
| ENSMUSG00000078517 | -0,283 | 0,036 | Emc1 | ENSMUSG00000043614 | -0,615 | 0,008 | Vps37d |
| ENSMUSG00000001034 | -0,283 | 0,001 | Mapk7 | ENSMUSG00000030254 | -0,616 | 0,003 | Rad18 |
| ENSMUSG00000025925 | -0,283 | 0,001 | Terf1 | ENSMUSG00000032352 | -0,617 | 0,002 | Lrrc1 |
| ENSMUSG00000021271 | -0,284 | 0,024 | Zfp839 | ENSMUSG00000029363 | -0,618 | 0,000 | Rfc5 |
| ENSMUSG00000049858 | -0,284 | 0,014 | Suox | ENSMUSG00000010064 | -0,618 | 0,001 | Slc38a3 |
| ENSMUSG00000070814 | -0,284 | 0,008 | Zswim9 | ENSMUSG00000038068 | -0,618 | 0,018 | Rnf144b |
| ENSMUSG00000039637 | -0,285 | 0,005 | Coro7 | ENSMUSG00000029869 | -0,619 | 0,004 | Ephb6 |
| ENSMUSG00000038718 | -0,285 | 0,013 | Pbx3 | ENSMUSG00000050022 | -0,621 | 0,000 | Amz1 |
| ENSMUSG00000030614 | -0,285 | 0,016 | Tmem126b | ENSMUSG00000027469 | -0,621 | 0,004 | Tpx2 |
| ENSMUSG00000022558 | -0,285 | 0,019 | Mroh1 | ENSMUSG00000019214 | -0,623 | 0,004 | Chtf18 |
| ENSMUSG00000031485 | -0,285 | 0,002 | Plpbp | ENSMUSG00000070390 | -0,624 | 0,038 | Nlrp1b |
| ENSMUSG00000052395 | -0,287 | 0,000 | Rft1 | ENSMUSG00000071176 | -0,625 | 0,008 | Arhgef10 |
| ENSMUSG00000046949 | -0,288 | 0,004 | Nqo2 | ENSMUSG00000041859 | -0,625 | 0,000 | Mcm3 |
| ENSMUSG00000062646 | -0,289 | 0,050 | Ganc | ENSMUSG00000036867 | -0,626 | 0,001 | Smad6 |
| ENSMUSG00000038260 | -0,289 | 0,030 | Trpm4 | ENSMUSG00000035024 | -0,626 | 0,004 | Ncapd3 |
| ENSMUSG00000038604 | -0,289 | 0,022 | Ripor1 | ENSMUSG00000021287 | -0,626 | 0,000 | Xrcc3 |
| ENSMUSG00000020196 | -0,289 | 0,015 | Cabin1 | ENSMUSG00000063445 | -0,627 | 0,000 | Nmral1 |
| ENSMUSG00000039781 | -0,290 | 0,045 | Cep131 | ENSMUSG00000022723 | -0,628 | 0,006 | Crybg3 |
| ENSMUSG00000033703 | -0,290 | 0,019 | Fuk | ENSMUSG00000042500 | -0,629 | 0,003 | Ago4 |
| ENSMUSG00000026134 | -0,290 | 0,008 | Prim2 | ENSMUSG00000038943 | -0,630 | 0,002 | Prc1 |
| ENSMUSG00000046753 | -0,290 | 0,048 | Ccdc66 | ENSMUSG00000031595 | -0,630 | 0,004 | Pdgfrl |
| ENSMUSG00000099519 | -0,291 | 0,000 | Gm29253 | ENSMUSG00000053062 | -0,630 | 0,013 | Jam2 |
| ENSMUSG00000029047 | -0,293 | 0,033 | Pex10 | ENSMUSG00000039963 | -0,630 | 0,025 | Ccdc40 |
| ENSMUSG00000025008 | -0,293 | 0,047 | Tctn3 | ENSMUSG00000056214 | -0,631 | 0,001 | Pard6g |
| ENSMUSG00000025366 | -0,294 | 0,013 | Esyt1 | ENSMUSG00000050390 | -0,632 | 0,014 | C77080 |
| ENSMUSG00000035399 | -0,295 | 0,046 | Oser1 | ENSMUSG00000020592 | -0,633 | 0,002 | Sdc1 |
| ENSMUSG00000038564 | -0,295 | 0,035 | Ift172 | ENSMUSG00000020593 | -0,634 | 0,006 | Lpin1 |

| | | | | | | | |
|---|---|---|---|---|---|---|---|
| ENSMUSG00000026925 | -0,296 | 0,023 | Inpp5e | ENSMUSG00000030319 | -0,634 | 0,002 | Cand2 |
| ENSMUSG00000061313 | -0,296 | 0,030 | Ddhd2 | ENSMUSG00000030978 | -0,635 | 0,000 | Rrm1 |
| ENSMUSG00000111409 | -0,296 | 0,036 | Rnf26 | ENSMUSG00000026355 | -0,636 | 0,000 | Mcm6 |
| ENSMUSG00000037826 | -0,296 | 0,047 | Ppm1k | ENSMUSG00000019732 | -0,638 | 0,042 | Calr3 |
| ENSMUSG00000042138 | -0,297 | 0,016 | Msantd2 | ENSMUSG00000020019 | -0,639 | 0,042 | Ntn4 |
| ENSMUSG00000038206 | -0,297 | 0,001 | Fbxo8 | ENSMUSG00000021575 | -0,639 | 0,001 | Ahrr |
| ENSMUSG00000062526 | -0,298 | 0,010 | Mppe1 | ENSMUSG00000029512 | -0,639 | 0,000 | Ulk1 |
| ENSMUSG00000027341 | -0,298 | 0,049 | Tmem230 | ENSMUSG00000039385 | -0,640 | 0,040 | Cdh6 |
| ENSMUSG00000026270 | -0,298 | 0,015 | Capn10 | ENSMUSG00000024901 | -0,640 | 0,001 | Peli3 |
| ENSMUSG00000046897 | -0,299 | 0,005 | Zfp740 | ENSMUSG00000027075 | -0,642 | 0,000 | Slc43a1 |
| ENSMUSG00000024561 | -0,299 | 0,000 | Mbd1 | ENSMUSG00000021176 | -0,643 | 0,001 | Efcab11 |
| ENSMUSG00000024556 | -0,299 | 0,032 | Me2 | ENSMUSG00000007646 | -0,644 | 0,000 | Rad51c |
| ENSMUSG00000034006 | -0,299 | 0,012 | Pqlc1 | ENSMUSG00000026955 | -0,644 | 0,004 | Sapcd2 |
| ENSMUSG00000020747 | -0,300 | 0,024 | Tmem94 | ENSMUSG00000032218 | -0,645 | 0,011 | Ccnb2 |
| ENSMUSG00000038241 | -0,300 | 0,018 | Cep250 | ENSMUSG00000024989 | -0,645 | 0,034 | Cep55 |
| ENSMUSG00000031453 | -0,301 | 0,049 | Rasa3 | ENSMUSG00000087408 | -0,645 | 0,001 | Cers1 |
| ENSMUSG00000010660 | -0,301 | 0,003 | Plcd1 | ENSMUSG00000041147 | -0,647 | 0,000 | Brca2 |
| ENSMUSG00000037894 | -0,302 | 0,031 | H2afz | ENSMUSG00000001039 | -0,647 | 0,006 | B9d1 |
| ENSMUSG00000026482 | -0,302 | 0,048 | Rgl1 | ENSMUSG00000036862 | -0,648 | 0,034 | Dchs1 |
| ENSMUSG00000040964 | -0,302 | 0,005 | Arhgef10l | ENSMUSG00000055884 | -0,650 | 0,016 | Fancm |
| ENSMUSG00000072847 | -0,303 | 0,043 | A530017D24Rik | ENSMUSG00000003824 | -0,650 | 0,024 | Syce2 |
| ENSMUSG00000047037 | -0,304 | 0,029 | Nipa1 | ENSMUSG00000078762 | -0,651 | 0,000 | Haus5 |
| ENSMUSG00000087385 | -0,304 | 0,045 | Frg2f1 | ENSMUSG00000046618 | -0,651 | 0,023 | Olfml2a |
| ENSMUSG00000030780 | -0,305 | 0,002 | BC017158 | ENSMUSG00000049624 | -0,655 | 0,000 | Slc17a5 |
| ENSMUSG00000032192 | -0,305 | 0,029 | Gnb5 | ENSMUSG00000021747 | -0,655 | 0,007 | 4930452B06Rik |
| ENSMUSG00000007589 | -0,305 | 0,009 | Tinf2 | ENSMUSG00000059013 | -0,655 | 0,000 | Sh2d3c |
| ENSMUSG00000032498 | -0,306 | 0,004 | Mlh1 | ENSMUSG00000037572 | -0,656 | 0,000 | Wdhd1 |
| ENSMUSG00000047514 | -0,307 | 0,007 | Tspyl1 | ENSMUSG00000048249 | -0,657 | 0,001 | Crebrf |
| ENSMUSG00000115074 | -0,307 | 0,000 | AL732309.2 | ENSMUSG00000021884 | -0,658 | 0,000 | Hacl1 |
| ENSMUSG00000039989 | -0,307 | 0,045 | Cbx4 | ENSMUSG00000039763 | -0,659 | 0,001 | Dnajc28 |
| ENSMUSG00000074738 | -0,307 | 0,024 | Fndc10 | ENSMUSG00000024795 | -0,663 | 0,020 | Kif20b |
| ENSMUSG00000028809 | -0,307 | 0,018 | Srrm1 | ENSMUSG00000020774 | -0,665 | 0,007 | Aspa |
| ENSMUSG00000009621 | -0,308 | 0,026 | Vav2 | ENSMUSG00000048965 | -0,666 | 0,000 | Mrgpre |
| ENSMUSG00000035863 | -0,308 | 0,034 | Palm | ENSMUSG00000025140 | -0,666 | 0,010 | Pycr1 |
| ENSMUSG00000039307 | -0,309 | 0,050 | Hexdc | ENSMUSG00000031756 | -0,667 | 0,000 | Cenpn |
| ENSMUSG00000044465 | -0,310 | 0,001 | Fam160a2 | ENSMUSG00000044702 | -0,667 | 0,000 | Palb2 |
| ENSMUSG00000027984 | -0,310 | 0,005 | Hadh | ENSMUSG00000115022 | -0,668 | 0,000 | Hacl1 |
| ENSMUSG00000006127 | -0,310 | 0,000 | Inpp5k | ENSMUSG00000028873 | -0,668 | 0,005 | Cdca8 |
| ENSMUSG00000021023 | -0,311 | 0,020 | 1110008L16Rik | ENSMUSG00000043290 | -0,669 | 0,000 | Zfp784 |
| ENSMUSG00000027291 | -0,313 | 0,013 | Vps39 | ENSMUSG00000047757 | -0,669 | 0,002 | Fancb |
| ENSMUSG00000021820 | -0,313 | 0,008 | Camk2g | ENSMUSG00000019699 | -0,669 | 0,029 | Akt3 |
| ENSMUSG00000020439 | -0,313 | 0,012 | Smtn | ENSMUSG00000054893 | -0,670 | 0,000 | Zfp667 |
| ENSMUSG00000028484 | -0,313 | 0,000 | Psip1 | ENSMUSG00000022091 | -0,670 | 0,000 | Sorbs3 |
| ENSMUSG00000022389 | -0,314 | 0,033 | Tef | ENSMUSG00000022033 | -0,670 | 0,002 | Pbk |
| ENSMUSG00000030929 | -0,314 | 0,050 | Eri2 | ENSMUSG00000037725 | -0,671 | 0,001 | Ckap2 |
| ENSMUSG00000059772 | -0,314 | 0,009 | Slx1b | ENSMUSG00000020914 | -0,672 | 0,007 | Top2a |
| ENSMUSG00000055436 | -0,314 | 0,016 | Srsf11 | ENSMUSG00000054850 | -0,672 | 0,001 | Smim10l2a |

| | | | | | | | |
|---|---|---|---|---|---|---|---|
| ENSMUSG00000025050 | -0,314 | 0,032 | Pcgf6 | ENSMUSG00000029413 | -0,673 | 0,038 | Naaa |
| ENSMUSG00000035891 | -0,316 | 0,019 | Cerk | ENSMUSG00000048458 | -0,673 | 0,001 | Fam212b |
| ENSMUSG00000043411 | -0,317 | 0,032 | Usp48 | ENSMUSG00000074194 | -0,674 | 0,019 | Zfp791 |
| ENSMUSG00000007850 | -0,318 | 0,002 | Hnrnph1 | ENSMUSG00000002997 | -0,675 | 0,003 | Prkar2b |
| ENSMUSG00000025283 | -0,318 | 0,030 | Sat1 | ENSMUSG00000073371 | -0,676 | 0,004 | Gm6594 |
| ENSMUSG00000057778 | -0,318 | 0,016 | Cyb5d2 | ENSMUSG00000027533 | -0,677 | 0,004 | Fabp5 |
| ENSMUSG00000016933 | -0,318 | 0,045 | Plcg1 | ENSMUSG00000006398 | -0,678 | 0,010 | Cdc20 |
| ENSMUSG00000022885 | -0,319 | 0,021 | St6gal1 | ENSMUSG00000019303 | -0,678 | 0,001 | Psmc3ip |
| ENSMUSG00000036678 | -0,319 | 0,022 | Aaas | ENSMUSG00000022756 | -0,679 | 0,000 | Slc7a4 |
| ENSMUSG00000024118 | -0,319 | 0,009 | Tedc2 | ENSMUSG00000032251 | -0,679 | 0,032 | Irak1bp1 |
| ENSMUSG00000062075 | -0,320 | 0,005 | Lmnb2 | ENSMUSG00000002409 | -0,680 | 0,000 | Dyrk1b |
| ENSMUSG00000024006 | -0,320 | 0,000 | Stk38 | ENSMUSG00000025001 | -0,680 | 0,010 | Hells |
| ENSMUSG00000028582 | -0,321 | 0,000 | Cc2d1b | ENSMUSG00000022203 | -0,681 | 0,018 | Efs |
| ENSMUSG00000022244 | -0,321 | 0,018 | Amacr | ENSMUSG00000071723 | -0,685 | 0,012 | Gspt2 |
| ENSMUSG00000029455 | -0,321 | 0,024 | Aldh2 | ENSMUSG00000022385 | -0,686 | 0,001 | Gtse1 |
| ENSMUSG00000026031 | -0,322 | 0,035 | Cflar | ENSMUSG00000048416 | -0,686 | 0,040 | Mlf1 |
| ENSMUSG00000022019 | -0,322 | 0,000 | Tdrd3 | ENSMUSG00000037568 | -0,686 | 0,003 | Vash2 |
| ENSMUSG00000029547 | -0,323 | 0,007 | Ints1 | ENSMUSG00000030380 | -0,687 | 0,004 | Mzf1 |
| ENSMUSG00000070044 | -0,323 | 0,030 | Fam149a | ENSMUSG00000001403 | -0,687 | 0,006 | Ube2c |
| ENSMUSG00000019878 | -0,324 | 0,026 | Hsf2 | ENSMUSG00000029910 | -0,690 | 0,000 | Mad2l1 |
| ENSMUSG00000030161 | -0,324 | 0,005 | Gabarapl1 | ENSMUSG00000025576 | -0,691 | 0,013 | Rbfox3 |
| ENSMUSG00000022537 | -0,324 | 0,025 | Tmem44 | ENSMUSG00000032855 | -0,691 | 0,036 | Pkd1 |
| ENSMUSG00000000751 | -0,325 | 0,000 | Rpa1 | ENSMUSG00000045751 | -0,693 | 0,000 | Mms22l |
| ENSMUSG00000028960 | -0,325 | 0,027 | Ube4b | ENSMUSG00000038370 | -0,694 | 0,012 | Pcp4l1 |
| ENSMUSG00000053460 | -0,326 | 0,020 | Ggcx | ENSMUSG00000017716 | -0,696 | 0,031 | Birc5 |
| ENSMUSG00000045730 | -0,327 | 0,035 | Adrb2 | ENSMUSG00000024301 | -0,697 | 0,009 | Kifc5b |
| ENSMUSG00000035776 | -0,327 | 0,000 | Cd99l2 | ENSMUSG00000047473 | -0,697 | 0,005 | Zfp30 |
| ENSMUSG00000006288 | -0,327 | 0,000 | Ttc5 | ENSMUSG00000021294 | -0,697 | 0,017 | Kif26a |
| ENSMUSG00000020263 | -0,327 | 0,049 | Appl2 | ENSMUSG00000047747 | -0,698 | 0,008 | Rnf150 |
| ENSMUSG00000026709 | -0,327 | 0,003 | Dars2 | ENSMUSG00000026683 | -0,698 | 0,000 | Nuf2 |
| ENSMUSG00000021007 | -0,328 | 0,031 | Spata7 | ENSMUSG00000038119 | -0,698 | 0,049 | Cdon |
| ENSMUSG00000074093 | -0,328 | 0,016 | Svip | ENSMUSG00000017550 | -0,699 | 0,036 | Atad5 |
| ENSMUSG00000037443 | -0,328 | 0,025 | Cep85 | ENSMUSG00000005043 | -0,700 | 0,000 | Sgsh |
| ENSMUSG00000019917 | -0,329 | 0,046 | Sept10 | ENSMUSG00000062510 | -0,701 | 0,000 | Nsl1 |
| ENSMUSG00000025486 | -0,329 | 0,041 | Sirt3 | ENSMUSG00000030110 | -0,703 | 0,018 | Ret |
| ENSMUSG00000050619 | -0,330 | 0,004 | Zscan29 | ENSMUSG00000020099 | -0,704 | 0,010 | Unc5b |
| ENSMUSG00000020190 | -0,330 | 0,001 | Mknk2 | ENSMUSG00000029299 | -0,706 | 0,007 | Abcg3 |
| ENSMUSG00000032329 | -0,331 | 0,002 | Hmg20a | ENSMUSG00000056832 | -0,706 | 0,010 | Ttc26 |
| ENSMUSG00000032264 | -0,331 | 0,000 | Zw10 | ENSMUSG00000020895 | -0,707 | 0,016 | Tmem107 |
| ENSMUSG00000002297 | -0,332 | 0,005 | Dbf4 | ENSMUSG00000037628 | -0,707 | 0,036 | Cdkn3 |
| ENSMUSG00000031641 | -0,332 | 0,002 | Cbr4 | ENSMUSG00000041836 | -0,707 | 0,012 | Ptpre |
| ENSMUSG00000018372 | -0,333 | 0,009 | Cep95 | ENSMUSG00000002870 | -0,709 | 0,000 | Mcm2 |
| ENSMUSG00000029428 | -0,334 | 0,000 | Stx2 | ENSMUSG00000031821 | -0,709 | 0,000 | Gins2 |
| ENSMUSG00000039713 | -0,334 | 0,031 | Plekhg5 | ENSMUSG00000024376 | -0,709 | 0,024 | Epb41l4a |
| ENSMUSG00000027180 | -0,334 | 0,020 | Fbxo3 | ENSMUSG00000020330 | -0,711 | 0,006 | Hmmr |
| ENSMUSG00000022106 | -0,334 | 0,009 | Rcbtb2 | ENSMUSG00000052852 | -0,711 | 0,000 | Reep1 |
| ENSMUSG00000042476 | -0,334 | 0,028 | Abcb4 | ENSMUSG00000052921 | -0,712 | 0,018 | Arhgef15 |

| | | | | | | | |
|---|---|---|---|---|---|---|---|
| ENSMUSG00000024974 | -0,335 | 0,011 | Smc3 | ENSMUSG00000035246 | -0,713 | 0,002 | Pcyt1b |
| ENSMUSG00000039347 | -0,335 | 0,000 | Atp6v0e2 | ENSMUSG00000023015 | -0,714 | 0,002 | Racgap1 |
| ENSMUSG00000031858 | -0,336 | 0,033 | Mau2 | ENSMUSG00000024727 | -0,714 | 0,036 | Trpm6 |
| ENSMUSG00000079109 | -0,336 | 0,000 | Pms2 | ENSMUSG00000045328 | -0,715 | 0,010 | Cenpe |
| ENSMUSG00000059482 | -0,336 | 0,035 | 2610301B20Rik | ENSMUSG00000085133 | -0,717 | 0,002 | B930095G15Rik |
| ENSMUSG00000022621 | -0,336 | 0,010 | Rabl2 | ENSMUSG00000038022 | -0,718 | 0,000 | Mindy4 |
| ENSMUSG00000041625 | -0,336 | 0,024 | Ggact | ENSMUSG00000108011 | -0,718 | 0,004 | Gm44596 |
| ENSMUSG00000031787 | -0,337 | 0,000 | Katnb1 | ENSMUSG00000029516 | -0,720 | 0,022 | Cit |
| ENSMUSG00000042787 | -0,337 | 0,004 | Exog | ENSMUSG00000031379 | -0,720 | 0,023 | Pir |
| ENSMUSG00000039958 | -0,337 | 0,015 | Etfbkmt | ENSMUSG00000038644 | -0,721 | 0,000 | Pold1 |
| ENSMUSG00000025795 | -0,337 | 0,045 | Rassf3 | ENSMUSG00000030528 | -0,722 | 0,000 | Blm |
| ENSMUSG00000109493 | -0,338 | 0,004 | Gm45208 | ENSMUSG00000047115 | -0,722 | 0,011 | Fam221a |
| ENSMUSG00000039914 | -0,338 | 0,050 | Coq10a | ENSMUSG00000049488 | -0,724 | 0,000 | Tmem67 |
| ENSMUSG00000030089 | -0,338 | 0,016 | Slc41a3 | ENSMUSG00000021175 | -0,724 | 0,000 | Cdca7l |
| ENSMUSG00000020823 | -0,338 | 0,000 | Sec14l1 | ENSMUSG00000020486 | -0,725 | 0,043 | Sept4 |
| ENSMUSG00000025151 | -0,339 | 0,000 | Maged1 | ENSMUSG00000016756 | -0,726 | 0,049 | Cmah |
| ENSMUSG00000044629 | -0,339 | 0,001 | Cnrip1 | ENSMUSG00000028445 | -0,726 | 0,007 | Enho |
| ENSMUSG00000046567 | -0,339 | 0,028 | 4930430F08Rik | ENSMUSG00000029730 | -0,728 | 0,000 | Mcm7 |
| ENSMUSG00000054728 | -0,340 | 0,023 | Phactr1 | ENSMUSG00000027306 | -0,728 | 0,003 | Nusap1 |
| ENSMUSG00000043190 | -0,340 | 0,030 | Rfesd | ENSMUSG00000032113 | -0,728 | 0,000 | Chek1 |
| ENSMUSG00000031684 | -0,340 | 0,003 | Slc10a7 | ENSMUSG00000004099 | -0,729 | 0,000 | Dnmt1 |
| ENSMUSG00000027452 | -0,340 | 0,023 | Acss1 | ENSMUSG00000028773 | -0,730 | 0,046 | Fabp3 |
| ENSMUSG00000006471 | -0,341 | 0,001 | Ndor1 | ENSMUSG00000034311 | -0,730 | 0,001 | Kif4 |
| ENSMUSG00000032714 | -0,341 | 0,028 | Syde1 | ENSMUSG00000027654 | -0,731 | 0,006 | Fam83d |
| ENSMUSG00000040410 | -0,342 | 0,002 | Fbxl4 | ENSMUSG00000037151 | -0,732 | 0,000 | Lrrc20 |
| ENSMUSG00000006527 | -0,342 | 0,038 | Sfmbt1 | ENSMUSG00000094595 | -0,733 | 0,001 | Fsbp |
| ENSMUSG00000009828 | -0,343 | 0,000 | Ick | ENSMUSG00000027203 | -0,735 | 0,001 | Dut |
| ENSMUSG00000060301 | -0,344 | 0,010 | 2610008E11Rik | ENSMUSG00000047534 | -0,735 | 0,002 | Mis18bp1 |
| ENSMUSG00000003778 | -0,344 | 0,001 | Brd8 | ENSMUSG00000040111 | -0,737 | 0,001 | Gramd1b |
| ENSMUSG00000018796 | -0,344 | 0,028 | Acsl1 | ENSMUSG00000036777 | -0,738 | 0,012 | Anln |
| ENSMUSG00000028089 | -0,344 | 0,001 | Chd1l | ENSMUSG00000041633 | -0,739 | 0,005 | Kctd12b |
| ENSMUSG00000040302 | -0,345 | 0,048 | Rbm48 | ENSMUSG00000015568 | -0,739 | 0,007 | Lpl |
| ENSMUSG00000015944 | -0,347 | 0,001 | Gatsl2 | ENSMUSG00000038252 | -0,740 | 0,000 | Ncapd2 |
| ENSMUSG00000031959 | -0,347 | 0,031 | Wdr59 | ENSMUSG00000025154 | -0,740 | 0,000 | Arhgap19 |
| ENSMUSG00000018548 | -0,348 | 0,035 | Trim37 | ENSMUSG00000044201 | -0,742 | 0,019 | Cdc25c |
| ENSMUSG00000056204 | -0,348 | 0,000 | Pgpep1 | ENSMUSG00000074212 | -0,742 | 0,001 | Dnajb14 |
| ENSMUSG00000039197 | -0,349 | 0,000 | Adk | ENSMUSG00000024525 | -0,744 | 0,000 | Impa2 |
| ENSMUSG00000031821 | -0,349 | 0,027 | Gins2 | ENSMUSG00000024791 | -0,744 | 0,001 | Cdca5 |
| ENSMUSG00000019338 | -0,349 | 0,048 | Zfp687 | ENSMUSG00000028587 | -0,747 | 0,005 | Orc1 |
| ENSMUSG00000021179 | -0,350 | 0,013 | Nrde2 | ENSMUSG00000107761 | -0,748 | 0,049 | 2010008C14Rik |
| ENSMUSG00000041471 | -0,350 | 0,049 | Fam35a | ENSMUSG00000048922 | -0,748 | 0,004 | Cdca2 |
| ENSMUSG00000034032 | -0,351 | 0,010 | Rpap1 | ENSMUSG00000030499 | -0,749 | 0,000 | Kctd15 |
| ENSMUSG00000026986 | -0,352 | 0,011 | Hnmt | ENSMUSG00000031486 | -0,751 | 0,000 | Adgra2 |
| ENSMUSG00000025791 | -0,353 | 0,000 | Pgm2 | ENSMUSG00000084989 | -0,751 | 0,042 | Crocc2 |
| ENSMUSG00000050382 | -0,353 | 0,010 | Kif7 | ENSMUSG00000030515 | -0,752 | 0,000 | Tarsl2 |
| ENSMUSG00000030067 | -0,353 | 0,025 | Foxp1 | ENSMUSG00000048327 | -0,753 | 0,001 | Ckap2l |
| ENSMUSG00000021830 | -0,353 | 0,048 | Txndc16 | ENSMUSG00000048424 | -0,753 | 0,034 | Ranbp3l |

| | | | | | |
|---|---|---|---|---|---|
| ENSMUSG00000035051 | -0,354 | 0,000 Dhx57 | ENSMUSG00000027715 | -0,754 | 0,001 Ccna2 |
| ENSMUSG00000028300 | -0,354 | 0,010 3110043O21Rik | ENSMUSG00000034206 | -0,754 | 0,002 Polq |
| ENSMUSG00000038545 | -0,356 | 0,000 Cul7 | ENSMUSG00000027699 | -0,755 | 0,030 Ect2 |
| ENSMUSG00000026707 | -0,356 | 0,013 Nsun6 | ENSMUSG00000036295 | -0,757 | 0,021 Lrrn3 |
| ENSMUSG00000026213 | -0,356 | 0,004 Stk11ip | ENSMUSG00000073910 | -0,758 | 0,040 Mob3b |
| ENSMUSG00000039382 | -0,357 | 0,010 Wdr45 | ENSMUSG00000040276 | -0,758 | 0,000 Pacsin1 |
| ENSMUSG00000032782 | -0,357 | 0,001 Cntrob | ENSMUSG00000006958 | -0,760 | 0,041 Chrd |
| ENSMUSG00000034248 | -0,358 | 0,006 Slc25a37 | ENSMUSG00000022177 | -0,761 | 0,000 Haus4 |
| ENSMUSG00000031295 | -0,359 | 0,003 Phka2 | ENSMUSG00000020098 | -0,761 | 0,009 Pcbd1 |
| ENSMUSG00000000278 | -0,359 | 0,000 Scpep1 | ENSMUSG00000049580 | -0,762 | 0,000 Tsku |
| ENSMUSG00000038365 | -0,359 | 0,000 Fbxo25 | ENSMUSG00000037466 | -0,764 | 0,000 Tedc1 |
| ENSMUSG00000041949 | -0,359 | 0,022 Tango6 | ENSMUSG00000040084 | -0,765 | 0,002 Bub1b |
| ENSMUSG00000048142 | -0,360 | 0,004 Nat8l | ENSMUSG00000114515 | -0,766 | 0,037 Aldoa |
| ENSMUSG00000041491 | -0,360 | 0,001 Cep78 | ENSMUSG00000032717 | -0,767 | 0,035 Mdfi |
| ENSMUSG00000043067 | -0,361 | 0,030 Dpy19l1 | ENSMUSG00000089837 | -0,768 | 0,043 Npcd |
| ENSMUSG00000018395 | -0,362 | 0,037 Kif3a | ENSMUSG00000003779 | -0,768 | 0,001 Kif20a |
| ENSMUSG00000042834 | -0,363 | 0,002 Nrep | ENSMUSG00000022322 | -0,768 | 0,001 Shcbp1 |
| ENSMUSG00000031668 | -0,364 | 0,035 Eif2ak3 | ENSMUSG00000072082 | -0,769 | 0,000 Ccnf |
| ENSMUSG00000039671 | -0,364 | 0,028 Zmynd8 | ENSMUSG00000072762 | -0,770 | 0,002 4930522L14Rik |
| ENSMUSG00000023191 | -0,364 | 0,000 P3h3 | ENSMUSG00000022604 | -0,773 | 0,000 Cep97 |
| ENSMUSG00000039231 | -0,365 | 0,000 Suv39h1 | ENSMUSG00000001517 | -0,776 | 0,000 Foxm1 |
| ENSMUSG00000033065 | -0,365 | 0,000 Pfkm | ENSMUSG00000071324 | -0,777 | 0,000 Armc2 |
| ENSMUSG00000035629 | -0,365 | 0,018 Rubcn | ENSMUSG00000054074 | -0,779 | 0,036 Skida1 |
| ENSMUSG00000032580 | -0,365 | 0,000 Rbm5 | ENSMUSG00000028803 | -0,779 | 0,000 Nipal3 |
| ENSMUSG00000030872 | -0,365 | 0,001 Gga2 | ENSMUSG00000035455 | -0,780 | 0,000 Fignl1 |
| ENSMUSG00000030782 | -0,365 | 0,029 Tgfb1i1 | ENSMUSG00000040658 | -0,780 | 0,001 Dnph1 |
| ENSMUSG00000031922 | -0,366 | 0,002 Cep57 | ENSMUSG00000032815 | -0,781 | 0,002 Fanca |
| ENSMUSG00000085793 | -0,366 | 0,000 Lin52 | ENSMUSG00000032400 | -0,783 | 0,000 Zwilch |
| ENSMUSG00000038637 | -0,366 | 0,013 Lrrc56 | ENSMUSG00000000782 | -0,784 | 0,000 Tcf7 |
| ENSMUSG00000001750 | -0,366 | 0,001 Tcirg1 | ENSMUSG00000034023 | -0,785 | 0,001 Fancd2 |
| ENSMUSG00000034327 | -0,367 | 0,016 Kctd9 | ENSMUSG00000020381 | -0,785 | 0,000 Mrnip |
| ENSMUSG00000037622 | -0,367 | 0,001 Wdtc1 | ENSMUSG00000030346 | -0,787 | 0,000 Rad51ap1 |
| ENSMUSG00000020532 | -0,367 | 0,018 Acaca | ENSMUSG00000075266 | -0,787 | 0,020 Cenpw |
| ENSMUSG00000040820 | -0,367 | 0,020 Hlcs | ENSMUSG00000037337 | -0,789 | 0,004 Map4k1 |
| ENSMUSG00000010307 | -0,368 | 0,012 Tmem86a | ENSMUSG00000031883 | -0,790 | 0,031 Car7 |
| ENSMUSG00000046605 | -0,368 | 0,000 B3gntl1 | ENSMUSG00000032666 | -0,791 | 0,048 1700025G04Rik |
| ENSMUSG00000028245 | -0,369 | 0,000 Nsmaf | ENSMUSG00000042029 | -0,791 | 0,028 Ncapg2 |
| ENSMUSG00000020474 | -0,371 | 0,004 Polm | ENSMUSG00000051278 | -0,792 | 0,015 Zgrf1 |
| ENSMUSG00000034243 | -0,371 | 0,031 Golgb1 | ENSMUSG00000002055 | -0,793 | 0,000 Spag5 |
| ENSMUSG00000006641 | -0,372 | 0,002 Slc5a6 | ENSMUSG00000033083 | -0,795 | 0,001 Tbc1d4 |
| ENSMUSG00000037656 | -0,373 | 0,005 Slc20a2 | ENSMUSG00000074802 | -0,796 | 0,016 Gas2l3 |
| ENSMUSG00000026355 | -0,373 | 0,030 Mcm6 | ENSMUSG00000028121 | -0,797 | 0,000 Bcar3 |
| ENSMUSG00000013236 | -0,373 | 0,024 Ptprs | ENSMUSG00000079685 | -0,797 | 0,004 Ulbp1 |
| ENSMUSG00000013033 | -0,374 | 0,017 Adgrl1 | ENSMUSG00000064043 | -0,798 | 0,002 Trerf1 |
| ENSMUSG00000055817 | -0,374 | 0,003 Mta3 | ENSMUSG00000049521 | -0,800 | 0,000 Cdc42ep1 |
| ENSMUSG00000026179 | -0,375 | 0,000 Pnkd | ENSMUSG00000027331 | -0,801 | 0,002 Knstrn |
| ENSMUSG00000027695 | -0,375 | 0,027 Pld1 | ENSMUSG00000015880 | -0,804 | 0,002 Ncapg |

| | | | | | | | |
|---|---|---|---|---|---|---|---|
| ENSMUSG00000021975 | -0,375 | 0,000 | Ints9 | ENSMUSG00000030865 | -0,807 | 0,018 | Chp2 |
| ENSMUSG00000002504 | -0,375 | 0,044 | Slc9a3r2 | ENSMUSG00000074476 | -0,808 | 0,003 | Spc24 |
| ENSMUSG00000050144 | -0,376 | 0,003 | Slc25a44 | ENSMUSG00000033060 | -0,809 | 0,034 | Lmo7 |
| ENSMUSG00000047407 | -0,376 | 0,021 | Tgif1 | ENSMUSG00000021714 | -0,810 | 0,004 | Cenpk |
| ENSMUSG00000009739 | -0,376 | 0,005 | Pou6f1 | ENSMUSG00000069910 | -0,811 | 0,000 | Spdl1 |
| ENSMUSG00000020086 | -0,376 | 0,043 | H2afy2 | ENSMUSG00000029591 | -0,811 | 0,000 | Ung |
| ENSMUSG00000020380 | -0,376 | 0,001 | Rad50 | ENSMUSG00000034917 | -0,812 | 0,001 | Tjp3 |
| ENSMUSG00000033439 | -0,376 | 0,026 | Trmt13 | ENSMUSG00000026605 | -0,812 | 0,005 | Cenpf |
| ENSMUSG00000074166 | -0,377 | 0,048 | AW146154 | ENSMUSG00000087406 | -0,813 | 0,008 | E130215H24Rik |
| ENSMUSG00000013629 | -0,377 | 0,022 | Cad | ENSMUSG00000017861 | -0,813 | 0,000 | Mybl2 |
| ENSMUSG00000022617 | -0,378 | 0,016 | Chkb | ENSMUSG00000030091 | -0,817 | 0,000 | Nup210 |
| ENSMUSG00000021245 | -0,378 | 0,010 | Mlh3 | ENSMUSG00000027074 | -0,818 | 0,000 | Slc43a3 |
| ENSMUSG00000019518 | -0,378 | 0,000 | Ap4m1 | ENSMUSG00000049588 | -0,818 | 0,000 | Ccdc69 |
| ENSMUSG00000030223 | -0,378 | 0,001 | Ptpro | ENSMUSG00000002032 | -0,819 | 0,002 | Tmem25 |
| ENSMUSG00000028164 | -0,379 | 0,000 | Manba | ENSMUSG00000012443 | -0,820 | 0,002 | Kif11 |
| ENSMUSG00000033059 | -0,379 | 0,000 | Pygb | ENSMUSG00000079553 | -0,827 | 0,001 | Kifc1 |
| ENSMUSG00000036944 | -0,380 | 0,022 | Tmem71 | ENSMUSG00000030123 | -0,828 | 0,014 | Plxnd1 |
| ENSMUSG00000091498 | -0,380 | 0,042 | Mpc1-ps | ENSMUSG00000039055 | -0,829 | 0,000 | Eme1 |
| ENSMUSG00000052949 | -0,380 | 0,006 | Rnf157 | ENSMUSG00000026888 | -0,829 | 0,015 | Grb14 |
| ENSMUSG00000029710 | -0,381 | 0,034 | Ephb4 | ENSMUSG00000030677 | -0,830 | 0,003 | Kif22 |
| ENSMUSG00000049739 | -0,381 | 0,015 | Zfp646 | ENSMUSG00000051022 | -0,831 | 0,001 | Hs3st1 |
| ENSMUSG00000004891 | -0,381 | 0,019 | Nes | ENSMUSG00000024245 | -0,832 | 0,004 | Tmem178 |
| ENSMUSG00000039298 | -0,383 | 0,044 | Cdk5rap2 | ENSMUSG00000027490 | -0,834 | 0,000 | E2f1 |
| ENSMUSG00000027963 | -0,383 | 0,001 | Extl2 | ENSMUSG00000028438 | -0,835 | 0,011 | Kif24 |
| ENSMUSG00000029578 | -0,383 | 0,000 | Wipi2 | ENSMUSG00000040631 | -0,835 | 0,001 | Dok4 |
| ENSMUSG00000044068 | -0,384 | 0,002 | Zrsr1 | ENSMUSG00000031004 | -0,837 | 0,004 | Mki67 |
| ENSMUSG00000038884 | -0,385 | 0,041 | A230050P20Rik | ENSMUSG00000030606 | -0,837 | 0,000 | Hapln3 |
| ENSMUSG00000046152 | -0,385 | 0,013 | Fut10 | ENSMUSG00000036768 | -0,837 | 0,000 | Kif15 |
| ENSMUSG00000032397 | -0,385 | 0,007 | Tipin | ENSMUSG00000045092 | -0,838 | 0,000 | S1pr1 |
| ENSMUSG00000054474 | -0,385 | 0,046 | Thnsl2 | ENSMUSG00000022034 | -0,840 | 0,001 | Esco2 |
| ENSMUSG00000030083 | -0,386 | 0,020 | Abtb1 | ENSMUSG00000027698 | -0,842 | 0,000 | Nceh1 |
| ENSMUSG00000031539 | -0,386 | 0,002 | Ap3m2 | ENSMUSG00000027323 | -0,846 | 0,000 | Rad51 |
| ENSMUSG00000039623 | -0,386 | 0,001 | Ap5z1 | ENSMUSG00000022422 | -0,848 | 0,000 | Dscc1 |
| ENSMUSG00000022369 | -0,386 | 0,000 | Mtbp | ENSMUSG00000006715 | -0,849 | 0,000 | Gmnn |
| ENSMUSG00000005986 | -0,387 | 0,010 | Ankrd13d | ENSMUSG00000017002 | -0,851 | 0,029 | Slpi |
| ENSMUSG00000050107 | -0,387 | 0,002 | Haspin | ENSMUSG00000022358 | -0,851 | 0,000 | Fbxo32 |
| ENSMUSG00000040040 | -0,387 | 0,007 | Ift88 | ENSMUSG00000049119 | -0,852 | 0,000 | Fam110b |
| ENSMUSG00000028943 | -0,387 | 0,034 | Espn | ENSMUSG00000041064 | -0,853 | 0,000 | Pif1 |
| ENSMUSG00000038520 | -0,388 | 0,021 | Tbc1d17 | ENSMUSG00000027792 | -0,853 | 0,005 | Bche |
| ENSMUSG00000025235 | -0,388 | 0,000 | Bbs4 | ENSMUSG00000051043 | -0,856 | 0,000 | Gprc5c |
| ENSMUSG00000071350 | -0,388 | 0,027 | Setdb2 | ENSMUSG00000027115 | -0,857 | 0,001 | Kif18a |
| ENSMUSG00000033326 | -0,388 | 0,006 | Kdm4a | ENSMUSG00000111184 | -0,858 | 0,000 | AC163703.1 |
| ENSMUSG00000021226 | -0,390 | 0,003 | Acot2 | ENSMUSG00000025577 | -0,859 | 0,000 | Cbx2 |
| ENSMUSG00000023055 | -0,390 | 0,000 | Calcoco1 | ENSMUSG00000030867 | -0,860 | 0,000 | Plk1 |
| ENSMUSG00000028675 | -0,390 | 0,000 | Pnrc2 | ENSMUSG00000026039 | -0,860 | 0,001 | Sgo2a |
| ENSMUSG00000025358 | -0,390 | 0,000 | Cdk2 | ENSMUSG00000023940 | -0,862 | 0,000 | Sgo1 |
| ENSMUSG00000030177 | -0,390 | 0,000 | Ccdc77 | ENSMUSG00000027379 | -0,862 | 0,000 | Bub1 |

| | | | | | | | |
|---|---|---|---|---|---|---|---|
| ENSMUSG00000031835 | -0,390 | 0,000 | Mbtps1 | ENSMUSG00000031629 | -0,862 | 0,000 | Cenpu |
| ENSMUSG00000027829 | -0,391 | 0,004 | Ccnl1 | ENSMUSG00000020974 | -0,863 | 0,000 | Pole2 |
| ENSMUSG00000024660 | -0,391 | 0,013 | Incenp | ENSMUSG00000045231 | -0,864 | 0,003 | BC106179 |
| ENSMUSG00000097715 | -0,391 | 0,038 | Gpr137b-ps | ENSMUSG00000026447 | -0,865 | 0,043 | Pik3c2b |
| ENSMUSG00000029603 | -0,393 | 0,037 | Dtx1 | ENSMUSG00000049538 | -0,865 | 0,014 | Adamts16 |
| ENSMUSG00000041975 | -0,393 | 0,017 | Mettl8 | ENSMUSG00000059479 | -0,865 | 0,004 | B3gnt8 |
| ENSMUSG00000029186 | -0,394 | 0,001 | Pi4k2b | ENSMUSG00000027811 | -0,865 | 0,000 | 4930579G24Rik |
| ENSMUSG00000024666 | -0,394 | 0,003 | Tmem138 | ENSMUSG00000034858 | -0,866 | 0,000 | Fam214a |
| ENSMUSG00000012017 | -0,394 | 0,004 | Scarf2 | ENSMUSG00000020808 | -0,868 | 0,000 | Pimreg |
| ENSMUSG00000042408 | -0,395 | 0,048 | Zmym6 | ENSMUSG00000040016 | -0,869 | 0,015 | Ptger3 |
| ENSMUSG00000056698 | -0,395 | 0,001 | Elmod3 | ENSMUSG00000050541 | -0,870 | 0,008 | Adra1b |
| ENSMUSG00000055493 | -0,395 | 0,028 | Epm2a | ENSMUSG00000030322 | -0,871 | 0,000 | Mbd4 |
| ENSMUSG00000061371 | -0,396 | 0,024 | Zfp873 | ENSMUSG00000039187 | -0,871 | 0,000 | Fanci |
| ENSMUSG00000054708 | -0,396 | 0,017 | Ankrd24 | ENSMUSG00000069830 | -0,874 | 0,030 | Nlrp1a |
| ENSMUSG00000032898 | -0,396 | 0,017 | Fbxo21 | ENSMUSG00000004110 | -0,874 | 0,038 | Cacna1e |
| ENSMUSG00000049657 | -0,396 | 0,006 | Zbtb5 | ENSMUSG00000032254 | -0,879 | 0,000 | Kif23 |
| ENSMUSG00000030323 | -0,397 | 0,016 | Ift122 | ENSMUSG00000038305 | -0,880 | 0,000 | Spats2l |
| ENSMUSG00000020668 | -0,397 | 0,015 | Kif3c | ENSMUSG00000020623 | -0,880 | 0,000 | Map2k6 |
| ENSMUSG00000025766 | -0,397 | 0,016 | D3Ertd751e | ENSMUSG00000028718 | -0,880 | 0,001 | Stil |
| ENSMUSG00000002222 | -0,398 | 0,035 | Rmnd5a | ENSMUSG00000024420 | -0,881 | 0,010 | Zfp521 |
| ENSMUSG00000038967 | -0,398 | 0,000 | Pdk2 | ENSMUSG00000027496 | -0,882 | 0,000 | Aurka |
| ENSMUSG00000031266 | -0,399 | 0,006 | Gla | ENSMUSG00000040204 | -0,886 | 0,000 | Pclaf |
| ENSMUSG00000021377 | -0,399 | 0,002 | Dek | ENSMUSG00000028175 | -0,888 | 0,016 | Depdc1a |
| ENSMUSG00000066306 | -0,400 | 0,038 | Numa1 | ENSMUSG00000086111 | -0,889 | 0,018 | Gm15326 |
| ENSMUSG00000023919 | -0,400 | 0,002 | Cenpq | ENSMUSG00000068101 | -0,891 | 0,004 | Cenpm |
| ENSMUSG00000035722 | -0,400 | 0,001 | Abca7 | ENSMUSG00000060402 | -0,891 | 0,012 | Chst8 |
| ENSMUSG00000041341 | -0,400 | 0,039 | Atg2b | ENSMUSG00000110631 | -0,892 | 0,041 | Gm42047 |
| ENSMUSG00000061455 | -0,402 | 0,000 | Stx17 | ENSMUSG00000009876 | -0,892 | 0,018 | Cox4i2 |
| ENSMUSG00000028497 | -0,402 | 0,050 | Hacd4 | ENSMUSG00000029177 | -0,893 | 0,001 | Cenpa |
| ENSMUSG00000045039 | -0,403 | 0,013 | Megf8 | ENSMUSG00000000028 | -0,893 | 0,000 | Cdc45 |
| ENSMUSG00000004952 | -0,403 | 0,005 | Rasa4 | ENSMUSG00000024056 | -0,893 | 0,000 | Ndc80 |
| ENSMUSG00000048200 | -0,403 | 0,038 | Cracr2b | ENSMUSG00000020682 | -0,895 | 0,003 | Mmp28 |
| ENSMUSG00000036249 | -0,404 | 0,002 | Rbm43 | ENSMUSG00000045287 | -0,895 | 0,000 | Rtn4rl1 |
| ENSMUSG00000017405 | -0,404 | 0,015 | Nek8 | ENSMUSG00000021707 | -0,898 | 0,000 | Dhfr |
| ENSMUSG00000027284 | -0,405 | 0,037 | Cdan1 | ENSMUSG00000050410 | -0,898 | 0,000 | Tcf19 |
| ENSMUSG00000057530 | -0,406 | 0,019 | Ece1 | ENSMUSG00000026622 | -0,898 | 0,001 | Nek2 |
| ENSMUSG00000034518 | -0,407 | 0,007 | Hmgxb4 | ENSMUSG00000106825 | -0,901 | 0,000 | 2510016D11Rik |
| ENSMUSG00000005225 | -0,407 | 0,012 | Plekha8 | ENSMUSG00000001228 | -0,901 | 0,000 | Uhrf1 |
| ENSMUSG00000008855 | -0,407 | 0,000 | Hdac5 | ENSMUSG00000036223 | -0,901 | 0,000 | Ska1 |
| ENSMUSG00000044712 | -0,408 | 0,021 | Slc38a6 | ENSMUSG00000023505 | -0,901 | 0,000 | Cdca3 |
| ENSMUSG00000041096 | -0,408 | 0,007 | Tspyl2 | ENSMUSG00000020185 | -0,906 | 0,005 | E2f7 |
| ENSMUSG00000003546 | -0,409 | 0,000 | Klc4 | ENSMUSG00000020897 | -0,907 | 0,000 | Aurkb |
| ENSMUSG00000033763 | -0,409 | 0,049 | Mtss1l | ENSMUSG00000005410 | -0,907 | 0,000 | Mcm5 |
| ENSMUSG00000065979 | -0,409 | 0,000 | Cpped1 | ENSMUSG00000033207 | -0,907 | 0,002 | Mamdc2 |
| ENSMUSG00000005362 | -0,409 | 0,047 | Crbn | ENSMUSG00000041431 | -0,907 | 0,000 | Ccnb1 |
| ENSMUSG00000037461 | -0,410 | 0,001 | Ints7 | ENSMUSG00000029070 | -0,908 | 0,001 | Mxra8 |
| ENSMUSG00000028101 | -0,410 | 0,000 | Pias3 | ENSMUSG00000079598 | -0,909 | 0,046 | Clec2l |

| | | | | | |
|---|---|---|---|---|---|
| ENSMUSG00000021713 | -0,410 | 0,020 Ppwd1 | ENSMUSG00000035413 | -0,911 | 0,000 Tmem98 |
| ENSMUSG00000037419 | -0,412 | 0,010 Endod1 | ENSMUSG00000031258 | -0,913 | 0,045 Xkrx |
| ENSMUSG00000033713 | -0,412 | 0,005 Foxn3 | ENSMUSG00000037544 | -0,916 | 0,000 Dlgap5 |
| ENSMUSG00000040327 | -0,414 | 0,001 Cul9 | ENSMUSG00000020649 | -0,916 | 0,000 Rrm2 |
| ENSMUSG00000038481 | -0,414 | 0,011 Cdk19 | ENSMUSG00000056367 | -0,918 | 0,000 Actr3b |
| ENSMUSG00000032842 | -0,414 | 0,011 Abcc10 | ENSMUSG00000040133 | -0,919 | 0,041 Gpr176 |
| ENSMUSG00000024018 | -0,414 | 0,005 Ccdc167 | ENSMUSG00000002835 | -0,919 | 0,000 Chaf1a |
| ENSMUSG00000034685 | -0,414 | 0,013 Fam171a2 | ENSMUSG00000046814 | -0,920 | 0,029 Gchfr |
| ENSMUSG00000021719 | -0,415 | 0,018 Rgs7bp | ENSMUSG00000067714 | -0,920 | 0,017 Lpar5 |
| ENSMUSG00000029163 | -0,415 | 0,019 Emilin1 | ENSMUSG00000022360 | -0,921 | 0,000 Atad2 |
| ENSMUSG00000018651 | -0,415 | 0,000 Tada2a | ENSMUSG00000039814 | -0,923 | 0,001 Xkr5 |
| ENSMUSG00000054717 | -0,416 | 0,016 Hmgb2 | ENSMUSG00000033752 | -0,924 | 0,044 Mnd1 |
| ENSMUSG00000053436 | -0,416 | 0,000 Mapk14 | ENSMUSG00000028278 | -0,925 | 0,000 Rragd |
| ENSMUSG00000071337 | -0,417 | 0,013 Tia1 | ENSMUSG00000063810 | -0,925 | 0,018 Alms1 |
| ENSMUSG00000109061 | -0,417 | 0,047 Map2k7 | ENSMUSG00000025586 | -0,927 | 0,022 Cpeb1 |
| ENSMUSG00000040618 | -0,417 | 0,012 Pck2 | ENSMUSG00000027326 | -0,928 | 0,023 Knl1 |
| ENSMUSG00000087174 | -0,417 | 0,002 5530601H04Rik | ENSMUSG00000075502 | -0,930 | 0,040 Kbtbd6 |
| ENSMUSG00000027454 | -0,417 | 0,016 Gins1 | ENSMUSG00000044770 | -0,931 | 0,006 Scml4 |
| ENSMUSG00000048965 | -0,418 | 0,035 Mrgpre | ENSMUSG00000033952 | -0,933 | 0,022 Aspm |
| ENSMUSG00000029591 | -0,418 | 0,045 Ung | ENSMUSG00000019773 | -0,933 | 0,000 Fbxo5 |
| ENSMUSG00000066838 | -0,418 | 0,017 Zfp772 | ENSMUSG00000034353 | -0,933 | 0,000 Ramp1 |
| ENSMUSG00000029191 | -0,419 | 0,000 Rfc1 | ENSMUSG00000025395 | -0,933 | 0,000 Prim1 |
| ENSMUSG00000036555 | -0,419 | 0,001 Iqce | ENSMUSG00000055116 | -0,935 | 0,000 Arntl |
| ENSMUSG00000028007 | -0,420 | 0,001 Snx7 | ENSMUSG00000029414 | -0,935 | 0,000 Kntc1 |
| ENSMUSG00000052056 | -0,421 | 0,047 Zfp217 | ENSMUSG00000036875 | -0,936 | 0,000 Dna2 |
| ENSMUSG00000002249 | -0,421 | 0,002 Tead3 | ENSMUSG00000053040 | -0,936 | 0,000 Aph1c |
| ENSMUSG00000025816 | -0,423 | 0,000 Sec61a2 | ENSMUSG00000047875 | -0,938 | 0,004 Gpr157 |
| ENSMUSG00000043557 | -0,423 | 0,035 Mdga1 | ENSMUSG00000038379 | -0,938 | 0,000 Ttk |
| ENSMUSG00000034121 | -0,424 | 0,000 Mks1 | ENSMUSG00000046591 | -0,944 | 0,000 Ticrr |
| ENSMUSG00000061536 | -0,425 | 0,027 Sec22c | ENSMUSG00000031262 | -0,950 | 0,000 Cenpi |
| ENSMUSG00000038042 | -0,425 | 0,000 Ptpdc1 | ENSMUSG00000020052 | -0,950 | 0,022 Ascl1 |
| ENSMUSG00000022853 | -0,425 | 0,016 Ehhadh | ENSMUSG00000028212 | -0,953 | 0,000 Ccne2 |
| ENSMUSG00000021185 | -0,426 | 0,004 Dglucy | ENSMUSG00000026779 | -0,954 | 0,000 Mastl |
| ENSMUSG00000039936 | -0,426 | 0,007 Pik3cd | ENSMUSG00000037474 | -0,954 | 0,000 Dtl |
| ENSMUSG00000061288 | -0,427 | 0,001 Taok3 | ENSMUSG00000037991 | -0,958 | 0,005 Rmi2 |
| ENSMUSG00000002870 | -0,427 | 0,005 Mcm2 | ENSMUSG00000030559 | -0,961 | 0,018 Rab38 |
| ENSMUSG00000063445 | -0,428 | 0,012 Nmral1 | ENSMUSG00000019027 | -0,962 | 0,019 Dnah1 |
| ENSMUSG00000071604 | -0,428 | 0,015 Fam189a2 | ENSMUSG00000045273 | -0,963 | 0,000 Cenph |
| ENSMUSG00000043015 | -0,428 | 0,000 Nemp2 | ENSMUSG00000032377 | -0,966 | 0,026 Plscr4 |
| ENSMUSG00000022421 | -0,428 | 0,020 Nptxr | ENSMUSG00000054134 | -0,968 | 0,010 Umodl1 |
| ENSMUSG00000110980 | -0,429 | 0,042 AC122428.2 | ENSMUSG00000031847 | -0,968 | 0,000 1700030J22Rik |
| ENSMUSG00000052911 | -0,430 | 0,000 Lamb2 | ENSMUSG00000078773 | -0,968 | 0,000 Rad54b |
| ENSMUSG00000028456 | -0,430 | 0,024 Unc13b | ENSMUSG00000050232 | -0,969 | 0,023 Cxcr3 |
| ENSMUSG00000056394 | -0,430 | 0,003 Lig1 | ENSMUSG00000051378 | -0,972 | 0,000 Kif18b |
| ENSMUSG00000047342 | -0,430 | 0,017 Zfp286 | ENSMUSG00000006219 | -0,974 | 0,001 Fblim1 |
| ENSMUSG00000017119 | -0,430 | 0,002 Nbr1 | ENSMUSG00000034329 | -0,978 | 0,000 Brip1 |
| ENSMUSG00000023908 | -0,430 | 0,020 Pkmyt1 | ENSMUSG00000041957 | -0,981 | 0,004 Pkp2 |

| | | | | | | | |
|---|---|---|---|---|---|---|---|
| ENSMUSG00000057229 | -0,431 | 0,002 | Dmac2 | ENSMUSG00000026196 | -0,981 | 0,001 | Bard1 |
| ENSMUSG00000022673 | -0,431 | 0,003 | Mcm4 | ENSMUSG00000033182 | -0,984 | 0,023 | Kbtbd12 |
| ENSMUSG00000029863 | -0,432 | 0,021 | Casp2 | ENSMUSG00000069743 | -0,988 | 0,002 | Zfp820 |
| ENSMUSG00000099481 | -0,433 | 0,000 | Xndc1 | ENSMUSG00000032313 | -0,988 | 0,022 | Tmem266 |
| ENSMUSG00000028199 | -0,433 | 0,001 | Cryz | ENSMUSG00000028551 | -0,992 | 0,000 | Cdkn2c |
| ENSMUSG00000051343 | -0,433 | 0,011 | Rab11fip5 | ENSMUSG00000021965 | -0,993 | 0,000 | Ska3 |
| ENSMUSG00000028517 | -0,433 | 0,012 | Plpp3 | ENSMUSG00000029283 | -0,993 | 0,000 | Cdc7 |
| ENSMUSG00000057315 | -0,433 | 0,002 | Arhgap24 | ENSMUSG00000031661 | -1,004 | 0,000 | Nkd1 |
| ENSMUSG00000029279 | -0,434 | 0,020 | Brdt | ENSMUSG00000038319 | -1,012 | 0,000 | Kcnh2 |
| ENSMUSG00000066357 | -0,434 | 0,000 | Wdr6 | ENSMUSG00000022945 | -1,014 | 0,000 | Chaf1b |
| ENSMUSG00000032913 | -0,435 | 0,029 | Lrig2 | ENSMUSG00000048096 | -1,020 | 0,049 | Lmod1 |
| ENSMUSG00000020156 | -0,435 | 0,003 | Mum1 | ENSMUSG00000026429 | -1,031 | 0,000 | Ube2t |
| ENSMUSG00000024240 | -0,436 | 0,003 | Epc1 | ENSMUSG00000022519 | -1,032 | 0,000 | Srl |
| ENSMUSG00000035958 | -0,436 | 0,019 | Tdp2 | ENSMUSG00000061175 | -1,036 | 0,002 | Fnip2 |
| ENSMUSG00000028035 | -0,437 | 0,002 | Dnajb4 | ENSMUSG00000039748 | -1,041 | 0,000 | Exo1 |
| ENSMUSG00000034639 | -0,437 | 0,034 | Setmar | ENSMUSG00000022332 | -1,044 | 0,000 | Khdrbs3 |
| ENSMUSG00000069255 | -0,437 | 0,000 | Dusp22 | ENSMUSG00000051220 | -1,050 | 0,000 | Ercc6l |
| ENSMUSG00000029270 | -0,437 | 0,000 | Fam69a | ENSMUSG00000025574 | -1,054 | 0,000 | Tk1 |
| ENSMUSG00000035877 | -0,438 | 0,001 | Zhx3 | ENSMUSG00000034883 | -1,055 | 0,010 | Lrr1 |
| ENSMUSG00000038860 | -0,438 | 0,001 | Garnl3 | ENSMUSG00000023968 | -1,055 | 0,045 | Crip3 |
| ENSMUSG00000019467 | -0,439 | 0,013 | Arhgef25 | ENSMUSG00000022357 | -1,058 | 0,007 | Klhl38 |
| ENSMUSG00000051721 | -0,439 | 0,012 | Wdcp | ENSMUSG00000028702 | -1,059 | 0,000 | Rad54l |
| ENSMUSG00000092356 | -0,439 | 0,010 | Gm20532 | ENSMUSG00000017146 | -1,060 | 0,000 | Brca1 |
| ENSMUSG00000005198 | -0,440 | 0,039 | Polr2a | ENSMUSG00000024600 | -1,066 | 0,003 | Slc27a6 |
| ENSMUSG00000027244 | -0,440 | 0,000 | Atg13 | ENSMUSG00000061577 | -1,068 | 0,023 | Adgrg5 |
| ENSMUSG00000034216 | -0,440 | 0,000 | Vps18 | ENSMUSG00000039994 | -1,077 | 0,000 | Timeless |
| ENSMUSG00000050605 | -0,440 | 0,011 | Zfp61 | ENSMUSG00000051606 | -1,078 | 0,001 | 2010001K21Rik |
| ENSMUSG00000038368 | -0,441 | 0,001 | Focad | ENSMUSG00000033033 | -1,080 | 0,000 | Calhm2 |
| ENSMUSG00000040687 | -0,441 | 0,000 | Madd | ENSMUSG00000068744 | -1,083 | 0,000 | Psrc1 |
| ENSMUSG00000032737 | -0,441 | 0,005 | Inppl1 | ENSMUSG00000006241 | -1,084 | 0,013 | Ccdc159 |
| ENSMUSG00000028933 | -0,442 | 0,000 | Xrcc2 | ENSMUSG00000044303 | -1,087 | 0,025 | Cdkn2a |
| ENSMUSG00000031770 | -0,442 | 0,004 | Herpud1 | ENSMUSG00000030670 | -1,088 | 0,010 | Cyp2r1 |
| ENSMUSG00000039050 | -0,442 | 0,000 | Osbpl2 | ENSMUSG00000034303 | -1,092 | 0,007 | Ccdc15 |
| ENSMUSG00000035936 | -0,442 | 0,023 | Aldh5a1 | ENSMUSG00000086040 | -1,094 | 0,032 | Wipf3 |
| ENSMUSG00000033762 | -0,443 | 0,024 | Recql4 | ENSMUSG00000006362 | -1,101 | 0,000 | Cbfa2t3 |
| ENSMUSG00000043895 | -0,443 | 0,000 | S1pr2 | ENSMUSG00000079502 | -1,101 | 0,023 | Cfap77 |
| ENSMUSG00000035401 | -0,443 | 0,000 | Emsy | ENSMUSG00000042707 | -1,109 | 0,001 | Dnali1 |
| ENSMUSG00000053470 | -0,443 | 0,017 | Kdm3a | ENSMUSG00000026274 | -1,110 | 0,000 | Pask |
| ENSMUSG00000022679 | -0,444 | 0,019 | Mpv17l | ENSMUSG00000057604 | -1,110 | 0,000 | Lmcd1 |
| ENSMUSG00000009905 | -0,444 | 0,000 | Kdsr | ENSMUSG00000110537 | -1,111 | 0,001 | Gm4316 |
| ENSMUSG00000041966 | -0,445 | 0,009 | Dcaf17 | ENSMUSG00000032783 | -1,114 | 0,000 | Troap |
| ENSMUSG00000045374 | -0,445 | 0,008 | Wdr81 | ENSMUSG00000019982 | -1,114 | 0,000 | Myb |
| ENSMUSG00000040557 | -0,445 | 0,002 | Mettl27 | ENSMUSG00000040710 | -1,119 | 0,003 | St8sia4 |
| ENSMUSG00000024665 | -0,446 | 0,028 | Fads2 | ENSMUSG00000007080 | -1,122 | 0,000 | Pole |
| ENSMUSG00000041859 | -0,447 | 0,001 | Mcm3 | ENSMUSG00000034773 | -1,125 | 0,000 | BC030867 |
| ENSMUSG00000029998 | -0,447 | 0,004 | Pcyox1 | ENSMUSG00000026589 | -1,126 | 0,002 | Sec16b |
| ENSMUSG00000050530 | -0,447 | 0,005 | Fam171a1 | ENSMUSG00000097429 | -1,127 | 0,008 | Gm26520 |

| | | | | | |
|---|---|---|---|---|---|
| ENSMUSG00000022881 | -0,448 | 0,019 Rfc4 | ENSMUSG00000009418 | -1,145 | 0,000 Nav1 |
| ENSMUSG00000029366 | -0,448 | 0,032 Dck | ENSMUSG00000026304 | -1,145 | 0,010 Rab17 |
| ENSMUSG00000025026 | -0,448 | 0,015 Add3 | ENSMUSG00000053706 | -1,153 | 0,012 B430305J03Rik |
| ENSMUSG00000028256 | -0,448 | 0,000 Odf2l | ENSMUSG00000031554 | -1,159 | 0,012 Adam5 |
| ENSMUSG00000042063 | -0,448 | 0,039 Zfp386 | ENSMUSG00000026809 | -1,166 | 0,001 Spaca9 |
| ENSMUSG00000033717 | -0,449 | 0,011 Adra2a | ENSMUSG00000041361 | -1,167 | 0,003 Myzap |
| ENSMUSG00000054252 | -0,449 | 0,038 Fgfr3 | ENSMUSG00000072980 | -1,170 | 0,000 Oip5 |
| ENSMUSG00000024462 | -0,449 | 0,035 Gabbr1 | ENSMUSG00000035681 | -1,178 | 0,015 Kcnc2 |
| ENSMUSG00000113204 | -0,450 | 0,010 Gm46430 | ENSMUSG00000019851 | -1,180 | 0,001 Perp |
| ENSMUSG00000005682 | -0,450 | 0,001 Pan2 | ENSMUSG00000028392 | -1,182 | 0,005 Bspry |
| ENSMUSG00000040746 | -0,450 | 0,002 Rnf167 | ENSMUSG00000081683 | -1,182 | 0,000 Fzd10 |
| ENSMUSG00000032352 | -0,451 | 0,027 Lrrc1 | ENSMUSG00000042489 | -1,184 | 0,000 Clspn |
| ENSMUSG00000096687 | -0,451 | 0,003 Mfsd4b4 | ENSMUSG00000046179 | -1,190 | 0,000 E2f8 |
| ENSMUSG00000050666 | -0,452 | 0,048 Vstm4 | ENSMUSG00000026669 | -1,198 | 0,000 Mcm10 |
| ENSMUSG00000063145 | -0,452 | 0,001 Bbs5 | ENSMUSG00000030708 | -1,202 | 0,000 Dnajb13 |
| ENSMUSG00000042606 | -0,452 | 0,002 Hirip3 | ENSMUSG00000018427 | -1,218 | 0,000 Ypel2 |
| ENSMUSG00000039473 | -0,452 | 0,006 Ubn1 | ENSMUSG00000022676 | -1,225 | 0,015 Snai2 |
| ENSMUSG00000004565 | -0,453 | 0,000 Pnpla6 | ENSMUSG00000086015 | -1,237 | 0,013 4833417C18Rik |
| ENSMUSG00000000078 | -0,453 | 0,032 Klf6 | ENSMUSG00000042988 | -1,242 | 0,000 Notum |
| ENSMUSG00000026991 | -0,453 | 0,003 Pkp4 | ENSMUSG00000052374 | -1,247 | 0,021 Actn2 |
| ENSMUSG00000029676 | -0,453 | 0,000 Pot1a | ENSMUSG00000043020 | -1,261 | 0,001 Wdr63 |
| ENSMUSG00000025203 | -0,454 | 0,039 Scd2 | ENSMUSG00000097519 | -1,269 | 0,004 4930558J18Rik |
| ENSMUSG00000037366 | -0,455 | 0,000 Pafah2 | ENSMUSG00000025504 | -1,282 | 0,004 Eps8l2 |
| ENSMUSG00000021306 | -0,455 | 0,005 Gpr137b | ENSMUSG00000024990 | -1,284 | 0,028 Rbp4 |
| ENSMUSG00000048109 | -0,456 | 0,013 Rbm15 | ENSMUSG00000025916 | -1,306 | 0,029 Ppp1r42 |
| ENSMUSG00000026017 | -0,457 | 0,028 Carf | ENSMUSG00000042498 | -1,311 | 0,029 D330045A20Rik |
| ENSMUSG00000035944 | -0,457 | 0,000 Ttc38 | ENSMUSG00000020151 | -1,314 | 0,003 Ptprr |
| ENSMUSG00000020290 | -0,457 | 0,029 Xpo1 | ENSMUSG00000026870 | -1,324 | 0,024 Cutal |
| ENSMUSG00000029647 | -0,458 | 0,002 Pan3 | ENSMUSG00000045498 | -1,336 | 0,019 Pcdhb3 |
| ENSMUSG00000024206 | -0,459 | 0,037 Rfx2 | ENSMUSG00000039252 | -1,343 | 0,004 Lgi2 |
| ENSMUSG00000022840 | -0,459 | 0,033 Adcy5 | ENSMUSG00000022780 | -1,346 | 0,033 Meltf |
| ENSMUSG00000042202 | -0,460 | 0,009 Slc35e2 | ENSMUSG00000070583 | -1,348 | 0,039 Fv1 |
| ENSMUSG00000023977 | -0,460 | 0,022 Ubr2 | ENSMUSG00000026778 | -1,353 | 0,000 Prkcq |
| ENSMUSG00000033446 | -0,460 | 0,027 Lpar6 | ENSMUSG00000105954 | -1,354 | 0,001 Gm42793 |
| ENSMUSG00000053799 | -0,461 | 0,006 Exoc6 | ENSMUSG00000036186 | -1,355 | 0,000 Fam69b |
| ENSMUSG00000024975 | -0,461 | 0,000 Pdcd4 | ENSMUSG00000037846 | -1,368 | 0,000 Rtkn2 |
| ENSMUSG00000064128 | -0,461 | 0,002 Cenpj | ENSMUSG00000046818 | -1,384 | 0,000 Ddit4l |
| ENSMUSG00000039089 | -0,462 | 0,010 L3mbtl3 | ENSMUSG00000017400 | -1,427 | 0,001 Stac2 |
| ENSMUSG00000062822 | -0,463 | 0,001 4833420G17Rik | ENSMUSG00000041479 | -1,427 | 0,000 Syt15 |
| ENSMUSG00000025758 | -0,464 | 0,030 Plk4 | ENSMUSG00000067547 | -1,428 | 0,041 Gm7666 |
| ENSMUSG00000040818 | -0,464 | 0,010 Dennd6a | ENSMUSG00000081600 | -1,461 | 0,022 Gm12286 |
| ENSMUSG00000034487 | -0,465 | 0,000 Kdelc2 | ENSMUSG00000111463 | -1,507 | 0,015 AC027700.1 |
| ENSMUSG00000038205 | -0,466 | 0,002 Prkab2 | ENSMUSG00000040231 | -1,538 | 0,022 Syngr4 |
| ENSMUSG00000031974 | -0,467 | 0,002 Abcb10 | ENSMUSG00000022262 | -1,541 | 0,025 Dnah5 |
| ENSMUSG00000050910 | -0,468 | 0,017 Cdr2l | ENSMUSG00000024697 | -1,542 | 0,013 Gna14 |
| ENSMUSG00000031767 | -0,468 | 0,028 Nudt7 | ENSMUSG00000090291 | -1,563 | 0,002 Lrrc10b |
| ENSMUSG00000057193 | -0,468 | 0,002 Slc44a2 | ENSMUSG00000021678 | -1,563 | 0,013 F2rl1 |

| | | | | | | | |
|---|---|---|---|---|---|---|---|
| ENSMUSG00000039787 | -0,469 | 0,001 | Cercam | ENSMUSG00000045259 | -1,563 | 0,014 | Klhdc9 |
| ENSMUSG00000068551 | -0,469 | 0,000 | Zfp467 | ENSMUSG00000005320 | -1,580 | 0,002 | Fgfr4 |
| ENSMUSG00000025277 | -0,469 | 0,000 | Abhd6 | ENSMUSG00000026890 | -1,597 | 0,000 | Lhx6 |
| ENSMUSG00000020070 | -0,469 | 0,031 | Rufy2 | ENSMUSG00000108841 | -1,597 | 0,015 | Frmpd2 |
| ENSMUSG00000033565 | -0,469 | 0,009 | Rbfox2 | ENSMUSG00000058743 | -1,620 | 0,000 | Kcnj14 |
| ENSMUSG00000024897 | -0,470 | 0,038 | Apba1 | ENSMUSG00000043719 | -1,620 | 0,004 | Col6a6 |
| ENSMUSG00000049588 | -0,470 | 0,039 | Ccdc69 | ENSMUSG00000038756 | -1,645 | 0,010 | Ttll6 |
| ENSMUSG00000048232 | -0,470 | 0,000 | Fbxo10 | ENSMUSG00000018126 | -1,647 | 0,000 | Baiap2l2 |
| ENSMUSG00000042213 | -0,470 | 0,040 | Zfand4 | ENSMUSG00000046480 | -1,662 | 0,006 | Scn4b |
| ENSMUSG00000032121 | -0,471 | 0,002 | Tmem218 | ENSMUSG00000045362 | -1,663 | 0,016 | Tnfrsf26 |
| ENSMUSG00000029385 | -0,471 | 0,007 | Ccng2 | ENSMUSG00000085555 | -1,682 | 0,048 | 2610035F20Rik |
| ENSMUSG00000026317 | -0,472 | 0,001 | Cln8 | ENSMUSG00000049872 | -1,703 | 0,003 | Fam26e |
| ENSMUSG00000021481 | -0,472 | 0,000 | Zfp346 | ENSMUSG00000035551 | -1,714 | 0,047 | Igfbpl1 |
| ENSMUSG00000031145 | -0,472 | 0,003 | Prickle3 | ENSMUSG00000097760 | -1,714 | 0,019 | 6030442K20Rik |
| ENSMUSG00000029521 | -0,472 | 0,001 | Chek2 | ENSMUSG00000039313 | -1,721 | 0,000 | AF529169 |
| ENSMUSG00000047003 | -0,473 | 0,036 | Zfp41 | ENSMUSG00000075256 | -1,814 | 0,003 | Cerkl |
| ENSMUSG00000019303 | -0,473 | 0,024 | Psmc3ip | ENSMUSG00000104347 | -1,861 | 0,026 | Gm37689 |
| ENSMUSG00000020130 | -0,473 | 0,010 | Tbc1d15 | ENSMUSG00000108778 | -1,890 | 0,020 | Gm20083 |
| ENSMUSG00000021120 | -0,473 | 0,000 | Pigh | ENSMUSG00000090399 | -1,898 | 0,001 | Gm38399 |
| ENSMUSG00000048271 | -0,474 | 0,045 | Rbm33 | ENSMUSG00000074461 | -1,954 | 0,022 | Gm10699 |
| ENSMUSG00000021806 | -0,474 | 0,015 | Nid2 | ENSMUSG00000033805 | -1,969 | 0,030 | Ephx4 |
| ENSMUSG00000067199 | -0,474 | 0,036 | Frat1 | ENSMUSG00000106590 | -2,010 | 0,015 | Gm42879 |
| ENSMUSG00000062198 | -0,474 | 0,012 | 2700097O09Rik | ENSMUSG00000024517 | -2,064 | 0,027 | Grp |
| ENSMUSG00000021597 | -0,474 | 0,016 | Slf1 | ENSMUSG00000046321 | -2,102 | 0,000 | Hs3st2 |
| ENSMUSG00000000282 | -0,476 | 0,003 | Mnt | ENSMUSG00000032291 | -2,165 | 0,001 | Crabp1 |
| ENSMUSG00000037466 | -0,476 | 0,005 | Tedc1 | ENSMUSG00000115388 | -2,217 | 0,031 | EPPK1 |
| ENSMUSG00000032594 | -0,477 | 0,000 | Ip6k1 | ENSMUSG00000112808 | -2,242 | 0,000 | Gm4739 |
| ENSMUSG00000051224 | -0,477 | 0,032 | Tceanc | ENSMUSG00000027224 | -2,429 | 0,047 | Duoxa1 |
| ENSMUSG00000022021 | -0,477 | 0,014 | Diaph3 | ENSMUSG00000014453 | -2,451 | 0,010 | Blk |
| ENSMUSG00000063605 | -0,478 | 0,011 | Ccdc102a | ENSMUSG00000026012 | -2,486 | 0,000 | Cd28 |
| ENSMUSG00000049411 | -0,478 | 0,005 | Tmem241 | ENSMUSG00000085783 | -2,546 | 0,008 | Gm9816 |
| ENSMUSG00000040435 | -0,478 | 0,003 | Ppp1r15a | ENSMUSG00000018486 | -2,674 | 0,012 | Wnt9b |
| ENSMUSG00000021250 | -0,479 | 0,043 | Fos | ENSMUSG00000109896 | -2,798 | 0,017 | Gm31409 |
| ENSMUSG00000082163 | -0,479 | 0,011 | Gm14276 | ENSMUSG00000038775 | -3,000 | 0,000 | Vill |
| ENSMUSG00000022799 | -0,480 | 0,041 | Arhgap31 | ENSMUSG00000029431 | -3,005 | 0,012 | B3gnt4 |
| ENSMUSG00000037885 | -0,480 | 0,019 | Stk35 | ENSMUSG00000085882 | -3,451 | 0,004 | 2610507I01Rik |
| ENSMUSG00000030522 | -0,481 | 0,001 | Mtmr10 | ENSMUSG00000039628 | -3,605 | 0,012 | Hs3st6 |
| ENSMUSG00000048458 | -0,481 | 0,017 | Fam212b | ENSMUSG00000022342 | -4,357 | 0,034 | Kcnv1 |
| ENSMUSG00000002814 | -0,483 | 0,001 | Top3a | ENSMUSG00000021590 | -5,224 | 0,022 | Spata9 |
| ENSMUSG00000061882 | -0,484 | 0,050 | Ccdc62 | | | | |
| ENSMUSG00000055923 | -0,485 | 0,001 | Aasdh | | | | |
| ENSMUSG00000022701 | -0,485 | 0,000 | Ccdc191 | | | | |
| ENSMUSG00000058486 | -0,486 | 0,000 | Wdr91 | | | | |
| ENSMUSG00000026792 | -0,486 | 0,000 | Lrsam1 | | | | |
| ENSMUSG00000051790 | -0,486 | 0,004 | Nlgn2 | | | | |
| ENSMUSG00000034247 | -0,486 | 0,000 | Plekhm1 | | | | |
| ENSMUSG00000005615 | -0,486 | 0,002 | Pcyt1a | | | | |

| | | | |
|---|---|---|---|
| ENSMUSG00000019564 | -0,487 | 0,010 | Arid3a |
| ENSMUSG00000006850 | -0,487 | 0,000 | Tmco6 |
| ENSMUSG00000033902 | -0,487 | 0,009 | Mapkbp1 |
| ENSMUSG00000020868 | -0,487 | 0,000 | Xylt2 |
| ENSMUSG00000029769 | -0,487 | 0,015 | Ccdc136 |
| ENSMUSG00000057110 | -0,488 | 0,000 | Cntrl |
| ENSMUSG00000056919 | -0,488 | 0,012 | Cep162 |
| ENSMUSG00000032826 | -0,488 | 0,043 | Ank2 |
| ENSMUSG00000027959 | -0,488 | 0,008 | Sass6 |
| ENSMUSG00000036561 | -0,489 | 0,000 | Ppp6r2 |
| ENSMUSG00000022521 | -0,489 | 0,048 | Crebbp |
| ENSMUSG00000043487 | -0,489 | 0,013 | Acot6 |
| ENSMUSG00000047907 | -0,489 | 0,045 | Tshz2 |
| ENSMUSG00000058290 | -0,490 | 0,005 | Espl1 |
| ENSMUSG00000047789 | -0,490 | 0,000 | Slc38a9 |
| ENSMUSG00000024525 | -0,490 | 0,000 | Impa2 |
| ENSMUSG00000024151 | -0,490 | 0,000 | Msh2 |
| ENSMUSG00000028696 | -0,490 | 0,027 | Ipp |
| ENSMUSG00000022179 | -0,490 | 0,000 | 4931414P19Rik |
| ENSMUSG00000043262 | -0,491 | 0,033 | Uevld |
| ENSMUSG00000038685 | -0,491 | 0,000 | Rtel1 |
| ENSMUSG00000025049 | -0,492 | 0,002 | Taf5 |
| ENSMUSG00000022453 | -0,493 | 0,017 | Naga |
| ENSMUSG00000029687 | -0,493 | 0,000 | Ezh2 |
| ENSMUSG00000030641 | -0,495 | 0,020 | Ddias |
| ENSMUSG00000026708 | -0,496 | 0,009 | Cenpl |
| ENSMUSG00000025648 | -0,497 | 0,000 | Pfkfb4 |
| ENSMUSG00000038949 | -0,497 | 0,004 | Cnst |
| ENSMUSG00000035172 | -0,498 | 0,000 | Plekhh3 |
| ENSMUSG00000035597 | -0,498 | 0,019 | Prpf39 |
| ENSMUSG00000045790 | -0,500 | 0,028 | Ccdc149 |
| ENSMUSG00000042616 | -0,500 | 0,017 | Oscp1 |
| ENSMUSG00000026042 | -0,501 | 0,046 | Col5a2 |
| ENSMUSG00000036854 | -0,501 | 0,000 | Hspb6 |
| ENSMUSG00000073791 | -0,501 | 0,007 | Efcab7 |
| ENSMUSG00000005958 | -0,501 | 0,004 | Ephb3 |
| ENSMUSG00000005686 | -0,501 | 0,049 | Ampd3 |
| ENSMUSG00000027784 | -0,501 | 0,035 | Ppm1l |
| ENSMUSG00000039328 | -0,502 | 0,000 | Rnf122 |
| ENSMUSG00000039349 | -0,502 | 0,026 | C130074G19Rik |
| ENSMUSG00000035125 | -0,503 | 0,000 | Gcfc2 |
| ENSMUSG00000033016 | -0,503 | 0,013 | Nfatc1 |
| ENSMUSG00000035967 | -0,503 | 0,012 | Ints6l |
| ENSMUSG00000056367 | -0,503 | 0,020 | Actr3b |
| ENSMUSG00000051705 | -0,503 | 0,025 | Senp8 |
| ENSMUSG00000039686 | -0,503 | 0,000 | Zer1 |
| ENSMUSG00000006464 | -0,504 | 0,029 | Bbs1 |

| | | | |
|---|---|---|---|
| ENSMUSG00000029468 | -0,504 | 0,004 | P2rx7 |
| ENSMUSG00000026840 | -0,504 | 0,047 | Lamc3 |
| ENSMUSG00000041762 | -0,504 | 0,019 | Gpr155 |
| ENSMUSG00000021725 | -0,505 | 0,002 | Parp8 |
| ENSMUSG00000019823 | -0,506 | 0,000 | Mical1 |
| ENSMUSG00000038422 | -0,506 | 0,031 | Hdhd3 |
| ENSMUSG00000041775 | -0,506 | 0,008 | Mapk1ip1 |
| ENSMUSG00000033124 | -0,506 | 0,000 | Atg9a |
| ENSMUSG00000040084 | -0,507 | 0,045 | Bub1b |
| ENSMUSG00000058997 | -0,507 | 0,001 | Vwa8 |
| ENSMUSG00000026955 | -0,508 | 0,017 | Sapcd2 |
| ENSMUSG00000029363 | -0,508 | 0,001 | Rfc5 |
| ENSMUSG00000029910 | -0,508 | 0,000 | Mad2l1 |
| ENSMUSG00000048546 | -0,509 | 0,015 | Tob2 |
| ENSMUSG00000025265 | -0,510 | 0,002 | Fgd1 |
| ENSMUSG00000038047 | -0,510 | 0,018 | Haus6 |
| ENSMUSG00000020212 | -0,510 | 0,043 | Mdm1 |
| ENSMUSG00000020100 | -0,510 | 0,040 | Slc29a3 |
| ENSMUSG00000038593 | -0,511 | 0,000 | Tctn1 |
| ENSMUSG00000028133 | -0,511 | 0,003 | Rwdd3 |
| ENSMUSG00000006920 | -0,511 | 0,000 | Ezh1 |
| ENSMUSG00000037526 | -0,511 | 0,001 | Atg14 |
| ENSMUSG00000023186 | -0,512 | 0,050 | Vwa5a |
| ENSMUSG00000036864 | -0,512 | 0,000 | Proser3 |
| ENSMUSG00000051586 | -0,512 | 0,007 | Mical3 |
| ENSMUSG00000062312 | -0,512 | 0,034 | Erbb2 |
| ENSMUSG00000069206 | -0,512 | 0,032 | Zfp874a |
| ENSMUSG00000085241 | -0,512 | 0,018 | Snhg3 |
| ENSMUSG00000041328 | -0,513 | 0,028 | Pcf11 |
| ENSMUSG00000020937 | -0,514 | 0,036 | Plcd3 |
| ENSMUSG00000036882 | -0,514 | 0,001 | Arhgap33 |
| ENSMUSG00000046311 | -0,514 | 0,013 | Zfp62 |
| ENSMUSG00000036528 | -0,514 | 0,000 | Ppfibp2 |
| ENSMUSG00000037415 | -0,515 | 0,000 | Ranbp10 |
| ENSMUSG00000074364 | -0,515 | 0,008 | Ehd2 |
| ENSMUSG00000038215 | -0,515 | 0,000 | Cep44 |
| ENSMUSG00000040621 | -0,515 | 0,000 | Gemin8 |
| ENSMUSG00000047881 | -0,515 | 0,000 | Rell1 |
| ENSMUSG00000056019 | -0,516 | 0,018 | Zfp709 |
| ENSMUSG00000047786 | -0,516 | 0,016 | Lix1 |
| ENSMUSG00000043987 | -0,516 | 0,002 | Cep164 |
| ENSMUSG00000040272 | -0,516 | 0,000 | Accs |
| ENSMUSG00000028082 | -0,517 | 0,033 | Sh3d19 |
| ENSMUSG00000050022 | -0,517 | 0,003 | Amz1 |
| ENSMUSG00000039485 | -0,518 | 0,015 | Tspyl4 |
| ENSMUSG00000074785 | -0,518 | 0,044 | Plxnc1 |
| ENSMUSG00000073705 | -0,518 | 0,036 | Cenps |

| | | | |
|---|---|---|---|
| ENSMUSG00000020653 | -0,519 | 0,013 | Klf11 |
| ENSMUSG00000039157 | -0,520 | 0,002 | Fam102a |
| ENSMUSG00000041000 | -0,520 | 0,000 | Trim62 |
| ENSMUSG00000019992 | -0,520 | 0,042 | Mtfr2 |
| ENSMUSG00000001521 | -0,520 | 0,001 | Tulp3 |
| ENSMUSG00000028068 | -0,521 | 0,008 | Iqgap3 |
| ENSMUSG00000027381 | -0,521 | 0,003 | Bcl2l11 |
| ENSMUSG00000024866 | -0,521 | 0,019 | Acy3 |
| ENSMUSG00000025352 | -0,522 | 0,039 | Gdf11 |
| ENSMUSG00000029670 | -0,522 | 0,000 | Ing3 |
| ENSMUSG00000043885 | -0,523 | 0,045 | Slc36a4 |
| ENSMUSG00000044548 | -0,523 | 0,039 | Dact1 |
| ENSMUSG00000053985 | -0,524 | 0,005 | Zfp14 |
| ENSMUSG00000021895 | -0,525 | 0,012 | Arhgef3 |
| ENSMUSG00000023972 | -0,525 | 0,003 | Ptk7 |
| ENSMUSG00000028698 | -0,525 | 0,010 | Pik3r3 |
| ENSMUSG00000050855 | -0,526 | 0,011 | Zfp940 |
| ENSMUSG00000029730 | -0,526 | 0,000 | Mcm7 |
| ENSMUSG00000026098 | -0,526 | 0,001 | Pms1 |
| ENSMUSG00000073557 | -0,527 | 0,024 | Ppp1r12b |
| ENSMUSG00000107499 | -0,527 | 0,016 | Ccdc142 |
| ENSMUSG00000026333 | -0,528 | 0,008 | Gin1 |
| ENSMUSG00000040204 | -0,528 | 0,027 | Pclaf |
| ENSMUSG00000030257 | -0,528 | 0,027 | Srgap3 |
| ENSMUSG00000009145 | -0,529 | 0,028 | Dqx1 |
| ENSMUSG00000021514 | -0,529 | 0,003 | Zfp369 |
| ENSMUSG00000026409 | -0,529 | 0,035 | Pfkfb2 |
| ENSMUSG00000002475 | -0,529 | 0,001 | Abhd3 |
| ENSMUSG00000041935 | -0,530 | 0,028 | AW549877 |
| ENSMUSG00000060657 | -0,530 | 0,010 | Marf1 |
| ENSMUSG00000035266 | -0,530 | 0,000 | Helq |
| ENSMUSG00000054737 | -0,530 | 0,018 | Zfp182 |
| ENSMUSG00000039239 | -0,531 | 0,045 | Tgfb2 |
| ENSMUSG00000040312 | -0,531 | 0,001 | Cchcr1 |
| ENSMUSG00000071281 | -0,531 | 0,036 | Zfp65 |
| ENSMUSG00000041594 | -0,531 | 0,001 | Tmtc4 |
| ENSMUSG00000025907 | -0,532 | 0,005 | Rb1cc1 |
| ENSMUSG00000000693 | -0,532 | 0,000 | Loxl3 |
| ENSMUSG00000025348 | -0,533 | 0,009 | Itga7 |
| ENSMUSG00000078794 | -0,534 | 0,000 | Dact3 |
| ENSMUSG00000028469 | -0,534 | 0,017 | Npr2 |
| ENSMUSG00000027353 | -0,534 | 0,008 | Mcm8 |
| ENSMUSG00000028212 | -0,534 | 0,042 | Ccne2 |
| ENSMUSG00000039103 | -0,535 | 0,046 | Nexn |
| ENSMUSG00000042744 | -0,536 | 0,045 | Hectd4 |
| ENSMUSG00000000276 | -0,536 | 0,037 | Dgke |
| ENSMUSG00000027203 | -0,536 | 0,017 | Dut |

| | | | |
|---|---|---|---|
| ENSMUSG00000034311 | -0,536 | 0,020 | Kif4 |
| ENSMUSG00000028051 | -0,537 | 0,034 | Hcn3 |
| ENSMUSG00000112095 | -0,537 | 0,000 | A130077B15Rik |
| ENSMUSG00000031669 | -0,538 | 0,000 | Gins3 |
| ENSMUSG00000033790 | -0,539 | 0,012 | Tubgcp5 |
| ENSMUSG00000010051 | -0,539 | 0,038 | Hyal1 |
| ENSMUSG00000064105 | -0,539 | 0,000 | Cnnm2 |
| ENSMUSG00000032741 | -0,539 | 0,000 | Tpcn1 |
| ENSMUSG00000046688 | -0,539 | 0,004 | Tifa |
| ENSMUSG00000022994 | -0,540 | 0,000 | Adcy6 |
| ENSMUSG00000027331 | -0,540 | 0,040 | Knstrn |
| ENSMUSG00000026456 | -0,540 | 0,005 | Cyb5r1 |
| ENSMUSG00000076435 | -0,541 | 0,003 | Acsf2 |
| ENSMUSG00000022843 | -0,542 | 0,003 | Clcn2 |
| ENSMUSG00000037455 | -0,542 | 0,000 | Slc18b1 |
| ENSMUSG00000029401 | -0,542 | 0,026 | Rilpl2 |
| ENSMUSG00000023947 | -0,542 | 0,004 | Nfkbie |
| ENSMUSG00000059713 | -0,542 | 0,000 | Rcan3 |
| ENSMUSG00000007646 | -0,542 | 0,001 | Rad51c |
| ENSMUSG00000070425 | -0,544 | 0,000 | Xntrpc |
| ENSMUSG00000056832 | -0,544 | 0,049 | Ttc26 |
| ENSMUSG00000014850 | -0,544 | 0,000 | Msh3 |
| ENSMUSG00000030337 | -0,544 | 0,005 | Vamp1 |
| ENSMUSG00000024036 | -0,544 | 0,012 | Slc37a1 |
| ENSMUSG00000033306 | -0,545 | 0,025 | Lpp |
| ENSMUSG00000032652 | -0,545 | 0,000 | Crebl2 |
| ENSMUSG00000037151 | -0,547 | 0,001 | Lrrc20 |
| ENSMUSG00000046157 | -0,547 | 0,001 | Tmem229b |
| ENSMUSG00000055200 | -0,547 | 0,000 | Sertad3 |
| ENSMUSG00000089715 | -0,547 | 0,000 | Cbx6 |
| ENSMUSG00000030978 | -0,548 | 0,000 | Rrm1 |
| ENSMUSG00000031310 | -0,548 | 0,001 | Zmym3 |
| ENSMUSG00000022822 | -0,548 | 0,001 | Abcc5 |
| ENSMUSG00000039831 | -0,549 | 0,039 | Arhgap29 |
| ENSMUSG00000078495 | -0,549 | 0,016 | Zfp984 |
| ENSMUSG00000055897 | -0,549 | 0,003 | Ppp4r1l-ps |
| ENSMUSG00000072653 | -0,549 | 0,001 | Zfp783 |
| ENSMUSG00000037818 | -0,549 | 0,008 | Abhd18 |
| ENSMUSG00000027715 | -0,550 | 0,017 | Ccna2 |
| ENSMUSG00000032718 | -0,550 | 0,043 | Mansc1 |
| ENSMUSG00000028621 | -0,550 | 0,000 | Cyb5rl |
| ENSMUSG00000001138 | -0,550 | 0,000 | Cnnm3 |
| ENSMUSG00000033857 | -0,550 | 0,000 | Engase |
| ENSMUSG00000038010 | -0,551 | 0,008 | Ccdc138 |
| ENSMUSG00000060227 | -0,551 | 0,044 | Casc4 |
| ENSMUSG00000038126 | -0,551 | 0,018 | Mphosph9 |
| ENSMUSG00000032555 | -0,552 | 0,008 | Topbp1 |

| | | | |
|---|---|---|---|
| ENSMUSG00000027217 | -0,552 | 0,027 | Tspan18 |
| ENSMUSG00000000823 | -0,552 | 0,002 | Zfp512b |
| ENSMUSG00000059708 | -0,552 | 0,042 | Akap17b |
| ENSMUSG00000034522 | -0,552 | 0,001 | Zfp395 |
| ENSMUSG00000052917 | -0,552 | 0,031 | Senp7 |
| ENSMUSG00000017639 | -0,553 | 0,005 | Rab11fip4 |
| ENSMUSG00000043496 | -0,553 | 0,024 | Tril |
| ENSMUSG00000035547 | -0,554 | 0,002 | Capn5 |
| ENSMUSG00000030643 | -0,554 | 0,049 | Rab30 |
| ENSMUSG00000028088 | -0,554 | 0,001 | Fmo5 |
| ENSMUSG00000039853 | -0,555 | 0,023 | Trim14 |
| ENSMUSG00000021747 | -0,555 | 0,023 | 4930452B06Rik |
| ENSMUSG00000045467 | -0,556 | 0,036 | Ttll13 |
| ENSMUSG00000018931 | -0,556 | 0,000 | Natd1 |
| ENSMUSG00000001741 | -0,557 | 0,020 | Il16 |
| ENSMUSG00000032850 | -0,557 | 0,002 | Rnft2 |
| ENSMUSG00000004085 | -0,557 | 0,002 | Map3k20 |
| ENSMUSG00000027589 | -0,557 | 0,047 | Pcmtd2 |
| ENSMUSG00000022756 | -0,558 | 0,002 | Slc7a4 |
| ENSMUSG00000047804 | -0,558 | 0,019 | Akap10 |
| ENSMUSG00000005410 | -0,558 | 0,001 | Mcm5 |
| ENSMUSG00000027330 | -0,558 | 0,001 | Cdc25b |
| ENSMUSG00000032531 | -0,558 | 0,002 | Amotl2 |
| ENSMUSG00000028944 | -0,558 | 0,000 | Prkag2 |
| ENSMUSG00000015709 | -0,558 | 0,032 | Arnt2 |
| ENSMUSG00000024423 | -0,559 | 0,001 | Impact |
| ENSMUSG00000028613 | -0,559 | 0,044 | Lrp8 |
| ENSMUSG00000019947 | -0,559 | 0,032 | Arid5b |
| ENSMUSG00000046111 | -0,559 | 0,026 | Cep295 |
| ENSMUSG00000043252 | -0,560 | 0,008 | Tmem64 |
| ENSMUSG00000023505 | -0,560 | 0,024 | Cdca3 |
| ENSMUSG00000055799 | -0,560 | 0,003 | Tcf7l1 |
| ENSMUSG00000033594 | -0,560 | 0,007 | Spata2l |
| ENSMUSG00000029009 | -0,560 | 0,001 | Mthfr |
| ENSMUSG00000027469 | -0,561 | 0,008 | Tpx2 |
| ENSMUSG00000055963 | -0,562 | 0,014 | Triqk |
| ENSMUSG00000038618 | -0,562 | 0,004 | Rassf7 |
| ENSMUSG00000025656 | -0,563 | 0,007 | Arhgef9 |
| ENSMUSG00000021466 | -0,563 | 0,037 | Ptch1 |
| ENSMUSG00000039497 | -0,563 | 0,007 | Dse |
| ENSMUSG00000020228 | -0,563 | 0,000 | Helb |
| ENSMUSG00000024121 | -0,563 | 0,002 | Atp6v0c |
| ENSMUSG00000026383 | -0,564 | 0,049 | Epb41l5 |
| ENSMUSG00000033350 | -0,564 | 0,006 | Chst2 |
| ENSMUSG00000020808 | -0,564 | 0,018 | Pimreg |
| ENSMUSG00000035683 | -0,564 | 0,014 | Melk |
| ENSMUSG00000028364 | -0,564 | 0,041 | Tnc |

| | | | |
|---|---|---|---|
| ENSMUSG00000020541 | -0,566 | 0,007 | Tom1l1 |
| ENSMUSG00000057835 | -0,566 | 0,029 | Zfp119a |
| ENSMUSG00000062510 | -0,566 | 0,002 | Nsl1 |
| ENSMUSG00000053004 | -0,566 | 0,009 | Hrh1 |
| ENSMUSG00000057914 | -0,566 | 0,041 | Cacnb2 |
| ENSMUSG00000027457 | -0,567 | 0,025 | Snph |
| ENSMUSG00000025507 | -0,567 | 0,002 | Pidd1 |
| ENSMUSG00000023015 | -0,567 | 0,015 | Racgap1 |
| ENSMUSG00000054115 | -0,568 | 0,000 | Skp2 |
| ENSMUSG00000020900 | -0,570 | 0,029 | Myh10 |
| ENSMUSG00000022758 | -0,570 | 0,013 | P2rx6 |
| ENSMUSG00000034906 | -0,571 | 0,000 | Ncaph |
| ENSMUSG00000020592 | -0,572 | 0,003 | Sdc1 |
| ENSMUSG00000107879 | -0,573 | 0,005 | 9330102E08Rik |
| ENSMUSG00000032086 | -0,573 | 0,001 | Bace1 |
| ENSMUSG00000022788 | -0,573 | 0,016 | Fgd4 |
| ENSMUSG00000031133 | -0,573 | 0,003 | Arhgef6 |
| ENSMUSG00000070000 | -0,573 | 0,000 | Fcho1 |
| ENSMUSG00000034917 | -0,574 | 0,019 | Tjp3 |
| ENSMUSG00000020914 | -0,574 | 0,020 | Top2a |
| ENSMUSG00000068299 | -0,575 | 0,000 | Nat8f4 |
| ENSMUSG00000019813 | -0,575 | 0,000 | Cep57l1 |
| ENSMUSG00000087408 | -0,575 | 0,003 | Cers1 |
| ENSMUSG00000002346 | -0,575 | 0,001 | Slc25a42 |
| ENSMUSG00000078942 | -0,576 | 0,010 | Naip6 |
| ENSMUSG00000079304 | -0,577 | 0,020 | Tex52 |
| ENSMUSG00000049323 | -0,577 | 0,040 | Smcr8 |
| ENSMUSG00000044338 | -0,577 | 0,049 | Aplnr |
| ENSMUSG00000070808 | -0,579 | 0,001 | Bicra |
| ENSMUSG00000038576 | -0,580 | 0,010 | Susd4 |
| ENSMUSG00000043290 | -0,580 | 0,000 | Zfp784 |
| ENSMUSG00000032221 | -0,580 | 0,042 | Mns1 |
| ENSMUSG00000036986 | -0,580 | 0,001 | Pml |
| ENSMUSG00000026034 | -0,581 | 0,000 | Clk1 |
| ENSMUSG00000027490 | -0,581 | 0,000 | E2f1 |
| ENSMUSG00000037243 | -0,581 | 0,008 | Zfp692 |
| ENSMUSG00000020176 | -0,581 | 0,006 | Grb10 |
| ENSMUSG00000040606 | -0,581 | 0,031 | Kazn |
| ENSMUSG00000051098 | -0,582 | 0,018 | Mblac2 |
| ENSMUSG00000032194 | -0,582 | 0,001 | Kank2 |
| ENSMUSG00000014496 | -0,582 | 0,019 | Ankrd28 |
| ENSMUSG00000031478 | -0,582 | 0,000 | Nek3 |
| ENSMUSG00000041777 | -0,582 | 0,001 | Cir1 |
| ENSMUSG00000034317 | -0,583 | 0,001 | Trim59 |
| ENSMUSG00000042647 | -0,583 | 0,010 | Acad12 |
| ENSMUSG00000034379 | -0,583 | 0,007 | Wdr5b |
| ENSMUSG00000027669 | -0,584 | 0,000 | Gnb4 |

| | | | |
|---|---|---|---|
| ENSMUSG00000044350 | -0,584 | 0,010 | Lacc1 |
| ENSMUSG00000031756 | -0,585 | 0,001 | Cenpn |
| ENSMUSG00000050394 | -0,585 | 0,000 | Armcx6 |
| ENSMUSG00000074863 | -0,585 | 0,005 | Platr25 |
| ENSMUSG00000030279 | -0,585 | 0,000 | C2cd5 |
| ENSMUSG00000066647 | -0,587 | 0,006 | Gm5113 |
| ENSMUSG00000035759 | -0,588 | 0,031 | Bbs10 |
| ENSMUSG00000028445 | -0,588 | 0,030 | Enho |
| ENSMUSG00000022091 | -0,588 | 0,000 | Sorbs3 |
| ENSMUSG00000059146 | -0,588 | 0,031 | Ntrk3 |
| ENSMUSG00000021838 | -0,589 | 0,038 | Samd4 |
| ENSMUSG00000068036 | -0,589 | 0,034 | Afdn |
| ENSMUSG00000025241 | -0,589 | 0,043 | Fyco1 |
| ENSMUSG00000038943 | -0,590 | 0,003 | Prc1 |
| ENSMUSG00000020492 | -0,590 | 0,000 | Ska2 |
| ENSMUSG00000027641 | -0,590 | 0,046 | Rbl1 |
| ENSMUSG00000038370 | -0,591 | 0,027 | Pcp4l1 |
| ENSMUSG00000032586 | -0,591 | 0,000 | Traip |
| ENSMUSG00000037295 | -0,592 | 0,000 | Ldlrap1 |
| ENSMUSG00000006235 | -0,592 | 0,000 | Epor |
| ENSMUSG00000058070 | -0,593 | 0,008 | Eml1 |
| ENSMUSG00000028073 | -0,593 | 0,000 | Pear1 |
| ENSMUSG00000003031 | -0,593 | 0,000 | Cdkn1b |
| ENSMUSG00000026170 | -0,595 | 0,008 | Cyp27a1 |
| ENSMUSG00000054850 | -0,595 | 0,003 | Smim10l2a |
| ENSMUSG00000025092 | -0,595 | 0,009 | Hspa12a |
| ENSMUSG00000050379 | -0,595 | 0,000 | Sept6 |
| ENSMUSG00000093456 | -0,596 | 0,040 | Gm20662 |
| ENSMUSG00000034647 | -0,596 | 0,012 | Ankrd12 |
| ENSMUSG00000026556 | -0,596 | 0,002 | Vangl2 |
| ENSMUSG00000079038 | -0,597 | 0,023 | D130040H23Rik |
| ENSMUSG00000001300 | -0,597 | 0,023 | Efnb2 |
| ENSMUSG00000039852 | -0,597 | 0,007 | Rere |
| ENSMUSG00000022945 | -0,598 | 0,002 | Chaf1b |
| ENSMUSG00000033031 | -0,598 | 0,016 | C330027C09Rik |
| ENSMUSG00000023079 | -0,599 | 0,000 | Gtf2ird1 |
| ENSMUSG00000042156 | -0,600 | 0,001 | Dzip1 |
| ENSMUSG00000026489 | -0,601 | 0,000 | Coq8a |
| ENSMUSG00000060012 | -0,601 | 0,044 | Kif13b |
| ENSMUSG00000030630 | -0,601 | 0,018 | Fah |
| ENSMUSG00000025271 | -0,601 | 0,012 | Pfkfb1 |
| ENSMUSG00000099689 | -0,601 | 0,021 | Zfp383 |
| ENSMUSG00000020888 | -0,602 | 0,000 | Dvl2 |
| ENSMUSG00000024039 | -0,602 | 0,048 | Cbs |
| ENSMUSG00000054939 | -0,602 | 0,029 | Zfp174 |
| ENSMUSG00000063488 | -0,602 | 0,001 | Zkscan7 |
| ENSMUSG00000020034 | -0,603 | 0,000 | Tcp11l2 |

| | | | |
|---|---|---|---|
| ENSMUSG00000061353 | -0,603 | 0,001 | Cxcl12 |
| ENSMUSG00000054555 | -0,603 | 0,023 | Adam12 |
| ENSMUSG00000028980 | -0,604 | 0,000 | H6pd |
| ENSMUSG00000002771 | -0,606 | 0,022 | Grin2d |
| ENSMUSG00000053062 | -0,607 | 0,013 | Jam2 |
| ENSMUSG00000026211 | -0,607 | 0,026 | Obsl1 |
| ENSMUSG00000024867 | -0,608 | 0,012 | Pip5k1b |
| ENSMUSG00000067629 | -0,608 | 0,010 | Syngap1 |
| ENSMUSG00000037235 | -0,608 | 0,000 | Mxd4 |
| ENSMUSG00000027306 | -0,608 | 0,013 | Nusap1 |
| ENSMUSG00000022422 | -0,609 | 0,006 | Dscc1 |
| ENSMUSG00000056735 | -0,609 | 0,010 | A930024E05Rik |
| ENSMUSG00000035104 | -0,610 | 0,003 | Eva1a |
| ENSMUSG00000048327 | -0,610 | 0,005 | Ckap2l |
| ENSMUSG00000028496 | -0,610 | 0,001 | Mllt3 |
| ENSMUSG00000029177 | -0,611 | 0,032 | Cenpa |
| ENSMUSG00000028312 | -0,611 | 0,035 | Smc2 |
| ENSMUSG00000028989 | -0,611 | 0,016 | Angptl7 |
| ENSMUSG00000032411 | -0,612 | 0,000 | Tfdp2 |
| ENSMUSG00000061524 | -0,612 | 0,037 | Zic2 |
| ENSMUSG00000030231 | -0,612 | 0,001 | Plekha5 |
| ENSMUSG00000032198 | -0,612 | 0,000 | Dock6 |
| ENSMUSG00000031134 | -0,614 | 0,000 | Rbmx |
| ENSMUSG00000026944 | -0,614 | 0,002 | Abca2 |
| ENSMUSG00000022385 | -0,615 | 0,003 | Gtse1 |
| ENSMUSG00000000275 | -0,615 | 0,001 | Trim25 |
| ENSMUSG00000001151 | -0,616 | 0,002 | Pcnt |
| ENSMUSG00000059013 | -0,616 | 0,000 | Sh2d3c |
| ENSMUSG00000026360 | -0,617 | 0,000 | Rgs2 |
| ENSMUSG00000049521 | -0,617 | 0,001 | Cdc42ep1 |
| ENSMUSG00000057133 | -0,617 | 0,024 | Chd6 |
| ENSMUSG00000032750 | -0,617 | 0,009 | Gab3 |
| ENSMUSG00000050410 | -0,618 | 0,000 | Tcf19 |
| ENSMUSG00000097842 | -0,618 | 0,026 | 9330104G04Rik |
| ENSMUSG00000021009 | -0,618 | 0,002 | Ptpn21 |
| ENSMUSG00000025085 | -0,619 | 0,015 | Ablim1 |
| ENSMUSG00000030729 | -0,619 | 0,004 | Pgm2l1 |
| ENSMUSG00000023411 | -0,620 | 0,016 | Nfatc4 |
| ENSMUSG00000068115 | -0,621 | 0,000 | Ninl |
| ENSMUSG00000047153 | -0,621 | 0,000 | Khnyn |
| ENSMUSG00000041406 | -0,621 | 0,000 | BC055324 |
| ENSMUSG00000035356 | -0,621 | 0,010 | Nfkbiz |
| ENSMUSG00000045333 | -0,621 | 0,016 | Zfp423 |
| ENSMUSG00000031480 | -0,622 | 0,013 | Thsd1 |
| ENSMUSG00000059810 | -0,622 | 0,000 | Rgs3 |
| ENSMUSG00000097119 | -0,623 | 0,012 | B230354K17Rik |
| ENSMUSG00000078153 | -0,624 | 0,014 | Psme2b |

| | | | |
|---|---|---|---|
| ENSMUSG00000059839 | -0,624 | 0,001 | Zfp874b |
| ENSMUSG00000047409 | -0,624 | 0,000 | Ctdspl |
| ENSMUSG00000029851 | -0,625 | 0,044 | Tcaf2 |
| ENSMUSG00000037725 | -0,625 | 0,001 | Ckap2 |
| ENSMUSG00000006675 | -0,625 | 0,036 | P4htm |
| ENSMUSG00000024044 | -0,625 | 0,008 | Epb41l3 |
| ENSMUSG00000025812 | -0,625 | 0,038 | Pard3 |
| ENSMUSG00000071181 | -0,625 | 0,016 | 3830408C21Rik |
| ENSMUSG00000026676 | -0,625 | 0,015 | Ccdc3 |
| ENSMUSG00000006219 | -0,626 | 0,044 | Fblim1 |
| ENSMUSG00000038766 | -0,626 | 0,011 | Gabpb2 |
| ENSMUSG00000022322 | -0,626 | 0,004 | Shcbp1 |
| ENSMUSG00000006800 | -0,627 | 0,000 | Sulf2 |
| ENSMUSG00000042810 | -0,627 | 0,000 | Krba1 |
| ENSMUSG00000035455 | -0,627 | 0,004 | Fignl1 |
| ENSMUSG00000005949 | -0,627 | 0,000 | Ctns |
| ENSMUSG00000047115 | -0,629 | 0,024 | Fam221a |
| ENSMUSG00000022203 | -0,629 | 0,023 | Efs |
| ENSMUSG00000044317 | -0,629 | 0,005 | Gpr4 |
| ENSMUSG00000035878 | -0,629 | 0,008 | Hykk |
| ENSMUSG00000016087 | -0,630 | 0,042 | Fli1 |
| ENSMUSG00000078762 | -0,630 | 0,000 | Haus5 |
| ENSMUSG00000020744 | -0,630 | 0,000 | Slc25a19 |
| ENSMUSG00000022070 | -0,630 | 0,000 | Bora |
| ENSMUSG00000045751 | -0,631 | 0,000 | Mms22l |
| ENSMUSG00000066894 | -0,631 | 0,000 | Vsig10 |
| ENSMUSG00000100254 | -0,631 | 0,007 | Trpc2 |
| ENSMUSG00000097993 | -0,633 | 0,020 | Ptprv |
| ENSMUSG00000115022 | -0,633 | 0,000 | Hacl1 |
| ENSMUSG00000037572 | -0,634 | 0,000 | Wdhd1 |
| ENSMUSG00000030867 | -0,634 | 0,007 | Plk1 |
| ENSMUSG00000041132 | -0,634 | 0,000 | N4bp2l1 |
| ENSMUSG00000027698 | -0,635 | 0,000 | Nceh1 |
| ENSMUSG00000024773 | -0,637 | 0,000 | Atg2a |
| ENSMUSG00000042510 | -0,637 | 0,000 | AA986860 |
| ENSMUSG00000033596 | -0,638 | 0,000 | Rfwd3 |
| ENSMUSG00000038156 | -0,638 | 0,038 | Spon1 |
| ENSMUSG00000042320 | -0,638 | 0,005 | Prox2 |
| ENSMUSG00000032641 | -0,639 | 0,000 | Gpr19 |
| ENSMUSG00000025020 | -0,640 | 0,045 | Slit1 |
| ENSMUSG00000030766 | -0,640 | 0,000 | Arhgap17 |
| ENSMUSG00000036526 | -0,640 | 0,013 | Card11 |
| ENSMUSG00000052684 | -0,641 | 0,000 | Jun |
| ENSMUSG00000069727 | -0,642 | 0,020 | Zfp975 |
| ENSMUSG00000027827 | -0,642 | 0,049 | Kcnab1 |
| ENSMUSG00000038143 | -0,643 | 0,044 | Stox2 |
| ENSMUSG00000030213 | -0,644 | 0,007 | Atf7ip |

| | | | |
|---|---|---|---|
| ENSMUSG00000038644 | -0,644 | 0,000 | Pold1 |
| ENSMUSG00000030930 | -0,644 | 0,000 | Chst15 |
| ENSMUSG00000057722 | -0,644 | 0,028 | Lepr |
| ENSMUSG00000071064 | -0,645 | 0,000 | Zfp827 |
| ENSMUSG00000059851 | -0,645 | 0,000 | Kmt5c |
| ENSMUSG00000061751 | -0,645 | 0,014 | Kalrn |
| ENSMUSG00000031075 | -0,646 | 0,045 | Ano1 |
| ENSMUSG00000021782 | -0,646 | 0,034 | Dlg5 |
| ENSMUSG00000010064 | -0,646 | 0,000 | Slc38a3 |
| ENSMUSG00000044337 | -0,646 | 0,000 | Ackr3 |
| ENSMUSG00000087368 | -0,647 | 0,017 | BC065397 |
| ENSMUSG00000040594 | -0,647 | 0,018 | Ranbp17 |
| ENSMUSG00000002835 | -0,650 | 0,000 | Chaf1a |
| ENSMUSG00000078429 | -0,650 | 0,000 | Ctdsp2 |
| ENSMUSG00000014329 | -0,650 | 0,025 | Bicc1 |
| ENSMUSG00000046027 | -0,650 | 0,000 | Stard5 |
| ENSMUSG00000062866 | -0,651 | 0,001 | Phactr2 |
| ENSMUSG00000022364 | -0,651 | 0,000 | Tbc1d31 |
| ENSMUSG00000014303 | -0,651 | 0,000 | Glis2 |
| ENSMUSG00000020974 | -0,651 | 0,000 | Pole2 |
| ENSMUSG00000001665 | -0,651 | 0,000 | Gstt3 |
| ENSMUSG00000022177 | -0,652 | 0,001 | Haus4 |
| ENSMUSG00000028005 | -0,653 | 0,016 | Gucy1b3 |
| ENSMUSG00000006678 | -0,653 | 0,003 | Pola1 |
| ENSMUSG00000071192 | -0,654 | 0,046 | Wfikkn1 |
| ENSMUSG00000053647 | -0,655 | 0,047 | Gper1 |
| ENSMUSG00000040181 | -0,655 | 0,002 | Fmo1 |
| ENSMUSG00000022836 | -0,656 | 0,042 | Mylk |
| ENSMUSG00000024883 | -0,656 | 0,027 | Rin1 |
| ENSMUSG00000031666 | -0,657 | 0,001 | Rbl2 |
| ENSMUSG00000064080 | -0,657 | 0,028 | Fbln2 |
| ENSMUSG00000018809 | -0,659 | 0,000 | Smyd4 |
| ENSMUSG00000037754 | -0,659 | 0,030 | Ppp1r16b |
| ENSMUSG00000021884 | -0,659 | 0,000 | Hacl1 |
| ENSMUSG00000029086 | -0,660 | 0,000 | Prom1 |
| ENSMUSG00000042216 | -0,660 | 0,036 | Sgsm1 |
| ENSMUSG00000026080 | -0,661 | 0,000 | Chst10 |
| ENSMUSG00000025608 | -0,661 | 0,010 | Podxl |
| ENSMUSG00000054893 | -0,661 | 0,000 | Zfp667 |
| ENSMUSG00000000901 | -0,661 | 0,000 | Mmp11 |
| ENSMUSG00000049624 | -0,662 | 0,000 | Slc17a5 |
| ENSMUSG00000030309 | -0,662 | 0,000 | Caprin2 |
| ENSMUSG00000031379 | -0,662 | 0,018 | Pir |
| ENSMUSG00000079553 | -0,663 | 0,007 | Kifc1 |
| ENSMUSG00000029869 | -0,663 | 0,001 | Ephb6 |
| ENSMUSG00000051517 | -0,663 | 0,004 | Arhgef39 |
| ENSMUSG00000034349 | -0,664 | 0,000 | Smc4 |

| | | | |
|---|---|---|---|
| ENSMUSG00000035829 | -0,664 | 0,004 | Ppp1r26 |
| ENSMUSG00000033458 | -0,664 | 0,022 | Fan1 |
| ENSMUSG00000038252 | -0,664 | 0,000 | Ncapd2 |
| ENSMUSG00000050397 | -0,666 | 0,042 | Foxl2 |
| ENSMUSG00000029299 | -0,666 | 0,009 | Abcg3 |
| ENSMUSG00000097242 | -0,666 | 0,018 | Gm16907 |
| ENSMUSG00000030515 | -0,667 | 0,000 | Tarsl2 |
| ENSMUSG00000014786 | -0,668 | 0,000 | Slc9a5 |
| ENSMUSG00000024696 | -0,668 | 0,000 | Lpxn |
| ENSMUSG00000079481 | -0,668 | 0,012 | Nhsl2 |
| ENSMUSG00000020652 | -0,668 | 0,000 | Cenpo |
| ENSMUSG00000037355 | -0,669 | 0,011 | Uvssa |
| ENSMUSG00000037736 | -0,669 | 0,004 | Limch1 |
| ENSMUSG00000038332 | -0,669 | 0,000 | Sesn1 |
| ENSMUSG00000027496 | -0,670 | 0,003 | Aurka |
| ENSMUSG00000028385 | -0,670 | 0,003 | Snx30 |
| ENSMUSG00000036437 | -0,670 | 0,022 | Npy1r |
| ENSMUSG00000040860 | -0,671 | 0,000 | Crocc |
| ENSMUSG00000002007 | -0,672 | 0,001 | Srpk3 |
| ENSMUSG00000025001 | -0,672 | 0,008 | Hells |
| ENSMUSG00000051344 | -0,673 | 0,011 | Plekhm3 |
| ENSMUSG00000030677 | -0,673 | 0,017 | Kif22 |
| ENSMUSG00000041219 | -0,673 | 0,025 | Arhgap11a |
| ENSMUSG00000024791 | -0,673 | 0,002 | Cdca5 |
| ENSMUSG00000035992 | -0,673 | 0,035 | Fnip1 |
| ENSMUSG00000031543 | -0,674 | 0,004 | Ank1 |
| ENSMUSG00000031150 | -0,674 | 0,005 | Ccdc120 |
| ENSMUSG00000032796 | -0,676 | 0,008 | Lama1 |
| ENSMUSG00000033904 | -0,676 | 0,010 | Ccp110 |
| ENSMUSG00000056458 | -0,677 | 0,000 | Mok |
| ENSMUSG00000039601 | -0,677 | 0,000 | Rcan2 |
| ENSMUSG00000038022 | -0,678 | 0,000 | Mindy4 |
| ENSMUSG00000030319 | -0,678 | 0,001 | Cand2 |
| ENSMUSG00000048118 | -0,678 | 0,011 | Arid4a |
| ENSMUSG00000113425 | -0,679 | 0,046 | AC159813.1 |
| ENSMUSG00000070822 | -0,679 | 0,000 | Zscan18 |
| ENSMUSG00000040875 | -0,679 | 0,047 | Osbpl10 |
| ENSMUSG00000108011 | -0,679 | 0,002 | Gm44596 |
| ENSMUSG00000022464 | -0,680 | 0,044 | Slc38a4 |
| ENSMUSG00000049404 | -0,681 | 0,018 | Rarres1 |
| ENSMUSG00000030551 | -0,681 | 0,001 | Nr2f2 |
| ENSMUSG00000028546 | -0,682 | 0,010 | Elavl4 |
| ENSMUSG00000030539 | -0,683 | 0,001 | Sema4b |
| ENSMUSG00000040717 | -0,683 | 0,029 | Il17rd |
| ENSMUSG00000024989 | -0,684 | 0,015 | Cep55 |
| ENSMUSG00000026683 | -0,684 | 0,000 | Nuf2 |
| ENSMUSG00000036617 | -0,684 | 0,032 | Etl4 |

| | | | |
|---|---|---|---|
| ENSMUSG00000027883 | -0,684 | 0,000 | Gpsm2 |
| ENSMUSG00000048058 | -0,685 | 0,008 | Ldlrad3 |
| ENSMUSG00000021175 | -0,685 | 0,000 | Cdca7l |
| ENSMUSG00000034773 | -0,685 | 0,007 | BC030867 |
| ENSMUSG00000022122 | -0,685 | 0,016 | Ednrb |
| ENSMUSG00000030380 | -0,687 | 0,003 | Mzf1 |
| ENSMUSG00000021835 | -0,687 | 0,042 | Bmp4 |
| ENSMUSG00000020649 | -0,687 | 0,000 | Rrm2 |
| ENSMUSG00000029673 | -0,688 | 0,015 | Auts2 |
| ENSMUSG00000034471 | -0,688 | 0,001 | Caskin2 |
| ENSMUSG00000027811 | -0,689 | 0,000 | 4930579G24Rik |
| ENSMUSG00000021198 | -0,689 | 0,022 | Unc79 |
| ENSMUSG00000043065 | -0,689 | 0,000 | Spice1 |
| ENSMUSG00000042010 | -0,690 | 0,017 | Acacb |
| ENSMUSG00000049916 | -0,691 | 0,002 | 2610318N02Rik |
| ENSMUSG00000033420 | -0,691 | 0,002 | Antxr1 |
| ENSMUSG00000020099 | -0,691 | 0,008 | Unc5b |
| ENSMUSG00000027075 | -0,695 | 0,000 | Slc43a1 |
| ENSMUSG00000021176 | -0,695 | 0,000 | Efcab11 |
| ENSMUSG00000026872 | -0,695 | 0,039 | Zeb2 |
| ENSMUSG00000055148 | -0,696 | 0,012 | Klf2 |
| ENSMUSG00000030102 | -0,696 | 0,037 | Itpr1 |
| ENSMUSG00000114584 | -0,696 | 0,028 | AC131687.1 |
| ENSMUSG00000050541 | -0,697 | 0,016 | Adra1b |
| ENSMUSG00000031262 | -0,697 | 0,000 | Cenpi |
| ENSMUSG00000040852 | -0,698 | 0,007 | Plekhh2 |
| ENSMUSG00000041961 | -0,698 | 0,017 | Znrf3 |
| ENSMUSG00000049303 | -0,698 | 0,042 | Syt12 |
| ENSMUSG00000087403 | -0,699 | 0,001 | Kantr |
| ENSMUSG00000107476 | -0,699 | 0,001 | Zfp862-ps |
| ENSMUSG00000045273 | -0,699 | 0,001 | Cenph |
| ENSMUSG00000059401 | -0,701 | 0,020 | Mamld1 |
| ENSMUSG00000097415 | -0,701 | 0,000 | AU020206 |
| ENSMUSG00000026014 | -0,701 | 0,012 | Raph1 |
| ENSMUSG00000020330 | -0,701 | 0,005 | Hmmr |
| ENSMUSG00000020642 | -0,702 | 0,006 | Rnf144a |
| ENSMUSG00000048445 | -0,702 | 0,000 | Ccdc57 |
| ENSMUSG00000074212 | -0,702 | 0,001 | Dnajb14 |
| ENSMUSG00000060548 | -0,702 | 0,000 | Tnfrsf19 |
| ENSMUSG00000061186 | -0,703 | 0,045 | Sfmbt2 |
| ENSMUSG00000115186 | -0,703 | 0,039 | AC151836.1 |
| ENSMUSG00000027323 | -0,704 | 0,000 | Rad51 |
| ENSMUSG00000027524 | -0,704 | 0,003 | Edn3 |
| ENSMUSG00000021707 | -0,704 | 0,000 | Dhfr |
| ENSMUSG00000046808 | -0,704 | 0,000 | Atp10d |
| ENSMUSG00000037337 | -0,705 | 0,009 | Map4k1 |
| ENSMUSG00000015314 | -0,705 | 0,013 | Slamf6 |

| | | | |
|---|---|---|---|
| ENSMUSG00000032179 | -0,706 | 0,024 | Bmp5 |
| ENSMUSG00000021572 | -0,706 | 0,000 | Cep72 |
| ENSMUSG00000070565 | -0,706 | 0,037 | Rasal2 |
| ENSMUSG00000020897 | -0,706 | 0,000 | Aurkb |
| ENSMUSG00000024376 | -0,707 | 0,015 | Epb41l4a |
| ENSMUSG00000062901 | -0,708 | 0,047 | Klhl24 |
| ENSMUSG00000028017 | -0,709 | 0,015 | Egf |
| ENSMUSG00000002996 | -0,709 | 0,000 | Hbp1 |
| ENSMUSG00000089853 | -0,710 | 0,037 | Gm15853 |
| ENSMUSG00000074802 | -0,711 | 0,027 | Gas2l3 |
| ENSMUSG00000019699 | -0,711 | 0,013 | Akt3 |
| ENSMUSG00000059173 | -0,711 | 0,011 | Pde1a |
| ENSMUSG00000033981 | -0,712 | 0,043 | Gria2 |
| ENSMUSG00000045954 | -0,712 | 0,001 | Cavin2 |
| ENSMUSG00000015880 | -0,713 | 0,005 | Ncapg |
| ENSMUSG00000063796 | -0,713 | 0,014 | Slc22a8 |
| ENSMUSG00000037533 | -0,713 | 0,017 | Rapgef6 |
| ENSMUSG00000075592 | -0,713 | 0,001 | Nynrin |
| ENSMUSG00000037890 | -0,714 | 0,001 | Wdr19 |
| ENSMUSG00000029438 | -0,714 | 0,000 | Bcl7a |
| ENSMUSG00000073434 | -0,714 | 0,000 | Wdr90 |
| ENSMUSG00000027015 | -0,715 | 0,019 | Cybrd1 |
| ENSMUSG00000107253 | -0,715 | 0,032 | Gm43634 |
| ENSMUSG00000072837 | -0,715 | 0,000 | Mir546 |
| ENSMUSG00000041431 | -0,716 | 0,001 | Ccnb1 |
| ENSMUSG00000036928 | -0,718 | 0,012 | Stag3 |
| ENSMUSG00000033253 | -0,718 | 0,000 | Szt2 |
| ENSMUSG00000024793 | -0,719 | 0,042 | Tnfrsf25 |
| ENSMUSG00000001228 | -0,719 | 0,000 | Uhrf1 |
| ENSMUSG00000030346 | -0,720 | 0,000 | Rad51ap1 |
| ENSMUSG00000009575 | -0,721 | 0,000 | Cbx5 |
| ENSMUSG00000027661 | -0,722 | 0,023 | Slc2a10 |
| ENSMUSG00000039774 | -0,722 | 0,013 | Galnt12 |
| ENSMUSG00000025140 | -0,722 | 0,002 | Pycr1 |
| ENSMUSG00000032113 | -0,722 | 0,000 | Chek1 |
| ENSMUSG00000038070 | -0,723 | 0,000 | Cntln |
| ENSMUSG00000063804 | -0,723 | 0,000 | Lin28b |
| ENSMUSG00000050199 | -0,723 | 0,011 | Lgr4 |
| ENSMUSG00000032009 | -0,725 | 0,008 | Sesn3 |
| ENSMUSG00000017861 | -0,726 | 0,000 | Mybl2 |
| ENSMUSG00000025815 | -0,726 | 0,026 | Dhtkd1 |
| ENSMUSG00000051497 | -0,727 | 0,022 | Kcnj16 |
| ENSMUSG00000041696 | -0,728 | 0,019 | Rasl12 |
| ENSMUSG00000028587 | -0,728 | 0,004 | Orc1 |
| ENSMUSG00000033906 | -0,729 | 0,013 | Zdhhc15 |
| ENSMUSG00000005951 | -0,729 | 0,000 | Shpk |
| ENSMUSG00000001507 | -0,729 | 0,002 | Itga3 |

| | | | |
|---|---|---|---|
| ENSMUSG00000003779 | -0,731 | 0,001 | Kif20a |
| ENSMUSG00000035045 | -0,731 | 0,047 | Zc3h12b |
| ENSMUSG00000033170 | -0,732 | 0,001 | Card10 |
| ENSMUSG00000028214 | -0,732 | 0,038 | Gem |
| ENSMUSG00000032717 | -0,733 | 0,034 | Mdfi |
| ENSMUSG00000029714 | -0,733 | 0,000 | Gigyf1 |
| ENSMUSG00000036223 | -0,734 | 0,001 | Ska1 |
| ENSMUSG00000024299 | -0,734 | 0,000 | Adamts10 |
| ENSMUSG00000045328 | -0,734 | 0,005 | Cenpe |
| ENSMUSG00000039763 | -0,734 | 0,000 | Dnajc28 |
| ENSMUSG00000040260 | -0,734 | 0,006 | Daam2 |
| ENSMUSG00000025574 | -0,734 | 0,004 | Tk1 |
| ENSMUSG00000052407 | -0,735 | 0,030 | Ccdc171 |
| ENSMUSG00000027489 | -0,735 | 0,008 | Necab3 |
| ENSMUSG00000038379 | -0,735 | 0,000 | Ttk |
| ENSMUSG00000028551 | -0,737 | 0,000 | Cdkn2c |
| ENSMUSG00000063415 | -0,738 | 0,031 | Cyp26b1 |
| ENSMUSG00000030255 | -0,738 | 0,009 | Sspn |
| ENSMUSG00000072762 | -0,742 | 0,001 | 4930522L14Rik |
| ENSMUSG00000087396 | -0,742 | 0,001 | 4933407K13Rik |
| ENSMUSG00000047757 | -0,743 | 0,000 | Fancb |
| ENSMUSG00000024301 | -0,743 | 0,002 | Kifc5b |
| ENSMUSG00000076275 | -0,745 | 0,003 | Mir675 |
| ENSMUSG00000051022 | -0,745 | 0,001 | Hs3st1 |
| ENSMUSG00000026809 | -0,745 | 0,035 | Spaca9 |
| ENSMUSG00000034883 | -0,746 | 0,044 | Lrr1 |
| ENSMUSG00000033715 | -0,746 | 0,030 | Akr1c14 |
| ENSMUSG00000042628 | -0,747 | 0,000 | Zfyve1 |
| ENSMUSG00000024174 | -0,747 | 0,000 | Pot1b |
| ENSMUSG00000055737 | -0,747 | 0,000 | Ghr |
| ENSMUSG00000037020 | -0,747 | 0,000 | Wdr62 |
| ENSMUSG00000016493 | -0,748 | 0,046 | Cd46 |
| ENSMUSG00000040055 | -0,748 | 0,012 | Gjb6 |
| ENSMUSG00000036862 | -0,749 | 0,007 | Dchs1 |
| ENSMUSG00000097337 | -0,749 | 0,034 | Gm26606 |
| ENSMUSG00000045231 | -0,750 | 0,010 | BC106179 |
| ENSMUSG00000005043 | -0,750 | 0,000 | Sgsh |
| ENSMUSG00000094595 | -0,750 | 0,000 | Fsbp |
| ENSMUSG00000044702 | -0,750 | 0,000 | Palb2 |
| ENSMUSG00000047888 | -0,751 | 0,010 | Tnrc6b |
| ENSMUSG00000068744 | -0,752 | 0,004 | Psrc1 |
| ENSMUSG00000052105 | -0,753 | 0,045 | Mtcl1 |
| ENSMUSG00000006715 | -0,753 | 0,001 | Gmnn |
| ENSMUSG00000039187 | -0,753 | 0,000 | Fanci |
| ENSMUSG00000037474 | -0,754 | 0,000 | Dtl |
| ENSMUSG00000026622 | -0,754 | 0,005 | Nek2 |
| ENSMUSG00000097243 | -0,754 | 0,045 | Gm26908 |

| | | | |
|---|---|---|---|
| ENSMUSG00000021767 | -0,755 | 0,011 | Kat6b |
| ENSMUSG00000019845 | -0,755 | 0,000 | Tube1 |
| ENSMUSG00000066440 | -0,756 | 0,003 | Zfyve26 |
| ENSMUSG00000026812 | -0,756 | 0,000 | Tsc1 |
| ENSMUSG00000039963 | -0,756 | 0,005 | Ccdc40 |
| ENSMUSG00000028024 | -0,756 | 0,015 | Enpep |
| ENSMUSG00000023066 | -0,756 | 0,012 | Rttn |
| ENSMUSG00000034684 | -0,757 | 0,006 | Sema3f |
| ENSMUSG00000072980 | -0,757 | 0,000 | Oip5 |
| ENSMUSG00000000056 | -0,757 | 0,000 | Narf |
| ENSMUSG00000022893 | -0,757 | 0,004 | Adamts1 |
| ENSMUSG00000021662 | -0,757 | 0,003 | Arhgef28 |
| ENSMUSG00000060176 | -0,757 | 0,037 | Kif27 |
| ENSMUSG00000032558 | -0,757 | 0,000 | Nphp3 |
| ENSMUSG00000025555 | -0,758 | 0,000 | Farp1 |
| ENSMUSG00000024901 | -0,758 | 0,000 | Peli3 |
| ENSMUSG00000028803 | -0,759 | 0,000 | Nipal3 |
| ENSMUSG00000110652 | -0,759 | 0,039 | Gm33023 |
| ENSMUSG00000000142 | -0,760 | 0,001 | Axin2 |
| ENSMUSG00000023034 | -0,760 | 0,042 | Nr4a1 |
| ENSMUSG00000026674 | -0,761 | 0,003 | Ddr2 |
| ENSMUSG00000019763 | -0,761 | 0,000 | Rmnd1 |
| ENSMUSG00000041147 | -0,761 | 0,000 | Brca2 |
| ENSMUSG00000027843 | -0,762 | 0,004 | Ptpn22 |
| ENSMUSG00000024713 | -0,763 | 0,002 | Pcsk5 |
| ENSMUSG00000097536 | -0,763 | 0,043 | 2610037D02Rik |
| ENSMUSG00000026255 | -0,763 | 0,023 | Efhd1 |
| ENSMUSG00000048001 | -0,763 | 0,047 | Hes5 |
| ENSMUSG00000030494 | -0,763 | 0,010 | Rhpn2 |
| ENSMUSG00000019773 | -0,764 | 0,000 | Fbxo5 |
| ENSMUSG00000090626 | -0,764 | 0,000 | Tex9 |
| ENSMUSG00000103400 | -0,764 | 0,000 | Gm15853 |
| ENSMUSG00000019214 | -0,764 | 0,000 | Chtf18 |
| ENSMUSG00000044783 | -0,765 | 0,000 | Hjurp |
| ENSMUSG00000036867 | -0,765 | 0,000 | Smad6 |
| ENSMUSG00000040276 | -0,765 | 0,000 | Pacsin1 |
| ENSMUSG00000025395 | -0,765 | 0,000 | Prim1 |
| ENSMUSG00000032238 | -0,766 | 0,004 | Rora |
| ENSMUSG00000024598 | -0,766 | 0,030 | Fbn2 |
| ENSMUSG00000022475 | -0,766 | 0,000 | Hdac7 |
| ENSMUSG00000037336 | -0,766 | 0,037 | Mfsd2b |
| ENSMUSG00000099032 | -0,766 | 0,050 | Tcf24 |
| ENSMUSG00000029414 | -0,767 | 0,001 | Kntc1 |
| ENSMUSG00000047534 | -0,767 | 0,001 | Mis18bp1 |
| ENSMUSG00000005774 | -0,767 | 0,000 | Rfx5 |
| ENSMUSG00000079685 | -0,768 | 0,004 | Ulbp1 |
| ENSMUSG00000039748 | -0,768 | 0,000 | Exo1 |

| | | | |
|---|---|---|---|
| ENSMUSG00000020098 | -0,768 | 0,005 | Pcbd1 |
| ENSMUSG00000069892 | -0,769 | 0,032 | 9930111J21Rik2 |
| ENSMUSG00000050751 | -0,769 | 0,009 | Pgbd5 |
| ENSMUSG00000054134 | -0,769 | 0,029 | Umodl1 |
| ENSMUSG00000067860 | -0,769 | 0,004 | Zic3 |
| ENSMUSG00000071324 | -0,770 | 0,000 | Armc2 |
| ENSMUSG00000033705 | -0,771 | 0,010 | Stard9 |
| ENSMUSG00000034903 | -0,771 | 0,017 | Cobll1 |
| ENSMUSG00000000037 | -0,772 | 0,007 | Scml2 |
| ENSMUSG00000023940 | -0,772 | 0,000 | Sgo1 |
| ENSMUSG00000020883 | -0,772 | 0,000 | Fbxl20 |
| ENSMUSG00000000028 | -0,772 | 0,000 | Cdc45 |
| ENSMUSG00000002076 | -0,773 | 0,024 | Hsf2bp |
| ENSMUSG00000035365 | -0,773 | 0,007 | Parpbp |
| ENSMUSG00000025776 | -0,773 | 0,028 | Crispld1 |
| ENSMUSG00000014164 | -0,773 | 0,000 | Klhl3 |
| ENSMUSG00000110393 | -0,774 | 0,010 | Gm36445 |
| ENSMUSG00000041957 | -0,774 | 0,030 | Pkp2 |
| ENSMUSG00000046280 | -0,775 | 0,011 | She |
| ENSMUSG00000056214 | -0,775 | 0,000 | Pard6g |
| ENSMUSG00000031355 | -0,775 | 0,030 | Arhgap6 |
| ENSMUSG00000021690 | -0,775 | 0,006 | Jmy |
| ENSMUSG00000020105 | -0,775 | 0,000 | Lrig3 |
| ENSMUSG00000049134 | -0,776 | 0,006 | Nrap |
| ENSMUSG00000025154 | -0,777 | 0,000 | Arhgap19 |
| ENSMUSG00000041064 | -0,777 | 0,000 | Pif1 |
| ENSMUSG00000004099 | -0,778 | 0,000 | Dnmt1 |
| ENSMUSG00000042097 | -0,779 | 0,005 | Zfp239 |
| ENSMUSG00000032633 | -0,779 | 0,000 | Flcn |
| ENSMUSG00000021180 | -0,779 | 0,000 | Rps6ka5 |
| ENSMUSG00000032925 | -0,781 | 0,022 | Itgbl1 |
| ENSMUSG00000020042 | -0,781 | 0,028 | Btbd11 |
| ENSMUSG00000089809 | -0,781 | 0,005 | Rasgef1b |
| ENSMUSG00000040631 | -0,782 | 0,001 | Dok4 |
| ENSMUSG00000002032 | -0,782 | 0,002 | Tmem25 |
| ENSMUSG00000072082 | -0,783 | 0,000 | Ccnf |
| ENSMUSG00000048939 | -0,783 | 0,044 | Atp13a5 |
| ENSMUSG00000032400 | -0,783 | 0,000 | Zwilch |
| ENSMUSG00000025993 | -0,784 | 0,018 | Slc40a1 |
| ENSMUSG00000001864 | -0,784 | 0,000 | Aif1l |
| ENSMUSG00000030606 | -0,785 | 0,000 | Hapln3 |
| ENSMUSG00000090115 | -0,785 | 0,000 | Usp49 |
| ENSMUSG00000012443 | -0,785 | 0,002 | Kif11 |
| ENSMUSG00000034601 | -0,786 | 0,000 | 2700049A03Rik |
| ENSMUSG00000030237 | -0,787 | 0,002 | Slco1a4 |
| ENSMUSG00000074024 | -0,788 | 0,008 | 4632427E13Rik |
| ENSMUSG00000021573 | -0,789 | 0,001 | Tppp |

| | | | |
|---|---|---|---|
| ENSMUSG00000020774 | -0,790 | 0,000 | Aspa |
| ENSMUSG00000040147 | -0,792 | 0,004 | Maob |
| ENSMUSG00000049265 | -0,792 | 0,048 | Kcnk3 |
| ENSMUSG00000097177 | -0,792 | 0,039 | 9330159M07Rik |
| ENSMUSG00000074194 | -0,793 | 0,002 | Zfp791 |
| ENSMUSG00000030499 | -0,797 | 0,000 | Kctd15 |
| ENSMUSG00000030134 | -0,797 | 0,011 | Rasgef1a |
| ENSMUSG00000049488 | -0,797 | 0,000 | Tmem67 |
| ENSMUSG00000044362 | -0,797 | 0,018 | Ccdc89 |
| ENSMUSG00000029641 | -0,798 | 0,022 | Rasl11a |
| ENSMUSG00000001517 | -0,799 | 0,000 | Foxm1 |
| ENSMUSG00000030865 | -0,800 | 0,013 | Chp2 |
| ENSMUSG00000114980 | -0,802 | 0,048 | AC102815.1 |
| ENSMUSG00000074415 | -0,803 | 0,035 | 2610203C20Rik |
| ENSMUSG00000041930 | -0,803 | 0,000 | Fam222a |
| ENSMUSG00000020493 | -0,806 | 0,008 | Prr11 |
| ENSMUSG00000039765 | -0,807 | 0,000 | Cc2d2a |
| ENSMUSG00000021294 | -0,808 | 0,002 | Kif26a |
| ENSMUSG00000109179 | -0,808 | 0,011 | Gm35339 |
| ENSMUSG00000034329 | -0,808 | 0,003 | Brip1 |
| ENSMUSG00000022206 | -0,808 | 0,021 | Npr3 |
| ENSMUSG00000108368 | -0,809 | 0,026 | Gm45053 |
| ENSMUSG00000031196 | -0,809 | 0,007 | F8 |
| ENSMUSG00000047867 | -0,810 | 0,001 | Gimap6 |
| ENSMUSG00000034911 | -0,811 | 0,031 | Ushbp1 |
| ENSMUSG00000052302 | -0,811 | 0,001 | Tbc1d30 |
| ENSMUSG00000040287 | -0,811 | 0,027 | Stac3 |
| ENSMUSG00000038119 | -0,812 | 0,010 | Cdon |
| ENSMUSG00000022833 | -0,812 | 0,000 | Ccdc14 |
| ENSMUSG00000051378 | -0,812 | 0,000 | Kif18b |
| ENSMUSG00000051220 | -0,813 | 0,000 | Ercc6l |
| ENSMUSG00000036777 | -0,814 | 0,003 | Anln |
| ENSMUSG00000026358 | -0,814 | 0,050 | Rgs1 |
| ENSMUSG00000026768 | -0,815 | 0,014 | Itga8 |
| ENSMUSG00000049580 | -0,815 | 0,000 | Tsku |
| ENSMUSG00000073437 | -0,815 | 0,000 | D330041H03Rik |
| ENSMUSG00000057137 | -0,816 | 0,000 | Tmem140 |
| ENSMUSG00000042029 | -0,816 | 0,016 | Ncapg2 |
| ENSMUSG00000045092 | -0,816 | 0,000 | S1pr1 |
| ENSMUSG00000027635 | -0,817 | 0,000 | Dsn1 |
| ENSMUSG00000038331 | -0,817 | 0,030 | Satb2 |
| ENSMUSG00000091586 | -0,817 | 0,002 | Cyp4f17 |
| ENSMUSG00000037544 | -0,819 | 0,000 | Dlgap5 |
| ENSMUSG00000000416 | -0,820 | 0,020 | Cttnbp2 |
| ENSMUSG00000063450 | -0,820 | 0,023 | Syne2 |
| ENSMUSG00000027379 | -0,821 | 0,000 | Bub1 |
| ENSMUSG00000087231 | -0,821 | 0,003 | E230016M11Rik |

| | | | |
|---|---|---|---|
| ENSMUSG00000041688 | -0,821 | 0,006 | Amot |
| ENSMUSG00000097519 | -0,821 | 0,042 | 4930558J18Rik |
| ENSMUSG00000048728 | -0,822 | 0,014 | Zfp454 |
| ENSMUSG00000091661 | -0,823 | 0,002 | Gm17021 |
| ENSMUSG00000021965 | -0,824 | 0,000 | Ska3 |
| ENSMUSG00000028278 | -0,825 | 0,000 | Rragd |
| ENSMUSG00000028555 | -0,825 | 0,039 | Ttc39a |
| ENSMUSG00000030254 | -0,825 | 0,000 | Rad18 |
| ENSMUSG00000061414 | -0,826 | 0,001 | Cracr2a |
| ENSMUSG00000031847 | -0,829 | 0,001 | 1700030J22Rik |
| ENSMUSG00000026648 | -0,830 | 0,000 | Dclre1c |
| ENSMUSG00000046470 | -0,830 | 0,030 | Sox18 |
| ENSMUSG00000006386 | -0,830 | 0,021 | Tek |
| ENSMUSG00000048922 | -0,830 | 0,001 | Cdca2 |
| ENSMUSG00000038732 | -0,830 | 0,000 | Mboat1 |
| ENSMUSG00000048924 | -0,831 | 0,000 | Ccdc125 |
| ENSMUSG00000029913 | -0,831 | 0,000 | Prdm5 |
| ENSMUSG00000026605 | -0,834 | 0,003 | Cenpf |
| ENSMUSG00000030020 | -0,835 | 0,023 | Prickle2 |
| ENSMUSG00000078937 | -0,836 | 0,000 | Cpt1b |
| ENSMUSG00000048402 | -0,837 | 0,021 | Gli2 |
| ENSMUSG00000030559 | -0,838 | 0,021 | Rab38 |
| ENSMUSG00000042961 | -0,838 | 0,000 | Egflam |
| ENSMUSG00000098090 | -0,840 | 0,009 | 2700099C18Rik |
| ENSMUSG00000075225 | -0,840 | 0,013 | Ccdc162 |
| ENSMUSG00000024727 | -0,840 | 0,007 | Trpm6 |
| ENSMUSG00000103260 | -0,841 | 0,016 | Gm8146 |
| ENSMUSG00000027699 | -0,841 | 0,009 | Ect2 |
| ENSMUSG00000039620 | -0,847 | 0,023 | 6430573F11Rik |
| ENSMUSG00000084896 | -0,850 | 0,013 | Gm11632 |
| ENSMUSG00000032783 | -0,851 | 0,000 | Troap |
| ENSMUSG00000002997 | -0,851 | 0,000 | Prkar2b |
| ENSMUSG00000030650 | -0,853 | 0,039 | Tmc5 |
| ENSMUSG00000039741 | -0,854 | 0,003 | Bahcc1 |
| ENSMUSG00000028702 | -0,854 | 0,000 | Rad54l |
| ENSMUSG00000027115 | -0,855 | 0,000 | Kif18a |
| ENSMUSG00000024795 | -0,855 | 0,001 | Kif20b |
| ENSMUSG00000023033 | -0,856 | 0,042 | Scn8a |
| ENSMUSG00000022332 | -0,856 | 0,000 | Khdrbs3 |
| ENSMUSG00000034023 | -0,857 | 0,000 | Fancd2 |
| ENSMUSG00000071855 | -0,858 | 0,000 | Ccdc112 |
| ENSMUSG00000060402 | -0,859 | 0,017 | Chst8 |
| ENSMUSG00000021575 | -0,862 | 0,000 | Ahrr |
| ENSMUSG00000030528 | -0,862 | 0,000 | Blm |
| ENSMUSG00000010175 | -0,863 | 0,002 | Prox1 |
| ENSMUSG00000042429 | -0,863 | 0,016 | Adora1 |
| ENSMUSG00000072915 | -0,864 | 0,000 | Gm12258 |

| | | | |
|---|---|---|---|
| ENSMUSG00000055240 | -0,864 | 0,000 | Zfp101 |
| ENSMUSG00000060275 | -0,864 | 0,023 | Nrg2 |
| ENSMUSG00000027242 | -0,864 | 0,000 | Wdr76 |
| ENSMUSG00000032855 | -0,865 | 0,004 | Pkd1 |
| ENSMUSG00000021611 | -0,866 | 0,006 | Tert |
| ENSMUSG00000016028 | -0,866 | 0,024 | Celsr1 |
| ENSMUSG00000022604 | -0,867 | 0,000 | Cep97 |
| ENSMUSG00000042988 | -0,870 | 0,001 | Notum |
| ENSMUSG00000051043 | -0,871 | 0,000 | Gprc5c |
| ENSMUSG00000090958 | -0,872 | 0,002 | Lrrc32 |
| ENSMUSG00000039814 | -0,874 | 0,001 | Xkr5 |
| ENSMUSG00000026039 | -0,874 | 0,000 | Sgo2a |
| ENSMUSG00000039055 | -0,876 | 0,000 | Eme1 |
| ENSMUSG00000078502 | -0,876 | 0,016 | Gm13212 |
| ENSMUSG00000051367 | -0,877 | 0,044 | Six1 |
| ENSMUSG00000048249 | -0,877 | 0,000 | Crebrf |
| ENSMUSG00000035024 | -0,878 | 0,000 | Ncapd3 |
| ENSMUSG00000086782 | -0,878 | 0,043 | E130102H24Rik |
| ENSMUSG00000043639 | -0,878 | 0,029 | Rbm20 |
| ENSMUSG00000047473 | -0,879 | 0,000 | Zfp30 |
| ENSMUSG00000027875 | -0,879 | 0,000 | Hmgcs2 |
| ENSMUSG00000034610 | -0,879 | 0,000 | Zcchc11 |
| ENSMUSG00000042851 | -0,879 | 0,008 | Zc3h6 |
| ENSMUSG00000048387 | -0,880 | 0,029 | Osr1 |
| ENSMUSG00000089837 | -0,880 | 0,011 | Npcd |
| ENSMUSG00000086962 | -0,880 | 0,007 | Gm12248 |
| ENSMUSG00000029406 | -0,880 | 0,000 | Pitpnm2 |
| ENSMUSG00000032815 | -0,881 | 0,000 | Fanca |
| ENSMUSG00000046352 | -0,883 | 0,000 | Gjb2 |
| ENSMUSG00000015659 | -0,884 | 0,003 | Serac1 |
| ENSMUSG00000056531 | -0,885 | 0,001 | Ccdc18 |
| ENSMUSG00000033149 | -0,885 | 0,001 | Phldb2 |
| ENSMUSG00000028978 | -0,885 | 0,006 | Nos3 |
| ENSMUSG00000028804 | -0,886 | 0,039 | Csmd2 |
| ENSMUSG00000113184 | -0,886 | 0,006 | AC125351.1 |
| ENSMUSG00000020758 | -0,887 | 0,002 | Itgb4 |
| ENSMUSG00000045930 | -0,888 | 0,000 | Clec14a |
| ENSMUSG00000040133 | -0,890 | 0,031 | Gpr176 |
| ENSMUSG00000047747 | -0,890 | 0,000 | Rnf150 |
| ENSMUSG00000074220 | -0,890 | 0,037 | Zfp382 |
| ENSMUSG00000030091 | -0,890 | 0,000 | Nup210 |
| ENSMUSG00000037568 | -0,892 | 0,000 | Vash2 |
| ENSMUSG00000041695 | -0,893 | 0,007 | Kcnj2 |
| ENSMUSG00000021186 | -0,893 | 0,000 | Fbln5 |
| ENSMUSG00000084989 | -0,894 | 0,012 | Crocc2 |
| ENSMUSG00000080463 | -0,895 | 0,031 | Gm22489 |
| ENSMUSG00000027074 | -0,895 | 0,000 | Slc43a3 |

| | | | |
|---|---|---|---|
| ENSMUSG00000021287 | -0,895 | 0,000 | Xrcc3 |
| ENSMUSG00000098183 | -0,895 | 0,038 | Gm27010 |
| ENSMUSG00000038068 | -0,895 | 0,000 | Rnf144b |
| ENSMUSG00000074793 | -0,896 | 0,000 | Hspa12b |
| ENSMUSG00000021714 | -0,897 | 0,001 | Cenpk |
| ENSMUSG00000049119 | -0,897 | 0,000 | Fam110b |
| ENSMUSG00000041836 | -0,897 | 0,000 | Ptpre |
| ENSMUSG00000069910 | -0,898 | 0,000 | Spdl1 |
| ENSMUSG00000026196 | -0,900 | 0,001 | Bard1 |
| ENSMUSG00000024056 | -0,901 | 0,000 | Ndc80 |
| ENSMUSG00000044667 | -0,902 | 0,037 | Plppr4 |
| ENSMUSG00000094942 | -0,903 | 0,018 | Gm3604 |
| ENSMUSG00000032254 | -0,904 | 0,000 | Kif23 |
| ENSMUSG00000051606 | -0,904 | 0,004 | 2010001K21Rik |
| ENSMUSG00000038151 | -0,908 | 0,002 | Prdm1 |
| ENSMUSG00000027977 | -0,908 | 0,016 | Ndst3 |
| ENSMUSG00000020593 | -0,909 | 0,000 | Lpin1 |
| ENSMUSG00000019027 | -0,909 | 0,020 | Dnah1 |
| ENSMUSG00000057604 | -0,911 | 0,000 | Lmcd1 |
| ENSMUSG00000036186 | -0,912 | 0,001 | Fam69b |
| ENSMUSG00000085133 | -0,913 | 0,000 | B930095G15Rik |
| ENSMUSG00000045287 | -0,913 | 0,000 | Rtn4rl1 |
| ENSMUSG00000018604 | -0,914 | 0,005 | Tbx3 |
| ENSMUSG00000039981 | -0,914 | 0,042 | Zc3h12d |
| ENSMUSG00000020256 | -0,914 | 0,006 | Aldh1l2 |
| ENSMUSG00000041633 | -0,914 | 0,000 | Kctd12b |
| ENSMUSG00000029070 | -0,915 | 0,001 | Mxra8 |
| ENSMUSG00000047146 | -0,916 | 0,014 | Tet1 |
| ENSMUSG00000114995 | -0,916 | 0,018 | AC137842.1 |
| ENSMUSG00000064493 | -0,917 | 0,024 | Snora28 |
| ENSMUSG00000053411 | -0,918 | 0,000 | Cbx7 |
| ENSMUSG00000024647 | -0,918 | 0,029 | Cbln2 |
| ENSMUSG00000073414 | -0,918 | 0,035 | Mpig6b |
| ENSMUSG00000036768 | -0,920 | 0,000 | Kif15 |
| ENSMUSG00000067430 | -0,923 | 0,001 | Zfp763 |
| ENSMUSG00000033453 | -0,927 | 0,003 | Adamts15 |
| ENSMUSG00000058740 | -0,930 | 0,021 | Kcnt1 |
| ENSMUSG00000020275 | -0,930 | 0,000 | Rel |
| ENSMUSG00000036875 | -0,930 | 0,000 | Dna2 |
| ENSMUSG00000040310 | -0,933 | 0,008 | Alx4 |
| ENSMUSG00000019982 | -0,934 | 0,000 | Myb |
| ENSMUSG00000020381 | -0,934 | 0,000 | Mrnip |
| ENSMUSG00000039385 | -0,936 | 0,000 | Cdh6 |
| ENSMUSG00000078866 | -0,938 | 0,018 | Zfp970 |
| ENSMUSG00000002409 | -0,938 | 0,000 | Dyrk1b |
| ENSMUSG00000029516 | -0,939 | 0,001 | Cit |
| ENSMUSG00000090665 | -0,940 | 0,023 | Gad1-ps |

| | | | |
|---|---|---|---|
| ENSMUSG00000038305 | -0,940 | 0,000 | Spats2l |
| ENSMUSG00000035246 | -0,941 | 0,000 | Pcyt1b |
| ENSMUSG00000044303 | -0,941 | 0,017 | Cdkn2a |
| ENSMUSG00000040543 | -0,942 | 0,006 | Pitpnm3 |
| ENSMUSG00000071723 | -0,942 | 0,000 | Gspt2 |
| ENSMUSG00000026628 | -0,944 | 0,006 | Atf3 |
| ENSMUSG00000042564 | -0,946 | 0,002 | Fam227a |
| ENSMUSG00000049001 | -0,946 | 0,000 | Ndnf |
| ENSMUSG00000034586 | -0,949 | 0,000 | Hid1 |
| ENSMUSG00000041757 | -0,950 | 0,000 | Plekha6 |
| ENSMUSG00000034206 | -0,952 | 0,000 | Polq |
| ENSMUSG00000071176 | -0,953 | 0,000 | Arhgef10 |
| ENSMUSG00000098986 | -0,954 | 0,025 | Gm28006 |
| ENSMUSG00000097750 | -0,954 | 0,002 | Gm4673 |
| ENSMUSG00000027792 | -0,955 | 0,001 | Bche |
| ENSMUSG00000088219 | -0,957 | 0,010 | Gm22077 |
| ENSMUSG00000011263 | -0,957 | 0,007 | Exoc3l2 |
| ENSMUSG00000021986 | -0,960 | 0,009 | Amer2 |
| ENSMUSG00000052921 | -0,961 | 0,000 | Arhgef15 |
| ENSMUSG00000055884 | -0,961 | 0,000 | Fancm |
| ENSMUSG00000030616 | -0,963 | 0,008 | Sytl2 |
| ENSMUSG00000039396 | -0,963 | 0,000 | Neil3 |
| ENSMUSG00000002055 | -0,965 | 0,000 | Spag5 |
| ENSMUSG00000030757 | -0,965 | 0,000 | Zkscan2 |
| ENSMUSG00000048096 | -0,965 | 0,024 | Lmod1 |
| ENSMUSG00000026589 | -0,966 | 0,008 | Sec16b |
| ENSMUSG00000052852 | -0,966 | 0,000 | Reep1 |
| ENSMUSG00000050390 | -0,968 | 0,000 | C77080 |
| ENSMUSG00000022358 | -0,971 | 0,000 | Fbxo32 |
| ENSMUSG00000025504 | -0,971 | 0,018 | Eps8l2 |
| ENSMUSG00000086266 | -0,973 | 0,026 | Igf2os |
| ENSMUSG00000031004 | -0,974 | 0,000 | Mki67 |
| ENSMUSG00000000392 | -0,975 | 0,001 | Fap |
| ENSMUSG00000066191 | -0,976 | 0,000 | Anks6 |
| ENSMUSG00000033952 | -0,977 | 0,011 | Aspm |
| ENSMUSG00000039994 | -0,978 | 0,000 | Timeless |
| ENSMUSG00000027654 | -0,980 | 0,000 | Fam83d |
| ENSMUSG00000020185 | -0,981 | 0,001 | E2f7 |
| ENSMUSG00000035413 | -0,982 | 0,000 | Tmem98 |
| ENSMUSG00000026779 | -0,982 | 0,000 | Mastl |
| ENSMUSG00000063810 | -0,983 | 0,008 | Alms1 |
| ENSMUSG00000017146 | -0,983 | 0,000 | Brca1 |
| ENSMUSG00000029512 | -0,987 | 0,000 | Ulk1 |
| ENSMUSG00000052353 | -0,988 | 0,009 | Cemip |
| ENSMUSG00000028525 | -0,989 | 0,000 | Pde4b |
| ENSMUSG00000071341 | -0,989 | 0,050 | Egr4 |
| ENSMUSG00000074867 | -0,992 | 0,009 | Zfp808 |

| | | | |
|---|---|---|---|
| ENSMUSG00000020151 | -0,993 | 0,034 | Ptprr |
| ENSMUSG00000055333 | -0,994 | 0,036 | Fat2 |
| ENSMUSG00000027386 | -0,996 | 0,002 | Fbln7 |
| ENSMUSG00000028718 | -0,997 | 0,000 | Stil |
| ENSMUSG00000030283 | -0,999 | 0,012 | St8sia1 |
| ENSMUSG00000049985 | -0,999 | 0,018 | Ankrd55 |
| ENSMUSG00000002578 | -0,999 | 0,013 | Ikzf4 |
| ENSMUSG00000020905 | -1,000 | 0,013 | Usp43 |
| ENSMUSG00000029283 | -1,003 | 0,000 | Cdc7 |
| ENSMUSG00000108095 | -1,003 | 0,020 | Gm44067 |
| ENSMUSG00000039908 | -1,006 | 0,000 | Slc26a11 |
| ENSMUSG00000026274 | -1,011 | 0,000 | Pask |
| ENSMUSG00000036295 | -1,011 | 0,000 | Lrrn3 |
| ENSMUSG00000016624 | -1,012 | 0,000 | Phf21b |
| ENSMUSG00000025270 | -1,014 | 0,048 | Alas2 |
| ENSMUSG00000064043 | -1,014 | 0,000 | Trerf1 |
| ENSMUSG00000022360 | -1,014 | 0,000 | Atad2 |
| ENSMUSG00000000782 | -1,015 | 0,000 | Tcf7 |
| ENSMUSG00000105368 | -1,015 | 0,006 | Gm43759 |
| ENSMUSG00000022766 | -1,015 | 0,010 | Serpind1 |
| ENSMUSG00000058385 | -1,018 | 0,003 | Hist1h2bg |
| ENSMUSG00000019789 | -1,020 | 0,000 | Hey2 |
| ENSMUSG00000024420 | -1,021 | 0,001 | Zfp521 |
| ENSMUSG00000030123 | -1,023 | 0,001 | Plxnd1 |
| ENSMUSG00000033083 | -1,024 | 0,000 | Tbc1d4 |
| ENSMUSG00000026669 | -1,026 | 0,000 | Mcm10 |
| ENSMUSG00000053040 | -1,028 | 0,000 | Aph1c |
| ENSMUSG00000047875 | -1,028 | 0,001 | Gpr157 |
| ENSMUSG00000022723 | -1,029 | 0,000 | Crybg3 |
| ENSMUSG00000101122 | -1,032 | 0,035 | Gm17971 |
| ENSMUSG00000032419 | -1,032 | 0,000 | Tbx18 |
| ENSMUSG00000043456 | -1,034 | 0,021 | Zfp536 |
| ENSMUSG00000020182 | -1,034 | 0,018 | Ddc |
| ENSMUSG00000054580 | -1,035 | 0,003 | Pla2r1 |
| ENSMUSG00000031486 | -1,036 | 0,000 | Adgra2 |
| ENSMUSG00000028438 | -1,036 | 0,001 | Kif24 |
| ENSMUSG00000085912 | -1,038 | 0,002 | Trp53cor1 |
| ENSMUSG00000043020 | -1,038 | 0,008 | Wdr63 |
| ENSMUSG00000047793 | -1,040 | 0,000 | Sned1 |
| ENSMUSG00000027326 | -1,041 | 0,006 | Knl1 |
| ENSMUSG00000017400 | -1,043 | 0,021 | Stac2 |
| ENSMUSG00000108137 | -1,043 | 0,036 | Gm44053 |
| ENSMUSG00000035509 | -1,046 | 0,002 | Fbxl21 |
| ENSMUSG00000031629 | -1,051 | 0,000 | Cenpu |
| ENSMUSG00000003949 | -1,053 | 0,000 | Hlf |
| ENSMUSG00000030699 | -1,060 | 0,009 | Tbx6 |
| ENSMUSG00000006958 | -1,060 | 0,001 | Chrd |

| | | | |
|---|---|---|---|
| ENSMUSG00000078773 | -1,061 | 0,000 | Rad54b |
| ENSMUSG00000009418 | -1,061 | 0,000 | Nav1 |
| ENSMUSG00000073910 | -1,061 | 0,001 | Mob3b |
| ENSMUSG00000017550 | -1,062 | 0,000 | Atad5 |
| ENSMUSG00000038319 | -1,064 | 0,000 | Kcnh2 |
| ENSMUSG00000040111 | -1,065 | 0,000 | Gramd1b |
| ENSMUSG00000095123 | -1,070 | 0,012 | Gm21781 |
| ENSMUSG00000102953 | -1,072 | 0,049 | Gm37019 |
| ENSMUSG00000019989 | -1,074 | 0,022 | Enpp3 |
| ENSMUSG00000026888 | -1,074 | 0,000 | Grb14 |
| ENSMUSG00000102302 | -1,075 | 0,016 | Gm38190 |
| ENSMUSG00000080741 | -1,076 | 0,044 | Gm11398 |
| ENSMUSG00000103046 | -1,079 | 0,023 | Gm37309 |
| ENSMUSG00000044037 | -1,080 | 0,000 | Als2cl |
| ENSMUSG00000112008 | -1,080 | 0,020 | AC124556.1 |
| ENSMUSG00000024274 | -1,080 | 0,043 | Zscan30 |
| ENSMUSG00000064065 | -1,080 | 0,018 | Ipcef1 |
| ENSMUSG00000018126 | -1,082 | 0,028 | Baiap2l2 |
| ENSMUSG00000025927 | -1,083 | 0,017 | Tfap2b |
| ENSMUSG00000022034 | -1,083 | 0,000 | Esco2 |
| ENSMUSG00000042500 | -1,089 | 0,000 | Ago4 |
| ENSMUSG00000000126 | -1,093 | 0,004 | Wnt9a |
| ENSMUSG00000059479 | -1,094 | 0,000 | B3gnt8 |
| ENSMUSG00000031258 | -1,095 | 0,007 | Xkrx |
| ENSMUSG00000087142 | -1,097 | 0,012 | Gm12454 |
| ENSMUSG00000025916 | -1,098 | 0,035 | Ppp1r42 |
| ENSMUSG00000108024 | -1,098 | 0,028 | Gm43912 |
| ENSMUSG00000033207 | -1,099 | 0,000 | Mamdc2 |
| ENSMUSG00000089542 | -1,100 | 0,005 | Gm25835 |
| ENSMUSG00000069743 | -1,101 | 0,000 | Zfp820 |
| ENSMUSG00000039783 | -1,101 | 0,020 | Kmo |
| ENSMUSG00000030353 | -1,103 | 0,021 | Tead4 |
| ENSMUSG00000046818 | -1,103 | 0,001 | Ddit4l |
| ENSMUSG00000046591 | -1,105 | 0,000 | Ticrr |
| ENSMUSG00000033730 | -1,107 | 0,021 | Egr3 |
| ENSMUSG00000061175 | -1,108 | 0,001 | Fnip2 |
| ENSMUSG00000052415 | -1,109 | 0,011 | Tchh |
| ENSMUSG00000021803 | -1,110 | 0,017 | Cdhr1 |
| ENSMUSG00000033060 | -1,111 | 0,001 | Lmo7 |
| ENSMUSG00000058396 | -1,114 | 0,003 | Gpr182 |
| ENSMUSG00000086741 | -1,119 | 0,009 | Gm15816 |
| ENSMUSG00000016756 | -1,120 | 0,000 | Cmah |
| ENSMUSG00000051278 | -1,126 | 0,000 | Zgrf1 |
| ENSMUSG00000037892 | -1,126 | 0,000 | Pcdh18 |
| ENSMUSG00000037661 | -1,126 | 0,012 | Gpr160 |
| ENSMUSG00000006362 | -1,128 | 0,000 | Cbfa2t3 |
| ENSMUSG00000025577 | -1,128 | 0,000 | Cbx2 |

| | | | |
|---|---|---|---|
| ENSMUSG00000031558 | -1,129 | 0,002 | Slit2 |
| ENSMUSG00000025932 | -1,131 | 0,000 | Eya1 |
| ENSMUSG00000111535 | -1,133 | 0,019 | Gm35154 |
| ENSMUSG00000032394 | -1,136 | 0,001 | Igdcc3 |
| ENSMUSG00000103928 | -1,137 | 0,010 | Gm37893 |
| ENSMUSG00000093452 | -1,141 | 0,018 | Zfhx2os |
| ENSMUSG00000100826 | -1,141 | 0,045 | Snhg14 |
| ENSMUSG00000104965 | -1,142 | 0,038 | Gm43437 |
| ENSMUSG00000028457 | -1,142 | 0,011 | Atp8b5 |
| ENSMUSG00000078700 | -1,142 | 0,005 | D030028A08Rik |
| ENSMUSG00000025912 | -1,144 | 0,003 | Mybl1 |
| ENSMUSG00000092341 | -1,150 | 0,000 | Malat1 |
| ENSMUSG00000026870 | -1,152 | 0,023 | Cutal |
| ENSMUSG00000032503 | -1,153 | 0,000 | Arpp21 |
| ENSMUSG00000041703 | -1,154 | 0,005 | Zic5 |
| ENSMUSG00000046618 | -1,156 | 0,000 | Olfml2a |
| ENSMUSG00000031661 | -1,157 | 0,000 | Nkd1 |
| ENSMUSG00000045875 | -1,161 | 0,038 | Adra1a |
| ENSMUSG00000098462 | -1,164 | 0,002 | Gm27694 |
| ENSMUSG00000112808 | -1,164 | 0,022 | Gm4739 |
| ENSMUSG00000086008 | -1,166 | 0,026 | Gm8817 |
| ENSMUSG00000007080 | -1,167 | 0,000 | Pole |
| ENSMUSG00000113515 | -1,167 | 0,011 | AL645745.1 |
| ENSMUSG00000090291 | -1,167 | 0,019 | Lrrc10b |
| ENSMUSG00000049538 | -1,170 | 0,000 | Adamts16 |
| ENSMUSG00000025083 | -1,172 | 0,000 | Afap1l2 |
| ENSMUSG00000026429 | -1,175 | 0,000 | Ube2t |
| ENSMUSG00000034303 | -1,178 | 0,002 | Ccdc15 |
| ENSMUSG00000108120 | -1,180 | 0,030 | 9930120I10Rik |
| ENSMUSG00000111264 | -1,184 | 0,005 | E030022I16Rik |
| ENSMUSG00000034648 | -1,185 | 0,001 | Lrrn1 |
| ENSMUSG00000048424 | -1,187 | 0,000 | Ranbp3l |
| ENSMUSG00000037379 | -1,193 | 0,004 | Spon2 |
| ENSMUSG00000060843 | -1,194 | 0,022 | Ctnna3 |
| ENSMUSG00000026447 | -1,195 | 0,002 | Pik3c2b |
| ENSMUSG00000071489 | -1,196 | 0,048 | Ptgdr |
| ENSMUSG00000030322 | -1,199 | 0,000 | Mbd4 |
| ENSMUSG00000114456 | -1,201 | 0,008 | Hist1h2bh |
| ENSMUSG00000030708 | -1,202 | 0,000 | Dnajb13 |
| ENSMUSG00000039193 | -1,203 | 0,023 | Nlrc4 |
| ENSMUSG00000037921 | -1,204 | 0,001 | Ddx60 |
| ENSMUSG00000045215 | -1,205 | 0,017 | Asxl3 |
| ENSMUSG00000114515 | -1,206 | 0,000 | Aldoa |
| ENSMUSG00000105311 | -1,213 | 0,001 | Gm44574 |
| ENSMUSG00000103238 | -1,217 | 0,002 | Gm36527 |
| ENSMUSG00000030110 | -1,221 | 0,000 | Ret |
| ENSMUSG00000022519 | -1,222 | 0,000 | Srl |

| | | | |
|---|---|---|---|
| ENSMUSG00000104814 | -1,225 | 0,001 | Gm42979 |
| ENSMUSG00000042489 | -1,228 | 0,000 | Clspn |
| ENSMUSG00000067714 | -1,234 | 0,000 | Lpar5 |
| ENSMUSG00000093133 | -1,238 | 0,000 | Gm25994 |
| ENSMUSG00000084821 | -1,240 | 0,007 | Gm15880 |
| ENSMUSG00000055116 | -1,242 | 0,000 | Arntl |
| ENSMUSG00000090799 | -1,244 | 0,000 | Klhl33 |
| ENSMUSG00000050334 | -1,247 | 0,000 | C130071C03Rik |
| ENSMUSG00000098955 | -1,248 | 0,001 | Mir6935 |
| ENSMUSG00000078144 | -1,251 | 0,000 | Capns2 |
| ENSMUSG00000061577 | -1,253 | 0,004 | Adgrg5 |
| ENSMUSG00000072812 | -1,256 | 0,044 | Ahnak2 |
| ENSMUSG00000021678 | -1,257 | 0,027 | F2rl1 |
| ENSMUSG00000054074 | -1,257 | 0,000 | Skida1 |
| ENSMUSG00000085995 | -1,261 | 0,008 | Gm2788 |
| ENSMUSG00000053626 | -1,261 | 0,012 | Tll1 |
| ENSMUSG00000075256 | -1,272 | 0,028 | Cerkl |
| ENSMUSG00000025432 | -1,274 | 0,004 | Avil |
| ENSMUSG00000028834 | -1,275 | 0,008 | Trim63 |
| ENSMUSG00000019359 | -1,276 | 0,005 | Gdpd2 |
| ENSMUSG00000024990 | -1,279 | 0,016 | Rbp4 |
| ENSMUSG00000033033 | -1,281 | 0,000 | Calhm2 |
| ENSMUSG00000059246 | -1,282 | 0,028 | Foxb1 |
| ENSMUSG00000105201 | -1,286 | 0,004 | Gm43362 |
| ENSMUSG00000110631 | -1,291 | 0,001 | Gm42047 |
| ENSMUSG00000042793 | -1,296 | 0,006 | Lgr6 |
| ENSMUSG00000110540 | -1,297 | 0,008 | Gm45743 |
| ENSMUSG00000040828 | -1,297 | 0,046 | Catsperd |
| ENSMUSG00000107877 | -1,297 | 0,045 | Gm43951 |
| ENSMUSG00000020080 | -1,304 | 0,035 | Hkdc1 |
| ENSMUSG00000026778 | -1,306 | 0,000 | Prkcq |
| ENSMUSG00000103593 | -1,308 | 0,007 | Gm37352 |
| ENSMUSG00000028392 | -1,308 | 0,000 | Bspry |
| ENSMUSG00000110087 | -1,317 | 0,045 | Gm45309 |
| ENSMUSG00000041361 | -1,319 | 0,000 | Myzap |
| ENSMUSG00000004110 | -1,320 | 0,000 | Cacna1e |
| ENSMUSG00000113771 | -1,322 | 0,029 | Gm36529 |
| ENSMUSG00000020682 | -1,326 | 0,000 | Mmp28 |
| ENSMUSG00000054641 | -1,329 | 0,000 | Mmrn1 |
| ENSMUSG00000090582 | -1,330 | 0,003 | Gm17024 |
| ENSMUSG00000105742 | -1,332 | 0,002 | Gm42748 |
| ENSMUSG00000115624 | -1,333 | 0,011 | AC154640.4 |
| ENSMUSG00000113894 | -1,336 | 0,008 | AC129597.1 |
| ENSMUSG00000106825 | -1,337 | 0,000 | 2510016D11Rik |
| ENSMUSG00000103285 | -1,337 | 0,010 | Gm37274 |
| ENSMUSG00000025586 | -1,338 | 0,000 | Cpeb1 |
| ENSMUSG00000046179 | -1,338 | 0,000 | E2f8 |

| | | | |
|---|---|---|---|
| ENSMUSG00000111030 | -1,344 | 0,033 | AC151971.2 |
| ENSMUSG00000032377 | -1,349 | 0,000 | Plscr4 |
| ENSMUSG00000074207 | -1,353 | 0,001 | Adh1 |
| ENSMUSG00000003545 | -1,356 | 0,000 | Fosb |
| ENSMUSG00000034858 | -1,357 | 0,000 | Fam214a |
| ENSMUSG00000024600 | -1,359 | 0,000 | Slc27a6 |
| ENSMUSG00000058897 | -1,362 | 0,000 | Col25a1 |
| ENSMUSG00000105950 | -1,372 | 0,035 | Gm43679 |
| ENSMUSG00000075770 | -1,373 | 0,047 | Gm22589 |
| ENSMUSG00000045498 | -1,377 | 0,018 | Pcdhb3 |
| ENSMUSG00000103558 | -1,381 | 0,031 | Gm38220 |
| ENSMUSG00000058743 | -1,382 | 0,002 | Kcnj14 |
| ENSMUSG00000020623 | -1,384 | 0,000 | Map2k6 |
| ENSMUSG00000070803 | -1,388 | 0,013 | Cited4 |
| ENSMUSG00000098267 | -1,388 | 0,002 | Mir7060 |
| ENSMUSG00000105632 | -1,391 | 0,004 | Gm43272 |
| ENSMUSG00000047592 | -1,393 | 0,038 | Nxpe5 |
| ENSMUSG00000020154 | -1,394 | 0,002 | Ptprb |
| ENSMUSG00000076618 | -1,395 | 0,016 | Ighj4 |
| ENSMUSG00000041479 | -1,396 | 0,000 | Syt15 |
| ENSMUSG00000063430 | -1,398 | 0,017 | Wscd2 |
| ENSMUSG00000035187 | -1,408 | 0,001 | Nkx6-1 |
| ENSMUSG00000040016 | -1,408 | 0,000 | Ptger3 |
| ENSMUSG00000025255 | -1,408 | 0,004 | Zfhx4 |
| ENSMUSG00000105769 | -1,412 | 0,009 | Gm43351 |
| ENSMUSG00000105748 | -1,414 | 0,009 | Gm43088 |
| ENSMUSG00000032719 | -1,415 | 0,008 | Sbspon |
| ENSMUSG00000115317 | -1,417 | 0,042 | AC139845.1 |
| ENSMUSG00000108551 | -1,419 | 0,007 | Gm20274 |
| ENSMUSG00000107000 | -1,423 | 0,000 | Gm43481 |
| ENSMUSG00000107200 | -1,425 | 0,015 | Gm43039 |
| ENSMUSG00000040710 | -1,426 | 0,000 | St8sia4 |
| ENSMUSG00000098790 | -1,431 | 0,040 | Mir6942 |
| ENSMUSG00000027939 | -1,432 | 0,001 | Nup210l |
| ENSMUSG00000107756 | -1,433 | 0,005 | Gm44164 |
| ENSMUSG00000004359 | -1,434 | 0,000 | Spic |
| ENSMUSG00000097151 | -1,440 | 0,034 | Gm26514 |
| ENSMUSG00000090399 | -1,441 | 0,010 | Gm38399 |
| ENSMUSG00000115417 | -1,443 | 0,009 | AC131586.4 |
| ENSMUSG00000079049 | -1,460 | 0,049 | Serpinb1c |
| ENSMUSG00000085399 | -1,466 | 0,002 | Foxd2os |
| ENSMUSG00000003882 | -1,475 | 0,002 | Il7r |
| ENSMUSG00000022676 | -1,482 | 0,000 | Snai2 |
| ENSMUSG00000081683 | -1,482 | 0,000 | Fzd10 |
| ENSMUSG00000087579 | -1,492 | 0,031 | Hectd2os |
| ENSMUSG00000006241 | -1,502 | 0,000 | Ccdc159 |
| ENSMUSG00000106416 | -1,505 | 0,009 | Gm5857 |

| | | | |
|---|---|---|---|
| ENSMUSG00000114278 | -1,514 | 0,015 | AC157931.1 |
| ENSMUSG00000022123 | -1,517 | 0,000 | Scel |
| ENSMUSG00000112358 | -1,519 | 0,001 | AC108802.1 |
| ENSMUSG00000049872 | -1,520 | 0,002 | Fam26e |
| ENSMUSG00000113047 | -1,526 | 0,000 | CT009718.2 |
| ENSMUSG00000105556 | -1,537 | 0,021 | Gm43080 |
| ENSMUSG00000108841 | -1,540 | 0,013 | Frmpd2 |
| ENSMUSG00000052951 | -1,551 | 0,033 | C130021I20Rik |
| ENSMUSG00000105929 | -1,551 | 0,039 | Gm43693 |
| ENSMUSG00000104546 | -1,553 | 0,000 | Gm43858 |
| ENSMUSG00000054999 | -1,555 | 0,000 | Naaladl1 |
| ENSMUSG00000102964 | -1,560 | 0,007 | 9430034N14Rik |
| ENSMUSG00000028838 | -1,562 | 0,020 | Extl1 |
| ENSMUSG00000069270 | -1,563 | 0,000 | Hist1h2ac |
| ENSMUSG00000046480 | -1,563 | 0,007 | Scn4b |
| ENSMUSG00000108825 | -1,563 | 0,000 | Gm45838 |
| ENSMUSG00000115149 | -1,575 | 0,041 | AC112142.1 |
| ENSMUSG00000104342 | -1,578 | 0,032 | Gm36401 |
| ENSMUSG00000106665 | -1,580 | 0,006 | Gm43389 |
| ENSMUSG00000092457 | -1,582 | 0,027 | Gm8835 |
| ENSMUSG00000043903 | -1,587 | 0,001 | Zfp469 |
| ENSMUSG00000035681 | -1,587 | 0,000 | Kcnc2 |
| ENSMUSG00000037846 | -1,593 | 0,000 | Rtkn2 |
| ENSMUSG00000004872 | -1,601 | 0,028 | Pax3 |
| ENSMUSG00000096351 | -1,612 | 0,004 | Samd11 |
| ENSMUSG00000110422 | -1,619 | 0,020 | Gm17999 |
| ENSMUSG00000108154 | -1,626 | 0,017 | Gm44033 |
| ENSMUSG00000032940 | -1,627 | 0,039 | Rbm11 |
| ENSMUSG00000104347 | -1,629 | 0,025 | Gm37689 |
| ENSMUSG00000005640 | -1,631 | 0,026 | Insrr |
| ENSMUSG00000026308 | -1,641 | 0,036 | Klhl30 |
| ENSMUSG00000022262 | -1,654 | 0,011 | Dnah5 |
| ENSMUSG00000102555 | -1,657 | 0,038 | 6430511E19Rik |
| ENSMUSG00000044156 | -1,659 | 0,027 | Hepacam2 |
| ENSMUSG00000021356 | -1,668 | 0,000 | Irf4 |
| ENSMUSG00000108064 | -1,673 | 0,000 | Gm44423 |
| ENSMUSG00000103509 | -1,676 | 0,021 | Gm38372 |
| ENSMUSG00000113836 | -1,693 | 0,028 | AC124426.1 |
| ENSMUSG00000103507 | -1,695 | 0,005 | Gm38375 |
| ENSMUSG00000062372 | -1,701 | 0,030 | Otof |
| ENSMUSG00000046321 | -1,707 | 0,000 | Hs3st2 |
| ENSMUSG00000018427 | -1,716 | 0,000 | Ypel2 |
| ENSMUSG00000057132 | -1,721 | 0,016 | Rpgrip1 |
| ENSMUSG00000049630 | -1,725 | 0,037 | C1ql3 |
| ENSMUSG00000084408 | -1,727 | 0,013 | Gm11870 |
| ENSMUSG00000035095 | -1,729 | 0,005 | Fam167a |
| ENSMUSG00000026012 | -1,745 | 0,008 | Cd28 |

| | | | |
|---|---|---|---|
| ENSMUSG00000074629 | -1,777 | 0,014 | 4930518I15Rik |
| ENSMUSG00000030324 | -1,778 | 0,004 | Rho |
| ENSMUSG00000106099 | -1,784 | 0,006 | Gm42664 |
| ENSMUSG00000022357 | -1,788 | 0,000 | Klhl38 |
| ENSMUSG00000099398 | -1,798 | 0,002 | Ms4a14 |
| ENSMUSG00000032291 | -1,827 | 0,005 | Crabp1 |
| ENSMUSG00000103047 | -1,831 | 0,035 | Gm37310 |
| ENSMUSG00000047419 | -1,839 | 0,000 | Cmya5 |
| ENSMUSG00000097876 | -1,847 | 0,034 | Gm16892 |
| ENSMUSG00000097081 | -1,854 | 0,005 | Gm10425 |
| ENSMUSG00000038276 | -1,869 | 0,002 | Asic3 |
| ENSMUSG00000035948 | -1,873 | 0,000 | Acss3 |
| ENSMUSG00000103174 | -1,875 | 0,021 | Gm37168 |
| ENSMUSG00000086296 | -1,879 | 0,041 | D030055H07Rik |
| ENSMUSG00000039252 | -1,882 | 0,000 | Lgi2 |
| ENSMUSG00000049115 | -1,890 | 0,035 | Agtr1a |
| ENSMUSG00000079177 | -1,891 | 0,038 | Fam228a |
| ENSMUSG00000064427 | -1,905 | 0,004 | Gm22748 |
| ENSMUSG00000038756 | -1,910 | 0,002 | Ttll6 |
| ENSMUSG00000058248 | -1,942 | 0,039 | Kcnh1 |
| ENSMUSG00000107215 | -1,951 | 0,007 | Gm43197 |
| ENSMUSG00000103133 | -1,961 | 0,005 | Gm37303 |
| ENSMUSG00000034810 | -1,963 | 0,009 | Scn7a |
| ENSMUSG00000057777 | -1,968 | 0,039 | Mab21l2 |
| ENSMUSG00000111390 | -2,030 | 0,000 | CT025660.2 |
| ENSMUSG00000020061 | -2,042 | 0,028 | Mybpc1 |
| ENSMUSG00000039313 | -2,069 | 0,000 | AF529169 |
| ENSMUSG00000058934 | -2,077 | 0,004 | Igf1os |
| ENSMUSG00000115388 | -2,150 | 0,018 | EPPK1 |
| ENSMUSG00000043719 | -2,177 | 0,000 | Col6a6 |
| ENSMUSG00000039691 | -2,178 | 0,000 | Tspan10 |
| ENSMUSG00000046634 | -2,193 | 0,018 | Pkd1l1 |
| ENSMUSG00000038775 | -2,212 | 0,000 | Vill |
| ENSMUSG00000085783 | -2,223 | 0,017 | Gm9816 |
| ENSMUSG00000005320 | -2,228 | 0,000 | Fgfr4 |
| ENSMUSG00000073733 | -2,275 | 0,005 | Rsg1 |
| ENSMUSG00000048782 | -2,314 | 0,001 | Insc |
| ENSMUSG00000093226 | -2,334 | 0,040 | Gm25749 |
| ENSMUSG00000017767 | -2,379 | 0,000 | Spata25 |
| ENSMUSG00000033634 | -2,389 | 0,024 | Nat8f2 |
| ENSMUSG00000108970 | -2,393 | 0,027 | Gm45906 |
| ENSMUSG00000097623 | -2,425 | 0,031 | B230323A14Rik |
| ENSMUSG00000079173 | -2,433 | 0,002 | Zan |
| ENSMUSG00000060284 | -2,440 | 0,001 | Sp7 |
| ENSMUSG00000109896 | -2,474 | 0,024 | Gm31409 |
| ENSMUSG00000092229 | -2,477 | 0,022 | Gm5977 |
| ENSMUSG00000040372 | -2,520 | 0,014 | Gpr63 |

| | | | |
|---|---|---|---|
| ENSMUSG00000100832 | -2,565 | 0,050 | Gm29260 |
| ENSMUSG00000018486 | -2,626 | 0,008 | Wnt9b |
| ENSMUSG00000073130 | -2,628 | 0,001 | Gm1141 |
| ENSMUSG00000097021 | -2,656 | 0,000 | 4933433G15Rik |
| ENSMUSG00000097336 | -2,693 | 0,000 | Fendrr |
| ENSMUSG00000029431 | -2,717 | 0,018 | B3gnt4 |
| ENSMUSG00000110997 | -2,719 | 0,008 | 2610028D06Rik |
| ENSMUSG00000110397 | -2,906 | 0,000 | Gm45540 |
| ENSMUSG00000067578 | -3,163 | 0,013 | Cbln4 |
| ENSMUSG00000085882 | -4,015 | 0,000 | 2610507I01Rik |
| ENSMUSG00000039628 | -4,018 | 0,002 | Hs3st6 |
| ENSMUSG00000021732 | -4,447 | 0,015 | Fgf10 |
| ENSMUSG00000021590 | -5,110 | 0,019 | Spata9 |

**Supplementary table 4: mRNA values of phenotype associated genes used to create Figure 6B.**

| | CEREBRUM IL1 vs PBS | | | | | CEREBELLUM IL1 vs PBS | | | | |
|---|---|---|---|---|---|---|---|---|---|---|
| | Mean CR PBS | Mean CR IL1 | log2FoldChange | Fold Change | p adj value | Mean CB PBS | Mean CB IL1 | log2FoldChange | Fold Change | p adj value |
| Ptgs2 | 634,872 | 852,267 | -0,037 | 0,974 | 9,623E-01 | 417,839 | 535,091 | 0,822 | 1,767 | 1,641E-01 |
| Nos2 | 85,514 | 323,142 | 1,867 | 3,648 | 4,745E-06 | 66,809 | 195,838 | 1,603 | 3,038 | 2,230E-04 |
| Tnf | 8296,125 | 9152,485 | -0,057 | 0,961 | 9,537E-01 | 9211,302 | 9984,824 | 0,315 | 1,244 | 7,799E-01 |
| Il6 | 55,840 | 179,689 | 1,741 | 3,342 | 2,680E-05 | 19,942 | 93,231 | 2,153 | 4,448 | 1,100E-06 |
| Cd86 | 5784,204 | 7087,705 | 0,258 | 1,196 | 6,782E-01 | 7673,717 | 8752,218 | 0,225 | 1,168 | 7,954E-01 |
| Cd206 | 77056,546 | 106254,064 | 0,516 | 1,430 | 4,149E-01 | 57893,656 | 77797,566 | 0,374 | 1,296 | 6,677E-01 |
| Arg1 | 64,446 | 31,973 | -1,097 | 0,467 | 3,926E-01 | 0,679 | 0,808 | 0,575 | 1,490 | 8,461E-01 |
| Igf1 | 3660,409 | 2819,899 | -0,755 | 0,593 | 1,213E-01 | 4844,813 | 3048,979 | -0,290 | 0,818 | 7,048E-01 |
| Lgals3 | 844,245 | 1699,667 | 1,001 | 2,001 | 1,040E-06 | 988,471 | 1558,167 | 0,666 | 1,587 | 3,286E-03 |
| Il4ra | 16328,818 | 42707,224 | 1,328 | 2,511 | 6,245E-12 | 15932,171 | 31384,115 | 1,037 | 2,052 | 3,085E-07 |
| Il1rn | 239,780 | 722,219 | 1,530 | 2,888 | 2,471E-08 | 344,748 | 527,274 | 0,673 | 1,595 | 4,428E-02 |
| Socs3 | 11252,499 | 14539,621 | 0,232 | 1,174 | 3,158E-01 | 6853,289 | 8752,910 | 0,491 | 1,405 | 2,016E-02 |
| Sphk1 | 77,297 | 97,485 | 0,202 | 1,150 | 7,120E-01 | 23,959 | 23,644 | 0,137 | 1,100 | 8,781E-01 |

**Supplementary table 5:**
**Gene list used to build the DEG heat map and analysis for clusters (Figure 6C).**

| *Gene name* | *CR_ PBS* | *CR_ IL1* | *CB_ PBS* | *CB_ IL1* |
|---|---|---|---|---|
| Nipal1 | 0 | 1,74229447 | 0 | 2,88945588 |
| Serpina3g | 42,5731542 | 1075,17095 | 30,163386 | 1222,10698 |
| Bmp8a | 0,27850422 | 3,17637015 | 0 | 2,03865217 |
| Iigp1 | 31,6976738 | 484,819951 | 16,2763238 | 448,538975 |
| Tgtp2 | 7,36592876 | 66,981111 | 3,09389379 | 67,9207943 |
| Prss16 | 11,5669603 | 285,256612 | 4,37629728 | 110,298806 |
| Ccl17 | 1,84085398 | 30,0343532 | 0,38720443 | 17,6883558 |
| Foxd3 | 1,7284899 | 0,58134746 | 0 | 6,35058097 |
| Plet1 | 5,19335138 | 130,522177 | 4,97306319 | 105,347239 |
| Spr-ps1 | 3,10636624 | 10,4280879 | 0,28883159 | 8,79422977 |
| Tgtp1 | 3,05235703 | 51,2585694 | 2,20222287 | 39,4145917 |
| Mirt2 | 11,1631276 | 42,3610414 | 0,93947794 | 16,9860902 |
| Mpz | 36,2167029 | 24,1848916 | 29,7358892 | 661,872512 |
| Slc28a3 | 0,77865717 | 17,3362508 | 0,67879482 | 5,2736339 |
| Serpina3h | 28,7795977 | 678,485681 | 33,0675781 | 399,97385 |
| Jaml | 37,2038877 | 1384,87321 | 46,5858409 | 414,415169 |
| Gzma | 16,7705398 | 41,9843723 | 2,56348181 | 60,124744 |
| Serpina3e-ps | 6,29640715 | 113,756834 | 5,66833071 | 71,4289031 |
| Steap4 | 61,6969308 | 641,15698 | 24,5707973 | 335,453198 |
| Ms4a4b | 10,9240359 | 29,0612765 | 2,2213256 | 22,2709444 |
| Gm5416 | 39,2335885 | 276,263002 | 11,7801628 | 124,502136 |
| Serpina3i | 13,4453085 | 285,056831 | 14,6687362 | 144,14846 |
| Apol10b | 5,88189453 | 18,1094061 | 2,29154998 | 19,7506508 |
| Serpina3f | 17,219472 | 251,143012 | 14,1760303 | 189,556769 |
| Gbp4 | 174,618399 | 842,081986 | 95,2382737 | 869,368465 |
| Cxcl11 | 4,12383461 | 36,9628224 | 4,71216654 | 32,6927362 |
| Gbp10 | 11,4501225 | 58,2070921 | 7,65733519 | 64,0796169 |
| Rnase2a | 27,3326586 | 71,2569522 | 1,32944117 | 14,0113612 |
| Il1r2 | 155,248679 | 1492,24788 | 85,1289719 | 811,413805 |
| Zbtb16 | 37,4882934 | 306,935168 | 26,0541389 | 186,980137 |
| Il2rb | 6,81741592 | 30,4850416 | 3,82070915 | 21,7319455 |
| Cxcl9 | 20,0566008 | 85,5313956 | 6,40915359 | 56,2005473 |
| Plet1os | 4,01147053 | 40,8248696 | 5,96310694 | 55,1909425 |
| Nrg4 | 4,54739971 | 29,5856199 | 3,44976391 | 27,5296112 |
| Gm807 | 1,75498534 | 16,6942587 | 0,67879482 | 6,79715055 |
| 2010005H15I | 90,1597164 | 456,234286 | 48,6723659 | 348,009932 |
| Rab44 | 59,457162 | 355,433334 | 27,2441582 | 181,41108 |
| Cpm | 11,5056263 | 85,775998 | 8,54603378 | 56,1483565 |
| Gm9733 | 16,1619347 | 47,7507923 | 4,63621105 | 32,3154703 |
| E230001N04I | 7,11391513 | 37,8398723 | 4,80204947 | 40,5366457 |
| Upp1 | 34,1031081 | 211,458844 | 16,5705365 | 136,96468 |
| Klf9 | 128,671324 | 932,525486 | 69,9512785 | 388,744881 |

| | | | | |
|---|---|---|---|---|
| Gng13 | 4,99087328 | 3,3177586 | 0,67879482 | 16,3308997 |
| Vcam1 | 736,957267 | 4706,6674 | 597,876984 | 3443,31195 |
| Orm1 | 6,00666617 | 50,4179385 | 3,55408139 | 24,41598 |
| Ifit3b | 75,9306222 | 95,6161885 | 58,9357295 | 469,295296 |
| Rnase1 | 0,50267065 | 4,64553595 | 0,72660184 | 3,56926364 |
| Areg | 12,797634 | 42,1800204 | 2,83045187 | 26,6863034 |
| Gm10143 | 5,53781576 | 3,74434351 | 0,28883159 | 4,17757697 |
| Sell | 151,644402 | 604,758021 | 70,9644479 | 427,638892 |
| Kcna5 | 8,93849694 | 70,9826187 | 7,3057062 | 45,3920957 |
| Jdp2 | 1021,5941 | 4858,33341 | 857,32504 | 5154,07072 |
| Nlrp12 | 25,4380327 | 109,171902 | 12,1865987 | 63,1941221 |
| Kcna1 | 0,53844688 | 0,94977853 | 1,08544428 | 5,9134593 |
| Gm10603 | 3,32969046 | 10,2903105 | 3,09389379 | 17,4336514 |
| Csf3 | 11,5730662 | 18,8957189 | 3,06574532 | 19,6557093 |
| Pla1a | 8,72808497 | 50,2420879 | 8,43428933 | 42,856053 |
| Il2ra | 2,21511669 | 10,7240757 | 0,96486761 | 6,52274917 |
| Retnlg | 653,07884 | 3475,90468 | 312,881314 | 1680,74137 |
| Mc5r | 1,84509935 | 7,06416052 | 1,44725904 | 3,62265592 |
| Arc | 101,333306 | 473,904458 | 186,003063 | 879,923684 |
| Fcnaos | 7,65717396 | 18,130271 | 2,46566479 | 10,9823988 |
| Ifit1 | 103,757141 | 150,25367 | 84,68938 | 453,34525 |
| Tomt | 15,2362607 | 12,9802032 | 5,99457003 | 23,9353022 |
| Stra6 | 285,789933 | 714,668677 | 51,0155916 | 292,936346 |
| Trim69 | 12,6179677 | 16,8516066 | 24,7218902 | 118,014918 |
| Gm10524 | 8,42616959 | 24,1246242 | 1,39966554 | 12,9497512 |
| Gm12250 | 32,1677243 | 104,120168 | 19,8471198 | 114,42695 |
| Hif3a | 871,493914 | 4336,38175 | 777,142449 | 3412,28419 |
| Mrgpra2b | 6,94297273 | 24,7443251 | 3,03738333 | 14,7569612 |
| Fpr2 | 157,978378 | 487,585847 | 62,9455802 | 302,722462 |
| Gm16194 | 5,83879835 | 31,102898 | 4,05578908 | 20,460223 |
| Cd3d | 22,4195391 | 90,1861908 | 9,77821376 | 51,3008745 |
| Fkbp5 | 3102,02343 | 17314,4976 | 3345,14613 | 15608,9787 |
| Gbp2 | 549,107564 | 2644,27788 | 548,184624 | 2278,06228 |
| Mrap | 15,0287472 | 79,4450541 | 12,436625 | 50,655431 |
| Trbc2 | 3,59215082 | 7,6305492 | 0,70121217 | 8,27457516 |
| Map3k6 | 110,579845 | 433,204385 | 74,1065271 | 367,92297 |
| Lbx1 | 301,58835 | 1261,69436 | 252,903589 | 1091,72999 |
| Gm26473 | 17,5253856 | 63,310579 | 16,8727914 | 74,1984365 |
| Il6 | 55,8399344 | 179,689106 | 19,9418797 | 93,2312525 |
| Gm15675 | 13,4505819 | 36,9911782 | 2,77967253 | 8,68176785 |
| Oas3 | 44,1171094 | 113,250343 | 24,5917607 | 114,183277 |
| Ccl5 | 32,3087444 | 61,3293374 | 11,3955884 | 52,3316736 |
| Capn11 | 0,7138678 | 3,43275072 | 0,67879482 | 2,98006767 |
| Vwc2 | 1,22330154 | 4,92041254 | 2,2216679 | 5,57009084 |
| Hdc | 176,059513 | 835,69382 | 107,332923 | 525,544672 |
| Rdh12 | 36,8464304 | 134,558704 | 15,2585022 | 67,3246392 |

| | | | | |
|---|---|---|---|---|
| Hs3st3b1 | 133,921682 | 1150,41557 | 141,598297 | 461,214076 |
| Serpinb10 | 23,119738 | 65,2244366 | 11,4390695 | 52,2168274 |
| Gm43181 | 16,905044 | 56,2686101 | 6,54387122 | 28,3010222 |
| Slfn1 | 84,295015 | 394,376496 | 42,364399 | 199,091922 |
| Cdh3 | 24,9990712 | 29,7526872 | 0 | 4,54063068 |
| Gbp2b | 52,1251823 | 120,033461 | 37,7907765 | 165,711514 |
| Il1f9 | 52,9167697 | 176,467072 | 26,2470628 | 106,676229 |
| Mefv | 24,9769637 | 67,984765 | 11,5170632 | 48,175395 |
| Creb5 | 110,462027 | 629,08735 | 53,8098605 | 197,525802 |
| 1700016G22F | 3,27904116 | 5,08049491 | 12,2658201 | 50,0262859 |
| Ly6i | 17,9165915 | 56,2883301 | 14,6051534 | 51,8042725 |
| Vdr | 45,8143253 | 396,694712 | 41,0236807 | 106,557401 |
| Atp1a4 | 2,56265565 | 5,01821496 | 0,38720443 | 6,01686584 |
| Gm43302 | 51,452626 | 165,860064 | 40,6113802 | 153,764694 |
| Ddit4 | 525,325662 | 2113,47128 | 488,865701 | 2122,91217 |
| Prss23 | 113,910226 | 386,621212 | 51,1769597 | 248,069077 |
| Gm20109 | 4,87543462 | 15,2690748 | 4,41760384 | 21,2339894 |
| Gm26586 | 4,18979478 | 18,2022405 | 3,40238393 | 14,4102481 |
| Ifi205 | 9,88408428 | 36,5076975 | 9,72420454 | 33,736999 |
| Gm45774 | 4,1793432 | 13,5029552 | 2,07846036 | 11,5481552 |
| Fam107a | 67,9963098 | 186,169968 | 22,8420849 | 97,5625698 |
| Gm8463 | 3,12721236 | 9,84582513 | 26,7329854 | 110,97232 |
| Ttc39c | 18,718941 | 68,2773855 | 11,2841862 | 52,2333627 |
| I830127L07Ri | 70,7217177 | 236,140645 | 40,2739409 | 157,279755 |
| S100a7a | 1,25231468 | 6,40720039 | 1,37724819 | 2,33788056 |
| Prss12 | 3,3989535 | 10,8169999 | 2,29706757 | 13,1386772 |
| Cd300lf | 732,03756 | 4211,67566 | 427,760059 | 1822,92826 |
| Gbp5 | 561,405472 | 2022,42425 | 571,006539 | 1668,53163 |
| B9d1os | 2,33055535 | 1,47503982 | 0,38720443 | 6,71717724 |
| Trim30b | 133,665134 | 312,536211 | 62,1355302 | 237,998315 |
| Ly6g | 42,8118046 | 171,654211 | 25,9648117 | 96,791479 |
| Adam8 | 548,525511 | 2225,35999 | 335,499981 | 1200,55811 |
| Ifnlr1 | 18,2712221 | 42,8264863 | 6,27117004 | 19,925569 |
| Gda | 230,482286 | 889,609063 | 136,48366 | 493,047999 |
| Galnt15 | 48,2479168 | 148,035679 | 30,3494625 | 112,477452 |
| Dok2 | 1194,16002 | 4843,66297 | 812,23535 | 3293,73843 |
| Ccl12 | 4643,11216 | 20025,656 | 3474,69142 | 14083,0814 |
| Cx3cl1 | 139,702223 | 722,274901 | 128,134107 | 470,9682 |
| Arg2 | 45,1192898 | 130,797254 | 23,9251567 | 103,381162 |
| Gm28592 | 1,04111753 | 5,80573085 | 1,86282547 | 4,18899515 |
| Mmp8 | 481,320609 | 1366,12867 | 264,979273 | 890,362597 |
| Cd55 | 192,627737 | 491,318853 | 58,4648197 | 225,158766 |
| Gbp6 | 59,1449622 | 169,881485 | 40,8244698 | 137,262244 |
| Ifit3 | 264,782988 | 362,961569 | 196,172068 | 732,076621 |
| Eomes | 3,26238339 | 5,4246778 | 0,74901919 | 2,07154942 |
| Ramp3 | 4,41357559 | 19,8196646 | 3,94128582 | 12,7257749 |

| | | | | |
|---|---|---|---|---|
| Htr2a | 7,74616452 | 13,1812058 | 1,0884166 | 5,89119943 |
| Il15ra | 325,298016 | 1196,55942 | 194,57547 | 631,279489 |
| Clec4d | 166,63836 | 823,131438 | 108,589317 | 406,191374 |
| Arhgef37 | 19,9000652 | 42,1656252 | 10,170807 | 41,53833 |
| Acod1 | 51,1997872 | 179,01235 | 25,6571757 | 108,221008 |
| Gpr141 | 55,9723928 | 148,8749 | 29,3672804 | 110,05239 |
| Chrm3 | 5,54168037 | 17,6154193 | 3,7726886 | 11,8539008 |
| Wfdc21 | 162,602591 | 381,764521 | 54,6985358 | 216,444479 |
| Clec4e | 230,976405 | 509,823798 | 110,259699 | 354,957609 |
| Nt5e | 100,564992 | 142,356447 | 14,3655061 | 57,5213969 |
| Chil5 | 35,9043317 | 74,5884233 | 19,9831423 | 63,6446946 |
| Aqp9 | 11,3952279 | 36,6875822 | 8,91413549 | 33,7032597 |
| Mir147 | 11,1089422 | 32,0150864 | 4,48761469 | 16,7705918 |
| Ifit1bl1 | 21,120492 | 28,1549714 | 16,7019756 | 80,4865472 |
| Esyt3 | 12,5875603 | 9,56376823 | 2,97663174 | 12,509087 |
| A930030B08F | 6,59235917 | 16,9274537 | 8,13938432 | 23,7592032 |
| Gm26448 | 9,01587482 | 11,7092641 | 4,76549116 | 16,2394878 |
| Socs2 | 151,888416 | 1361,10911 | 206,175705 | 453,159004 |
| Stfa2l1 | 736,301286 | 1999,07973 | 308,783301 | 1087,66128 |
| Syde2 | 4,58200514 | 12,5227761 | 2,3448746 | 6,27769277 |
| Ocstamp | 13,002249 | 84,3507842 | 14,3304865 | 42,0738044 |
| Tarm1 | 45,1404979 | 147,485326 | 19,7881928 | 67,104628 |
| A330023F24F | 10,0919786 | 16,2671103 | 2,63073386 | 14,3877914 |
| Ms4a8a | 14,0814322 | 36,0577475 | 7,9714717 | 27,959638 |
| Abtb2 | 68,0319196 | 327,960351 | 58,7977493 | 163,049185 |
| Slc16a14 | 23,1135743 | 66,0103795 | 17,593433 | 56,1335173 |
| Pdzd3 | 19,1899854 | 41,3269013 | 16,1683494 | 67,0433075 |
| Tmem252 | 178,78303 | 550,804219 | 92,1883965 | 333,806404 |
| Ifi44 | 22,8483776 | 30,448843 | 16,6022492 | 50,5236688 |
| F10 | 94,2237225 | 322,822611 | 50,5325079 | 188,831655 |
| Runx3 | 55,8000269 | 222,841674 | 19,675635 | 64,9199973 |
| Prok2 | 65,9997092 | 198,471123 | 27,3836131 | 99,1407606 |
| Hspa1b | 993,758933 | 4120,49528 | 374,466821 | 934,13305 |
| Gm38150 | 3,5457469 | 13,1530294 | 4,12877225 | 12,6818392 |
| Gm42928 | 1,61416984 | 3,69206367 | 1,01543343 | 3,64845835 |
| Gm5483 | 940,424101 | 2899,28086 | 392,767903 | 1517,27028 |
| Sh3pxd2b | 853,566234 | 3410,20722 | 795,65829 | 2183,44501 |
| Mx1 | 356,81148 | 328,72851 | 240,114331 | 677,118484 |
| Gm14964 | 1,80680543 | 8,75624284 | 2,61128883 | 6,60961297 |
| Gm11620 | 14,6810273 | 11,1558586 | 13,8863446 | 40,5783993 |
| Scn2b | 21,211396 | 52,8476655 | 13,7470075 | 44,4851648 |
| Slc39a14 | 882,251475 | 3546,32304 | 640,98472 | 1597,73537 |
| Ccl8 | 58,6202316 | 113,927412 | 16,5190726 | 54,6148627 |
| Chdh | 4,02247899 | 8,60910223 | 0,28883159 | 2,5833381 |
| 1700047M11 | 13,9108135 | 34,4574049 | 4,1767928 | 16,3716973 |
| Vcan | 144,410079 | 428,047248 | 153,653702 | 441,156426 |

| | | | | |
|---|---|---|---|---|
| Cmpk2 | 116,356348 | 158,149036 | 86,348931 | 344,290445 |
| Gm23935 | 191,418799 | 754,256358 | 402,607981 | 1336,21251 |
| Prnd | 144,358531 | 366,577647 | 62,7417608 | 188,723927 |
| Il12rb1 | 17,7391575 | 37,380594 | 11,4498959 | 40,3322266 |
| Oasl1 | 101,852327 | 165,654533 | 66,3161901 | 200,730004 |
| Camp | 616,443338 | 1984,45646 | 314,533292 | 1218,70588 |
| Cldn1 | 22,106039 | 48,2100639 | 4,12842995 | 17,1758024 |
| Mgp | 370,759898 | 479,739424 | 47,5764202 | 204,959649 |
| Arl9 | 14,2661812 | 16,3996212 | 12,846162 | 42,036956 |
| Gjc3 | 2,80421286 | 1,37636344 | 8,91668076 | 39,4089594 |
| Nos2 | 85,513575 | 323,142343 | 66,8091616 | 195,837819 |
| Ltb4r1 | 144,663378 | 333,187533 | 73,166223 | 194,980794 |
| Lefty1 | 26,2079283 | 102,379453 | 23,5522988 | 71,5656229 |
| Per2 | 334,319251 | 911,92495 | 223,749654 | 593,18745 |
| Adora2b | 24,7686031 | 87,7540654 | 22,1171505 | 59,2775313 |
| 1700014D04I | 2,98583998 | 8,5028039 | 4,95947805 | 13,2484053 |
| Pdgfc | 66,8861908 | 225,44773 | 35,1711735 | 96,4444561 |
| Ly6c2 | 71,4215213 | 259,833818 | 45,5195718 | 164,072251 |
| Slc28a2 | 28,1751668 | 76,2266757 | 21,776855 | 66,6625259 |
| Gjc2 | 8,48480009 | 9,73734655 | 7,13002431 | 27,6911172 |
| Hhipl2 | 5,71771037 | 4,90029046 | 6,91736174 | 31,7487299 |
| Gm4759 | 21,1845768 | 20,6652125 | 11,2310751 | 31,9796139 |
| Dhrs9 | 65,9994526 | 189,490618 | 34,0220871 | 108,857476 |
| Slc10a6 | 69,2207348 | 189,969816 | 38,768689 | 119,196614 |
| Gm9899 | 1,7310076 | 4,58837791 | 1,40263786 | 6,17131046 |
| Cyp4f18 | 144,430459 | 329,210308 | 77,3504882 | 175,837407 |
| Mcemp1 | 222,12398 | 512,229028 | 96,6832417 | 287,796324 |
| Tacstd2 | 26,1216977 | 69,3383864 | 9,0900309 | 37,0729707 |
| Rsad2 | 197,080271 | 207,155844 | 138,657479 | 512,266604 |
| Sema6b | 440,642253 | 1723,99155 | 324,648277 | 865,617447 |
| Cdc42ep2 | 137,969567 | 381,113073 | 146,761193 | 428,037913 |
| Cdr2 | 322,548712 | 1031,42569 | 259,001162 | 747,897569 |
| Plin4 | 2,53498942 | 6,43684885 | 1,83136238 | 5,08845803 |
| Trem1 | 403,257281 | 1168,86678 | 201,831127 | 623,948364 |
| Pawr | 35,7177353 | 69,7507028 | 8,22926725 | 28,8385984 |
| Asprv1 | 409,65719 | 863,666431 | 169,889899 | 502,043443 |
| 3930402G23I | 6,1468726 | 17,5686937 | 1,2536992 | 6,48806561 |
| Hp | 774,693641 | 1892,10144 | 353,586985 | 1211,30712 |
| Gja1 | 1721,69873 | 3840,23922 | 875,967611 | 2475,01689 |
| Usp18 | 166,589737 | 207,912908 | 156,746224 | 483,082703 |
| Cnksr1 | 6,12042934 | 7,3377308 | 8,07977276 | 21,8365845 |
| Map2k3os | 31,4619893 | 85,2763788 | 34,550968 | 93,9214159 |
| Klra2 | 58,3169309 | 143,61177 | 27,5487055 | 74,288815 |
| AA467197 | 27,8808763 | 65,9771983 | 14,3292101 | 44,096727 |
| Egfr | 467,47305 | 1283,42264 | 403,178147 | 1064,83717 |
| Cish | 131,043163 | 640,371934 | 144,900959 | 305,204283 |

| | | | | |
|---|---|---|---|---|
| G0s2 | 86,2035519 | 244,41516 | 33,2661073 | 99,8462137 |
| Timp1 | 32,7283112 | 95,8790431 | 12,8518932 | 41,7698474 |
| Ehd1 | 5252,71068 | 15273,0302 | 3730,70671 | 10784,9934 |
| Tcap | 15,1709047 | 24,2241399 | 25,387998 | 62,269059 |
| Gm8430 | 7,67578771 | 12,8158729 | 6,75993316 | 20,750766 |
| Igfbp3 | 409,671546 | 1045,81351 | 258,093069 | 712,512347 |
| Hspa1l | 17,9886911 | 25,5866969 | 25,8662254 | 80,7049528 |
| Met | 55,8858581 | 130,389002 | 51,8245334 | 128,520677 |
| Gm44777 | 5,81598656 | 6,63976515 | 2,92253777 | 11,8172556 |
| Ifit1bl2 | 7,58505978 | 8,11512473 | 5,19498502 | 15,1284637 |
| Tbx21 | 2,83187909 | 7,87811412 | 3,11928346 | 7,50178421 |
| Emp1 | 347,23382 | 1002,44444 | 317,78272 | 799,603322 |
| Atp8b4 | 73,8629642 | 203,654924 | 43,9943678 | 123,185201 |
| Mt2 | 1334,23454 | 7932,85109 | 1657,11598 | 4033,48635 |
| Snord55 | 5,85796898 | 22,5735438 | 7,03462379 | 22,1233892 |
| Adrb1 | 236,431789 | 608,928763 | 126,991779 | 307,402363 |
| Mir17hg | 101,216517 | 195,676963 | 63,7202827 | 145,154941 |
| Isg15 | 254,797983 | 412,016918 | 272,688448 | 814,0845 |
| Far2 | 78,7204807 | 201,138373 | 56,088163 | 144,460125 |
| Gm43802 | 603,065396 | 1383,49541 | 615,768213 | 1328,74489 |
| C1qtnf9 | 7,39186732 | 14,4600452 | 7,92099063 | 24,179766 |
| Shisa2 | 19,8864006 | 45,1441047 | 15,0144206 | 32,9681077 |
| Gm24958 | 26,1780681 | 26,6098225 | 9,61479183 | 20,9589739 |
| AL645522.1 | 27,2154971 | 26,9264154 | 9,61479183 | 20,9589739 |
| Igtp | 612,955118 | 1263,63208 | 677,109821 | 1697,35592 |
| Snord37 | 26,3960186 | 21,7674515 | 10,1257588 | 21,6531746 |
| Zbp1 | 183,77714 | 372,198189 | 136,860017 | 360,71348 |
| 6030468B19F | 16,5483815 | 49,7193452 | 13,5731909 | 38,702091 |
| 1700109H08F | 29,397917 | 32,9194277 | 11,4759701 | 26,930351 |
| Ifit2 | 448,249132 | 562,547052 | 468,504384 | 1414,77346 |
| Syt6 | 9,55431685 | 24,1322189 | 11,1260731 | 28,2373615 |
| Sapcd1 | 65,2582526 | 100,226399 | 46,3571611 | 159,057717 |
| Gbp8 | 115,877409 | 154,303426 | 54,3028728 | 138,280608 |
| Ucp1 | 11,7742406 | 12,475064 | 11,2554819 | 31,8387663 |
| Tgm2 | 979,056758 | 2108,05367 | 585,652686 | 1478,45087 |
| Meox1 | 25,7442749 | 77,1501501 | 16,3813542 | 41,6419935 |
| Ascl2 | 49,7447205 | 254,099784 | 66,9794467 | 164,527372 |
| Gpat3 | 47,0288271 | 151,349445 | 42,6965545 | 93,6063072 |
| Socs1 | 95,4482521 | 248,570357 | 139,212533 | 314,96945 |
| Chil1 | 648,40531 | 1482,29106 | 301,431807 | 873,592243 |
| Plbd1 | 359,858879 | 713,339429 | 190,325562 | 514,750405 |
| C5ar1 | 3755,9652 | 13234,8455 | 3949,97546 | 10141,4018 |
| Rpl7a-ps7 | 9,12744888 | 16,0509511 | 8,06971577 | 22,0627289 |
| S100a6 | 420,041626 | 1119,72651 | 230,669805 | 648,69778 |
| Ldhd | 10,4565797 | 20,3314207 | 9,78338905 | 27,8412407 |
| Batf2 | 136,045264 | 267,68909 | 81,6936989 | 176,650985 |

| | | | | |
|---|---|---|---|---|
| Gm16287 | 14,1529798 | 5,44992179 | 12,0334189 | 39,6882187 |
| Xdh | 468,815033 | 876,671232 | 241,859212 | 588,17486 |
| Gm26797 | 23,8453236 | 23,4091237 | 10,313801 | 29,4339201 |
| Ifi204 | 563,85187 | 921,8189 | 380,506141 | 944,831733 |
| Napsa | 211,310834 | 488,882311 | 115,167059 | 275,274601 |
| Gm29264 | 21,5399878 | 25,6032818 | 16,9614339 | 42,4732765 |
| Efna2 | 29,9726156 | 35,176974 | 75,5934765 | 233,641344 |
| Ifi47 | 283,900248 | 587,518647 | 244,244995 | 592,099719 |
| Prn | 188,039382 | 420,310913 | 100,213877 | 231,853006 |
| Bbs12 | 69,2770904 | 179,56031 | 72,53163 | 140,726594 |
| Snord14e | 94,9369056 | 98,9164863 | 25,9530072 | 69,5698753 |
| Ppef2 | 49,6054174 | 47,9867385 | 25,010984 | 63,3783106 |
| Erich3 | 8,56814618 | 28,2499355 | 8,93655284 | 20,2712917 |
| Trpc6 | 13,8394845 | 25,5271101 | 6,38156094 | 21,0537274 |
| Cd209b | 460,093907 | 892,13502 | 309,174461 | 738,605694 |
| CT009713.7 | 43,7547073 | 50,6553933 | 26,8272789 | 62,5834159 |
| Cfap69 | 9,89084737 | 35,2092351 | 12,9228869 | 23,2517594 |
| Nox4 | 19,1292083 | 26,7628371 | 9,76781446 | 23,747785 |
| Irf7 | 508,035419 | 731,29063 | 492,363435 | 1170,22981 |
| Cldn19 | 8,21865608 | 10,0486196 | 13,806575 | 40,4544278 |
| Kcnn4 | 53,6148757 | 184,951449 | 58,42901 | 135,201001 |
| A530076I17R | 9,88040064 | 17,7444158 | 6,68970878 | 19,1744614 |
| Gm31718 | 64,550962 | 129,320144 | 40,1073408 | 99,0059904 |
| Rasd1 | 68,3768927 | 205,918218 | 61,030738 | 110,269883 |
| Rhov | 31,0705886 | 83,2781109 | 23,9987364 | 61,7048868 |
| Rnd3 | 93,5988503 | 330,799298 | 112,865332 | 209,252465 |
| Afap1l1 | 2118,27837 | 4415,8913 | 1489,71246 | 3503,63486 |
| Vgf | 12,999503 | 20,1110135 | 11,0179008 | 27,9478187 |
| Runx1 | 4575,31194 | 13385,8525 | 4828,60497 | 10648,2407 |
| Thbs1 | 2154,56118 | 4814,95775 | 1767,51096 | 3450,62007 |
| Fabp4 | 12,3808368 | 39,6533336 | 9,64095085 | 22,5018123 |
| 4933406C10F | 5,50847404 | 12,2374992 | 5,90659648 | 12,7210429 |
| S100a10 | 237,780353 | 526,537304 | 144,346406 | 355,036122 |
| Srgn | 2574,00714 | 7545,6005 | 2179,17952 | 4526,16553 |
| Mmp19 | 536,289312 | 1810,6063 | 543,945946 | 1032,62178 |
| Kit | 189,054456 | 306,426684 | 79,9251449 | 171,905204 |
| Il18rap | 206,369824 | 567,686856 | 142,645629 | 323,701275 |
| Sult1a1 | 1046,98403 | 1947,51718 | 809,367829 | 1860,09688 |
| Klf4 | 6256,16741 | 9798,10278 | 5042,32988 | 9466,54222 |
| Mlph | 167,028403 | 352,34281 | 243,513941 | 693,778838 |
| Hs3st3a1 | 9,56980868 | 29,6655244 | 10,1227865 | 18,9930335 |
| Ctla2b | 668,774096 | 1709,00091 | 469,012161 | 1007,24586 |
| Ankrd35 | 5,53804406 | 4,48207959 | 3,53145052 | 15,7562655 |
| Adamtsl4 | 145,394522 | 379,463711 | 77,7897598 | 156,626425 |
| Gm7972 | 12,7753366 | 19,9484311 | 9,45545387 | 22,6785082 |
| Serpinb1a | 149,888575 | 315,072381 | 77,6549985 | 193,562187 |

| | | | | |
|---|---|---|---|---|
| Gm18407 | 21,9508119 | 26,1208772 | 28,2943253 | 54,9551962 |
| Zglp1 | 24,9759357 | 37,2862714 | 43,0275908 | 88,3393477 |
| Txndc2 | 7,93511645 | 7,69898154 | 7,22266604 | 17,1304729 |
| AC160962.1 | 758,675318 | 1864,45523 | 506,931762 | 1095,74555 |
| Cyp1b1 | 85,2331871 | 173,11154 | 61,1540814 | 114,046186 |
| Irgm2 | 475,383163 | 733,753749 | 522,769186 | 1040,82137 |
| Coro6 | 99,7708241 | 101,416701 | 70,4646366 | 174,143809 |
| Cdh1 | 369,334229 | 275,210424 | 29,4375802 | 72,0448426 |
| Hes1 | 390,449105 | 731,146457 | 420,600794 | 917,548507 |
| Gbp3 | 832,069066 | 1771,76929 | 884,959374 | 1679,06909 |
| Madcam1 | 27,4040871 | 81,2175332 | 10,7389974 | 30,3203753 |
| Tmem236 | 26,1254099 | 59,3440871 | 18,6441999 | 37,0519142 |
| Trem3 | 86,9549277 | 176,876886 | 45,0452479 | 99,498182 |
| Nr1h3 | 73,7560929 | 117,895606 | 60,0931774 | 164,468657 |
| Cabyr | 15,8209305 | 28,4248294 | 17,8199946 | 38,765513 |
| Sdcbp2 | 13,408376 | 15,0121765 | 6,7878681 | 24,0018893 |
| Wnt5b | 53,5765624 | 82,0166036 | 30,0931977 | 79,2539093 |
| Ldlr | 2425,9418 | 4429,21996 | 1060,38806 | 2209,49328 |
| Glipr2 | 243,996349 | 541,129009 | 153,707375 | 340,711816 |
| Cidea | 3,89463278 | 5,7103066 | 3,0179383 | 9,70866088 |
| Sult2b1 | 12,6117141 | 21,4007269 | 9,894207 | 24,06535 |
| Anxa1 | 2312,95471 | 3988,31038 | 1186,44923 | 2627,97165 |
| Cdk6 | 1710,09643 | 3968,62199 | 1175,07224 | 2456,89512 |
| C5ar2 | 294,281841 | 836,724666 | 368,733987 | 836,002421 |
| Kcne4 | 38,2925702 | 30,4827667 | 5,78877861 | 22,2161681 |
| Gm16201 | 15,3824924 | 77,5067984 | 17,8198658 | 32,9834824 |
| C030034L19R | 47,1903696 | 91,4549195 | 35,8731928 | 82,6821318 |
| Hormad2 | 30,6816338 | 39,7596412 | 18,7165425 | 40,8193765 |
| Ctrl | 22,3914204 | 25,8223159 | 23,7667576 | 52,5056346 |
| Mall | 14,8879268 | 15,538712 | 7,80372858 | 19,4118111 |
| Tspoap1 | 554,72594 | 453,718704 | 612,167134 | 1539,11077 |
| Nudt10 | 2,44912078 | 8,93581703 | 5,93750375 | 10,1166333 |
| 9130230N09R | 21,643504 | 32,6226464 | 27,9989445 | 67,9786111 |
| Gbp7 | 3221,61348 | 7099,1866 | 3704,33822 | 6505,27141 |
| Slc5a5 | 57,4283085 | 40,71327 | 28,2455354 | 81,0045598 |
| Slc16a12 | 149,090813 | 309,789814 | 109,403011 | 199,06664 |
| Nlrc5 | 303,677038 | 510,908737 | 278,588114 | 540,174273 |
| Tubb6 | 2555,16078 | 7259,28185 | 2820,76844 | 5632,29322 |
| Noct | 225,210743 | 635,364275 | 245,924224 | 466,195349 |
| Wdr38 | 20,3224359 | 30,9744953 | 17,3778421 | 41,0904482 |
| Gm3052 | 18,7124111 | 17,12787 | 17,3651958 | 39,6155596 |
| Rps27rt | 21,2218476 | 44,9959657 | 25,4631888 | 62,2542661 |
| Gm43062 | 14,1955239 | 17,5314233 | 11,1925274 | 22,3256907 |
| Gprc5a | 27,7599644 | 32,6358617 | 10,5575844 | 23,764132 |
| Rnf225 | 83,1154033 | 108,638577 | 83,3089379 | 162,946332 |
| Gm29650 | 15,3486721 | 26,3935414 | 13,8200754 | 24,8887542 |

| | | | | |
|---|---|---|---|---|
| Ankrd33b | 48,3817595 | 98,2197807 | 29,4368642 | 56,9806719 |
| Rhou | 46,349003 | 107,150032 | 37,8771469 | 73,0040956 |
| Oasl2 | 492,392986 | 649,418717 | 466,334919 | 941,907552 |
| Olfm4 | 767,121367 | 1588,90961 | 685,754756 | 1280,80874 |
| Plac8 | 522,419052 | 1021,63072 | 338,999285 | 747,145164 |
| Mx2 | 404,809116 | 331,377434 | 414,208012 | 878,399593 |
| Ctgf | 156,768905 | 218,142888 | 40,4156174 | 78,7513965 |
| Slc41a2 | 12,0538154 | 23,9044075 | 12,7320011 | 28,5407562 |
| Itgb7 | 82,5512712 | 140,648169 | 53,339338 | 125,92211 |
| Rufy4 | 15,9403382 | 28,1747737 | 14,1339544 | 29,8420604 |
| Cd44 | 1293,9648 | 2908,0699 | 858,679918 | 1759,68883 |
| AW046200 | 18,0654599 | 25,4627673 | 13,9318199 | 30,6183992 |
| Mpzl3 | 42,8678693 | 106,75379 | 47,8626548 | 84,6434035 |
| Gm42732 | 26,5017337 | 23,40543 | 16,953371 | 32,7102393 |
| Slc4a11 | 11,92378 | 24,4063707 | 8,88245887 | 16,6340836 |
| Ctla2a | 212,725719 | 428,916193 | 111,480597 | 226,136872 |
| Pou3f1 | 8,08890123 | 11,680922 | 7,76729904 | 16,9287532 |
| Myc | 2236,87466 | 4725,1311 | 1281,90835 | 2311,07397 |
| Tgm1 | 554,940267 | 1198,83337 | 457,675391 | 880,891946 |
| Ctsw | 13,8257349 | 30,2544938 | 13,9968202 | 28,9248717 |
| L3mbtl1 | 3,88361947 | 5,60206245 | 5,94973533 | 14,1112015 |
| Bace2 | 31,504734 | 46,7483363 | 13,3604436 | 37,9369917 |
| Il4ra | 16328,8183 | 42707,2242 | 15932,1713 | 31384,1146 |
| 9130213A22F | 5,36742539 | 6,38989811 | 3,67155696 | 10,599448 |
| Slc1a1 | 36,7300632 | 68,3319515 | 29,2421644 | 57,5361673 |
| Adam33 | 1145,84652 | 1194,09512 | 392,599067 | 717,63705 |
| Igf2bp1 | 82,838485 | 176,459406 | 102,525322 | 191,629736 |
| Uck2 | 446,294277 | 1012,99038 | 340,759207 | 699,828631 |
| Ccr2 | 176,29267 | 312,966061 | 119,594749 | 243,743673 |
| Phyhip | 12,0659376 | 13,4843027 | 7,63194552 | 24,7900399 |
| Thbd | 777,391755 | 1425,08005 | 237,617546 | 419,292895 |
| Gm31597 | 38,931992 | 42,5349123 | 36,4635052 | 76,1810552 |
| Il17ra | 4967,43011 | 10248,3574 | 4112,99877 | 7466,75599 |
| Mb21d2 | 219,797851 | 407,004309 | 271,538734 | 552,315332 |
| Gm37607 | 12,6511789 | 13,5794925 | 12,9382479 | 24,5836269 |
| Chka | 2865,43586 | 4782,33711 | 2859,25781 | 6137,65938 |
| Tnfrsf13c | 12,1109995 | 23,6514542 | 14,8414535 | 34,7038038 |
| Helz2 | 302,454748 | 366,272068 | 199,678245 | 411,112139 |
| Papss2 | 38,5575435 | 78,0935555 | 32,1201657 | 61,1615807 |
| Srgap1 | 116,887218 | 197,283194 | 76,7247714 | 148,802238 |
| Tmod1 | 282,406421 | 577,683836 | 309,909729 | 626,834596 |
| Stx11 | 98,0280456 | 199,553256 | 65,663889 | 130,319442 |
| Acpp | 32,4093336 | 58,2236991 | 20,0793121 | 39,0216477 |
| 1700034H15F | 21,4021278 | 36,6418919 | 18,3397937 | 33,304562 |
| Nkd1 | 330,655758 | 137,798424 | 237,903468 | 129,749886 |
| Kcnh2 | 142,658079 | 69,6677767 | 196,989465 | 94,8081237 |

| | | | | |
|---|---|---|---|---|
| Chaf1b | 839,994181 | 497,301671 | 573,111032 | 318,071914 |
| Lmod1 | 22,5313937 | 11,4478835 | 10,6666548 | 5,68711052 |
| Ube2t | 141,015072 | 54,6752437 | 92,1535857 | 52,5955979 |
| Srl | 311,052023 | 134,602047 | 279,091292 | 134,726621 |
| Fnip2 | 4117,72572 | 2202,45789 | 5792,10411 | 2450,58156 |
| Exo1 | 439,782656 | 253,310783 | 231,535519 | 114,972193 |
| Khdrbs3 | 152,163839 | 82,120306 | 147,659665 | 73,160662 |
| Ercc6l | 578,962312 | 321,297023 | 443,294356 | 219,520844 |
| Tk1 | 1441,18899 | 782,2617 | 633,998223 | 339,744198 |
| Lrr1 | 69,3782748 | 39,1197075 | 30,3577105 | 15,6815996 |
| Crip3 | 8,17108137 | 3,94931822 | 9,42178781 | 6,455577 |
| Klhl38 | 172,480499 | 52,9763399 | 315,618143 | 143,690862 |
| Rad54l | 586,807644 | 315,712308 | 398,888375 | 197,331284 |
| Brca1 | 907,604507 | 469,972449 | 606,10459 | 283,975775 |
| Slc27a6 | 737,126378 | 310,642062 | 749,498348 | 331,19801 |
| Adgrg5 | 1453,22163 | 583,304844 | 1372,17231 | 684,934232 |
| Timeless | 1632,72564 | 820,183395 | 1122,91149 | 537,787166 |
| 2010001K21F | 37,4139092 | 16,878909 | 37,5938406 | 20,7149091 |
| Calhm2 | 302,892504 | 123,917647 | 198,419526 | 94,2886488 |
| Psrc1 | 284,513527 | 161,921427 | 208,706517 | 103,377133 |
| Ccdc159 | 17,9963542 | 5,21938333 | 9,75863995 | 6,13112268 |
| Cdkn2a | 53,4091705 | 31,1578758 | 25,5318745 | 10,3132273 |
| Cyp2r1 | 36,5174766 | 34,033847 | 39,1602994 | 18,7861829 |
| Ccdc15 | 82,6164838 | 35,977671 | 63,6751734 | 30,3439989 |
| Wipf3 | 61,5056674 | 57,0673716 | 141,336386 | 55,7130629 |
| Cbfa2t3 | 2165,39117 | 942,484795 | 1940,6167 | 951,479328 |
| Cfap77 | 5,61580271 | 3,90660615 | 11,991343 | 5,10126138 |
| Dnali1 | 10,5577022 | 7,61867931 | 21,3090269 | 9,32375064 |
| Pask | 536,082228 | 269,253276 | 367,85033 | 168,275348 |
| Lmcd1 | 36,1718888 | 15,5289419 | 32,2029076 | 19,3034758 |
| Gm4316 | 110,553554 | 81,5295422 | 80,0903318 | 35,1982422 |
| Troap | 843,934767 | 490,901578 | 474,827673 | 208,759439 |
| Myb | 121,226253 | 73,1953926 | 71,0630937 | 26,8952975 |
| St8sia4 | 831,810381 | 346,319025 | 721,444904 | 296,776676 |
| Pole | 1543,25751 | 701,4749 | 920,884967 | 414,340028 |
| BC030867 | 293,627753 | 169,120205 | 157,157335 | 78,8000502 |
| Sec16b | 42,5814842 | 24,3548175 | 61,3370251 | 24,6752238 |
| Gm26520 | 60,34293 | 58,6691072 | 99,3639885 | 48,4403087 |
| Nav1 | 2579,50326 | 1352,47814 | 1916,21664 | 791,359357 |
| Rab17 | 100,430031 | 46,1067419 | 126,370536 | 73,8997696 |
| B430305J03R | 41,8880288 | 35,7626962 | 74,9032805 | 26,2874003 |
| Adam5 | 6,38087669 | 3,18061809 | 14,1591305 | 6,73116034 |
| Spaca9 | 33,4251265 | 16,6783846 | 30,1599379 | 16,7197591 |
| Myzap | 48,6640713 | 19,1428929 | 14,7715714 | 6,82059128 |
| Oip5 | 163,225041 | 95,6565204 | 112,542564 | 50,5164095 |
| Kcnc2 | 53,8937419 | 16,4851579 | 28,1971048 | 13,7615502 |

| | | | | |
|---|---|---|---|---|
| Perp | 253,469423 | 171,908498 | 312,169559 | 119,764123 |
| Bspry | 23,076456 | 12,3722615 | 22,1919944 | 6,71066531 |
| Fzd10 | 27,4439514 | 9,23260756 | 44,7765536 | 20,5829201 |
| Clspn | 1237,01319 | 548,017696 | 687,810605 | 291,260099 |
| E2f8 | 1594,0215 | 667,158596 | 825,384749 | 341,303541 |
| Mcm10 | 1008,71692 | 451,607013 | 551,32013 | 264,926804 |
| Dnajb13 | 160,984082 | 52,5910042 | 134,391642 | 77,4918782 |
| Ypel2 | 435,064007 | 130,802935 | 493,701087 | 214,935782 |
| Snai2 | 17,8970067 | 7,66269769 | 8,67056565 | 2,61032138 |
| 4833417C18R | 37,058822 | 17,6575332 | 27,0717312 | 15,1022538 |
| Notum | 79,2147031 | 40,0805211 | 66,0157832 | 31,3220413 |
| Actn2 | 19,863318 | 15,9437788 | 19,4784339 | 7,24136703 |
| Wdr63 | 17,5265516 | 8,61422414 | 23,2752357 | 10,2308837 |
| 4930558J18R | 18,0602059 | 11,4843213 | 13,3941944 | 4,30148714 |
| Eps8l2 | 63,6394275 | 32,2257812 | 36,2159078 | 15,4276846 |
| Rbp4 | 11,7947582 | 5,76314064 | 11,7104942 | 4,08872253 |
| Ppp1r42 | 14,374405 | 5,39796171 | 7,43265456 | 4,15984589 |
| D330045A20I | 56,4351219 | 28,3598335 | 34,1104546 | 13,2188549 |
| Ptprr | 16,9534993 | 7,10556629 | 21,9771288 | 10,5712828 |
| Cutal | 16,5423417 | 7,26032884 | 10,0087103 | 4,22485956 |
| Pcdhb3 | 5,52372787 | 3,69793769 | 13,2840611 | 3,40438598 |
| Lgi2 | 207,073154 | 52,1135748 | 89,0256301 | 37,6182502 |
| Meltf | 4,70425907 | 3,55274291 | 11,189128 | 4,07041495 |
| Fv1 | 29,9808446 | 16,5740321 | 32,5751657 | 10,6861161 |
| Prkcq | 213,301167 | 89,0313391 | 191,884975 | 72,1788588 |
| Gm42793 | 677,322104 | 488,402471 | 950,448136 | 351,369168 |
| Fam69b | 37,9055617 | 18,473711 | 41,5071067 | 17,760061 |
| Rtkn2 | 201,328987 | 62,845468 | 148,796248 | 61,2765635 |
| Ddit4l | 25,5012134 | 11,3722708 | 18,5051564 | 7,66095443 |
| Stac2 | 53,0014441 | 23,9121148 | 89,736183 | 35,5099007 |
| Syt15 | 211,226346 | 71,0121294 | 161,329543 | 68,3626901 |
| Gm7666 | 98,9434785 | 61,1950667 | 87,8342951 | 36,1575566 |
| Gm12286 | 8,43974308 | 7,83659577 | 12,6805371 | 4,80341817 |
| AC027700.1 | 14,569401 | 6,88562352 | 22,2912166 | 8,07916306 |
| Syngr4 | 15,9788082 | 10,9992721 | 8,89469046 | 2,42849235 |
| Dnah5 | 12,8356426 | 4,1910505 | 15,3026523 | 5,24269832 |
| Gna14 | 7,28257782 | 2,8590252 | 6,57201969 | 2,80948146 |
| Lrrc10b | 8,52561172 | 4,12114862 | 8,0860597 | 2,70428001 |
| F2rl1 | 20,7419742 | 9,24935565 | 16,1151535 | 5,01990069 |
| Klhdc9 | 4,9937766 | 2,73978512 | 7,3941068 | 2,24963048 |
| Fgfr4 | 46,458012 | 8,82245097 | 15,0455414 | 6,3763759 |
| Lhx6 | 61,4872534 | 39,583441 | 40,5035833 | 14,6326435 |
| Frmpd2 | 8,12177414 | 2,95595366 | 8,16830214 | 2,6197779 |
| Kcnj14 | 55,2813314 | 22,0377696 | 50,3790345 | 15,5208699 |
| Col6a6 | 32,2772708 | 10,963308 | 33,3158237 | 6,59425109 |
| Ttll6 | 6,94950266 | 0,89456632 | 6,40363599 | 3,39729226 |

| | | | | |
|---|---|---|---|---|
| Baiap2l2 | 537,491123 | 229,900974 | 745,772132 | 262,210689 |
| Scn4b | 9,76686576 | 2,53643648 | 8,82425256 | 3,54346121 |
| Tnfrsf26 | 137,814076 | 80,9694362 | 183,489279 | 41,1315449 |
| 2610035F20R | 5,47671489 | 3,44787275 | 4,85240177 | 0,31104661 |
| Fam26e | 13,0699368 | 5,50426003 | 6,31099427 | 1,25975264 |
| Igfbpl1 | 32,5851541 | 26,9365373 | 58,8363807 | 16,2446231 |
| 6030442K20R | 4,77794845 | 3,64641923 | 10,8462872 | 1,73588123 |
| AF529169 | 161,302577 | 41,1915805 | 234,466894 | 66,4747485 |
| Cerkl | 15,5171017 | 9,34227987 | 19,0371355 | 3,25132764 |
| Gm37689 | 40,7841405 | 15,1303887 | 19,5649018 | 4,47345852 |
| Gm20083 | 2,1596081 | 0,9472785 | 5,75298964 | 1,59089314 |
| Gm38399 | 147,487467 | 74,6445631 | 128,427097 | 24,4479007 |
| Gm10699 | 3,10496714 | 2,4290663 | 5,08026824 | 0,26633114 |
| Ephx4 | 3,31364659 | 0,53113531 | 3,90960921 | 1,59542082 |
| Gm42879 | 8,88494918 | 4,69856789 | 6,97591035 | 1,80187144 |
| Grp | 12,3591436 | 6,49337664 | 7,50309246 | 1,86175197 |
| Hs3st2 | 62,537848 | 17,3306254 | 70,2249685 | 17,7360448 |
| Crabp1 | 7,66220452 | 2,05813521 | 7,24452756 | 1,93149015 |
| EPPK1 | 5,03481168 | 0,94977853 | 2,63094739 | 0,88271582 |
| Gm4739 | 20,3035409 | 10,0612416 | 14,0725622 | 2,5088679 |
| Duoxa1 | 3,96580446 | 1,06564461 | 3,72862325 | 0,57737775 |
| Blk | 31,9164192 | 7,77768982 | 36,1799133 | 8,70991051 |
| Cd28 | 114,928327 | 28,9073606 | 139,665013 | 29,3738202 |
| Gm9816 | 30,8573883 | 6,01439931 | 28,3929794 | 5,36306483 |
| Wnt9b | 4,6988429 | 0,26388066 | 2,41212665 | 0,93313983 |
| Gm31409 | 3,44401051 | 0,94477847 | 3,54206333 | 0,31104661 |
| Vill | 322,840007 | 79,601341 | 292,22003 | 32,0481412 |
| B3gnt4 | 4,03427748 | 0,63068566 | 2,05604301 | 0,3311405 |
| 2610507I01Ri | 30,9258091 | 2,27680377 | 27,9560585 | 2,1621612 |
| Hs3st6 | 7,55352894 | 0,5345093 | 1,42484169 | 0 |
| Kcnv1 | 2,33307305 | 0,26388066 | 2,08121915 | 0 |
| Spata9 | 2,19790203 | 0 | 1,06599925 | 0 |
| Gm15601 | 0,2402103 | 0,85110214 | 0 | 1,46009357 |
| Gm16325 | 0,48042059 | 4,27785699 | 0 | 0,3311405 |
| Prss50 | 0,2402103 | 1,32615129 | 0,67603602 | 3,86298031 |
| Cxcl3 | 8,97394458 | 64,1073125 | 6,24586512 | 24,8739903 |
| Ccl22 | 8,76754969 | 57,8559651 | 24,4234617 | 36,6713563 |
| Gm37194 | 1,74453376 | 9,13848961 | 6,2548261 | 11,2233287 |
| Dcpp2 | 45,4164427 | 191,970443 | 71,7193733 | 101,130766 |
| Tnfsf14 | 27,5429034 | 131,601897 | 28,8392002 | 66,8565733 |
| Mmp13 | 14,5508395 | 65,2556083 | 22,4189707 | 34,9018164 |
| 1700023F06R | 1,26159547 | 2,90661547 | 1,79260109 | 6,24814236 |
| Selp | 26,1893947 | 124,284327 | 13,4151658 | 41,4203864 |
| Gpr15 | 6,93117424 | 22,3146953 | 3,28138021 | 8,62266704 |
| 2210008F06R | 2,28298064 | 6,19222558 | 3,57968459 | 9,85148409 |
| 3110082J24R | 0,78793796 | 2,37548016 | 2,24705758 | 5,8730661 |

| | | | | |
|---|---|---|---|---|
| Stfa2 | 1072,93971 | 2792,30788 | 479,29907 | 1415,59177 |
| Rep15 | 1,49756038 | 7,2159907 | 1,66883857 | 3,94886758 |
| B230303A05F | 29,5384131 | 81,8288142 | 15,1456948 | 56,2019444 |
| BC048644 | 2,02052028 | 10,0582999 | 2,08121915 | 4,11039841 |
| Kcna7 | 6,4698721 | 25,1961887 | 11,4076064 | 21,904545 |
| Ch25h | 108,626712 | 672,992707 | 246,37358 | 377,593256 |
| Gm21887 | 14,1417479 | 40,2372523 | 7,85607032 | 27,6911334 |
| Ngp | 3050,23015 | 8100,67366 | 1515,9129 | 4886,60986 |
| Lcn2 | 1650,05846 | 4983,01356 | 861,310428 | 2609,88878 |
| Gm11707 | 11,0825226 | 38,2451322 | 6,08721176 | 15,1952272 |
| Vaultrc5 | 5,39107455 | 10,3535862 | 9,25905049 | 23,1357226 |
| P2ry2 | 289,579395 | 1203,39329 | 307,141606 | 469,854638 |
| Ppbp | 25,067178 | 92,2011285 | 40,9962168 | 77,2528207 |
| Bcl2a1a | 115,05107 | 370,870999 | 124,969448 | 236,512362 |
| Rab33a | 6,41185066 | 18,7092403 | 10,1326015 | 20,0037786 |
| Klrb1c | 23,0138606 | 59,9516304 | 20,0406356 | 50,5832343 |
| Msr1 | 2591,16591 | 6435,34263 | 2040,3187 | 4199,1407 |
| Rpl28-ps1 | 36,4584842 | 92,390779 | 51,6965894 | 112,040025 |
| Gm5879 | 3,85578197 | 12,2796157 | 7,70492869 | 17,3203722 |
| Cnga2 | 4,33251406 | 14,8997698 | 9,91046619 | 17,04519 |
| Ltf | 2814,52776 | 7356,5105 | 1638,1959 | 4572,87408 |
| Ffar2 | 12,9658401 | 35,4252564 | 10,083091 | 26,3150127 |
| Rtn4rl2 | 2,34777001 | 5,87269109 | 0,33939741 | 2,14206731 |
| Hc | 1,4882796 | 6,11362989 | 2,33914348 | 3,50445428 |
| AC238811.2 | 10,0995317 | 30,0725415 | 8,38393703 | 15,5445086 |
| Dmkn | 3,79775568 | 13,0936818 | 8,32819592 | 13,4477622 |
| Lor | 2,76960743 | 7,13100818 | 2,58589916 | 4,80656068 |
| Gm10073 | 74,5920205 | 194,569289 | 98,6131573 | 212,135331 |
| Il1rn | 239,779829 | 722,219488 | 344,747979 | 527,273557 |
| Palm3 | 2,95179143 | 7,49999344 | 3,21171166 | 7,39580414 |
| Serpina3n | 26,5399561 | 95,6861844 | 78,2358548 | 123,719905 |
| Ackr1 | 146,032614 | 433,987475 | 111,44797 | 255,297455 |
| Hrct1 | 15,7969528 | 36,7184151 | 15,429943 | 33,0797802 |
| Fam83f | 2,56148486 | 6,18353177 | 4,85813288 | 9,54655577 |
| Tulp1 | 3,37167288 | 7,43715929 | 1,81225965 | 5,20251054 |
| Dmp1 | 4,80365387 | 11,6616739 | 1,73906295 | 5,23935148 |
| Rpl10-ps3 | 22,465824 | 55,6546614 | 39,3003728 | 60,6202609 |
| Gm14963 | 2,79917745 | 5,51263407 | 2,05328421 | 6,57199122 |
| Pdgfa | 343,095712 | 931,692657 | 366,148715 | 632,213621 |
| Gm32899 | 6,86615657 | 15,1684526 | 9,32893256 | 12,0144551 |
| Gm11808 | 7,66533613 | 22,0490346 | 13,9012062 | 22,0507064 |
| Bcl2l1 | 4372,89148 | 11937,3807 | 4088,40134 | 7574,66677 |
| Gm12696 | 5,98464925 | 12,3500717 | 6,8610648 | 15,553964 |
| Gm26902 | 12,3069428 | 40,2176132 | 16,9551186 | 21,0038937 |
| Gm5781 | 44,9050001 | 100,87307 | 51,1675873 | 96,9135395 |
| Ms4a6c | 2966,91649 | 6423,18915 | 2365,04465 | 4909,11758 |

| | | | | |
|---|---|---|---|---|
| Gm45855 | 11,0648465 | 27,4701015 | 18,0943429 | 21,6738664 |
| C3 | 1066,03247 | 1953,53608 | 607,130355 | 1419,99521 |
| Fank1 | 1,67666981 | 3,79618173 | 3,57615644 | 7,45706984 |
| Gm8210 | 8,70780058 | 20,0687337 | 11,4836907 | 23,263185 |
| Gm43703 | 17,358736 | 38,2340236 | 22,8835766 | 42,8075746 |
| Plaur | 1704,05932 | 4705,48944 | 2534,59661 | 3549,54579 |
| B3gnt7 | 527,847991 | 1452,51684 | 546,470303 | 776,516743 |
| Olfm1 | 174,530135 | 478,853717 | 143,594479 | 222,021735 |
| AC127578.1 | 40,3282538 | 80,9177782 | 54,261376 | 90,7388811 |
| Ccnd1 | 5646,67402 | 12955,1716 | 4579,50966 | 9110,99495 |
| Pparg | 66,7270706 | 159,172016 | 78,4040093 | 148,344189 |
| AL845457.1 | 26,7979573 | 66,5918581 | 30,4319985 | 44,7981783 |
| H2-Ab1 | 464,348874 | 1150,92097 | 533,930416 | 875,708755 |
| Fam241a | 67,4218248 | 158,011092 | 58,7430554 | 114,775304 |
| Entpd3 | 25,733904 | 68,2041764 | 13,5190122 | 16,2221589 |
| Dnajb3 | 8,00069585 | 17,1559338 | 5,94676301 | 11,7465333 |
| Cxcr2 | 732,281479 | 1637,61567 | 673,865484 | 1369,40809 |
| Klc3 | 6,07028474 | 12,3490759 | 6,95646532 | 13,282278 |
| Smim3 | 522,099027 | 1300,75235 | 569,763647 | 717,616265 |
| Ccdc73 | 19,1345152 | 31,7550701 | 19,8483115 | 57,492822 |
| Cd177 | 745,780102 | 1442,96991 | 408,609815 | 896,81521 |
| Piwil2 | 41,5406227 | 106,082099 | 37,4107524 | 57,8389608 |
| H2-M2 | 10,7258032 | 24,4561919 | 16,1897838 | 21,3813381 |
| Hcar2 | 621,850821 | 1423,67462 | 612,922904 | 836,9098 |
| Fgr | 583,904663 | 1149,07697 | 386,093525 | 762,957404 |
| Gm10051 | 11,9891263 | 21,2573927 | 13,4244658 | 32,6487757 |
| Gm15501 | 47,1870904 | 99,5893201 | 63,2507951 | 109,262471 |
| Hivep2 | 1209,23937 | 3085,34538 | 997,906878 | 1707,81896 |
| Gm4754 | 17,7803548 | 38,1312924 | 18,5485859 | 37,0385344 |
| Slc35e4 | 227,503266 | 510,179516 | 186,14588 | 307,864014 |
| Nfil3 | 1359,16771 | 3360,64302 | 1009,8742 | 1585,49854 |
| Hmcn2 | 5,92158755 | 11,0434883 | 2,70668936 | 6,86530674 |
| Ass1 | 43,0925641 | 82,156309 | 27,2608567 | 53,6308246 |
| H2-Aa | 533,852686 | 1330,22901 | 591,642126 | 977,550822 |
| Gpt2 | 672,043181 | 1544,27324 | 774,616989 | 1301,58257 |
| Fhl3 | 1065,2092 | 2277,71821 | 1030,33652 | 1960,05676 |
| Mras | 1070,83663 | 2386,83785 | 1718,24418 | 3159,24794 |
| Rpph1 | 77,5715916 | 133,14825 | 117,631427 | 217,956175 |
| Rpl9-ps6 | 74,5568721 | 140,52098 | 94,7577539 | 154,770827 |
| Rbm47 | 1243,2801 | 2791,91386 | 1112,01867 | 1840,07397 |
| H2-Eb1 | 431,520679 | 942,985591 | 447,150684 | 771,381084 |
| Mt1 | 6577,96165 | 13263,8709 | 7222,49847 | 9856,77931 |
| Plin5 | 5,25198673 | 8,46358768 | 6,28263227 | 10,0894468 |
| Stk40 | 1951,05191 | 4366,53488 | 2163,07571 | 3620,41024 |
| CT010498.2 | 331,190768 | 707,34522 | 231,048412 | 402,991264 |
| Gm2000 | 51,5343968 | 85,8402584 | 44,8314453 | 98,3564956 |

| | | | | |
|---|---|---|---|---|
| Fgf2 | 10,9437634 | 33,4654413 | 17,440853 | 18,705019 |
| Gm10288 | 23,8560557 | 44,4012788 | 29,1548314 | 41,8137583 |
| Cdkn2b | 19,568156 | 40,7974199 | 18,5799202 | 28,7544748 |
| Spint2 | 61,1799554 | 112,452245 | 18,554317 | 30,7734439 |
| Trmt61a | 429,978953 | 870,344457 | 480,840649 | 737,335503 |
| Gm11539 | 42,9084053 | 77,8161881 | 46,827445 | 93,2158358 |
| Pglyrp1 | 524,27519 | 1020,96941 | 321,488378 | 680,875183 |
| Glipr1 | 492,951002 | 1031,58095 | 759,72066 | 1185,98826 |
| Tma16 | 259,783179 | 559,885589 | 246,836327 | 430,219483 |
| Fbp1 | 109,673246 | 238,408009 | 101,488359 | 174,848836 |
| Lipc | 14,0808704 | 29,762584 | 12,3760022 | 19,8075932 |
| Ptpn7 | 1111,97366 | 2133,34392 | 1064,8612 | 1461,53668 |
| Gm15536 | 39,1776184 | 69,9610524 | 51,1692629 | 96,9746502 |
| Rps18-ps3 | 25,869256 | 49,4656626 | 29,5865722 | 42,5899251 |
| Rpl36a-ps2 | 22,5098815 | 40,1201652 | 30,1936887 | 50,694853 |
| Rpl39-ps | 57,5056912 | 95,8032264 | 62,2471222 | 110,571907 |
| Rrs1 | 937,945767 | 2012,96498 | 871,327452 | 1557,8756 |
| Emilin2 | 2083,30707 | 4158,80871 | 1412,46808 | 2597,32802 |
| Lgals3 | 844,244652 | 1699,66691 | 988,471294 | 1558,16653 |
| Bex4 | 11,2338329 | 21,3524167 | 10,6504396 | 17,5811991 |
| 2510016D11F | 255,004622 | 114,756566 | 268,612568 | 125,564429 |
| 2610028D06F | 15,8131491 | 3,33232642 | 2,99297567 | 0 |
| 4930518I15Ri | 9,09922576 | 2,85390329 | 7,55654585 | 3,58956185 |
| 4933433G15F | 12,467767 | 1,65036607 | 11,0848953 | 12,7754192 |
| 6430511E19R | 10,7162941 | 3,26929437 | 9,99371987 | 8,69004352 |
| 9430034N14F | 55,6845612 | 19,6320269 | 23,6080038 | 18,209986 |
| 9930120I10Ri | 74,9379835 | 29,614682 | 39,1728577 | 28,417459 |
| AC108802.1 | 221,484204 | 82,0456248 | 81,2666656 | 63,6359822 |
| AC112142.1 | 2,98756766 | 1,06227062 | 4,85558761 | 1,91139626 |
| AC124426.1 | 109,95196 | 44,2866161 | 104,976996 | 41,4071474 |
| AC124556.1 | 48,4721075 | 25,5019835 | 16,5395805 | 20,1710265 |
| AC129597.1 | 58,3991452 | 22,9730471 | 30,9919249 | 18,7369322 |
| AC131586.4 | 27,8390275 | 10,3483425 | 9,19413486 | 5,86124788 |
| AC139845.1 | 5,56028928 | 1,78663261 | 2,48808214 | 1,56016188 |
| AC151971.2 | 26,57183 | 13,5303887 | 15,2005094 | 12,6381389 |
| AC154640.4 | 67,0797598 | 30,8413727 | 34,2584795 | 23,6871022 |
| AC157931.1 | 16,5167936 | 5,08061676 | 10,8211111 | 9,49454116 |
| Acss3 | 15,0470756 | 3,11953187 | 11,5639721 | 8,78676499 |
| Adamts16 | 777,840437 | 376,506928 | 864,41033 | 435,162153 |
| Adgra2 | 1244,66417 | 667,889023 | 1042,16596 | 562,991314 |
| Adh1 | 77,3239947 | 31,32035 | 38,5723734 | 28,2421107 |
| Adra1a | 10,7708553 | 4,60207178 | 5,21145773 | 3,91931718 |
| Afap1l2 | 230,455182 | 90,7464019 | 138,170778 | 109,608968 |
| Ago4 | 1048,64587 | 546,784857 | 1185,73772 | 691,376467 |
| Agtr1a | 4,65747439 | 2,05388726 | 1,90710434 | 0 |
| Ahnak2 | 166,408943 | 55,3521096 | 199,575075 | 141,785736 |

| | | | | |
|---|---|---|---|---|
| AL645745.1 | 10,9197856 | 5,57078793 | 8,26051682 | 3,69908667 |
| Alas2 | 107,876803 | 38,1274514 | 47,0056845 | 37,929898 |
| Aldoa | 48,4683041 | 21,4702286 | 25,924017 | 14,7122157 |
| Als2cl | 827,732166 | 421,733847 | 575,030766 | 350,5543 |
| Aph1c | 401,108831 | 190,188876 | 342,604104 | 186,146439 |
| Arntl | 528,602677 | 227,443243 | 505,513154 | 259,619516 |
| Arpp21 | 57,9666225 | 22,3820651 | 43,1623809 | 32,8452041 |
| Asic3 | 19,0926469 | 6,50282255 | 10,1519178 | 7,48091061 |
| Asxl3 | 14,3512125 | 5,49751205 | 9,63466391 | 7,47736805 |
| Atad2 | 2369,48444 | 1187,14254 | 1670,3569 | 871,362105 |
| Atad5 | 911,556571 | 468,823005 | 629,329732 | 360,252803 |
| Atp8b5 | 14,5756121 | 7,59855724 | 22,4682237 | 10,7483809 |
| Avil | 29,0998225 | 12,839369 | 41,627583 | 36,8557009 |
| B230323A14F | 1,06644218 | 0,31409281 | 3,63003688 | 0,82736298 |
| B3gnt8 | 147,844049 | 57,8758757 | 107,62126 | 71,1052586 |
| C130021I20R | 7,02357766 | 3,22320831 | 5,92137334 | 3,06497091 |
| C130071C03F | 28,0676579 | 15,0644564 | 30,8535583 | 14,9878302 |
| C1ql3 | 3,24751032 | 1,6402441 | 17,1298176 | 5,16410907 |
| Cacna1e | 130,875076 | 59,0279978 | 72,5396226 | 35,1166858 |
| Capns2 | 99,7050832 | 43,4158652 | 46,1442286 | 33,754116 |
| Catsperd | 6,40452585 | 2,37635412 | 4,80799411 | 2,92136799 |
| Cbln4 | 0,70710472 | 0,31659284 | 3,62672226 | 0,9934215 |
| Cbx2 | 153,082971 | 64,7420994 | 94,9873209 | 56,5945174 |
| Cdc7 | 466,309963 | 250,905622 | 236,818256 | 108,728238 |
| Cdhr1 | 29,4136796 | 15,0058702 | 15,1913789 | 13,1843756 |
| Cenpu | 221,300006 | 100,7248 | 131,816246 | 77,4150505 |
| Chrd | 36,817317 | 18,5816261 | 24,4831613 | 13,9547963 |
| Cited4 | 18,3269638 | 5,38883556 | 10,1546766 | 9,03279792 |
| Cmah | 634,197239 | 285,943219 | 389,384251 | 240,718441 |
| Cmya5 | 96,7567422 | 24,6417298 | 93,1928181 | 51,4497333 |
| Col25a1 | 67,2846742 | 31,1222315 | 47,7067603 | 29,554234 |
| Cpeb1 | 24,3496071 | 11,8643464 | 22,8196392 | 9,37890666 |
| Crybg3 | 1279,22775 | 703,340828 | 1249,11912 | 720,397824 |
| CT009718.2 | 56,9826603 | 23,9587229 | 39,8391506 | 20,9485409 |
| CT025660.2 | 61,864986 | 17,4173905 | 32,7368917 | 17,0802521 |
| Ctnna3 | 24,1378106 | 11,2373545 | 41,0857164 | 21,7603666 |
| D030028A08I | 782,770288 | 416,409948 | 894,428464 | 440,516177 |
| D030055H07I | 4,09230377 | 1,16519496 | 1,42505521 | 0,5421188 |
| Ddc | 13,6739631 | 9,11761544 | 15,9380664 | 6,61336095 |
| Ddx60 | 1000,50124 | 527,937628 | 804,168591 | 614,696808 |
| E030022I16Ri | 96,9879815 | 45,9960671 | 69,7553899 | 63,0589669 |
| Egr3 | 1060,9474 | 626,233097 | 2562,85252 | 1422,74037 |
| Enpp3 | 13,5632745 | 7,68659399 | 9,56711358 | 6,04662057 |
| Esco2 | 387,629598 | 184,25728 | 243,115223 | 134,745074 |
| Extl1 | 7,50651595 | 3,38753863 | 2,87473075 | 0,26633114 |
| Eya1 | 41,3216589 | 18,4654496 | 12,9701381 | 9,71754201 |

| | | | | |
|---|---|---|---|---|
| Fam167a | 468,827752 | 115,965616 | 515,737088 | 270,368493 |
| Fam214a | 990,821447 | 400,766626 | 965,340183 | 510,805362 |
| Fam228a | 2,30947608 | 0,53113531 | 2,46566479 | 2,32960489 |
| Fbxl21 | 31,5708606 | 15,2254127 | 16,1435155 | 12,868597 |
| Fendrr | 40,1130828 | 7,59898958 | 10,246549 | 4,1056739 |
| Fgf10 | 1,42773561 | 0 | 2,03086686 | 0,27578766 |
| Fosb | 33185,7303 | 15990,9033 | 24076,0197 | 16343,4801 |
| Foxb1 | 0,79045566 | 0,26725465 | 15,5140901 | 14,4271694 |
| Foxd2os | 18,94563 | 5,35024959 | 1,74203527 | 3,40674878 |
| Gdpd2 | 154,194351 | 54,4475962 | 71,5969948 | 56,643581 |
| Gm10425 | 11,8119777 | 3,75184359 | 23,3906731 | 22,9293021 |
| Gm11398 | 14,2596994 | 7,21631046 | 8,19382059 | 4,56150428 |
| Gm1141 | 7,30482788 | 1,15594695 | 2,85286922 | 0,92861214 |
| Gm11870 | 17,5443377 | 6,80334336 | 4,83592906 | 1,89268755 |
| Gm12454 | 34,8437053 | 15,0949788 | 17,6287665 | 27,0300193 |
| Gm15816 | 90,8753429 | 40,9232175 | 76,166649 | 51,3802198 |
| Gm15880 | 103,472893 | 50,0007335 | 126,018272 | 58,5341443 |
| Gm16892 | 4,41586015 | 0,84522813 | 6,01995971 | 2,99563387 |
| Gm17024 | 359,538328 | 192,152062 | 298,584683 | 238,656168 |
| Gm17971 | 77,8041827 | 48,4925005 | 51,7306152 | 54,3417755 |
| Gm17999 | 34,0222283 | 13,2150657 | 9,90417924 | 6,64882422 |
| Gm20274 | 25,164431 | 10,6612416 | 18,8244729 | 12,8804238 |
| Gm21781 | 83,884243 | 48,9209806 | 62,5430745 | 46,5768231 |
| Gm22589 | 14,1461645 | 5,17385147 | 12,1168862 | 6,10099688 |
| Gm22748 | 16,7101768 | 5,48239003 | 4,1511896 | 3,66953627 |
| Gm25749 | 2,05881421 | 0,52776132 | 2,31375381 | 0,62209322 |
| Gm25835 | 177,016015 | 96,0441743 | 59,640244 | 45,371606 |
| Gm25994 | 643,184946 | 291,645302 | 213,692847 | 192,772447 |
| Gm26514 | 23,9077233 | 10,3184961 | 10,4755555 | 10,398736 |
| Gm27694 | 231,979265 | 125,964853 | 97,9491631 | 84,056837 |
| Gm2788 | 23,6464241 | 9,08796698 | 18,6246701 | 13,0594814 |
| Gm29260 | 3,07768168 | 0,84860212 | 1,35483084 | 0 |
| Gm35154 | 53,8932893 | 25,4518172 | 51,2151652 | 26,3817451 |
| Gm36401 | 16,2841838 | 3,90867383 | 9,21909748 | 6,63916338 |
| Gm36527 | 58,1303778 | 25,5381913 | 42,8850522 | 25,5841131 |
| Gm36529 | 9,53034396 | 3,64479317 | 9,76152752 | 5,06343531 |
| Gm37019 | 30,0441207 | 17,288624 | 9,36514858 | 13,0311119 |
| Gm37168 | 19,5644487 | 6,37957824 | 35,8332067 | 23,2901769 |
| Gm37274 | 26,9798649 | 10,5127946 | 13,2936186 | 13,6478762 |
| Gm37303 | 15,9077417 | 4,85257834 | 8,91413549 | 3,31238902 |
| Gm37309 | 37,494519 | 19,6029326 | 16,2194994 | 14,1058941 |
| Gm37310 | 68,6110153 | 30,5330456 | 51,9120798 | 11,3850413 |
| Gm37352 | 436,714899 | 202,000365 | 165,503498 | 129,547763 |
| Gm37893 | 62,2894651 | 28,3818508 | 23,7469703 | 28,7062522 |
| Gm38190 | 68,4882344 | 40,7152919 | 23,2164735 | 22,1048848 |
| Gm38220 | 49,4575884 | 20,5280306 | 7,87671175 | 7,29671987 |

| | | | | |
|---|---|---|---|---|
| Gm38372 | 91,1919744 | 39,9082174 | 60,7964481 | 17,7319032 |
| Gm38375 | 18,39963 | 7,7258516 | 10,3579511 | 5,25313887 |
| Gm42047 | 1031,51603 | 389,102679 | 480,645872 | 281,002143 |
| Gm42664 | 88,9645128 | 33,6697444 | 26,2421371 | 12,856582 |
| Gm42748 | 83,5400265 | 43,9870102 | 31,6719885 | 21,5801185 |
| Gm42979 | 78,4916357 | 38,8463351 | 32,6841637 | 32,221316 |
| Gm43039 | 18,9233278 | 7,39969099 | 5,99159771 | 3,85450782 |
| Gm43080 | 33,1652034 | 12,9919512 | 9,62948862 | 5,7412825 |
| Gm43088 | 27,972342 | 10,1732599 | 11,2306481 | 7,44051961 |
| Gm43197 | 8,61611135 | 3,69413136 | 6,45420181 | 3,29604311 |
| Gm43272 | 79,4061218 | 31,6578931 | 39,8672427 | 23,8677515 |
| Gm43351 | 34,1419646 | 12,4893213 | 30,3839056 | 20,3112826 |
| Gm43362 | 37,1920419 | 14,9082657 | 26,1050029 | 19,6088138 |
| Gm43389 | 27,6242413 | 10,5110467 | 9,15847467 | 7,03038873 |
| Gm43437 | 8,87645842 | 2,90248937 | 9,5557361 | 6,40947636 |
| Gm43481 | 68,8471438 | 33,621478 | 42,4423642 | 22,8000641 |
| Gm43679 | 28,3641668 | 11,5679975 | 20,066154 | 12,3654926 |
| Gm43693 | 12,2534953 | 4,44143521 | 4,728724 | 3,48179437 |
| Gm43759 | 99,4348129 | 53,4573853 | 46,0228542 | 56,0648244 |
| Gm43858 | 65,1670729 | 30,014275 | 27,8860571 | 15,2705072 |
| Gm43912 | 32,3018477 | 16,078019 | 16,0639031 | 16,7987559 |
| Gm43951 | 16,3626851 | 7,66889147 | 11,0346718 | 6,7804035 |
| Gm44033 | 13,6683138 | 4,27123087 | 9,89534703 | 5,31834937 |
| Gm44053 | 14,4021423 | 6,60574688 | 14,4113237 | 11,1254253 |
| Gm44067 | 55,7118077 | 27,1882816 | 46,6097766 | 53,1446652 |
| Gm44164 | 66,441685 | 33,800889 | 41,255402 | 26,4497022 |
| Gm44423 | 366,304651 | 131,145911 | 176,54424 | 106,511761 |
| Gm44574 | 72,0935543 | 35,7102668 | 76,2813501 | 42,983695 |
| Gm45309 | 17,3293281 | 7,26707681 | 14,8205184 | 5,60555411 |
| Gm45540 | 72,9250985 | 10,5171645 | 25,7279072 | 11,5989878 |
| Gm45743 | 30,2092661 | 13,6009209 | 17,9170423 | 11,7796349 |
| Gm45838 | 38,9798431 | 16,5139324 | 10,3165598 | 6,58873937 |
| Gm45906 | 3,72699326 | 0,31659284 | 1,64069011 | 1,60034965 |
| Gm5857 | 11,515088 | 3,44199873 | 13,7187302 | 7,0682148 |
| Gm5977 | 3,98843044 | 0 | 4,20846941 | 2,10228068 |
| Gm8817 | 43,3229367 | 23,299217 | 23,8606758 | 26,7434189 |
| Gm8835 | 14,2189399 | 4,42261944 | 8,68571318 | 6,1157608 |
| Gpr157 | 1417,02686 | 740,060853 | 1759,74188 | 861,748776 |
| Gpr160 | 1643,62898 | 758,716898 | 1490,80508 | 827,240904 |
| Gpr182 | 63,5702027 | 30,3469764 | 42,7596866 | 24,1782066 |
| Gpr63 | 6,35863148 | 1,15932094 | 5,93508725 | 1,46955009 |
| Gramd1b | 3987,67021 | 1939,30248 | 5190,80852 | 3061,93236 |
| Grb14 | 56,4653685 | 21,7569337 | 30,706105 | 22,2268269 |
| Hectd2os | 7,59998504 | 2,21484359 | 18,8285569 | 9,50065835 |
| Hepacam2 | 5,60786884 | 1,96271096 | 5,20071614 | 4,48271072 |
| Hey2 | 55,5098865 | 23,6152508 | 32,1848678 | 29,5300339 |

| | | | | |
|---|---|---|---|---|
| Hist1h2ac | 65,9362927 | 21,2830004 | 22,1808257 | 16,9102413 |
| Hist1h2bg | 142,01267 | 69,6583426 | 52,1938268 | 44,7952101 |
| Hist1h2bh | 34,0863035 | 14,0339019 | 14,167198 | 16,2378994 |
| Hkdc1 | 7,84634935 | 1,52862596 | 4,13747569 | 3,81806802 |
| Hlf | 169,101391 | 75,3211228 | 73,2861104 | 61,407961 |
| Igdcc3 | 22,6505208 | 9,23391387 | 11,977122 | 9,78903754 |
| Igf1os | 16,0864703 | 4,05918843 | 16,2157293 | 11,6986765 |
| Ighj4 | 49,5765582 | 18,3462188 | 43,9094183 | 21,1171729 |
| Il7r | 144,522313 | 54,2822495 | 165,658601 | 81,1476065 |
| Insc | 18,0261477 | 2,37722808 | 9,58078347 | 9,41770926 |
| Insrr | 7,68987075 | 1,84771883 | 0,72362952 | 1,33165571 |
| Ipcef1 | 508,539564 | 245,931388 | 436,090904 | 239,603387 |
| Irf4 | 683,452648 | 257,902362 | 571,205706 | 302,600571 |
| Kcnh1 | 5,82386825 | 1,43244961 | 1,39966554 | 0,60692816 |
| Kif24 | 276,610531 | 141,66842 | 191,466131 | 102,040551 |
| Klhl30 | 16,3891757 | 4,27091111 | 7,31560135 | 5,24033551 |
| Klhl33 | 127,668377 | 47,7142098 | 41,9854989 | 37,1961 |
| Kmo | 120,309278 | 52,2663453 | 85,0943197 | 56,3224961 |
| Knl1 | 704,518268 | 378,069968 | 423,713151 | 201,358453 |
| Lgr6 | 14,489891 | 4,64565781 | 5,64888568 | 4,06666697 |
| Lmo7 | 75,4812468 | 31,2754577 | 51,9800165 | 33,4806878 |
| Lpar5 | 664,551265 | 319,395223 | 709,261982 | 331,039031 |
| Lrrn1 | 50,0294253 | 19,5045439 | 93,4097658 | 71,5783802 |
| Lrrn3 | 68,9812002 | 32,7299752 | 43,7960991 | 27,2853345 |
| Mab21l2 | 1,08617454 | 0 | 8,01639115 | 5,24033551 |
| Malat1 | 15919,6863 | 8011,8064 | 6203,4372 | 5874,01206 |
| Mamdc2 | 132,988642 | 62,1266238 | 294,159814 | 155,667736 |
| Map2k6 | 546,99372 | 207,015175 | 619,491096 | 340,775732 |
| Mbd4 | 478,49675 | 216,348161 | 315,730368 | 165,300919 |
| Mir6935 | 72,8690088 | 37,3482707 | 41,9093346 | 29,5818517 |
| Mir6942 | 11,0690967 | 4,89409669 | 14,2797072 | 13,9439546 |
| Mir7060 | 46,8582084 | 15,9123824 | 34,4848323 | 29,6474344 |
| Mmp28 | 168,135781 | 69,8337375 | 149,544721 | 76,9942898 |
| Mmrn1 | 197,209136 | 95,9013941 | 80,929683 | 49,4087462 |
| Mob3b | 241,163357 | 125,328187 | 183,994044 | 100,126096 |
| Ms4a14 | 563,470309 | 207,91754 | 288,582176 | 147,853124 |
| Mybl1 | 83,9923464 | 42,1199307 | 55,5318055 | 27,334584 |
| Mybpc1 | 51,8886655 | 13,8171791 | 31,4247287 | 15,1014793 |
| Naaladl1 | 25,1260416 | 9,02556517 | 12,3617812 | 5,65165475 |
| Nat8f2 | 4,40485654 | 0,89206629 | 0,91706059 | 0,31104661 |
| Nkx6-1 | 14,318563 | 3,90660615 | 10,6506532 | 8,92898164 |
| Nlrc4 | 541,879351 | 260,253513 | 385,632116 | 268,555212 |
| Nup210l | 44,9954093 | 15,2752686 | 22,2166147 | 19,462675 |
| Nxpe5 | 206,270685 | 76,0288515 | 125,381133 | 54,8050428 |
| Olfml2a | 264,190482 | 114,493906 | 142,159778 | 94,5959195 |
| Otof | 5,67517106 | 0,90044033 | 2,14537011 | 2,01166889 |

| | | | | |
|---|---|---|---|---|
| Pax3 | 0,99913511 | 0,31659284 | 14,4451185 | 10,4371375 |
| Pcdh18 | 140,637182 | 59,7388731 | 92,8107733 | 76,0713444 |
| Phf21b | 144,007627 | 66,3980544 | 104,626768 | 91,7984305 |
| Pik3c2b | 670,948378 | 317,358861 | 684,425225 | 346,679767 |
| Pkd1l1 | 12,0662565 | 1,59003195 | 8,05519647 | 5,62190002 |
| Pla2r1 | 28,4574027 | 11,0596062 | 13,1157621 | 16,9244309 |
| Plscr4 | 19,3809936 | 8,24654518 | 13,5789221 | 6,05391861 |
| Plxnd1 | 6326,03541 | 3158,90919 | 4340,08654 | 2408,78117 |
| Ptgdr | 12,6846657 | 4,1910505 | 1,13622362 | 2,33788056 |
| Ptger3 | 59,5189584 | 22,6595178 | 79,0871723 | 42,7104358 |
| Ptprb | 1754,48128 | 714,01575 | 1296,26624 | 613,720691 |
| Rad54b | 443,552358 | 222,076955 | 234,463406 | 113,485319 |
| Ranbp3l | 167,086071 | 90,8691886 | 222,501561 | 107,039476 |
| Rbm11 | 3,95036966 | 0,84860212 | 3,26193518 | 2,0412193 |
| Ret | 119,332078 | 57,5940853 | 100,51669 | 55,1986288 |
| Rho | 7,93455473 | 2,74315911 | 7,3972079 | 5,0766398 |
| Rpgrip1 | 17,2206003 | 5,72055042 | 9,85008528 | 9,121048 |
| Rsg1 | 6,10550408 | 1,79338059 | 3,8934788 | 0,58683427 |
| Samd11 | 74,4039881 | 26,9783803 | 61,5666018 | 24,6401241 |
| Sbspon | 9,50753217 | 3,81217771 | 12,8751646 | 8,45049134 |
| Scel | 32,6284173 | 15,1141143 | 32,56175 | 13,583468 |
| Scn7a | 6,49636269 | 1,69545631 | 5,79506555 | 2,51832442 |
| Serpinb1c | 21,3466479 | 6,98724155 | 17,9626463 | 14,5010405 |
| Serpind1 | 61,8449544 | 26,3146673 | 19,5686875 | 26,312764 |
| Skida1 | 23,8914463 | 6,87637552 | 17,7078231 | 14,016062 |
| Slc26a11 | 715,563065 | 329,807893 | 464,505814 | 329,093357 |
| Slit2 | 125,738137 | 51,6407722 | 40,5640002 | 42,6383532 |
| Sned1 | 389,143296 | 211,502709 | 165,562771 | 118,166785 |
| Snhg14 | 13,3909233 | 6,63746303 | 10,4251184 | 9,44077142 |
| Sp7 | 7,81034482 | 1,21203312 | 4,15989304 | 1,89130237 |
| Spata25 | 16,4162329 | 3,12115794 | 13,8000745 | 7,4225906 |
| Spic | 95,2719511 | 36,2248748 | 77,9131123 | 74,3297803 |
| Spon2 | 34,6757135 | 15,8619724 | 12,1086051 | 6,33264446 |
| Tbc1d4 | 2441,16796 | 1256,56111 | 2411,50135 | 1327,02173 |
| Tbx18 | 125,955679 | 69,3897855 | 55,2799152 | 38,1792842 |
| Tbx6 | 186,099944 | 99,5309528 | 157,636581 | 130,522294 |
| Tcf7 | 228,319299 | 114,034372 | 125,598086 | 72,6729067 |
| Tchh | 13,5273222 | 6,34286204 | 8,6021172 | 8,77552216 |
| Tead4 | 31,9417299 | 12,522024 | 15,5368498 | 16,1918213 |
| Tfap2b | 25,4174393 | 15,3140797 | 26,6402919 | 19,6903766 |
| Ticrr | 823,984985 | 385,363535 | 446,600535 | 230,562898 |
| Tll1 | 27,8841173 | 15,6051502 | 7,71065981 | 6,60272358 |
| Trerf1 | 179,669216 | 96,0970887 | 166,525823 | 88,5261023 |
| Trim63 | 92,6392088 | 36,4819709 | 137,12591 | 90,5851183 |
| Trp53cor1 | 95,3105168 | 46,2344532 | 119,519452 | 90,3594642 |
| Tspan10 | 27,7402272 | 7,48856517 | 35,5692937 | 17,9395057 |

| | | | | |
|---|---|---|---|---|
| Wnt9a | 30,8222161 | 11,5325876 | 11,5480552 | 10,5789756 |
| Wscd2 | 9,61794028 | 3,79542963 | 5,60715208 | 2,96037492 |
| Xkrx | 32,6406154 | 16,5983929 | 32,1407177 | 15,8214771 |
| Zan | 6,7236134 | 0,90044033 | 2,31693965 | 1,61551471 |
| Zfhx2os | 9,21046638 | 5,06092703 | 7,82584766 | 4,33181726 |
| Zfhx4 | 39,5555126 | 12,3878251 | 15,3686796 | 15,8555552 |
| Zfp469 | 9,76803171 | 3,69663139 | 12,1570967 | 5,39339495 |
| Zfp521 | 127,540117 | 68,8589885 | 106,018174 | 52,446893 |
| Zfp536 | 15,8597813 | 7,06503448 | 11,1074414 | 8,46093189 |
| Zfp820 | 49,0258646 | 25,8591861 | 53,828745 | 23,8785932 |
| Zgrf1 | 522,274368 | 242,876248 | 295,127121 | 167,64788 |
| Zic5 | 49,1014528 | 22,8546049 | 27,7732937 | 17,756897 |
| Zscan30 | 26,0877634 | 12,6704802 | 16,4955152 | 15,9501268 |

### Supplementary table 6: Outputs from the cluster-based gene analysis.

| CLUSTER 1 | | | | | | | | |
|---|---|---|---|---|---|---|---|---|
| Term | Count | % | PValue | Genes | List Total | Pop Hits | Fold Enrichment | Benjamini Pvalue |
| GO:0006954~inflammatory response | 32 | 8,840 | 6,37E-15 | CCL12, CXCL9, | 290 | 344 | 5,800160385 | 5,89E-12 |
| GO:0002376~immune system process | 30 | 8,287 | 4,06E-12 | HP, NLRC5, IL4 | 290 | 383 | 4,88394706 | 2,51E-09 |
| GO:0006955~immune response | 25 | 6,906 | 1,31E-11 | CCL12, CXCL9, | 290 | 272 | 5,730856998 | 6,07E-09 |
| GO:0008284~positive regulation of cell proliferation | 22 | 6,077 | 1,82E-04 | CSF3, PDGFA, | 290 | 542 | 2,530881792 | 1,35E-02 |
| GO:0070374~positive regulation of ERK1 and ERK2 cascade | 20 | 5,525 | 1,88E-10 | CCL12, CCL22, | 290 | 188 | 6,633162142 | 6,96E-08 |
| GO:0071346~cellular response to interferon-gamma | 19 | 5,249 | 1,63E-17 | GBP6, GBP5, C | 290 | 68 | 17,42180527 | 3,01E-14 |
| GO:0010628~positive regulation of gene expression | 18 | 4,972 | 2,49E-04 | VDR, PLAUR, P | 290 | 399 | 2,812859736 | 1,62E-02 |
| GO:0045087~innate immune response | 18 | 4,972 | 2,53E-04 | ZBP1, ANXA1, | 290 | 400 | 2,805827586 | 1,62E-02 |
| GO:0006935~chemotaxis | 16 | 4,420 | 6,53E-10 | CCL12, CXCL9, | 290 | 118 | 8,45447107 | 2,01E-07 |
| GO:0071356~cellular response to tumor necrosis factor | 15 | 4,144 | 2,41E-09 | CCL12, CCL22, | 290 | 110 | 8,502507837 | 4,96E-07 |
| GO:0019221~cytokine-mediated signaling pathway | 14 | 3,867 | 6,46E-07 | CCL12, IL1RN, | 290 | 146 | 5,978932452 | 7,97E-05 |
| GO:0030593~neutrophil chemotaxis | 13 | 3,591 | 8,66E-10 | CCL12, CCL22, | 290 | 69 | 11,74742629 | 2,13E-07 |
| GO:0045766~positive regulation of angiogenesis | 13 | 3,591 | 5,50E-07 | C5AR1, THBS1, | 290 | 121 | 6,698945569 | 7,26E-05 |
| GO:0042127~regulation of cell proliferation | 13 | 3,591 | 3,04E-04 | CXCL9, ANXA1, | 290 | 227 | 3,570803585 | 1,76E-02 |
| GO:0070098~chemokine-mediated signaling pathway | 12 | 3,315 | 9,22E-10 | CXCL9, CXCL11 | 290 | 55 | 13,60401254 | 2,13E-07 |
| GO:0071347~cellular response to interleukin-1 | 12 | 3,315 | 5,67E-08 | IL6, CCL12, CCL | 290 | 80 | 9,352758621 | 8,06E-06 |
| GO:0032496~response to lipopolysaccharide | 12 | 3,315 | 3,39E-04 | THBD, CXCL9, | 290 | 197 | 3,798074567 | 1,85E-02 |
| GO:0050729~positive regulation of inflammatory response | 11 | 3,039 | 5,64E-08 | CCL12, CCL8, F | 290 | 63 | 10,88680898 | 8,06E-06 |
| GO:0035458~cellular response to interferon-beta | 10 | 2,762 | 1,35E-08 | GBP6, TGTP2, T | 290 | 41 | 15,20773759 | 2,49E-06 |
| GO:0060326~cell chemotaxis | 10 | 2,762 | 4,18E-06 | CXCL9, CXCL11 | 290 | 78 | 7,993810787 | 4,54E-04 |
| GO:0051897~positive regulation of protein kinase B signaling | 10 | 2,762 | 2,51E-05 | IL6, CSF3, F10, | 290 | 97 | 6,428012798 | 2,45E-03 |
| GO:0042832~defense response to protozoan | 9 | 2,486 | 1,67E-08 | GBP6, BATF2, IL | 290 | 30 | 18,70551724 | 2,80E-06 |
| GO:0050728~negative regulation of inflammatory response | 9 | 2,486 | 7,57E-05 | MIR147, CTLA2 | 290 | 87 | 6,450178359 | 6,37E-03 |
| GO:0050830~defense response to Gram-positive bacterium | 9 | 2,486 | 1,22E-04 | GBP6, FGR, GB | 290 | 93 | 6,03403782 | 9,77E-03 |
| GO:0002548~monocyte chemotaxis | 8 | 2,210 | 2,94E-06 | LGALS3, CCL12 | 290 | 40 | 12,47034483 | 3,40E-04 |
| GO:0007200~phospholipase C-activating G-protein coupled rec. sign. pathway | 8 | 2,210 | 4,64E-05 | C5AR2, P2RY2, | 290 | 60 | 8,313563218 | 4,29E-03 |
| GO:0043434~response to peptide hormone | 8 | 2,210 | 1,64E-04 | GJA1, ANXA1, S | 290 | 73 | 6,833065659 | 1,26E-02 |

| | | | | | | | | |
|---|---|---|---|---|---|---|---|---|
| GO:0048661~positive regulation of smooth muscle cell proliferation | 8 | 2,210 | 2,91E-04 | IL6, CCL5, MYC | 290 | 80 | 6,235172414 | 1,73E-02 |
| GO:0009615~response to virus | 8 | 2,210 | 3,93E-04 | TGTP2, CDK6, T | 290 | 84 | 5,938259442 | 2,07E-02 |
| GO:0042060~wound healing | 8 | 2,210 | 7,75E-04 | IL6, CDH3, WNT | 290 | 94 | 5,306529714 | 3,50E-02 |
| GO:0048247~lymphocyte chemotaxis | 7 | 1,934 | 1,22E-05 | CCL12, CCL8, C | 290 | 33 | 13,2261233 | 1,25E-03 |
| GO:0043406~positive regulation of MAP kinase activity | 7 | 1,934 | 2,15E-04 | PDGFC, KIT, NC | 290 | 54 | 8,082630907 | 1,47E-02 |
| GO:0070301~cellular response to hydrogen peroxide | 7 | 1,934 | 2,90E-04 | IL6, TRPC6, AN | 290 | 57 | 7,65722928 | 1,73E-02 |
| GO:0034341~response to interferon-gamma | 6 | 1,657 | 5,03E-05 | TGTP2, H2-EB1 | 290 | 26 | 14,38885942 | 4,43E-03 |
| GO:0045740~positive regulation of DNA replication | 6 | 1,657 | 5,90E-04 | IL6, PDGFC, PD | 290 | 43 | 8,700240577 | 2,95E-02 |
| GO:0045600~positive regulation of fat cell differentiation | 6 | 1,657 | 0,00108 | WNT5B, NOCT, | 290 | 49 | 7,634904996 | 4,76E-02 |
| GO:1901741~positive regulation of myoblast fusion | 5 | 1,381 | 2,05E-04 | EHD1, CXCL9, C | 290 | 19 | 16,40834846 | 1,46E-02 |
| GO:0010575~positive regulation of vascular endothelial growth factor production | 5 | 1,381 | 5,28E-04 | C3, ADORA2B, C | 290 | 24 | 12,98994253 | 2,71E-02 |
| GO:0001895~retina homeostasis | 5 | 1,381 | 6,20E-04 | CDH3, TULP1, S | 290 | 25 | 12,47034483 | 3,02E-02 |
| GO:0044406~adhesion of symbiont to host | 4 | 1,105 | 3,16E-04 | GBP6, GBP7, G | 290 | 9 | 27,71187739 | 1,77E-02 |
| GO:2000660~negative regulation of interleukin-1-mediated signaling pathway | 3 | 0,829 | 7,56E-04 | IL6, IL1RN, IL1R | 290 | 3 | 62,35172414 | 3,49E-02 |
| GO:0033031~positive regulation of neutrophil apoptotic process | 3 | 0,829 | 7,56E-04 | HCAR2, ANXA1, | 290 | 3 | 62,35172414 | 3,49E-02 |

CLUSTER 2

| Term | Count | % | PValue | Genes | List Total | Pop Hits | Fold Enrichment | Benjamini |
|---|---|---|---|---|---|---|---|---|
| GO:0051607~defense response to virus | 13 | 10,32 | 4,81E-12 | GBP2B, RSAD2, | 77 | 167 | 18,28027063 | 2,11E-09 |
| GO:0002376~immune system process | 11 | 8,73 | 4,45E-06 | GBP2B, RSAD2, | 77 | 383 | 6,744498322 | 6,49E-04 |
| GO:0045087~innate immune response | 9 | 7,143 | 2,71E-04 | RSAD2, MX2, M | 77 | 400 | 5,283701299 | 1,98E-02 |
| GO:0009615~response to virus | 8 | 6,349 | 6,11E-08 | RSAD2, MX2, M | 77 | 84 | 22,36487322 | 1,34E-05 |
| GO:0035458~cellular response to interferon-beta | 5 | 3,968 | 2,59E-05 | GBP2B, IGTP, IF | 77 | 41 | 28,63794742 | 2,27E-03 |
| GO:0035457~cellular response to interferon-alpha | 4 | 3,175 | 1,53E-05 | IFIT3B, IFIT1, IF | 77 | 12 | 78,27705628 | 1,67E-03 |
| GO:0045071~negative regulation of viral genome replication | 4 | 3,175 | 3,24E-04 | RSAD2, MX2, IS | 77 | 32 | 29,3538961 | 2,03E-02 |

**Supplementary table 7:**
**Specific gene lists used to build the predicted protein networks in STRING.**

| *Cerebrum only* | Cerebellum only | SHARED GENES |
|---|---|---|
| | | Nipal1 |
| Gm16325 | Foxd3 | Serpina3g |
| Prss50 | Spr-ps1 | Bmp8a |
| Cxcl3 | Mpz | Iigp1 |
| Gm37194 | Rnase2a | Tgtp2 |
| Dcpp2 | Gng13 | Prss16 |
| Tnfsf14 | Ifit3b | Ccl17 |
| 1700023F06Rik | Gm10143 | Plet1 |
| Selp | Kcna1 | Tgtp1 |
| 2210008F06Rik | Csf3 | Mirt2 |
| 3110082J24Rik | Mc5r | Slc28a3 |
| Stfa2 | Fcnaos | Serpina3h |
| Rep15 | Ifit1 | Jaml |
| B230303A05Rik | Tomt | Gzma |
| BC048644 | Trim69 | Serpina3e-ps |
| Kcna7 | Vwc2 | Steap4 |
| Gm21887 | Cdh3 | Ms4a4b |
| Ngp | 1700016G22Rik | Gm5416 |
| Lcn2 | Ly6i | Serpina3i |
| Gm11707 | Gm8463 | Apol10b |
| Vaultrc5 | B9d1os | Serpina3f |
| Rab33a | Ifnlr1 | Gbp4 |
| Klrb1c | Ifit3 | Cxcl11 |
| Msr1 | Eomes | Gbp10 |
| Rpl28-ps1 | Htr2a | Il1r2 |
| Cnga2 | Clec4e | Zbtb16 |
| Ltf | Nt5e | Il2rb |
| Ffar2 | Chil5 | Cxcl9 |
| Hc | Ifit1bl1 | Plet1os |
| Lor | Esyt3 | Nrg4 |
| Gm10073 | A930030B08Rik | Gm807 |
| Serpina3n | Gm26448 | 2010005H15Rik |
| Hrct1 | A330023F24Rik | Rab44 |
| Fam83f | Ms4a8a | Cpm |
| Rpl10-ps3 | Ifi44 | Gm9733 |
| Gm32899 | Gm42928 | E230001N04Rik |
| Gm11808 | Mx1 | Upp1 |
| Gm12696 | Gm11620 | Klf9 |
| Gm26902 | Ccl8 | Vcam1 |
| Gm45855 | Chdh | Orm1 |
| Gm43703 | Cmpk2 | Rnase1 |
| AC127578.1 | Oasl1 | Areg |

| | | |
|---|---|---|
| AL845457.1 | Mgp | Sell |
| Dnajb3 | Arl9 | Kcna5 |
| Klc3 | Gjc3 | Jdp2 |
| Smim3 | 1700014D04Rik | Nlrp12 |
| Ccdc73 | Gjc2 | Gm10603 |
| Cd177 | Hhipl2 | Pla1a |
| Piwil2 | Gm4759 | Il2ra |
| Hcar2 | Cyp4f18 | Retnlg |
| Gm10051 | Rsad2 | Arc |
| Rpph1 | Plin4 | Stra6 |
| Rpl9-ps6 | Usp18 | Gm10524 |
| Mt1 | Cnksr1 | Gm12250 |
| Gm2000 | Tcap | Hif3a |
| Fgf2 | Gm8430 | Mrgpra2b |
| Gm11539 | Hspa1l | Fpr2 |
| Lipc | Gm44777 | Gm16194 |
| Rps18-ps3 | Ifit1bl2 | Cd3d |
| Rpl36a-ps2 | Mir17hg | Fkbp5 |
| Bex4 | Shisa2 | Gbp2 |
| Rps16-ps2 | Gm24958 | Mrap |
| Crybb1 | AL645522.1 | Trbc2 |
| Ptger4 | Snord37 | Map3k6 |
| Cd209a | 1700109H08Rik | Lbx1 |
| Gm19680 | Ifit2 | Gm26473 |
| Scimp | Sapcd1 | Il6 |
| Gm26619 | Gbp8 | Gm15675 |
| Rps10-ps1 | Ucp1 | Oas3 |
| Rps28 | Rpl7a-ps7 | Ccl5 |
| Shisa8 | Gm16287 | Capn11 |
| Gm5134 | Xdh | Hdc |
| Bcl3 | Gm26797 | Rdh12 |
| Rgcc | Ifi204 | Hs3st3b1 |
| Llph | Gm29264 | Serpinb10 |
| Gm45629 | Snord14e | Gm43181 |
| Gm27494 | Ppef2 | Slfn1 |
| Oaf | Cd209b | Gbp2b |
| Smkr-ps | CT009713.7 | Il1f9 |
| Rmrp | Nox4 | Mefv |
| AL732506.1 | Cldn19 | Creb5 |
| Scand1 | Vgf | Vdr |
| Gm4149 | 4933406C10Rik | Atp1a4 |
| Gm6807 | Klf4 | Gm43302 |
| Ildr1 | Ankrd35 | Ddit4 |
| Epb41l4aos | Gm7972 | Prss23 |
| Slc39a4 | Gm18407 | Gm20109 |
| Pstpip1 | Zglp1 | Gm26586 |

| | | |
|---|---|---|
| Rpl31-ps8 | Txndc2 | Ifi205 |
| Tlr7 | Irgm2 | Gm45774 |
| n-R5-8s1 | Coro6 | Fam107a |
| Cryba4 | Cdh1 | Ttc39c |
| Casp4 | Cabyr | I830127L07Rik |
| Cd40 | Sdcbp2 | S100a7a |
| Ecscr | Sult2b1 | Prss12 |
| Gm9843 | Anxa1 | Cd300lf |
| Rpl10a-ps1 | Kcne4 | Gbp5 |
| Gm17455 | Hormad2 | Trim30b |
| E430024P14Rik | Ctrl | Ly6g |
| Sec61g | Mall | Adam8 |
| Ccno | Tspoap1 | Gda |
| 1500009C09Rik | 9130230N09Rik | Galnt15 |
| Crip1 | Slc5a5 | Dok2 |
| Gm38079 | Nlrc5 | Ccl12 |
| Prrt1 | Gm3052 | Cx3cl1 |
| Eif4ebp1 | Gm43062 | Arg2 |
| Gm26881 | Gprc5a | Gm28592 |
| Dcxr | Rnf225 | Mmp8 |
| Mcoln2 | Gm29650 | Cd55 |
| 3010003L21Rik | Oasl2 | Gbp6 |
| Ninj1 | Mx2 | Ramp3 |
| Polr2l | Ctgf | Il15ra |
| Zfp335os | Itgb7 | Clec4d |
| Il2rg | AW046200 | Arhgef37 |
| Cxcl16 | Gm42732 | Acod1 |
| Lrrc51 | Pou3f1 | Gpr141 |
| Pik3ip1 | L3mbtl1 | Chrm3 |
| Sla | 9130213A22Rik | Wfdc21 |
| Timm8a1 | Slc1a1 | Aqp9 |
| Chad | Adam33 | Mir147 |
| Endog | Phyhip | Socs2 |
| Id1 | Thbd | Stfa2l1 |
| S100a4 | Gm31597 | Syde2 |
| Macrod1 | Gm37607 | Ocstamp |
| Gm14586 | Helz2 | Tarm1 |
| Wdr74 | 1700034H15Rik | Abtb2 |
| Lck | Gimap4 | Slc16a14 |
| Cxcl14 | Il18bp | Pdzd3 |
| Wdr89 | Proser2 | Tmem252 |
| Fam60a | Slfn5 | F10 |
| St3gal1 | Oas1b | Runx3 |
| Havcr2 | Gm27390 | Prok2 |
| Arhgap15 | AC129931.1 | Hspa1b |
| Mdfic | Strip2 | Gm38150 |

| | | |
|---|---|---|
| Gm42418 | Klf5 | Gm5483 |
| Hcst | Adgrg3 | Sh3pxd2b |
| Tnfrsf12a | Slc5a3 | Gm14964 |
| Gmppb | Gm26737 | Scn2b |
| Phb | AC154636.3 | Slc39a14 |
| Sugct | Nhs | 1700047M11Rik |
| Lrrc8c | Adgrg6 | Vcan |
| Prr7 | Gfpt2 | Gm23935 |
| Tmem158 | Otub2 | Prnd |
| Polr1d | Tbx15 | Il12rb1 |
| 1500009L16Rik | Adm | Camp |
| Ebi3 | Rnf223 | Cldn1 |
| Trir | Doc2g | Nos2 |
| Ncf1 | Slc45a3 | Ltb4r1 |
| Samsn1 | Fam19a3 | Lefty1 |
| Crlf2 | Ccdc141 | Per2 |
| Slc22a15 | Shroom2 | Adora2b |
| Tmem130 | Olfr56 | Pdgfc |
| Gm11478 | Gm11175 | Ly6c2 |
| Tes | AC124577.2 | Slc28a2 |
| Lrrc3 | Ikbke | Dhrs9 |
| AC113060.1 | Gt(ROSA)26Sor | Slc10a6 |
| Asns | Pnma2 | Gm9899 |
| Gm5874 | Gm31024 | Mcemp1 |
| Gm10110 | Tmem121b | Tacstd2 |
| Rps10-ps2 | Itih2 | Sema6b |
| Galnt7 | Zfp708 | Cdc42ep2 |
| Tmem65 | Gm9856 | Cdr2 |
| Mrto4 | Cd200 | Trem1 |
| Cdc34 | Cpeb2 | Pawr |
| Nol12 | Gm16973 | Asprv1 |
| Cdh13 | Runx2 | 3930402G23Rik |
| Eef1akmt4 | Lifr | Hp |
| Emc6 | Snai1 | Gja1 |
| Gm9794 | Mir7115 | Map2k3os |
| Mocs3 | Oas1a | Klra2 |
| Eif1b | C430049B03Rik | AA467197 |
| Gm21987 | Errfi1 | Egfr |
| Hivep3 | Dnah2 | Cish |
| 2310043P16Rik | Ppp1r3b | G0s2 |
| Slco3a1 | Irgm1 | Timp1 |
| Ifitm10 | Zfp948 | Ehd1 |
| Tpm3-rs7 | Rtp4 | Igfbp3 |
| Denr | Dgka | Met |
| Rpl13a | Zfp40 | Tbx21 |
| Tmem120a | Mlxip | Emp1 |

| | | |
|---|---|---|
| Pycr2 | Sik1 | Atp8b4 |
| Bop1 | Inpp5j | Mt2 |
| Rps2 | D330050G23Rik | Snord55 |
| Nr2f6 | Mal | Adrb1 |
| Nip7 | Zmynd15 | Isg15 |
| Rpl18a | Samd9l | Far2 |
| Pctp | Fas | Gm43802 |
| B3galt5 | Col27a1 | C1qtnf9 |
| Tpt1-ps3 | Dmpk | Igtp |
| Prdx5 | Rrad | Zbp1 |
| Eif6 | Casp12 | 6030468B19Rik |
| Itpkb | Clec12a | Syt6 |
| Ube2v2 | F3 | Tgm2 |
| R74862 | Gm28177 | Meox1 |
| Cd3eap | Foxred2 | Ascl2 |
| Med21 | Sqle | Gpat3 |
| Rps29 | Cp | Socs1 |
| Marcksl1 | Synpo | Chil1 |
| Rps10 | Cdc42ep3 | Plbd1 |
| Psmb10 | Lama4 | C5ar1 |
| Prnp | Ankrd23 | S100a6 |
| Elp5 | Celf5 | Ldhd |
| Fth1 | D930016D06Rik | Batf2 |
| Pik3r5 | Col7a1 | Napsa |
| Spred1 | Stat1 | Efna2 |
| Grin1 | Tal1 | Ifi47 |
| 1300002E11Rik | Eif2s3y | Prn |
| Hmga1 | AC163667.1 | Bbs12 |
| Metrnl | Nedd9 | Erich3 |
| Sac3d1 | Rbm12b1 | Trpc6 |
| Gnl1 | Stat2 | Cfap69 |
| Eif3f | Pla2g3 | Irf7 |
| Fut4 | Nr6a1 | Kcnn4 |
| Cox5b | Homer2 | A530076I17Rik |
| Fdx1l | Ubp1 | Gm31718 |
| Spred2 | Gm17435 | Rasd1 |
| Bend3 | Gm28038 | Rhov |
| Use1 | Simc1 | Rnd3 |
| B4galt6 | Bach2os | Afap1l1 |
| Rps5 | Fam129c | Runx1 |
| Eef1b2 | Taf1d | Thbs1 |
| Lyar | Aoc2 | Fabp4 |
| Trnau1ap | Bcl6b | S100a10 |
| Gm38843 | Pmel | Srgn |
| Rps9 | Plekhb1 | Mmp19 |
| Gm5239 | Pnisr | Kit |

| | | |
|---|---|---|
| Atf6 | Irf1 | Il18rap |
| Pvt1 | Gm28035 | Sult1a1 |
| Apoe | Gm28496 | Mlph |
| Rpl23 | Prr18 | Hs3st3a1 |
| Uba52 | Arid5a | Ctla2b |
| Pfn1 | Parp14 | Adamtsl4 |
| Ntmt1 | Pvr | Serpinb1a |
| Spryd4 | Ifih1 | AC160962.1 |
| BC031181 | Relt | Cyp1b1 |
| Gm17084 | Foxc1 | Hes1 |
| Nr2c2ap | AI606181 | Gbp3 |
| Taf10 | 2810403A07Rik | Madcam1 |
| Pus1 | Mlxipl | Tmem236 |
| Rps7 | Pabpn1 | Trem3 |
| Slc12a7 | Sybu | Nr1h3 |
| Igsf8 | Col15a1 | Wnt5b |
| Nxt1 | Gm15446 | Ldlr |
| Nufip1 | Tob1 | Glipr2 |
| Eid2 | Socs3 | Cidea |
| Fam120aos | Plk2 | Cdk6 |
| Nek6 | Gcc1 | C5ar2 |
| Cmklr1 | Myo19 | Gm16201 |
| Myo1e | Zfp202 | C030034L19Rik |
| Bag1 | Pde12 | Nudt10 |
| Gemin7 | Paxbp1 | Gbp7 |
| Orai1 | Crebzf | Slc16a12 |
| Eif2b3 | Lrif1 | Tubb6 |
| Hspa8 | Insig1 | Noct |
| Tysnd1 | Bcl2 | Wdr38 |
| Dubr | Nckap5l | Rps27rt |
| Polr2h | Zcchc7 | Ankrd33b |
| Chmp4b | E2f3 | Rhou |
| Higd1a | Zbtb48 | Olfm4 |
| Tomm5 | Fbxl7 | Plac8 |
| Mrps6 | Gm20521 | Slc41a2 |
| Rpl19 | Ddx58 | Rufy4 |
| Gtpbp6 | Fgfr1 | Cd44 |
| Gemin4 | Arl4a | Mpzl3 |
| Ppp1r11 | Mag | Slc4a11 |
| Timm23 | Dnajb9 | Ctla2a |
| Ctnnbip1 | Pde1b | Myc |
| H13 | Irf9 | Tgm1 |
| Jagn1 | Sema4a | Ctsw |
| Ado | Trmt10a | Bace2 |
| Gm6472 | Grk4 | Il4ra |
| Lypla2 | Slc25a25 | Igf2bp1 |

| | | |
|---|---|---|
| Flot1 | Il13ra1 | Uck2 |
| Ctdnep1 | Gm20708 | Ccr2 |
| Lpcat2 | Taf13 | Il17ra |
| Eif3g | Col11a2 | Mb21d2 |
| Psmc5 | Dnajb1 | Chka |
| Dnajc2 | Rbm26 | Tnfrsf13c |
| Inafm1 | Gfap | Papss2 |
| Ube2s | Gm42417 | Srgap1 |
| Rpl5 | Fubp1 | Tmod1 |
| Gm20458 | Mtrf1l | Stx11 |
| D430040D24Rik | Rccd1 | Acpp |
| Fbxl14 | Dcun1d3 | AC168059.2 |
| Kctd11 | Zfp276 | Pglyrp1 |
| Cnn2 | Rassf1 | Gstm6 |
| Eif3b | Mafk | Cd74 |
| Dda1 | Agap2 | Cxcr2 |
| Parl | Akap8l | Nabp1 |
| Hnrnpab | Lrrc8a | CT010498.1 |
| Dnajb2 | Rnf19b | Bcl2l1 |
| Slc25a10 | Cyp4f13 | Hivep2 |
| Psmd7 | B3galnt2 | Ly6a |
| Rcl1 | Parp3 | Fam241a |
| Rpl10a | Chordc1 | Fam78b |
| Cmtm8 | Paqr8 | Bst1 |
| Rras | Parp10 | Phlda1 |
| E130309D02Rik | Trit1 | Cebpb |
| Sarnp | Luc7l | Isg20 |
| Id2 | Hk2 | Ccnd1 |
| Gm20509 | Ccnyl1 | Ciita |
| Magoh | Rabggtb | Rcan1 |
| Kif3b | Susd1 | P2ry2 |
| Zpr1 | Hs6st1 | Pparg |
| Dxo | Csrnp2 | Acap1 |
| Mrpl20 | Tap1 | Mras |
| Hvcn1 | Ccnl2 | 1700001P01Rik |
| Hpcal1 | Ppig | Rrs1 |
| Sod2 | Gsdmd | Ifitm3 |
| Ppat | C330018D20Rik | H2-Aa |
| Sephs2 | Cry2 | Gadd45a |
| Dbndd2 | Hspa2 | Osmr |
| Tsg101 | Rcc1 | Timp3 |
| Bloc1s4 | Gramd1a | Fhl3 |
| Tle3 | 1810011H11Rik | Bcl2a1a |
| Ucp2 | Nae1 | Rnf169 |
| Rbm8a | Taf1a | Nop58 |
| Sys1 | Rbm38 | Ell2 |

| | | |
|---|---|---|
| Arpc2 | Cdyl | Hunk |
| Fra10ac1 | Rhbdf1 | Hbegf |
| Ddx49 | Papd7 | Gm42595 |
| Ralb | Tmem63a | Grhl1 |
| Gm6969 | Smad7 | 1810032O08Rik |
| Gm20427 | Tnip2 | Lmna |
| Ssbp4 | Sept7 | Tma16 |
| Med8 | Zfp623 | Nfe2 |
| Tm2d1 | Thoc1 | Polr1b |
| Pdf | 0610010F05Rik | Hcar1 |
| Rbpms | Fam133b | Cables1 |
| Slc39a7 | Akap8 | Fbp1 |
| Gfod2 | Sugp2 | Tmem52 |
| Utp23 | Tmem39a | Arhgap26 |
| Mrpl50 | Tbkbp1 | Emilin2 |
| Map2k2 | Slc25a22 | Tsr1 |
| Rcc1l | Gm44250 | Pdgfa |
| Adprhl2 | Tardbp | Man2a1 |
| Csnk2b | Parp12 | Cdc42bpg |
| Rtn4 | Sirt7 | Gtpbp4 |
| Chst7 | Med26 | S100a11 |
| Slc7a6 | Prpf40b | Traf1 |
| 0610037L13Rik | Unc45a | Snora65 |
| Ogfr | BC017643 | Gpt2 |
| Plekhj1 | Kansl3 | Ston2 |
| Fam207a | Dvl1 | Stk40 |
| Taf12 | BC024978 | H2-Eb1 |
| Snx3 | Mcl1 | Olfm1 |
| Ssu72 | Ing5 | CT010498.2 |
| 4833439L19Rik | Oxsr1 | Nfil3 |
| Me1 | Rab18 | Peg13 |
| Ptges2 | Ppp1r18 | Slc30a1 |
| Rbm42 | 9430015G10Rik | 37681 |
| Rabepk | Uckl1 | BC049352 |
| Cbfb | Bag3 | Rbm47 |
| Pdxk | Ggnbp2 | Ptpn12 |
| Snta1 | Sltm | Mir5107 |
| 1700052K11Rik | Ppp1r12c | H2-Ab1 |
| Med10 | Klc2 | Fgr |
| Rpl7 | Pgrmc1 | Atp1b1 |
| Exosc4 | Lgals8 | Klf13 |
| Nop9 | Cstf3 | Pnp2 |
| Sept9 | Ints13 | Rgs9 |
| Mtmr11 | Srfbp1 | Col14a1 |
| Ak1 | Fchsd1 | Spata5l1 |
| Nt5c3 | Ppp4r2 | E030030I06Rik |

| | | |
|---|---|---|
| Slc9a3r1 | Dnajc18 | Ass1 |
| Mrpl3 | Snx14 | AC115037.1 |
| Rsl1d1 | Rnaseh1 | Gm5905 |
| Zfp213 | Morf4l2 | Cfap74 |
| Tex30 | Bms1 | Dusp16 |
| Cct2 | Rcn2 | P2ry14 |
| Tprn | Rnf34 | Slc35e4 |
| Pitpnb | Gpsm1 | Phospho1 |
| Fbxo4 | Tial1 | Rxrg |
| Larp7 | Zfp865 | Pag1 |
| Lsm1 | Ahsa2 | Snhg15 |
| Dus1l | Srsf2 | Gprc5b |
| Eif4a3 | Msl1 | Riox1 |
| Plpp2 | Srsf7 | Nolc1 |
| Pald1 | Sfswap | Sbno2 |
| Npc2 | Ssb | Cd300lb |
| Ski | Ergic2 | P2ry1 |
| Smn1 | Dhx34 | Ppp1r3d |
| Wdr55 | Tbk1 | Ppan |
| Zfp566 | Pdcd7 | Sgms1 |
| Phc2 | Dcaf6 | Garem2 |
| Eapp | Srpr | Peg10 |
| Vamp3 | Mia3 | Cebpd |
| Tmem70 | Mta2 | Eif1a |
| Abt1 | Cables2 | Nab1 |
| Ybx3 | Rbm6 | Marveld1 |
| Ube2e1 | Tor1b | Apoo |
| Crls1 | Sf3b2 | Snhg17 |
| Ldha | Pigt | Ugcg |
| Faap100 | Raver1 | Pdp1 |
| Mindy1 | Rnpepl1 | Gadd45g |
| Map4 | Lrpap1 | Vav3 |
| Lrrc59 | Cog7 | Sema7a |
| Fem1a | Sumo3 | B3gnt7 |
| Cdkn2aipnl | Vac14 | Tmem2 |
| Ears2 | Snd1 | Nop2 |
| Timm29 | Uqcrc2 | Gm15501 |
| Zadh2 | Cap1 | Ccdc86 |
| Srprb | Crat | Mrm3 |
| Mrps10 | Gusb | Polr3d |
| Rer1 | Pi4kb | Bmf |
| Nsun2 | Exoc7 | Mphosph10 |
| Prdx6 | Mpv17 | Lrp12 |
| Tymp | Dis3l2 | Mgst1 |
| Tm9sf3 | Retreg2 | Trmt61b |
| Rae1 | Tkfc | Itsn1 |

| | | |
|---|---|---|
| Slc39a1 | Zfp385a | Tcf7l2 |
| Dnajb6 | Smarca4 | Slc2a3 |
| Xrcc4 | Ddb1 | Zhx2 |
| Psmc2 | Spg21 | Aen |
| Rab2a | Nt5c2 | Plaur |
| Irf2bp1 | Jmjd8 | Mak16 |
| Ahsa1 | Ddost | Gm20712 |
| Mlf2 | Tfip11 | Hinfp |
| Wdr1 | Bnip3l | Il1rn |
| Heatr3 | Prkaca | Gpatch4 |
| Ube2k | Hsdl1 | Ifi35 |
| Slc11a2 | Gdi2 | Crispld2 |
| 2610001J05Rik | Eif2a | Klhl2 |
| Aars | Fig4 | Glipr1 |
| Siah2 | Pccb | Ankrd9 |
| Grn | Hsd17b4 | Unc119 |
| Rnase4 | Ap3s2 | Lgals3 |
| Csnk2a1 | Sdhd | Rpp38 |
| Cox10 | Tex261 | 5430416N02Rik |
| Rtn1 | Ifngr2 | Ahdc1 |
| Atg3 | Mfsd10 | Rnd1 |
| AC151967.1 | Arsa | Wdr43 |
| Ube2f | Ipo11 | Ctu1 |
| Crcp | Lrrc61 | Pinx1 |
| Slc52a2 | Mrpl39 | Spns3 |
| Cct8 | Tmx2 | BC037034 |
| Rhog | Ryk | Plp2 |
| Snx17 | Agtrap | Pdlim7 |
| Tmbim4 | Gm2a | Ier5l |
| Gars | Sppl3 | Nifk |
| Sec23b | Iba57 | Hgsnat |
| Ncbp1 | Wdr73 | Icosl |
| Bag5 | Vhl | Hilpda |
| Litaf | Trap1 | Eef1e1 |
| Utp6 | Srebf1 | Ppp1r13b |
| Arfgap3 | Usp30 | B4galt1 |
| Gabarapl2 | Cptp | Fhod1 |
| Atp1a1 | Plin3 | Grwd1 |
| Nom1 | Mthfd1 | Fhad1 |
| Mrps17 | Cux1 | Relb |
| Rnf41 | Gnas | Utp4 |
| Prmt5 | Kbtbd4 | Stk38l |
| Snrnp40 | Fadd | Homer1 |
| Prkag1 | Slc35f6 | Gm17066 |
| Mrpl16 | Nat6 | Rbms1 |
| Psmd12 | Rab9 | Srxn1 |

| | | |
|---|---|---|
| Smim7 | Ddx59 | Irf2bp2 |
| Ube2z | Idh1 | Galnt18 |
| Bloc1s5 | Tmem97 | Mafb |
| Anxa7 | Gtf3c5 | Pgs1 |
| Med22 | 2310022A10Rik | Kti12 |
| Nras | St6galnac4 | Ccnd2 |
| Asnsd1 | Emc10 | Donson |
| Pgd | Borcs7 | Zbtb46 |
| Thoc3 | Fam219a | Spns2 |
| Cdca4 | Zkscan17 | Zfp593 |
| Adipor2 | Got2 | Gm9625 |
| Elovl1 | Atxn10 | Nol9 |
| Taf5l | Echs1 | 4833413G10Rik |
| Arpc5 | Hps6 | 6330403L08Rik |
| Tmem38b | Pgam1 | H2-T23 |
| Nupl2 | Naglu | D16Ertd472e |
| Prkar1a | Ttll12 | Lrrfip1 |
| Hspd1 | Mrpl10 | Mir1668 |
| Hdgf | Gna12 | Pnp |
| Cmpk1 | Abhd4 | B4galt5 |
| Agpat5 | Ctsl | Trip10 |
| Ippk | Gatd1 | Pnp |
| Cisd2 | Homez | Prr5l |
| Tmem167 | Cln6 | Trmt61a |
| Ap2b1 | Abhd8 | Zdhhc2 |
| Ticam1 | Ttyh2 | Lrrc4 |
| Clptm1l | Acp1 | Eif3j1 |
| Sdhaf2 | Pef1 | Adgre5 |
| Wars | Jpt2 | St5 |
| Wdr77 | Cd302 | Ttpal |
| Smarca5 | Smpd1 | Vim |
| Gps1 | Morn4 | Usp36 |
| Spop | Oxa1l | Snx18 |
| Ncdn | Rpusd1 | Nkrf |
| Snx24 | Abcb6 | Znrf1 |
| Vps4a | Armc9 | Lonrf1 |
| Ctsa | Cep19 | Gpr146 |
| Abhd17c | Bckdha | Ndufaf4 |
| Tmem185b | Bckdk | Gnl3 |
| Ints5 | Ift46 | Exosc6 |
| Mettl16 | Slc39a11 | Rhoj |
| Rpl7l1 | Dtnb | Ogfrl1 |
| Stx12 | Dnaaf2 | Mmp14 |
| Rab5c | Rit1 | Ddx21 |
| Pa2g4 | Nudt19 | Art3 |
| Tdrd7 | Wbp2 | Pus3 |

| | | |
|---|---|---|
| Wsb2 | Siae | Ppp1r21 |
| Usp39 | Pebp1 | Bysl |
| Mapkapk2 | Fbxw4 | Taf4b |
| Csnk2a2 | Arl2bp | Picalm |
| Hnrnpf | Vps33a | Slc12a4 |
| Stard3nl | Maf1 | Nsun5 |
| Dnttip2 | Fundc2 | Xbp1 |
| Surf4 | Cln5 | Il1rl1 |
| Tmbim6 | Cdk4 | Mars2 |
| Smu1 | Gins4 | Lgi4 |
| Memo1 | Slc48a1 | Pacsin2 |
| Cdc16 | Supt7l | Tmem198b |
| Desi1 | Coil | Larp1b |
| Serbp1 | Shmt1 | Hsph1 |
| Prpf4 | Hagh | Mat2a |
| Cactin | Mis18a | Ctps |
| Smyd5 | Dnal4 | Chchd4 |
| Ubl3 | Wrap53 | Trim46 |
| Cog8 | Rfc3 | Rpf2 |
| Atp6v1h | Ncs1 | Dusp2 |
| Tmem19 | Mrpl49 | Nop56 |
| Pttg1ip | Eno1b | Slc2a1 |
| Eif4e2 | Ap5s1 | Mplkip |
| Tpd52l2 | Csrp1 | Adpgk |
| Ss18 | Lrfn3 | Vasp |
| Mesd | Orai2 | Ecm1 |
| Rela | Ripk3 | Rcsd1 |
| Fam204a | Ccm2 | Nlrx1 |
| Dazap2 | Ybey | Tsc22d3 |
| Rab8a | Got1 | Ccnd3 |
| Sqor | Zfyve21 | Napepld |
| Ykt6 | Msantd3 | AC131081.1 |
| Srp68 | Ckb | Alkbh1 |
| Ywhaz | Srd5a1 | Mtdh |
| Mettl9 | Galk1 | Rac3 |
| Rnf4 | Furin | Cbarp |
| Brd7 | Ypel1 | Asic1 |
| Tmem189 | Nt5dc2 | Nle1 |
| Rac1 | Otud3 | Dus4l |
| Hgs | Poc1a | Tbl2 |
| Agpat1 | Cby1 | Ddx17 |
| Rab11b | Ephx1 | Snu13 |
| Letm1 | Prdx4 | Rbpj |
| Calm1 | B9d2 | Gm15832 |
| Dars | Fam216a | Urb2 |
| Becn1 | Commd9 | Scn1b |

| | | |
|---|---|---|
| Sf3b4 | Dnajc9 | Slc29a2 |
| Ndrg3 | Mthfd1l | Rgmb |
| Ubl4a | Pdk3 | Msmo1 |
| Tmem127 | Bmyc | Man1a |
| Ppp2ca | Haus1 | Trmt10c |
| Cdc42 | Chd3os | Pim3 |
| Thoc5 | Dusp28 | Sptlc2 |
| Mboat7 | Kcnn1 | Klf16 |
| Rnf114 | Cox8a | Arhgap22 |
| Mrps2 | Acaa2 | Zfp930 |
| Trmt6 | Dusp18 | Rhoc |
| Plod3 | Cks1b | Gripap1 |
| Bcl10 | Selenoh | Ddx28 |
| Mark4 | Pdia5 | Agfg1 |
| 9530068E07Rik | Tacc3 | Mgat2 |
| Gpat4 | Scrn1 | Arrb2 |
| Xpnpep1 | Phf19 | Atg16l1 |
| Sec22b | Slc46a3 | Pex11a |
| Spcs2 | Rpa2 | Clic4 |
| Ubiad1 | Nipsnap1 | Creld2 |
| Imp4 | Ift27 | Cyp51 |
| Fam8a1 | Tmem17 | Lfng |
| Rbm22 | Ppih | Pprc1 |
| Tbc1d20 | B3glct | Cycs |
| Pcbp1 | Tyms | Nfyb |
| Get4 | Ak4 | Naa25 |
| Psmd5 | Nup37 | Ctu2 |
| Arl1 | Rftn1 | Polg |
| Dync1i2 | Gmpr | Tbc1d8 |
| Arf1 | Zfp691 | Slc31a2 |
| Metap2 | Chchd6 | Nus1 |
| Vapb | Ptgfrn | Zfp622 |
| Josd1 | Gm16580 | Usp16 |
| Ccdc43 | Cdk1 | Noc3l |
| Micu1 | Tmem204 | Sema4c |
| Kat14 | Tctn2 | Osbpl9 |
| Itm2c | Tmem175 | Nt5dc3 |
| Alg14 | 1700008J07Rik | Zmynd19 |
| Arl8a | Nim1k | Anxa3 |
| Rbm45 | Arl11 | Ftsj3 |
| Atp6ap1 | Fbxo17 | Nob1 |
| Polr2m | Pmf1 | Gpx3 |
| Dync1li1 | Depdc1b | Brix1 |
| M6pr | Efhc1 | Kri1 |
| P4hb | Ccne1 | Yars2 |
| Lamp1 | Cdt1 | Iffo2 |

| | | |
|---|---|---|
| Nmt1 | Atp5g2 | Tmem81 |
| Chtop | Mgst3 | Ppp2r5b |
| Exosc10 | AI427809 | Itga5 |
| Grb2 | Fen1 | Wdr75 |
| Clptm1 | Gcat | Scamp1 |
| Hnrnpl | Asf1b | Chst1 |
| Smc1a | Hmgb3 | Nab2 |
| Pitrm1 | E2f2 | Tank |
| Nup88 | Ighm | Snx2 |
| Pold3 | Gm6055 | Dnajc21 |
| Pomt1 | Pttg1 | Atp23 |
| Orc5 | Colca2 | Tpm4 |
| Primpol | Pigz | Tpst2 |
| Vars2 | Gpr162 | Gm6277 |
| Fbf1 | Gdf1 | Rrp9 |
| Vrk1 | AI467606 | Abce1 |
| Med24 | Ulk4 | Dph2 |
| Tmem110 | Rab7b | Hyou1 |
| Prr14 | Azin2 | Pigw |
| Eml3 | Vps37d | Wdr4 |
| Smpd4 | Nlrp1b | Dimt1 |
| Mccc1 | Pdgfrl | Hsbp1 |
| Abcd4 | Calr3 | 8430429K09Rik |
| Fam149b | Ntn4 | Fam129b |
| E130307A14Rik | Ccnb2 | Dram1 |
| Cnot3 | B9d1 | Fam220a |
| Zfp651 | Syce2 | Trp53rka |
| C87436 | Cdca8 | Gtf2a2 |
| Hps1 | Pbk | Ddx51 |
| Acap3 | Naaa | Ifrd2 |
| Odf2 | Gm6594 | Sox13 |
| Stx5a | Fabp5 | Sdad1 |
| Abcd1 | Cdc20 | Smox |
| Parp16 | Irak1bp1 | Utp11 |
| Atpaf2 | Mlf1 | Tlnrd1 |
| Ankmy2 | Ube2c | Tut1 |
| Gm43064 | Rbfox3 | Ptpn1 |
| Mkl1 | Birc5 | Mex3a |
| Hp1bp3 | Tmem107 | Dnajc25 |
| Stau1 | Cdkn3 | Heatr1 |
| D6Wsu163e | Sept4 | Itpripl1 |
| Lamtor3 | Fabp3 | Exosc1 |
| AL732309.1 | Lpl | B3galt6 |
| Tmpo | Cdc25c | Sipa1 |
| Lrch4 | 2010008C14Rik | Ccdc137 |
| Ap4b1 | Dnph1 | Banp |

| | | |
|---|---|---|
| Ctage5 | Cenpw | Rrp12 |
| Git2 | Car7 | Gm20537 |
| Acin1 | 1700025G04Rik | Ifngr1 |
| Gstm4 | Bcar3 | Mvp |
| Taok2 | Spc24 | Card19 |
| Ankrd10 | E130215H24Rik | Nmd3 |
| Limk2 | Tmem178 | 1110038B12Rik |
| Evi5l | Slpi | Rrp1b |
| Tmem129 | AC163703.1 | Map2k3 |
| Ugdh | Nlrp1a | Ppid |
| Aldh4a1 | Depdc1a | Utp18 |
| Mcoln1 | Gm15326 | Polr3e |
| Ptpn23 | Cenpm | Rbm34 |
| Clk4 | Cox4i2 | Cebpz |
| Zbtb44 | Clec2l | Nhlrc1 |
| Map3k14 | Gchfr | 1600014C10Rik |
| Exoc1 | Mnd1 | D1Ertd622e |
| Trafd1 | Kbtbd6 | Spata5 |
| Hars2 | Scml4 | Usp2 |
| Stoml1 | Ramp1 | Tmbim1 |
| Scrn3 | Ascl1 | Hgh1 |
| Wdr11 | Rmi2 | Gm45736 |
| Dgcr8 | Cxcr3 | 2410002F23Rik |
| Rbm39 | Kbtbd12 | Gnl2 |
| Scfd2 | Tmem266 | Ccdc134 |
| Cdadc1 | Crip3 | Gatsl3 |
| Rnf123 | Cyp2r1 | Rrp8 |
| Tmem143 | Wipf3 | Gdap2 |
| Gpbp1l1 | Cfap77 | Plekha3 |
| Recql5 | Dnali1 | Trim27 |
| Dcun1d2 | Gm4316 | Ccdc71l |
| Plekhm2 | Gm26520 | Fam217b |
| Tmem55a | Rab17 | Syngr2 |
| Zfp27 | B430305J03Rik | Srebf2 |
| Poli | Adam5 | Wfs1 |
| Unk | Perp | Dnaja2 |
| Uaca | 4833417C18Rik | Sco1 |
| Mavs | Actn2 | Arhgef7 |
| Terf1 | D330045A20Rik | Stat3 |
| Zfp839 | Meltf | Xkr8 |
| Suox | Fv1 | Fam118a |
| Zswim9 | Gm42793 | Lsm11 |
| Pbx3 | Gm7666 | Anapc16 |
| Mroh1 | Gm12286 | Slc7a5 |
| Ganc | AC027700.1 | Gm45867 |
| Trpm4 | Syngr4 | Dlg3 |

| | | |
|---|---|---|
| Ripor1 | Gna14 | Spryd3 |
| Cabin1 | Klhdc9 | Med7 |
| Cep131 | Lhx6 | Dnaja1 |
| Fuk | Tnfrsf26 | Zyx |
| Tctn3 | 2610035F20Rik | Ube2j2 |
| Esyt1 | Igfbpl1 | Spryd7 |
| Oser1 | 6030442K20Rik | Snupn |
| Ift172 | Gm20083 | Rab11a |
| Inpp5e | Gm10699 | Rrp15 |
| Ddhd2 | Ephx4 | Slc7a6os |
| Mppe1 | Gm42879 | Tapbpl |
| Tmem230 | Grp | Fez2 |
| Capn10 | Duoxa1 | Src |
| Zfp740 | Blk | Fbxo22 |
| Mbd1 | Kcnv1 | Tigd5 |
| Cep250 | | Surf6 |
| Rasa3 | | Ece2 |
| Plcd1 | | Znhit6 |
| Rgl1 | | Exoc8 |
| Arhgef10l | | Ncoa5 |
| Nipa1 | | Stim2 |
| Frg2f1 | | Uspl1 |
| BC017158 | | Sh3bp2 |
| Tinf2 | | Kras |
| Mlh1 | | Noc4l |
| Tspyl1 | | Aebp2 |
| AL732309.2 | | Irf2 |
| Cbx4 | | AC134869.3 |
| Srrm1 | | Cnih4 |
| Vav2 | | Zfp330 |
| Hexdc | | Gtf2f2 |
| Fam160a2 | | Naf1 |
| Vps39 | | Ube2w |
| Camk2g | | Lrrfip2 |
| Smtn | | Rgl2 |
| Eri2 | | Pik3cb |
| Slx1b | | Cd82 |
| Srsf11 | | Ap2a2 |
| Usp48 | | Isca1 |
| Hnrnph1 | | Zc3h15 |
| Aaas | | Sdccag3 |
| Lmnb2 | | Txnrd1 |
| Stk38 | | Diexf |
| Cc2d1b | | Cdv3 |
| Tdrd3 | | Tmem104 |
| Hsf2 | | Cnbp |

| | |
|---|---|
| Tmem44 | Rbm15b |
| Ube4b | Stat6 |
| Adrb2 | Slc37a3 |
| Appl2 | Ciapin1 |
| Spata7 | 2410004B18Rik |
| Cep85 | Rnf31 |
| Sept10 | Manf |
| Zscan29 | Yrdc |
| Mknk2 | Nop14 |
| Hmg20a | Mrps18b |
| Cbr4 | Diaph1 |
| Cep95 | Wdr3 |
| Plekhg5 | Dnaja4 |
| Smc3 | Ubtd1 |
| Mau2 | Slc25a45 |
| 2610301B20Rik | Smim15 |
| Rabl2 | Pigx |
| Exog | Caml |
| Rassf3 | St7l |
| Slc41a3 | Knop1 |
| 4930430F08Rik | Ptges3 |
| Phactr1 | N4bp1 |
| Rfesd | Nol10 |
| Ndor1 | Glrx |
| Syde1 | Vapa |
| Fbxl4 | Rrbp1 |
| Sfmbt1 | Tcof1 |
| 2610008E11Rik | Btbd1 |
| Brd8 | Eif2s2 |
| Acsl1 | Vkorc1l1 |
| Wdr59 | Atpaf1 |
| Fam35a | Eed |
| Rpap1 | Pomk |
| Hnmt | Ddx39 |
| Kif7 | Dlgap4 |
| Foxp1 | Fam131a |
| Txndc16 | Carhsp1 |
| Dhx57 | Hspa4 |
| 3110043O21Rik | Ddx50 |
| Nsun6 | Mbd2 |
| Stk11ip | Rnf149 |
| Cntrob | Taf15 |
| Slc25a37 | Ythdf2 |
| Phka2 | Btbd19 |
| Fbxo25 | Set |
| Kif3a | Pwp2 |

| | |
|---|---|
| Nrep | Ddah1 |
| Eif2ak3 | Zbtb7b |
| Zmynd8 | Ddx27 |
| P3h3 | G3bp1 |
| Rubcn | Ltv1 |
| Rbm5 | Csad |
| Cep57 | Senp6 |
| Tcirg1 | Zfp598 |
| Kctd9 | Nadk |
| Hlcs | Ube2h |
| Nsmaf | 4921524J17Rik |
| Polm | Laptm4b |
| Golgb1 | Ppp4r1 |
| Slc5a6 | Actr3 |
| Ptprs | Por |
| Adgrl1 | Rsl24d1 |
| Mta3 | Inpp5a |
| Pld1 | Rnmt |
| Tgif1 | Ddx31 |
| Pou6f1 | Ube2g2 |
| H2afy2 | Tigd2 |
| Rad50 | Stip1 |
| Trmt13 | Hcfc2 |
| Chkb | Kcmf1 |
| Mlh3 | Syncrip |
| Ap4m1 | Eif4e |
| Ptpro | Txndc11 |
| Tmem71 | Thumpd1 |
| Ephb4 | Hspa5 |
| Zfp646 | Acvr1b |
| Cdk5rap2 | Hnrnpu |
| Zrsr1 | Esf1 |
| A230050P20Rik | Nol8 |
| Thnsl2 | Rala |
| Abtb1 | Ywhag |
| Ap5z1 | Hyal2 |
| Ift88 | Ube2q2 |
| Setdb2 | Pspc1 |
| Calcoco1 | Paqr7 |
| Cdk2 | Ttll1 |
| Ccdc77 | Anks1 |
| Ccnl1 | E2f4 |
| Incenp | Atp6v0a1 |
| Gpr137b-ps | Tvp23b |
| Dtx1 | Ube2n |
| Mettl8 | Zfand3 |

| | |
|---|---|
| Scarf2 | 1600012H06Rik |
| Zmym6 | Bnip2 |
| Epm2a | Ube2d1 |
| Zfp873 | Fam98a |
| Ift122 | Tomm70a |
| Kif3c | Prmt6 |
| D3Ertd751e | Stx6 |
| Rmnd5a | Pak1ip1 |
| Gla | Hibch |
| Atg2b | Cab39 |
| Hacd4 | Sbds |
| Megf8 | Riok1 |
| Rasa4 | Fam104a |
| Nek8 | Ube2d2a |
| Cdan1 | Ndfip2 |
| Ece1 | Mcfd2 |
| Hmgxb4 | Rap1a |
| Plekha8 | Ptpn2 |
| Slc38a6 | Ppp2cb |
| Tspyl2 | Senp2 |
| Mtss1l | Uba5 |
| Crbn | Larp4b |
| Ppwd1 | Eif2s1 |
| Foxn3 | Lonp2 |
| Cul9 | Ercc2 |
| Cdk19 | Selenos |
| Abcc10 | Ppp3r1 |
| Tia1 | Utp14a |
| Map2k7 | Smndc1 |
| Iqce | Nol11 |
| Snx7 | Strap |
| Zfp217 | Fpgs |
| Tead3 | Gpbp1 |
| Sec61a2 | Zfp553 |
| Mdga1 | Psme3 |
| Mks1 | Ddx10 |
| Sec22c | Mogs |
| Ehhadh | Zc3h7a |
| Nemp2 | Dcakd |
| Nptxr | Dcaf13 |
| AC122428.2 | Fam214b |
| Zfp286 | Tbrg4 |
| Nbr1 | Riok3 |
| Plpp3 | Fam136a |
| Arhgap24 | Ubxn4 |
| Brdt | Mkln1 |

| | |
|---|---|
| Lrig2 | Yaf2 |
| Mum1 | Ufm1 |
| Epc1 | Noc2l |
| Tdp2 | Ppp2r2a |
| Dnajb4 | Tars |
| Setmar | Vta1 |
| Arhgef25 | 2510039O18Rik |
| Wdcp | Pex13 |
| Gm20532 | Tcn2 |
| Polr2a | Ddx24 |
| Zfp61 | Rap1gds1 |
| Madd | D11Wsu47e |
| Inppl1 | Farsa |
| Recql4 | Ubqln4 |
| Kdm3a | Rad23a |
| Dcaf17 | Pxn |
| Wdr81 | Ghdc |
| Fads2 | Ywhah |
| Zfp386 | Ppp1r2 |
| Adra2a | Slc31a1 |
| Fgfr3 | Dnaja3 |
| Gabbr1 | AC147560.1 |
| Pan2 | Pitpna |
| Mfsd4b4 | Rmdn3 |
| Vstm4 | Ddx18 |
| Bbs5 | Whamm |
| Ubn1 | Blcap |
| Klf6 | Dnajb11 |
| Scd2 | Kmt5a |
| Pafah2 | Phf10 |
| Rbm15 | Rap1b |
| Xpo1 | Efhd2 |
| Pan3 | Ankrd49 |
| Rfx2 | D10Wsu102e |
| Ubr2 | Lsg1 |
| Exoc6 | Map7d1 |
| Cenpj | Elavl1 |
| L3mbtl3 | Acot9 |
| 4833420G17Rik | Tmem248 |
| Rufy2 | Gskip |
| Rbfox2 | Minpp1 |
| Apba1 | Odc1 |
| Zfand4 | Nrf1 |
| Slf1 | Rufy3 |
| Mnt | Ythdf1 |
| Tmem241 | Rab21 |

| | |
|---|---|
| Ppp1r15a | Wdr5 |
| Fos | Arih2 |
| Gm14276 | Atad3a |
| Stk35 | Fbxo45 |
| Gm15601 | Capza2 |
| Ccl22 | Rbm18 |
| Mmp13 | Hsf1 |
| Gpr15 | Cers2 |
| Ch25h | Derl2 |
| Ppbp | Adam15 |
| Gm5879 | Tsr2 |
| Rtn4rl2 | Mgrn1 |
| AC238811.2 | Metap1 |
| Dmkn | Phf14 |
| Palm3 | Pum3 |
| Ackr1 | Hspa9 |
| Tulp1 | Elac2 |
| Dmp1 | Strn4 |
| Gm14963 | Pno1 |
| Gm5781 | Rwdd4a |
| Ms4a6c | Zcchc9 |
| C3 | 2810004N23Rik |
| Fank1 | Coro1c |
| Gm8210 | Senp3 |
| Entpd3 | Ncl |
| H2-M2 | Wdr36 |
| Gm4754 | Krr1 |
| Hmcn2 | Atrip |
| Plin5 | Usp4 |
| Gm10288 | Pabpc4 |
| Cdkn2b | 38412 |
| Spint2 | Mrps5 |
| Ptpn7 | Axin1 |
| Gm15536 | Zdhhc18 |
| Rpl39-ps | Efl1 |
| Ccnb1ip1 | Fbxw7 |
| Ms4a6d | Ubqln1 |
| Prr15 | Gnb1 |
| C1s1 | Lrrc47 |
| H2-DMb1 | Cct6a |
| 1700016A09Rik | Pi4k2a |
| Rnaset2a | Tsnax |
| Crry-ps | Pgam5 |
| Sox3 | Aida |
| Capg | Mark3 |
| Gm10076 | Sar1a |

| | |
|---|---|
| Plscr1 | Akirin1 |
| Gm15772 | Maea |
| Gpr85 | Ppm1a |
| 9330162G02Rik | D17Wsu92e |
| Slc25a29 | Smap2 |
| Gm15417 | Strip1 |
| Necab2 | Pwp1 |
| Gm12166 | Ppp3ca |
| Kcnmb4 | Wbp1l |
| Timm10 | Sec62 |
| Tlr13 | Pmpca |
| Gar1 | Fip1l1 |
| H2afj | Dnm2 |
| a | Afg3l2 |
| Eva1b | Nap1l4 |
| Trp53rkb | Ash2l |
| Rps24 | Cdc42se1 |
| Nop16 | Gorasp2 |
| Rps14 | Spats2 |
| Kcnk6 | Ldah |
| Hck | Morf4l1 |
| Fau | Nars |
| Epop | Rab35 |
| Rhbdf2 | Rab7 |
| Srm | Ehmt2 |
| Igfbp7 | Dcaf8 |
| A430035B10Rik | Ralbp1 |
| Rplp2 | Lta4h |
| Lcp2 | Rufy1 |
| Rps19bp1 | Nde1 |
| Gm9025 | Ints10 |
| Eps8 | Nub1 |
| Gm45713 | Rft1 |
| Wnk1 | Taf6 |
| Sirpb1c | Dok1 |
| Cdkn1a | Etfdh |
| Abl2 | Gatc |
| Trappc4 | Gorasp1 |
| Cox17 | Tpra1 |
| Rpl23a | Zw10 |
| Mrpl12 | Mat2b |
| Kcnab3 | Akirin2 |
| Mir5121 | Mccc2 |
| Tusc1 | Mfsd11 |
| Pgp | Ewsr1 |
| Rplp0 | Coro7 |

| | |
|---|---|
| Epo | Hdac6 |
| Abhd17b | Wipi2 |
| Gm7785 | Mapk7 |
| 2310039H08Rik | Klhl22 |
| Anxa2 | Sppl2b |
| Fhit | Slc12a9 |
| Tpt1 | Ttc5 |
| Tmem11 | Lrsam1 |
| Psma6 | Sh3bp5l |
| Aprt | Cul7 |
| Rps3 | Paics |
| Cox7a2l | Mast2 |
| Tbcc | Nup50 |
| Kyat1 | Med16 |
| Aimp2 | Fkbp9 |
| Lyn | Suv39h1 |
| Rps11 | Hk1 |
| Zdhhc14 | Rfxank |
| 1810037I17Rik | Slc25a12 |
| Tmem37 | Dscr3 |
| Tmed2 | Cnot9 |
| Fscn1 | Tada2a |
| Eif3d | Tab1 |
| Rps16 | Rad51d |
| Npl | Hmgn2 |
| Pmp22 | Cep78 |
| Kxd1 | Zfp362 |
| Faim | Zfp467 |
| Hnrnpa0 | Pik3c3 |
| 2010111I01Rik | Dars2 |
| Atf5 | Phtf1 |
| Dapk3 | Btd |
| Fitm2 | Tmem126b |
| 2310011J03Rik | Lin52 |
| Ostf1 | Hdac5 |
| Rps4x | Gga2 |
| Leprotl1 | Dbf4 |
| Mrpl17 | Ap3m2 |
| Ap4s1 | Abcb8 |
| Cct3 | Rnf122 |
| Rrp7a | Ppp6r2 |
| Gas2l1 | Acadl |
| Etv6 | Pfkm |
| BC003965 | Rragc |
| Eif2b4 | Chd1l |
| 2310015A10Rik | Slc10a7 |

| | |
|---|---|
| Tusc2 | Ints1 |
| Twistnb | Fbxo8 |
| Nudt18 | Psip1 |
| Ebna1bp2 | Sord |
| Cadm4 | Btbd6 |
| Cox18 | Gm29253 |
| Eloc | Atp13a2 |
| Polr1e | Gle1 |
| Dph3 | Manba |
| Rpp30 | Svip |
| Gpn1 | Wdtc1 |
| Ddx54 | Kdelc2 |
| Tomm20 | Xntrpc |
| Vopp1 | Pdcd4 |
| Rrp1 | Cyb5d2 |
| Alg3 | Hs1bp3 |
| Rpl22 | Me2 |
| Zdhhc20 | Fam69a |
| Myl12a | Trmt2b |
| Ccdc25 | 1110008L16Rik |
| Limd2 | Orai3 |
| Gpn2 | Dusp22 |
| St3gal5 | Arhgap17 |
| Stk11 | Pqlc1 |
| Rars | Fech |
| Tspan5 | Emc1 |
| Rassf2 | Ezh1 |
| Def6 | Ext2 |
| Nudt3 | Rcbtb2 |
| Aatf | Pex10 |
| Nubp2 | Rnf26 |
| Ccdc115 | Mtbp |
| Shisa5 | Cdkn1b |
| Pin1 | Lrrc40 |
| Sco2 | Ggact |
| Rbm17 | Selenon |
| Rnf126 | Stx2 |
| AU019823 | Nqo2 |
| Txnl1 | Pot1a |
| Shkbp1 | Nrde2 |
| Yif1a | Cenpo |
| Pknox1 | Pgpep1 |
| Aste1 | Ints7 |
| Dbnl | Gpr89 |
| Trabd | Cyb5rl |
| Larp1 | Aagab |

| | |
|---|---|
| Pou2f2 | Lrrc49 |
| Lsm12 | Zfp346 |
| Kdelr2 | Mapk14 |
| Mfsd13a | Ick |
| Ctsf | Abca7 |
| Pea15a | Ip6k1 |
| Trdmt1 | Pms2 |
| Txnl4a | Mbtps1 |
| Ptpn9 | Fbxo3 |
| Eid2b | Accs |
| Timm17a | Sec14l1 |
| Rfk | Extl2 |
| Lrrc14 | Slc20a2 |
| Ppm1g | Etfbkmt |
| Zfp787 | Xndc1 |
| Ovca2 | Ranbp10 |
| C1qbp | Tmco6 |
| Txn2 | Pnkd |
| Fam3c | Fbxo10 |
| Ctsd | Rpa1 |
| Lins1 | Prim2 |
| Smg9 | Pi4k2b |
| Sar1b | Zbtb5 |
| Cacybp | Rfc1 |
| Tmem5 | Sirt3 |
| Rtfdc1 | Ints9 |
| Arv1 | Dpy19l1 |
| Cops4 | Spice1 |
| Itm2b | Ninl |
| Atp6ap2 | Trim62 |
| Prelid3b | Kdsr |
| Ilk | Garnl3 |
| Trmt11 | Pfkfb4 |
| Gatad1 | Endod1 |
| Eif3h | Helq |
| F8a | Vwa8 |
| Itfg2 | Top3a |
| Pafah1b2 | Cdk20 |
| Etv5 | Fam171a1 |
| Tnfrsf1a | Cep57l1 |
| Rraga | Pck2 |
| Cars | Maged1 |
| Rrn3 | Zhx3 |
| Abcf3 | Arhgap33 |
| Ppm1b | Zfp687 |
| Ubtf | Sertad3 |

| | |
|---|---|
| Ppp1r9b | Amacr |
| Tcp1 | Lrrc56 |
| Mapkapk3 | Pigh |
| Ubash3b | Elmod3 |
| Lemd2 | Pygb |
| Thap4 | Focad |
| Mybbp1a | Rnf157 |
| Mob3a | Ezh2 |
| Commd7 | Emsy |
| Arf6 | Trim37 |
| Fam49b | Cchcr1 |
| Hdac2 | Bbs4 |
| Dhx29 | Erg28 |
| St7 | 5530601H04Rik |
| Cdipt | Coq10a |
| Itgb5 | Pik3cd |
| Saraf | Katnb1 |
| Smarce1 | Zfp395 |
| Rfwd2 | Atg9a |
| Zfp706 | Loxl3 |
| Ercc8 | Wdr45 |
| Lap3 | Gnb5 |
| Agap3 | Cerk |
| Eif3m | Pxmp4 |
| Erp44 | Atg2a |
| Bcas3 | Osbpl2 |
| Mtap | Atg14 |
| Gabpb1 | Lmnb1 |
| Lrp10 | Cnrip1 |
| Dusp3 | Atg13 |
| Capzb | Pcna |
| St3gal3 | Cd99l2 |
| Frmd8 | Slc25a44 |
| Rab14 | Wdr91 |
| Cenpb | Pgm2 |
| Pcmt1 | Cad |
| Os9 | Pcyox1 |
| Wdr61 | St6gal1 |
| Cd151 | Pnrc2 |
| Chtf8 | Map3k20 |
| Rab1a | Abcb4 |
| Golga7 | Dglucy |
| Egln1 | Cryz |
| Sec61a1 | Chek2 |
| Hsp90ab1 | Crebl2 |
| Dpagt1 | Bace1 |

| | |
|---|---|
| Csnk1d | Tmem86a |
| B4galt7 | Rgs2 |
| Gm28551 | Wdr6 |
| Copb2 | Scpep1 |
| Kif1bp | Tbc1d17 |
| Adprh | Pola2 |
| Ctbp1 | Gm45208 |
| Cd81 | Klc4 |
| Rtn3 | Cracr2b |
| Eef1a1 | Armcx6 |
| Rhoa | Rbmx |
| Txndc9 | Mpv17l |
| Fam98b | Fndc10 |
| Actr1a | Hadh |
| Lman2 | Lrig3 |
| Acadm | Cercam |
| Casc3 | Vangl2 |
| Raf1 | Hirip3 |
| Vps51 | Zer1 |
| Birc2 | Gtf2ird1 |
| Rsrc2 | Trim59 |
| Tjap1 | Adk |
| Trim41 | Ccdc167 |
| B4galt4 | Mettl27 |
| Rps6kc1 | Ccdc125 |
| Naa40 | Emilin1 |
| Fut8 | Plekhh3 |
| Ap1g2 | Gabarapl1 |
| Dclre1a | Slc44a2 |
| Ano8 | Lamb2 |
| Hdac10 | Pole3 |
| Pank4 | Msh2 |
| Acox3 | Ctdspl |
| Msl3 | Hspb6 |
| Plpbp | Tedc2 |
| Ccdc66 | Usp49 |
| Ppm1k | Dmac2 |
| Msantd2 | Rnf167 |
| Tmem94 | Tfdp2 |
| A530017D24Rik | Xrcc2 |
| Inpp5k | Nfkbie |
| Tef | H2afz |
| Pcgf6 | N4bp2l1 |
| Sat1 | Casp2 |
| Plcg1 | Fam222a |
| Cflar | Gpr137b |

| | |
|---|---|
| Fam149a | Cnnm3 |
| Ggcx | Slc18b1 |
| Acss1 | Cep72 |
| Rbm48 | Tmem140 |
| Tgfb1i1 | Mtmr10 |
| B3gntl1 | Acot2 |
| Slc9a3r2 | Ptpn21 |
| AW146154 | Aldh2 |
| Ankrd13d | Smc4 |
| Kdm4a | Tnfrsf19 |
| Ankrd24 | Cnnm2 |
| Dek | Dvl2 |
| Numa1 | Fbxo21 |
| Stx17 | Ccdc102a |
| Pias3 | Gnb4 |
| Zfp772 | Acot6 |
| Ptpdc1 | Fbxl20 |
| Taok3 | Bora |
| Unc13b | Xylt2 |
| Herpud1 | Plekhm1 |
| Aldh5a1 | Lpxn |
| S1pr2 | Ctns |
| Dck | Rfwd3 |
| Add3 | Fut10 |
| Odf2l | Vps18 |
| Gm46430 | Zfp709 |
| Pnpla6 | Pdk2 |
| Pkp4 | Capn5 |
| Carf | Rab11fip5 |
| Adcy5 | Abcb10 |
| Slc35e2 | Crocc |
| Lpar6 | Plk4 |
| Dennd6a | Tipin |
| Prkab2 | Nat8f4 |
| Cdr2l | Shpk |
| Ccng2 | Fam189a2 |
| Prickle3 | Rbl2 |
| Tbc1d15 | Mmp11 |
| Rbm33 | Pcnt |
| Nid2 | Cenpq |
| 2700097O09Rik | Mcm8 |
| Tceanc | Stard5 |
| Ccdc62 | Narf |
| Aasdh | Oscp1 |
| Ccdc191 | Acsf2 |
| Nlgn2 | Ldlrap1 |

| | |
|---|---|
| Arid3a | Dact3 |
| Mapkbp1 | Cpped1 |
| Ccdc136 | Glis2 |
| Cntrl | Acy3 |
| Cep162 | Abhd6 |
| Ank2 | Fmo5 |
| Sass6 | 9330102E08Rik |
| Crebbp | Rfc4 |
| Tshz2 | Smyd4 |
| Slc38a9 | Nat8l |
| 4931414P19Rik | Pgm2l1 |
| Rtel1 | Mxd4 |
| Taf5 | Tpcn1 |
| Ddias | Tcp11l2 |
| Cenpl | H6pd |
| Cnst | Haspin |
| Prpf39 | Vsig10 |
| Ccdc149 | Frat1 |
| Col5a2 | Pear1 |
| Ephb3 | Pcyt1a |
| Ampd3 | Cir1 |
| C130074G19Rik | Ccdc14 |
| Gcfc2 | Cdc25b |
| Nfatc1 | Ipp |
| Ints6l | Tube1 |
| Senp8 | Tom1l1 |
| Bbs1 | Dsn1 |
| P2rx7 | Acaca |
| Lamc3 | Abca2 |
| Gpr155 | Hdac7 |
| Parp8 | Arhgap31 |
| Hdhd3 | Cntln |
| Mapk1ip1 | Ttc38 |
| Tob2 | Mok |
| Fgd1 | Cln8 |
| Haus6 | Coq8a |
| Mdm1 | Mcm4 |
| Slc29a3 | Afap1l2 |
| Tctn1 | Prom1 |
| Rwdd3 | Wdr62 |
| Vwa5a | Ppm1l |
| Proser3 | Zfp827 |
| Mical3 | Traip |
| Erbb2 | Nek3 |
| Zfp874a | Atp6v0e2 |
| Snhg3 | Gatsl2 |

| | |
|---|---|
| Pcf11 | Tmem229b |
| Plcd3 | Hbp1 |
| Zfp62 | Gpsm2 |
| Ppfibp2 | Cyb5r1 |
| Ehd2 | Pkmyt1 |
| Cep164 | Fam171a2 |
| Sh3d19 | Hmgb2 |
| Tspyl4 | Rgs7bp |
| Plxnc1 | Uevld |
| Klf11 | Espl1 |
| Fam102a | Tango6 |
| Mtfr2 | Naga |
| Tulp3 | Wdr19 |
| Iqgap3 | Nes |
| Bcl2l11 | Gpr4 |
| Gdf11 | Mblac2 |
| Ing3 | C330027C09Rik |
| Slc36a4 | Lin28b |
| Dact1 | Palm |
| Zfp14 | Nudt7 |
| Arhgef3 | BC055324 |
| Pik3r3 | Sulf2 |
| Zfp940 | Hid1 |
| Pms1 | Pola1 |
| Ppp1r12b | Zfp101 |
| Ccdc142 | Sesn1 |
| Gin1 | Zfp41 |
| Srgap3 | Hmgcs2 |
| Dqx1 | Klhl3 |
| Zfp369 | Wdr90 |
| Pfkfb2 | Mical1 |
| Abhd3 | Fbln5 |
| AW549877 | Rilpl2 |
| Marf1 | Aif1l |
| Zfp182 | Rbm43 |
| Tgfb2 | Rassf7 |
| Zfp65 | Ska2 |
| Tmtc4 | Flcn |
| Rb1cc1 | Chst15 |
| Itga7 | Ptk7 |
| Npr2 | Cbx5 |
| Nexn | Trim14 |
| Hectd4 | Efcab7 |
| Dgke | Hspa12b |
| Hcn3 | Cenps |
| A130077B15Rik | Maob |

| | |
|---|---|
| Tubgcp5 | Epor |
| Hyal1 | Cbx6 |
| Tifa | Tbc1d30 |
| Adcy6 | Wdr76 |
| Clcn2 | Espn |
| Msh3 | Rgs3 |
| Vamp1 | Ccdc120 |
| Slc37a1 | Zfyve1 |
| Lpp | Tmem138 |
| Zmym3 | Arhgef39 |
| Abcc5 | Gins3 |
| Arhgap29 | Tex52 |
| Zfp984 | Gemin8 |
| Ppp4r1l-ps | Mir546 |
| Zfp783 | Naip6 |
| Abhd18 | Lix1 |
| Mansc1 | Cep44 |
| Engase | Ctdsp2 |
| Ccdc138 | Rps6ka5 |
| Casc4 | Pitpnm2 |
| Mphosph9 | Card11 |
| Topbp1 | Farp1 |
| Tspan18 | Melk |
| Zfp512b | Bcl7a |
| Akap17b | Hjurp |
| Senp7 | Mpc1-ps |
| Rab11fip4 | Rell1 |
| Tril | Gins1 |
| Rab30 | Zfp862-ps |
| Ttll13 | Ncaph |
| Il16 | AU020206 |
| Rnft2 | Tppp |
| Pcmtd2 | Skp2 |
| Akap10 | Clec14a |
| Amotl2 | Lig1 |
| Prkag2 | Als2cl |
| Arnt2 | 2700049A03Rik |
| Impact | 38961 |
| Lrp8 | Chst10 |
| Arid5b | Tmem218 |
| Cep295 | Rcan3 |
| Tmem64 | Anks6 |
| Tcf7l1 | Diaph3 |
| Spata2l | Natd1 |
| Mthfr | Slc26a11 |
| Triqk | Slc25a19 |

| | |
|---|---|
| Arhgef9 | Rad18 |
| Ptch1 | Lrrc1 |
| Dse | Rfc5 |
| Helb | Slc38a3 |
| Atp6v0c | Rnf144b |
| Epb41l5 | Ephb6 |
| Chst2 | Amz1 |
| Tnc | Tpx2 |
| Zfp119a | Chtf18 |
| Hrh1 | Arhgef10 |
| Cacnb2 | Mcm3 |
| Snph | Smad6 |
| Pidd1 | Ncapd3 |
| Myh10 | Xrcc3 |
| P2rx6 | Nmral1 |
| Fgd4 | Crybg3 |
| Arhgef6 | Ago4 |
| Fcho1 | Prc1 |
| Slc25a42 | Jam2 |
| Smcr8 | Ccdc40 |
| Aplnr | Pard6g |
| Bicra | C77080 |
| Susd4 | Sdc1 |
| Mns1 | Lpin1 |
| Pml | Cand2 |
| Clk1 | Rrm1 |
| Zfp692 | Mcm6 |
| Grb10 | Ahrr |
| Kazn | Ulk1 |
| Kank2 | Cdh6 |
| Ankrd28 | Peli3 |
| Acad12 | Slc43a1 |
| Wdr5b | Efcab11 |
| Lacc1 | Rad51c |
| Platr25 | Sapcd2 |
| C2cd5 | Cep55 |
| Gm5113 | Cers1 |
| Bbs10 | Brca2 |
| Ntrk3 | Dchs1 |
| Samd4 | Fancm |
| Afdn | Haus5 |
| Fyco1 | Olfml2a |
| Rbl1 | Slc17a5 |
| Eml1 | 4930452B06Rik |
| Cyp27a1 | Sh2d3c |
| Hspa12a | Wdhd1 |

| | |
|---|---|
| Gm20662 | Crebrf |
| Ankrd12 | Hacl1 |
| D130040H23Rik | Dnajc28 |
| Efnb2 | Kif20b |
| Rere | Aspa |
| Dzip1 | Mrgpre |
| Kif13b | Pycr1 |
| Fah | Cenpn |
| Pfkfb1 | Palb2 |
| Zfp383 | Hacl1 |
| Cbs | Zfp784 |
| Zfp174 | Fancb |
| Zkscan7 | Akt3 |
| Cxcl12 | Zfp667 |
| Adam12 | Sorbs3 |
| Grin2d | Ckap2 |
| Obsl1 | Top2a |
| Pip5k1b | Smim10l2a |
| Syngap1 | Fam212b |
| A930024E05Rik | Zfp791 |
| Eva1a | Prkar2b |
| Mllt3 | Psmc3ip |
| Smc2 | Slc7a4 |
| Angptl7 | Dyrk1b |
| Zic2 | Hells |
| Plekha5 | Efs |
| Dock6 | Gspt2 |
| Trim25 | Gtse1 |
| Chd6 | Vash2 |
| Gab3 | Mzf1 |
| 9330104G04Rik | Mad2l1 |
| Ablim1 | Pkd1 |
| Nfatc4 | Mms22l |
| Khnyn | Pcp4l1 |
| Nfkbiz | Kifc5b |
| Zfp423 | Zfp30 |
| Thsd1 | Kif26a |
| B230354K17Rik | Rnf150 |
| Psme2b | Nuf2 |
| Zfp874b | Cdon |
| Tcaf2 | Atad5 |
| P4htm | Sgsh |
| Epb41l3 | Nsl1 |
| Pard3 | Ret |
| 3830408C21Rik | Unc5b |
| Ccdc3 | Abcg3 |

| | |
|---|---|
| Gabpb2 | Ttc26 |
| Krba1 | Ptpre |
| Hykk | Mcm2 |
| Fli1 | Gins2 |
| Trpc2 | Epb41l4a |
| Ptprv | Hmmr |
| AA986860 | Reep1 |
| Spon1 | Arhgef15 |
| Prox2 | Pcyt1b |
| Gpr19 | Racgap1 |
| Slit1 | Trpm6 |
| Jun | Cenpe |
| Zfp975 | B930095G15Rik |
| Kcnab1 | Mindy4 |
| Stox2 | Gm44596 |
| Atf7ip | Cit |
| Lepr | Pir |
| Kmt5c | Pold1 |
| Kalrn | Blm |
| Ano1 | Fam221a |
| Dlg5 | Tmem67 |
| Ackr3 | Cdca7l |
| BC065397 | Cmah |
| Ranbp17 | Enho |
| Bicc1 | Mcm7 |
| Phactr2 | Nusap1 |
| Tbc1d31 | Chek1 |
| Gstt3 | Dnmt1 |
| Gucy1b3 | Kif4 |
| Wfikkn1 | Fam83d |
| Gper1 | Lrrc20 |
| Fmo1 | Fsbp |
| Mylk | Dut |
| Rin1 | Mis18bp1 |
| Fbln2 | Gramd1b |
| Ppp1r16b | Anln |
| Sgsm1 | Kctd12b |
| Podxl | Ncapd2 |
| Caprin2 | Arhgap19 |
| Ppp1r26 | Dnajb14 |
| Fan1 | Impa2 |
| Foxl2 | Cdca5 |
| Gm16907 | Orc1 |
| Slc9a5 | Cdca2 |
| Nhsl2 | Kctd15 |
| Uvssa | Adgra2 |

| | |
|---|---|
| Limch1 | Crocc2 |
| Snx30 | Tarsl2 |
| Npy1r | Ckap2l |
| Srpk3 | Ranbp3l |
| Plekhm3 | Ccna2 |
| Arhgap11a | Polq |
| Fnip1 | Ect2 |
| Ank1 | Lrrn3 |
| Lama1 | Mob3b |
| Ccp110 | Pacsin1 |
| Rcan2 | Chrd |
| Arid4a | Haus4 |
| AC159813.1 | Pcbd1 |
| Zscan18 | Tsku |
| Osbpl10 | Tedc1 |
| Slc38a4 | Bub1b |
| Rarres1 | Aldoa |
| Nr2f2 | Mdfi |
| Elavl4 | Npcd |
| Sema4b | Kif20a |
| Il17rd | Shcbp1 |
| Etl4 | Ccnf |
| Ldlrad3 | 4930522L14Rik |
| Ednrb | Cep97 |
| Bmp4 | Foxm1 |
| Auts2 | Armc2 |
| Caskin2 | Skida1 |
| Unc79 | Nipal3 |
| Acacb | Fignl1 |
| 2610318N02Rik | Fanca |
| Antxr1 | Zwilch |
| Zeb2 | Tcf7 |
| Klf2 | Fancd2 |
| Itpr1 | Mrnip |
| AC131687.1 | Rad51ap1 |
| Plekhh2 | Map4k1 |
| Znrf3 | Ncapg2 |
| Syt12 | Zgrf1 |
| Kantr | Spag5 |
| Mamld1 | Tbc1d4 |
| Raph1 | Gas2l3 |
| Rnf144a | Ulbp1 |
| Ccdc57 | Trerf1 |
| Sfmbt2 | Cdc42ep1 |
| AC151836.1 | Knstrn |
| Edn3 | Ncapg |

| | |
|---|---|
| Atp10d | Chp2 |
| Slamf6 | Lmo7 |
| Bmp5 | Cenpk |
| Rasal2 | Spdl1 |
| Klhl24 | Ung |
| Egf | Tjp3 |
| Pde1a | Cenpf |
| Gria2 | Mybl2 |
| Cavin2 | Nup210 |
| Slc22a8 | Slc43a3 |
| Rapgef6 | Ccdc69 |
| Nynrin | Tmem25 |
| Cybrd1 | Kif11 |
| Gm43634 | Kifc1 |
| Stag3 | Plxnd1 |
| Szt2 | Eme1 |
| Tnfrsf25 | Grb14 |
| Slc2a10 | Kif22 |
| Galnt12 | Hs3st1 |
| Lgr4 | E2f1 |
| Sesn3 | Kif24 |
| Dhtkd1 | Dok4 |
| Kcnj16 | Mki67 |
| Rasl12 | Hapln3 |
| Zdhhc15 | Kif15 |
| Itga3 | S1pr1 |
| Zc3h12b | Esco2 |
| Card10 | Nceh1 |
| Gem | Rad51 |
| Gigyf1 | Dscc1 |
| Adamts10 | Gmnn |
| Daam2 | Fbxo32 |
| Ccdc171 | Fam110b |
| Necab3 | Pif1 |
| Cyp26b1 | Bche |
| Sspn | Gprc5c |
| 4933407K13Rik | Kif18a |
| Mir675 | Cbx2 |
| Akr1c14 | Plk1 |
| Pot1b | Sgo2a |
| Ghr | Sgo1 |
| Cd46 | Bub1 |
| Gjb6 | Cenpu |
| Gm26606 | Pole2 |
| Tnrc6b | BC106179 |
| Mtcl1 | Pik3c2b |

| | |
|---|---|
| Gm26908 | Adamts16 |
| Kat6b | B3gnt8 |
| Zfyve26 | 4930579G24Rik |
| Tsc1 | Fam214a |
| Enpep | Pimreg |
| Rttn | Ptger3 |
| Sema3f | Adra1b |
| Adamts1 | Mbd4 |
| Arhgef28 | Fanci |
| Kif27 | Cacna1e |
| Nphp3 | Kif23 |
| Gm33023 | Spats2l |
| Axin2 | Map2k6 |
| Nr4a1 | Stil |
| Ddr2 | Zfp521 |
| Rmnd1 | Aurka |
| Ptpn22 | Pclaf |
| Pcsk5 | Chst8 |
| 2610037D02Rik | Gm42047 |
| Efhd1 | Cenpa |
| Hes5 | Cdc45 |
| Rhpn2 | Ndc80 |
| Tex9 | Mmp28 |
| Rora | Rtn4rl1 |
| Fbn2 | Dhfr |
| Mfsd2b | Tcf19 |
| Tcf24 | Nek2 |
| Rfx5 | 2510016D11Rik |
| 9930111J21Rik2 | Uhrf1 |
| Pgbd5 | Ska1 |
| Zic3 | Cdca3 |
| Stard9 | E2f7 |
| Cobll1 | Aurkb |
| Scml2 | Mcm5 |
| Hsf2bp | Mamdc2 |
| Parpbp | Ccnb1 |
| Crispld1 | Mxra8 |
| Gm36445 | Tmem98 |
| She | Xkrx |
| Arhgap6 | Dlgap5 |
| Jmy | Rrm2 |
| Nrap | Actr3b |
| Zfp239 | Gpr176 |
| Itgbl1 | Chaf1a |
| Btbd11 | Lpar5 |
| Rasgef1b | Atad2 |

| | |
|---|---|
| Atp13a5 | Xkr5 |
| Slc40a1 | Rragd |
| Slco1a4 | Alms1 |
| 4632427E13Rik | Cpeb1 |
| Kcnk3 | Knl1 |
| 9330159M07Rik | Aspm |
| Rasgef1a | Fbxo5 |
| Ccdc89 | Prim1 |
| Rasl11a | Arntl |
| AC102815.1 | Kntc1 |
| 2610203C20Rik | Dna2 |
| Prr11 | Aph1c |
| Cc2d2a | Gpr157 |
| Gm35339 | Ttk |
| Npr3 | Ticrr |
| Gm45053 | Cenpi |
| F8 | Ccne2 |
| Gimap6 | Mastl |
| Ushbp1 | Dtl |
| Stac3 | Rab38 |
| Rgs1 | Dnah1 |
| Itga8 | Cenph |
| D330041H03Rik | Plscr4 |
| Satb2 | Umodl1 |
| Cyp4f17 | 1700030J22Rik |
| Cttnbp2 | Rad54b |
| Syne2 | Kif18b |
| E230016M11Rik | Fblim1 |
| Amot | Brip1 |
| Zfp454 | Pkp2 |
| Gm17021 | Bard1 |
| Ttc39a | Zfp820 |
| Cracr2a | Cdkn2c |
| Dclre1c | Ska3 |
| Sox18 | Cdc7 |
| Tek | Nkd1 |
| Mboat1 | Kcnh2 |
| Prdm5 | Chaf1b |
| Prickle2 | Lmod1 |
| Cpt1b | Ube2t |
| Gli2 | Srl |
| Egflam | Fnip2 |
| 2700099C18Rik | Exo1 |
| Ccdc162 | Khdrbs3 |
| Gm8146 | Ercc6l |
| 6430573F11Rik | Tk1 |

| | |
|---|---|
| Gm11632 | Lrr1 |
| Tmc5 | Klhl38 |
| Bahcc1 | Rad54l |
| Scn8a | Brca1 |
| Ccdc112 | Slc27a6 |
| Prox1 | Adgrg5 |
| Adora1 | Timeless |
| Gm12258 | 2010001K21Rik |
| Nrg2 | Calhm2 |
| Tert | Psrc1 |
| Celsr1 | Ccdc159 |
| Lrrc32 | Cdkn2a |
| Gm13212 | Ccdc15 |
| Six1 | Cbfa2t3 |
| E130102H24Rik | Pask |
| Rbm20 | Lmcd1 |
| Zcchc11 | Troap |
| Zc3h6 | Myb |
| Osr1 | St8sia4 |
| Gm12248 | Pole |
| Gjb2 | BC030867 |
| Serac1 | Sec16b |
| Ccdc18 | Nav1 |
| Phldb2 | Spaca9 |
| Nos3 | Myzap |
| Csmd2 | Oip5 |
| AC125351.1 | Kcnc2 |
| Itgb4 | Bspry |
| Zfp382 | Fzd10 |
| Kcnj2 | Clspn |
| Gm22489 | E2f8 |
| Gm27010 | Mcm10 |
| Plppr4 | Dnajb13 |
| Gm3604 | Ypel2 |
| Prdm1 | Snai2 |
| Ndst3 | Notum |
| Tbx3 | Wdr63 |
| Zc3h12d | 4930558J18Rik |
| Aldh1l2 | Eps8l2 |
| Tet1 | Rbp4 |
| AC137842.1 | Ppp1r42 |
| Snora28 | Ptprr |
| Cbx7 | Cutal |
| Cbln2 | Pcdhb3 |
| Mpig6b | Lgi2 |
| Zfp763 | Prkcq |

| | |
|---|---|
| Adamts15 | Fam69b |
| Kcnt1 | Rtkn2 |
| Rel | Ddit4l |
| Alx4 | Stac2 |
| Zfp970 | Syt15 |
| Gad1-ps | Dnah5 |
| Pitpnm3 | Lrrc10b |
| Atf3 | F2rl1 |
| Fam227a | Fgfr4 |
| Ndnf | Frmpd2 |
| Plekha6 | Kcnj14 |
| Gm28006 | Col6a6 |
| Gm4673 | Ttll6 |
| Gm22077 | Baiap2l2 |
| Exoc3l2 | Scn4b |
| Amer2 | Fam26e |
| Sytl2 | AF529169 |
| Neil3 | Cerkl |
| Zkscan2 | Gm37689 |
| Igf2os | Gm38399 |
| Fap | Hs3st2 |
| Cemip | Crabp1 |
| Pde4b | EPPK1 |
| Egr4 | Gm4739 |
| Zfp808 | Cd28 |
| Fat2 | Gm9816 |
| Fbln7 | Wnt9b |
| St8sia1 | Gm31409 |
| Ankrd55 | Vill |
| Ikzf4 | B3gnt4 |
| Usp43 | 2610507I01Rik |
| Gm44067 | Hs3st6 |
| Phf21b | Spata9 |
| Alas2 | |
| Gm43759 | |
| Serpind1 | |
| Hist1h2bg | |
| Hey2 | |
| Gm17971 | |
| Tbx18 | |
| Zfp536 | |
| Ddc | |
| Pla2r1 | |
| Trp53cor1 | |
| Sned1 | |
| Gm44053 | |

Fbxl21
Hlf
Tbx6
Gm21781
Gm37019
Enpp3
Gm38190
Gm11398
Gm37309
AC124556.1
Zscan30
Ipcef1
Tfap2b
Wnt9a
Gm12454
Gm43912
Gm25835
Kmo
Tead4
Egr3
Tchh
Cdhr1
Gpr182
Gm15816
Pcdh18
Gpr160
Slit2
Eya1
Gm35154
Igdcc3
Gm37893
Zfhx2os
Snhg14
Gm43437
Atp8b5
D030028A08Rik
Mybl1
Malat1
Arpp21
Zic5
Adra1a
Gm27694
Gm8817
AL645745.1
9930120I10Rik
E030022I16Rik

Lrrn1
Spon2
Ctnna3
Ptgdr
Hist1h2bh
Nlrc4
Ddx60
Asxl3
Gm44574
Gm36527
Gm42979
Gm25994
Gm15880
Klhl33
C130071C03Rik
Mir6935
Capns2
Ahnak2
Gm2788
Tll1
Avil
Trim63
Gdpd2
Foxb1
Gm43362
Lgr6
Gm45743
Catsperd
Gm43951
Hkdc1
Gm37352
Gm45309
Gm36529
Mmrn1
Gm17024
Gm42748
AC154640.4
AC129597.1
Gm37274
AC151971.2
Adh1
Fosb
Col25a1
Gm43679
Gm22589
Gm38220

Cited4
Mir7060
Gm43272
Nxpe5
Ptprb
Ighj4
Wscd2
Nkx6-1
Zfhx4
Gm43351
Gm43088
Sbspon
AC139845.1
Gm20274
Gm43481
Gm43039
Mir6942
Nup210l
Gm44164
Spic
Gm26514
AC131586.4
Serpinb1c
Foxd2os
Il7r
Hectd2os
Gm5857
AC157931.1
Scel
AC108802.1
CT009718.2
Gm43080
C130021I20Rik
Gm43693
Gm43858
Naaladl1
9430034N14Rik
Extl1
Hist1h2ac
Gm45838
AC112142.1
Gm36401
Gm43389
Gm8835
Zfp469
Pax3

Samd11
Gm17999
Gm44033
Rbm11
Insrr
Klhl30
6430511E19Rik
Hepacam2
Irf4
Gm44423
Gm38372
AC124426.1
Gm38375
Otof
Rpgrip1
C1ql3
Gm11870
Fam167a
4930518I15Rik
Rho
Gm42664
Ms4a14
Gm37310
Cmya5
Gm16892
Gm10425
Asic3
Acss3
Gm37168
D030055H07Rik
Agtr1a
Fam228a
Gm22748
Kcnh1
Gm43197
Gm37303
Scn7a
Mab21l2
CT025660.2
Mybpc1
Igf1os
Tspan10
Pkd1l1
Rsg1
Insc
Gm25749

Spata25
Nat8f2
Gm45906
B230323A14Rik
Zan
Sp7
Gm5977
Gpr63
Gm29260
Gm1141
4933433G15Rik
Fendrr
2610028D06Rik
Gm45540
Cbln4
Fgf10

**Supplementary Table 8.** Wiki pathways outputs from the cerebellum only, cerebrum only and shared genes lists presenting the significantly enriched pathways.

| **Cerebrum** exclusive data query | | | | | | |
|---|---|---|---|---|---|---|
| **Term** | **Overlap** | **P-value** | **Adjusted P-value** | **Odds Ratio** | **Combined Score** | **Genes** |
| Cytoplasmic Ribosomal Proteins WP163 | 22/92 | 1.07E-04 | 0.017 | 2.79 | 25.55 | RPS7;RPS5;RPLP0;RPL22;RPL13A;RPL23A;RPL10A;RPL7;RPS4X;RPS14;RPS28;RPS16;RPL18A;RPS29;RPS3;RPLP2;RPS2;FAU;RPS11;RPS10;RPS24;RPL19 |
| **Cerebellum** exclusive data query | | | | | | |
| **Term** | **Overlap** | **P-value** | **Adjusted P-value** | **Odds Ratio** | **Combined Score** | **Genes** |
| Type II interferon signalling (IFNG) WP1253 | 08/34 | 1.75E-05 | 0.002 | 8.57 | 93.86 | SOCS3;STAT1;STAT2;IFNGR2;IRF1;TAP1;IRF9;IFIT2 |
| **Shared** gene list data query | | | | | | |
| Term | Overlap | P-value | Adjusted P-value | Odds Ratio | Combined Score | Genes |
| DNA Replication WP150 | 24/41 | 3.00E-15 | 4.68E-13 | 14.268487 | 477.1506625 | RFC5;PRIM2;PCNA;RFC4;MCM7;RFC1;PRIM1;GMNN;RPA1;MCM10;CDC7;POLA1;POLA2;CDC45;DBF4;ORC1;POLD1;POLE2;MCM3;MCM4;MCM5;MCM6;POLE;MCM2 |
| G1 to S cell cycle control WP413 | 29/61 | 6.45E-15 | 5.03E-13 | 9.1772984 | 299.866371 | PRIM2;CDKN1B;PCNA;MCM7;PRIM1;PKMYT1;CCND3;CCND2;CDC45;CCND1;ORC1;MYC;E2F1;E2F4;POLE;CDKN2C;CDKN2A;GADD45A;RPA1;POLA2;CDK6;TFDP2;CCNE2;POLE2;MCM3;MCM4;MCM5;MCM6;MCM2 |
| Mismatch repair WP1257 | 7-Sep | 1.57E-06 | 8.14E-05 | 35.071389 | 468.7926985 | PCNA;LIG1;RFC1;MSH2;EXO1;POLD1;RPA1 |
| p53 signaling WP2902 | 18/67 | 2.07E-05 | 8.07E-04 | 3.6938436 | 39.84178608 | RRM2;CD82;CDKN2A;GADD45A;IGFBP3;THBS1;GADD45G;CCND3;CCNB1;CCND2;CDK6;CCND1;CCNE2;CHEK2;SESN1;CHEK1;CYCS;GTSE1 |
| mir-193a and MVP in colon cancer metastasis WP3979 | 5-Jun | 3.45E-05 | 0.00108 | 50.049559 | 514.1953177 | CCND2;CCND1;MVP;MYC;G3BP1 |
